# Sensitive and quantitative detection of MHC-I displayed neoepitopes using a semi-automated workflow and TOMAHAQ mass spectrometry

**DOI:** 10.1101/2020.12.16.423097

**Authors:** Samuel B. Pollock, Christopher M. Rose, Martine Darwish, Romain Bouziat, Lélia Delamarre, Craig Blanchette, Jennie R. Lill

**Affiliations:** Genentech, South San Francisco, 94080, California, USA

**Author notes:** **Corresponding author:** Jennie R. Lill, 1 DNA way, South San Francisco, CA 94080.

## Abstract

Advances in several key technologies, including MHC peptidomics, has helped fuel our understanding of basic immune regulatory mechanisms and identify T cell receptor targets for the development of immunotherapeutics. Isolating and accurately quantifying MHC-bound peptides from cells and tissues enables characterization of dynamic changes in the ligandome due to cellular perturbations. This multi-step analytical process remains challenging, and throughput and reproducibility are paramount for rapidly characterizing multiple conditions in parallel. Here, we describe a robust and quantitative method whereby peptides derived from MHC-I complexes from a variety of cell lines, including challenging adherent lines, can be enriched in a semi-automated fashion on reusable, dry-storage, customized antibody cartridges. TOMAHAQ, a targeted mass spectrometry technique that combines sample multiplexing and high sensitivity, was employed to characterize neoepitopes displayed on MHC-I by tumor cells and to quantitatively assess the influence of neoantigen expression and induced degradation on neoepitope presentation.

## INTRODUCTION

As more cancer immunotherapeutic modalities enter the clinic, next generation sequencing and immunopeptidomics have played a key role in furthering our understanding of the mechanisms behind major histocompatibility complex (MHC) -I and -II peptide generation and display (Fig. 1*A*). A variety of cancer antigens (1–4) have been used in personalized cancer vaccines (5–7), as the targets for autologous T cell based therapeutics (8), or in engineered T cell approaches (9). Despite advances across *in silico* methods for predicting the epitopes presented from various human leukocyte antigen (HLA) alleles, many of these algorithms perform poorly for rare alleles and are often associated with a high error rate (10–12). The alternative approach is to use mass spectrometry to directly identify peptides presented on MHC-I. This approach also offers the opportunity to understand ligandome dynamics associated with cellular perturbation.

**Fig. 1.**
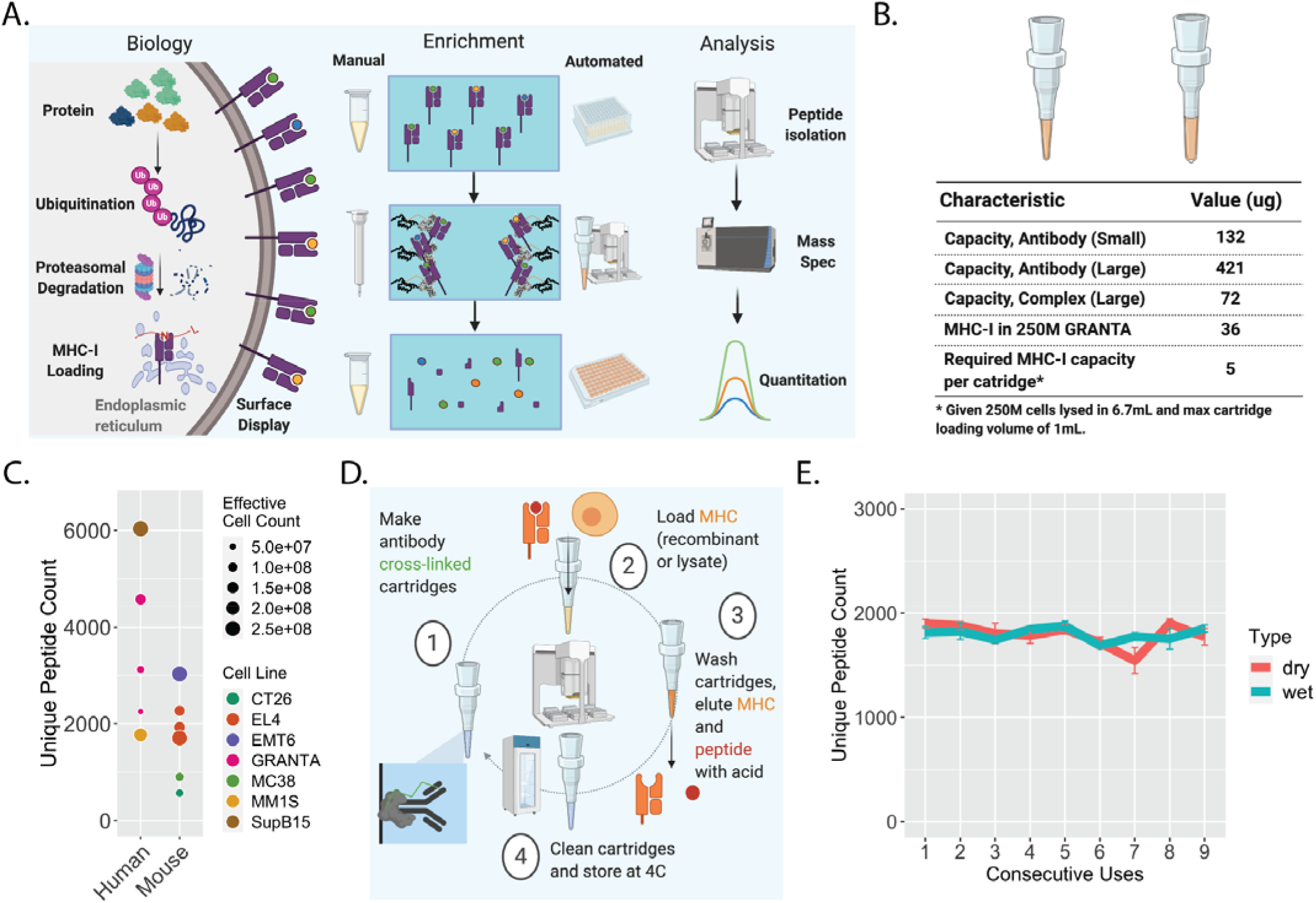
MHC-I enrichment is amenable to automation, reusable reagents, and dry storage of cartridges. A) Overview of the stages in peptide processing and MHC-I presentation, enrichment, and mass spectrometry-based analysis and quantification. In canonical presentation, intracellular proteins are polyubiquitinated within the cell and translocated to the proteasome for degradation. The resulting peptide fragments are transported into the endoplasmic reticulum via transporter associated with antigen processing (TAP) proteins. These fragments are then loaded into MHC-I complexes (consisting of an MHC-I heavy chain and ß2M), further trimmed by endoplasmic reticulum aminopeptidases (ERAP), and the stable MHC-I peptide complex is translocated to the cell surface. The display level of each unique peptide is determined by its abundance, degradation rate, and affinity for the MHC-I allele(s) in a given cell (25). Enrichment involves applying lysate containing MHC-I complexes to a solid support containing anti-MHC-I antibodies, washing away unbound contaminants, and then eluting the MHC-I proteins and associated peptides via acid treatment. Analysis involves peptide isolation and desalting, chromatographic separation, and analysis by mass spectrometry followed by data processing involving identification and quantification steps. B) Table of small and large cartridge characteristics, including recombinant protein capacities, cellular abundance, and amount to be loaded on customized antibody cartridges. C) Unique MHC-I peptides identified using a standard, single-use AssayMAP enrichment workflow, separated by species and effective cell count. D) Antibody cartridge reuse scheme and circular enrichment workflow. Protein A cartridges are loaded with antibody and cross-linked, used to enrich MHC-I complexes, and washed before acid elution. Cartridges are then cleaned via priming with acid and TBS, stored at 4°C, and re-used in a similar manner. E) Unique peptide count observed after nine consecutive uses of custom antibody cartridges, using GRANTA lysate (50 million cell equivalents each), on always-wet vs. dried then re-wetted cartridges.

Two approaches for isolating MHC-I ligandomes include mild acid elution (13), where peptides are released directly from the surface of intact cells, and the more commonly employed immunoprecipitation, where antibodies are used to enrich MHC-I complexes from cell lysate before the release of peptides. Although the latter method is more sensitive than acid elution, typically yields more MHC peptide identifications, and allows for allele-specific enrichment (14, 15), it comes with significant drawbacks. First, it is difficult to automate a workflow that involves many steps, distinct and occasionally detergent-rich buffers, and inputs as unstable as membrane protein complexes within lysates. Second, given the relatively low abundance of MHC-I peptides, standard, untargeted mass spectrometric methods are inefficient in detecting all targets of interest. Yet the field of immunopeptidomics has an ever increasing demand for data from more treatment conditions and cell types along with better detection (16), necessitating ever higher-throughput sample processing and ever more sensitive and quantitative techniques.

Efforts to address throughput have been made by Chong and colleagues (17), who described an elegant and high-throughput system for enriching MHC-I and MHC-II complexes in cell lysates using a stacked-plate-based bead system. However, their approach does not allow for immediate use by an untrained operator, or reuse of the enrichment reagents. In the quantitation space, a number of recent publications have described the use of targeted mass spectrometry and multiplexing to increase the sensitivity and throughput of MHC-I peptidomics. Stopfer and colleagues (18) used tandem mass tag (TMT) labels and heavy isotope peptide-MHC complexes for absolute quantitation of peptides of interest on cell surfaces, whereas Pfamatter and colleagues (19) explored the effects of TMT tagging on MHC-I peptide detection in synchronous precursor selection (SPS) and high-field asymmetric waveform ion mobility spectrometry (FAIMS).

We believed that several steps of this well-established but cumbersome antibody-based enrichment process could be automated to yield a simpler, more reproducible workflow, by combining reusable and easily shareable AssayMAP Bravo-based (20) MHC-I enrichment cartridges with Bravo-based C18 peptide enrichment and modification. We also hypothesized that Triggered by Offset, Multiplexed, Accurate-mass, High-resolution, and Absolute Quantification (TOMAHAQ) mass spectrometry (MS) (21, 22), an internal standard-triggered parallel reaction monitoring (IS-PRM) technique (23) that uses synthetic “trigger” peptides and long MS2 and MS3 collection times for very low abundance target detection, could be applied to MHC-I peptidomics for multiplexed, quantitative, and sensitive detection of high value targets like neoepitopes.

Our method allows for both the routine analysis of >4,000 unique MHC-I peptides from 250 million cells using non-targeted methods, as well as quantitative sensitivity down to the low amol/µL level using TOMAHAQ targeted MS. We demonstrate the utility of this method by monitoring predicted MHC-I neoepitopes in a murine tumor cell line engineered to inducibly express and degrade a single neoantigen, ADP-dependent glucokinase mutant protein (Adpgk(R304M)).

## EXPERIMENTAL PROCEDURES

### Reagents

Reagents were purchased from Sigma-Aldrich unless otherwise specified. Antibodies were purchased from Cell Signaling Technology (Danvers, MA), Abcam (Burlingame, CA), BioLegend (San Diego, CA), ThermoFisher (Waltham, MA), or generated in-house. The dTAG-13 degrader compound was purchased from Tocris (Minneapolis, MN). General plasticware was purchased from Corning and AssayMAP plasticware from Agilent as specified for use with AssayMAP Bravo. Peptides were purchased from JPT Peptide Technologies (Berlin, Germany), Peptides were dissolved in 50% ethylene glycol (Sigma) and stored at −20°C.

### HLA and β2M purification

Recombinant human HLA alleles and beta-2 microglobulin (β2M) were over-expressed in *E. coli*, purified from inclusion bodies, and stored under denaturing conditions (6 M guanidine HCl, 25 mM Tris, pH 8.0) at −80°C. Briefly, β2M and HLA biomass pellets were resuspended in lysis buffer (PBS containing 1% Triton X-114) at a concentration of 5 mL/g, then subjected to microfluidization at 1000 bar homogenization pressure, twice. The resulting suspension was centrifuged at 30,000 × g for 20 min in an ultracentrifuge. The pellets were collected and washed twice with 500 mL of lysis buffer and centrifuged at 30,000 × g for 20 min. The purified inclusion bodies were dissolved in denaturing buffer (20 mM MES, pH 6.0, 6 M guanidine HCl) at a concentration of 10 mL/g and stirred at 4°C overnight. The dissolved pellet was centrifuged at 40,000 × g for 60 min and the supernatant collected and filtered through a 0.22 μm filter. Protein concentration was determined using a bicinchoninic acid (BCA) assay. Samples were flash frozen and stored at −80°C prior to use in generating the MHC-I complexes.

### Generation of recombinant MHC-I complexes

In a 5 L reaction, the selected peptide (0.01 mM), oxidized and reduced glutathione (0.5 mM and 4.0 mM, respectively), recombinant HLA alleles (0.03 mg/mL) and β2M (0.01 mg/mL) were all combined in refold buffer (100 mM Tris, pH 8.0, 400 mM L-arginine, 2 mM EDTA). The refold mixtures were stirred for 4 days at 4°C. The refold solution was filtered through a 0.22 μm filter, concentrated and buffer exchanged by tangential flow filtration (TFF) (Millipore-P2C010C01) into 25 mM Tris, pH 7.5. The protein components were then analyzed by LC/MS to ensure that the HLA was in the appropriate reduced state. The refolded MHC-I complex was purified by ion exchange chromatography using a 5 mL HiTrap Q HP column on an AKTA Pure FPLC. The column was equilibrated with 10 column volumes of 25 mM Tris, pH 7.5 at 5 mL/min flow rate. The MHC-I complex was loaded on the column at a 5 mL/min flow rate and eluted using a 0-60% 25 mM Tris, pH 7.5, 1 M NaCl gradient over 30 column volumes. Samples were analyzed by SDS-PAGE and fractions containing both β2M and HLA bands were pooled and exchanged into storage buffer (25 mM Tris, pH 8.0, 150 mM NaCl). Protein concentration was determined by UV absorbance at 280 nm and samples were flash frozen and stored at −80°C.

### Adherent and suspension cell culture

The murine colon adenocarcinoma cell line MC38 and the human mantle cell lymphoma cell line GRANTA-519 (GRANTA) were obtained from the Genentech cell line bank. MC38 cells were grown in RPMI-1640 supplemented with 10% FBS, 2 mM glutamine, and 25 mM HEPES. Cells were passaged with an 18 h doubling time, at 37°C and 5% CO_2_. GRANTA cells were cultured in RPMI-1640, 10% FBS, 2 mM glutamine and passaged with a 48 h doubling time, at 37°C and 5% CO_2_ in an Infors HT Minitron incubator shaker at 110 rpm.

### Cell transfection and selection

MC38 cells were transfected with the idAdpgkG plasmid (Genscript) using the piggyBAC vector/transposase system with Lipofectamine 2000 (Thermo, see supplemental data for plasmid map). Briefly, growth media on 50% confluent MC38 cells in each well of a 6-well plate was replaced with 1.5 mL OptiMEM. Lipofectamine 2000 in OptiMEM and plasmid DNA in OptiMEM were combined in a 1:1 ratio to yield a 50X-diluted Lipofectamine 2000, 250 ng pBO transposase, and 750 ng vector in 300 μL OptiMEM solution per well. This mixture was incubated at room temperature for 15 min, before distributing evenly across wells. The plate was rocked gently to mix and incubated at 37°C for 2 hr. A 1.5 mL volume of growth media was then added to the OptiMEM-DNA mixture to create a 1:1 OptiMEM:growth media mixture, and the cells were incubated for an additional 24 h before replacing the 1:1 media with full growth media (day 2). After an additional 24 h (day 3), cells were expanded to T75 flasks. After an additional 72 h (day 6), cells were trypsinized and passaged 1:2 into growth media containing 5 μg/mL puromycin. After an additional 72-96 h (day 9-10), cells were tested using flow cytometry and immunoblot for the presence of expressed antigen, and cells with high expression of the idAdpgkG construct were sorted into selection media to yield the idAdpgkG MC38 cell line.

### Cell treatment and harvest

idAdpgkG MC38 cells were treated with 20 ng/mL mouse interferon γ (mIFNγ, R&D Systems, Inc.) alone for 51 h prior to harvest (“control” condition), mIFNγ for 48 h followed by 1 μM dTAG-13 addition for 3 h (“dTAG” condition), mIFNγ and 1 μg/mL doxycycline for 51 h (“dox” condition), or mIFNγ and doxycycline for 48 h followed by dTAG-13 addition for 3 h (“both” condition). The 3 h degrader treatment time was chosen as previous work had shown maximal MHC-I display post-degradation at around 3-6 h (24, 25).

Suspension GRANTA cells and MC38 cells detached using Accutase (Innovative Cell Technologies, Inc.) were counted using a Vi-Cell XR cell counter (Beckman Coulter). Cells were then pelleted by centrifugation flash frozen in liquid nitrogen and stored at −80°C.

### Cell lysis and storage

GRANTA pellets (250 million cells) were lysed in 5 mL non-denaturing OG detergent buffer (PBS, 0.25% sodium deoxycholate, 0.2 mM iodoacetamide (IAA), 1 mM EDTA, 1% octyl-beta-d glucopyranoside (OG), and 1X protease and phosphatase inhibitors (Sigma)), as described previously (17). MC38 pellets (250 million cells) were lysed in 10 mL OG buffer. Lysates were placed on ice for 30 mins then centrifuged at 20,000 x g for 60 min at 4°C. Clarified lysates were added to 50 mL vacuum filters (0.45 μm, Corning) and filtered under gentle vacuum. Filtered solutions were transferred to 15 mL tubes on ice in 4 mL aliquots (an entire GRANTA 250M cell lysis or half of a MC38 250M cell lysis), followed by 1.33 mL each of 50% glycerol and 1 M sucrose to a final concentration of 10% glycerol and 200 mM sucrose, respectively. Samples were then flash frozen, and placed at −80°C.

### Custom cartridge creation, storage, and re-use

Cartridge cross-linking was carried out as described by Purcell et al. (15). Briefly, dry protein A cartridges (Agilent) were primed in PBS at 300 μL/min before loading 1 mg of antibody at 1 mg/mL, followed by washing with PBS. Washing and loading small cartridges occurred at 10 μL and 5 μL/min respectively. Washing and loading large cartridges occurred at 20 μL/min. Cartridges were then equilibrated into 200 mM triethanolamine (TEA, Sigma), loaded with 5 mM dimethyl pimelimidate (DMP, Sigma) in TEA, pH 8.2, over 40 min at room temperature, then washed sequentially with Tris-buffered saline (TBS), 25 mM Tris, pH 8.0 (Tris buffer), 1% acetic acid, Tris buffer, and TBS. Cartridges were placed in a storage rack filled with TBS, 1 mM EDTA, 0.025% sodium azide, sealed with parafilm, and stored at 4°C. To make dried cartridges, an additional wash was performed with 20 mM histidine, 200 mM trehalose, pH 6.0, the cartridges were placed at 37°C for 1 h, and then left at room temperature in the dark for at least 18 h to complete drying. Dried cartridges were then reconstituted in the same manner as dry protein A cartridges.

Re-used wet cartridges were transferred to the AssayMAP Bravo, the neck of each cartridge was dried with a cotton swab, then primed with water, primed with 1% acetic acid, and washed with water before reuse.

### MHC-I enrichment

Frozen cell lysates (6.7 mL) were quickly thawed in a 37°C water bath, then placed on wet ice. Cross-linked cartridges were transferred to the AssayMAP and primed with water, primed with 1% acetic acid, and washed with water. In general, six cartridges were used per lysate as the maximum amount that can be loaded on a cartridge is currently 1 mL (with 0.1 mL dead volume). For human cell lines like GRANTA, six mouse monoclonal anti-HLA Class I antibody (W6/32, Abcam) cross-linked cartridges were used, while for MC38 six 1:1 anti-Db (clone B22-249):anti-Kb (clone Y-3) mixed antibody cartridges were used.

Six 1.1 mL aliquots of the cell lysate were transferred to individual wells in ice-cooled AssayMAP Deepwell plate. The AssayMAP affinity purification method used was: TBS priming (150 μL at 300 μL/min), TBS equilibration (100 μL at 20 μL/min), sample loading (1 ml at 20 μL/min), TBS wash (250 μL at 25 μL/min), Tris wash (250 μL at 25 μL/min), and 1% acetate elution (50 μL at 10 μL/min). The 6 x 50 μL eluates were combined into a single, 1.5 mL LoBind microtube (Eppendorf), flash frozen, and placed at −80°C. Cartridges were then primed with 1% acetic acid, primed with Tris buffer, and washed with TBS before storage in a cartridge rack filled with TBS, 1 mM EDTA, and 0.025% sodium azide, sealed with parafilm, and stored at 4°C for up to three months (before re-use or replacement of storage solution is necessary).

### On-cartridge peptide modification

C18 cartridges were primed using 70% acetonitrile (ACN) with 0.1% formic acid (FA) (100 μL, 300 μL/min), equilibrated in 2% ACN with 0.1% FA (50 μL, 5 μL/min), and eluates described above were loaded (280 μL, 5 μL/min). For performing reduction and alkylation, peptides were reduced in HEPES buffer containing 5 mM Tris (2-carboxyethyl) phosphine (TCEP; 100 μL, 5 μL/min), alkylated with 40 mM IAA (100 μL, 2 μL/min), and washed with HEPES buffer/5 mM TCEP (100 μL, 10 μL/min) followed by HEPES buffer alone (100 μL, 10 μL/min). For performing oxidation, the oxidant solution (5% formic acid, 1X H_2_O_2_) was prepared by mixing (1:1) 10% formic acid with 2X H_2_O_2_ (concentration varies). H_2_O_2_ was diluted with water from fresh 30% H_2_O_2_ (Sigma) to the appropriate concentration. Peptides were then washed with this oxidant solution (150 μL, 5 μL/min) followed by HEPES buffer (100 μL, 10 μL/min). For TMT tagging, 85 μg of TMT6plex reagent (ThermoFisher) in HEPES containing 8% acetonitrile (ACN) was loaded over 25 min (50 μL at 2 μL/min). Finally, peptides were washed with 2% ACN with 0.1% FA (100 μL, 10 μL/min) and eluted in 30% ACN with 0.1% FA (50 μL, 5 μL/min). Fractions were transferred to glass LCMS vials (Agilent), dried for 10 min by speedvac and stored at −80°C.

### In-gel reduction, alkylation, and digest

Cell lysates in 1X lithium dodecylsulfate (LDS) loading buffer containing 1 mM dithiothreitol (DTT) were boiled 5 min and centrifuged at 16,000 x g for 3 min. Approximately 20 μg protein/well was loaded on a 12-well 10% NuPage gel and electrophoresed for 5 mins at 150V. The gel was washed with water three times for 5 min, then stained for 1 h using SimplyBlue at room temperature, and destained in water overnight. Gel bands were excised, cut into small pieces, and transferred to LoBind microtubes. Gel pieces were washed with 50% ACN until the blue dye was fully extracted, then rinsed with 100% ACN and dried in a speedvac for 10 min without heat and stored at −80°C. In-gel digestion was performed by rehydrating gel pieces in 50 mM ammonium bicarbonate pH 8 (AMBIC), reducing with 6 mM TCEP in AMBIC at 37°C for 15 min, alkylating with 40 mM IAA in AMBIC for 30 min at room temperature in the dark, and quenching with 40 mM TCEP in AMBIC at 37°C for 15 min. Gel pieces were washed with water, 50% ACN, and 100% ACN before drying via speedvac. Dry gel pieces were digested with 200 μL of mass spectrometry-grade trypsin solution (4 ng/μL, Promega) in AMBIC and incubated at 37°C for 18 h. A 22 μL aliquot of 10% formic acid was added to quench the reaction and precipitate protein. The gel pieces were incubated in 100 μL of 50% ACN with 5% formic acid for 45 min, then sonicated for 5 min, and the supernatant transferred to a microtube. The extraction was repeated, followed by a final extraction with 100 μL of 90% ACN with 5% formic acid and 5 min sonication. The entire supernatant was dried down by speedvac and stored at −80°C.

### Untargeted mass spectrometry

For untargeted MHC peptidomics, unless otherwise specified, dried peptide samples were re-suspended in 10 μL 2% B (A = 2% ACN with 0.1% FA, B = 98% ACN with 0.1% FA). A 2.5 μL aliquot was injected using an Ultimate 3000 (sample height 1 mm, puncture depth 9 mm, ThermoFisher) onto an IonOpticks C18 column (25 cm Aurora Series, Parkville, Australia), and chromatographically separated using a 2-42% B in 80 min gradient.

The stream was ionized using nanospray ionization (1500 V positive, 600 V negative) and injected onto an Orbitrap Fusion Lumos Tribrid mass spectrometer (MS1 resolution = 240000 m/z, mass range = 350-1350 m/z, AGC target = 1E6, max injection time = 50 ms, cycle time = 1 sec, charge state = 2-3, dynamic exclusion = after 1 time, 30 sec, 7.5 ppm low and high mass tolerance, MS2 isolation window = 0.7 m/z, activation type = HCD, collision energy = 35%, detector type = orbitrap, scan range = normal, resolution = 50000, first mass = 110, AGC target = 2E5, max injection time = 86 ms, with advanced peak determination.)

For global proteomics, peptides were fractionated using the Pierce High pH Reversed-Phase Peptide Fractionation Kit. Briefly, columns were washed with ACN, then 0.1% TFA. Peptides were loaded on-column and washed with water (no acid) and eluted with 5, 7.5, 10, 12.5, 15, 17.5, 20 and 50% ACN in 0.1% triethylamine. Fractions were frozen, lyophilized and reconstituted in 0.1% TFA. Peptides (50% per fraction) were analyzed by nano LC/MS/MS/MS using a Waters NanoAcquity HPLC system interfaced to a ThermoFisher Fusion Lumos mass spectrometer. Peptides were loaded on a trapping column and eluted over a 75 μm analytical column at 350 nL/min; both columns were packed with Luna C18 resin (Phenomenex). Each high pH fraction was separated over a 2 h gradient (16 h instrument time total). The mass spectrometer was operated in data-dependent mode, with MS and MS/MS performed in the Orbitrap at 120,000 full width at half maximum (FWHM) resolution and 50,000 FWHM resolution, respectively. The isolation window was adjusted based on the charge state of the precursor. A 2 s cycle time was employed for all steps.

### Untargeted spectral search

Peptide-spectrum matching was performed in PEAKS v8.5 on imported .raw files. For the untargeted MHC-I peptidomics, searches were performed with the enzyme selection set to “none” in Orbi-Orbi mode under higher-energy collisional dissociation (HCD) fragmentation. Data was filtered using the recommended “mass only” setting. De novo followed by database searches were performed using 15.0 ppm parent and 0.02 Da fragment mass tolerances. Oxidation (+15.99) was set as a variable modification, with up to 3 variable post-translational modifications (PTMs) per peptide, and searched against a Uniprot-derived *Homo sapiens* human proteome (UP000005640, downloaded October 12, 2020, 20,600 genes) or a Uniprot-derived *Mus musculus* database (UP000000589, downloaded March 12, 2019, 22,286 genes). A contaminant database derived from the Contaminant Repository for Affinity Purification (v2012-01-01) was also used to remove nonspecific identifications. A peptide false discovery rate (FDR) cutoff of 1% was used.

For other untargeted MHC-I experiments, the same settings as above were used except constant modifications were set as appropriate, for example constant carbamidomethylation after reduction-alkylation (+57.02) or constant TMT0 (+224.15) or TMT-6plex (+229.16). To investigate TMT labeling efficiency, TMT tags were set as variable modifications. For database search, the idAdpgkG construct sequence was appended to the Uniprot mouse proteome database.

For global proteomics data from MC38-idAdpgk, spectral matching was performed in PEAKS v8.5 on imported .raw files, enzyme selection was set to “Trypsin” in Orbi-Trap mode under CID fragmentation. Data was filtered using the recommended “mass only” setting. De novo followed by database searches were performed using 15.0 ppm parent and 0.5 Da fragment mass tolerances, nonspecific cleavage from both ends with up to two missed cleavages, oxidation (+15.99) and N-terminal acetylation (+42.01) were set as a variable modifications, carbamidomethylation (+57.02) and TMT-6plex (+229.16) were set as a constant modifications, with up to 3 variable PTMs per peptide. For database search, the idAdpgkG construct sequence was appended to the same Uniprot-derived *Mus musculus* database mentioned above. A quantification search was then performed with a mass tolerance of 0.1 Da, MS2 reporter ions, and peptide and protein FDR cutoffs of 1%.

### Tomahaq targeted mass spectrometry and data analysis

The trigger peptides required for the TOMAHAQ method were generated by labeling with TMT-super heavy tags (TMTsh, +235.18, ThermoFisher) on the AssayMAP Bravo. Enriched MHC-I peptide mixtures were labeled with TMT6-plex tags. The TOMAHAQ assay was then performed as described by Rose et al. (22), using the TOMAHAQ companion software to generate the specific methods. Briefly, a target list was generated against the 223 synthesized neoepitopes (NEO223) plus eight controls (NEO223plus8) which include a high abundance peptide derived from CCS protein, two ß2M-derived peptides, two turboGFP-derived peptides, two Adpgk(R304M) protein-derived peptides (outside the neoepitope site), and the wild-type version of the neoepitope peptide. With oxidation states included, this list comes to a total of 315 targets (see “1_MC38_NEO223plus8_OX.csv” in the GitHub repository (https://github.com/sbpollock/NeoToma2021), within the folder Fig4/TOMAHAQ/). This target list was uploaded into TomahaqCompanion (which can be downloaded from https://github.com/CMRose3355/TomahaqCompanionProgram) by clicking “Browse” under the “Load Target Peptides” section. The appropriate PTMs under the “Modifications” section were selected as follows: trigger = SH-TMT, CAM, Ox; target = TMT11, CAM, Ox. The method target list was then formatted and generated by selecting the “Priming Target List” button to generate “2_MC38_NEO223plus8_OX_primingRunInclusionList.csv”. A template priming run file was opened in the Tune method editor, and the method target list uploaded in the inclusion list tab. The priming run was saved (“3_Priming_70min_CID30_NEO223plus8.meth”, this can also be used as a template for future runs) and 500 fmol per peptide of NEO223plus8 trigger peptides was injected and run using the priming run method. The priming run was then uploaded into TomahaqCompanion by selecting “Browse” beside “Priming Run Raw File” under the “Create Tomahaq Method” section, leaving the “Template Method” field blank (allowing for an analysis of targets without yet creating the method).

Selecting “Create Method” produced the priming run analysis, and a list of those targets that did not contain at least three selected MS2 peaks for MS3 was compiled (83 targets made up this list of “dropouts”, leaving a total of 232 targets, and 167 neoantigens plus 4 controls from which those targets are derived). These “dropouts” were removed from the initial target list, and the remaining targets were split between three target lists, such that none of the lists exceeded 100 targets (“4_targets_1.csv” = 100 targets, “4_targets_2.csv” = 99 targets, and “4_targets_3.csv” = 33 targets). No further manual refinement of scans chosen was used, although this is an option in the software. Each of the three final methods were then created by replacing the respective target list under “Load Target Peptides” section, adding the priming run file path to “Priming Run Raw File”, adding a template TOMAHAQ method (“5_Tomahaq_70min_CID_template.meth”) to “Template Method”, then leaving the default settings as is except for setting the method length to 70 min, targeting RT window to 10 min, and selecting “choose best charge state”, before clicking “Create Method”.

The priming and TOMAHAQ gradient were as follows: 2% B, 10 min; 2-42% B, 40 min; 70% B, 15 min, 2% B, 10 min (flow rate 0.3 μL/min).

Details of the template TOMAHAQ mass spectrometry method are as follows: nanospray ionization = 1500V positive, 600 V negative, no advanced peak determination, MS1 resolution = 60000 m/z, mass range = 350-1350 m/z, AGC target = 1E6, max injection time = 50ms, charge state = 2-3, data dependent mode = number of scans (10), targeted mass = inclusion list, mass tolerance = 10 ppm low and high, MS2 isolation window = 0.4 m/z, activation type = CID, collision energy = 30%, detector type = Orbitrap, resolution = 15000, AGC target = 2E5, max injection time = 100 ms, targeted mass trigger tolerance = 10 ppm low and high, trigger only with detection of at least three ions, triggered MS2 isolation window = 0.4 m/z, isolation offset = 5.0099 m/z, activation type = CID, collision energy = 30%, detector type = Orbitrap, resolution = 60000, AGC target = 2E5, max injection time = 900 ms, MS2 data dependent mode = number of scans (1), MS3 precursor selection range = 400-2000, precursor ion exclusion mass width = 5 m/z low and high, isobaric tag loss exclusion = TMT, targeted mass tolerance = 15 ppm low and high, data dependent mode = scans per outcome, MS3 synchronous precursor selection count = 6, MS isolation window = 0.4 m/z, activation type = HCD, activation energy = 55%, detector type = Orbitrap, detector resolution = 60000, first mass = 100, AGC target = 1E6, maximum injection time = 2500 ms. The exact method files used (including .meth, and .xml files) can be found on the Github page (https://github.com/sbpollock/NeoToma2021).

To make up the final mixture for injection, 10 μL of TMTsh-trigger peptide mix in 2% ACN + 0.1% FA in a glass LCMS vial (500 fmol/μL) was transferred to a glass LCMS vial containing the dried idAdpgkG 4-plex peptide mixture. Two replicates were run for each of three final TOMAHAQ methods, and the data was analyzed and exported in TomahaqCompanion using the “Analyze Tomahaq Run” section with the appropriate parameters selected in the other sections as described above. The data was then processed in R by selecting the 4 channels used, grouping by peptide (summing signal across MS2 scans), and filtering signal over noise (S/N) ≥15 per channel times the number of replicates (given the four channels and two replicates, S/N >120 was used).

### RNA sequencing

Cell pellets (1 million cells) underwent RNA extraction, library preparation, and 150 bp paired end sequencing at GeneWiz (South Plainfield, NJ). Raw sequencing data was aligned to mm10 using the R package Rsubread (26) and the data processed using DEseq2 (27).

### Immunoblotting

Cell lysates containing ~30 μg protein were loaded on a 15-well 4-12% NuPage gel and electrophoresed for 45 min at 150V. Proteins were transferred to nitrocellulose using the Transblot Turbo (BioRad). Blots were rinsed with water and dried, reconstituted with TBS, and blocked with Odyssey TBS blocking buffer for 1 h. Antibodies were added at 1:1000 dilution (anti-Adpgk, Abcam, ab228633) or 1:5000 dilution (anti-actin, Cell Signaling Technology, 3700) in TBS blocking buffer, 0.2% Tween-20, and rocked overnight at 4°C. Blots were then rinsed four times with TBS, 0.1% TBST. Secondary antibodies were added at 1:15000 dilution (LI-COR, goat anti-mouse 680, goat anti-rabbit 800) for 1 h at room temperature. Blots were rinsed three times with TBS, 0.1% TBST, then once with TBS. Images were obtained using an Odyssey imager (LI-COR).

### Flow cytometry

Cells from 6-well plates were filtered through a 70 μM filter into flow-cytometer-compatible tubes and analyzed using a FACSCanto II cell analyzer (BD Biosciences). Where applicable, cells were stained with anti-Kb antibody (BioLegend, 116523) or anti-MHC-II (IA/IE) (Thermo, 56-5321-82).

### Differential scanning fluorimetry (DSF)

Differential scanning fluorimetry was performed on a Bio-Rad CFX96RT C1000 Touch qPCR machine monitoring fluorescence at Ex/Em = 587/607 nm. 24 uL of 50 ug/mL solutions of MHC-I-peptide complexes were added to a Bio-Rad hard-shell 96-well PCR plate to wells containing 1uL of 25X SYPRO Orange. MHC-I-peptide complexes were formulated in 25mM Tris pH 8.0, 150mM NaCl, 4mM EDTA, 5% ethylene glycol. The plate was sealed with a Bio-Rad Microseal “C” sealing film. Thermostability measurements were acquired over a temperature gradient of 20□ to 100□ at a rate of 0.2□ per 10 seconds. Data analysis was performed on CFX Manager software and Tm was calculated using the negative first derivative of RFU values over temperature.

### Software

Figure schemes were created using BioRender.com. Plots were created using tidyverse (28) R (29) packages in Rstudio.

### Data Availability

Code and data used to make figures, as well as additional figures, can be found at https://github.com/sbpollock/NeoToma2021. Raw mass spectrometry data can be found at MassIVE (https://massive.ucsd.edu/ProteoSAFe/static/massive.jsp) under accession number MSV000086582. Raw sequencing data can be found at GEO (https://www.ncbi.nlm.nih.gov/geo/) under accession number GSE163326.

### Experimental Design and Statistical Rationale

Statistical confidence is annotated as follows: ns = P > 0.05; * = P ≤ 0.05; ** = P ≤ 0.01; *** = P ≤ 0.001; **** = P ≤ 0.0001. Significant differences between groups were calculated using the paired t-test function in R.

For the experiment described in Fig. 1*B*, values were determined using a multi-point curve with single technical replicates for each concentration (because approximate values were sufficient for demonstrating whether the number of cartridges used in experiments would be able to completely pull down the MHC-I complex loaded). No controls were used in this experiment (because we only sought enrichment values that fell within the calibration curve).

For the experiment described in Fig. 1*C*, between one and three replicates at different effective cell counts were used for all human and mouse samples (because approximate numbers of peptides detected was sufficient for demonstrating expected results and peptide count comparisons). For mouse samples, multiple enrichments were performed based on the number of alleles present (Db and Kb, or Dd, Kd, Ld) and then combined to yield the displayed count. No enrichment controls were used (because detection from, say, isotype control enrichments identifies very few MHC-I-sized peptides (8-10 mers)).

For the experiment described in Fig. 1*E*, three replicates were performed for each cartridge type, per re-use (to allow for determination of inter-cartridge variation and to get an estimate of the approximate amount of noise inherent in each enrichment). Each replicate was searched separately, and peptide-to-spectrum matches (PSMs) that did not meet the peptide FDR cutoff of 1% were excluded.

For the experiments described in Fig. 2, single replicates of each treatment or oxidation condition were used (because the exact number of peptides detected in each condition was less important than the condition-to-condition comparisons (Fig. 2*B*) or the detection trend with increasing oxidant (Figs. 2*C-*2*E*). Each replicate was searched separately, and PSMs that did not meet the peptide FDR cutoff of 1% were excluded.

**Fig. 2.**
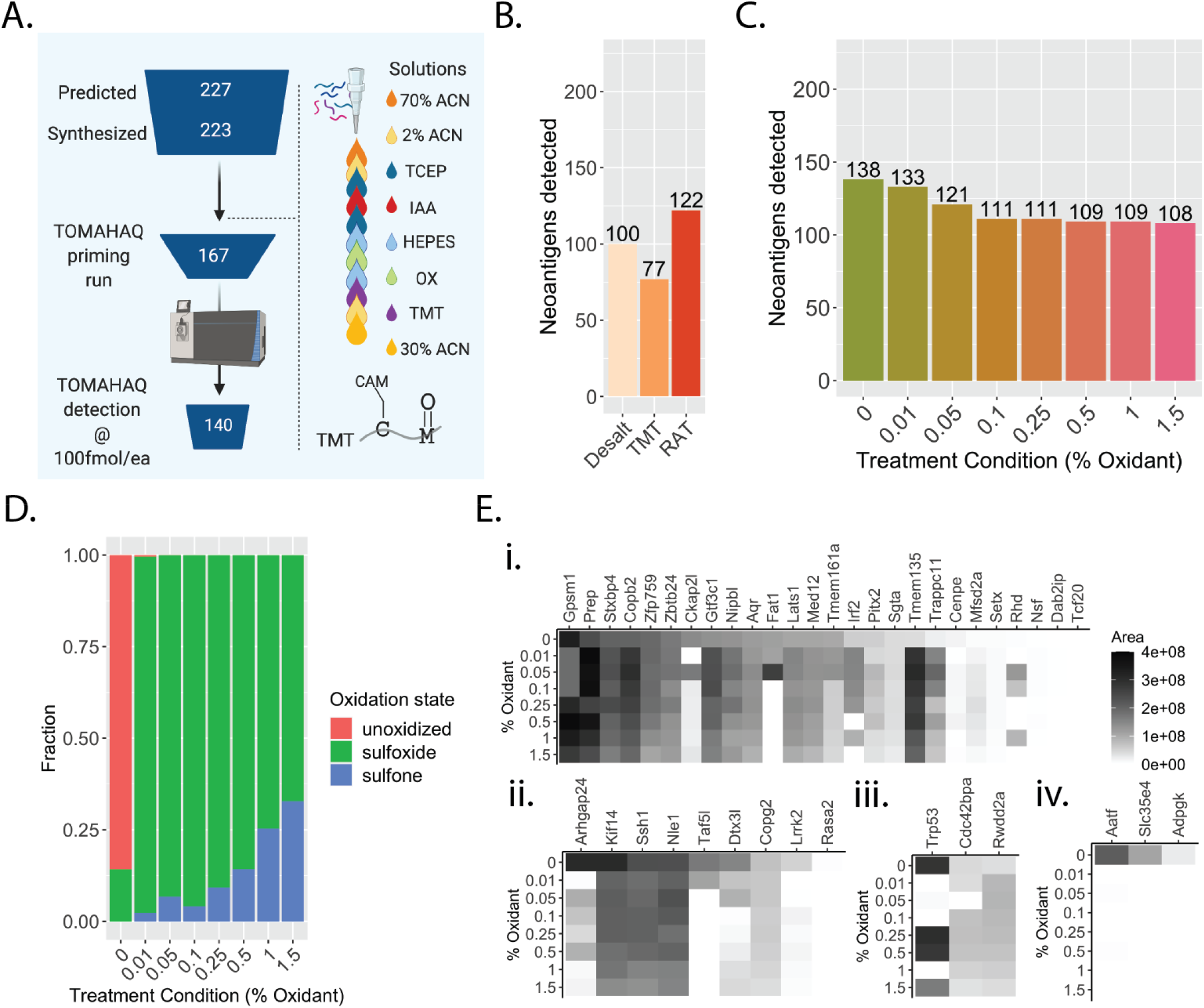
On-cartridge peptide modification is amenable to automation. A) Schematic overview of peptide attrition from ordering to mass spec detection, and full peptide modification using sequential on-cartridge treatment (70% ACN = 70% ACN with 0.1% FA; 2% ACN = 2% ACN with 0.1% FA; IAA = iodoacetamide; OX = variable concentrations of H_2_O_2_ + 5% FA; TMT = tandem mass tag solution; 30% ACN = 30% ACN with 0.1% FA; CAM = carbamidomethyl; C = cysteine; M = methionine). B) Neoantigens detected (out of 223 total) using desalting alone, TMT tagging, or RAT (reduction-alkylation-TMT tagging). C) Neoantigens detected (out of 223 total) using the reduction-alkylation-oxidation-TMT tagging workflow and increasing amounts of oxidant from 0% to 1.5%. D) Distribution of peptide oxidation states (as a fraction of total signal) with increasing amounts of oxidant. E) Intensity of the predominant oxidation state (including unoxidized, by signal) by neoepitope with increasing oxidant for (i) single methionine peptides where detection was improved by treatment with any concentration of oxidant, (ii) single methionine peptides where detection was hampered by oxidant, (iii) double or triple methionine peptides where detection was improved by oxidant, and (iv) double or triple methionine peptides where detection was hampered by oxidant.

For both the MS2 and TOMAHAQ parts of the dilution series experiment described in Fig. 3, only a single replicate of each concentration was run (because comparing typical, single run detection between the two techniques was of greatest interest). However, as described in the methods, the TOMAHAQ part is split between three methods with <= 100 targets each, so each concentration was run three times with each method run once. Additionally, the TOMAHAQ data was filtered for S/N >= 15 × # of channels used (which in this case led to a filter of S/N >= 90 with six channels used), the coefficient of variation (CV) values were calculated for each peptide detected, and an additional filter of CV < 100% was used. For the MS2 data, CV values were similarly calculated for each peptide detected, and a filter of CV < 100% used.

**Fig. 3.**
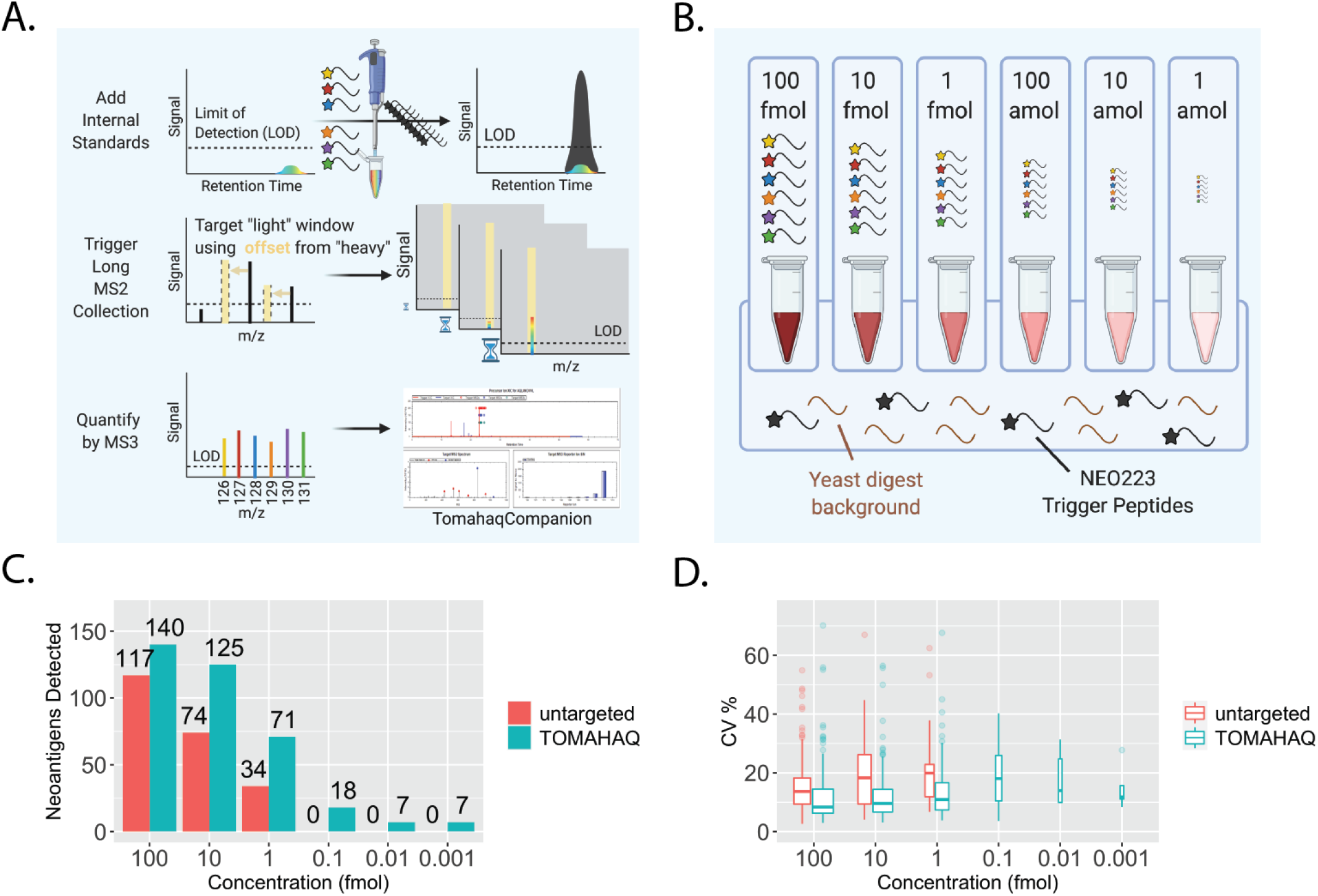
TOMAHAQ is more sensitive and quantitative than untargeted MS against a synthetic neoepitope mixture. A) Scheme showing how TOMAHAQ uses the detection of added internal standard peptides to trigger detection of endogenous TMT-tagged targets. Internal standard triggers (black lines) are first added to the multiplexed mixture (multi-color). MS1 detection of the trigger peptide m/z leads to fragmentation which, if the correct fragment ions are observed on the MS2 level, causes collection and fragmentation of the endogenous peptide via m/z offset followed by a long MS2 fragment collection time (900 ms) to build up signal. The most intense ions are then selected and sent to MS3 for quantification and demultiplexing over a very long collection time (2500 ms). B) Scheme illustrating the 6-plex equimolar dilution series experimental set-up where 100 fmol/μL per NEO223 peptide across each TMT-6plex channel was added to a constant background of 50 ng/mL yeast digest and 500 fmol/μL per NEO223 TMT-SH trigger peptide. This mixture was then 10-fold serially diluted in constant background (yeast digest plus triggers) down to 1 amol/μL per NEO223 peptide across each TMT-6plex channel. C) Comparison of total neoantigens detected with decreasing concentration of target peptides in constant background, between TOMAHAQ and an untargeted method. D) Percent coefficient of variation (CV) for the experiment in C.

For the experiment described in Figs. 4*C* and 4*D*, two technical replicates were run for each of the three TOMAHAQ methods for a total of six runs. As described above, data was filtered by selecting the four channels used, grouping by peptide (summing signal across MS2 scans), and filtering signal over noise (S/N) ≥15 per channel times the number of replicates (given the four channels and two replicates, S/N>120 was used).

**Fig. 4.**
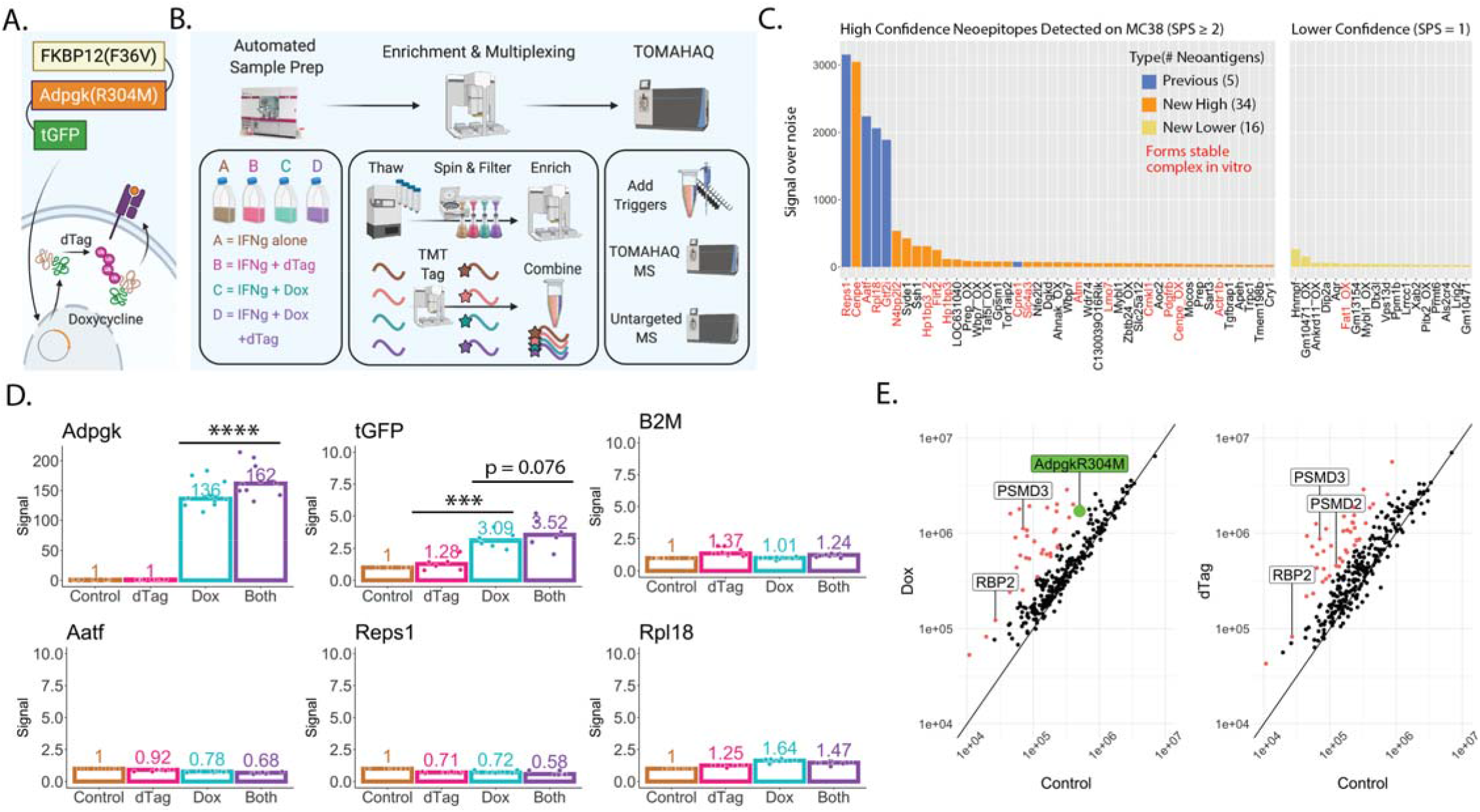
TOMAHAQ detection of neoepitopes in MC38 cells with and without induced expression and/or degradation of the Adpgk(R304M) neoantigen construct idAdpgkG. A) Illustration of the inducible expression and degradation construct idAdgpkG, containing an FKBP12(F36V) degron on the N-terminus and turboGFP protein on the C-terminus of the MC38 neoantigen Adpgk(R304M). Upon treatment with doxycycline, the idAdgpkG neoantigen construct is over-expressed. Treatment with dTAG-13 (dTAG) (24) leads to increased ubiquitination and degradation of the neoantigen, and a potential increase in presentation of neoantigen-derived MHC-I peptides. Flow cytometry and immunoblot confirm the expected changes on the proteomic level (supplemental Fig. S10). B) Scheme showing the idAdpgkG experimental set-up. In the first phase, cells were grown and treated using the automated CompacT SelecT cell culture system under −/+ doxycycline, −/+ dTAG conditions in a 20 ng/mL IFNγ background. Next, RNA sequencing and global proteomic analysis were carried out (supplemental Fig. S11), along with MHC-I enrichment and TMT tagging using the AssayMAP Bravo before being combined into a single, multiplexed sample. Finally, NEO223 synthetic triggers were added to the endogenous, multiplexed sample that was then assayed using TOMAHAQ and untargeted mass spectrometry. C) Neoepitopes detected on the surface of IFNγ-treated idAdpgkG MC38 cells (no dox or dTAG) using the described MHC-I enrichment, peptide modification, and TOMAHAQ workflow. Target peptides identified using signal over noise (S/N) ≥15 per channel per replicate were separated into high and lower confidence categories, based on the number of MS2 fragment ions selected for MS3 (at least two for high confidence, one for lower confidence). Five previously observed neoantigens were detected with high confidence, along with 34 novel neoantigens with high confidence and 16 with lower confidence. To support the identification of novel neoantigens, the names of peptide-MHC complexes found to form stable complexes *in vitro* (melting temperatures > 40°C in a DSF assay, data not shown), are colored red. D) TMT-quantified, treatment specific fold-changes in abundance for high interest targets, normalized to 1 for the control condition. Peptides from AdpgkR304M and turboGFP show significantly increased abundance upon dox treatment, and Adpgk(R304M) peptides show a further significant increase upon additional dTAG treatment. Non-construct targets, including those derived from ß2M and other neoantigens, do not show an increase upon treatment (see Github repository for data on all targets). E) To determine the specificity of dox and/or dTAG treatment, untargeted MHC-I peptidomics was performed on the same four samples that were assayed using TOMAHAQ. The AdpgkR304M neoepitope was detected in doxycycline-containing conditions only, while both dox and dTAG showed increases in numerous peptides (>3-fold upregulation in red), including E3 ligases and proteasomal subunits (labeled). Units on both axes represent signal intensity.

For the experiment described in Fig. 4*E*, one technical replicate of an untargeted MS2 method was run. Data was processed and exported as described in the methods, then filtered for those peptides with MS signal <1E4 in the TMT-126 channel and MS signal < 0.1 × TMT-127 in the TMT-128 channel. The list of proteins was then filtered for those with a total number of spectra greater than 1 (through, for example, two peptides with one spectra each or one peptide with two spectra).

### Development and Analytical Validation Targeted MS Assays/Measurements

TOMAHAQ targeted mass spectrometry (21) was used in the experiments as described in Fig. 3 and Fig. 4. Our implementation of TOMAHAQ can be described as a Tier 3 assay that uses non-manual and real time transition selection. The targets used in the experiment were selected by identifying all peptides that overlap with transcript coding variants in the MC38 line predicted to bind the C57BL/6 mouse resident H2-Db and/or H2-Kb alleles with high affinity, as described by Capietto et al. (14). Transitions are selected for each of these targets during priming run analysis by TomahaqCompanion (22) with optional fragment ion tolerance and SPS m/z > precursor m/z settings to minimize interference. Quantitative signal is the product of MS3 fragmentation of selected transitions and is a product of AGC targets and maximum collection times determined in real time. The transitions selected for each analyte are contained within the final methods (6_targets_1/2/3). The response to concentration, limits of detection, and quantitativeness of each analyte is explored in Fig. 3 and supplemental Fig. S7. Characterization of the internal standard/trigger peptides was performed via LCMS and is included as a supplemental file.

## RESULTS

### Adapting MHC-I enrichment to the AssayMAP Bravo

Automation of the MHC-I immunoaffinity enrichment workflow was achieved by adapting each step of the standard enrichment procedure (15) to the protein A cartridges and liquid handling protocols of Agilent’s AssayMAP Bravo system (Fig. 1*A*). We found that both small (5 μL) and large (25 μL) cartridges had ample capacity for complete immunoaffinity enrichment of MHC-I complex from a typical 250 million cell lysis (using GRANTA cells; Fig. 1*B*, supplemental Fig. S1). We performed single-use cartridge enrichments on a variety of human and mouse cell lines (Fig. 1*C*), and found that the automated workflow yielded a comparable number of unique peptides detected to published, manual workflows (3,000-5,000 for human lines and 1,000-2,000 for mouse lines from 100-500 million cell enrichments)(15, 30).

We hypothesized that the MHC-I enrichment workflow could be made more accessible by taking advantage of the covalent nature of antibody cross-linking to enable re-use of antibody-bound cartridges (Fig. 1*D*, supplemental Fig. S2). Initially we found that clogging of the cartridges beyond the first use prevented our ability to re-use them. However, by reducing the amount of crosslinker, using a large, vacuum lysate filter, diluting the filtered lysate 1.66-fold with glycerol and sucrose, and lowering the temperature of the lysate during enrichment (supplemental Fig. S3), we found that precipitation was minimized and the cartridges were no longer obstructed on repeat use. Using this optimized workflow we demonstrated that cartridges could be used to enrich lysate at least nine times with little decrease in the number of unique peptides observed or change in the composition of peptides detected (Fig. 1*E*, supplemental Fig. S4). This finding held true even if the antibody cross linked cartridges had been dried after cross-linking and then re-wetted.

### On-cartridge peptide modification to enhance the detection of MHC-I peptides

To increase our ability to detect cysteine and methionine-containing peptides, we investigated an on-C18-cartridge peptide derivatization step that included both reduction and alkylation of cysteine residues and an oxidation step (to drive the oxidation of methionine toward the sulfoxide form, Fig. 2*A*). For our test mixture, we set out to synthesize all 227 MC38 neoepitopes predicted to bind the C57BL/6 mouse resident H2-Db and/or H2-Kb alleles with high affinity, as described by Capietto et al. (14). Of the 227 peptides, 223 were readily synthesized at the desired >70% purity and 1 mg scale, and these were mixed to form “NEO223”. The mixture contains 41 cysteine-containing peptides and 60 methionine-containing peptides (which make up 18% and 27% of the complete mixture, respectively).

We first compared neoepitope detection using a standard desalting protocol (17) (supplemental Fig. S5) to one with added TMT labeling or TMT labeling plus reduction and alkylation. We found that peptides could be efficiently labeled with TMT and alkylated (supplemental Fig. S6), and that the number of detected neoepitopes compared to desalting alone decreased upon treatment with TMT, but increased above desalting alone with reduction-alkylation plus TMT (Fig. 2*B*).

We then sought to increase the detection of methionine-containing neoepitopes by driving each neoepitope’s multiple oxidized forms to a single sulfoxide form via peroxide treatment (31). We measured the change in number of neoepitopes detected using untargeted mass spectrometry with increasing amounts of oxidant and observed that higher oxidant led to fewer neoepitopes detected (Fig. 2*C*). When we examined the overall distribution of oxidation states with increasing oxidant, we observed the expected shift from the unoxidized to the sulfoxide state initially, followed by a gradual increase in the sulfone state (Fig. 2*D*). Plotting the maximum signal among each neoepitope’s oxidant forms across oxidant concentrations revealed that although single methionine peptides showed improved detection upon treatment, some or all signal for double and triple methionine peptides was lost with increasing oxidant (Fig. 2*E*).

### Benchmarking the sensitivity of TOMAHAQ mass spectrometry using a synthetic neoepitope mixture

For sensitive quantitation of the MHC-I ligandome, we employed TOMAHAQ mass spectrometry to target the neoepitopes included in NEO223. A summary of the method is illustrated in Fig. 3*A* and consists of the addition of TMT-sh labeled synthetic neoepitope standards to a TMT multiplexed sample of interest, followed by internal-standard-triggered, long duration collection of MS2 and MS3 spectra.

We benchmarked an optimized TOMAHAQ method (supplemental Fig. S7, S8) against an untargeted, MS2 method using a synthetic NEO223 mixture at a variety of concentrations, in a constant yeast digest background (Fig. 3*B*). The untargeted method detected 117, 74, and 34 neoantigens at 100, 10, and 1 fmol respectively, but was unable to detect targets below 1 fmol (Fig. 3*C*). TOMAHAQ analysis of the same samples yielded both greater breadth and sensitivity, with a larger number of neoantigens detected at all fmol concentrations, and 18, 7, and 7 neoantigens detected at 100, 10, and 1 amol, respectively (Fig. 3*C*, supplemental Fig. S9). In addition to identifying more neoantigens, we observed that TOMAHAQ maintained a low median CV % at all peptide concentrations (Fig. 3*D*).

### MC38 model cell line engineering and baseline detection of neoepitopes using TOMAHAQ

In order to demonstrate the sensitivity and quantitative potential for assaying hundreds of MHC-I neoepitopes with TOMAHAQ, we constructed a model system consisting of MC38 cells transfected with an inducible expression and degradation construct called idAdgpkG, consisting of the Adpgk(R304M) neoantigen sequence linked to an FKBP12(F36V) domain on the N-terminus (allowing for inducible degradation via treatment with dTAG-13) and a turboGFP moiety on the C-terminus (allowing for detection and cell sorting by flow cytometry), all under the control of a doxycycline-on (dox) promoter (Fig. 4*A*, supplemental Fig. S10, S11) (32). Using this model system, we designed a four-plex experiment where we could use the NEO223plus8 TOMAHAQ method (NEO223 plus construct and control peptides, see methods) to quantify the abundance of neoepitopes under IFNγ alone (“control”), IFNγ plus dTAG-13 (“dTAG”), IFNγ plus dox (“dox”), and IFNγ plus dox and dTAG (“both”, Fig. 4*B*).

We first examined data from the IFNγ-only, control condition, which represents an endogenous-like state. Neoepitopes were separated into high and lower confidence lists, based on the number of MS2 fragment ions selected for MS3 (at least two for high confidence, one for lower confidence, supplemental Fig. S12). After consolidating singly oxidized and/or unoxidized forms of each neoepitope, we were able to detect five previously observed MC38 neoantigens (7, 33) with high confidence: Aatf, Cpne1, Reps1, Gtf2i, and Rpl18. We also detected a total of 50 unique neoantigens (34 high and 16 lower confidence) that, to the best of our knowledge, have never before been directly observed by mass spectrometry (Fig. 4*C*). For all 227 predicted MC38 neoepitopes, we tested the stability of peptide-MHC complexes by Differential Scanning Fluorimetry (DSF) and found that a number of high confidence neoepitopes and one lower confidence neoepitope (albeit in the oxidized state) showed melting temperatures above 40°C.

### Assaying the magnitude and specificity of changes in neoepitope abundance in the induced neoantigen expression and degradation cell line

We also sought to isolate the effects of dox-based expression induction and dTAG-based degradation of the target neoantigen on MHC-I surface display of its neoepitope. We hypothesized that, due to the correlation of protein abundance and degradation with MHC-I display (34), the Adpgk(R304M)-derived neoepitope would show an increase upon dox induction as the neoantigen abundance increases, and a further increase upon addition of dTAG as the neoantigen degradation rate is increased. As expected, quantifying the TMT signal corresponding to each condition revealed that the presentation of the Adgpk(R304M)-derived neoepitope by MHC-I increased upon dox treatment, and further increased upon dTAG addition (Fig. 4*D*). This pattern was also observed with other construct-derived peptides like those derived from turboGFP (although the dTAG treatment in this case did not lead to a statistically significant increase), but not among control peptides like those derived from ß2M or other neoantigens.

This data gave us confidence that we could measure low abundance quantitative changes in MHC-I epitopes of interest using the TOMAHAQ approach. We also examined the specificity of dox and dTAG treatment by collecting global transcriptomic, proteomic, and peptidomic data. While Adpgk-derived transcripts were the only transcripts to show significant expression increases upon dox treatment, and little change was observed on the proteomic level for all proteins (supplemental Fig. S13), significant increases in MHC-I display were observed not only for the idAdpgkG construct, but for a number of other proteins in a dox and dTAG-dependent manner, including proteasomal and ubiquitination pathway-related proteins (Fig. 4*E*, supplemental Fig. S14).

## DISCUSSION

With the continued growth in the field of cancer immunotherapy and a need to understand the MHC-I ligandome dynamics associated with disease states, further development of robust, high-throughput, and quantitative immunopeptidomics techniques is needed. Understanding the cellular conditions that can influence neoantigen expression, induce alternative epitopes from a tumor antigen (35), or change the ligandome to induce a higher level of immunogenicity (36) are important steps forward in further developing cancer vaccines and T cell based therapeutics. Here we describe a semi-automated method developed for the enrichment, multiplexing, and sensitive detection of hundreds of neoepitopes using reusable cartridges, and applied it to demonstrate neoantigen expression and degradation-induced changes of the MHC-I peptidome.

The AssayMAP Bravo cartridge-based enrichment system was chosen due to its ease-of-use, robustness, and the availability of a wide range of cartridge matrices. Most of the cross-linking and enrichment described would be amenable to gravity flow columns or even magnetic bead-based systems, but one major advantage of the cartridges is that the antibody-resin is immobilized within the plastic cartridge, which is easy to store, handle, and share, especially considering that antibody cross-linked cartridges can be dried and later reconstituted without a decrease in performance. The antibodies used to immunoprecipitate MHC-I complexes are relatively expensive, so reusable cartridges can lead to substantial cost savings and remove the need to perform a cross-linking step every time an enrichment is performed.

In addition to enrichment, we sought to apply automation to the peptide isolation and desalting step, and successfully incorporated additional on-cartridge modifications where reduction-alkylation, TMT tagging, and/or oxidation steps could be applied. A number of clinically relevant MHC-I displayed neoepitopes are known to contain cysteine and/or methionine residues (37, 38), so chemical treatments that maximize the detection of such peptides are desired. We found, in contrast with work from others (19), that TMT tagging decreased the number of neoepitopes detected in our synthetic mixture, however this could be due to the much simpler or hydrophilic mixture with which we were working. On-cartridge labeling and chemical derivatization minimizes the independent number of clean up steps required in the protocol, reducing sample loss and enhancing overall sensitivity.

Several neoepitopes were already known to be presented on MC38 cells, either by direct observation by mass spectrometry (7, 33) or via slowed tumor growth upon vaccination with that neoepitope (39). We sought a targeted mass spectrometry approach that would allow us to quantitatively monitor changes in high value neoepitope targets with the greatest sensitivity possible. TOMAHAQ mass spectrometry allowed us to achieve this sensitive quantitation, to confirm the presence of a number of known neoepitopes, and to reveal dozens of novel neoepitopes that have never before been observed on the MC38 cell surface. At least nine of these novel neoepitopes have been tested and found to be immunogenic by our group, using peptide vaccination (14) and/or RNA lipoplex vaccination (data not shown). This represents a dramatic increase in our ability to detect neoepitopes, as within our untargeted MHC-I assay only two endogenous neoepitopes were observed (not counting those from the over-expressed neoantigen), a result that is typical for detection of neoepitopes in untargeted analyses performed in our lab and others (7, 33, 40). Our synthetic mixture experiments also allow us to assign approximate limits of detection for each neoepitope, providing information on the relative detectability of each, and therefore whether the absence of a given neoepitope may be due to lack of abundance or difficulty of detection.

Beyond improvements to the enrichment and mass spectrometry-based detection steps, we sought to monitor how perturbations in expression and degradation affect the ligandome. Our results indicated that increasing the expression of the neoantigen of interest dramatically increased the presentation of its neoepitope, and that boosting degradation further increased this presentation, albeit subtly, as has been observed previously (24, 25). Other doxycycline-induced changes in presentation were observed as well, lending support to the hypothesis that expression systems often thought to be specific have background effects (41). We quantified these low level abundance changes utilizing instrument methods that surveyed ~100 peptides at a time, utilizing multiple runs to cover our 165 neoantigens-worth of targets (or approximately 250 peptide targets with multiple oxidation forms).

This limitation was due to the current iteration of the mass spectrometer software itself, and improvements to the instrument software and implementation of TOMAHAQ through the instrument API (42) are two active areas of research that would allow for the simultaneous detection of hundreds of targets simultaneously. Targets of interest for MHC-I TOMAHAQ analysis include not only neoepitopes, but small open reading frames (43), viral peptides (44), and transposable elements (45).

As described above, we observed that treatment of the neoantigen construct with degrader led to a slight increase in abundance of its neoepitope on the cell surface. There is evidence that increasing immunogenic peptide abundance has a positive effect on tumor killing efficacy (46), which leads us to hypothesize that, if surface abundance and killing are positively correlated, degraders could potentially be used as a co-treatment to enhance therapeutic efficacy in a cancer vaccine setting where both the neoantigen is known and it possesses an available degrader. Although currently there are only a few dozen commercially available degraders of protein targets, large-scale efforts at converting protein inhibitors to degraders in a modular fashion (47), as well as target agnostic degrader technologies (48), may bring such a treatment strategy within reach in the near future.

Further automation improvements could allow for the seamless connection between complex enrichment and peptide modification stages of the workflow, using, for instance, an external plate hotel and arm for exchanging plates. A more difficult task still would be to apply automation to the filtration and lysate clearing step, although some progress has already been made in this area (49). Further innovation in the cartridges themselves need not be restricted to MHC complexes, as virtually any immunoaffinity enrichment is amenable to this strategy. Other MS improvements include expanding multiplexing capacity to 16-plex and beyond (50).

In summary, we have described a simplified MHC enrichment workflow where an operator with minimal training could, with very little hands-on time and in a single day, perform up to 96 simultaneous enrichments at a similar level of quality as a manual workflow. We also describe methods for modifying peptides for multiplexing and targeted mass spectrometry on the same AssayMAP Bravo system, as well as a TOMAHAQ assay for sensitive detection of hundreds of neoepitope targets. We believe this workflow will prove highly enabling to the peptidomics field.

## Abbreviations

MHC: major histocompatibility complex
TMT: tandem mass tag
SPS: synchronous precursor selection
FAIMS: high-field asymmetric waveform ion mobility spectrometry
TOMAHAQ: Triggered by Offset, Multiplexed, Accurate-mass, High-resolution, and Absolute Quantification
IS-PRM: internal standard-triggered parallel reaction monitoring
MS: mass spectrometer
Adpgk: ADP-dependent glucokinase
β2M: beta-2 microglobulin
BCA: bicinchoninic acid
TFF: tangential flow filtration
mIFNγ: mouse interferon γ
IAA: iodoacetamide
OG: octyl-beta-d glucopyranoside
DMP: dimethyl pimelimidate
TBS: Tris-buffered saline
TEA: triethanolamine
TCEP: Tris (2-carboxyethyl) phosphine
ACN: acetonitrile
LDS: lithium dodecylsulfate
DTT: dithiothreitol
AMBIC: ammonium bicarbonate, pH 8
TFA: trifluoroacetic acid
FWHM: full width at half maximum
HCD: higher-energy collisional dissociation
PTMs: post-translational modifications
FDR: false discovery rate
PSMs: peptide-spectrum matches
CV: coefficient of variation
TAP: transporter associated with antigen processing
ERAP: endoplasmic reticulum aminopeptidases
DSF: differential scanning fluorimetry

## Acknowledgements

We would like to thank Shuai Wu, Steve Murphy, and colleagues at Agilent that so willingly shared advice and prototype reagents. We would also like to thank Rajini Srinivasan and Keith Anderson of the Genentech ACE Hub for help with cloning and transfection, and Saundra Clausen for help with CompacT SelecT-based automated cell growth.

## Notes

https://massive.ucsd.edu/ProteoSAFe/dataset.jsp?accession=MSV000086582

https://www.ncbi.nlm.nih.gov/geo/query/acc.cgi?acc=GSE163326

https://github.com/sbpollock/NeoToma2021

### Competing Interest Statement

The authors have declared no competing interest.

## References

1. Al-Khadairi, G., and Decock, J. (2019) Cancer Testis Antigens and Immunotherapy: Where Do We Stand in the Targeting of PRAME? Cancers (Basel) 11

2. Cheever, M. A., Allison, J. P., Ferris, A. S., Finn, O. J., Hastings, B. M., Hecht, T. T., Mellman, I., Prindiville, S. A., Viner, J. L., Weiner, L. M., and Matrisian, L. M. (2009) The prioritization of cancer antigens: a national cancer institute pilot project for the acceleration of translational research. Clin Cancer Res 15, 5323–5337

3. Finn, O. J. (2017) Human Tumor Antigens Yesterday, Today, and Tomorrow. Cancer Immunol Res 5, 347–354

4. Khong, H. T., Wang, Q. J., and Rosenberg, S. A. (2004) Identification of multiple antigens recognized by tumor-infiltrating lymphocytes from a single patient: tumor escape by antigen loss and loss of MHC expression. J Immunother 27, 184–190

5. Ott, P. A., Hu, Z., Keskin, D. B., Shukla, S. A., Sun, J., Bozym, D. J., Zhang, W., Luoma, A., Giobbie-Hurder, A., Peter, L., Chen, C., Olive, O., Carter, T. A., Li, S., Lieb, D. J., Eisenhaure, T., Gjini, E., Stevens, J., Lane, W. J., Javeri, I., Nellaiappan, K., Salazar, A. M., Daley, H., Seaman, M., Buchbinder, E. I., Yoon, C. H., Harden, M., Lennon, N., Gabriel, S., Rodig, S. J., Barouch, D. H., Aster, J. C., Getz, G., Wucherpfennig, K., Neuberg, D., Ritz, J., Lander, E. S., Fritsch, E. F., Hacohen, N., and Wu, C. J. (2017) An immunogenic personal neoantigen vaccine for patients with melanoma. Nature 547, 217–221

6. Sahin, U., Derhovanessian, E., Miller, M., Kloke, B. P., Simon, P., Lower, M., Bukur, V., Tadmor, A. D., Luxemburger, U., Schrors, B., Omokoko, T., Vormehr, M., Albrecht, C., Paruzynski, A., Kuhn, A. N., Buck, J., Heesch, S., Schreeb, K. H., Muller, F., Ortseifer, I., Vogler, I., Godehardt, E., Attig, S., Rae, R., Breitkreuz, A., Tolliver, C., Suchan, M., Martic, G., Hohberger, A., Sorn, P., Diekmann, J., Ciesla, J., Waksmann, O., Bruck, A. K., Witt, M., Zillgen, M., Rothermel, A., Kasemann, B., Langer, D., Bolte, S., Diken, M., Kreiter, S., Nemecek, R., Gebhardt, C., Grabbe, S., Holler, C., Utikal, J., Huber, C., Loquai, C., and Tureci, O. (2017) Personalized RNA mutanome vaccines mobilize poly-specific therapeutic immunity against cancer. Nature 547, 222–226

7. Yadav, M., Jhunjhunwala, S., Phung, Q. T., Lupardus, P., Tanguay, J., Bumbaca, S., Franci, C., Cheung, T. K., Fritsche, J., Weinschenk, T., Modrusan, Z., Mellman, I., Lill, J. R., and Delamarre, L. (2014) Predicting immunogenic tumour mutations by combining mass spectrometry and exome sequencing. Nature 515, 572–576

8. Fernandez-Poma, S. M., Salas-Benito, D., Lozano, T., Casares, N., Riezu-Boj, J. I., Mancheno, U., Elizalde, E., Alignani, D., Zubeldia, N., Otano, I., Conde, E., Sarobe, P., Lasarte, J. J., and Hervas-Stubbs, S. (2017) Expansion of Tumor-Infiltrating CD8(+) T cells Expressing PD-1 Improves the Efficacy of Adoptive T-cell Therapy. Cancer Res 77, 3672–3684

9. Locke, F. L., Neelapu, S. S., Bartlett, N. L., Siddiqi, T., Chavez, J. C., Hosing, C. M., Ghobadi, A., Budde, L. E., Bot, A., Rossi, J. M., Jiang, Y., Xue, A. X., Elias, M., Aycock, J., Wiezorek, J., and Go, W. Y. (2017) Phase 1 Results of ZUMA-1: A Multicenter Study of KTE-C19 Anti-CD19 CAR T Cell Therapy in Refractory Aggressive Lymphoma. Mol Ther 25, 285–295

10. Lundegaard, C., Lamberth, K., Harndahl, M., Buus, S., Lund, O., and Nielsen, M. (2008) NetMHC-3.0: accurate web accessible predictions of human, mouse and monkey MHC class I affinities for peptides of length 8–11. Nucleic Acids Res 36, W509–512

11. O’Donnell, T. J., Rubinsteyn, A., Bonsack, M., Riemer, A. B., Laserson, U., and Hammerbacher, J. (2018) MHCflurry: Open-Source Class I MHC Binding Affinity Prediction. Cell Syst 7, 129–132 e124

12. Peters, B., Nielsen, M., and Sette, A. (2020) T Cell Epitope Predictions. Annu Rev Immunol 38, 123–145

13. Lanoix, J., Durette, C., Courcelles, M., Cossette, E., Comtois-Marotte, S., Hardy, M. P., Cote, C., Perreault, C., and Thibault, P. (2018) Comparison of the MHC I Immunopeptidome Repertoire of B-Cell Lymphoblasts Using Two Isolation Methods. Proteomics 18, e1700251

14. Capietto, A. H., Jhunjhunwala, S., Pollock, S. B., Lupardus, P., Wong, J., Hansch, L., Cevallos, J., Chestnut, Y., Fernandez, A., Lounsbury, N., Nozawa, T., Singh, M., Fan, Z., de la Cruz, C. C., Phung, Q. T., Taraborrelli, L., Haley, B., Lill, J. R., Mellman, I., Bourgon, R., and Delamarre, L. (2020) Mutation position is an important determinant for predicting cancer neoantigens. J Exp Med 217

15. Purcell, A. W., Ramarathinam, S. H., and Ternette, N. (2019) Mass spectrometry-based identification of MHC-bound peptides for immunopeptidomics. Nat Protoc 14, 1687–1707

16. Vizcaino, J. A., Kubiniok, P., Kovalchik, K. A., Ma, Q., Duquette, J. D., Mongrain, I., Deutsch, E. W., Peters, B., Sette, A., Sirois, I., and Caron, E. (2020) The Human Immunopeptidome Project: A Roadmap to Predict and Treat Immune Diseases. Mol Cell Proteomics 19, 31–49

17. Chong, C., Marino, F., Pak, H., Racle, J., Daniel, R. T., Muller, M., Gfeller, D., Coukos, G., and Bassani-Sternberg, M. (2018) High-throughput and Sensitive Immunopeptidomics Platform Reveals Profound Interferongamma-Mediated Remodeling of the Human Leukocyte Antigen (HLA) Ligandome. Mol Cell Proteomics 17, 533–548

18. Stopfer, L. E., Mesfin, J. M., Joughin, B. A., Lauffenburger, D. A., and White, F. M. (2020) Multiplexed relative and absolute quantitative immunopeptidomics reveals MHC I repertoire alterations induced by CDK4/6 inhibition. Nat Commun 11, 2760

19. Pfammatter, S., Bonneil, E., Lanoix, J., Vincent, K., Hardy, M. P., Courcelles, M., Perreault, C., and Thibault, P. (2020) Extending the Comprehensiveness of Immunopeptidome Analyses Using Isobaric Peptide Labeling. Anal Chem 92, 9194–9204

20. Fulton, S., Murphy, S., Reich, J., Van Den Heuvel, Z., Sakowski, R., Smith, R., and Agee, S. (2011) A high-throughput microchromatography platform for quantitative analytical scale protein sample preparation. J Lab Autom 16, 457–467

21. Erickson, B. K., Rose, C. M., Braun, C. R., Erickson, A. R., Knott, J., McAlister, G. C., Wuhr, M., Paulo, J. A., Everley, R. A., and Gygi, S. P. (2017) A Strategy to Combine Sample Multiplexing with Targeted Proteomics Assays for High-Throughput Protein Signature Characterization. Mol Cell 65, 361–370

22. Rose, C. M., Erickson, B. K., Schweppe, D. K., Viner, R., Choi, J., Rogers, J., Bomgarden, R., Gygi, S. P., and Kirkpatrick, D. S. (2019) TomahaqCompanion: A Tool for the Creation and Analysis of Isobaric Label Based Multiplexed Targeted Assays. J Proteome Res 18, 594–605

23. Gallien, S., Kim, S. Y., and Domon, B. (2015) Large-Scale Targeted Proteomics Using Internal Standard Triggered-Parallel Reaction Monitoring (IS-PRM). Mol Cell Proteomics 14, 1630–1644

24. Jensen, S. M., Potts, G. K., Ready, D. B., and Patterson, M. J. (2018) Specific MHC-I Peptides Are Induced Using PROTACs. Front Immunol 9, 2697

25. Moser, S. C., Voerman, J. S. A., Buckley, D. L., Winter, G. E., and Schliehe, C. (2017) Acute Pharmacologic Degradation of a Stable Antigen Enhances Its Direct Presentation on MHC Class I Molecules. Front Immunol 8, 1920

26. Liao, Y., Smyth, G. K., and Shi, W. (2019) The R package Rsubread is easier, faster, cheaper and better for alignment and quantification of RNA sequencing reads. Nucleic Acids Res 47, e47

27. Love, M. I., Huber, W., and Anders, S. (2014) Moderated estimation of fold change and dispersion for RNA-seq data with DESeq2. Genome Biol 15, 550

28. Wickham, H., Averick, M., Bryan, J., Chang, W., McGowan, L. D. A., François, R., Grolemund, G., Hayes, A., Henry, L., Hester, J., Kuhn, M., Pedersen, T. L., Miller, E., Bache, S. M., Müller, K., Ooms, J., Robinson, D., Seidel, D. P., Spinu, V., Takahashi, K., Vaughan, D., Wilke, C., Woo, K., and Yutani, H. (2019) Welcome to the Tidyverse. Journal of Open Source Software 4, 1686

29. R Development Core Team (2010) R: A language and environment for statistical computing., R Foundation for Statistical Computing, Vienna, Austria

30. Laumont, C. M., Vincent, K., Hesnard, L., Audemard, E., Bonneil, E., Laverdure, J. P., Gendron, P., Courcelles, M., Hardy, M. P., Cote, C., Durette, C., St-Pierre, C., Benhammadi, M., Lanoix, J., Vobecky, S., Haddad, E., Lemieux, S., Thibault, P., and Perreault, C. (2018) Noncoding regions are the main source of targetable tumor-specific antigens. Sci Transl Med 10

31. Phu, L., Izrael-Tomasevic, A., Matsumoto, M. L., Bustos, D., Dynek, J. N., Fedorova, A. V., Bakalarski, C. E., Arnott, D., Deshayes, K., Dixit, V. M., Kelley, R. F., Vucic, D., and Kirkpatrick, D. S. (2011) Improved quantitative mass spectrometry methods for characterizing complex ubiquitin signals. Mol Cell Proteomics 10, M110 003756

32. Nabet, B., Roberts, J. M., Buckley, D. L., Paulk, J., Dastjerdi, S., Yang, A., Leggett, A. L., Erb, M. A., Lawlor, M. A., Souza, A., Scott, T. G., Vittori, S., Perry, J. A., Qi, J., Winter, G. E., Wong, K. K., Gray, N. S., and Bradner, J. E. (2018) The dTAG system for immediate and target-specific protein degradation. Nat Chem Biol 14, 431–441

33. Hos, B. J., Camps, M. G. M., van den Bulk, J., Tondini, E., van den Ende, T. C., Ruano, D., Franken, K., Janssen, G. M. C., Ru, A., Filippov, D. V., Arens, R., van Veelen, P. A., Miranda, N., and Ossendorp, F. (2019) Identification of a neo-epitope dominating endogenous CD8 T cell responses to MC-38 colorectal cancer. Oncoimmunology 9, 1673125

34. Bassani-Sternberg, M., Pletscher-Frankild, S., Jensen, L. J., and Mann, M. (2015) Mass spectrometry of human leukocyte antigen class I peptidomes reveals strong effects of protein abundance and turnover on antigen presentation. Mol Cell Proteomics 14, 658–673

35. Blachere, N. E., Darnell, R. B., and Albert, M. L. (2005) Apoptotic cells deliver processed antigen to dendritic cells for cross-presentation. PLoS Biol 3, e185

36. Zervoudi, E., Saridakis, E., Birtley, J. R., Seregin, S. S., Reeves, E., Kokkala, P., Aldhamen, Y. A., Amalfitano, A., Mavridis, I. M., James, E., Georgiadis, D., and Stratikos, E. (2013) Rationally designed inhibitor targeting antigen-trimming aminopeptidases enhances antigen presentation and cytotoxic T-cell responses. Proc Natl Acad Sci U S A 110, 19890–19895

37. Dao, T., Korontsvit, T., Zakhaleva, V., Jarvis, C., Mondello, P., Oh, C., and Scheinberg, D. A. (2017) An immunogenic WT1-derived peptide that induces T cell response in the context of HLA-A*02:01 and HLA-A*24:02 molecules. Oncoimmunology 6, e1252895

38. Dutoit, V., Taub, R. N., Papadopoulos, K. P., Talbot, S., Keohan, M. L., Brehm, M., Gnjatic, S., Harris, P. E., Bisikirska, B., Guillaume, P., Cerottini, J. C., Hesdorffer, C. S., Old, L. J., and Valmori, D. (2002) Multiepitope CD8(+) T cell response to a NY-ESO-1 peptide vaccine results in imprecise tumor targeting. J Clin Invest 110, 1813–1822

39. Aurisicchio, L., Salvatori, E., Lione, L., Bandini, S., Pallocca, M., Maggio, R., Fanciulli, M., De Nicola, F., Goeman, F., Ciliberto, G., Conforti, A., Luberto, L., and Palombo, F. (2019) Poly-specific neoantigen-targeted cancer vaccines delay patient derived tumor growth. J Exp Clin Cancer Res 38, 78

40. Wickstrom, S. L., Lovgren, T., Volkmar, M., Reinhold, B., Duke-Cohan, J. S., Hartmann, L., Rebmann, J., Mueller, A., Melief, J., Maas, R., Ligtenberg, M., Hansson, J., Offringa, R., Seliger, B., Poschke, I., Reinherz, E. L., and Kiessling, R. (2019) Cancer Neoepitopes for Immunotherapy: Discordance Between Tumor-Infiltrating T Cell Reactivity and Tumor MHC Peptidome Display. Front Immunol 10, 2766

41. Vogel, R., Al-Daccak, R., Drews, O., Alonzeau, J., Mester, G., Charron, D., Stevanovic, S., and Mallet, J. (2013) Mass spectrometry reveals changes in MHC I antigen presentation after lentivector expression of a gene regulation system. Mol Ther Nucleic Acids 2, e75

42. Yu, Q., Xiao, H., Jedrychowski, M. P., Schweppe, D. K., Navarrete-Perea, J., Knott, J., Rogers, J., Chouchani, E. T., and Gygi, S. P. (2020) Sample multiplexing for targeted pathway proteomics in aging mice. Proc Natl Acad Sci U S A 117, 9723–9732

43. Martinez, T. F., Chu, Q., Donaldson, C., Tan, D., Shokhirev, M. N., and Saghatelian, A. (2020) Accurate annotation of human protein-coding small open reading frames. Nat Chem Biol 16, 458–468

44. Smith, C. C., Beckermann, K. E., Bortone, D. S., De Cubas, A. A., Bixby, L. M., Lee, S. J., Panda, A., Ganesan, S., Bhanot, G., Wallen, E. M., Milowsky, M. I., Kim, W. Y., Rathmell, W. K., Swanstrom, R., Parker, J. S., Serody, J. S., Selitsky, S. R., and Vincent, B. G. (2018) Endogenous retroviral signatures predict immunotherapy response in clear cell renal cell carcinoma. J Clin Invest 128, 4804–4820

45. Kong, Y., Rose, C. M., Cass, A. A., Williams, A. G., Darwish, M., Lianoglou, S., Haverty, P. M., Tong, A. J., Blanchette, C., Albert, M. L., Mellman, I., Bourgon, R., Greally, J., Jhunjhunwala, S., and Chen-Harris, H. (2019) Transposable element expression in tumors is associated with immune infiltration and increased antigenicity. Nat Commun 10, 5228

46. Wang, G., Chow, R. D., Bai, Z., Zhu, L., Errami, Y., Dai, X., Dong, M. B., Ye, L., Zhang, X., Renauer, P. A., Park, J. J., Shen, L., Ye, H., Fuchs, C. S., and Chen, S. (2019) Multiplexed activation of endogenous genes by CRISPRa elicits potent antitumor immunity. Nat Immunol 20, 1494–1505

47. Krajcovicova, S., Jorda, R., Hendrychova, D., Krystof, V., and Soural, M. (2019) Solid-phase synthesis for thalidomide-based proteolysis-targeting chimeras (PROTAC). Chem Commun (Camb) 55, 929–932

48. Clift, D., McEwan, W. A., Labzin, L. I., Konieczny, V., Mogessie, B., James, L. C., and Schuh, M. (2017) A Method for the Acute and Rapid Degradation of Endogenous Proteins. Cell 171, 1692–1706 e1618

49. Schlaeppi, J. M., Henke, M., Mahnke, M., Hartmann, S., Schmitz, R., Pouliquen, Y., Kerins, B., Weber, E., Kolbinger, F., and Kocher, H. P. (2006) A semi-automated large-scale process for the production of recombinant tagged proteins in the Baculovirus expression system. Protein Expr Purif 50, 185–195

50. Thompson, A., Wolmer, N., Koncarevic, S., Selzer, S., Bohm, G., Legner, H., Schmid, P., Kienle, S., Penning, P., Hohle, C., Berfelde, A., Martinez-Pinna, R., Farztdinov, V., Jung, S., Kuhn, K., and Pike, I. (2019) TMTpro: Design, Synthesis, and Initial Evaluation of a Proline-Based Isobaric 16-Plex Tandem Mass Tag Reagent Set. Anal Chem 91, 15941–15950

