## Supplemental Information for "Sensitive and quantitative detection of MHC-I displayed neoepitopes using a semi-automated workflow and TOMAHAQ mass spectrometry"

**Supplemental Figures**

**Fig. S1 - Gel data for cartridge capacities**

Coomassie gel data used to create calibration curves and estimated capacity for antibody on protein A cartridges, and for recombinant MHC-I complex on antibody cross-linked cartridges.

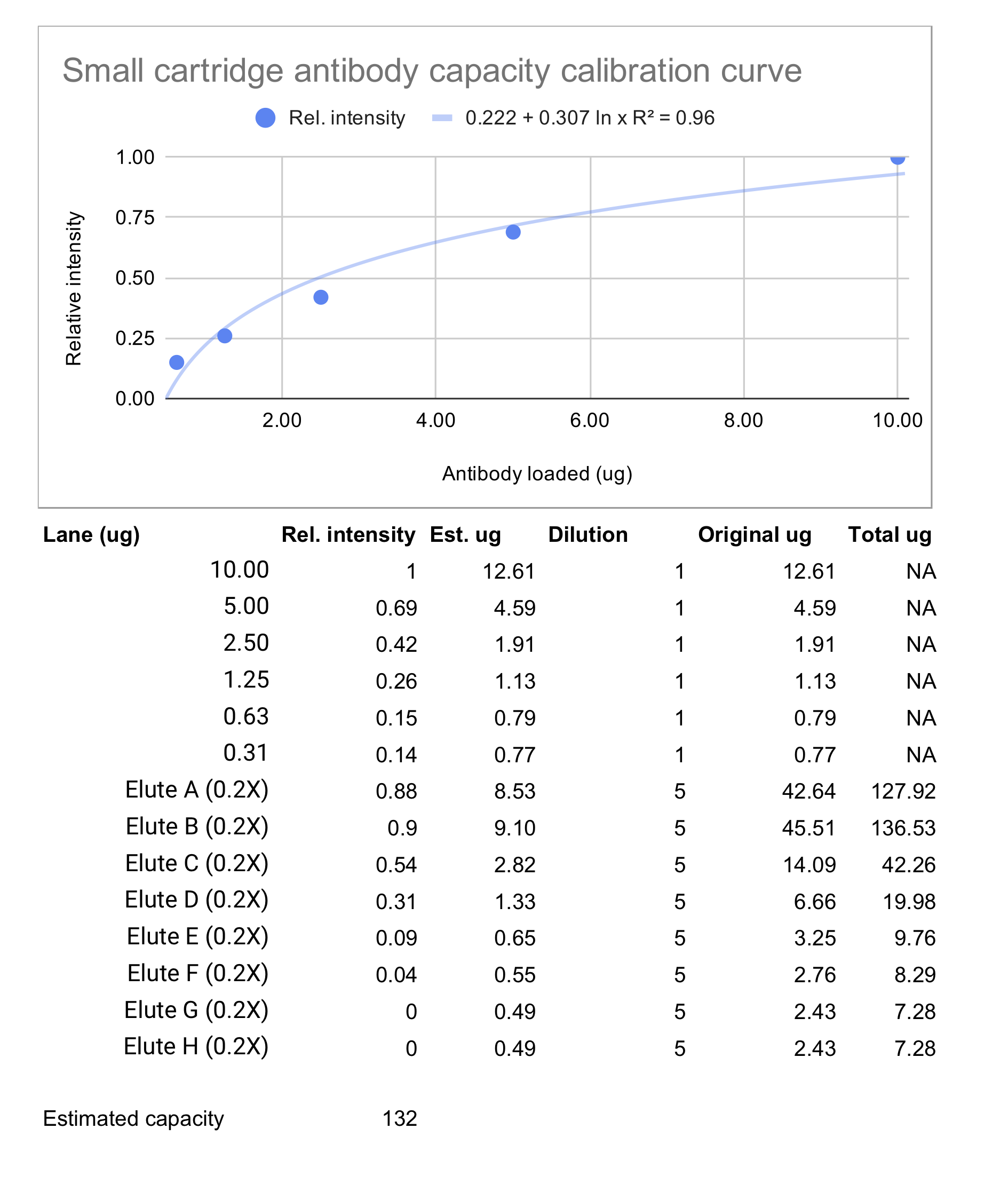

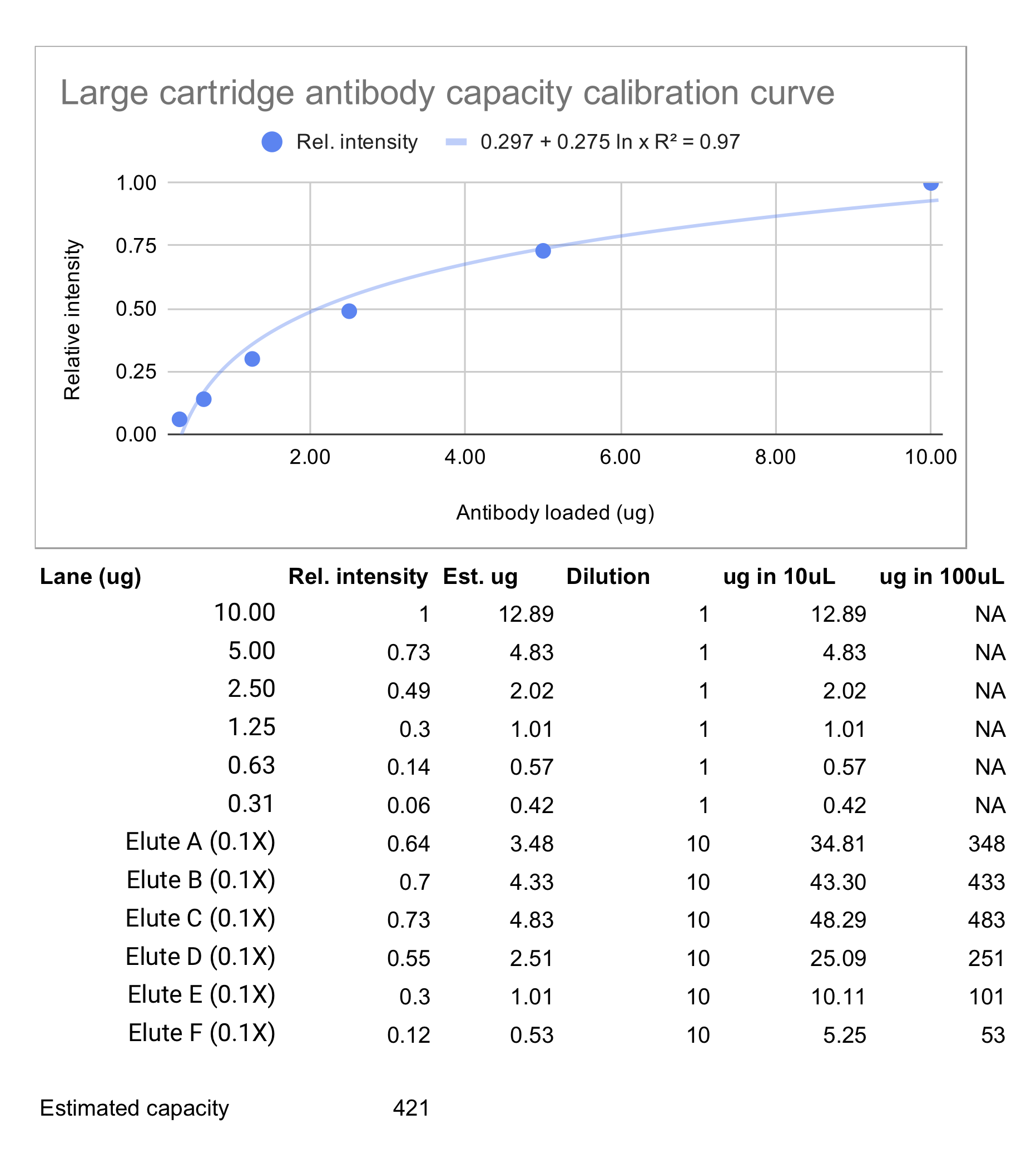

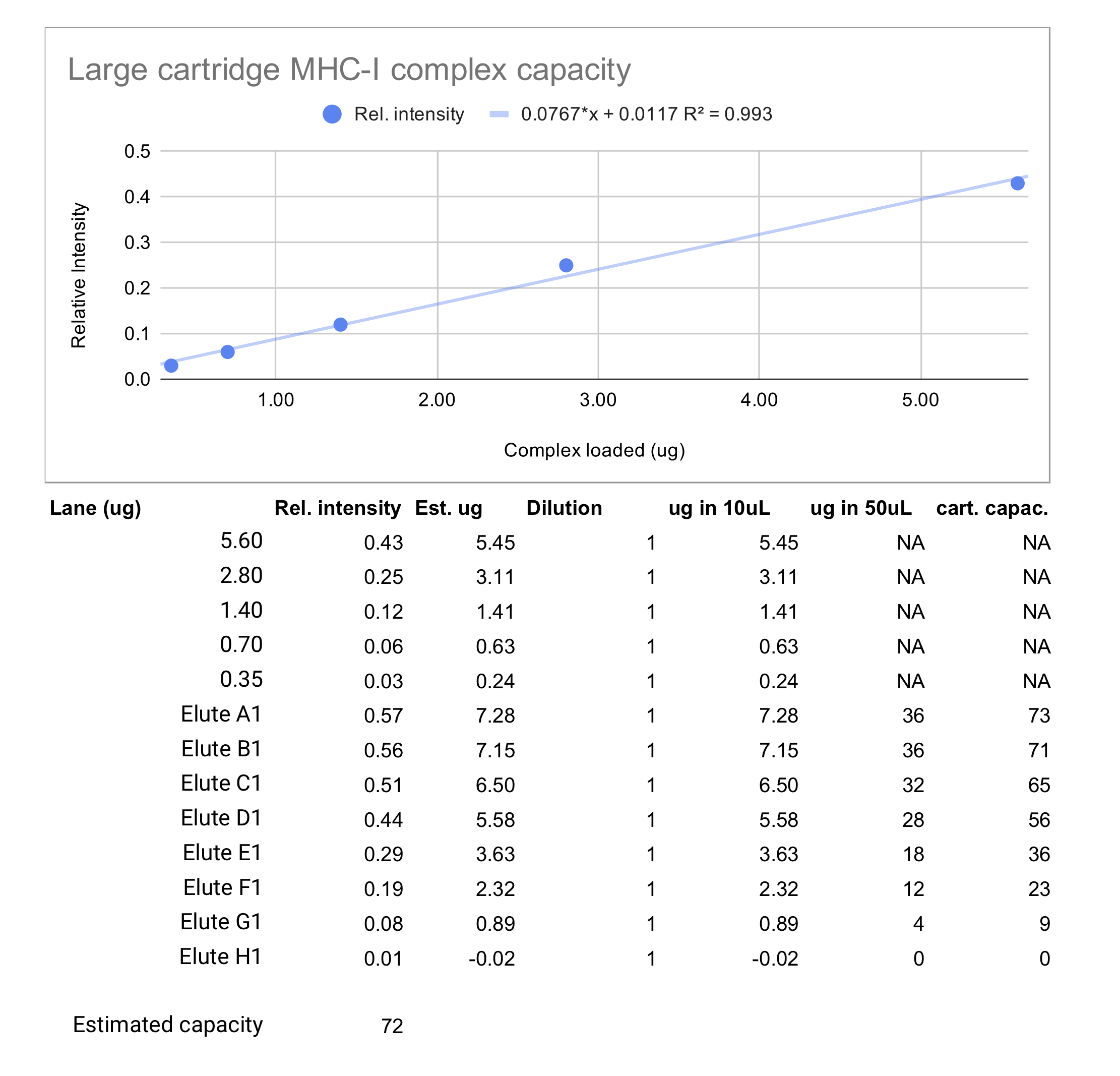

**Fig. S2 - Cartridge characteristics: the effect of cross-linking on MHC-I pulldown, and antibody specificity for dissimilar alleles**

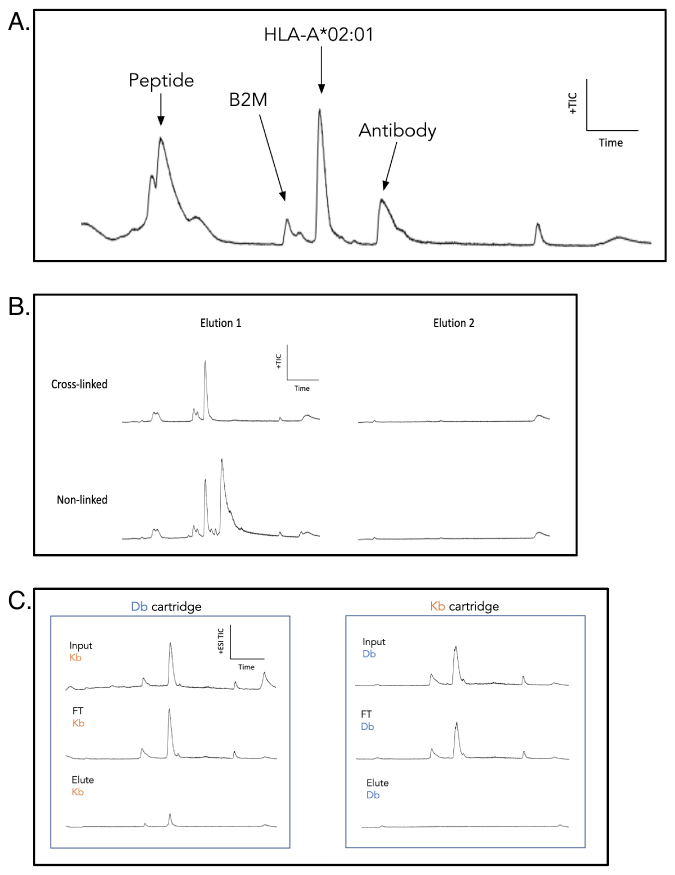
(A) Control run showing at what retention time doped in peptide, recombinant HLA and ß2M, and antibody elute on LCMS. (B) Comparison of cross-linked and non-cross-linked MHC-I enrichment showing that cross-linking has no apparent effect on enrichment. A second elution shows that cross-linking also does not lead to additional non-specific binding. (C) Test of non-specific binding of MHC complex to different cartridges. Kb complex shows some non-specific binding to an anti-Db antibody cartridge, while Db complex shows no non-specific binding to an anti-Kb antibody cartridge.

**Fig. S3 - Lysate viscosity assay using filtrate retained on top of a Costar 0.45 µm spin filter**

Analysis of the viscosity of lysate based on volume retained on top of a spin filter. Under the conditions described (500 µL pipetted onto a 0.45 µm Costar filter and spun at 16,000 x g for 1 min at 4°C), water and lysis buffers show no volume retained (negative control), while room temperature storage, day-old MC38 lysate (50M cells per mL of B-buffer, positive control), shows retention of over 400 µL. Fresh GRANTA lysate (50M cells per mL of B-buffer) shows no retention, while 4 h at room temperature (typical, un-optimized loading condition), shows a major increase in viscosity. Lowering the temperature to 4°C reduces retention in a manner that is maintained over 18 h. If lysate is flash frozen with PBS, sucrose (1 M starting, 200 mM final), or glycerol (50% glycerol starting, 10% final) and then thawed, viscosity is low, even after 4 h at room temperature (however 24 h at room temperature leads to a large increase in viscosity). Combining sucrose and glycerol additives (leading to an even more dilute lysate solution) allows for a freeze thaw with no apparent retention immediately, or after 4 h at room temperature.

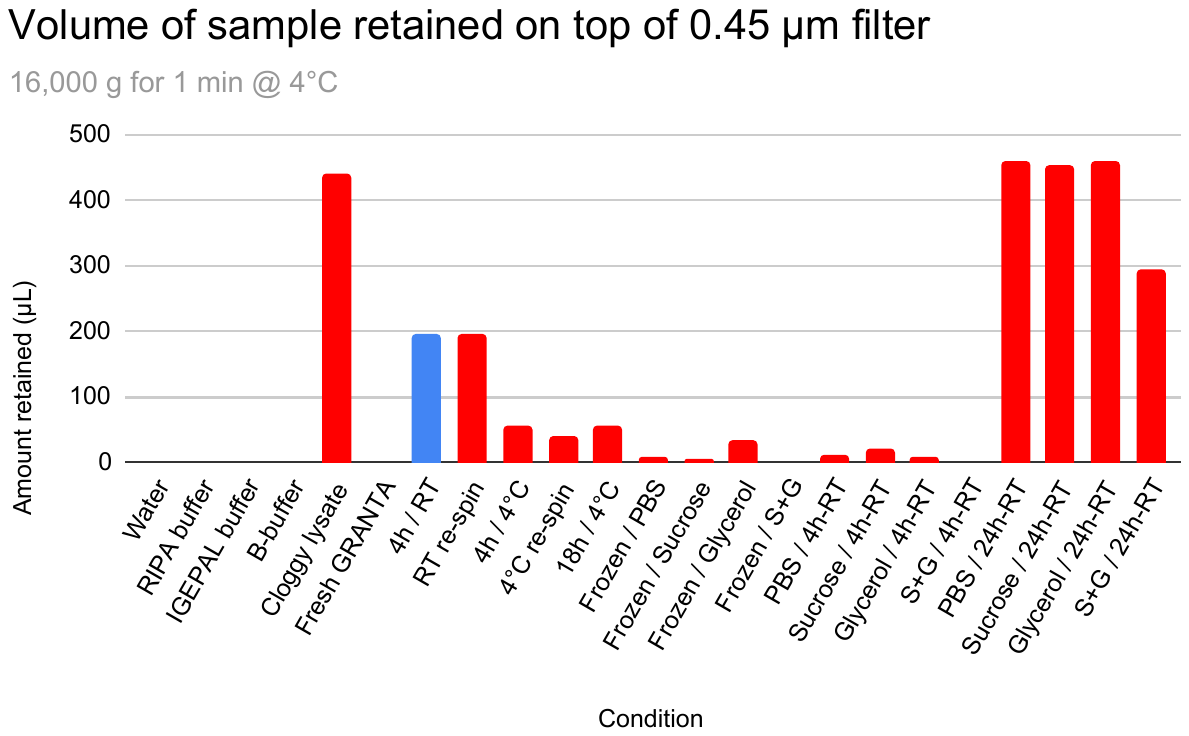

**Fig. S4 - Abundance corrected composition overlap between cartridges, uses, and wet vs. dried-then-rewetted cartridges**

Venn diagrams were used to show the overlap in composition between different cartridge enrichments of the same batch of GRANTA lysate. Peptides in each data set were weighted for abundance using total ion chromatogram (TIC) signal (so that far more abundant peptides contributed more to the calculated overlap) but were not corrected for abundance (so that different amounts of signal between runs was preserved in the calculation). Comparisons made here include between wet and dried-then-rewetted cartridges, between replicate cartridges of the same type, and over re-uses.

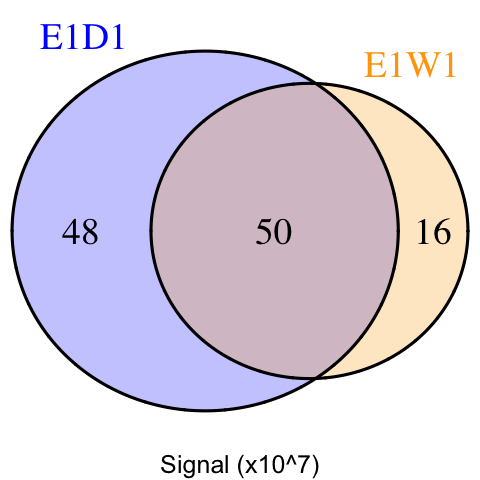

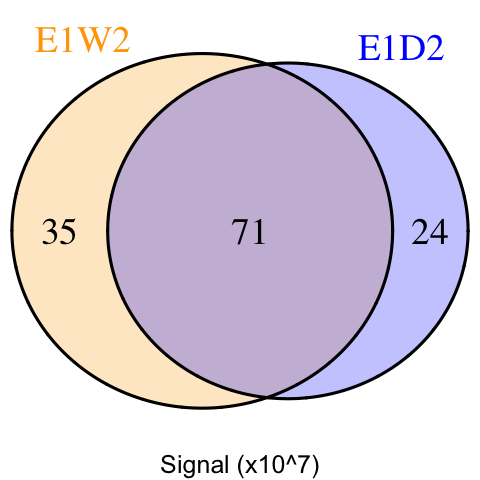

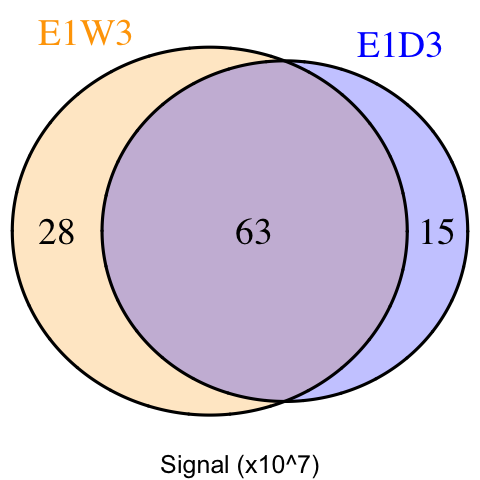

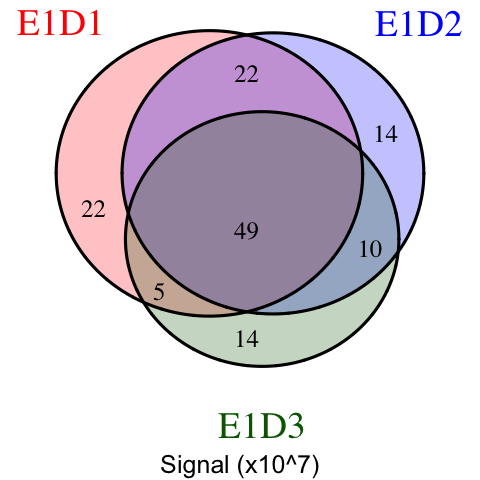

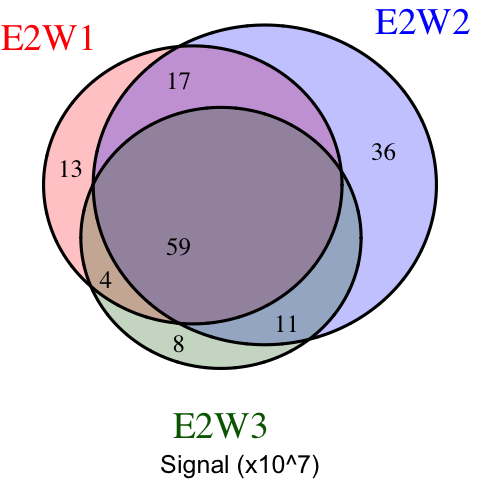

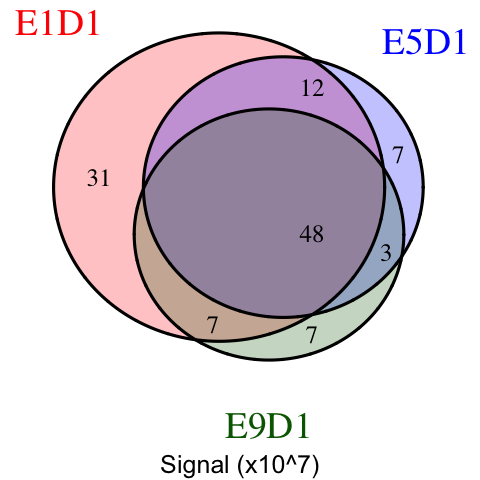

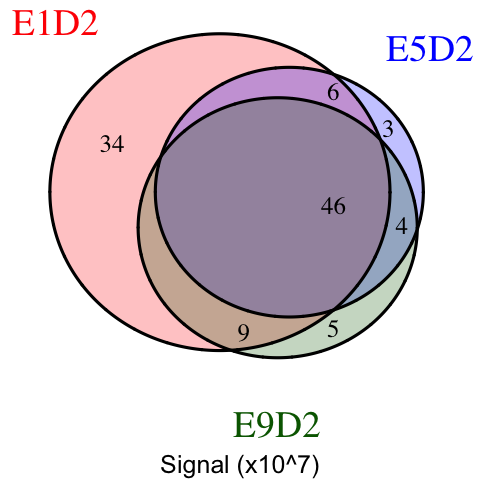

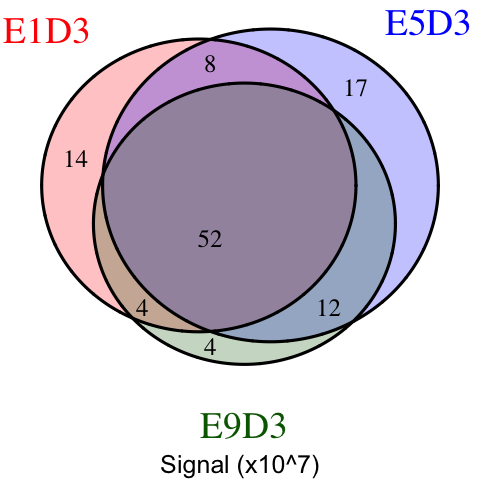

**Fig. S5 - Optimization of preferential peptide elution from a C18 cartridge under different % acetonitrile conditions**

LCMS TIC chromatograms showing input, flow-through, and eluates after washing with different % acetonitrile solution (% acetonitrile in water, 0.01% formic acid). Almost all peptide is absent from the flow through, with some protein carryover. Peptide is preferentially eluted from 30% acetonitrile, matching recommendations in the literature.

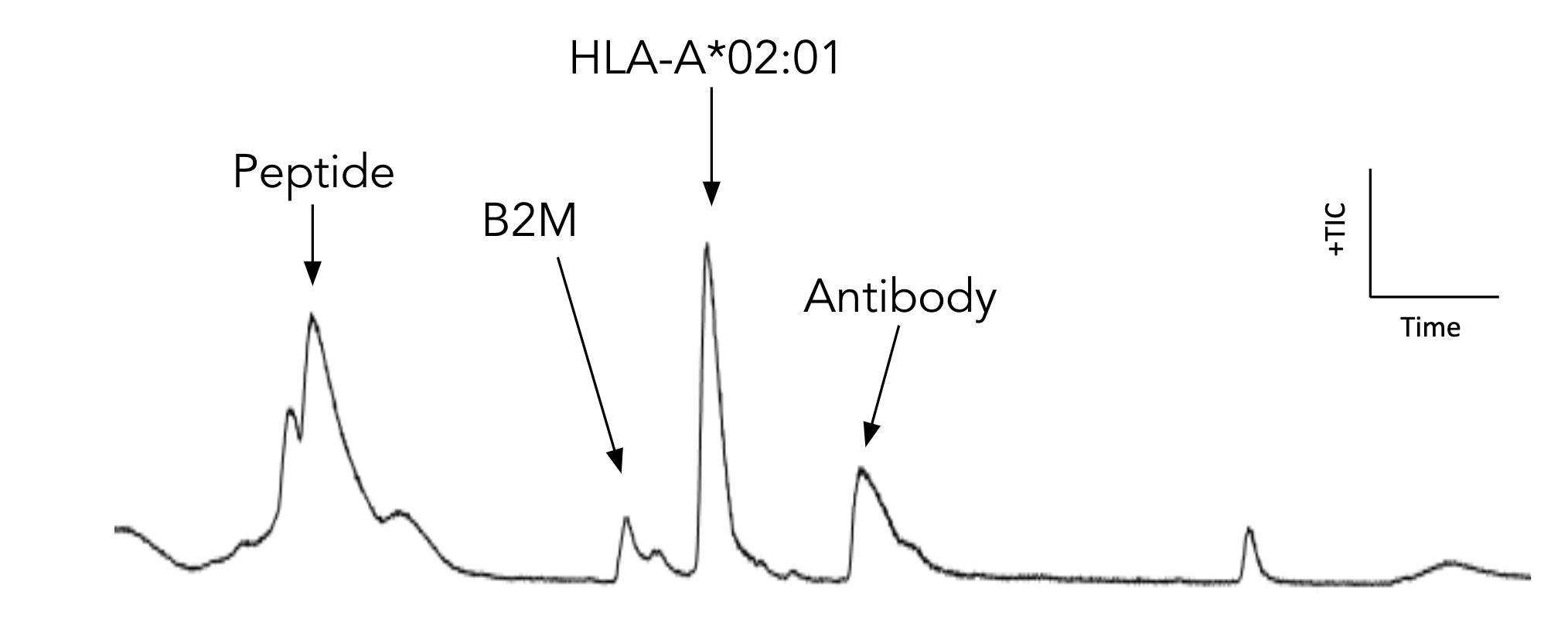

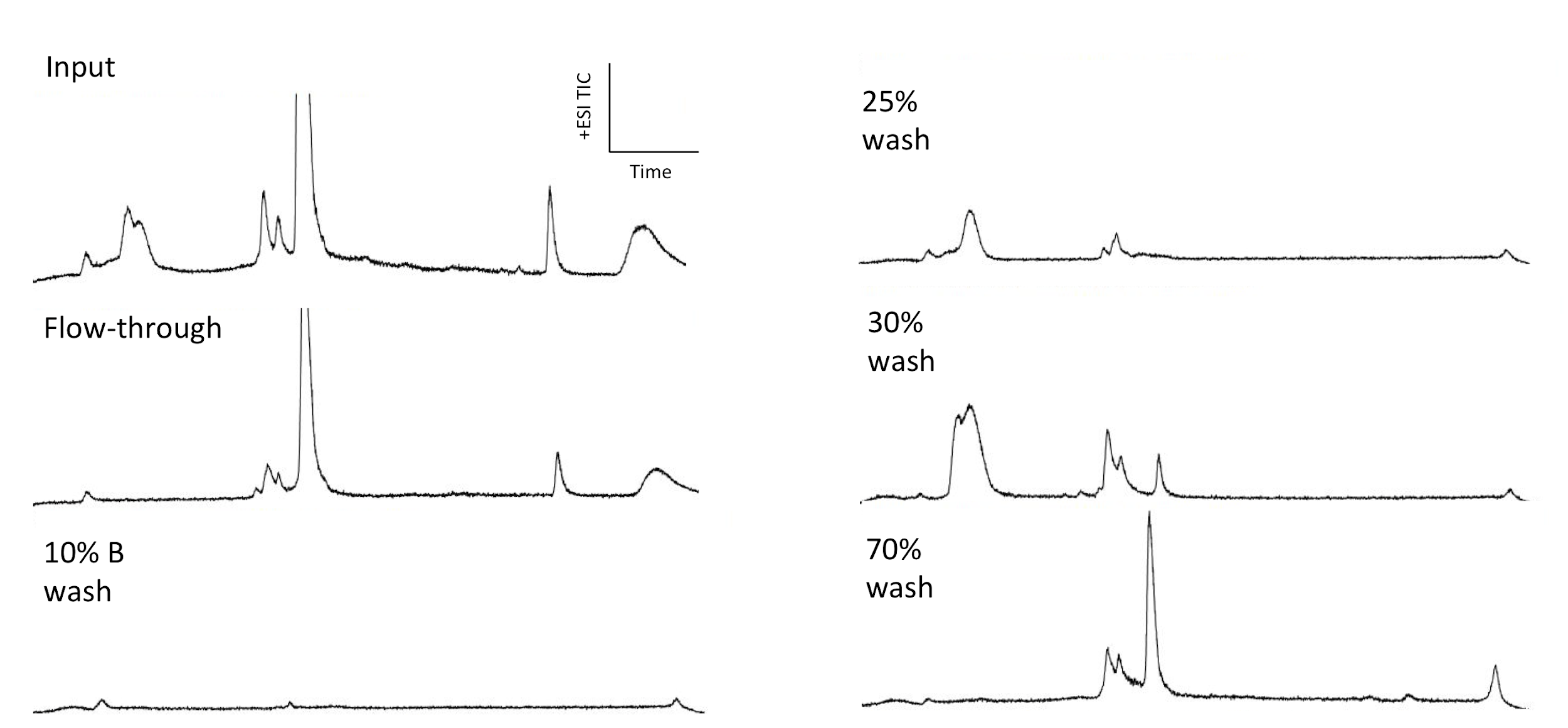

**Fig. S6 - Efficiency of peptide modification on C18 cartridges**

Modification efficiency of N-termini with TMT or cysteines with carbamidomethyl among NEO223 peptides modified on-C18-cartridge under the “desalt” condition (no reduction-alkylation with TCEP-iodoacetamide, no TMT treatment), the “TMT” condition (TMT treatment only), or the “RAT” condition (reduction-alkylation and TMT treatment). Note the y-axis scale is from 0.90 to 1.00.

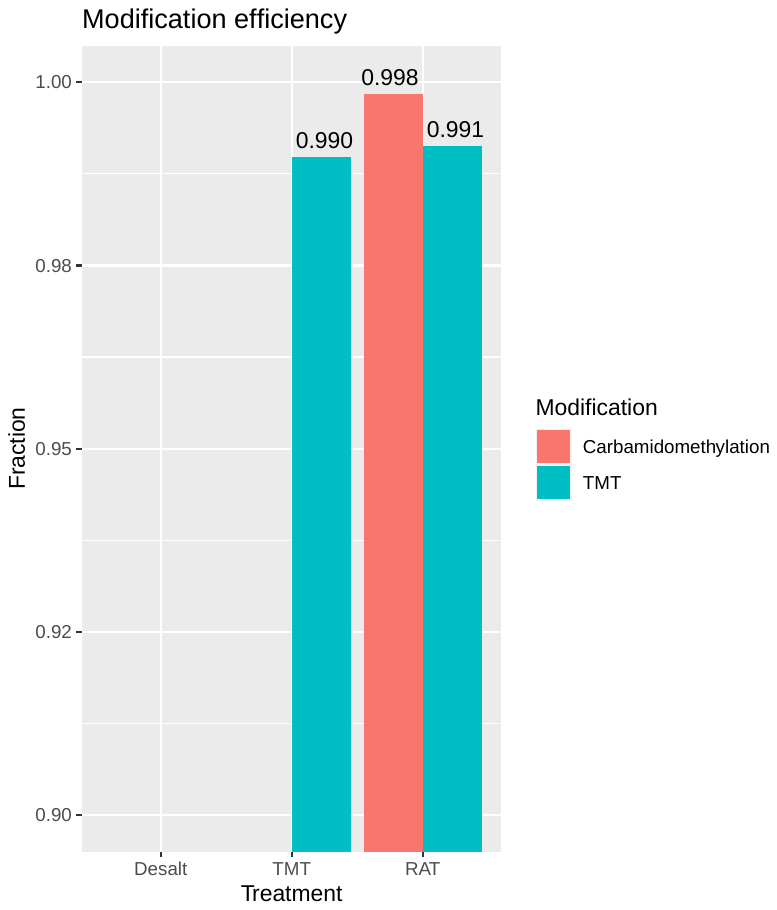

**Fig. S7- TOMAHAQ optimization: CID vs. HCD for Adpgk peptides**

Both collision-induced dissociation (CID) and higher-energy collisional dissociation (HCD) fragmentation was tested in the TOMAHAQ method to determine which produced higher signal and a greater number of fragment ions on the MS2 level for two Adpgk neoepitopes, ASMTNMELM (top panels) and ASM(Ox)TNMELM (bottom panels). CID and HCD produced similar fragmentation patterns for ASMTNMELM, however ASM(Ox)TNMELM had an apparently cleaner spectrum with higher maximum signal using CID. Method development and data analysis were carried out in TomahaqCompanion as described.

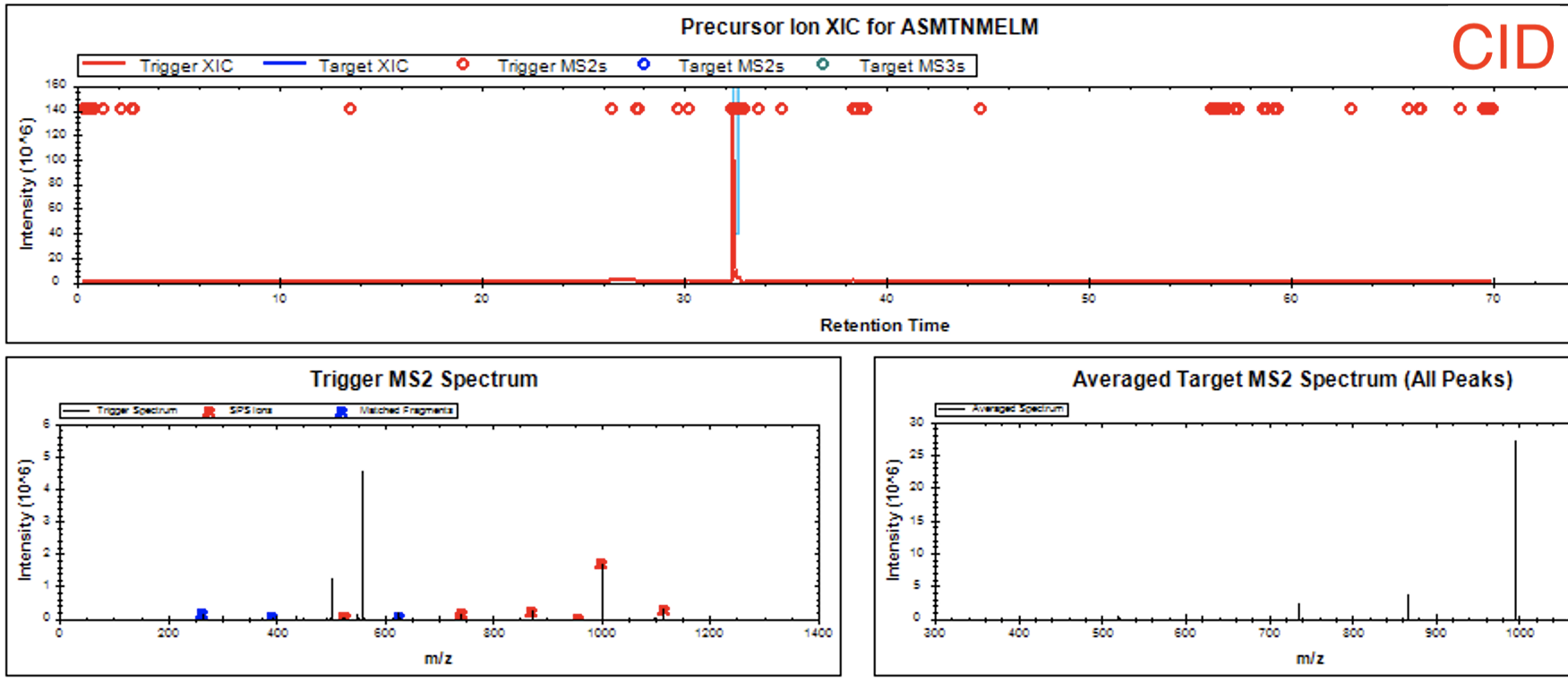

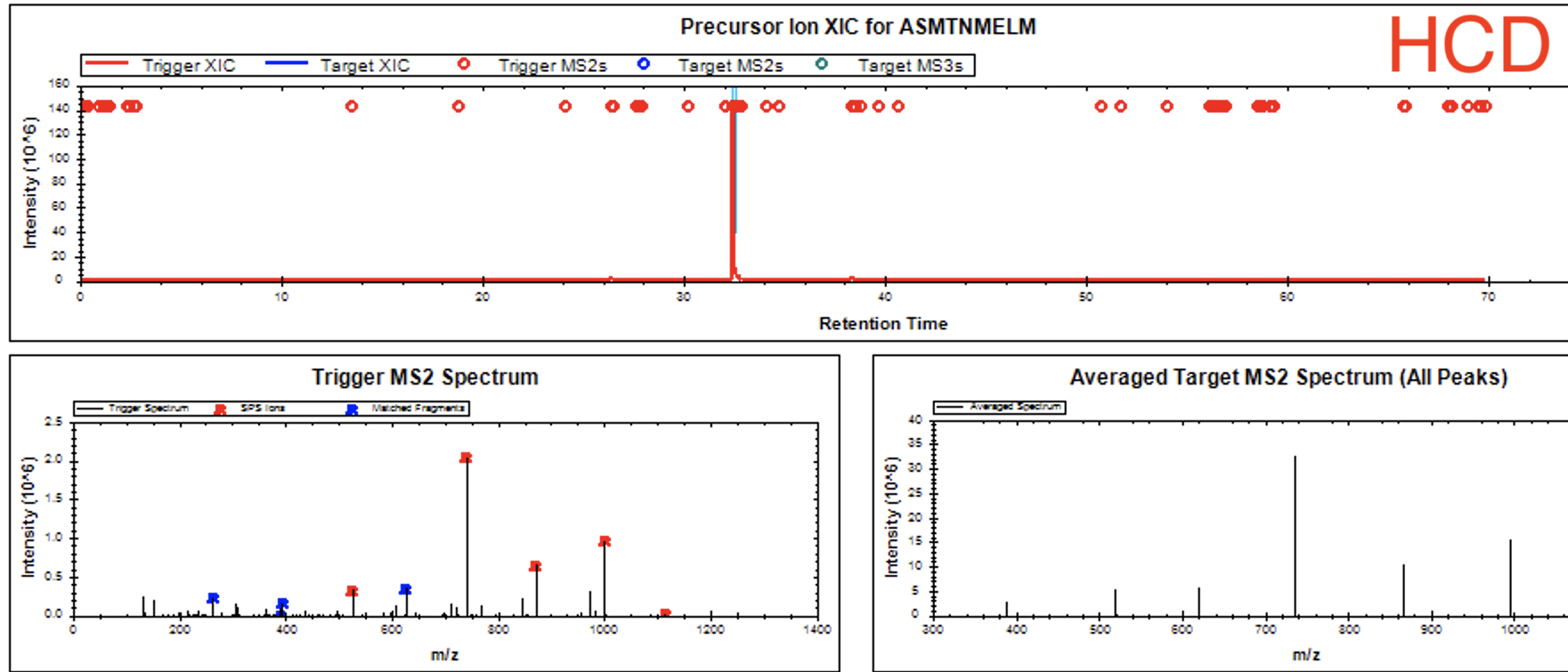

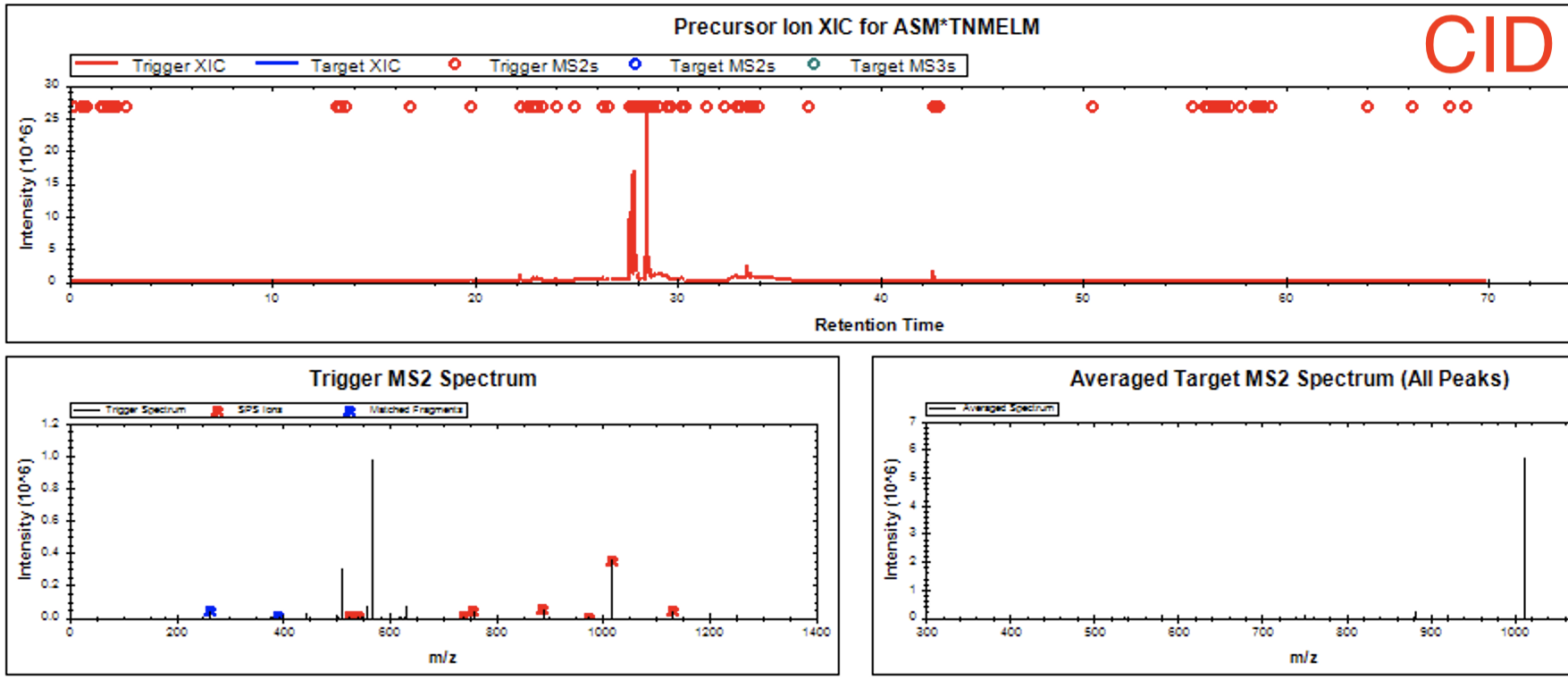

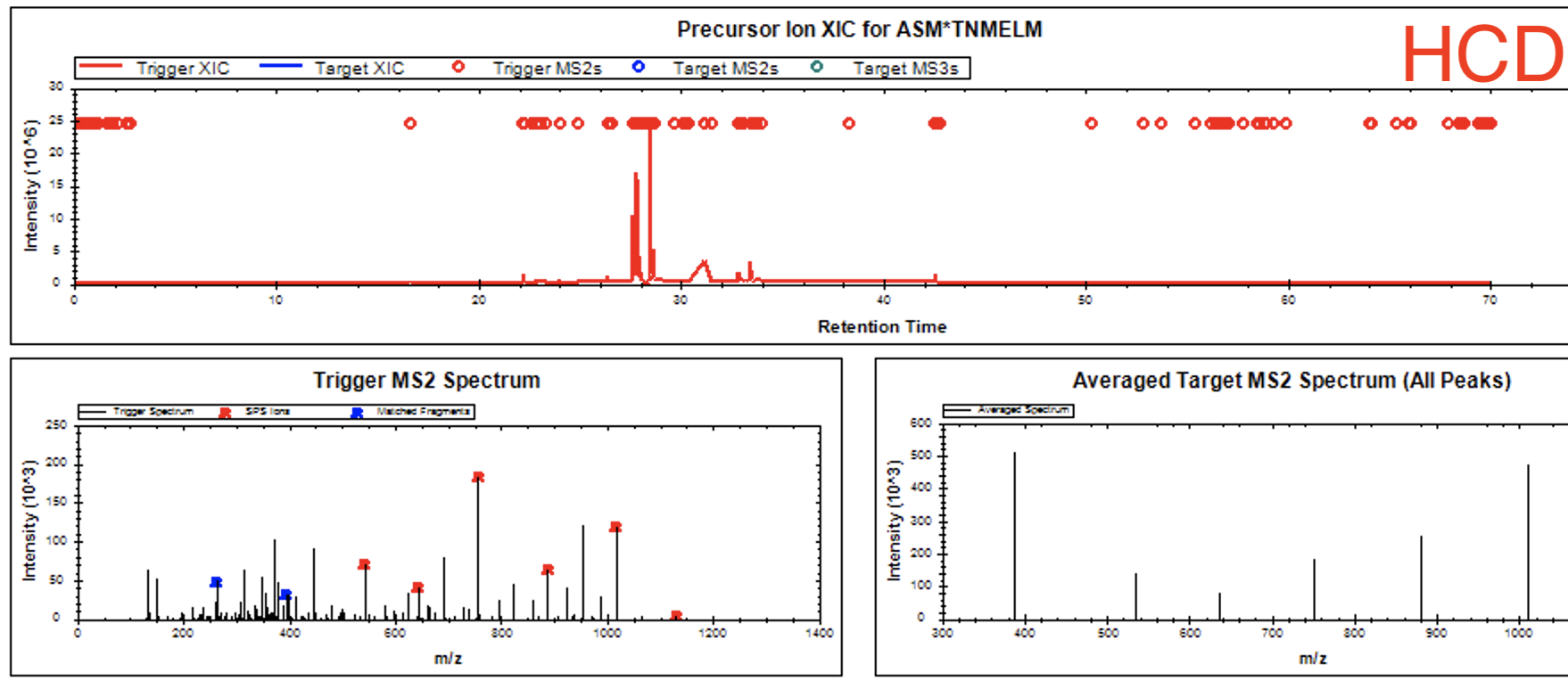

**Fig. S8 - TOMAHAQ optimization: CID vs. HCD comparisons across target number and run length for the NEO223 mixture**

Optimization of additional parameters within the TOMAHAQ method using the NEO223 synthetic mix, with the following syntax on the x-axis: TARGETS_FRAGMENTATION_GRADIENT. In the first two conditions (red and brown bars, far left), the previously published seven neoepitopes (PUB7) only were targeted (using the NEO223 synthetic mixture) with a 40 min 2-42 %B gradient, which led to the detection of 6 targets within the synthetic mix under CID and 5 targets under HCD fragmentation. Changing the number of targets to all 223 neoepitopes and oxidized forms with the same 40 min gradient led to almost identical CID and HCD detection at 159 and 161 targets respectively. Increasing the length of the gradient to 80 min caused a decrease in the number of targets detected, especially for the CID fragmentation condition. Because of this data and the Adpgk fragmentation results above (supplemental Fig. S7), we chose to use a 40 min, CID gradient.

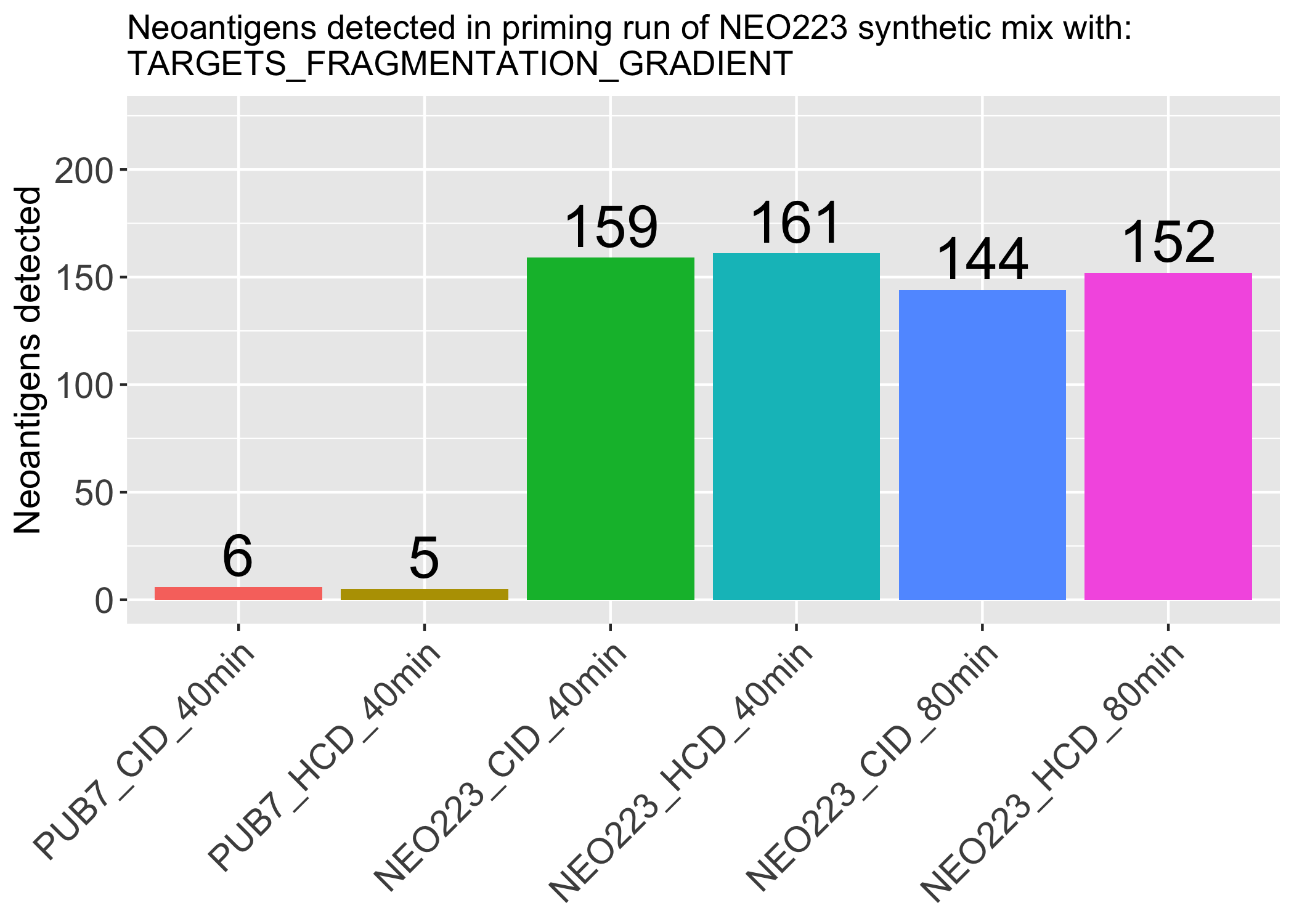

**Fig. S9 - Limit of detection and quantitativeness of NEO223 targets determined using Tomahaq vs. MS2 against a synthetic target mixture**

Sensitivity and quantitativeness of TOMAHAQ by neoantigen across decreasing synthetic, multiplexed target mixture concentrations. The size of each dot corresponds to extracted ion chromatogram (XIC) signal for respective methods, while the color corresponds to the percent coefficient of variation. The results using the TOMAHAQ-MS method are shown in the top panel, while results for the untargeted MS2 method are shown in the bottom panel.

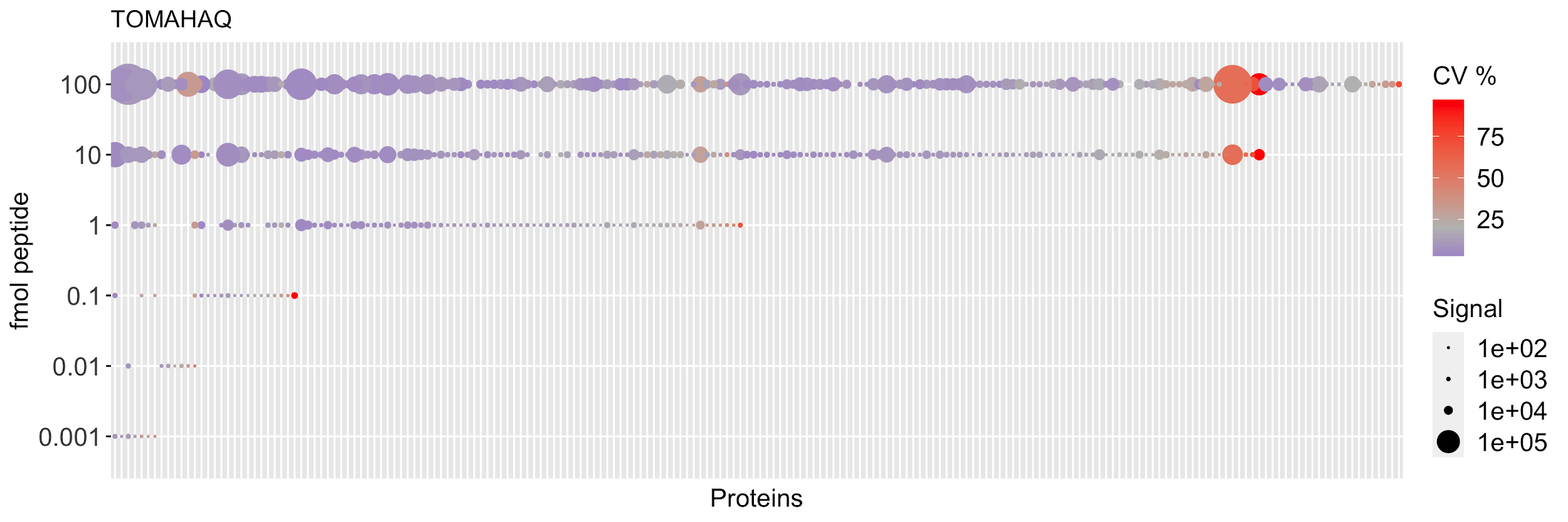

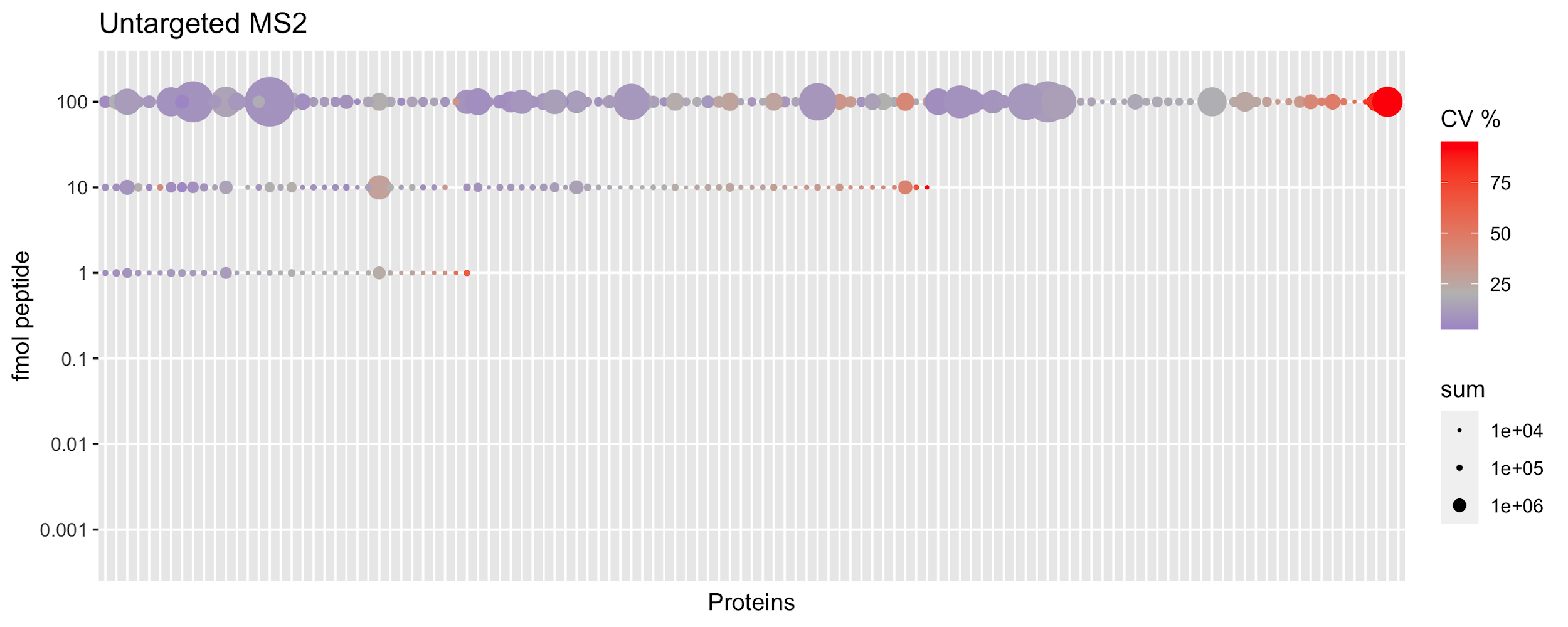

**Fig. S10 - idAdpgkG engineered cell line validation**

We confirmed the expected protein abundance changes within the MC38-idAdpgkG engineered cell line via GFP fluorescence in flow cytometry and anti-Adpgk western blot. In the left panel, flow cytometry shows that the 488 channel signal increases upon treatment with dox, then decreases upon dox+dTAG treatment, consistent with increased expression of the GFP-containing construct, followed by its degradation with dTAG treatment. In the right panel, western blot intensity for anti-Adpgk antibody (Abcam, ab228633) corrected for actin abundance (CST, 3700) under different conditions is shown for the mutant allele construct (right side) and wild-type endogenous allele (left side). The wild-type Adpgk allele is also present in the MC38-idAdpgkG engineered cell line, and is distinguishable from the larger mutant construct on western blot, so its relative abundance is also shown and does not differ significantly between conditions, as expected.

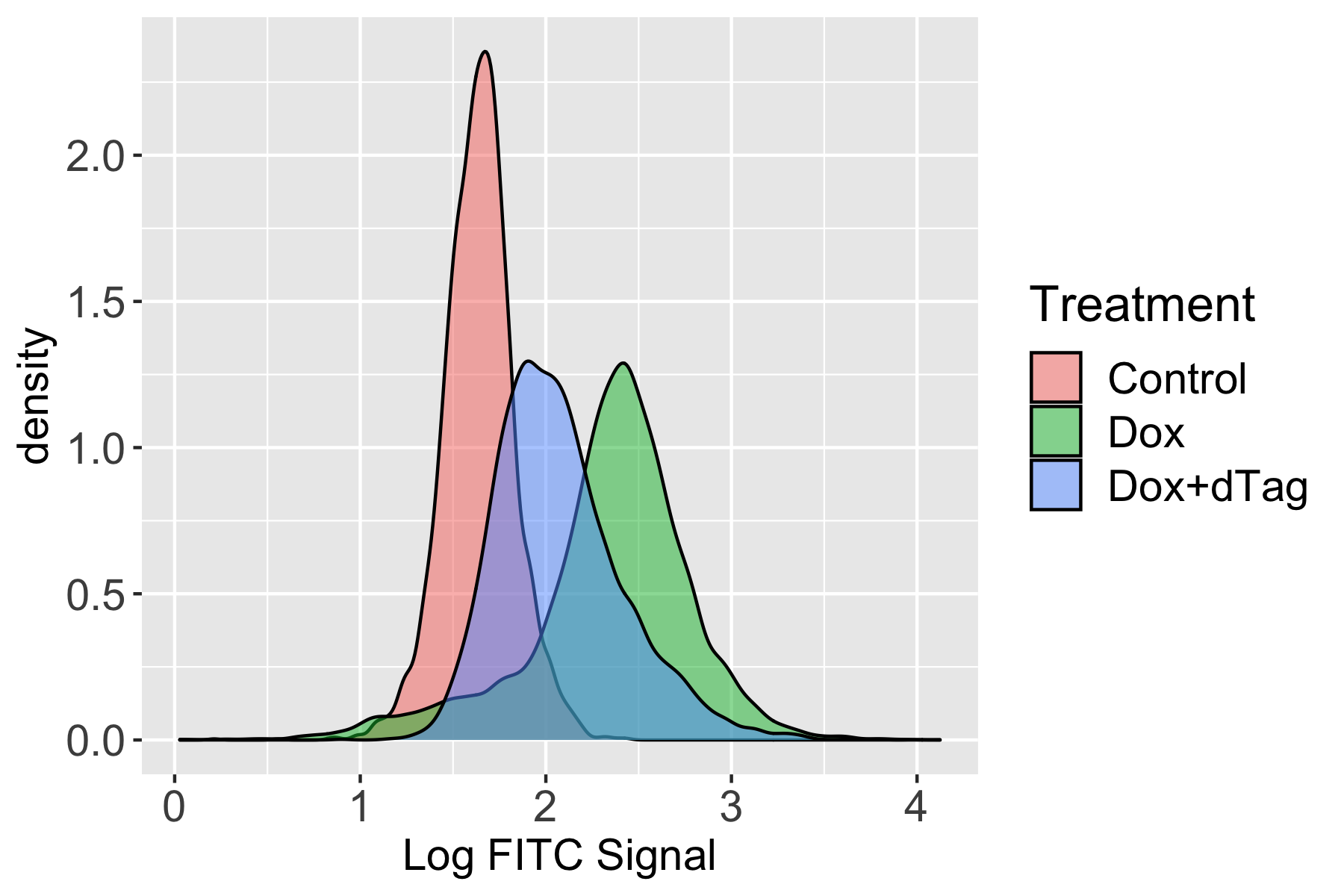

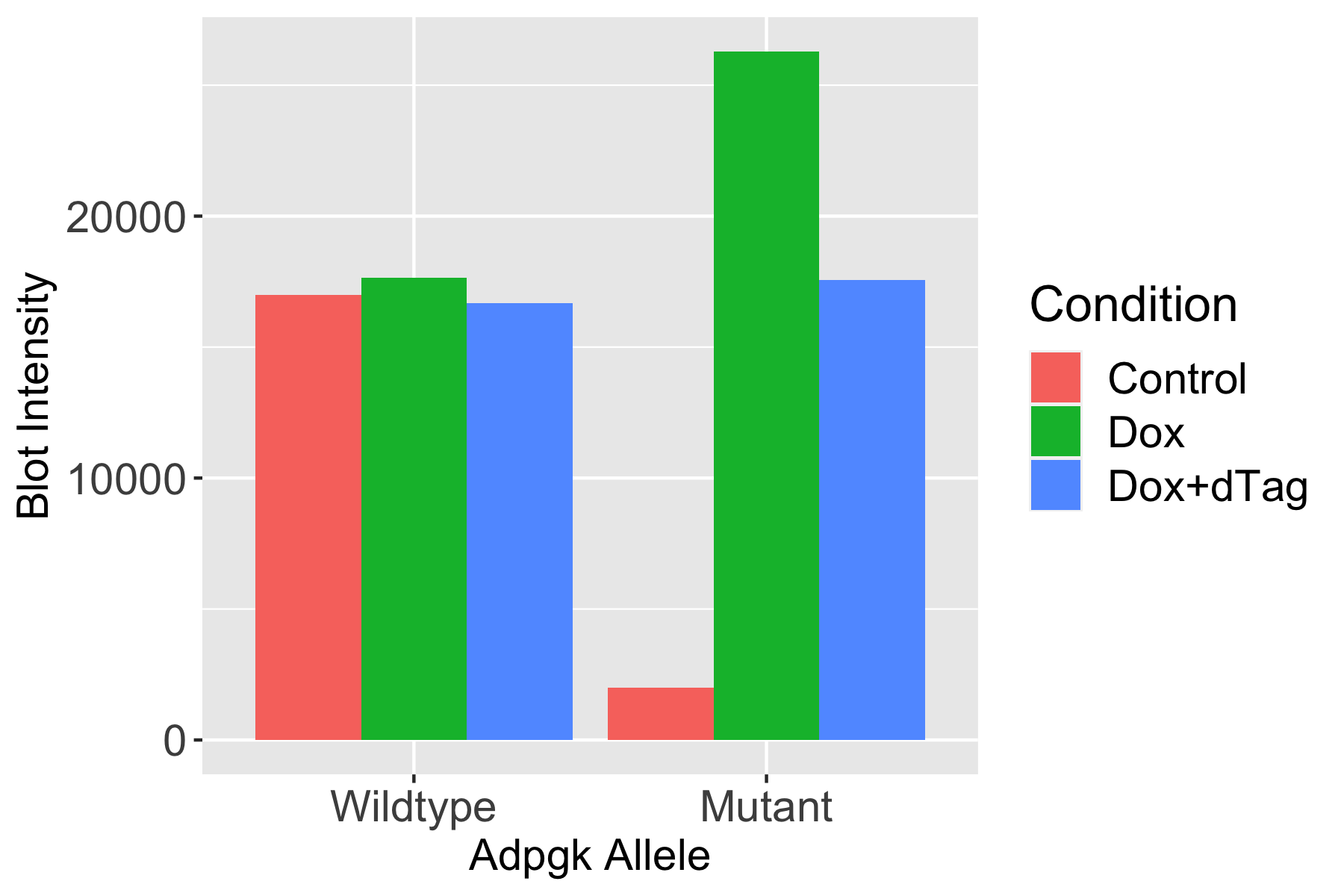

**Fig. S11 – idAdpgkG tGFP, MHC-I, and MHC-II signal abundance under 4plex (-/+ dox, -/+ dTAG) conditions**

The geometric mean fluorescence intensity (gMFI) for turboGFP, or antibody-stained MHC-I or MHC-II (see methods), after 3 h or 22 h, under -/+ doxycycline, -/+ dTAG, and -/+ IFNg conditions.

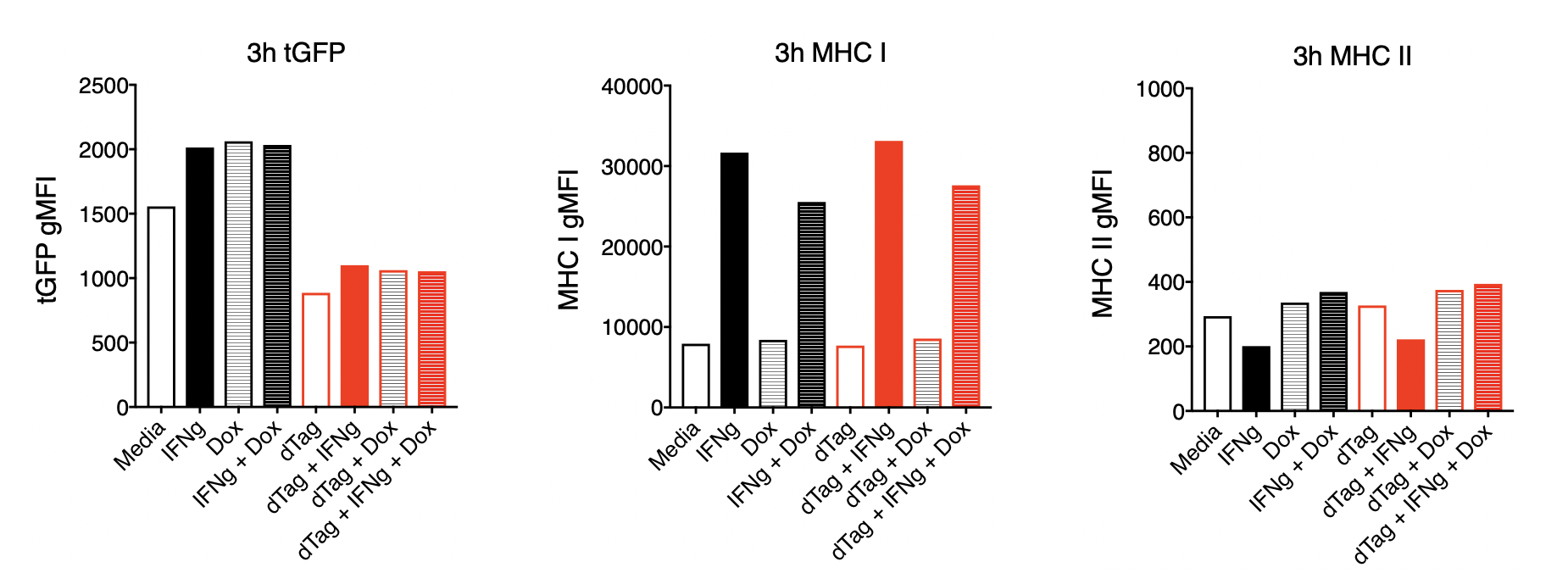

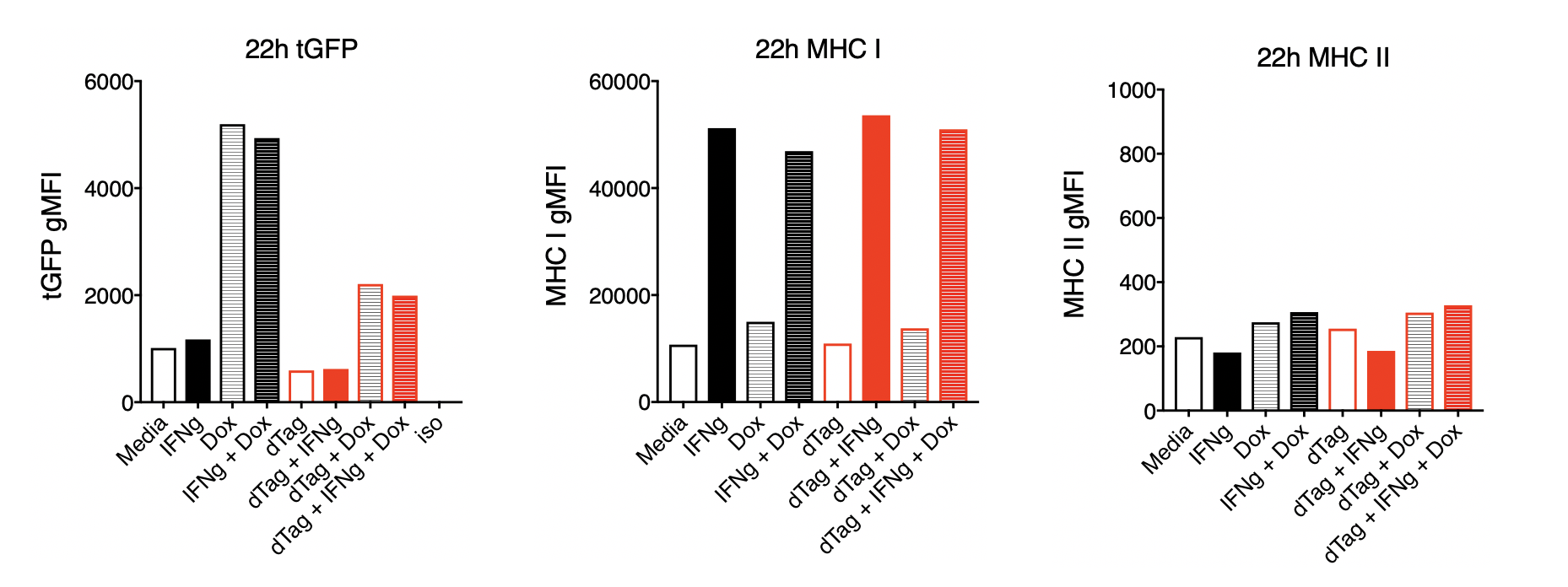

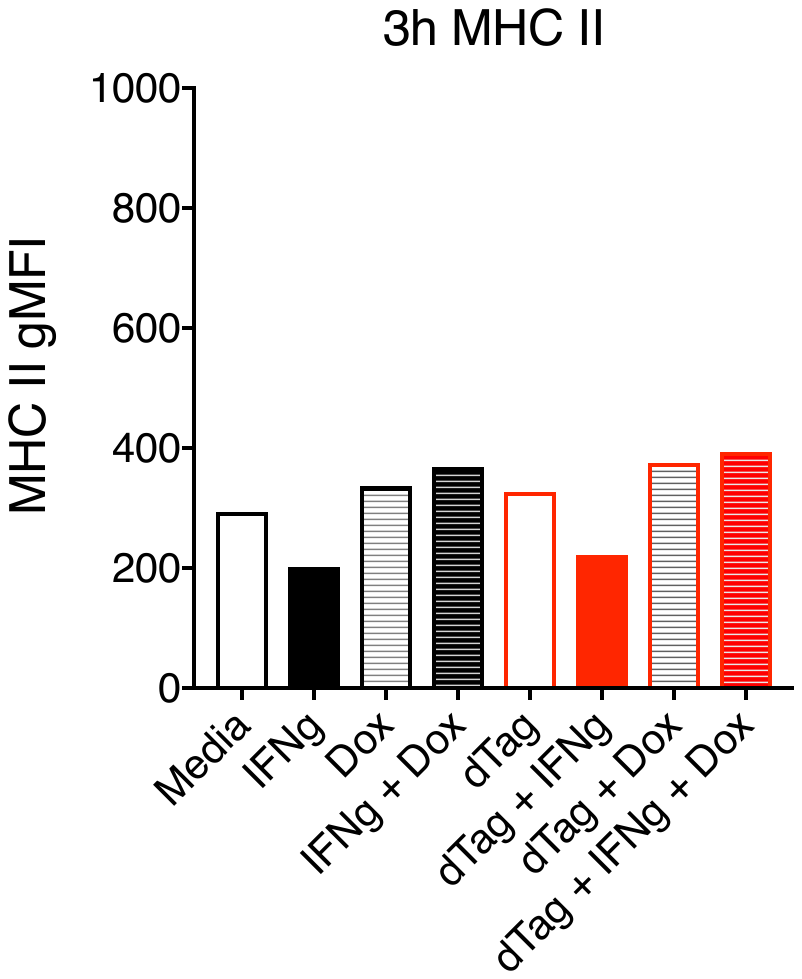

**Fig. S12 – Visual inspection of spectral quality using the TomahaqCompanion interface**

Presented are four examples of spectra of differing quality, all of which passed signal over noise filters (see methods). The Gtf2i spectrum shows a number of different SPS ions, most of which do not have much noise around them, and the signal is good (excellent). The Act1rb spectrum has more than one SPS ion without much noise around the peaks, although the signal is fairly weak (good). The Als2cr4 spectrum has only one SPS ion, but there is not much noise around it and the signal is decent (adequate). The Fat1_OX spectrum does not have a visible SPS ion by TomahaqCompanion (its SPS ion is lost in the noise) and its signal is low and noisy (bad).

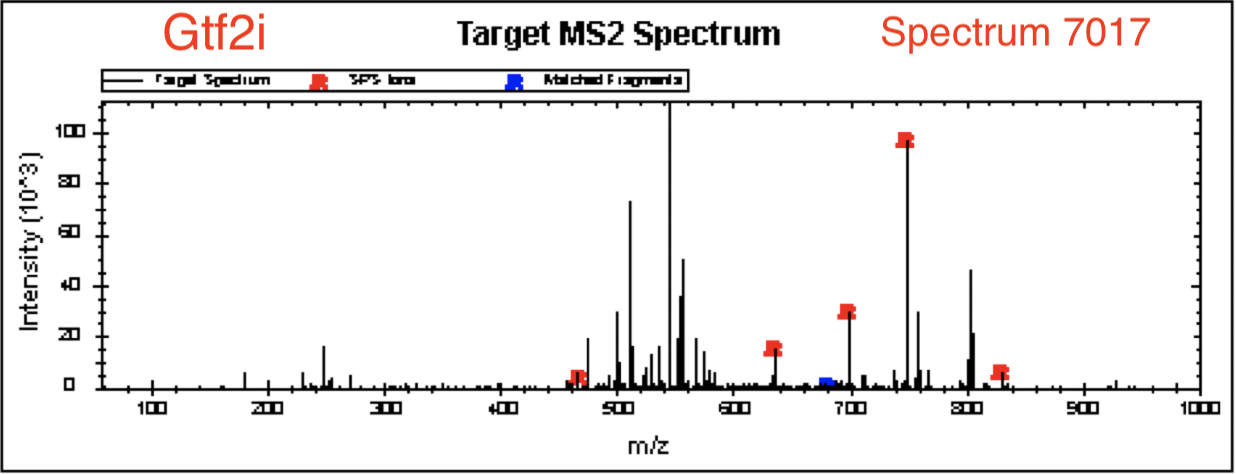

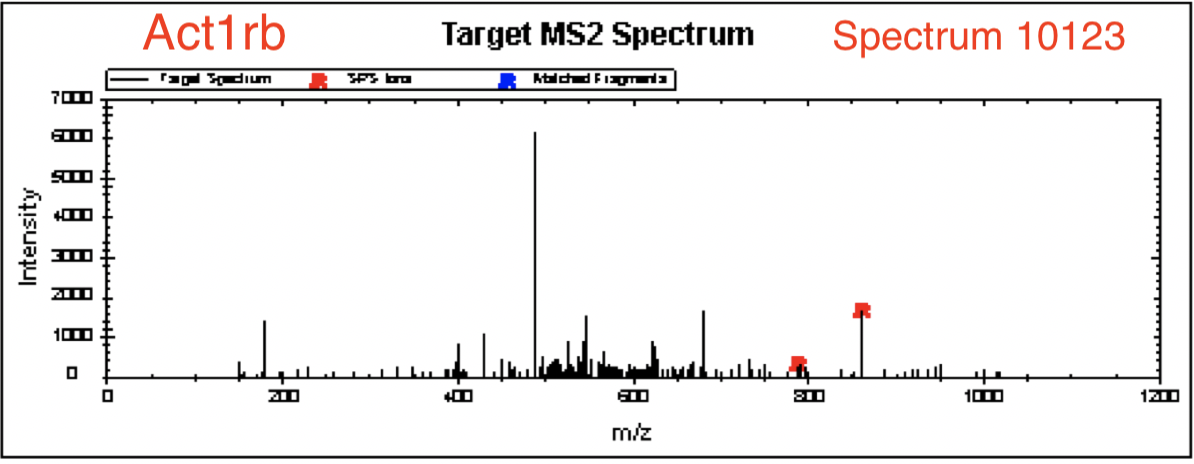

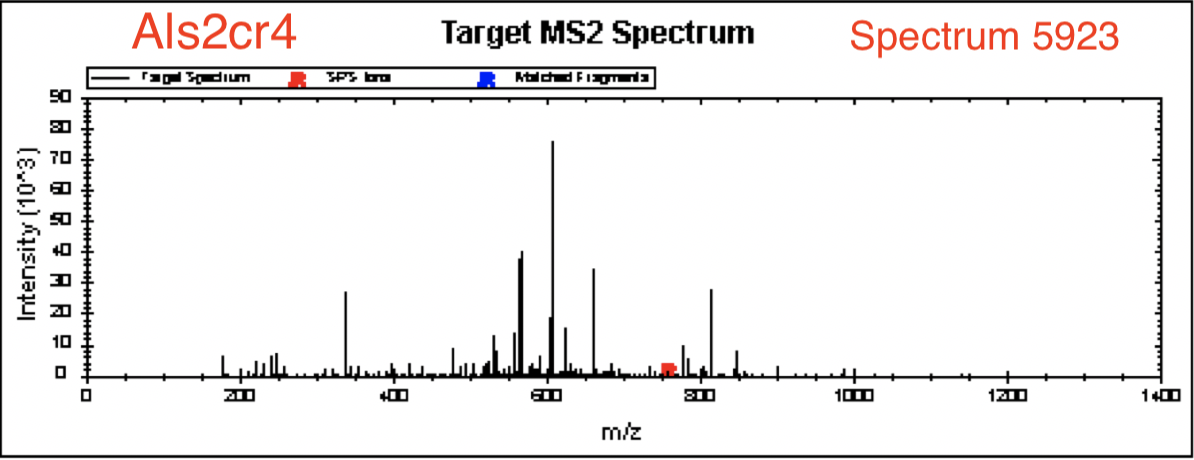

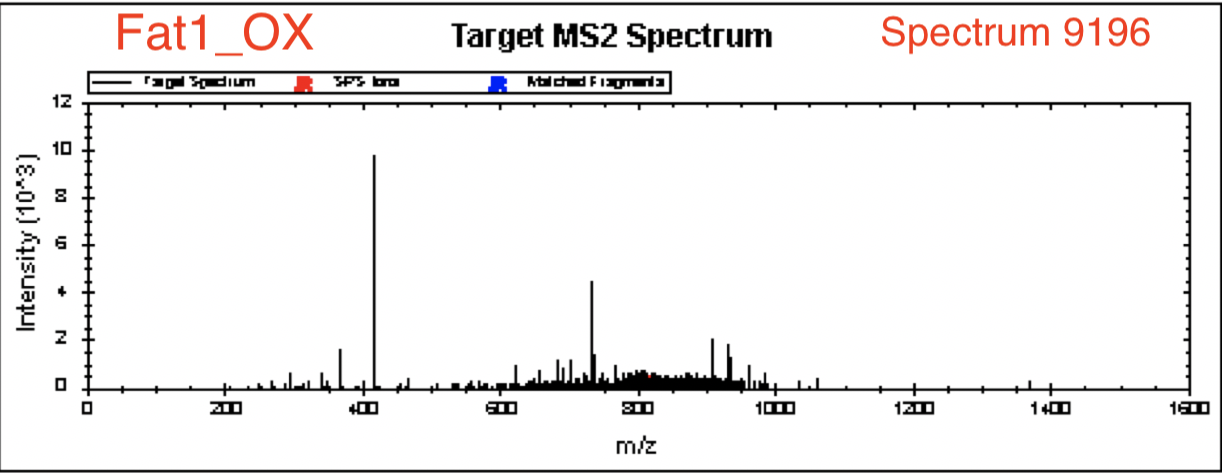

**Fig. S13 - Global multiomic heatmap comparisons**

Untargeted transcriptomics, proteomics, and peptidomics was performed on MC38-idAdpgkG cells and those proteins detected on all three omic levels were quantified (111 total, including Adpgk). Log2 changes in abundance were calculated between conditions compared to the control condition and normalized to dTAG treatment alone. Proteins were then ordered based on dox transcriptomic changes. Very little change, aside from Adpgk transcript expression, is observed on the transcript level. Subtle changes in abundance both up and down are observed on the proteomic level. Increases in abundance on the peptide level are broadly observed, even for those transcripts and proteins exhibiting little change.

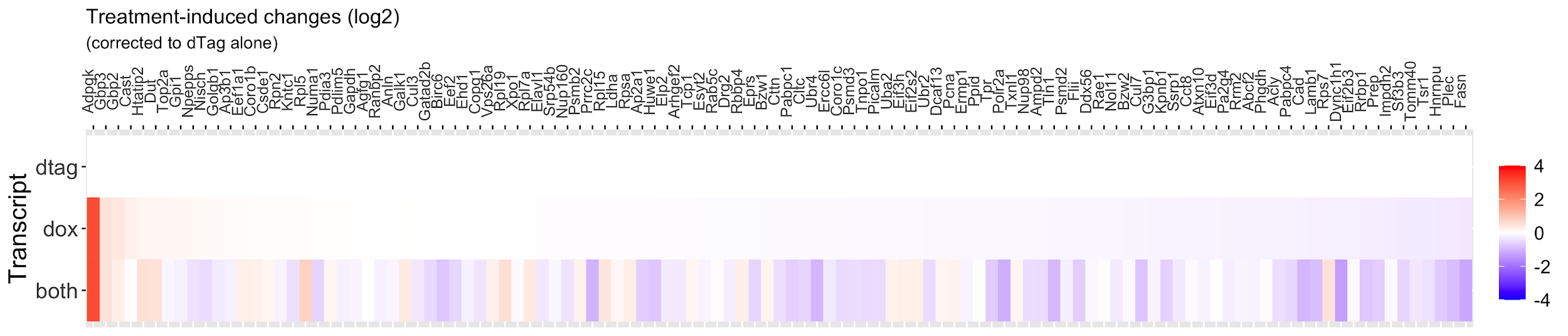

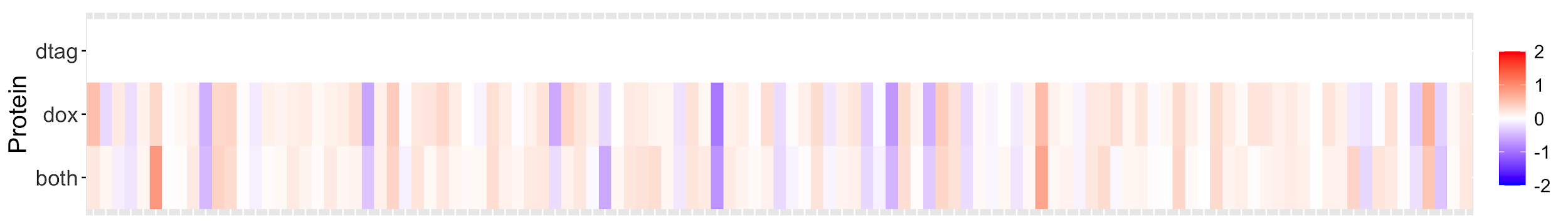

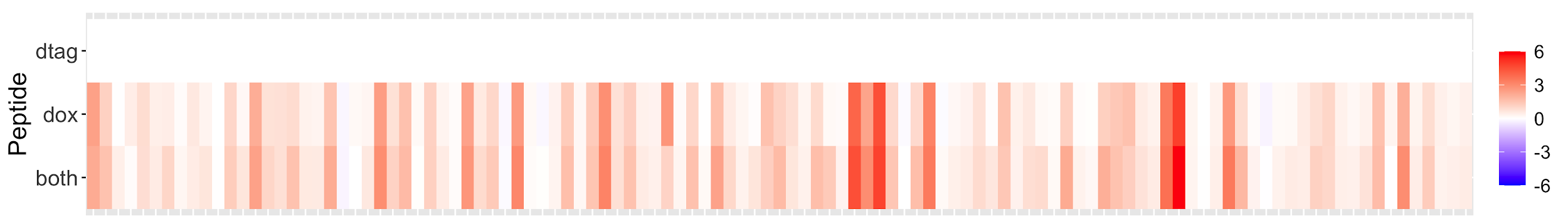

**Fig. S14 – Doxycycline vs. Both (dox+dTAG) untargeted surface peptidome changes**

Untargeted MHC-I peptidomics was performed as described in Fig. 4E. The dox vs. dox plus dTAG abundances of detected peptides is shown here, to demonstrate that the additive effects of dox and dTAG on the surface peptidome are small, compared to changes induced by dox or dTAG upon the control condition (IFNg only) as shown in Fig. 4E. Peptides with > 2-fold upregulation are colored red and labeled. Units on both axes represent signal intensity.

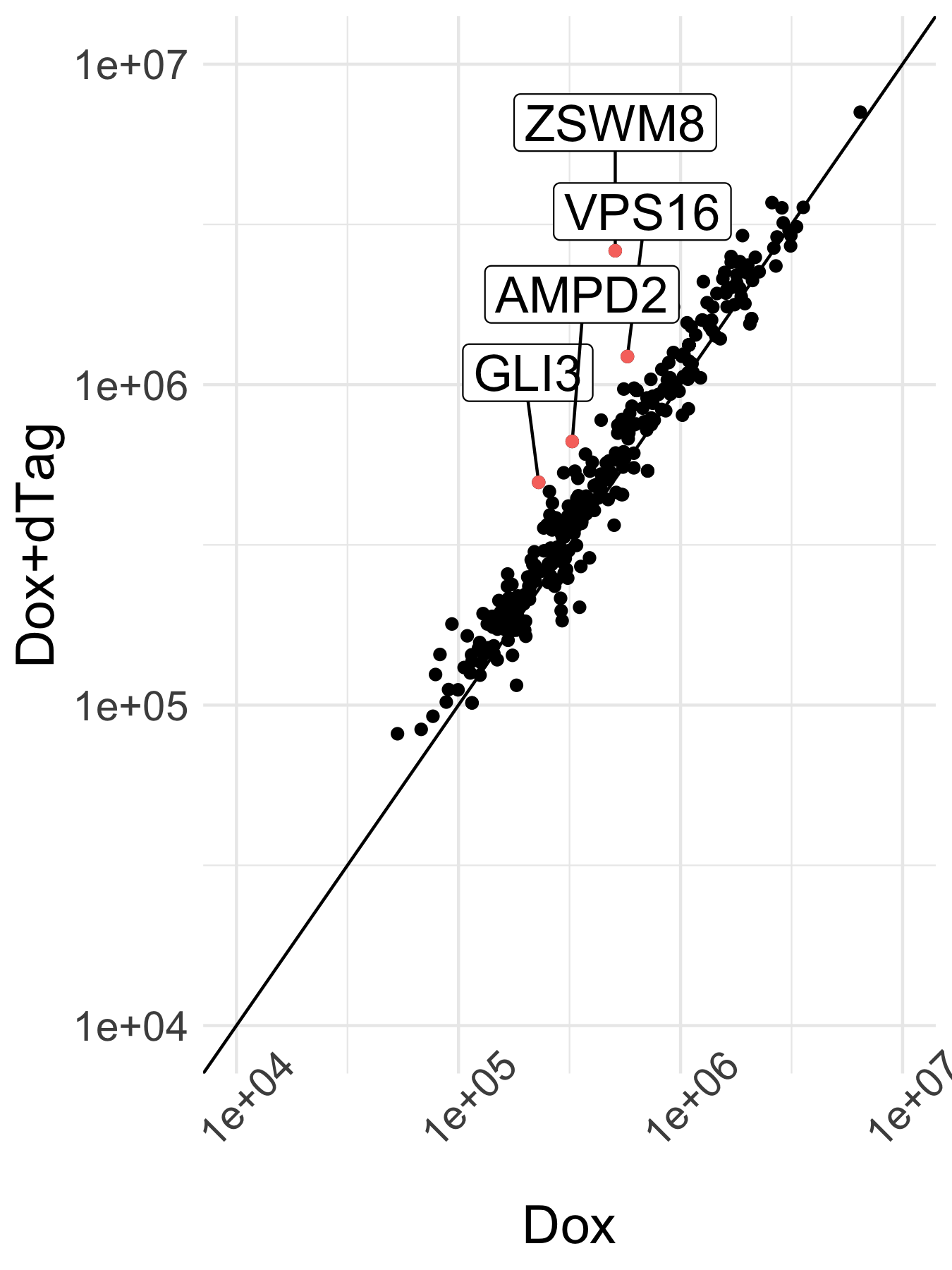
