## Supplementary material for "Sensitive and quantitative detection of MHC-I displayed neoepitopes using a semi-automated workflow and TOMAHAQ mass spectrometry": Peptide_traces_42454_Darwish.pdf

Data File : S:/MSD-07/2020/02-FEB/040220/1427/42454DAR001_FR3->

Injection Date : 04-Feb-20, 14:51:24
Sample Name : 42454DAR001_FR3
Remarks : Baseline corrected
Acq. Method : D:/METHODS/5-95-6M.M
column used : Je_559
Analysis Method : C:/CHEM32/1/METHODS/JPT-LCMS.M

Sample Info : 42454_001-Dr. Martine Darwish H-EVLAFAIL-OH

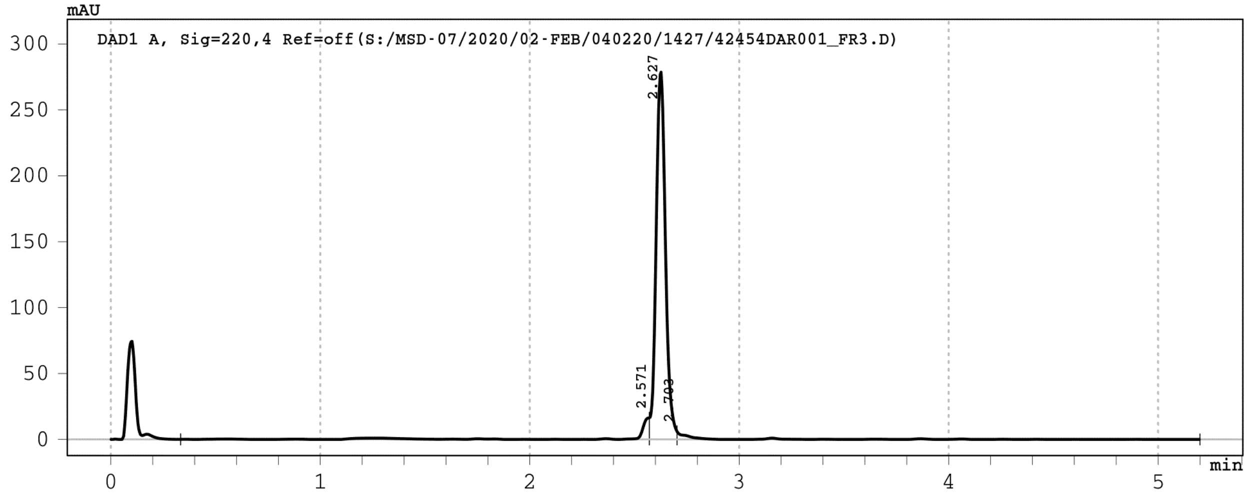


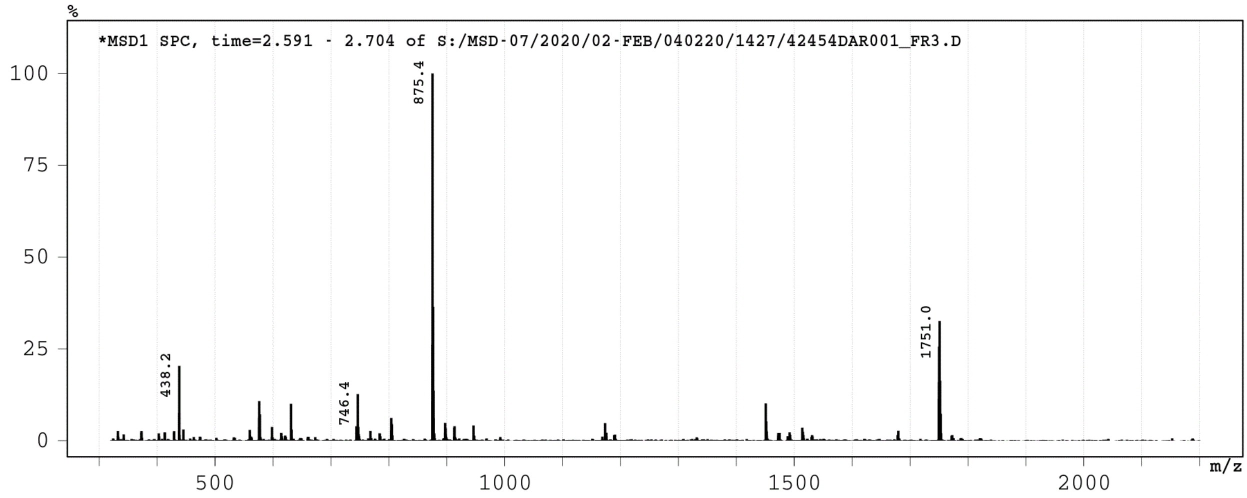


Signal 1: DAD1 A, Sig=220,4 Ref=off

| Peak # | RT [min] | Width [min] | Area | Area % |
|---|---|---|---|---|
| 1 | 2.571 | 2.24 | 60.2 | 6.3 |
| 2 | 2.627 | 0.13 | 866.1 | 90.8 |
| 3 | 2.703 | 2.50 | 27.6 | 2.9 |

Data File : S:/MSD-07/2020/02-FEB/040220/1427/42454DAR002_FR1->

Injection Date : 04-Feb-20, 14:59:48
Sample Name : 42454DAR002_FR11-12
Remarks : Baseline corrected
Acq. Method : D:/METHODS/5-95-6M.M
column used : Je_559
Analysis Method : C:/CHEM32/1/METHODS/JPT-LCMS.M

Sample Info : 42454_002-Dr. Martine Darwish H-ASMTNMELM-OH

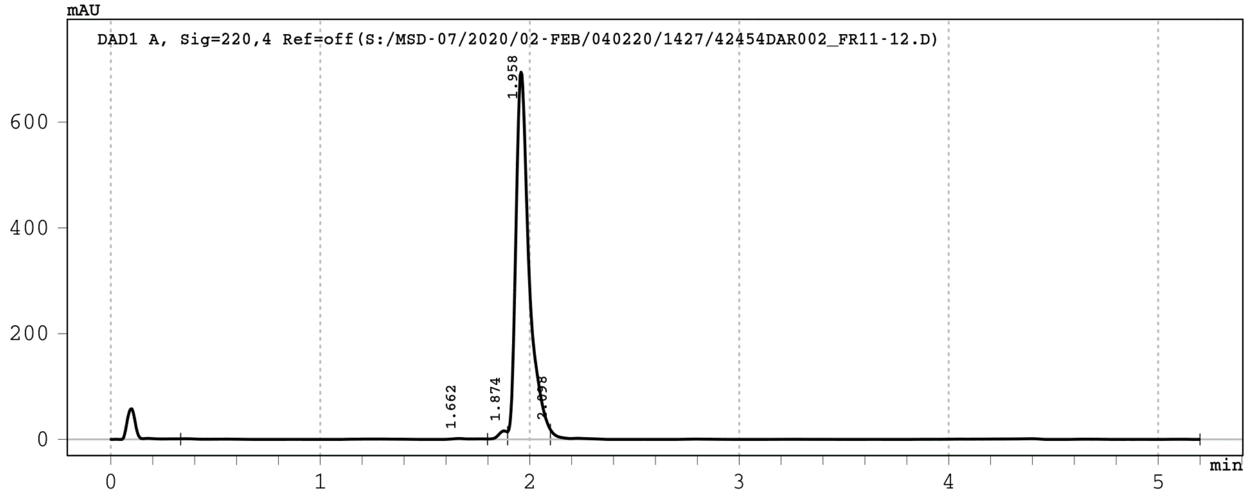


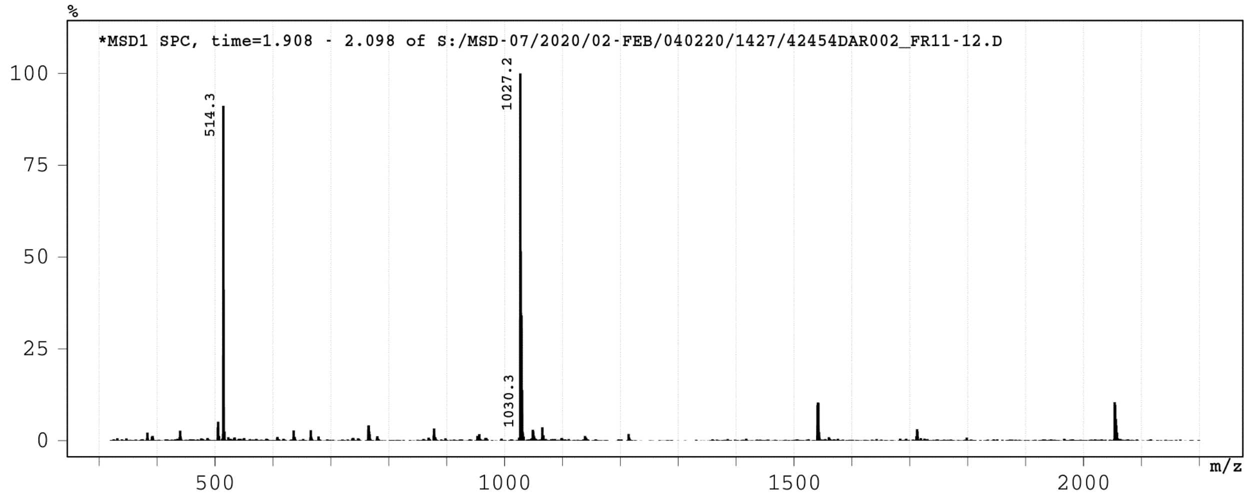


Signal 1: DAD1 A, Sig=220,4 Ref=off

| Peak # | RT [min] | Width [min] | Area | Area % |
|---|---|---|---|---|
| 1 | 1.662 | 1.46 | 33.5 | 1.0 |
| 2 | 1.874 | 0.10 | 45.2 | 1.4 |
| 3 | 1.958 | 0.20 | 3067.8 | 95.0 |
| 4 | 2.098 | 3.10 | 82.9 | 2.6 |

Data File : S:/MSD-07/2020/02-FEB/040220/1427/42454DAR003_FR1->

Injection Date : 04-Feb-20, 15:08:12
Sample Name : 42454DAR003_FR19-21
Remarks : Baseline corrected
Acq. Method : D:/METHODS/5-95-6M.M
column used : Je_559
Analysis Method : C:/CHEM32/1/METHODS/JPT-LCMS.M

Sample Info : 42454_003-Dr. Martine Darwish H-ANFQLCPL-OH

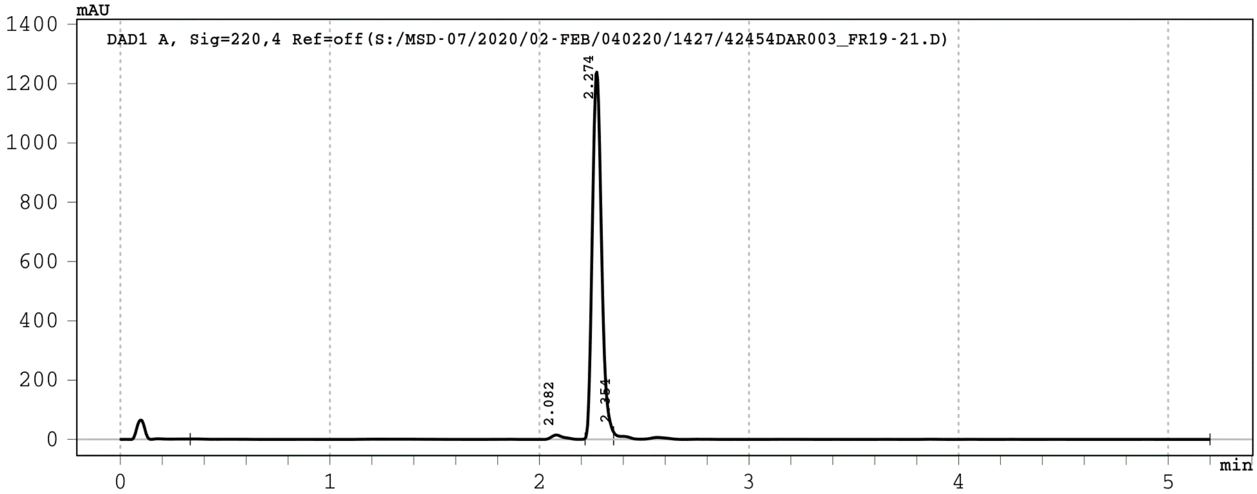


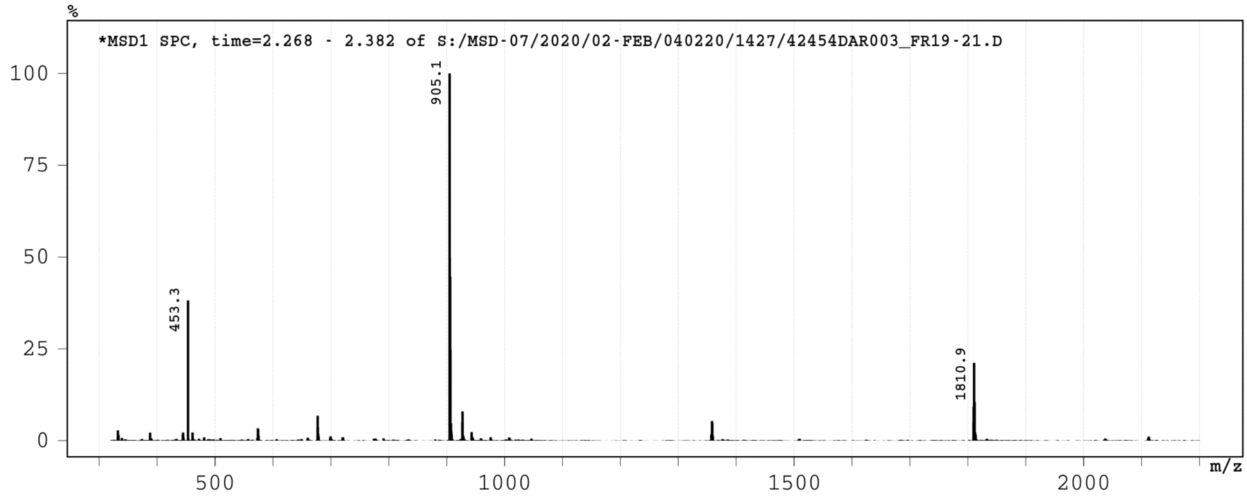


Signal 1: DAD1 A, Sig=220,4 Ref=off

| Peak # | RT [min] | Width [min] | Area | Area % |
|---|---|---|---|---|
| 1 | 2.082 | 1.88 | 89.2 | 2.2 |
| 2 | 2.274 | 0.14 | 3803.6 | 95.1 |
| 3 | 2.354 | 2.85 | 108.0 | 2.7 |

Data File : S:/MSD-07/2020/02-FEB/040220/1427/42454DAR004_FR2->

Injection Date : 04-Feb-20, 15:16:36
Sample Name : 42454DAR004_FR28
Remarks : Baseline corrected
Acq. Method : D:/METHODS/5-95-6M.M
column used : Je_559
Analysis Method : C:/CHEM32/1/METHODS/JPT-LCMS.M

Sample Info : 42454_004-Dr. Martine Darwish H-FSFFNPKCLL-OH

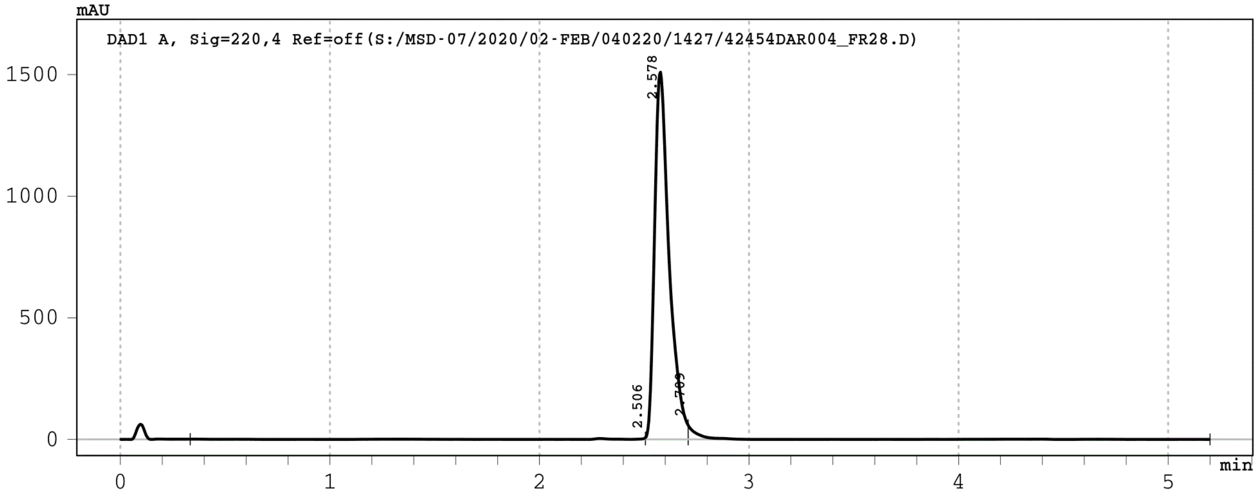


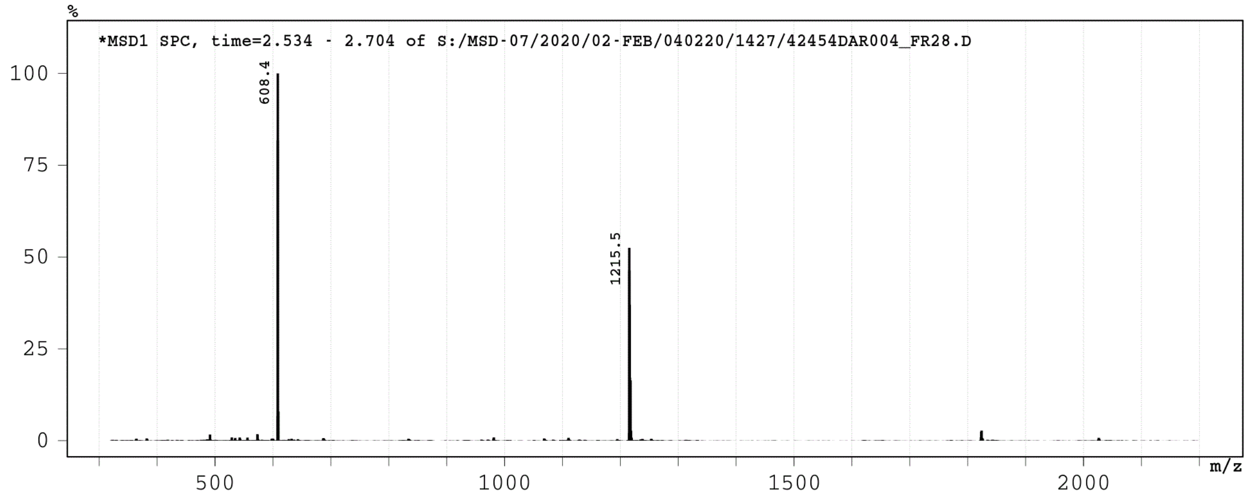


Signal 1: DAD1 A, Sig=220,4 Ref=off

| Peak # | RT [min] | Width [min] | Area | Area % |
|---|---|---|---|---|
| 1 | 2.506 | 2.17 | 56.8 | 0.7 |
| 2 | 2.578 | 0.20 | 7321.3 | 96.6 |
| 3 | 2.709 | 2.49 | 201.4 | 2.7 |

Data File : S:/MSD-07/2020/02-FEB/040220/1427/42454DAR005_FR4->

Injection Date : 04-Feb-20, 15:25:00
Sample Name : 42454DAR005_FR42-43
Remarks : Baseline corrected
Acq. Method : D:/METHODS/5-95-6M.M
column used : Je_559
Analysis Method : C:/CHEM32/1/METHODS/JPT-LCMS.M

Sample Info : 42454_005-Dr. Martine Darwish H-STYVIPRL-OH

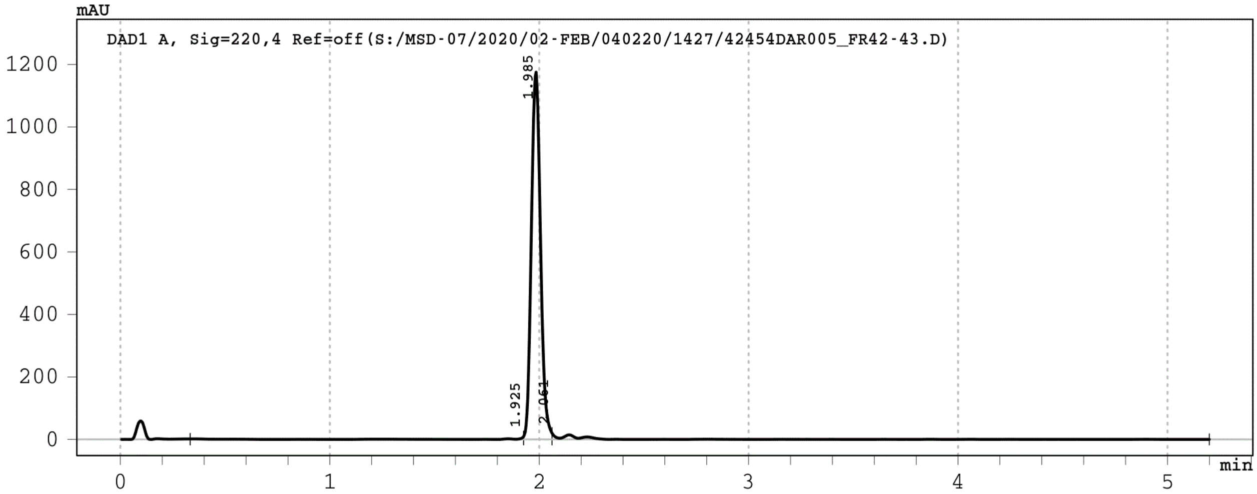


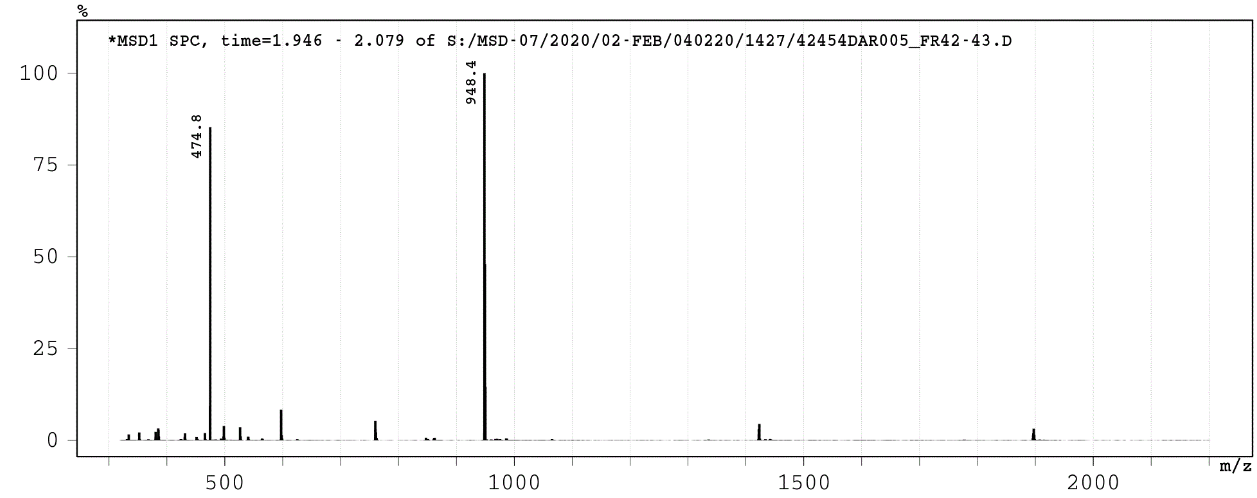


Signal 1: DAD1 A, Sig=220,4 Ref=off

| Peak # | RT [min] | Width [min] | Area | Area % |
|---|---|---|---|---|
| 1 | 1.925 | 1.59 | 40.8 | 1.1 |
| 2 | 1.985 | 0.14 | 3582.7 | 95.8 |
| 3 | 2.061 | 3.14 | 115.1 | 3.1 |

Data File : S:/MSD-07/2020/02-FEB/040220/1427/42454DAR006_FR5->

Injection Date : 04-Feb-20, 15:33:22
Sample Name : 42454DAR006_FR56
Remarks : Baseline corrected
Acq. Method : D:/METHODS/5-95-6M.M
column used : Je_559
Analysis Method : C:/CHEM32/1/METHODS/JPT-LCMS.M

Sample Info : 42454_006-Dr. Martine Darwish H-YTICNPCVPV-OH

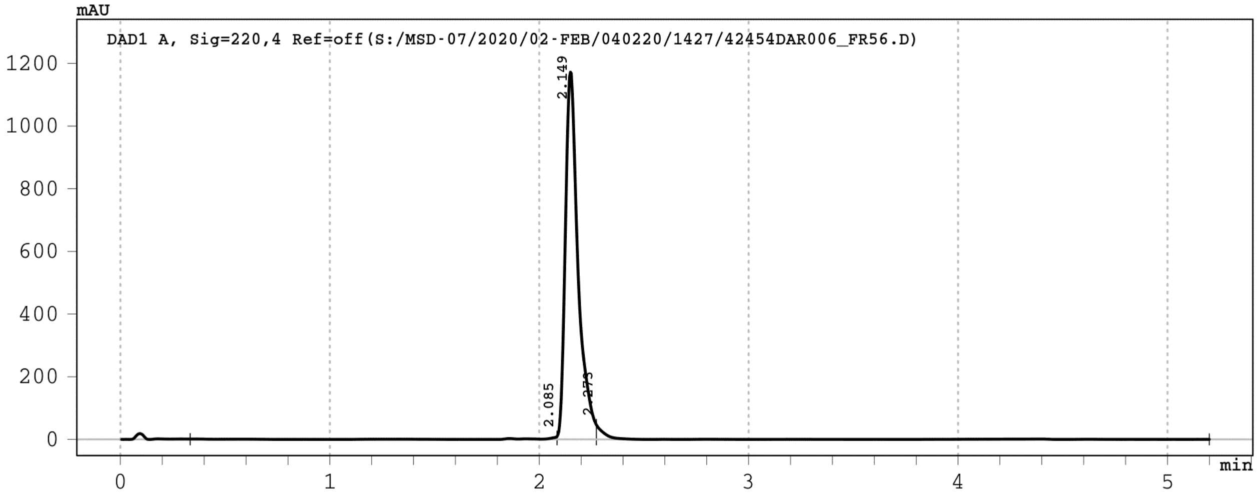


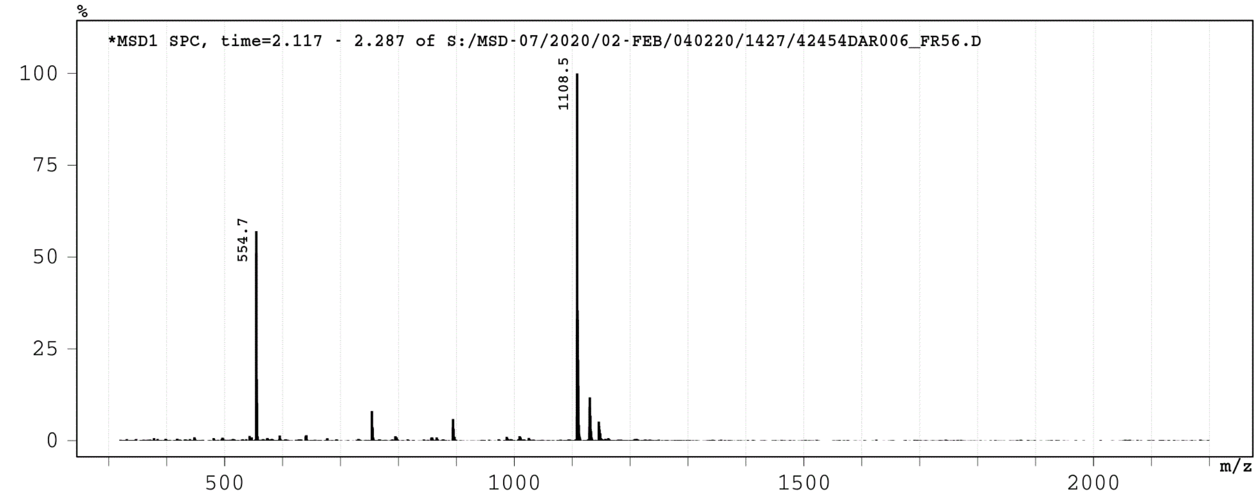


Signal 1: DAD1 A, Sig=220,4 Ref=off

| Peak # | RT [min] | Width [min] | Area | Area % |
|---|---|---|---|---|
| 1 | 2.085 | 1.75 | 62.6 | 1.2 |
| 2 | 2.149 | 0.19 | 4963.1 | 95.9 |
| 3 | 2.273 | 2.93 | 148.1 | 2.9 |

Data File : S:/MSD-07/2020/02-FEB/040220/1427/42454DAR007_FR6->

Injection Date : 04-Feb-20, 15:41:47
Sample Name : 42454DAR007_FR68-70
Remarks : Baseline corrected
Acq. Method : D:/METHODS/5-95-6M.M
column used : Je_559
Analysis Method : C:/CHEM32/1/METHODS/JPT-LCMS.M

Sample Info : 42454_007-Dr. Martine Darwish H-ASYNGFLPV-OH

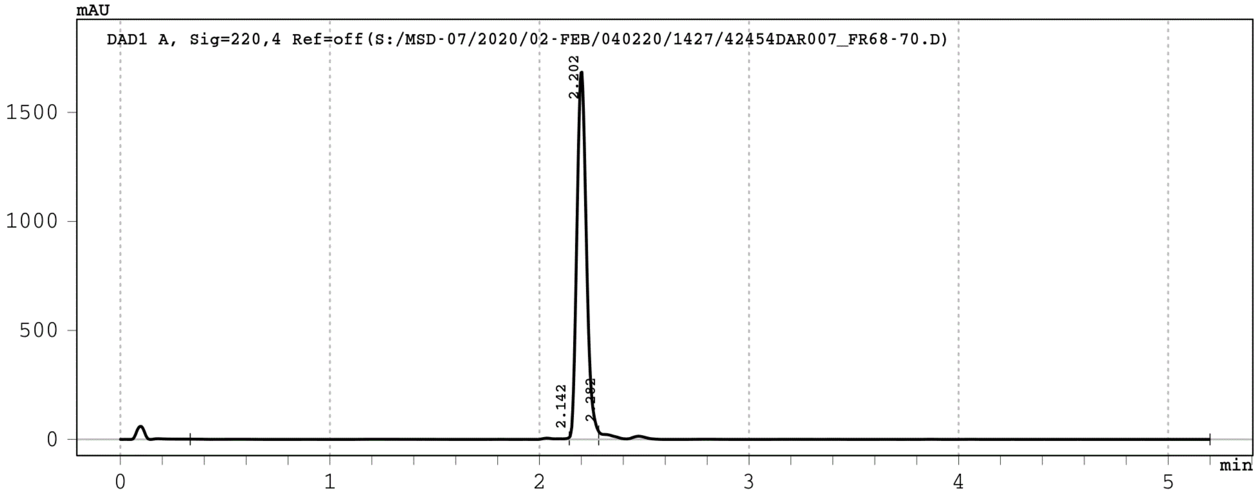


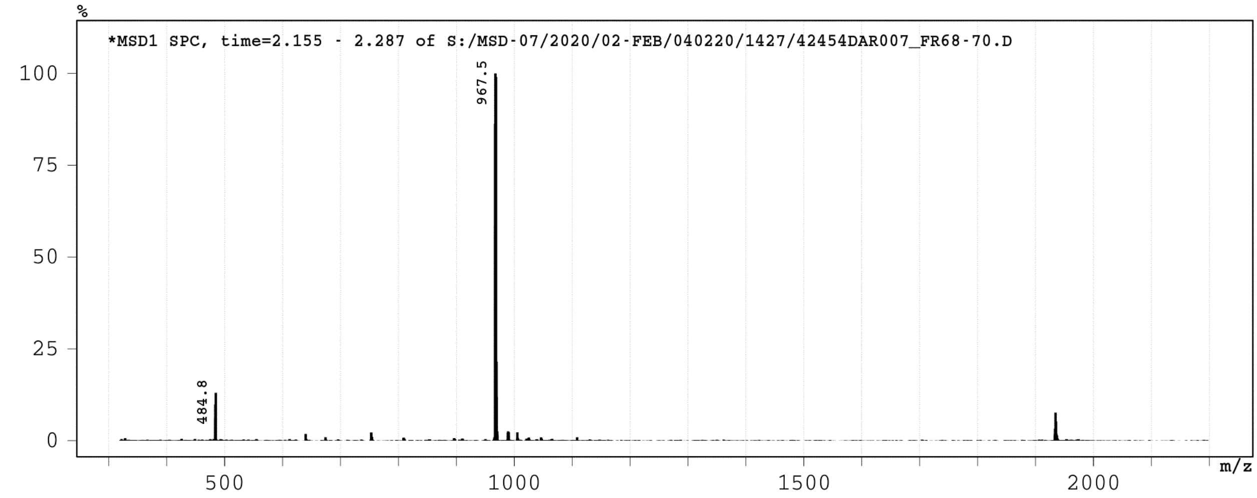


Signal 1: DAD1 A, Sig=220,4 Ref=off

| Peak # | RT [min] | Width [min] | Area | Area % |
|---|---|---|---|---|
| 1 | 2.142 | 1.81 | 51.9 | 0.9 |
| 2 | 2.202 | 0.14 | 5359.4 | 95.5 |
| 3 | 2.282 | 2.92 | 200.0 | 3.6 |

Data File : S:/MSD-07/2020/02-FEB/140220/1435/42454DAR008_FR7->

Injection Date : 14-Feb-20, 23:28:03
Sample Name : 42454DAR008_FR7-9
Remarks : Baseline corrected
Acq. Method : D:/METHODS/5-95-6M.M
column used : Je_559
Analysis Method : C:/CHEM32/1/METHODS/JPT-LCMS.M

Sample Info : 42454_008-Dr. Martine Darwish H-FVMDYIPV-OH

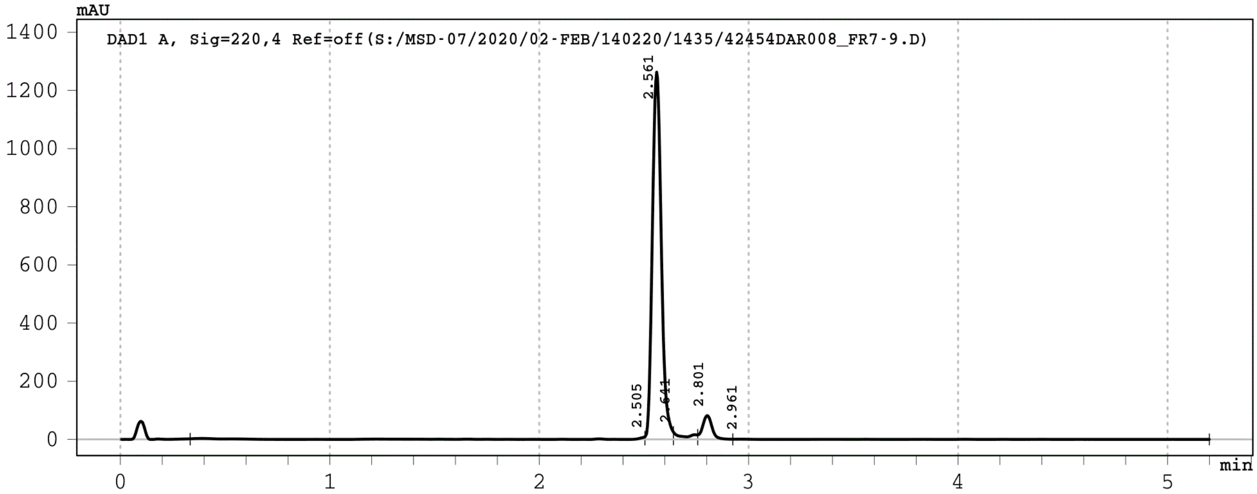


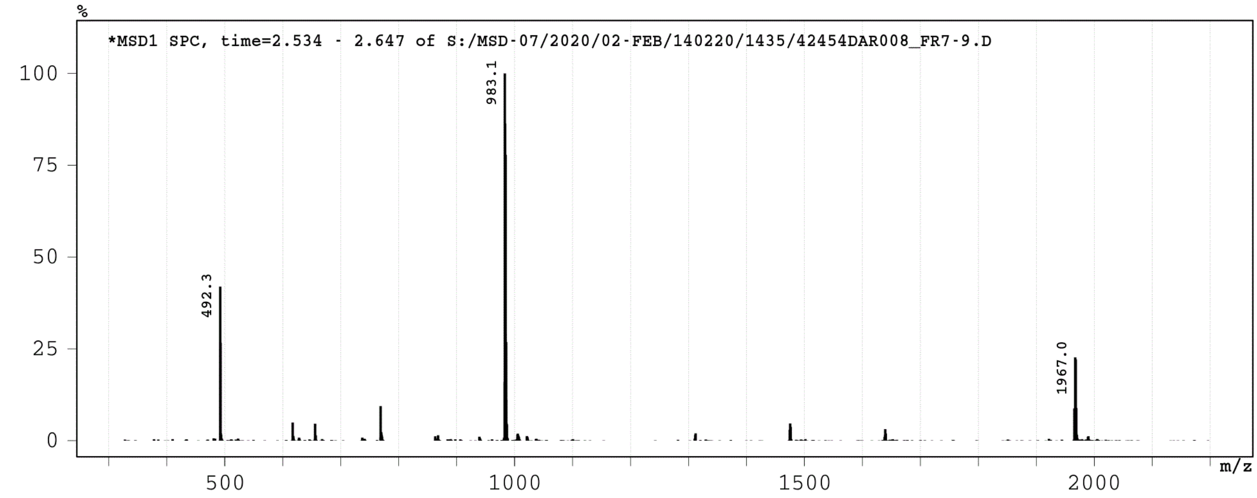


Signal 1: DAD1 A, Sig=220,4 Ref=off

| Peak # | RT [min] | Width [min] | Area | Area % |
|---|---|---|---|---|
| 1 | 2.505 | 2.17 | 84.8 | 2.0 |
| 2 | 2.561 | 0.14 | 3760.5 | 89.4 |
| 3 | 2.641 | 0.12 | 90.3 | 2.1 |
| 4 | 2.801 | 0.17 | 258.9 | 6.2 |
| 5 | 2.961 | 2.28 | 12.6 | 0.3 |

Data File : S:/MSD-07/2020/02-FEB/070220/1430/42454DAR009_FR2->

Injection Date : 07-Feb-20, 13:12:50
Sample Name : 42454DAR009_FR2-4
Remarks : Baseline corrected
Acq. Method : D:/METHODS/5-95-6M.M
column used : Je_559
Analysis Method : C:/CHEM32/1/METHODS/JPT-LCMS.M

Sample Info : 42454_009-Dr. Martine Darwish H-QSYVPAPL-OH

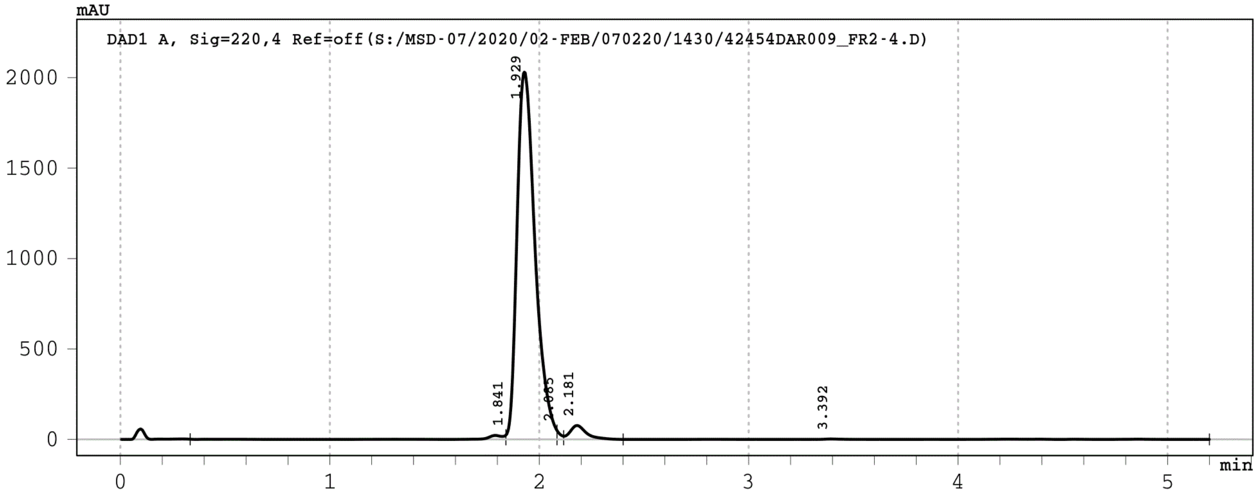


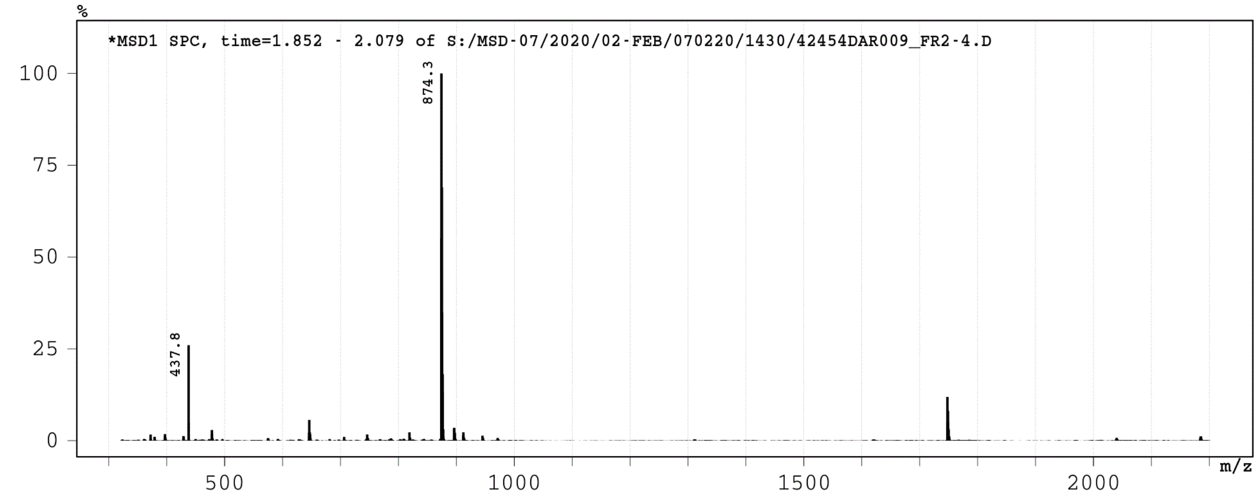


Signal 1: DAD1 A, Sig=220,4 Ref=off

| Peak # | RT [min] | Width [min] | Area | Area % |
|---|---|---|---|---|
| 1 | 1.841 | 1.51 | 131.5 | 1.0 |
| 2 | 1.929 | 0.24 | 12220.0 | 94.9 |
| 3 | 2.085 | 0.03 | 56.0 | 0.4 |
| 4 | 2.181 | 0.28 | 422.9 | 3.3 |
| 5 | 3.392 | 2.80 | 46.2 | 0.4 |

Data File : S:/MSD-07/2020/02-FEB/040220/1427/42454DAR010_FR8->

Injection Date : 04-Feb-20, 15:50:10
Sample Name : 42454DAR010_FR81-82
Remarks : Baseline corrected
Acq. Method : D:/METHODS/5-95-6M.M
column used : Je_559
Analysis Method : C:/CHEM32/1/METHODS/JPT-LCMS.M

Sample Info : 42454_010-Dr. Martine Darwish H-KNHWNSTML-OH

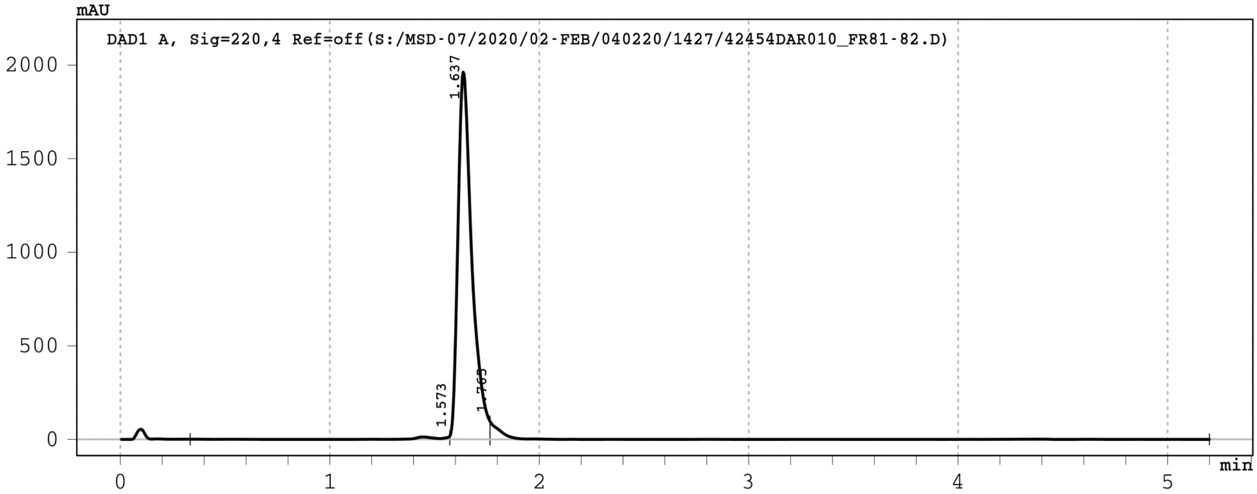


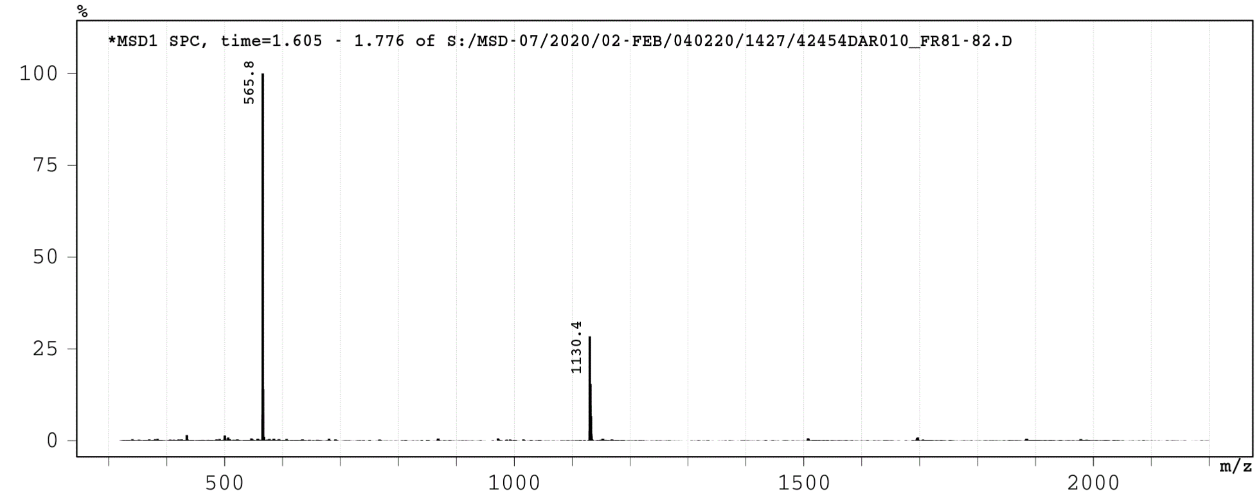


Signal 1: DAD1 A, Sig=220,4 Ref=off

| Peak # | RT [min] | Width [min] | Area | Area % |
|---|---|---|---|---|
| 1 | 1.573 | 1.24 | 115.7 | 1.2 |
| 2 | 1.637 | 0.19 | 9130.6 | 95.2 |
| 3 | 1.765 | 3.44 | 348.1 | 3.6 |

Data File : S:/MSD-07/2020/02-FEB/070220/1430/42454DAR011_FR1->

Injection Date : 07-Feb-20, 13:21:13
Sample Name : 42454DAR011_FR18-21
Remarks : Baseline corrected
Acq. Method : D:/METHODS/5-95-6M.M
column used : Je_559
Analysis Method : C:/CHEM32/1/METHODS/JPT-LCMS.M

Sample Info : 42454_011-Dr. Martine Darwish H-FMLPFFMAF-OH

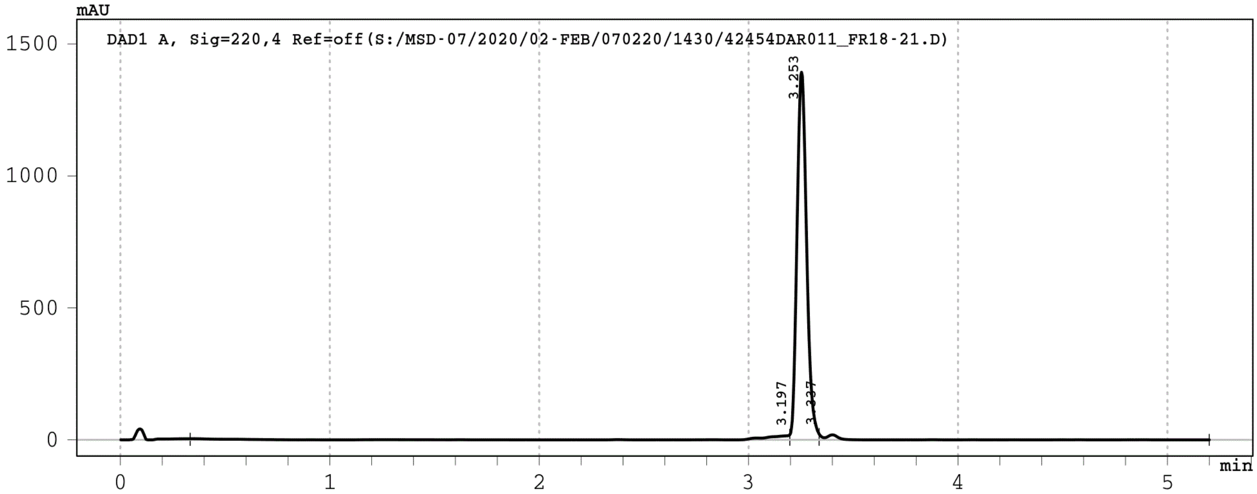


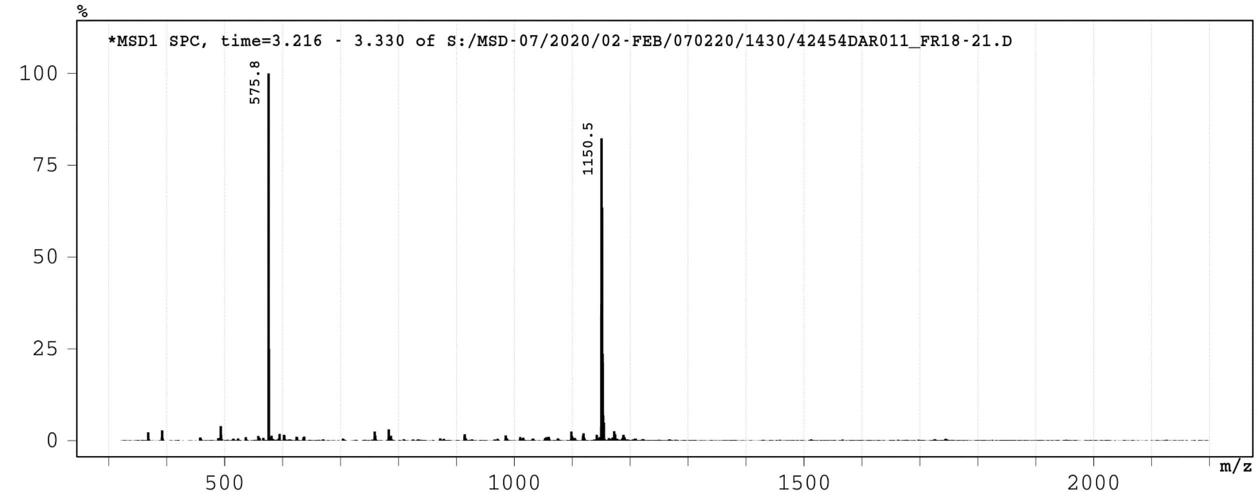


Signal 1: DAD1 A, Sig=220,4 Ref=off

| Peak # | RT [min] | Width [min] | Area | Area % |
|---|---|---|---|---|
| 1 | 3.197 | 2.86 | 202.8 | 4.3 |
| 2 | 3.253 | 0.14 | 4463.7 | 93.8 |
| 3 | 3.337 | 1.86 | 93.4 | 2.0 |

Data File : S:/MSD-07/2020/02-FEB/040220/1427/42454DAR012_FR1->

Injection Date : 04-Feb-20, 15:58:34
Sample Name : 42454DAR012_FR17-18
Remarks : Baseline corrected
Acq. Method : D:/METHODS/5-95-6M.M
column used : Je_559
Analysis Method : C:/CHEM32/1/METHODS/JPT-LCMS.M

Sample Info : 42454_012-Dr. Martine Darwish H-FVIDFKPL-OH

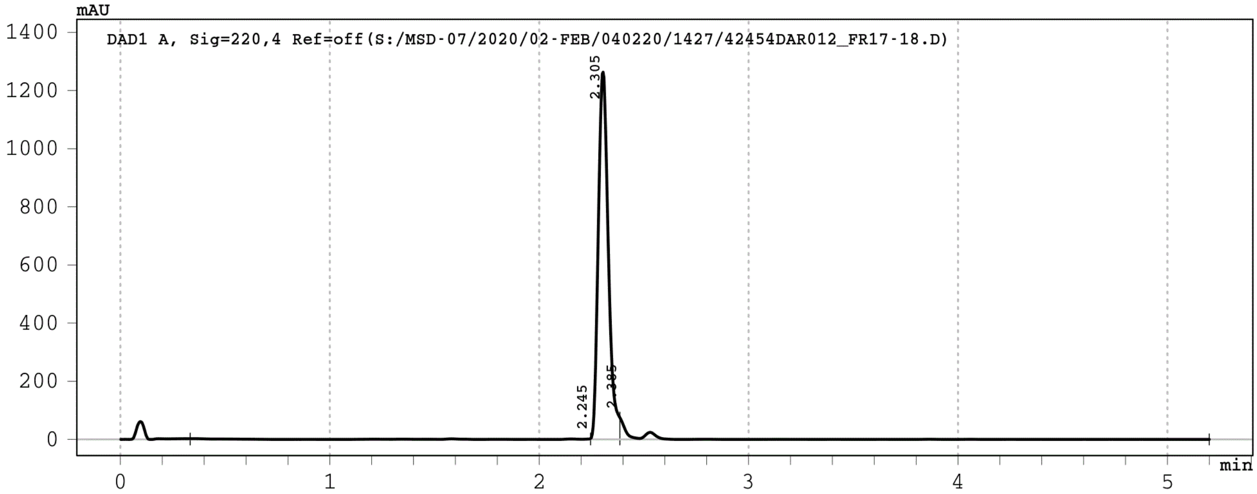


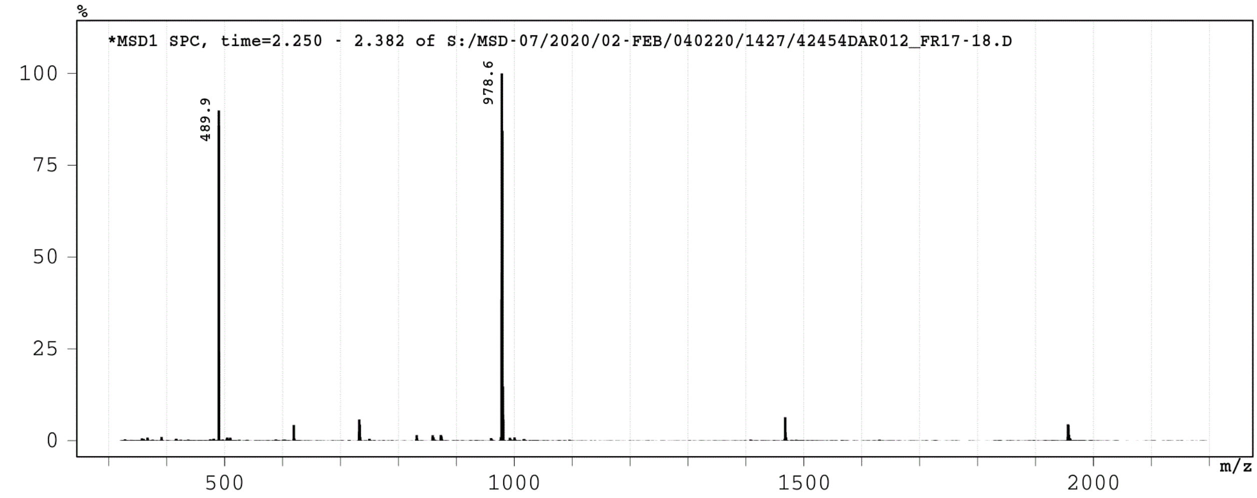


Signal 1: DAD1 A, Sig=220,4 Ref=off

| Peak # | RT [min] | Width [min] | Area | Area % |
|---|---|---|---|---|
| 1 | 2.245 | 1.91 | 60.4 | 1.3 |
| 2 | 2.305 | 0.14 | 4217.8 | 93.7 |
| 3 | 2.385 | 2.81 | 224.1 | 5.0 |

Data File : S:/MSD-07/2020/02-FEB/040220/1427/42454DAR013_FR3->

Injection Date : 04-Feb-20, 16:06:56
Sample Name : 42454DAR013_FR36-37
Remarks : Baseline corrected
Acq. Method : D:/METHODS/5-95-6M.M
column used : Je_559
Analysis Method : C:/CHEM32/1/METHODS/JPT-LCMS.M

Sample Info : 42454_013-Dr. Martine Darwish H-CMAFWIPFI-OH

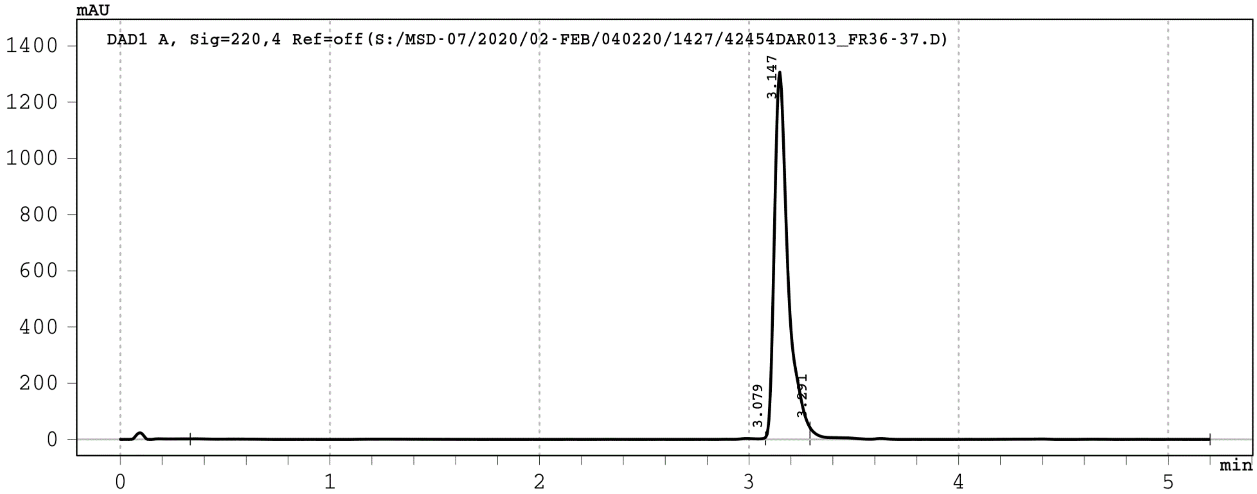


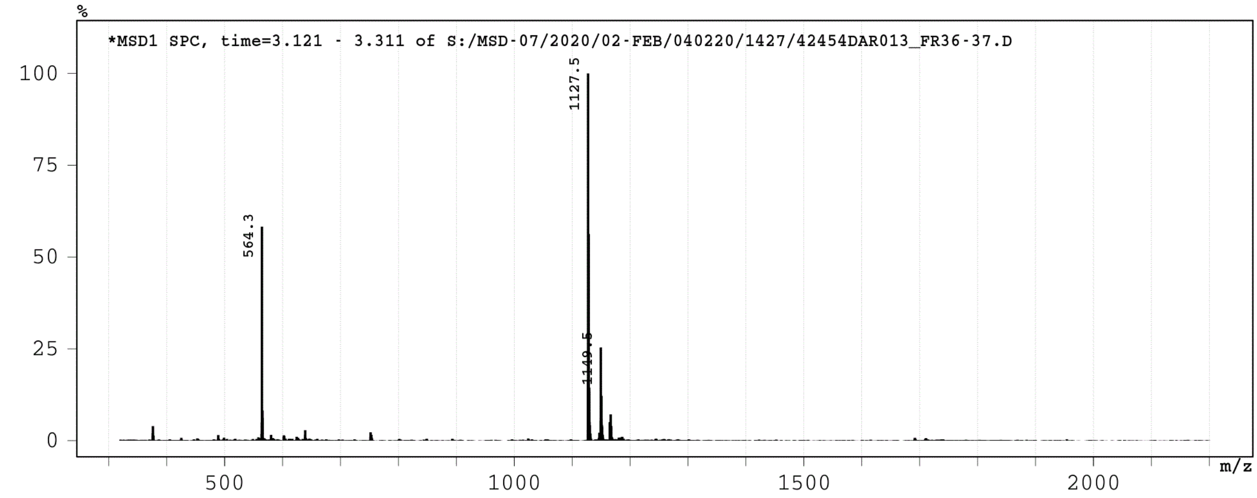


Signal 1: DAD1 A, Sig=220,4 Ref=off

| Peak # | RT [min] | Width [min] | Area | Area % |
|---|---|---|---|---|
| 1 | 3.079 | 2.75 | 67.2 | 1.1 |
| 2 | 3.147 | 0.21 | 5937.5 | 96.2 |
| 3 | 3.291 | 1.91 | 170.4 | 2.8 |

| | |
|---|---|
| Data File | : S:/MSD-07/2020/02-FEB/040220/1427/42454DAR014_FR5-> |
| Injection Date | : 04-Feb-20, 16:15:19 |
| Sample Name | : 42454DAR014_FR55-56 |
| Remarks | : Baseline corrected |
| Acq. Method | : D:/METHODS/5-95-6M.M |
| column used | : Je_559 |
| Analysis Method | : C:/CHEM32/1/METHODS/JPT-LCMS.M |
| Sample Info | : 42454_014-Dr. Martine Darwish H-VNSIHALL-OH |

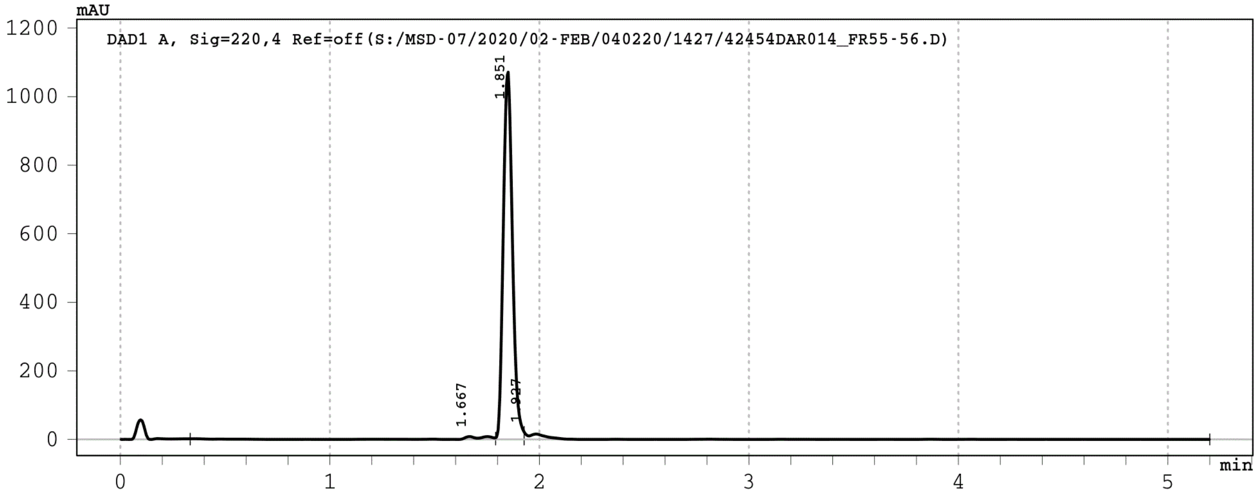


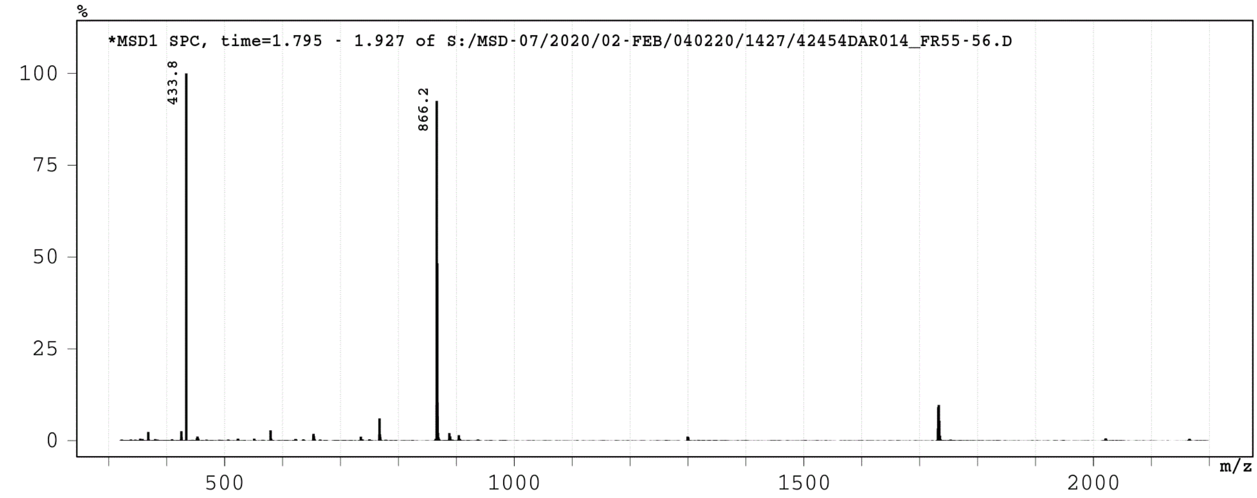


Signal 1: DAD1 A, Sig=220,4 Ref=off

| Peak # | RT [min] | Width [min] | Area | Area % |
|---|---|---|---|---|
| 1 | 1.667 | 1.46 | 83.7 | 2.4 |
| 2 | 1.851 | 0.14 | 3274.4 | 94.2 |
| 3 | 1.927 | 3.27 | 117.2 | 3.4 |

Data File : S:/MSD-07/2020/02-FEB/040220/1427/42454DAR015_FR6->

Injection Date : 04-Feb-20, 16:23:43
Sample Name : 42454DAR015_FR60
Remarks : Baseline corrected
Acq. Method : D:/METHODS/5-95-6M.M
column used : Je_559
Analysis Method : C:/CHEM32/1/METHODS/JPT-LCMS.M

Sample Info : 42454_015-Dr. Martine Darwish H-AQLANDVVL-OH

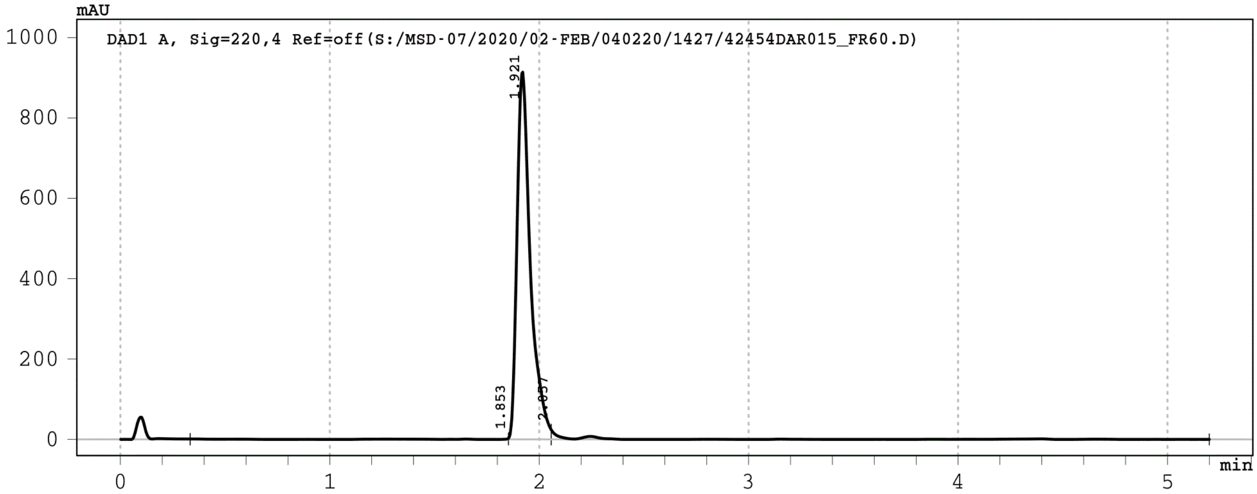


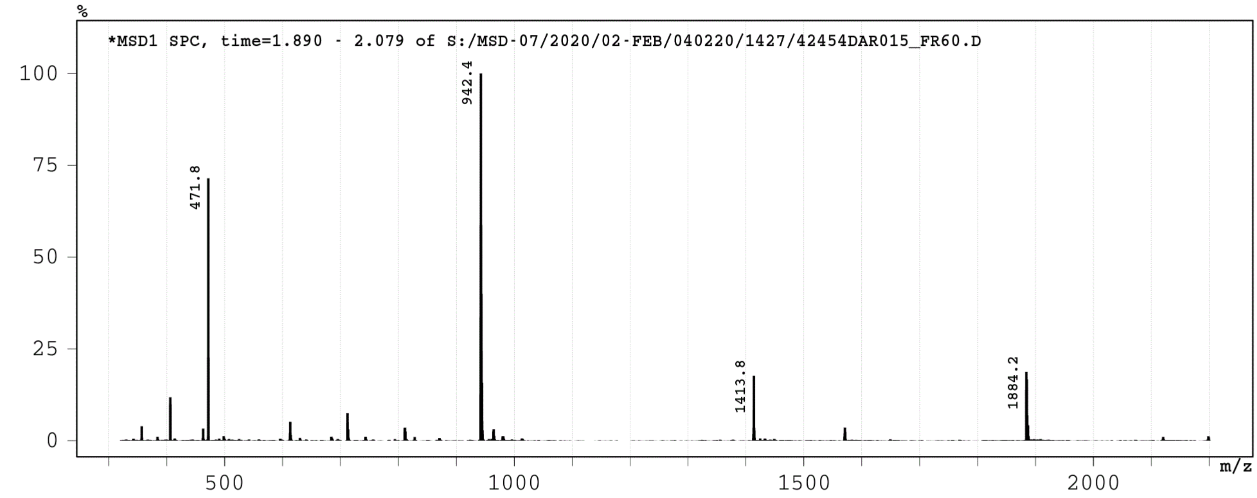


Signal 1: DAD1 A, Sig=220,4 Ref=off

| Peak # | RT [min] | Width [min] | Area | Area % |
|---|---|---|---|---|
| 1 | 1.853 | 1.52 | 30.3 | 0.7 |
| 2 | 1.921 | 0.20 | 4085.3 | 96.4 |
| 3 | 2.057 | 3.14 | 121.8 | 2.9 |

Data File : S:/MSD-07/2020/02-FEB/040220/1427/42454DAR016_FR7->

Injection Date : 04-Feb-20, 16:32:07
Sample Name : 42454DAR016_FR71-72
Remarks : Baseline corrected
Acq. Method : D:/METHODS/5-95-6M.M
column used : Je_559
Analysis Method : C:/CHEM32/1/METHODS/JPT-LCMS.M

Sample Info : 42454_016-Dr. Martine Darwish H-KILTFDRL-OH

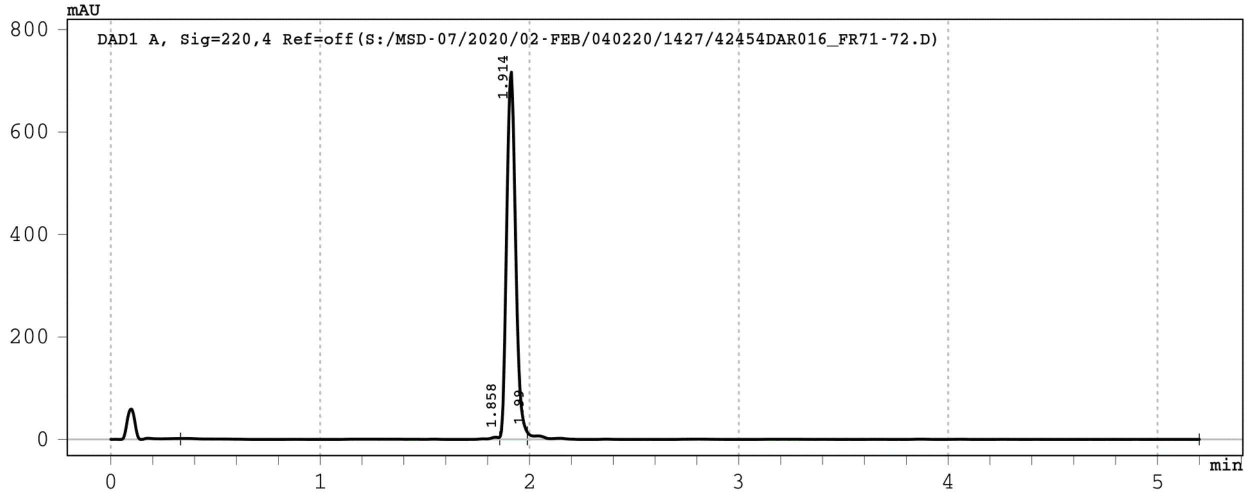


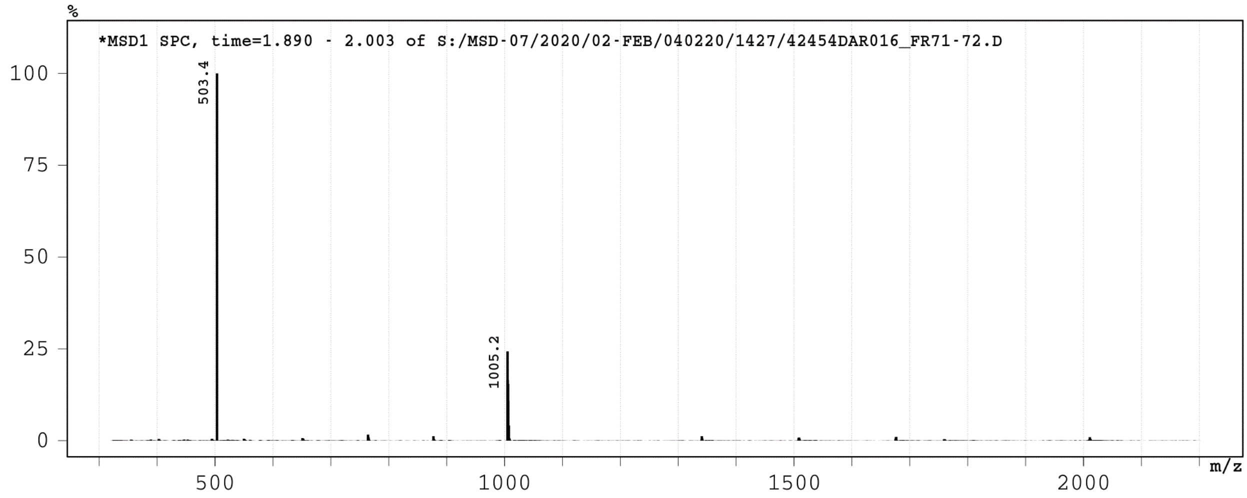


Signal 1: DAD1 A, Sig=220,4 Ref=off

| Peak # | RT [min] | Width [min] | Area | Area % |
|---|---|---|---|---|
| 1 | 1.858 | 1.52 | 40.9 | 1.8 |
| 2 | 1.914 | 0.13 | 2143.4 | 95.4 |
| 3 | 1.990 | 3.21 | 61.7 | 2.7 |

Data File : S:/MSD-07/2020/02-FEB/040220/1427/42454DAR017_FR7->

Injection Date : 04-Feb-20, 16:40:33
Sample Name : 42454DAR017_FR79
Remarks : Baseline corrected
Acq. Method : D:/METHODS/5-95-6M.M
column used : Je_559
Analysis Method : C:/CHEM32/1/METHODS/JPT-LCMS.M

Sample Info : 42454_017-Dr. Martine Darwish H-VMMTSAEAV-OH

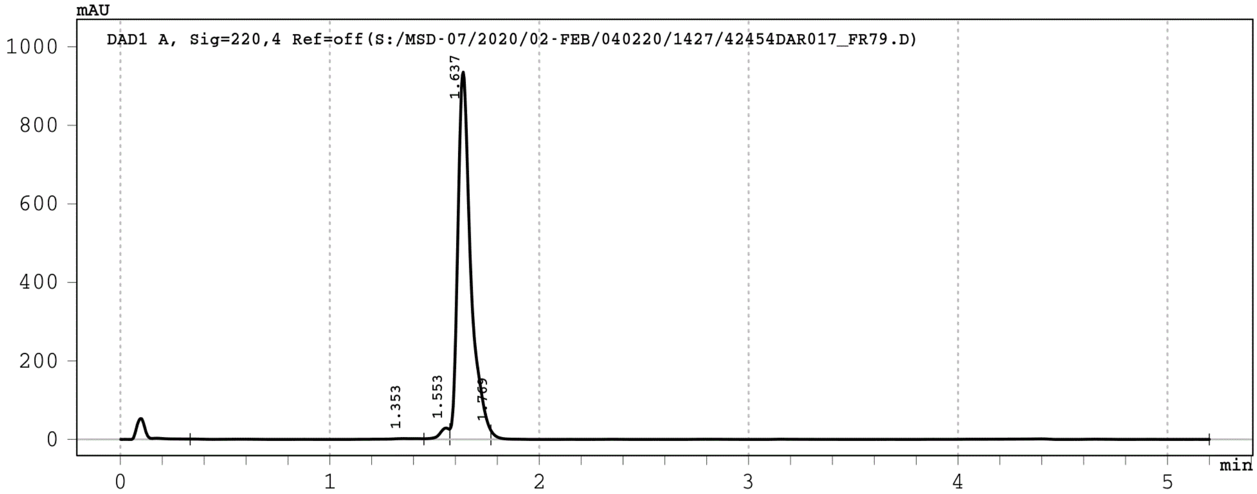


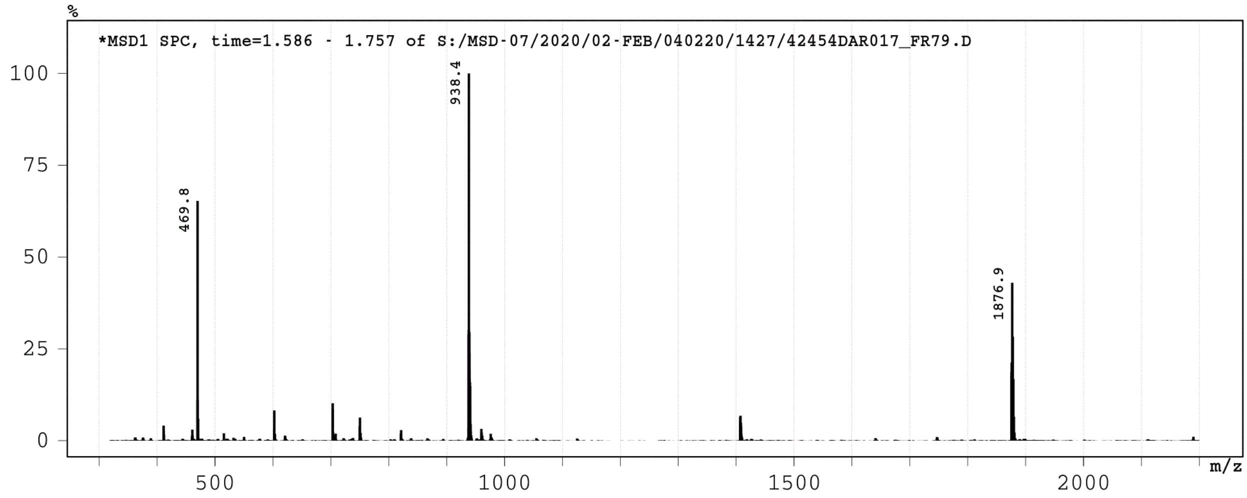


Signal 1: DAD1 A, Sig=220,4 Ref=off

| Peak # | RT [min] | Width [min] | Area | Area % |
|---|---|---|---|---|
| 1 | 1.353 | 1.12 | 32.2 | 0.8 |
| 2 | 1.553 | 0.12 | 93.1 | 2.2 |
| 3 | 1.637 | 0.20 | 4035.9 | 95.4 |
| 4 | 1.769 | 3.43 | 68.8 | 1.6 |

Data File : S:/MSD-07/2020/02-FEB/040220/1427/42454DAR018_FR8->

Injection Date : 04-Feb-20, 16:48:57
Sample Name : 42454DAR018_FR8-9
Remarks : Baseline corrected
Acq. Method : D:/METHODS/5-95-6M.M
column used : Je_559
Analysis Method : C:/CHEM32/1/METHODS/JPT-LCMS.M

Sample Info : 42454_018-Dr. Martine Darwish H-SAIRSYQYV-OH

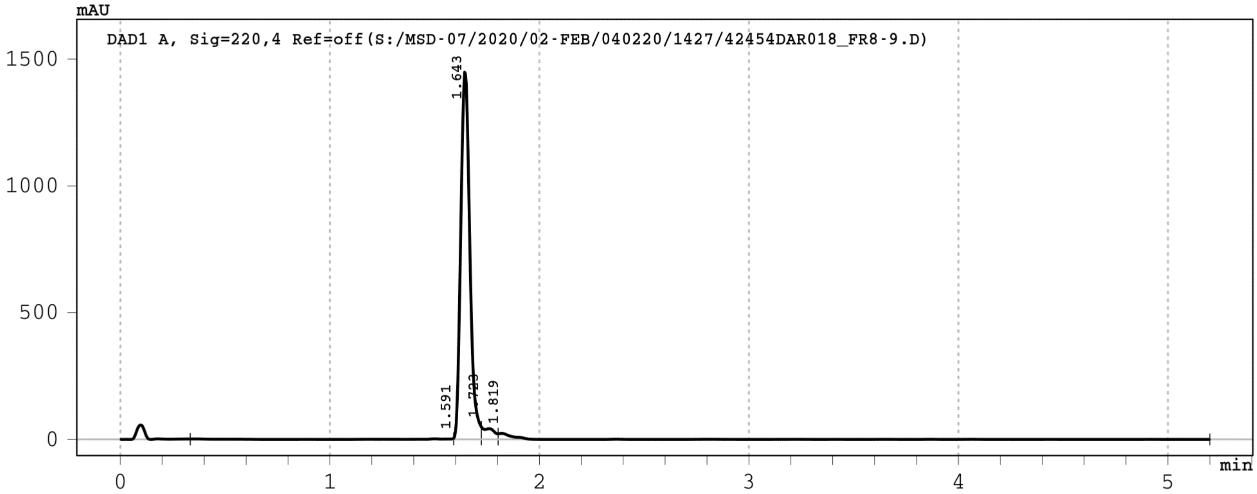


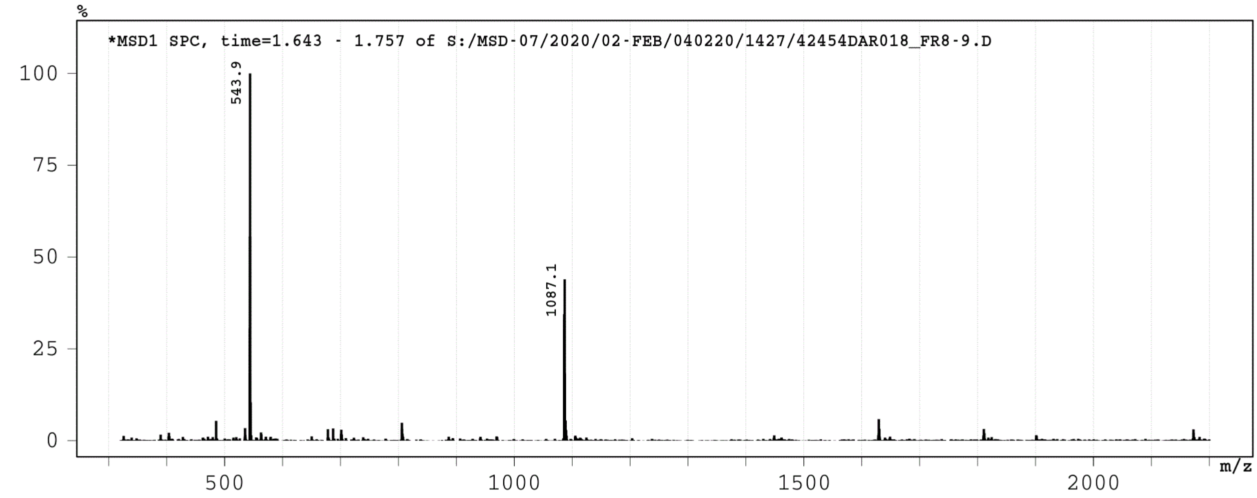


Signal 1: DAD1 A, Sig=220,4 Ref=off

| Peak # | RT [min] | Width [min] | Area | Area % |
|---|---|---|---|---|
| 1 | 1.591 | 1.26 | 37.6 | 0.8 |
| 2 | 1.643 | 0.13 | 4371.4 | 92.8 |
| 3 | 1.723 | 0.08 | 176.1 | 3.7 |
| 4 | 1.819 | 3.40 | 125.1 | 2.7 |

Data File : S:/MSD-07/2020/02-FEB/040220/1427/42454DAR019_FR1->

Injection Date : 04-Feb-20, 16:57:23
Sample Name : 42454DAR019_FR16-17
Remarks : Baseline corrected
Acq. Method : D:/METHODS/5-95-6M.M
column used : Je_559
Analysis Method : C:/CHEM32/1/METHODS/JPT-LCMS.M

Sample Info : 42454_019-Dr. Martine Darwish H-FALMNLKAL-OH

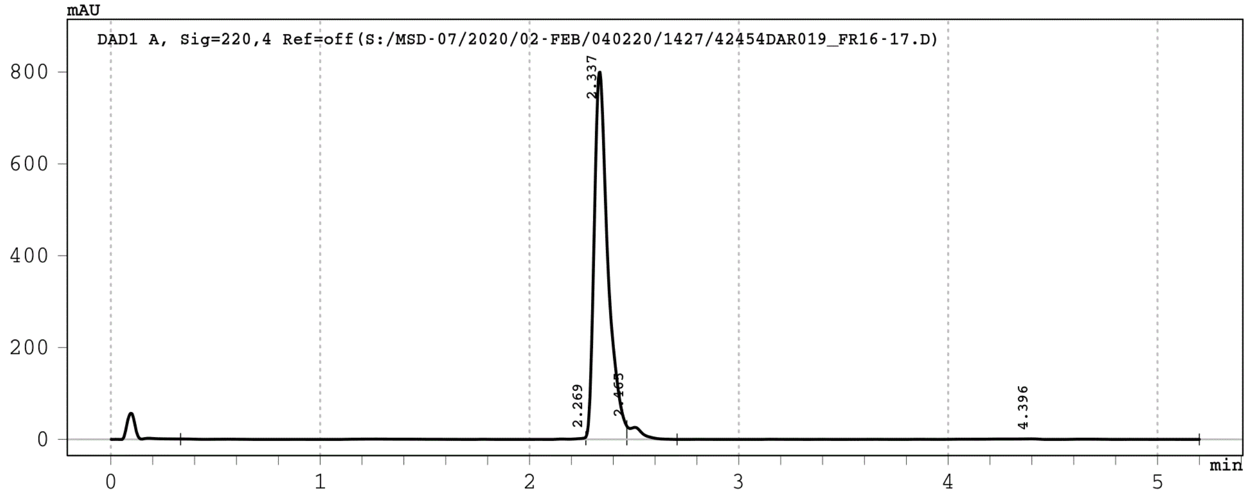


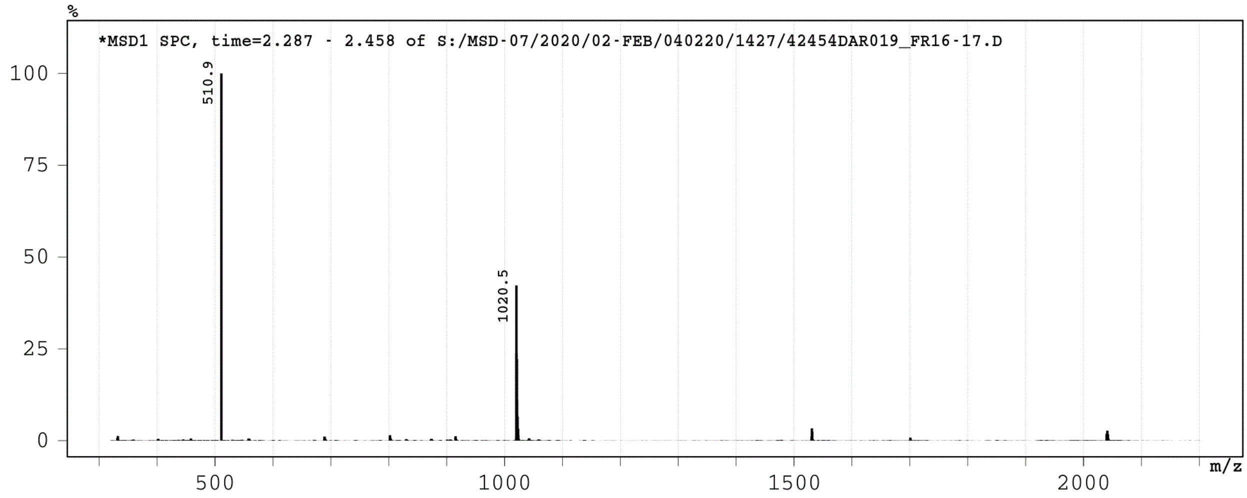


Signal 1: DAD1 A, Sig=220,4 Ref=off

| Peak # | RT [min] | Width [min] | Area | Area % |
|---|---|---|---|---|
| 1 | 2.269 | 1.94 | 34.1 | 0.9 |
| 2 | 2.337 | 0.20 | 3663.3 | 95.0 |
| 3 | 2.465 | 0.24 | 131.1 | 3.4 |
| 4 | 4.396 | 2.50 | 29.2 | 0.8 |

Data File : S:/MSD-07/2020/02-FEB/040220/1427/42454DAR020_FR2->

Injection Date : 04-Feb-20, 17:05:47
Sample Name : 42454DAR020_FR26-27
Remarks : Baseline corrected
Acq. Method : D:/METHODS/5-95-6M.M
column used : Je_559
Analysis Method : C:/CHEM32/1/METHODS/JPT-LCMS.M

Sample Info : 42454_020-Dr. Martine Darwish H-SNFHFMCAL-OH

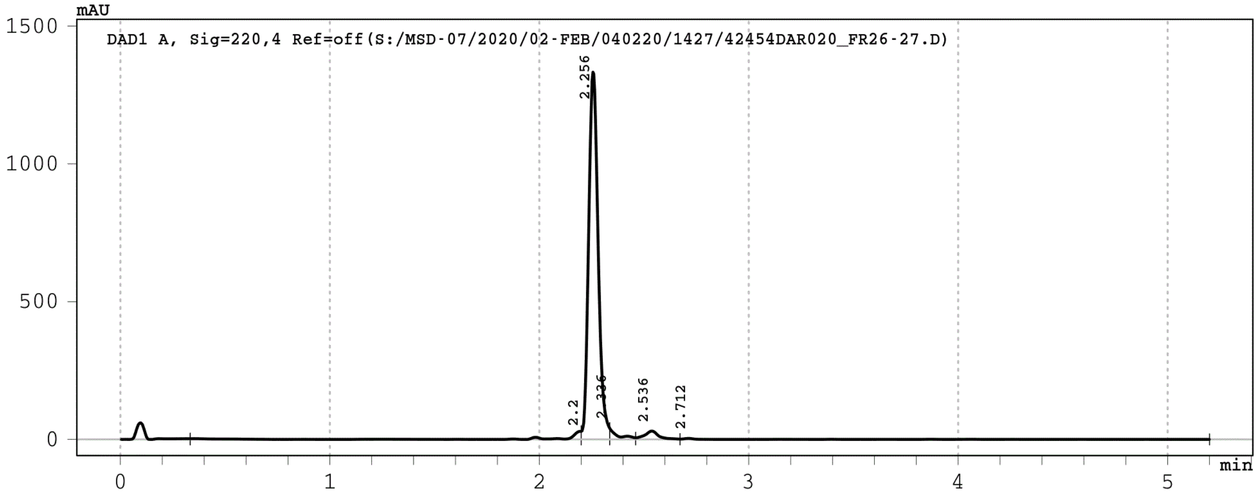


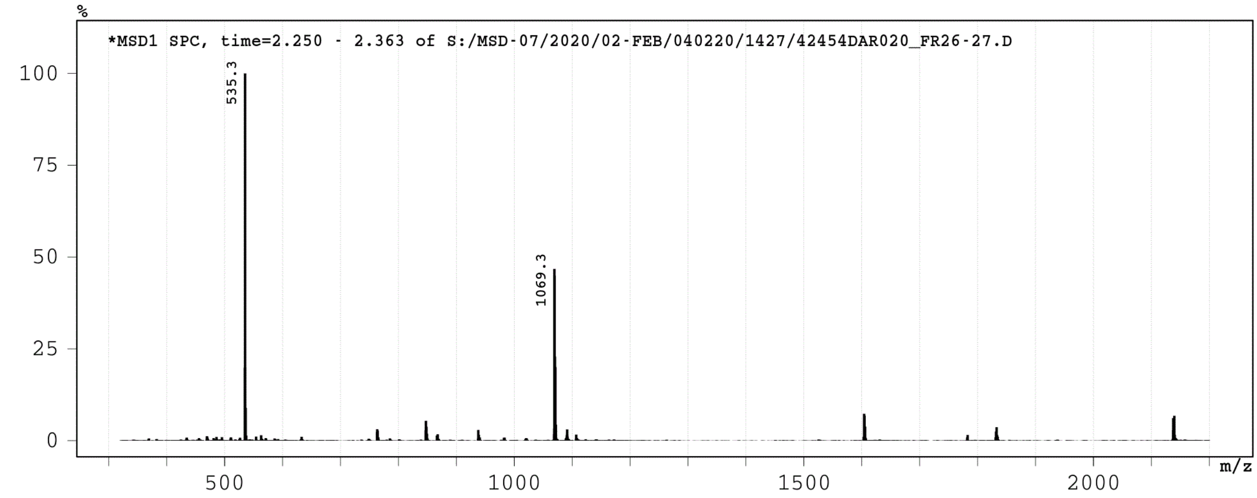


Signal 1: DAD1 A, Sig=220,4 Ref=off

| Peak # | RT [min] | Width [min] | Area | Area % |
|---|---|---|---|---|
| 1 | 2.200 | 1.87 | 142.8 | 3.1 |
| 2 | 2.256 | 0.14 | 4209.8 | 91.1 |
| 3 | 2.336 | 0.12 | 104.4 | 2.3 |
| 4 | 2.536 | 0.21 | 145.2 | 3.1 |
| 5 | 2.712 | 2.53 | 20.2 | 0.4 |

Data File : S:/MSD-07/2020/02-FEB/040220/1427/42454DAR021_FR3->

Injection Date : 04-Feb-20, 17:14:11
Sample Name : 42454DAR021_FR39
Remarks : Baseline corrected
Acq. Method : D:/METHODS/5-95-6M.M
column used : Je_559
Analysis Method : C:/CHEM32/1/METHODS/JPT-LCMS.M

Sample Info : 42454_021-Dr. Martine Darwish H-IGYSLLPLIL-OH

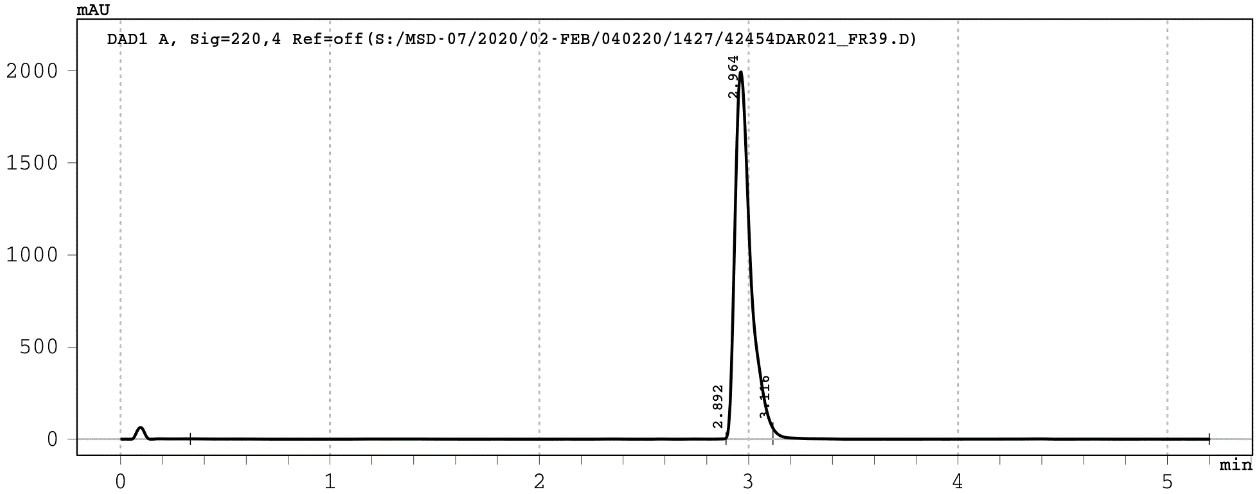


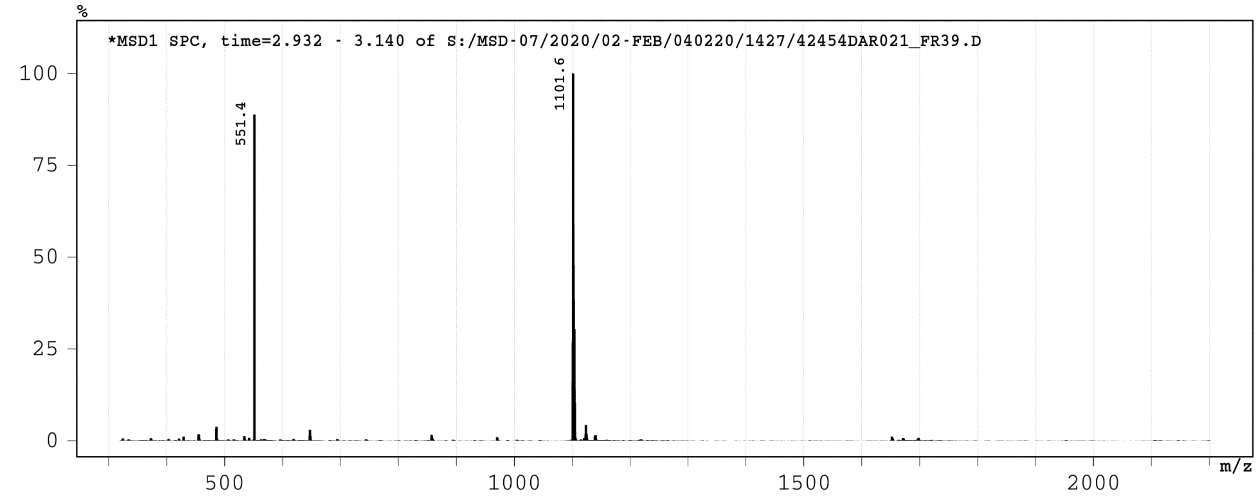


Signal 1: DAD1 A, Sig=220,4 Ref=off

| Peak # | RT [min] | Width [min] | Area | Area % |
|---|---|---|---|---|
| 1 | 2.892 | 2.56 | 46.6 | 0.4 |
| 2 | 2.964 | 0.22 | 10399.1 | 98.0 |
| 3 | 3.116 | 2.08 | 166.0 | 1.6 |

Data File : S:/MSD-07/2020/02-FEB/040220/1427/42454DAR022_FR5->

Injection Date : 04-Feb-20, 17:22:35
Sample Name : 42454DAR022_FR52
Remarks : Baseline corrected
Acq. Method : D:/METHODS/5-95-6M.M
column used : Je_559
Analysis Method : C:/CHEM32/1/METHODS/JPT-LCMS.M

Sample Info : 42454_022-Dr. Martine Darwish H-KSFHFYCPL-OH

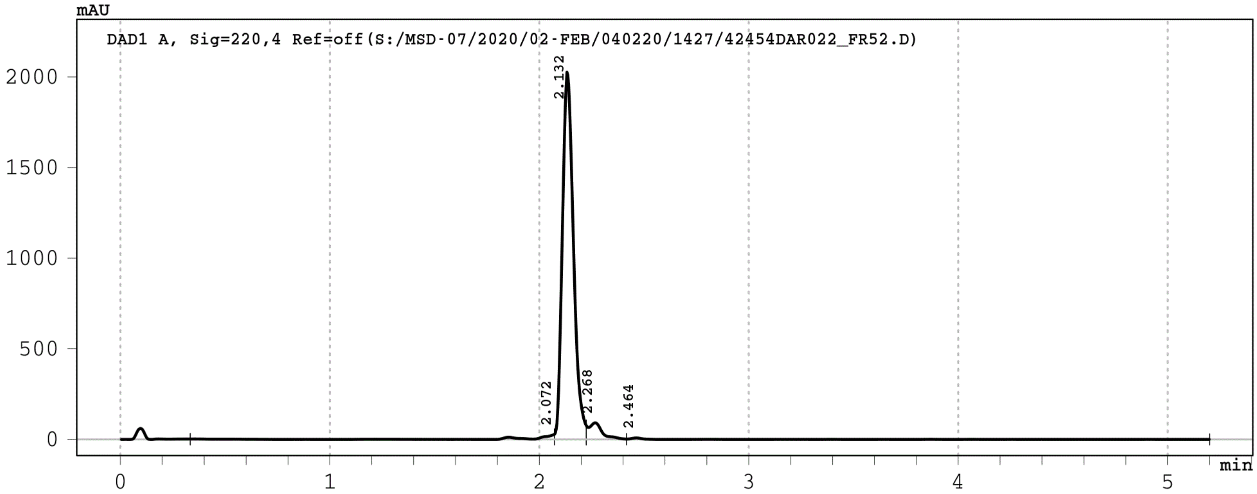


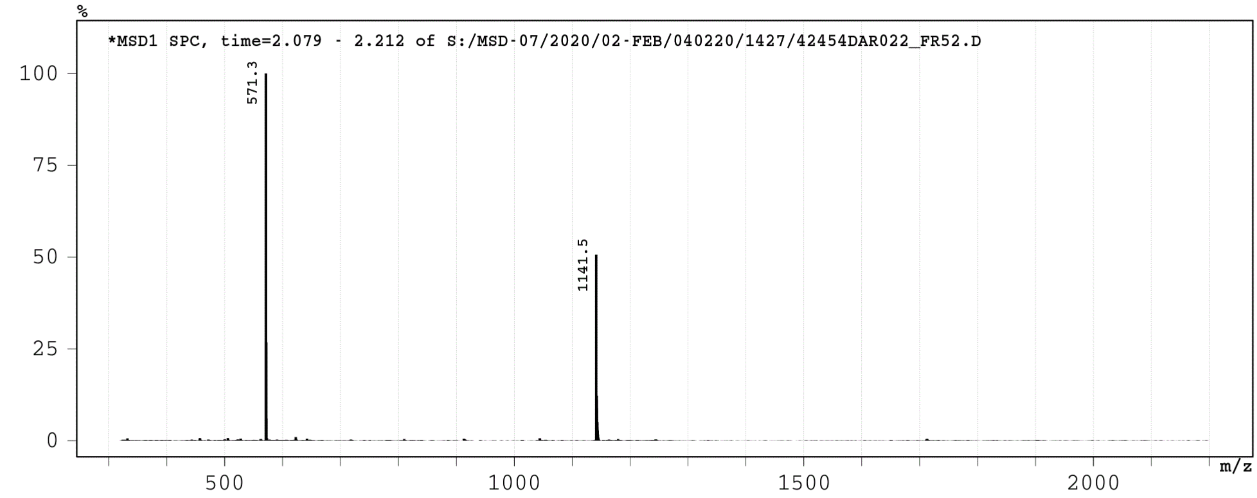


Signal 1: DAD1 A, Sig=220,4 Ref=off

| Peak # | RT [min] | Width [min] | Area | Area % |
|---|---|---|---|---|
| 1 | 2.072 | 1.74 | 176.4 | 2.2 |
| 2 | 2.132 | 0.15 | 7476.5 | 92.1 |
| 3 | 2.268 | 0.19 | 426.4 | 5.3 |
| 4 | 2.464 | 2.78 | 39.4 | 0.5 |

| | | |
|---|---|---|
| Data File | : | S:/MSD-07/2020/02-FEB/040220/1427/42454DAR023_FR1-> |
| Injection Date | : | 04-Feb-20, 19:37:02 |
| Sample Name | : | 42454DAR023_FR18 |
| Remarks | : | Baseline corrected |
| Acq. Method | : | D:/METHODS/5-95-6M.M |
| column used | : | Je_559 |
| Analysis Method | : | C:/CHEM32/1/METHODS/JPT-LCMS.M |
| Sample Info | : | 42454_023-Dr. Martine Darwish H-MMKYYYESV-OH |

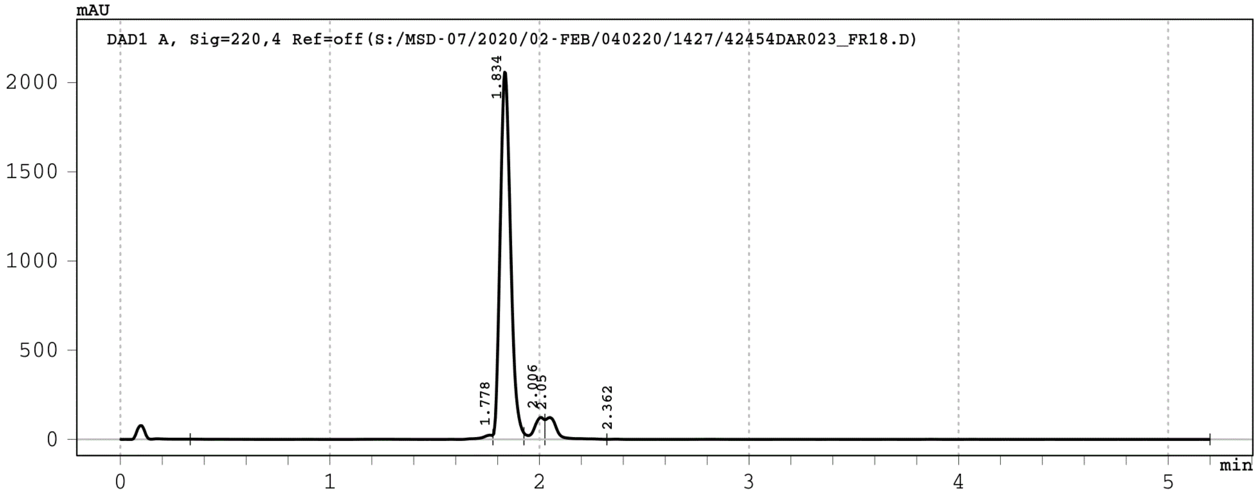


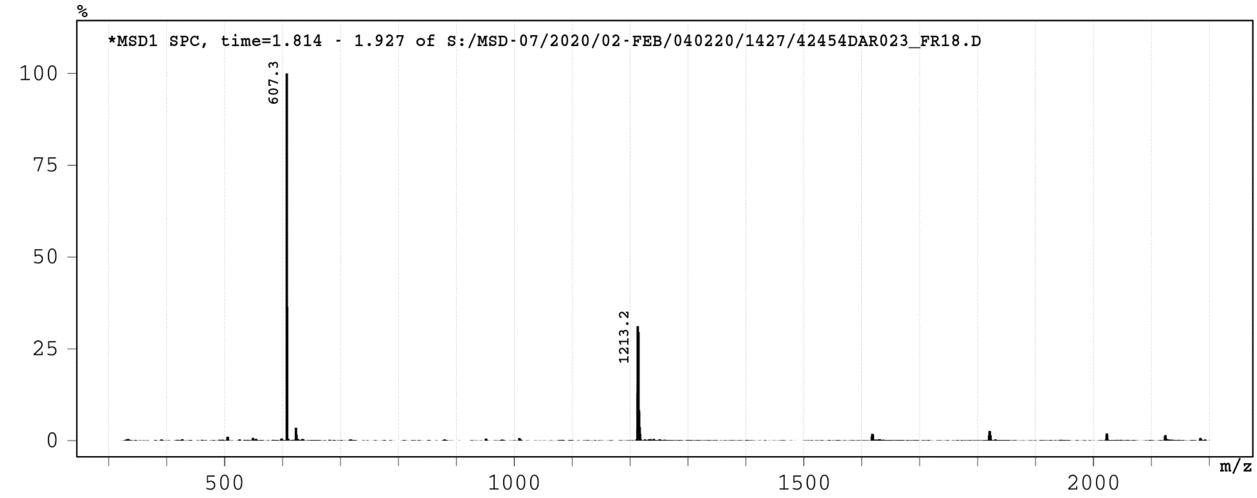


Signal 1: DAD1 A, Sig=220,4 Ref=off

| Peak # | RT [min] | Width [min] | Area | Area % |
|---|---|---|---|---|
| 1 | 1.778 | 1.44 | 104.0 | 1.3 |
| 2 | 1.834 | 0.15 | 7119.1 | 87.7 |
| 3 | 2.006 | 0.10 | 417.4 | 5.1 |
| 4 | 2.050 | 0.30 | 465.1 | 5.7 |
| 5 | 2.362 | 2.88 | 15.2 | 0.2 |

Data File : S:/MSD-07/2020/02-FEB/180220/1438/42454DAR024_FR6->

Injection Date : 18-Feb-20, 12:52:34
Sample Name : 42454DAR024_FR65-68
Remarks : Baseline corrected
Acq. Method : D:/METHODS/5-95-6M.M
column used : Je_559
Analysis Method : C:/CHEM32/1/METHODS/JPT-LCMS.M

Sample Info : 42454_024-Dr. Martine Darwish H-QILVFLILL-OH

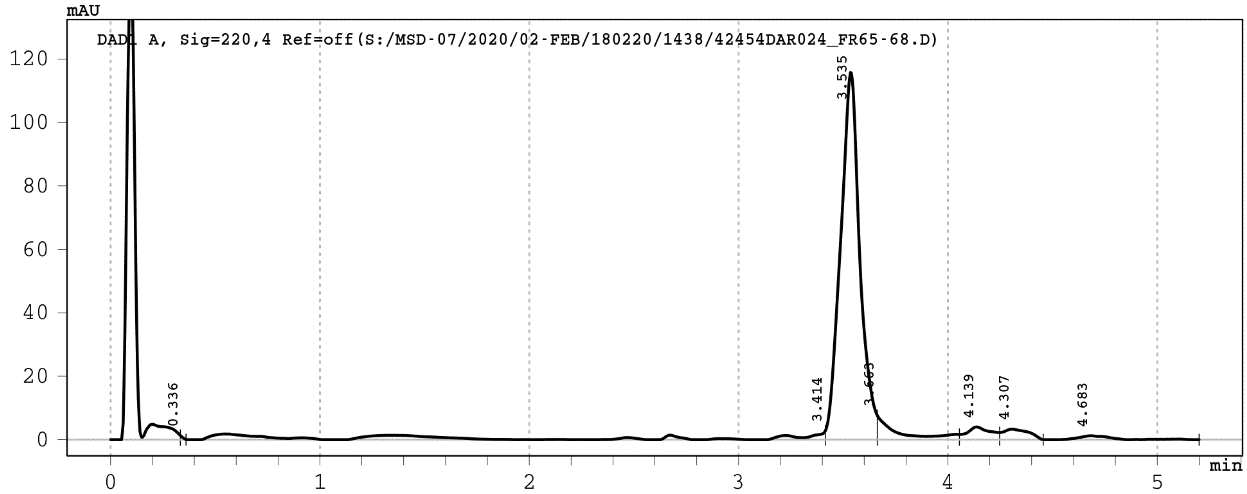


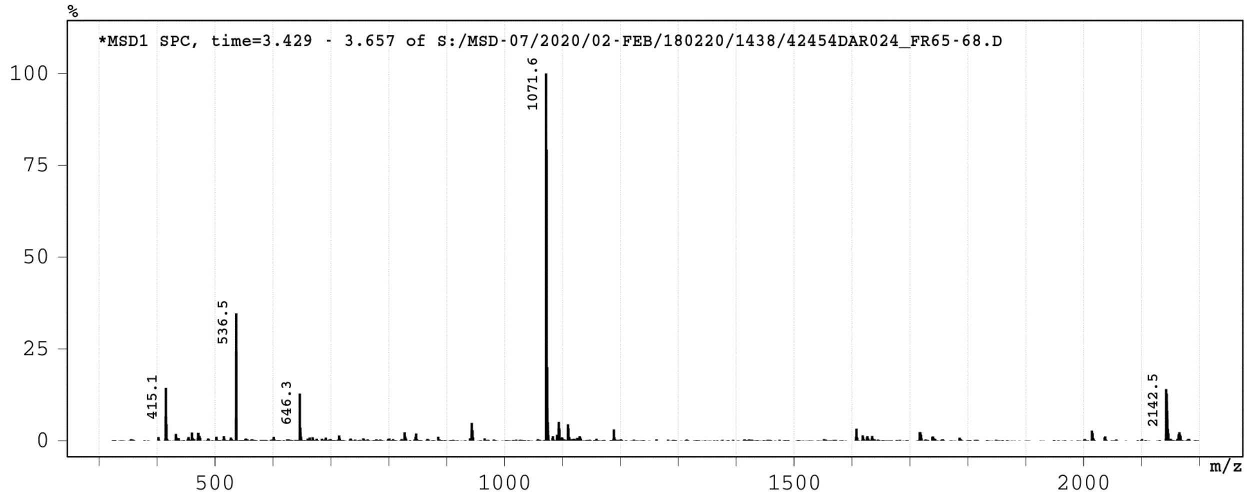


Signal 1: DAD1 A, Sig=220,4 Ref=off

| Peak # | RT [min] | Width [min] | Area | Area % |
|---|---|---|---|---|
| 1 | 0.336 | 0.03 | 0.8 | 0.1 |
| 2 | 3.414 | 3.05 | 92.9 | 9.7 |
| 3 | 3.535 | 0.25 | 742.6 | 77.5 |
| 4 | 3.663 | 0.39 | 48.8 | 5.1 |
| 5 | 4.139 | 0.19 | 31.8 | 3.3 |
| 6 | 4.307 | 0.21 | 28.5 | 3.0 |
| 7 | 4.683 | 0.74 | 13.1 | 1.4 |

Data File : S:/MSD-07/2020/02-FEB/040220/1427/42454DAR025_FR4->

Injection Date : 04-Feb-20, 19:53:52
Sample Name : 42454DAR025_FR44-45
Remarks : Baseline corrected
Acq. Method : D:/METHODS/5-95-6M.M
column used : Je_559
Analysis Method : C:/CHEM32/1/METHODS/JPT-LCMS.M

Sample Info : 42454_025-Dr. Martine Darwish H-TATRNDFTL-OH

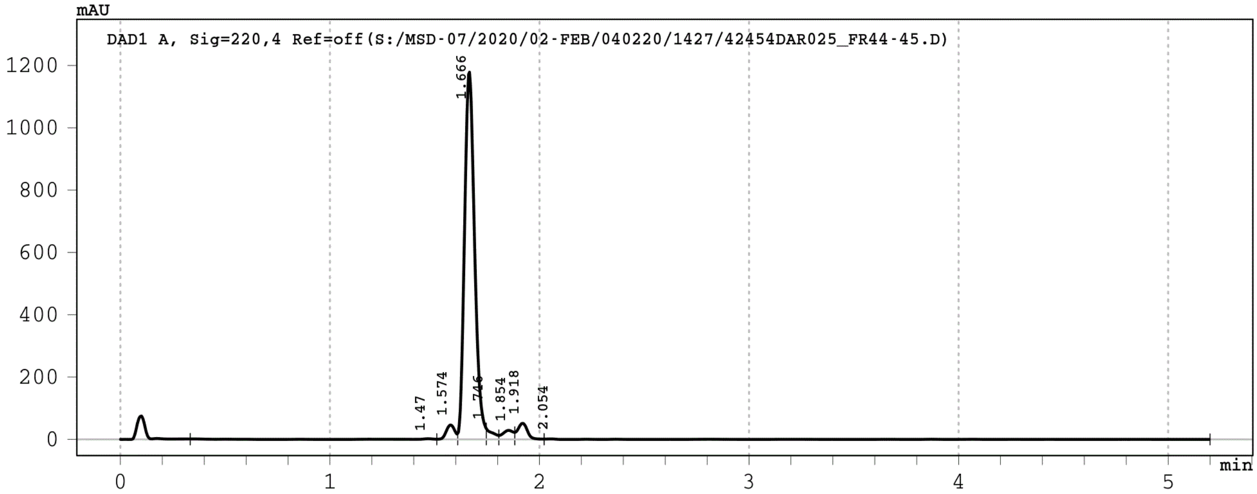


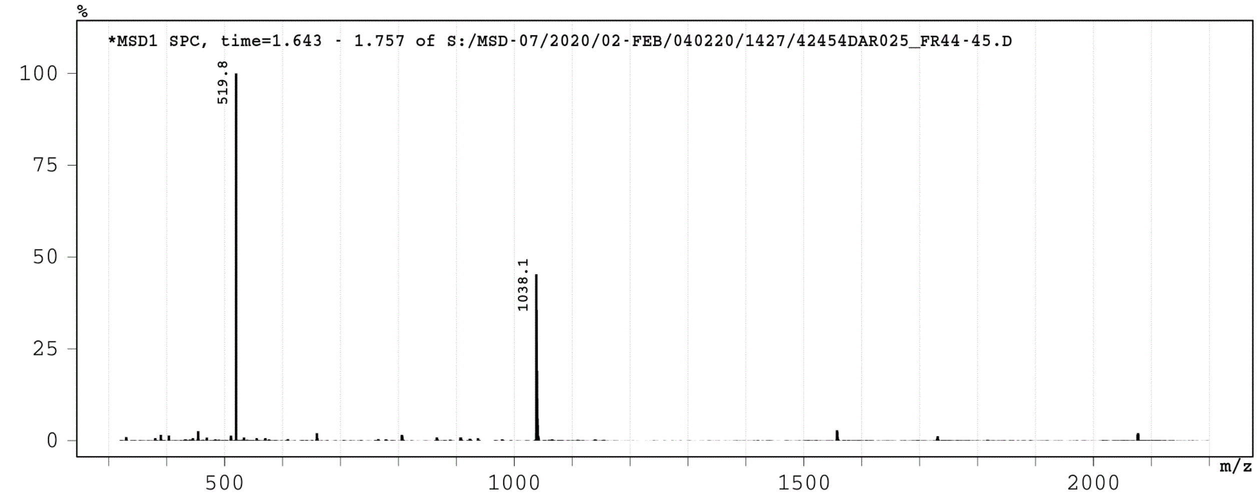


Signal 1: DAD1 A, Sig=220,4 Ref=off

| Peak # | RT [min] | Width [min] | Area | Area % |
|---|---|---|---|---|
| 1 | 1.470 | 1.18 | 28.6 | 0.6 |
| 2 | 1.574 | 0.10 | 132.0 | 3.0 |
| 3 | 1.666 | 0.14 | 3937.2 | 88.2 |
| 4 | 1.746 | 0.06 | 75.8 | 1.7 |
| 5 | 1.854 | 0.08 | 104.0 | 2.3 |
| 6 | 1.918 | 0.14 | 168.4 | 3.8 |
| 7 | 2.054 | 3.18 | 19.5 | 0.4 |

Data File : S:/MSD-07/2020/02-FEB/040220/1427/42454DAR026_FR5->

Injection Date : 04-Feb-20, 20:02:18
Sample Name : 42454DAR026_FR58-59
Remarks : Baseline corrected
Acq. Method : D:/METHODS/5-95-6M.M
column used : Je_559
Analysis Method : C:/CHEM32/1/METHODS/JPT-LCMS.M

Sample Info : 42454_026-Dr. Martine Darwish H-MAPIDHTTM-OH

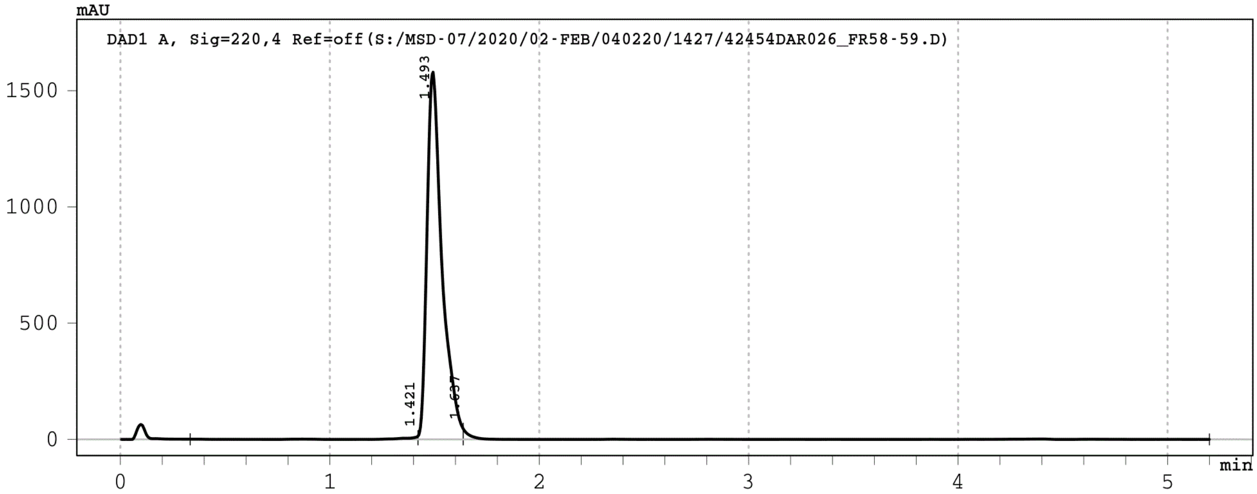


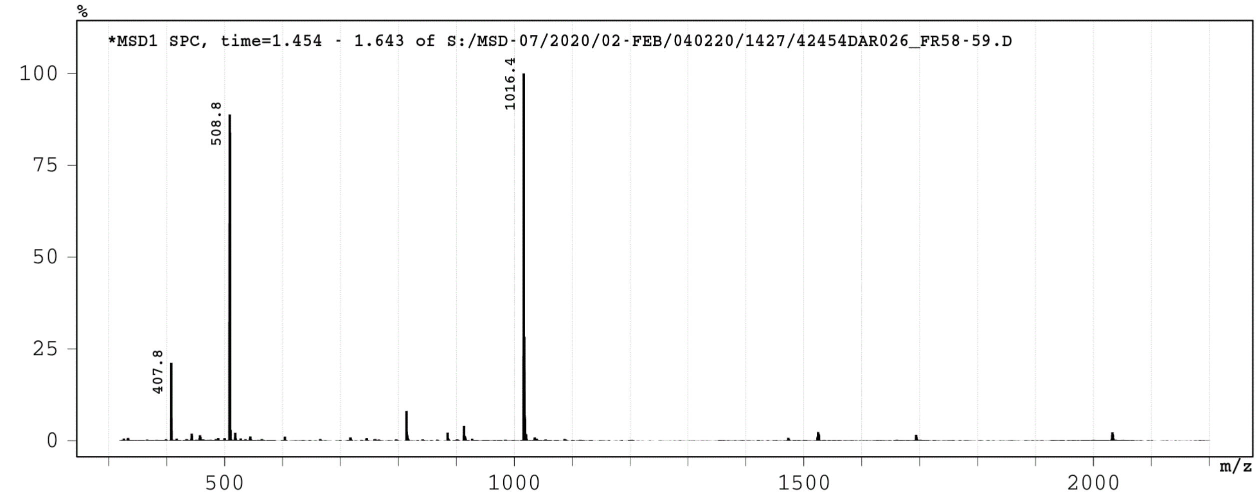


Signal 1: DAD1 A, Sig=220,4 Ref=off

| Peak # | RT [min] | Width [min] | Area | Area % |
|---|---|---|---|---|
| 1 | 1.421 | 1.09 | 62.8 | 0.8 |
| 2 | 1.493 | 0.22 | 7887.2 | 97.6 |
| 3 | 1.637 | 3.56 | 129.3 | 1.6 |

Data File : S:/MSD-07/2020/02-FEB/140220/1435/42454DAR028_FR1->

Injection Date : 14-Feb-20, 23:36:26
Sample Name : 42454DAR028_FR12-14
Remarks : Baseline corrected
Acq. Method : D:/METHODS/5-95-6M.M
column used : Je_559
Analysis Method : C:/CHEM32/1/METHODS/JPT-LCMS.M

Sample Info : 42454_028-Dr. Martine Darwish H-IVYKRSAV-OH

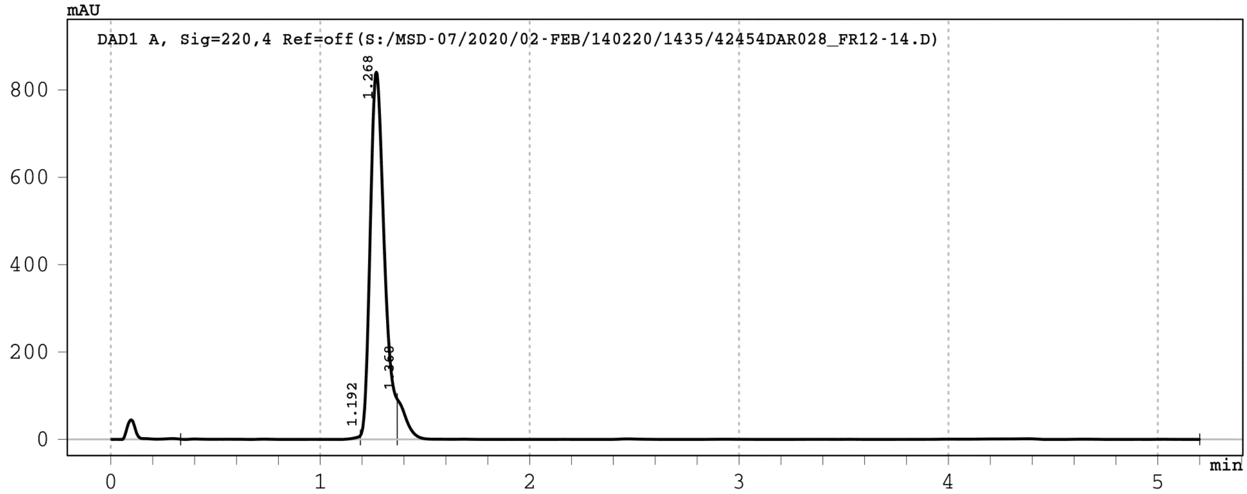


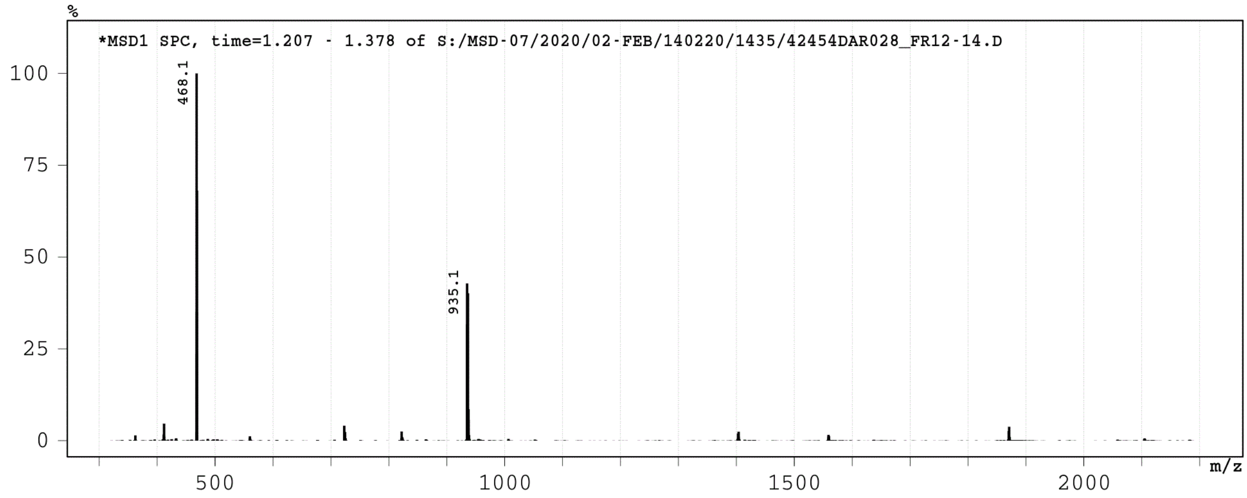


Signal 1: DAD1 A, Sig=220,4 Ref=off

| Peak # | RT [min] | Width [min] | Area | Area % |
|---|---|---|---|---|
| 1 | 1.192 | 0.86 | 22.2 | 0.5 |
| 2 | 1.268 | 0.18 | 3959.6 | 92.1 |
| 3 | 1.368 | 3.83 | 318.6 | 7.4 |

Data File : S:/MSD-07/2020/02-FEB/040220/1427/42454DAR029_FR7->

Injection Date : 04-Feb-20, 20:10:43
Sample Name : 42454DAR029_FR79
Remarks : Baseline corrected
Acq. Method : D:/METHODS/5-95-6M.M
column used : Je_559
Analysis Method : C:/CHEM32/1/METHODS/JPT-LCMS.M

Sample Info : 42454_029-Dr. Martine Darwish H-YQPLYSSSL-OH

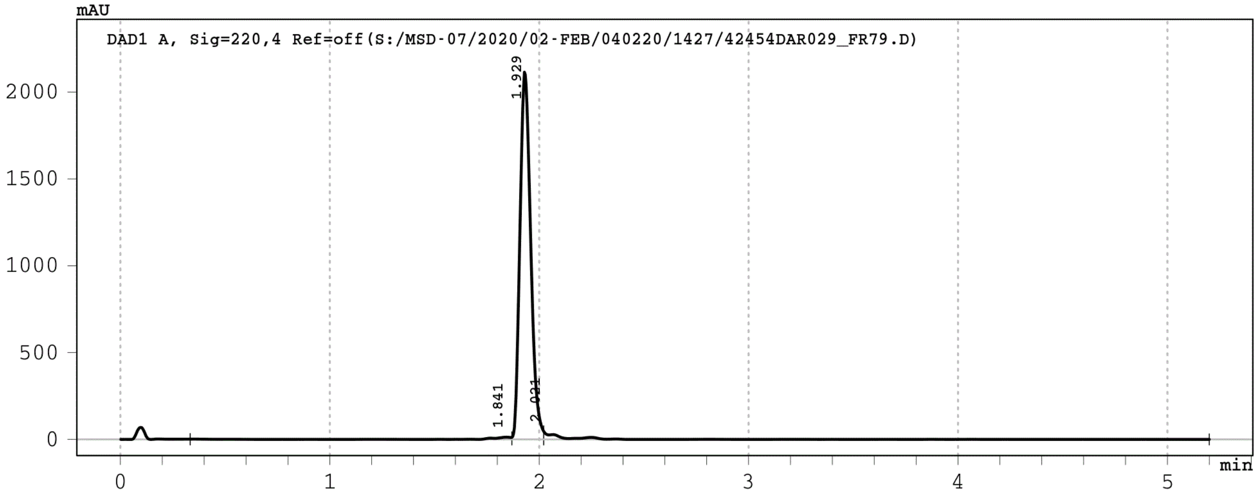


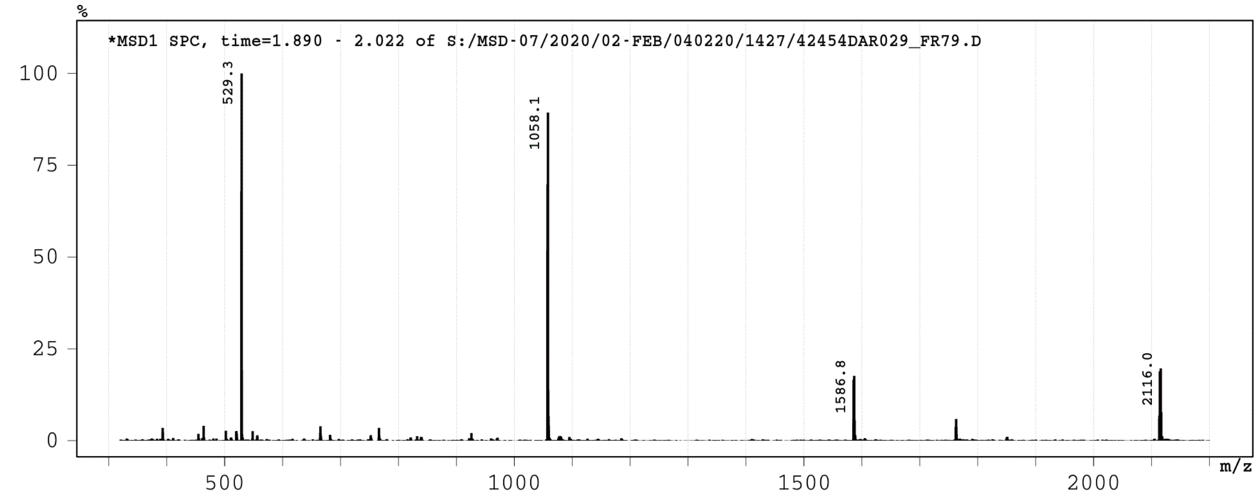


Signal 1: DAD1 A, Sig=220,4 Ref=off

| Peak # | RT [min] | Width [min] | Area | Area % |
|---|---|---|---|---|
| 1 | 1.841 | 1.54 | 94.9 | 1.2 |
| 2 | 1.929 | 0.15 | 7548.7 | 95.8 |
| 3 | 2.021 | 3.18 | 238.6 | 3.0 |

Data File : S:/MSD-07/2020/02-FEB/040220/1427/42454DAR030_FR9->

Injection Date : 04-Feb-20, 20:19:07
Sample Name : 42454DAR030_FR99-100
Remarks : Baseline corrected
Acq. Method : D:/METHODS/5-95-6M.M
column used : Je_559
Analysis Method : C:/CHEM32/1/METHODS/JPT-LCMS.M

Sample Info : 42454_030-Dr. Martine Darwish H-KIPKFSMPGF-OH

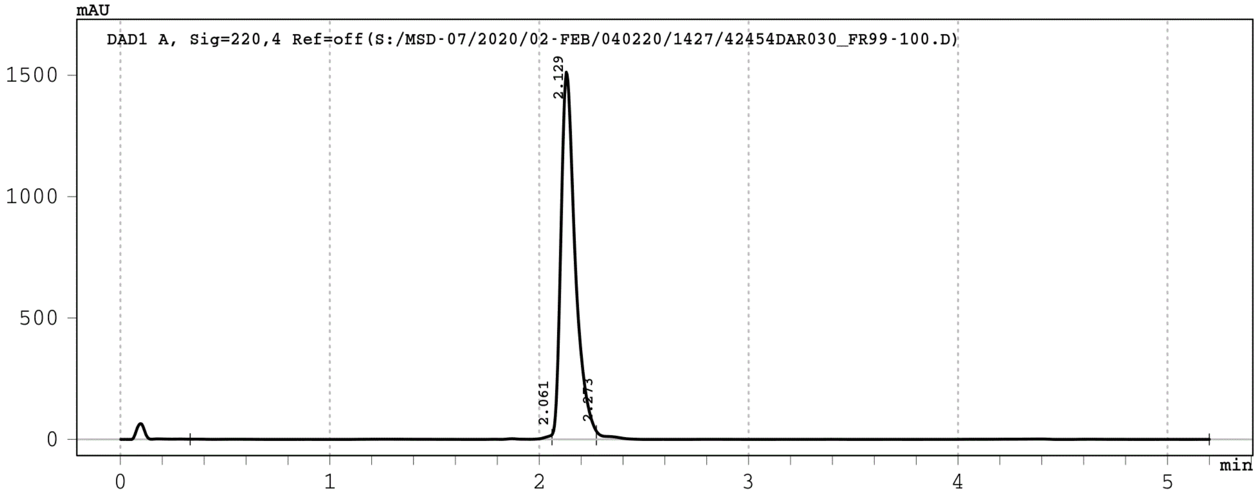


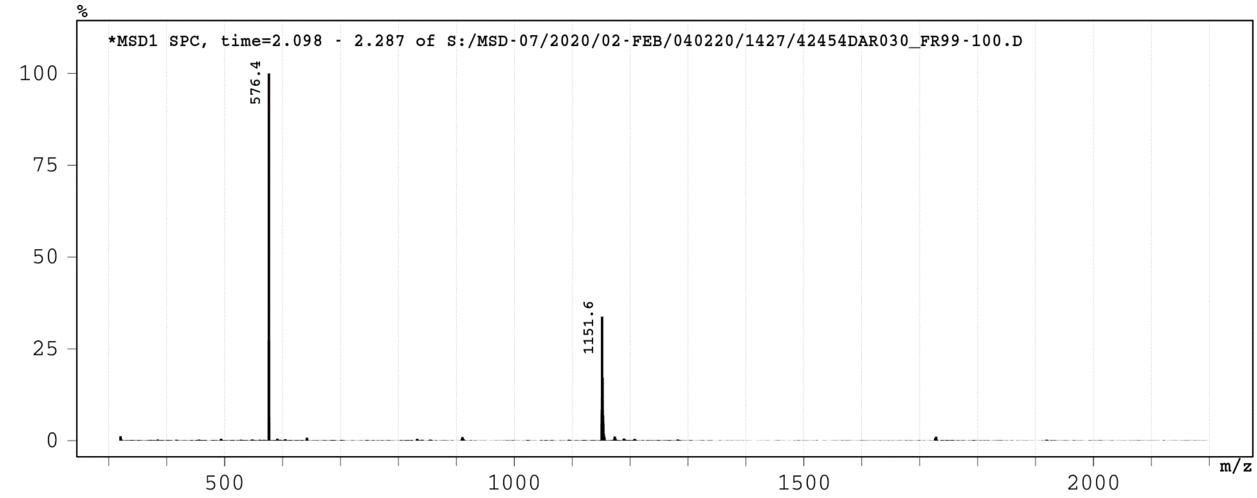


Signal 1: DAD1 A, Sig=220,4 Ref=off

| Peak # | RT [min] | Width [min] | Area | Area % |
|---|---|---|---|---|
| 1 | 2.061 | 1.73 | 75.6 | 1.0 |
| 2 | 2.129 | 0.21 | 7191.2 | 97.0 |
| 3 | 2.273 | 2.93 | 144.8 | 2.0 |

Data File : S:/MSD-07/2020/02-FEB/040220/1427/42454DAR031_FR4->

Injection Date : 04-Feb-20, 20:27:31
Sample Name : 42454DAR031_FR4-5
Remarks : Baseline corrected
Acq. Method : D:/METHODS/5-95-6M.M
column used : Je_559
Analysis Method : C:/CHEM32/1/METHODS/JPT-LCMS.M

Sample Info : 42454_031-Dr. Martine Darwish H-SAIQDNIPL-OH

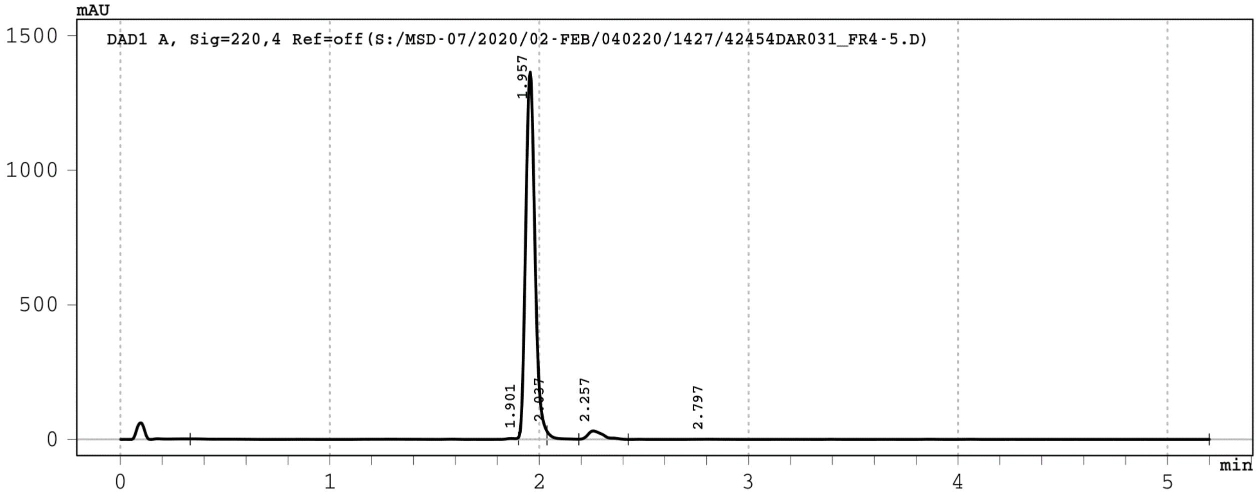


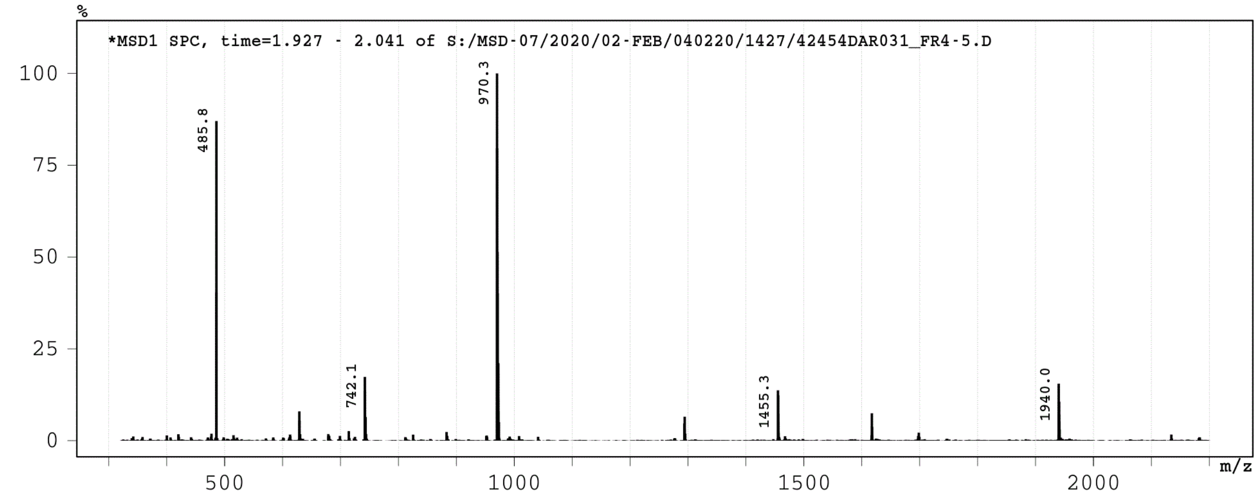


Signal 1: DAD1 A, Sig=220,4 Ref=off

| Peak # | RT [min] | Width [min] | Area | Area % |
|---|---|---|---|---|
| 1 | 1.901 | 1.57 | 45.8 | 1.0 |
| 2 | 1.957 | 0.14 | 4190.9 | 94.0 |
| 3 | 2.037 | 0.15 | 47.8 | 1.1 |
| 4 | 2.257 | 0.24 | 159.2 | 3.6 |
| 5 | 2.797 | 2.77 | 12.8 | 0.3 |

Data File : S:/MSD-07/2020/02-FEB/040220/1427/42454DAR032_FR1->

Injection Date : 04-Feb-20, 20:35:57
Sample Name : 42454DAR032_FR12-13
Remarks : Baseline corrected
Acq. Method : D:/METHODS/5-95-6M.M
column used : Je_559
Analysis Method : C:/CHEM32/1/METHODS/JPT-LCMS.M

Sample Info : 42454_032-Dr. Martine Darwish H-CVVPFTDLL-OH

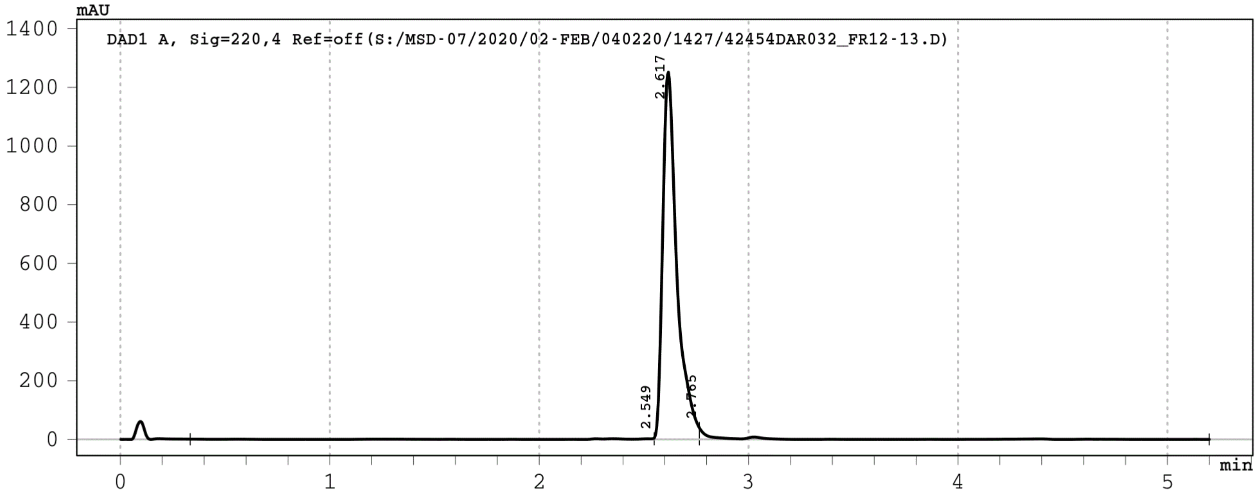


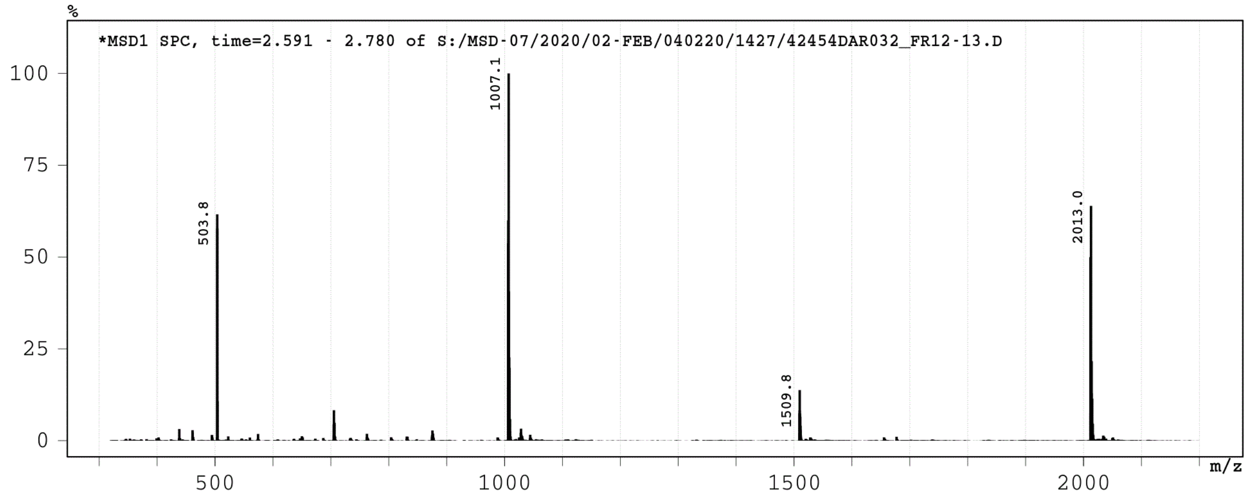


Signal 1: DAD1 A, Sig=220,4 Ref=off

| Peak # | RT [min] | Width [min] | Area | Area % |
|---|---|---|---|---|
| 1 | 2.549 | 2.22 | 55.7 | 0.9 |
| 2 | 2.617 | 0.22 | 5904.9 | 96.3 |
| 3 | 2.765 | 2.43 | 174.0 | 2.8 |

Data File : S:/MSD-07/2020/02-FEB/040220/1427/42454DAR033_FR2->

Injection Date : 04-Feb-20, 20:44:23
Sample Name : 42454DAR033_FR26-28
Remarks : Baseline corrected
Acq. Method : D:/METHODS/5-95-6M.M
column used : Je_559
Analysis Method : C:/CHEM32/1/METHODS/JPT-LCMS.M

Sample Info : 42454_033-Dr. Martine Darwish H-SAHTMLPGM-OH

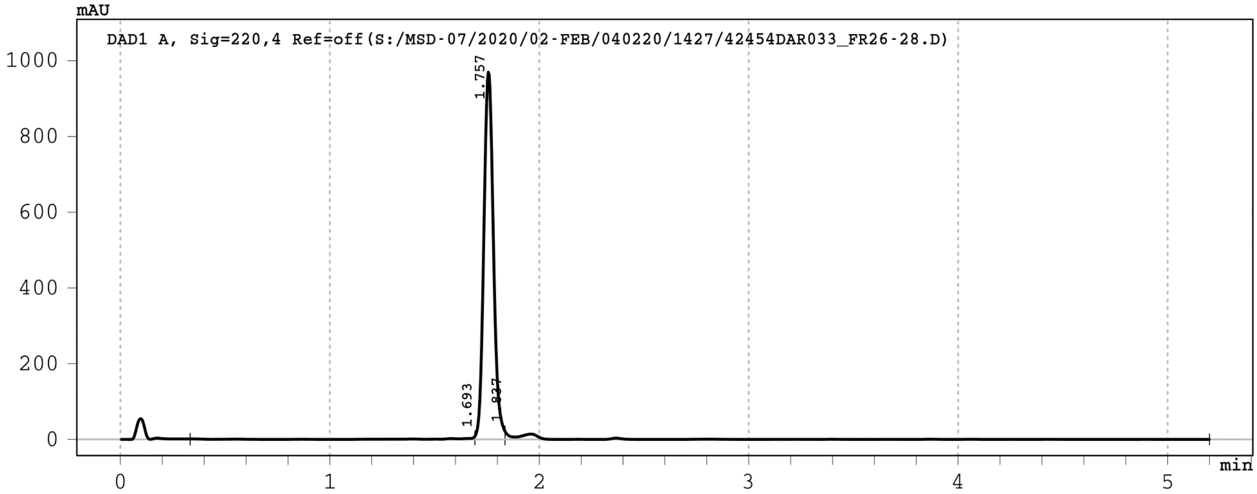


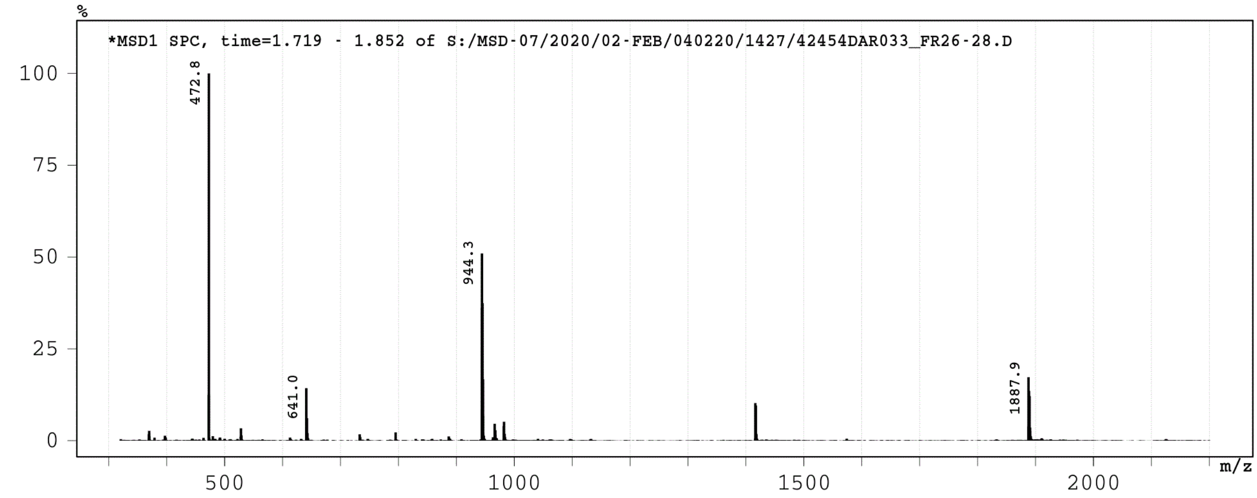


Signal 1: DAD1 A, Sig=220,4 Ref=off

| Peak # | RT [min] | Width [min] | Area | Area % |
|---|---|---|---|---|
| 1 | 1.693 | 1.36 | 47.0 | 1.4 |
| 2 | 1.757 | 0.14 | 3094.6 | 94.6 |
| 3 | 1.837 | 3.36 | 129.7 | 4.0 |

Data File : S:/MSD-07/2020/02-FEB/040220/1427/42454DAR034_FR3->

Injection Date : 04-Feb-20, 20:52:47
Sample Name : 42454DAR034_FR38-40
Remarks : Baseline corrected
Acq. Method : D:/METHODS/5-95-6M.M
column used : Je_559
Analysis Method : C:/CHEM32/1/METHODS/JPT-LCMS.M

Sample Info : 42454_034-Dr. Martine Darwish H-SSSTAAAL-OH

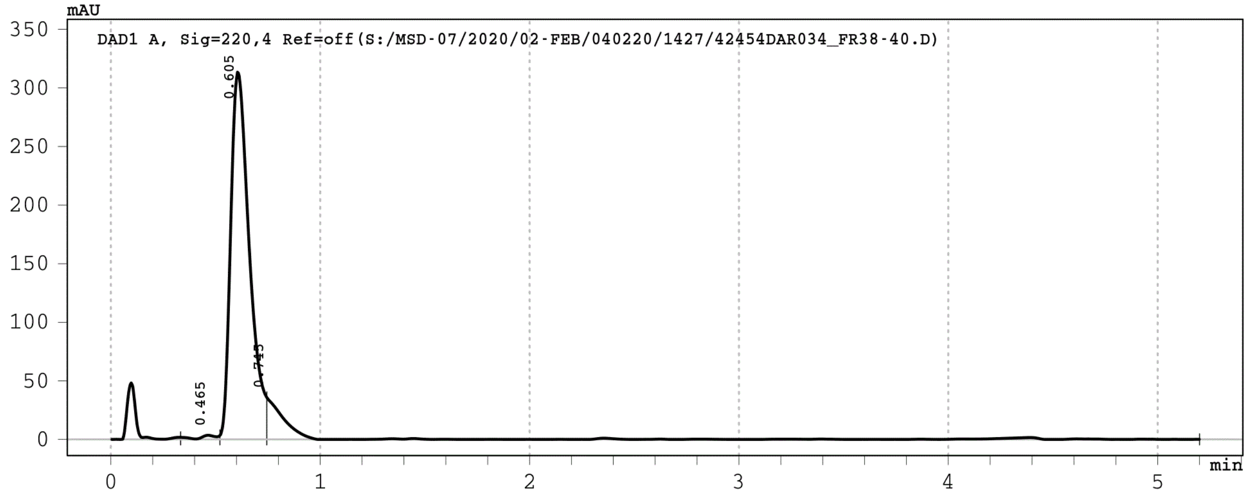


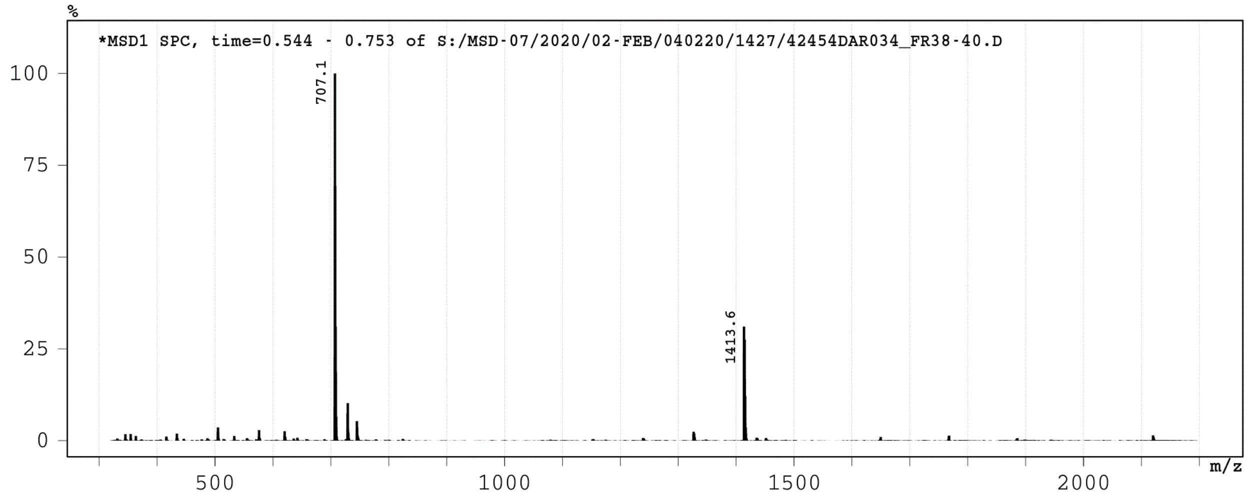


Signal 1: DAD1 A, Sig=220,4 Ref=off

| Peak # | RT [min] | Width [min] | Area | Area % |
|---|---|---|---|---|
| 1 | 0.465 | 0.19 | 20.9 | 0.9 |
| 2 | 0.605 | 0.22 | 1946.2 | 88.3 |
| 3 | 0.745 | 4.46 | 237.1 | 10.8 |

Data File : S:/MSD-07/2020/02-FEB/040220/1427/42454DAR035_FR5->

Injection Date : 04-Feb-20, 21:01:12
Sample Name : 42454DAR035_FR58-59
Remarks : Baseline corrected
Acq. Method : D:/METHODS/5-95-6M.M
column used : Je_559
Analysis Method : C:/CHEM32/1/METHODS/JPT-LCMS.M

Sample Info : 42454_035-Dr. Martine Darwish H-IVGHFYGGL-OH

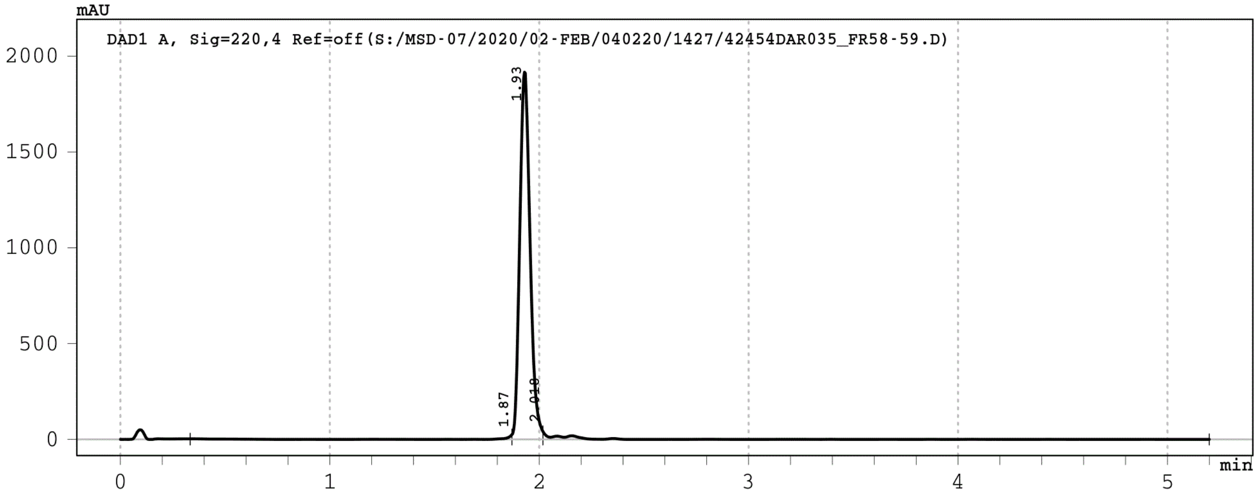


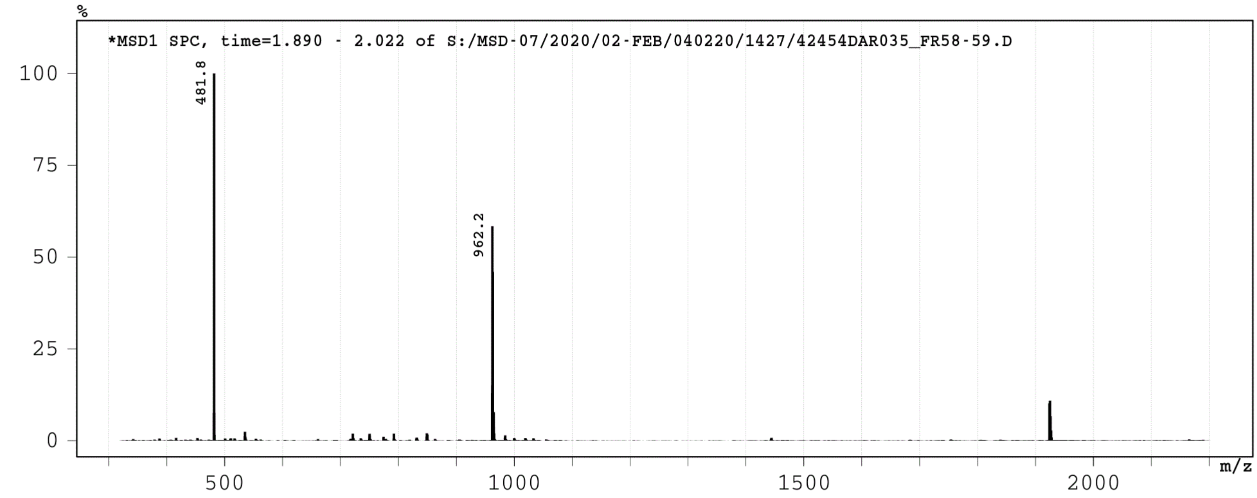


Signal 1: DAD1 A, Sig=220,4 Ref=off

| Peak # | RT [min] | Width [min] | Area | Area % |
|---|---|---|---|---|
| 1 | 1.870 | 1.54 | 79.6 | 1.2 |
| 2 | 1.930 | 0.15 | 6575.5 | 95.7 |
| 3 | 2.018 | 3.18 | 215.7 | 3.1 |

Data File : S:/MSD-07/2020/02-FEB/040220/1427/42454DAR036_FR7->

Injection Date : 04-Feb-20, 21:09:35
Sample Name : 42454DAR036_FR72-73
Remarks : Baseline corrected
Acq. Method : D:/METHODS/5-95-6M.M
column used : Je_559
Analysis Method : C:/CHEM32/1/METHODS/JPT-LCMS.M

Sample Info : 42454_036-Dr. Martine Darwish H-VVFDSVQRI-OH

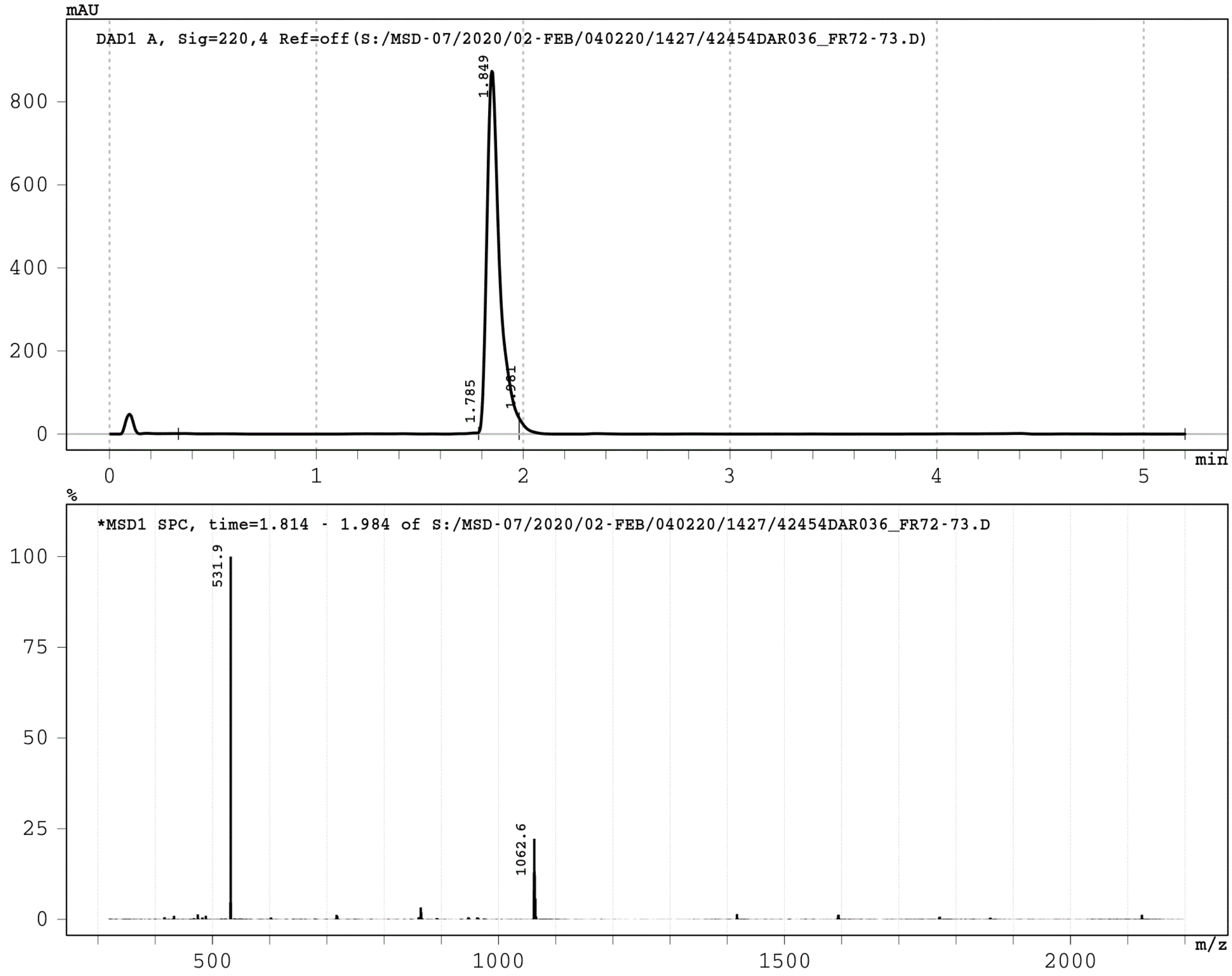


Signal 1: DAD1 A, Sig=220,4 Ref=off

| Peak # | RT [min] | Width [min] | Area | Area % |
|---|---|---|---|---|
| 1 | 1.785 | 1.45 | 38.4 | 1.0 |
| 2 | 1.849 | 0.20 | 3779.4 | 96.0 |
| 3 | 1.981 | 3.22 | 120.7 | 3.1 |

Data File : S:/MSD-07/2020/02-FEB/040220/1427/42454DAR037_FR7->

Injection Date : 04-Feb-20, 21:18:01
Sample Name : 42454DAR037_FR76-77
Remarks : Baseline corrected
Acq. Method : D:/METHODS/5-95-6M.M
column used : Je_559
Analysis Method : C:/CHEM32/1/METHODS/JPT-LCMS.M

Sample Info : 42454_037-Dr. Martine Darwish H-LSLVRTMAV-OH

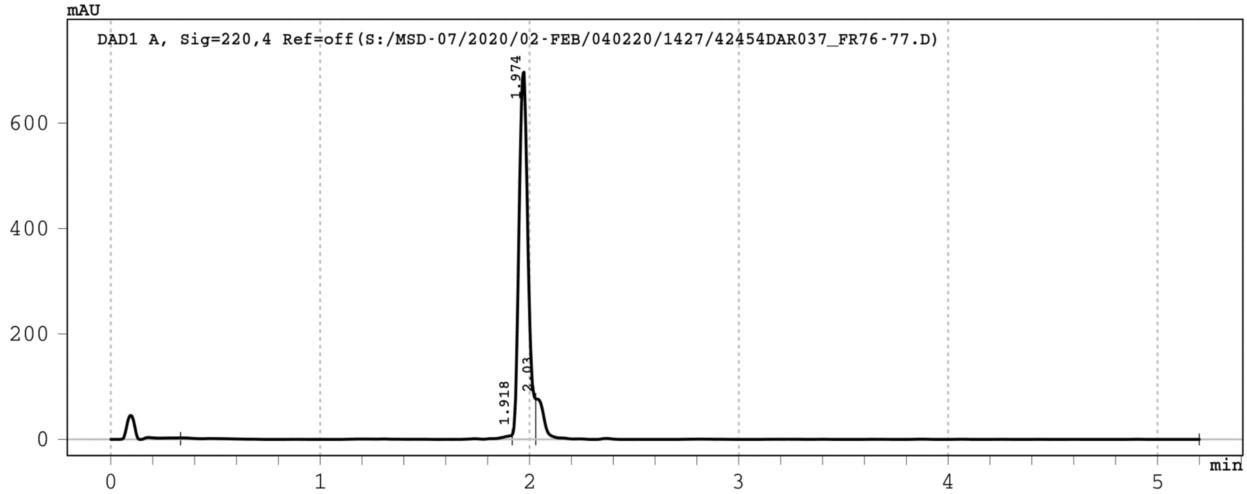


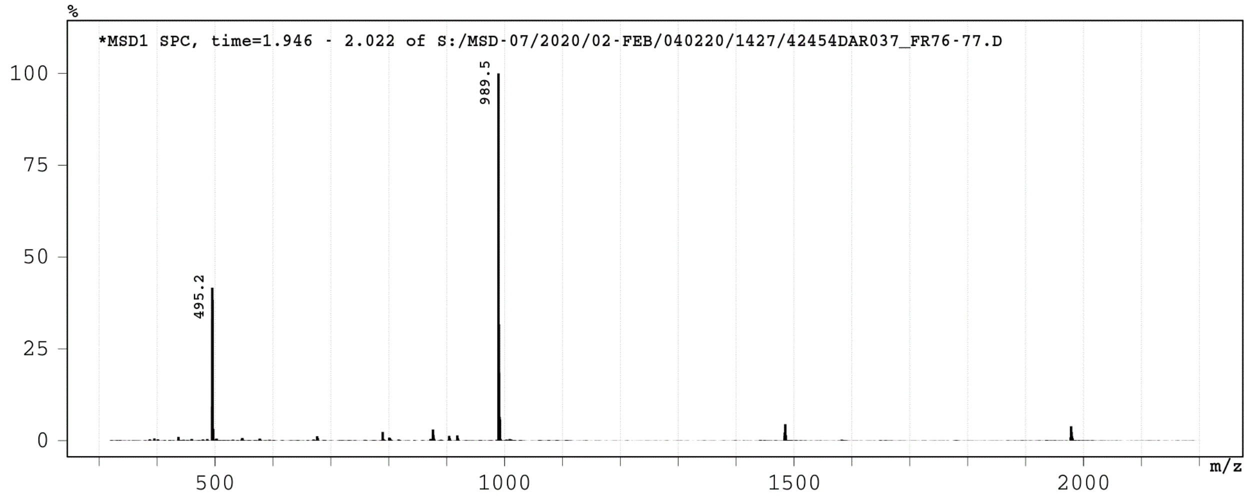


Signal 1: DAD1 A, Sig=220,4 Ref=off

| Peak # | RT [min] | Width [min] | Area | Area % |
|---|---|---|---|---|
| 1 | 1.918 | 1.58 | 70.4 | 2.9 |
| 2 | 1.974 | 0.11 | 2102.8 | 87.2 |
| 3 | 2.030 | 3.17 | 237.3 | 9.8 |

Data File : S:/MSD-07/2020/02-FEB/040220/1427/42454DAR038_FR9->

Injection Date : 04-Feb-20, 21:26:25
Sample Name : 42454DAR038_FR90
Remarks : Baseline corrected
Acq. Method : D:/METHODS/5-95-6M.M
column used : Je_559
Analysis Method : C:/CHEM32/1/METHODS/JPT-LCMS.M

Sample Info : 42454_038-Dr. Martine Darwish H-MSTQNLATV-OH

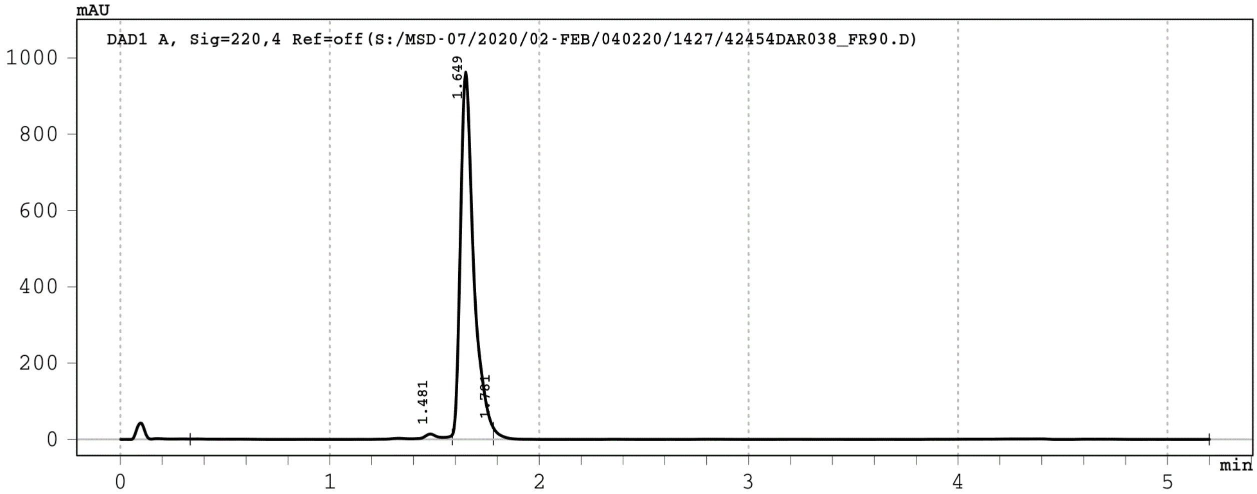


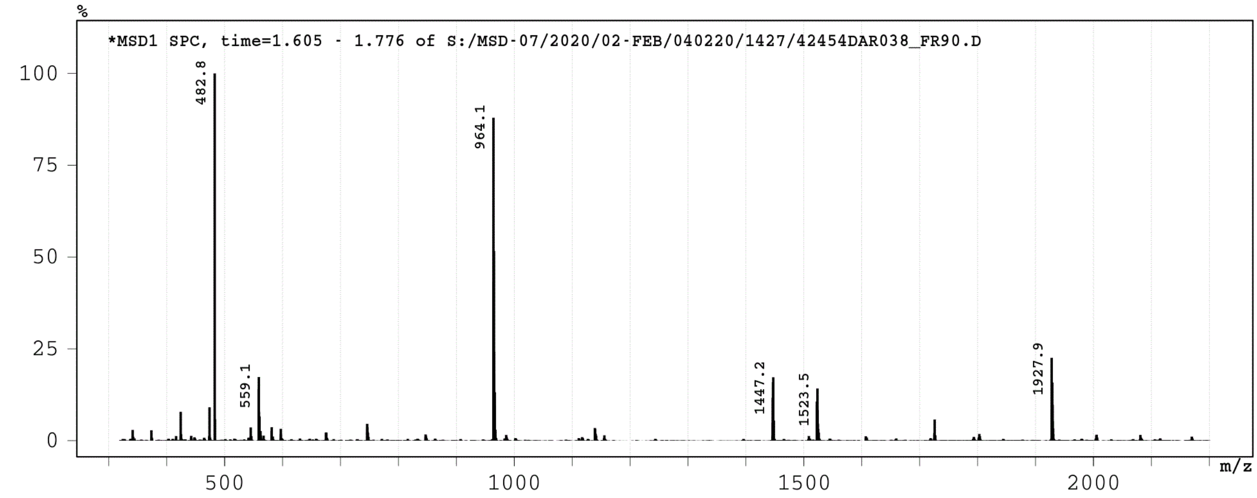


Signal 1: DAD1 A, Sig=220,4 Ref=off

| Peak # | RT [min] | Width [min] | Area | Area % |
|---|---|---|---|---|
| 1 | 1.481 | 1.25 | 108.7 | 2.4 |
| 2 | 1.649 | 0.20 | 4244.5 | 95.3 |
| 3 | 1.781 | 3.42 | 101.7 | 2.3 |

Data File : S:/MSD-07/2020/02-FEB/050220/1428/42454DAR039_FR8->

Injection Date : 06-Feb-20, 01:00:47
Sample Name : 42454DAR039_FR8
Remarks : Baseline corrected
Acq. Method : D:/METHODS/5-95-6M.M
column used : Je_559
Analysis Method : C:/CHEM32/1/METHODS/JPT-LCMS.M

Sample Info : 42454_039-Dr. Martine Darwish H-VIYSECLRV-OH

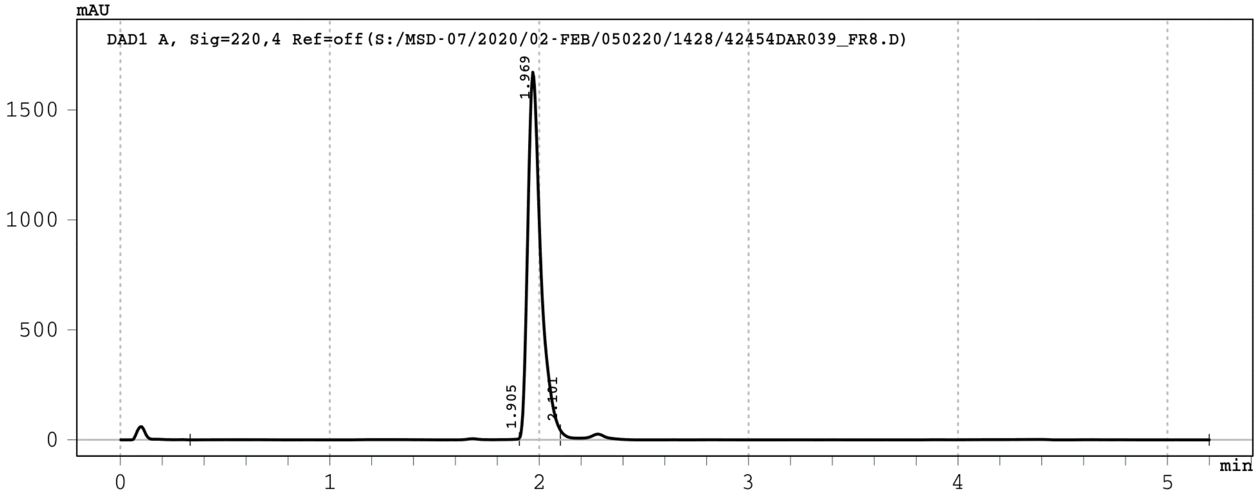


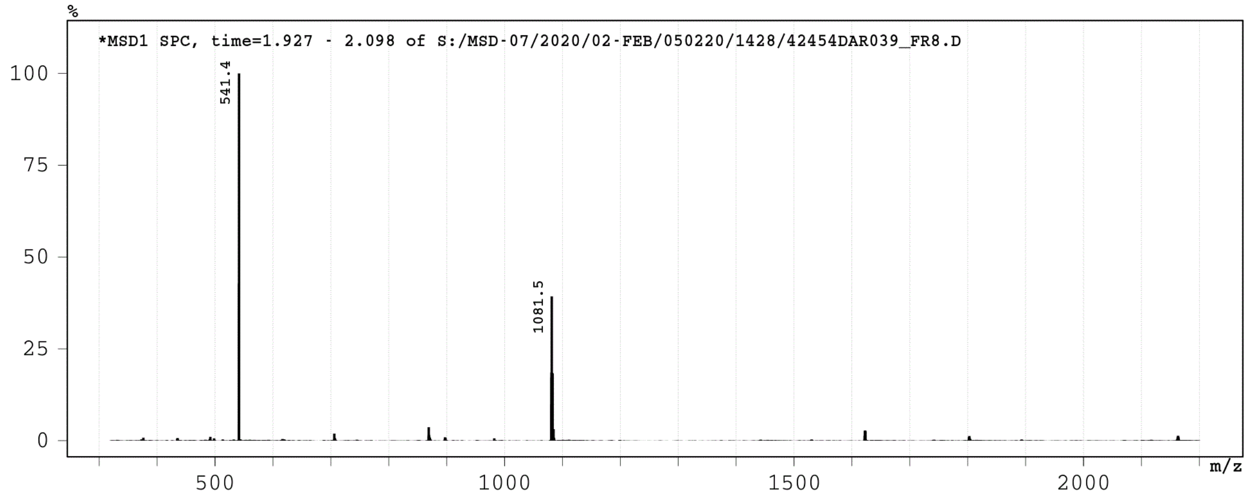


Signal 1: DAD1 A, Sig=220,4 Ref=off

| Peak # | RT [min] | Width [min] | Area | Area % |
|---|---|---|---|---|
| 1 | 1.905 | 1.57 | 60.9 | 0.8 |
| 2 | 1.969 | 0.20 | 7291.2 | 95.5 |
| 3 | 2.101 | 3.10 | 279.5 | 3.7 |

Data File : S:/MSD-07/2020/02-FEB/050220/1428/42454DAR040_FR1->

Injection Date : 06-Feb-20, 01:09:14
Sample Name : 42454DAR040_FR18-19
Remarks : Baseline corrected
Acq. Method : D:/METHODS/5-95-6M.M
column used : Je_559
Analysis Method : C:/CHEM32/1/METHODS/JPT-LCMS.M

Sample Info : 42454_040-Dr. Martine Darwish H-VKRTRFLRL-OH

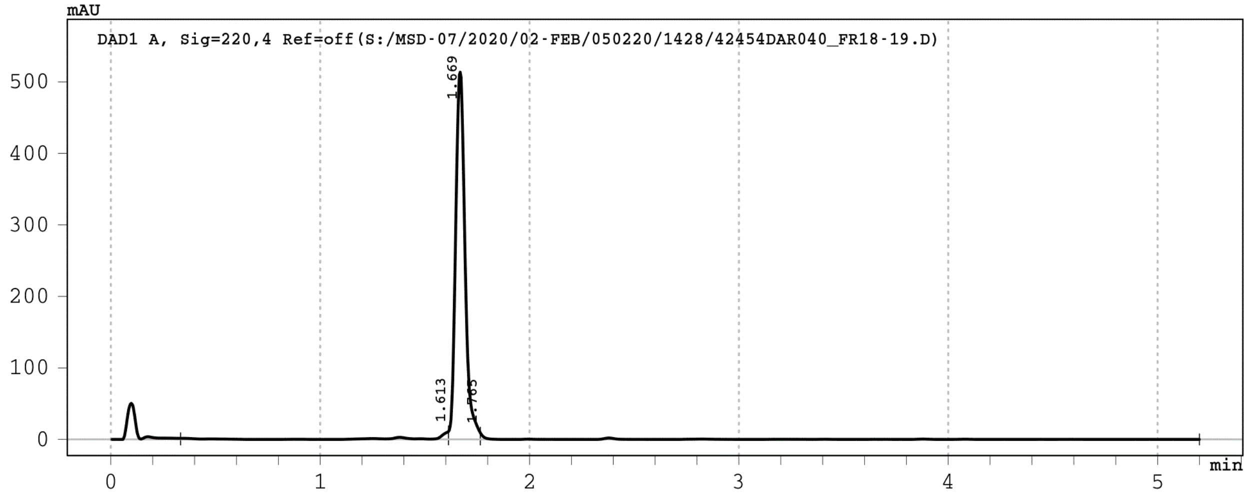


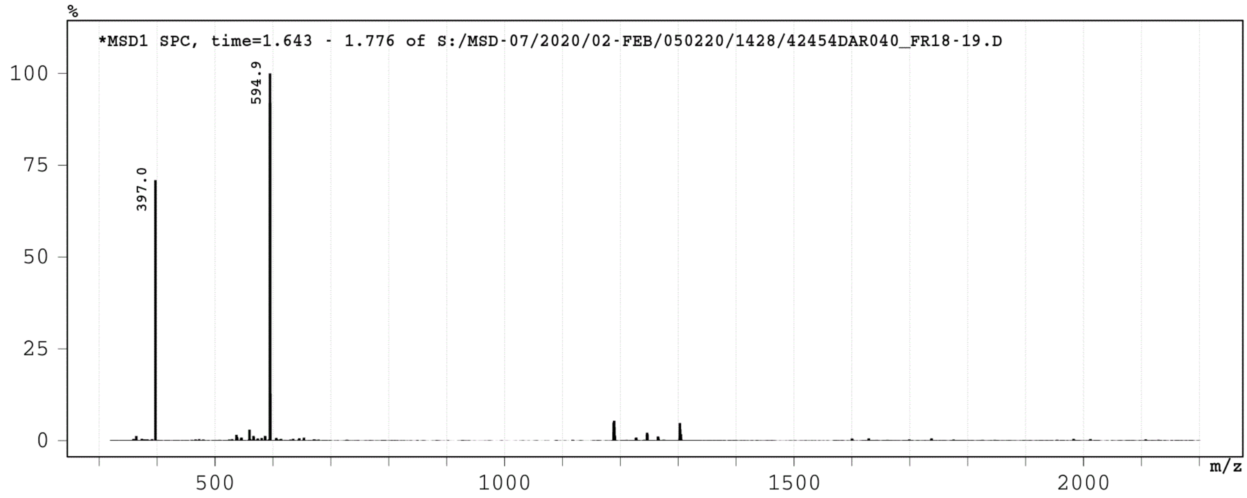


Signal 1: DAD1 A, Sig=220,4 Ref=off

| Peak # | RT [min] | Width [min] | Area | Area % |
|---|---|---|---|---|
| 1 | 1.613 | 1.28 | 60.5 | 3.6 |
| 2 | 1.669 | 0.15 | 1607.5 | 94.8 |
| 3 | 1.765 | 3.44 | 27.4 | 1.6 |

Data File : S:/MSD-07/2020/02-FEB/050220/1428/42454DAR041_FR4->

Injection Date : 06-Feb-20, 01:26:08
Sample Name : 42454DAR041_FR40
Remarks : Baseline corrected
Acq. Method : D:/METHODS/5-95-6M.M
column used : Je_559
Analysis Method : C:/CHEM32/1/METHODS/JPT-LCMS.M

Sample Info : 42454_041-Dr. Martine Darwish H-YFFEHGSSKL-OH

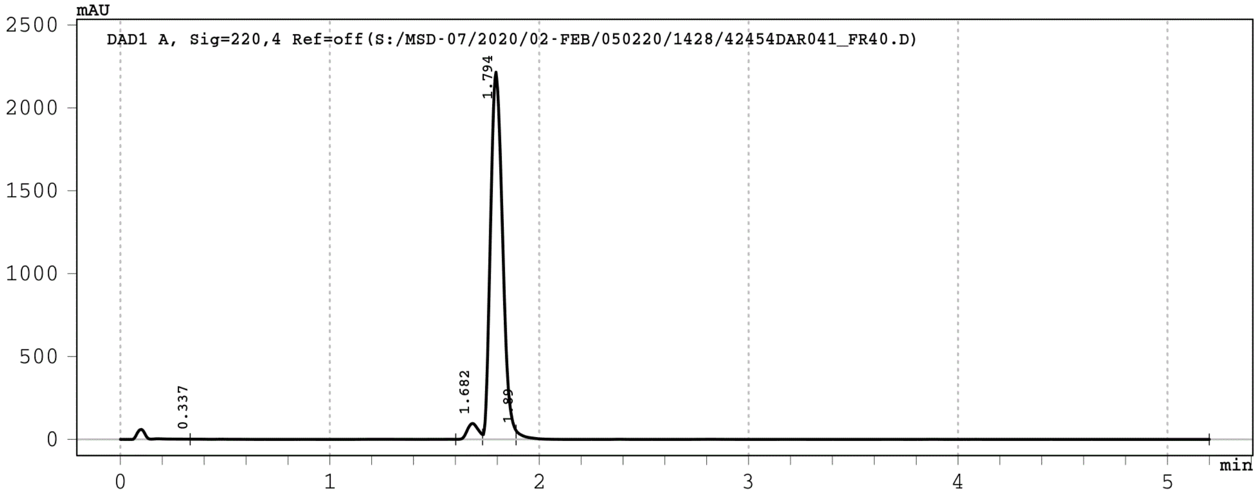


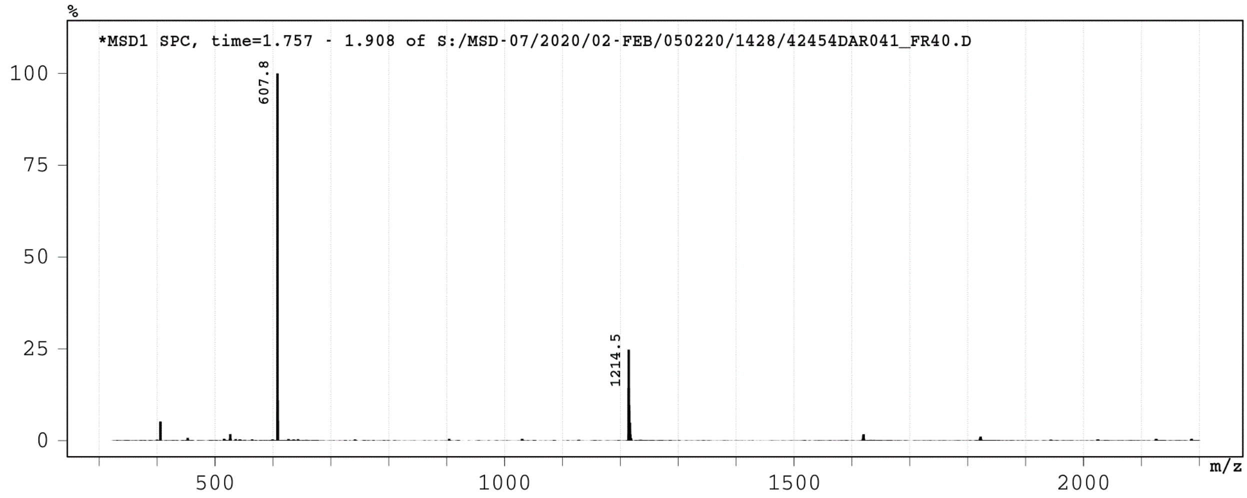


Signal 1: DAD1 A, Sig=220,4 Ref=off

| Peak # | RT [min] | Width [min] | Area | Area % |
|---|---|---|---|---|
| 1 | 0.337 | 1.27 | 27.2 | 0.3 |
| 2 | 1.682 | 0.13 | 349.5 | 3.7 |
| 3 | 1.794 | 0.16 | 8837.7 | 94.5 |
| 4 | 1.890 | 3.31 | 135.1 | 1.4 |

Data File : S:/MSD-07/2020/02-FEB/050220/1428/42454DAR042_FR5->

Injection Date : 06-Feb-20, 01:34:38
Sample Name : 42454DAR042_FR55-57
Remarks : Baseline corrected
Acq. Method : D:/METHODS/5-95-6M.M
column used : Je_559
Analysis Method : C:/CHEM32/1/METHODS/JPT-LCMS.M

Sample Info : 42454_042-Dr. Martine Darwish H-FQLAFGHL-OH

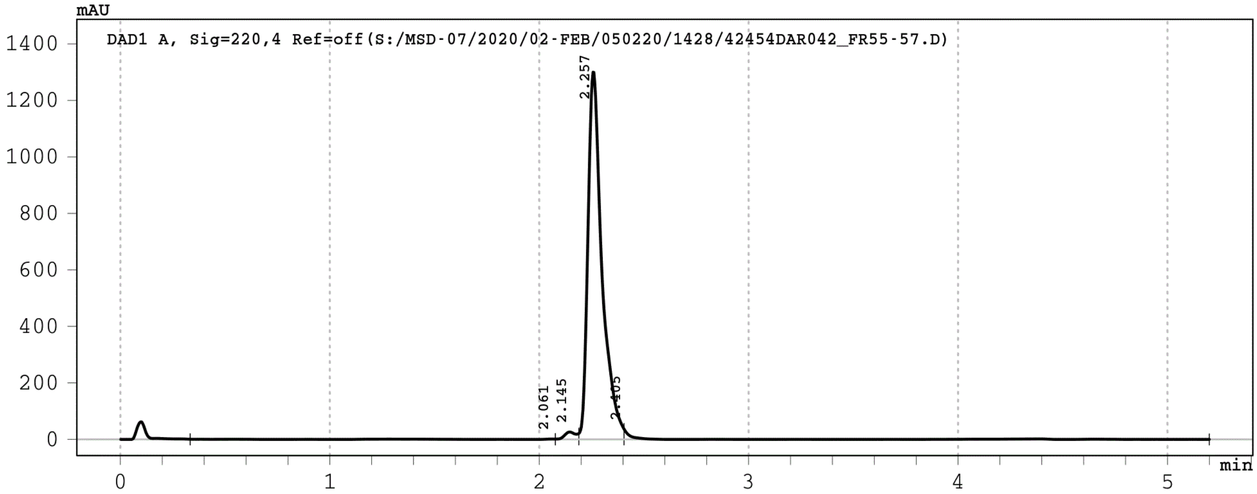


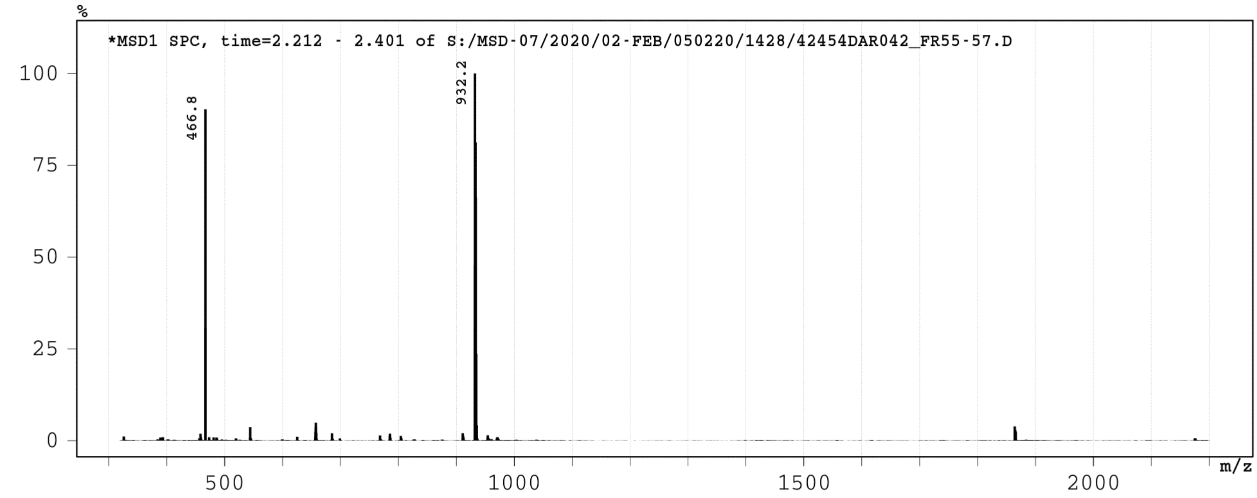


Signal 1: DAD1 A, Sig=220,4 Ref=off

| Peak # | RT [min] | Width [min] | Area | Area % |
|---|---|---|---|---|
| 1 | 2.061 | 1.74 | 34.8 | 0.6 |
| 2 | 2.145 | 0.11 | 100.8 | 1.6 |
| 3 | 2.257 | 0.22 | 6083.3 | 96.1 |
| 4 | 2.405 | 2.79 | 110.8 | 1.8 |

Data File : S:/MSD-07/2020/02-FEB/050220/1428/42454DAR043_FR6->

Injection Date : 06-Feb-20, 01:43:10
Sample Name : 42454DAR043_FR69-70
Remarks : Baseline corrected
Acq. Method : D:/METHODS/5-95-6M.M
column used : Je_559
Analysis Method : C:/CHEM32/1/METHODS/JPT-LCMS.M

Sample Info : 42454_043-Dr. Martine Darwish H-DAYSHHSGF-OH

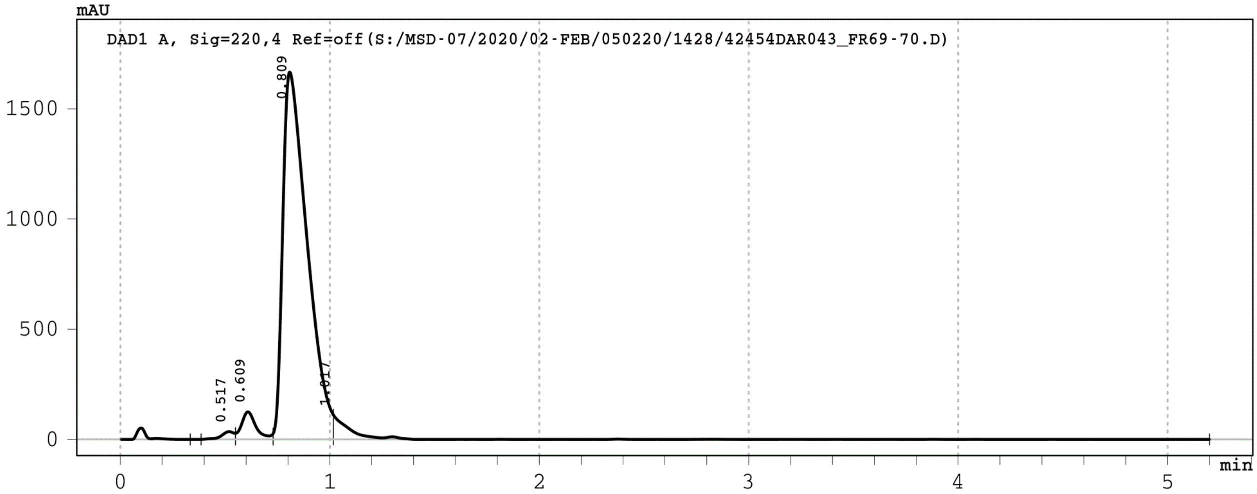


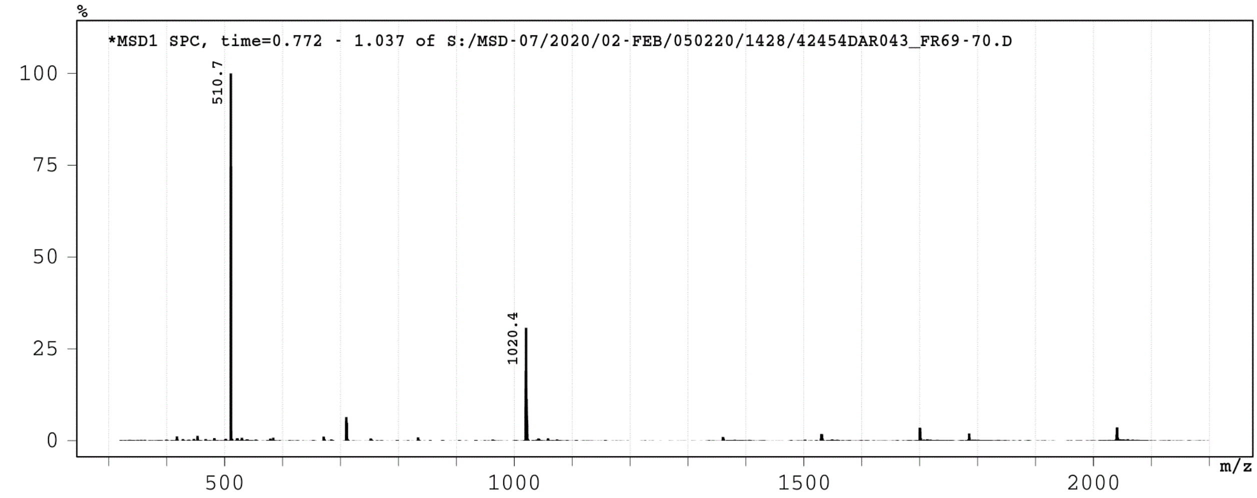


Signal 1: DAD1 A, Sig=220,4 Ref=off

| Peak # | RT [min] | Width [min] | Area | Area % |
|---|---|---|---|---|
| 1 | 0.337 | 0.05 | 0.0 | 0.0 |
| 2 | 0.517 | 0.16 | 146.3 | 1.0 |
| 3 | 0.609 | 0.18 | 612.1 | 4.1 |
| 4 | 0.809 | 0.29 | 13475.4 | 90.8 |
| 5 | 1.017 | 4.18 | 600.3 | 4.0 |

Data File : S:/MSD-07/2020/02-FEB/050220/1428/42454DAR044_FR8->

Injection Date : 06-Feb-20, 01:51:37
Sample Name : 42454DAR044_FR85
Remarks : Baseline corrected
Acq. Method : D:/METHODS/5-95-6M.M
column used : Je_559
Analysis Method : C:/CHEM32/1/METHODS/JPT-LCMS.M

Sample Info : 42454_044-Dr. Martine Darwish H-AYLGYLAML-OH

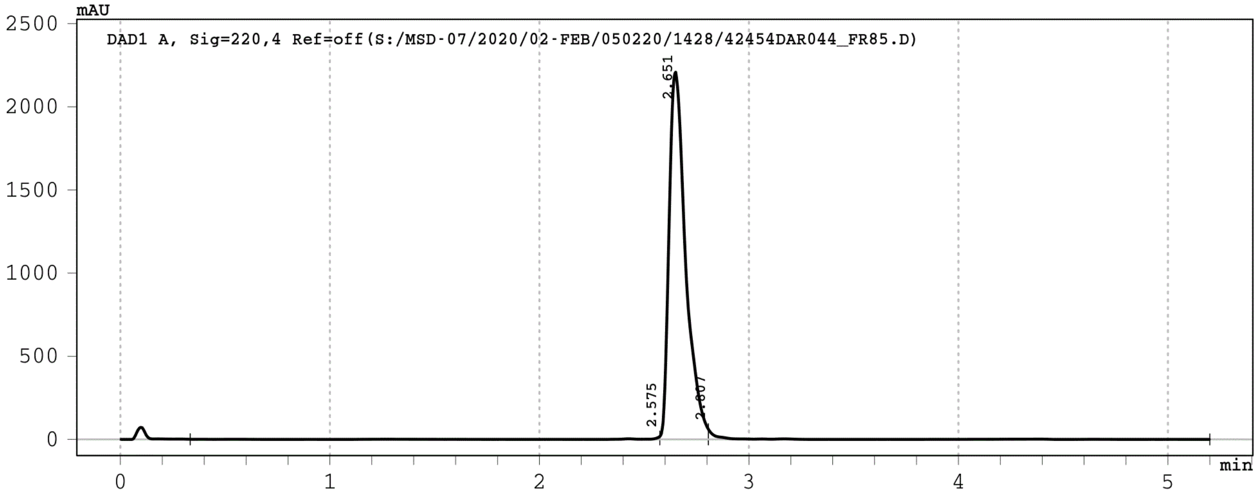


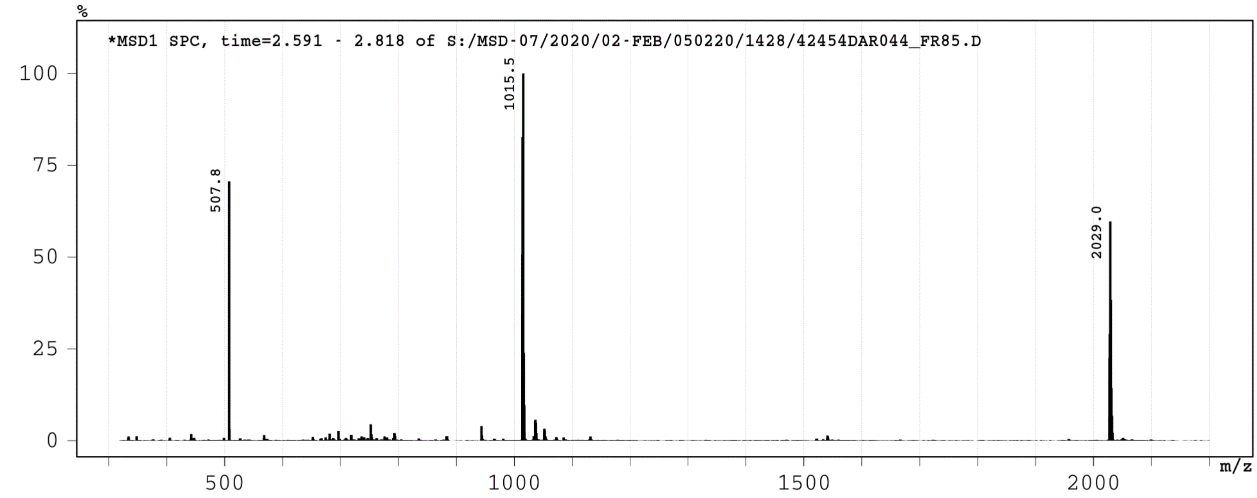


Signal 1: DAD1 A, Sig=220,4 Ref=off

| Peak # | RT [min] | Width [min] | Area | Area % |
|---|---|---|---|---|
| 1 | 2.575 | 2.24 | 71.7 | 0.6 |
| 2 | 2.651 | 0.23 | 12183.8 | 97.7 |
| 3 | 2.807 | 2.39 | 210.7 | 1.7 |

Data File : S:/MSD-07/2020/02-FEB/050220/1428/42454DAR045_FR9->

Injection Date : 06-Feb-20, 02:00:03
Sample Name : 42454DAR045_FR90
Remarks : Baseline corrected
Acq. Method : D:/METHODS/5-95-6M.M
column used : Je_559
Analysis Method : C:/CHEM32/1/METHODS/JPT-LCMS.M

Sample Info : 42454_045-Dr. Martine Darwish H-LSFHLDNIL-OH

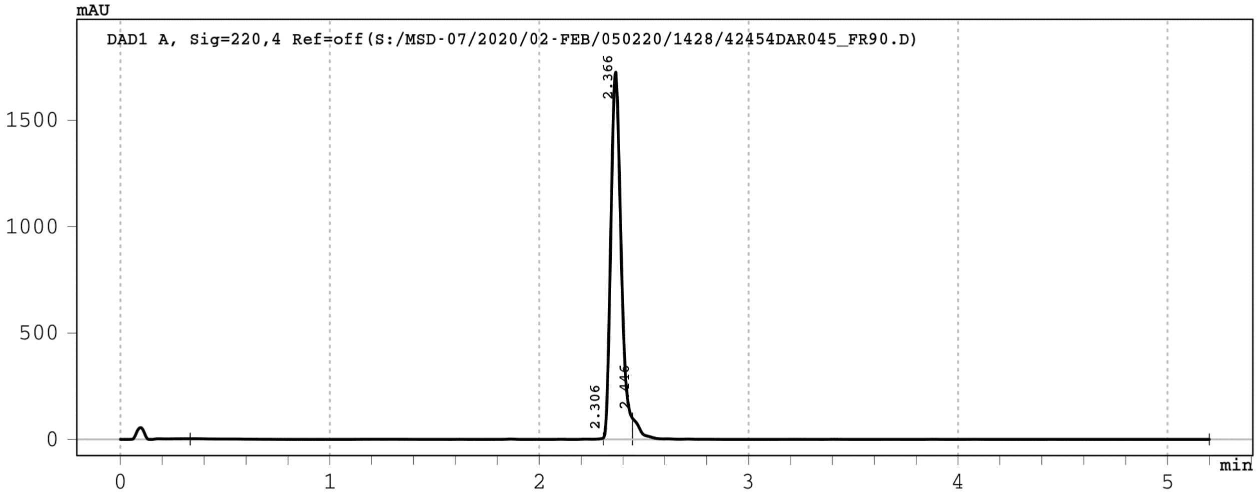


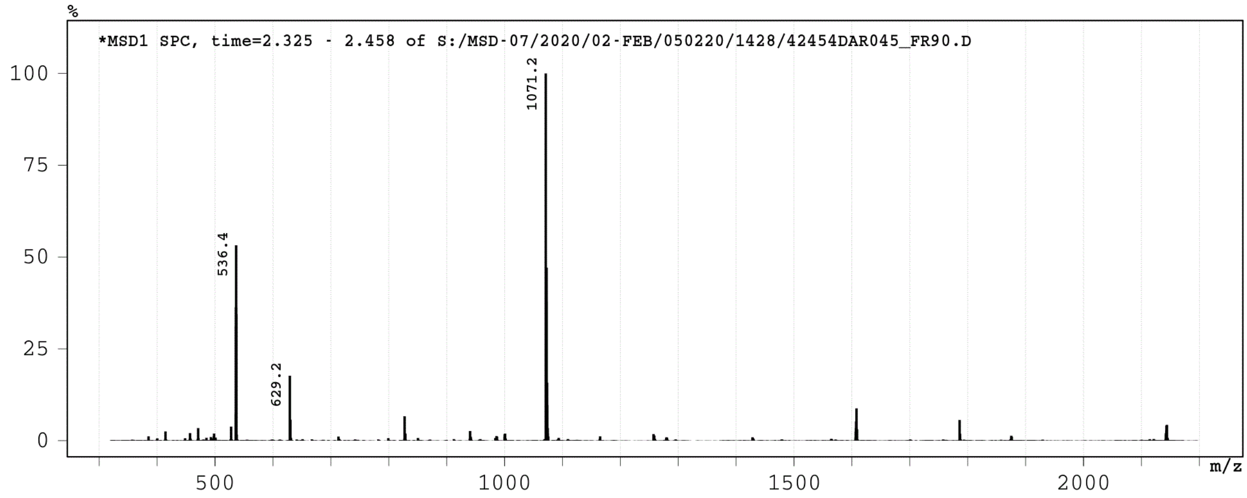


Signal 1: DAD1 A, Sig=220,4 Ref=off

| Peak # | RT [min] | Width [min] | Area | Area % |
|---|---|---|---|---|
| 1 | 2.306 | 1.97 | 71.4 | 1.1 |
| 2 | 2.366 | 0.14 | 5870.8 | 94.4 |
| 3 | 2.446 | 2.75 | 274.2 | 4.4 |

Data File : S:/MSD-07/2020/02-FEB/040220/1427/42454DAR046_FR7->

Injection Date : 04-Feb-20, 17:30:58
Sample Name : 42454DAR046_FR73-75
Remarks : Baseline corrected
Acq. Method : D:/METHODS/5-95-6M.M
column used : Je_559
Analysis Method : C:/CHEM32/1/METHODS/JPT-LCMS.M

Sample Info : 42454_046-Dr. Martine Darwish H-ASFHAAFL-OH

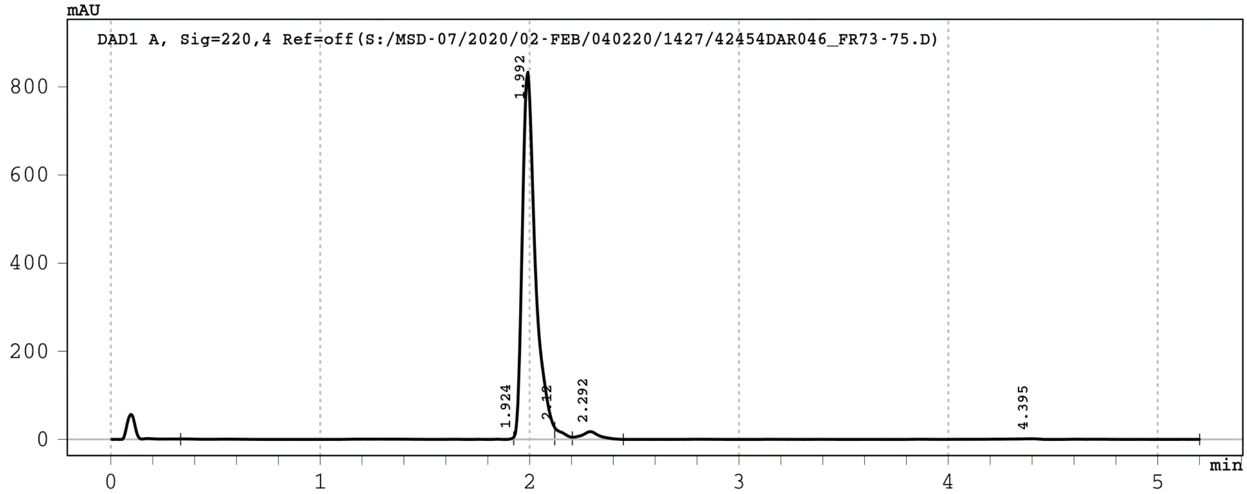


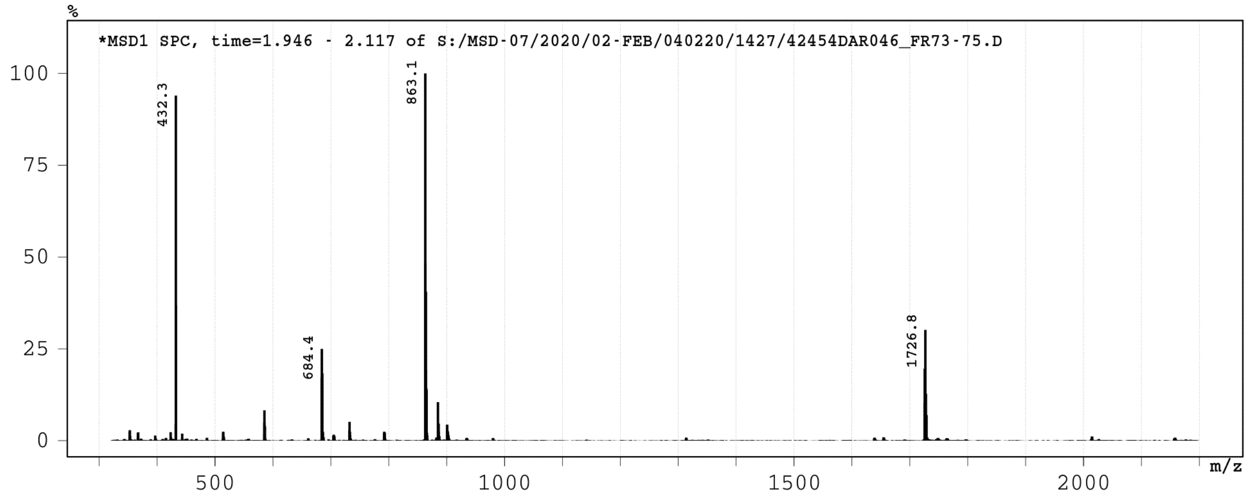


Signal 1: DAD1 A, Sig=220,4 Ref=off

| Peak # | RT [min] | Width [min] | Area | Area % |
|---|---|---|---|---|
| 1 | 1.924 | 1.59 | 26.5 | 0.7 |
| 2 | 1.992 | 0.20 | 3645.5 | 94.1 |
| 3 | 2.120 | 0.08 | 68.9 | 1.8 |
| 4 | 2.292 | 0.24 | 105.6 | 2.7 |
| 5 | 4.395 | 2.75 | 28.3 | 0.7 |

Data File : S:/MSD-07/2020/02-FEB/050220/1428/42454DAR047_FR9->

Injection Date : 05-Feb-20, 15:29:11
Sample Name : 42454DAR047_FR9-10
Remarks : Baseline corrected
Acq. Method : D:/METHODS/5-95-6M.M
column used : Je_559
Analysis Method : C:/CHEM32/1/METHODS/JPT-LCMS.M

Sample Info : 42454_047-Dr. Martine Darwish H-LGVCMYGML-OH

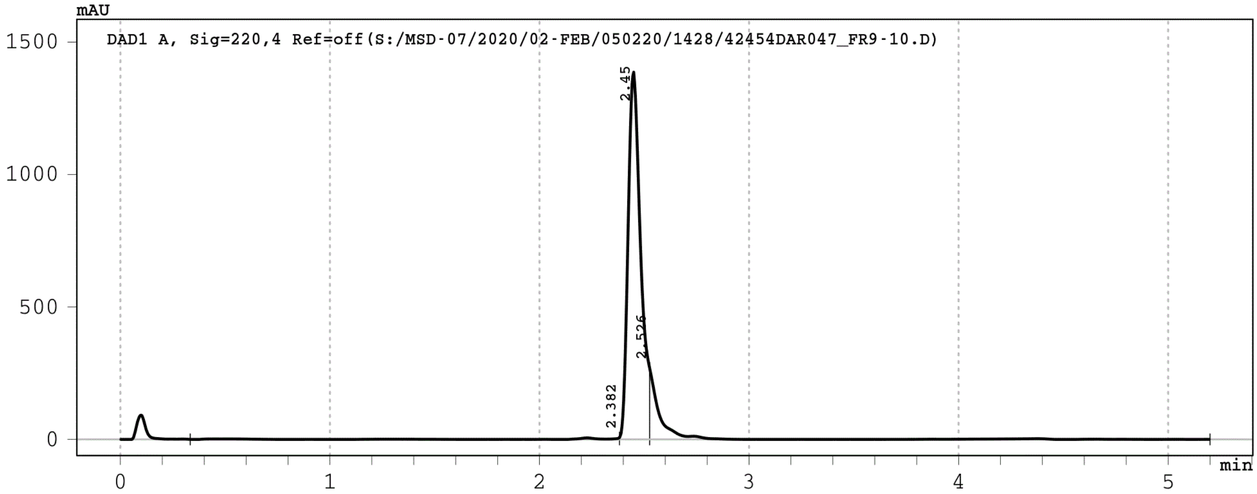


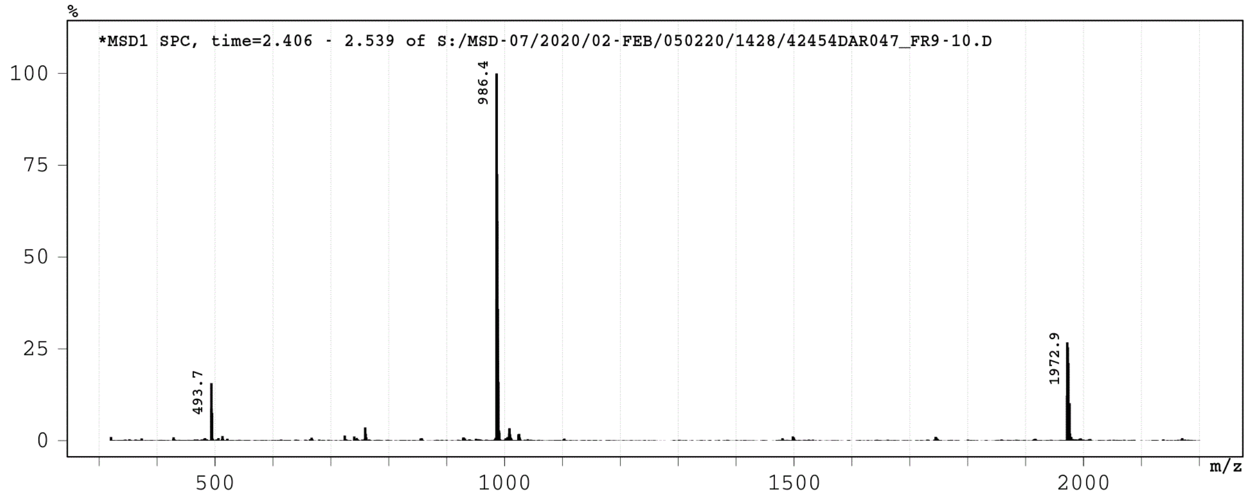


Signal 1: DAD1 A, Sig=220,4 Ref=off

| Peak # | RT [min] | Width [min] | Area | Area % |
|---|---|---|---|---|
| 1 | 2.382 | 2.05 | 81.0 | 1.2 |
| 2 | 2.450 | 0.14 | 5762.4 | 85.4 |
| 3 | 2.526 | 2.67 | 902.3 | 13.4 |

Data File : S:/MSD-07/2020/02-FEB/050220/1428/42454DAR048_FR1->

Injection Date : 05-Feb-20, 15:37:35
Sample Name : 42454DAR048_FR16-17
Remarks : Baseline corrected
Acq. Method : D:/METHODS/5-95-6M.M
column used : Je_559
Analysis Method : C:/CHEM32/1/METHODS/JPT-LCMS.M

Sample Info : 42454_048-Dr. Martine Darwish H-TAYHARIL-OH

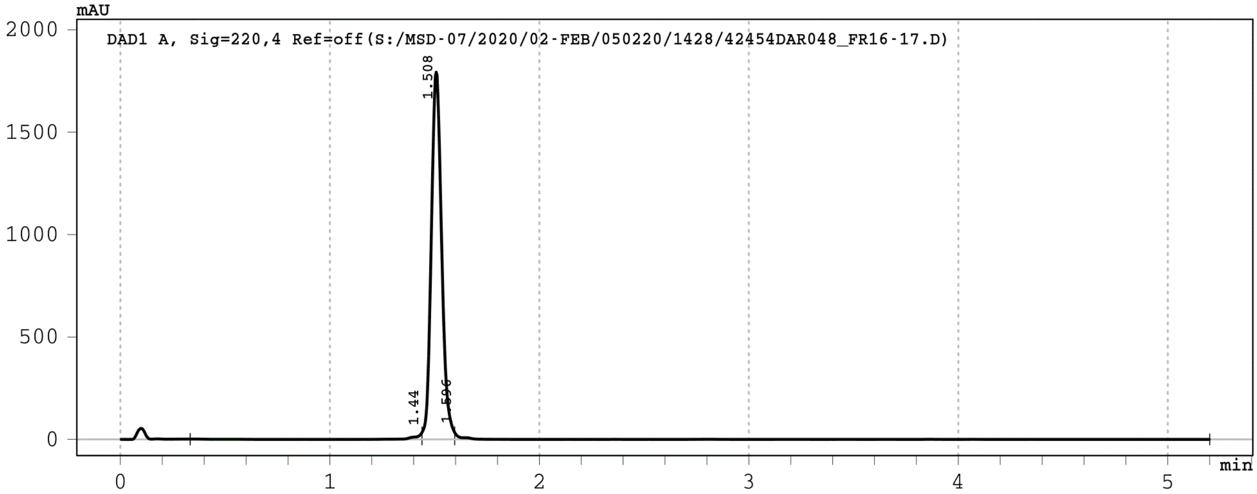


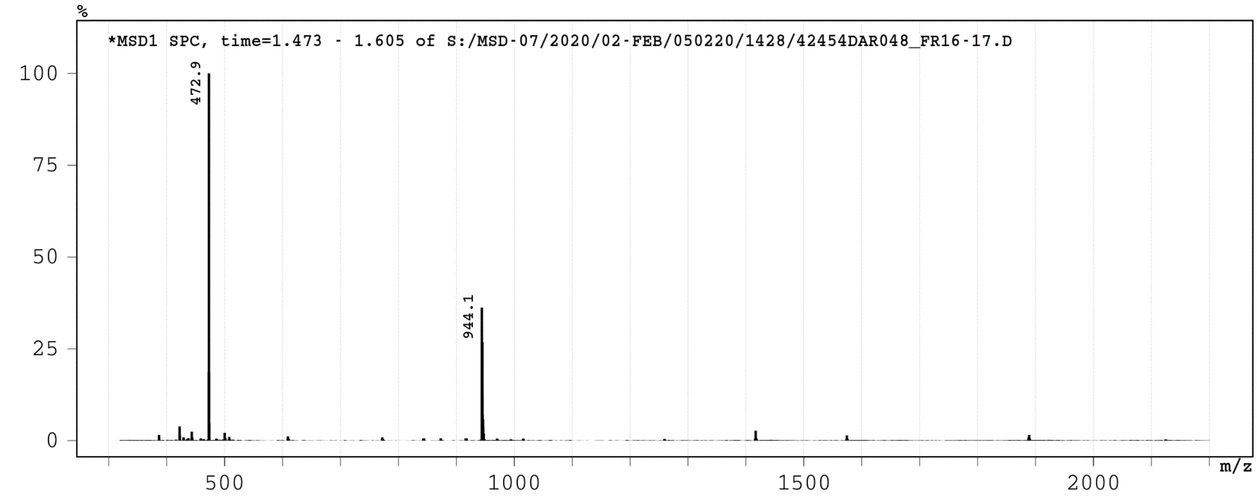


Signal 1: DAD1 A, Sig=220,4 Ref=off

| Peak # | RT [min] | Width [min] | Area | Area % |
|---|---|---|---|---|
| 1 | 1.440 | 1.11 | 82.0 | 1.3 |
| 2 | 1.508 | 0.16 | 6172.1 | 97.5 |
| 3 | 1.596 | 3.60 | 73.7 | 1.2 |

Data File : S:/MSD-07/2020/02-FEB/050220/1428/42454DAR049_FR3->

Injection Date : 05-Feb-20, 15:45:59
Sample Name : 42454DAR049_FR32-33
Remarks : Baseline corrected
Acq. Method : D:/METHODS/5-95-6M.M
column used : Je_559
Analysis Method : C:/CHEM32/1/METHODS/JPT-LCMS.M

Sample Info : 42454_049-Dr. Martine Darwish H-LAPYFLSEPL-OH

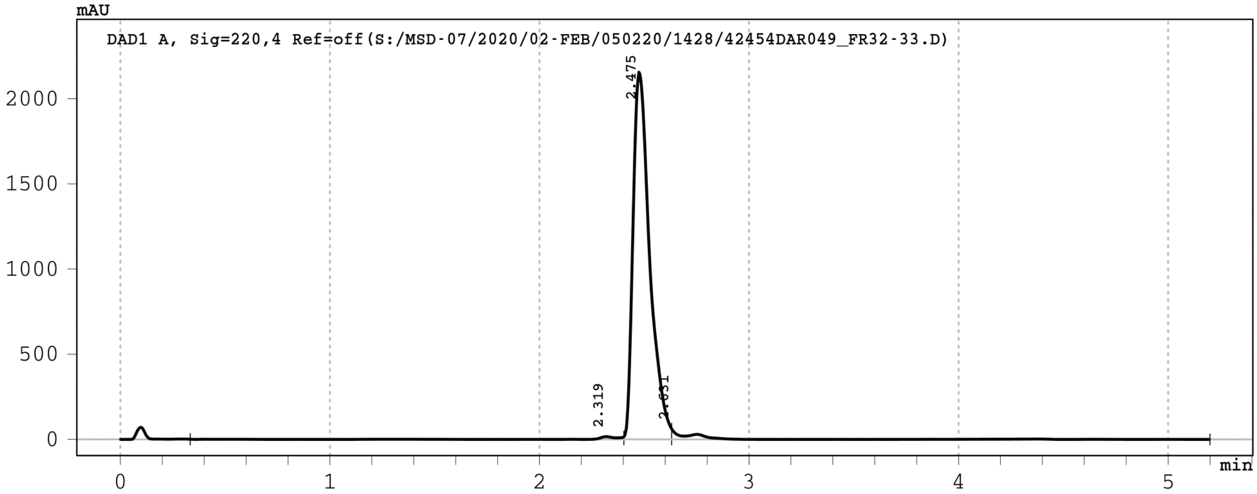


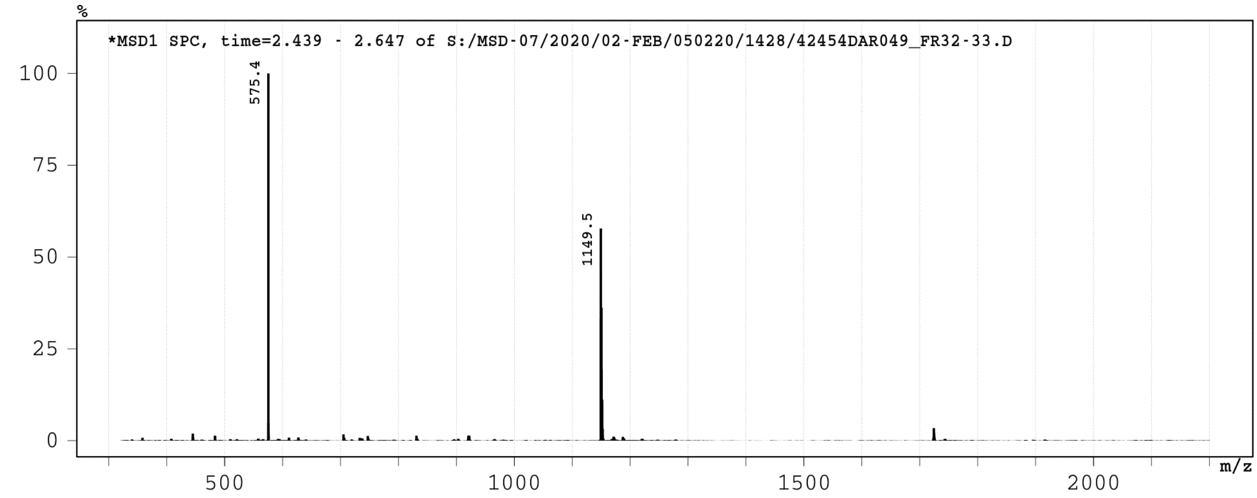


Signal 1: DAD1 A, Sig=220,4 Ref=off

| Peak # | RT [min] | Width [min] | Area | Area % |
|---|---|---|---|---|
| 1 | 2.319 | 2.07 | 125.4 | 1.0 |
| 2 | 2.475 | 0.23 | 11946.7 | 96.1 |
| 3 | 2.631 | 2.57 | 365.0 | 2.9 |

Data File : S:/MSD-07/2020/02-FEB/050220/1428/42454DAR050_FR4->

Injection Date : 05-Feb-20, 15:54:23
Sample Name : 42454DAR050_FR48-49
Remarks : Baseline corrected
Acq. Method : D:/METHODS/5-95-6M.M
column used : Je_559
Analysis Method : C:/CHEM32/1/METHODS/JPT-LCMS.M

Sample Info : 42454_050-Dr. Martine Darwish H-KIHQNLEEM-OH

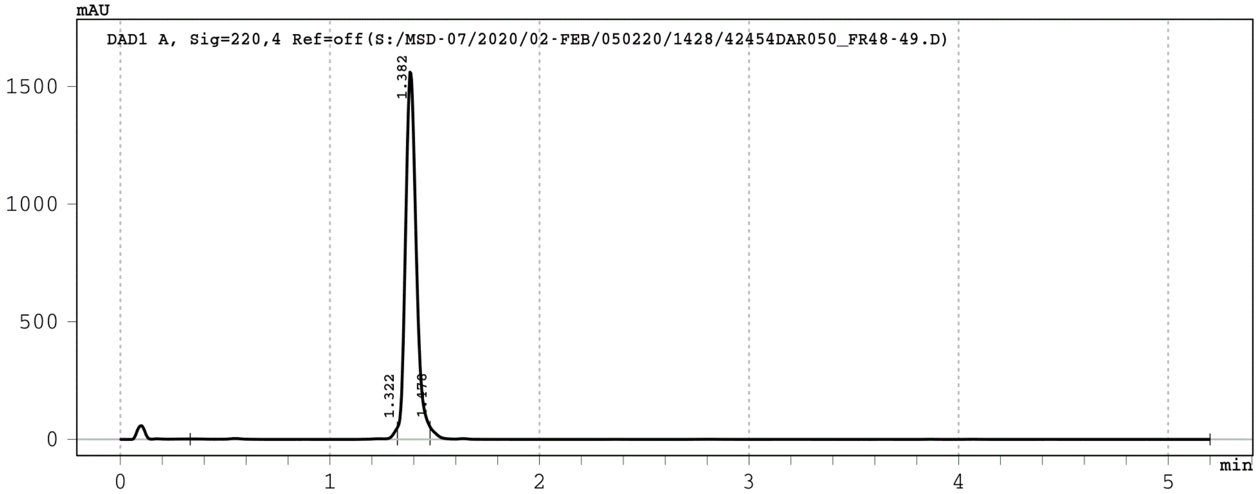


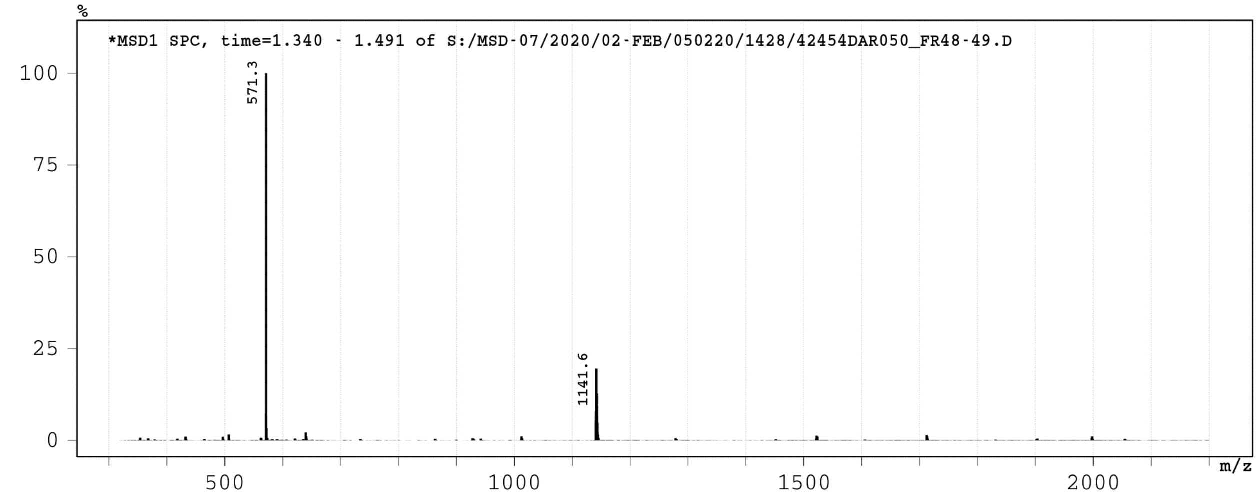


Signal 1: DAD1 A, Sig=220,4 Ref=off

| Peak # | RT [min] | Width [min] | Area | Area % |
|---|---|---|---|---|
| 1 | 1.322 | 0.99 | 102.8 | 1.8 |
| 2 | 1.382 | 0.16 | 5570.2 | 96.1 |
| 3 | 1.478 | 3.72 | 123.6 | 2.1 |

Data File : S:/MSD-07/2020/02-FEB/050220/1428/42454DAR051_FR5->

jpt Innovative Peptide Solutions

Injection Date : 05-Feb-20, 16:02:47
Sample Name : 42454DAR051_FR56-57
Remarks : Baseline corrected
Acq. Method : D:/METHODS/5-95-6M.M
column used : Je_559
Analysis Method : C:/CHEM32/1/METHODS/JPT-LCMS.M

Sample Info : 42454_051-Dr. Martine Darwish H-CSYLPELPL-OH

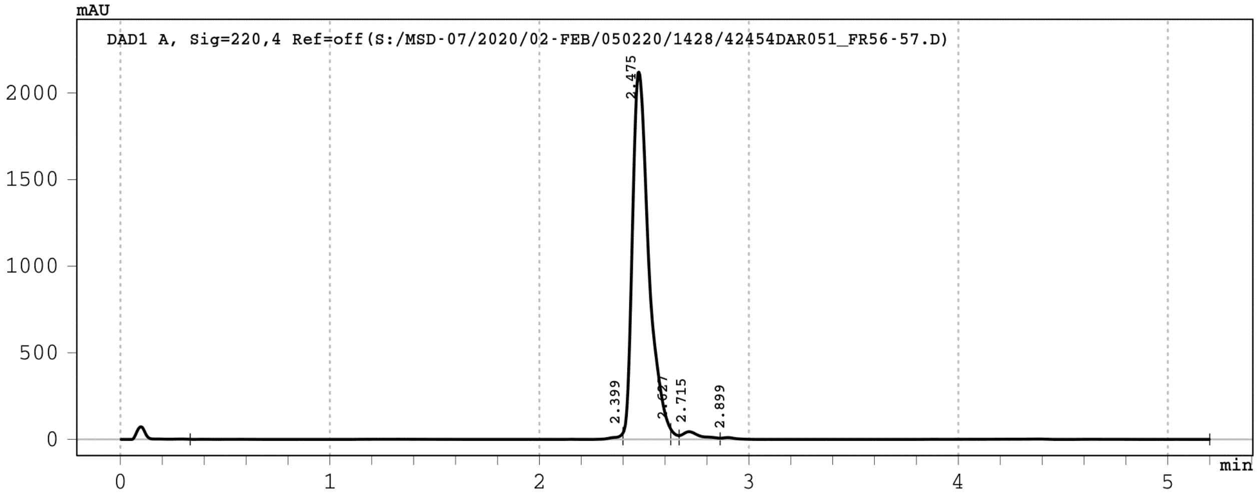


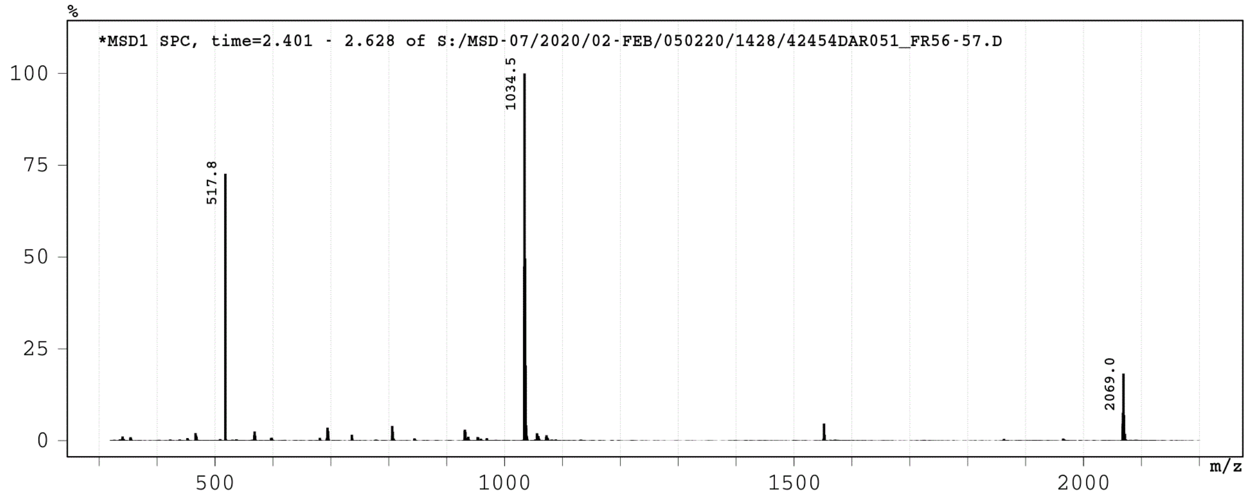


Signal 1: DAD1 A, Sig=220,4 Ref=off

| Peak # | RT [min] | Width [min] | Area | Area % |
|---|---|---|---|---|
| 1 | 2.399 | 2.07 | 98.0 | 0.8 |
| 2 | 2.475 | 0.23 | 11530.1 | 95.5 |
| 3 | 2.627 | 0.04 | 82.2 | 0.7 |
| 4 | 2.715 | 0.20 | 272.8 | 2.3 |
| 5 | 2.899 | 2.34 | 92.0 | 0.8 |

Data File : S:/MSD-07/2020/02-FEB/050220/1428/42454DAR052_FR6->

Injection Date : 05-Feb-20, 16:11:11
Sample Name : 42454DAR052_FR64-66
Remarks : Baseline corrected
Acq. Method : D:/METHODS/5-95-6M.M
column used : Je_559
Analysis Method : C:/CHEM32/1/METHODS/JPT-LCMS.M

Sample Info : 42454_052-Dr. Martine Darwish H-VSPVNDLDV-OH

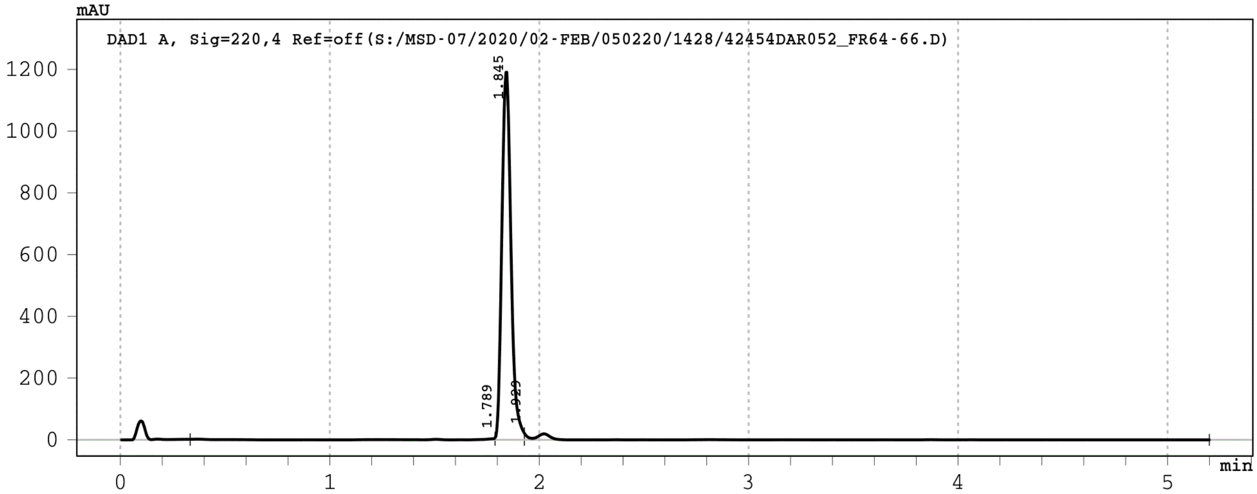


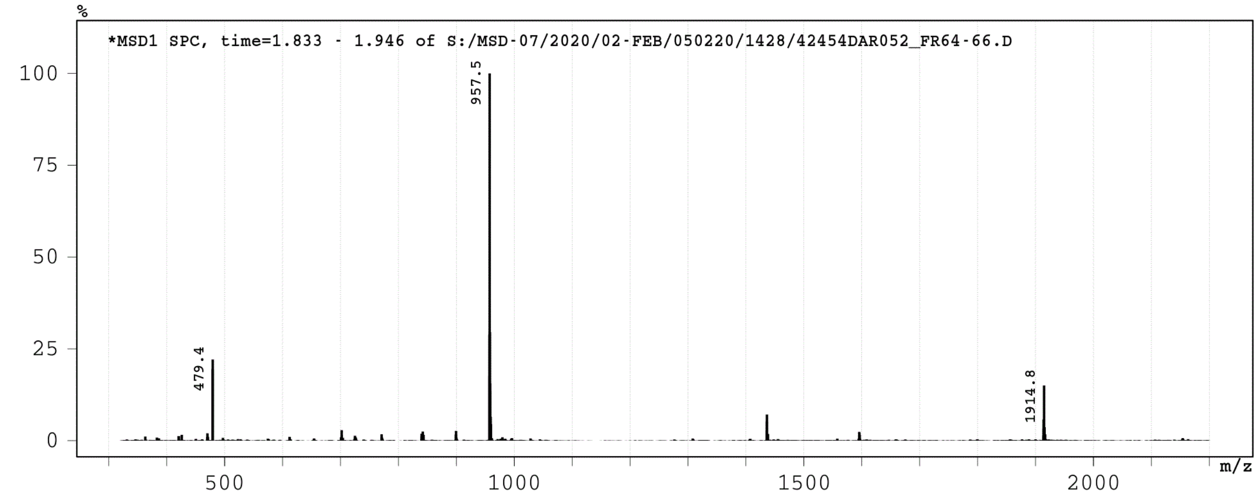


Signal 1: DAD1 A, Sig=220,4 Ref=off

| Peak # | RT [min] | Width [min] | Area | Area % |
|---|---|---|---|---|
| 1 | 1.789 | 1.46 | 48.3 | 1.3 |
| 2 | 1.845 | 0.14 | 3612.3 | 95.7 |
| 3 | 1.929 | 3.27 | 112.9 | 3.0 |

Data File : S:/MSD-07/2020/02-FEB/050220/1428/42454DAR053_FR7->

Injection Date : 05-Feb-20, 16:19:35
Sample Name : 42454DAR053_FR70-71
Remarks : Baseline corrected
Acq. Method : D:/METHODS/5-95-6M.M
column used : Je_559
Analysis Method : C:/CHEM32/1/METHODS/JPT-LCMS.M

Sample Info : 42454_053-Dr. Martine Darwish H-PMTKNTQYI-OH

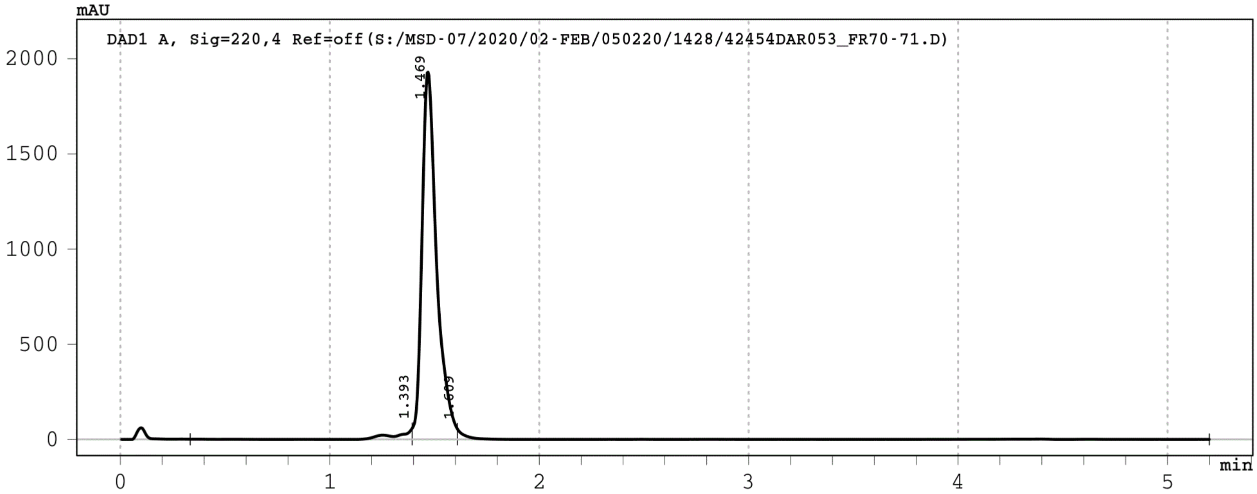


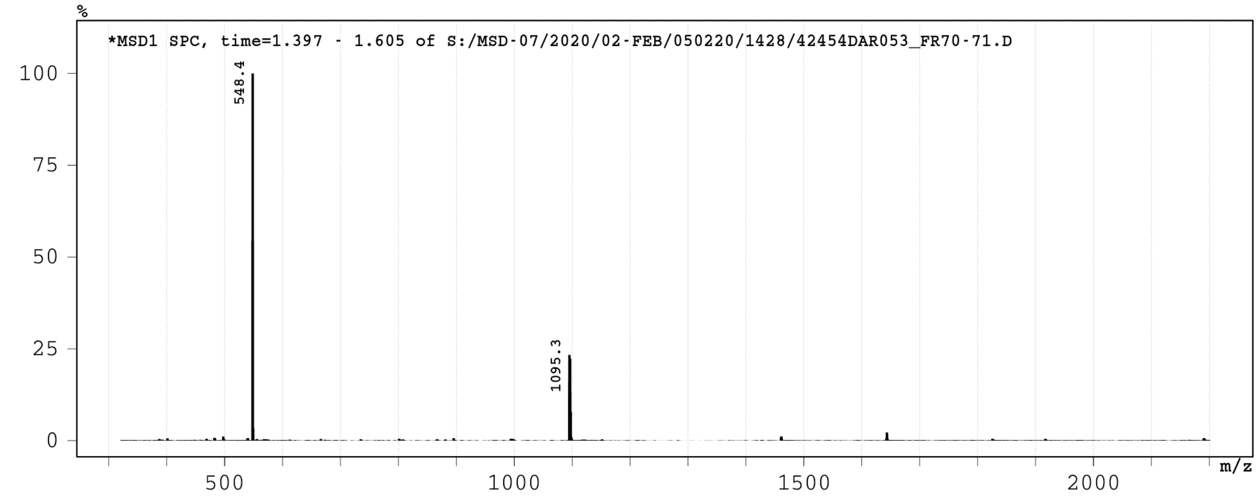


Signal 1: DAD1 A, Sig=220,4 Ref=off

| Peak # | RT [min] | Width [min] | Area | Area % |
|---|---|---|---|---|
| 1 | 1.393 | 1.06 | 285.6 | 2.9 |
| 2 | 1.469 | 0.22 | 9430.6 | 95.4 |
| 3 | 1.609 | 3.59 | 168.2 | 1.7 |

Data File : S:/MSD-07/2020/02-FEB/050220/1428/42454DAR054_FR2->

Injection Date : 05-Feb-20, 16:27:59
Sample Name : 42454DAR054_FR2-3
Remarks : Baseline corrected
Acq. Method : D:/METHODS/5-95-6M.M
column used : Je_559
Analysis Method : C:/CHEM32/1/METHODS/JPT-LCMS.M

Sample Info : 42454_054-Dr. Martine Darwish H-LALTDLQYI-OH

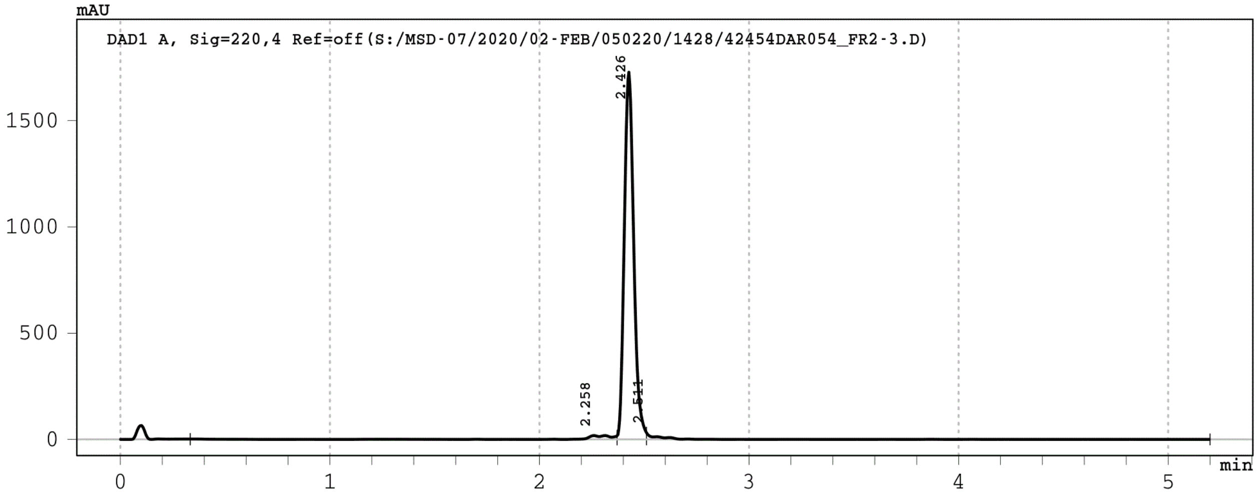


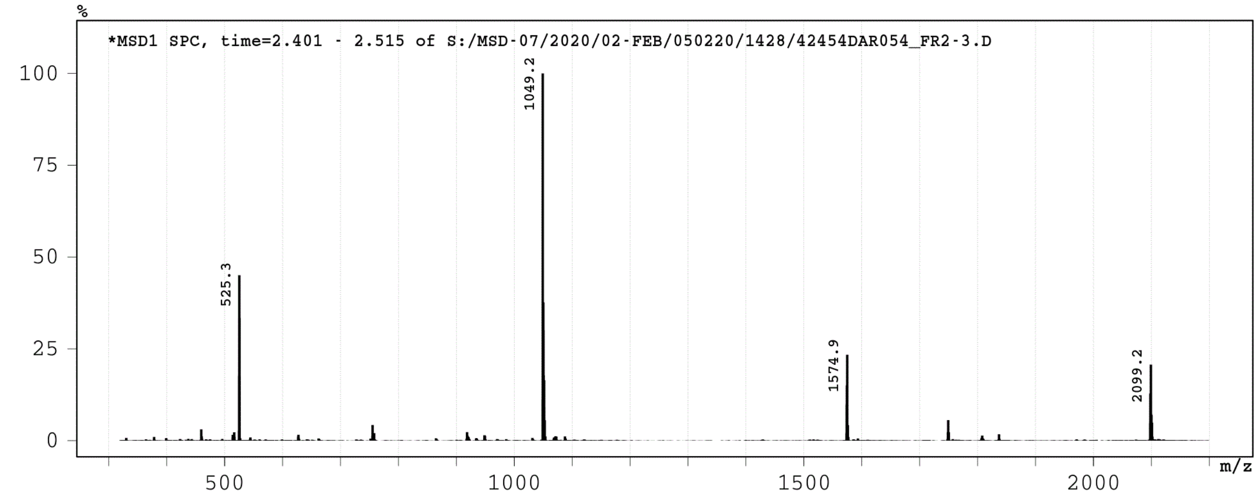


Signal 1: DAD1 A, Sig=220,4 Ref=off

| Peak # | RT [min] | Width [min] | Area | Area % |
|---|---|---|---|---|
| 1 | 2.258 | 2.04 | 164.2 | 2.9 |
| 2 | 2.426 | 0.14 | 5424.0 | 95.2 |
| 3 | 2.511 | 2.69 | 109.3 | 1.9 |

| | |
|---|---|
| Data File | : S:/MSD-07/2020/02-FEB/070220/1430/42454DAR055_FR3-> |
| Injection Date | : 07-Feb-20, 13:29:38 |
| Sample Name | : 42454DAR055_FR30-32 |
| Remarks | : Baseline corrected |
| Acq. Method | : D:/METHODS/5-95-6M.M |
| column used | : Je_559 |
| Analysis Method | : C:/CHEM32/1/METHODS/JPT-LCMS.M |
| Sample Info | : 42454_055-Dr. Martine Darwish H-QLYLLCCQL-OH |

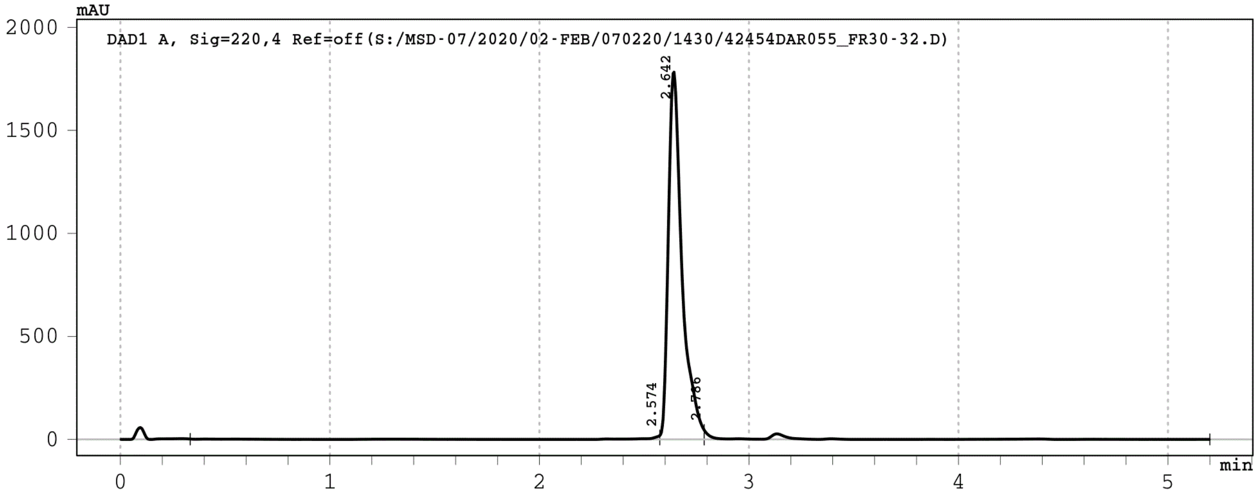


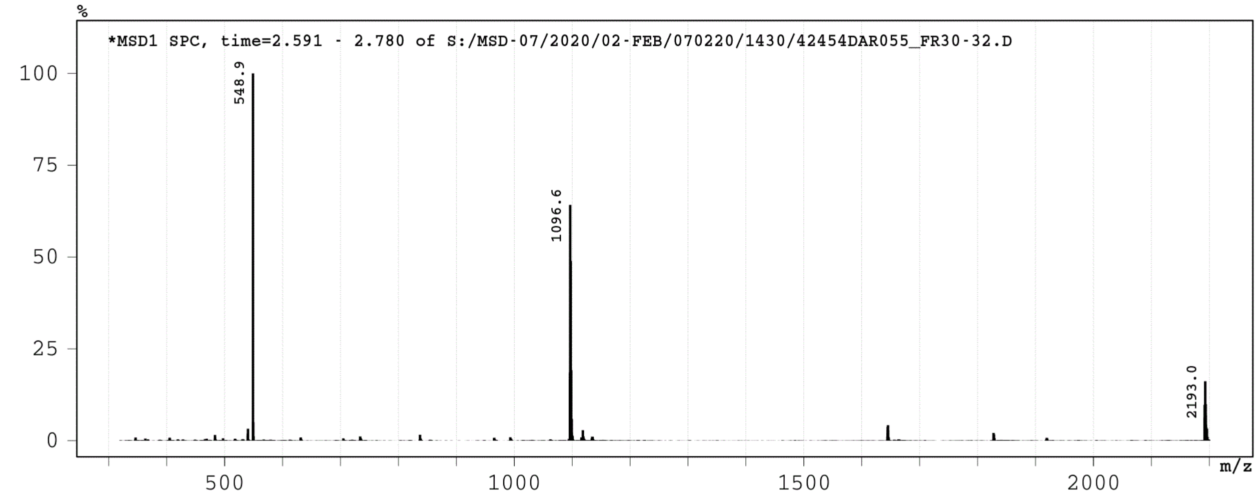


Signal 1: DAD1 A, Sig=220,4 Ref=off

| Peak # | RT [min] | Width [min] | Area | Area % |
|---|---|---|---|---|
| 1 | 2.574 | 2.24 | 111.5 | 1.3 |
| 2 | 2.642 | 0.21 | 8114.5 | 95.4 |
| 3 | 2.786 | 2.41 | 277.0 | 3.3 |

Data File : S:/MSD-07/2020/02-FEB/050220/1428/42454DAR056_FR1->

Injection Date : 05-Feb-20, 16:36:23
Sample Name : 42454DAR056_FR15-16
Remarks : Baseline corrected
Acq. Method : D:/METHODS/5-95-6M.M
column used : Je_559
Analysis Method : C:/CHEM32/1/METHODS/JPT-LCMS.M

Sample Info : 42454_056-Dr. Martine Darwish H-MSYFLQGTL-OH

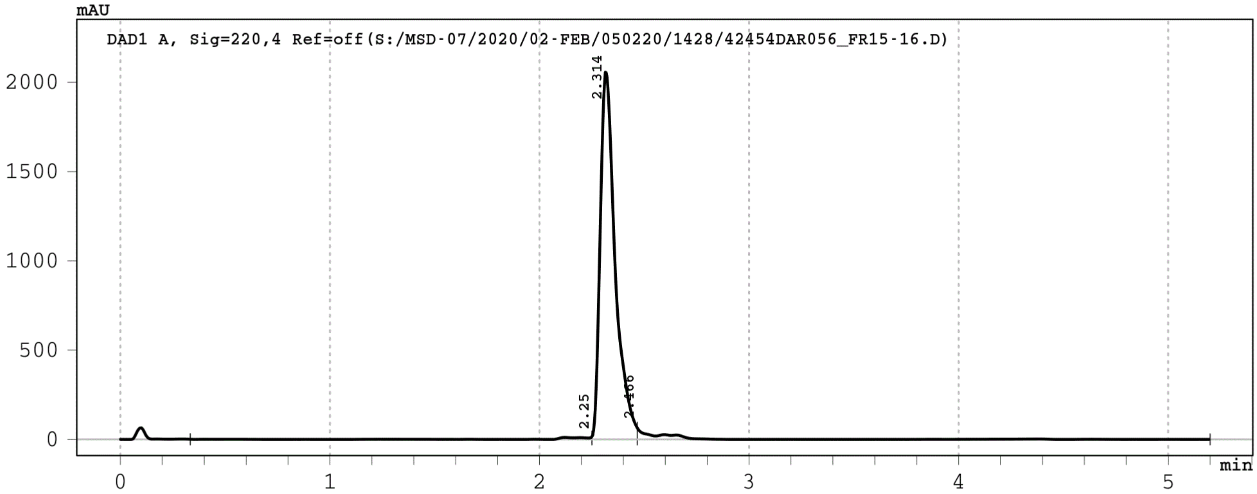


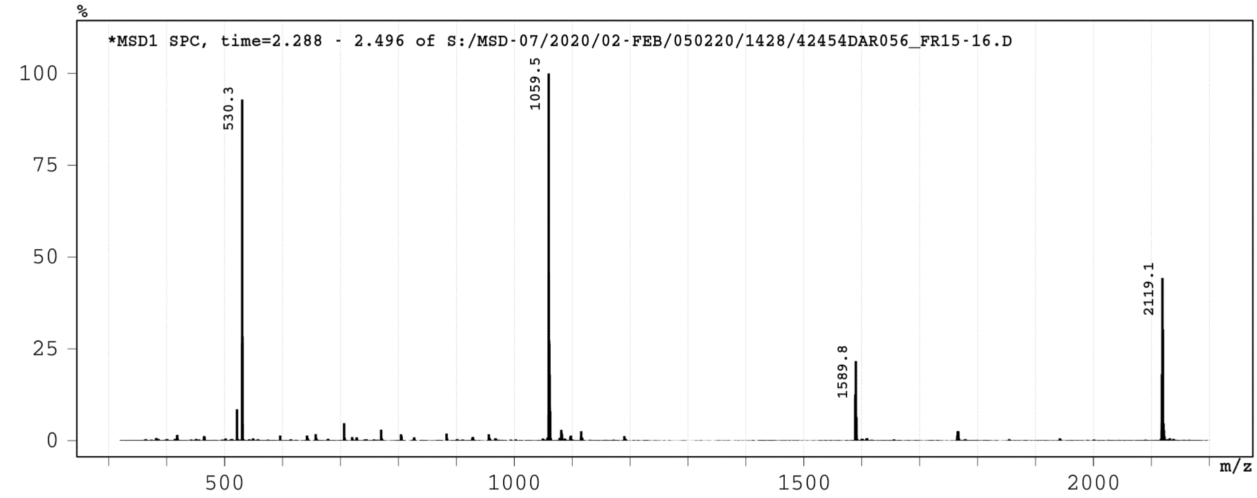


Signal 1: DAD1 A, Sig=220,4 Ref=off

| Peak # | RT [min] | Width [min] | Area | Area % |
|---|---|---|---|---|
| 1 | 2.250 | 1.92 | 124.6 | 1.2 |
| 2 | 2.314 | 0.22 | 10264.9 | 94.8 |
| 3 | 2.466 | 2.73 | 440.9 | 4.1 |

Data File : S:/MSD-07/2020/02-FEB/050220/1428/42454DAR057_FR1->

Injection Date : 05-Feb-20, 16:44:46
Sample Name : 42454DAR057_FR12-14
Remarks : Baseline corrected
Acq. Method : D:/METHODS/5-95-6M.M
column used : Je_559
Analysis Method : C:/CHEM32/1/METHODS/JPT-LCMS.M

Sample Info : 42454_057-Dr. Martine Darwish H-IMLQYHAL-OH

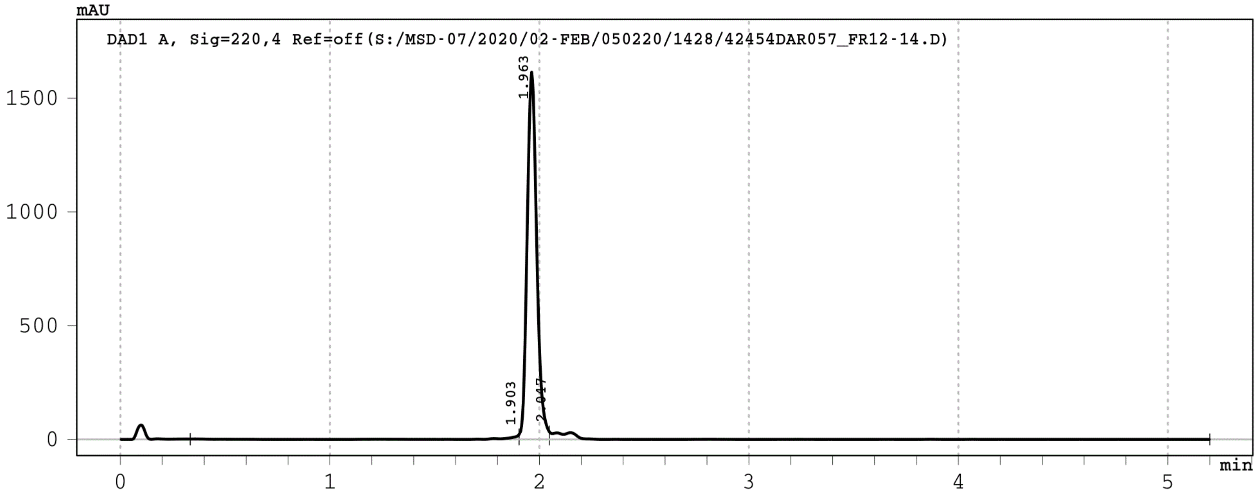


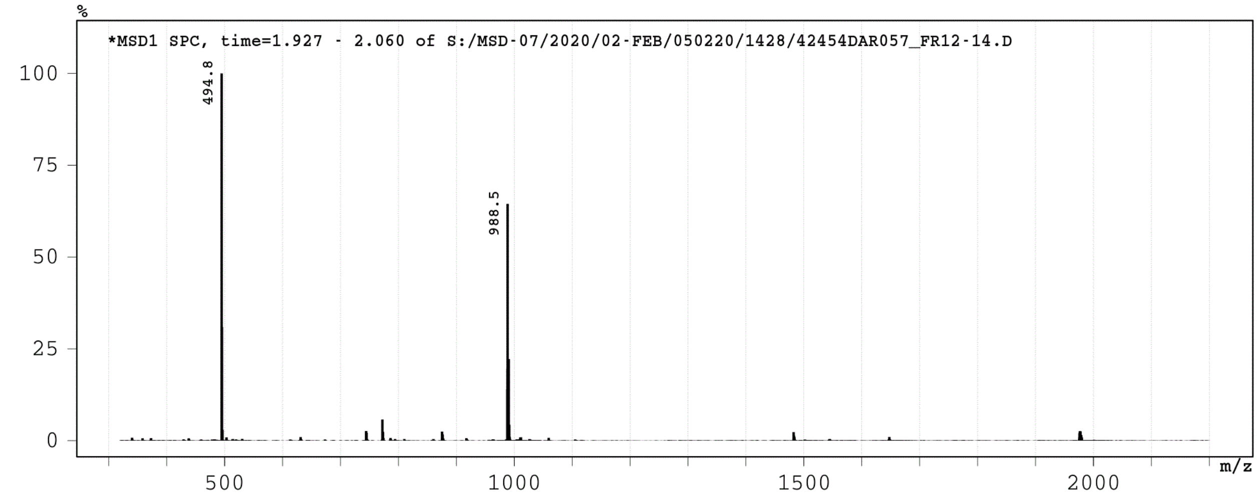


Signal 1: DAD1 A, Sig=220,4 Ref=off

| Peak # | RT [min] | Width [min] | Area | Area % |
|---|---|---|---|---|
| 1 | 1.903 | 1.57 | 81.5 | 1.5 |
| 2 | 1.963 | 0.14 | 5140.0 | 94.2 |
| 3 | 2.047 | 3.15 | 236.5 | 4.3 |

Data File : S:/MSD-07/2020/02-FEB/110220/1433/42454DAR058_FR3->

Injection Date : 11-Feb-20, 23:44:51
Sample Name : 42454DAR058_FR3-5
Remarks : Baseline corrected
Acq. Method : D:/METHODS/5-95-6M.M
column used : Je_559
Analysis Method : C:/CHEM32/1/METHODS/JPT-LCMS.M

Sample Info : 42454_058-Dr. Martine Darwish H-VIHRYREV-OH

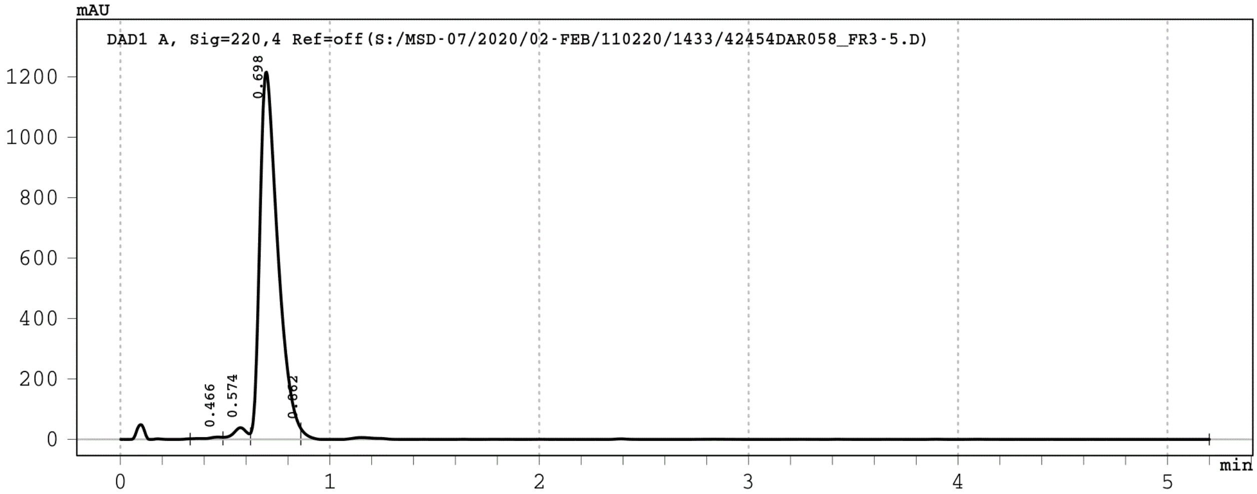


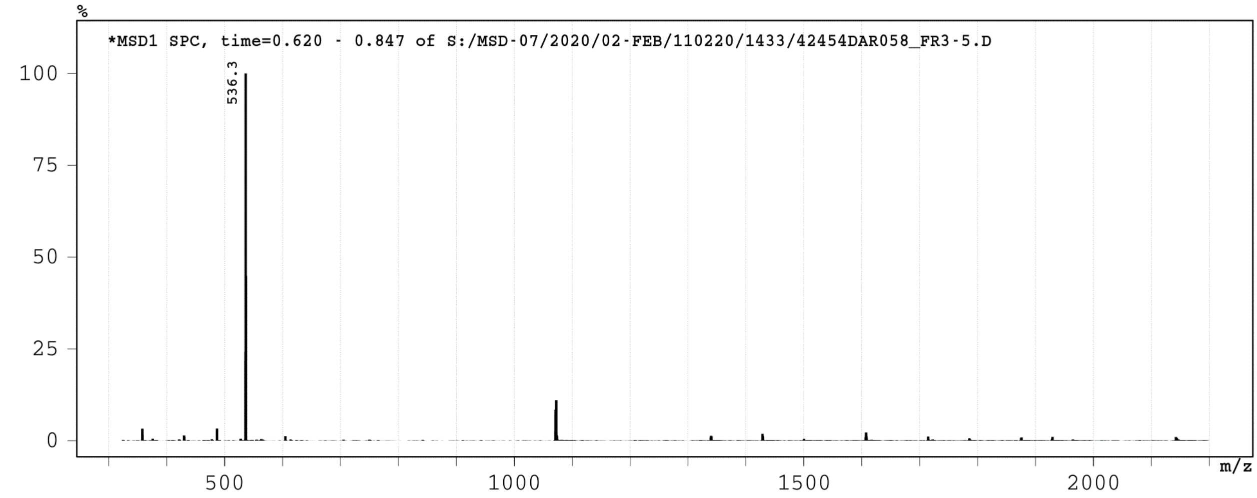


Signal 1: DAD1 A, Sig=220,4 Ref=off

| Peak # | RT [min] | Width [min] | Area | Area % |
|---|---|---|---|---|
| 1 | 0.466 | 0.16 | 42.2 | 0.6 |
| 2 | 0.574 | 0.13 | 176.7 | 2.3 |
| 3 | 0.698 | 0.24 | 7262.8 | 95.4 |
| 4 | 0.862 | 4.34 | 130.0 | 1.7 |

Data File : S:/MSD-07/2020/02-FEB/050220/1428/42454DAR059_FR3->

Injection Date : 05-Feb-20, 16:53:09
Sample Name : 42454DAR059_FR30-31
Remarks : Baseline corrected
Acq. Method : D:/METHODS/5-95-6M.M
column used : Je_559
Analysis Method : C:/CHEM32/1/METHODS/JPT-LCMS.M

Sample Info : 42454_059-Dr. Martine Darwish H-SSPYSLHYL-OH

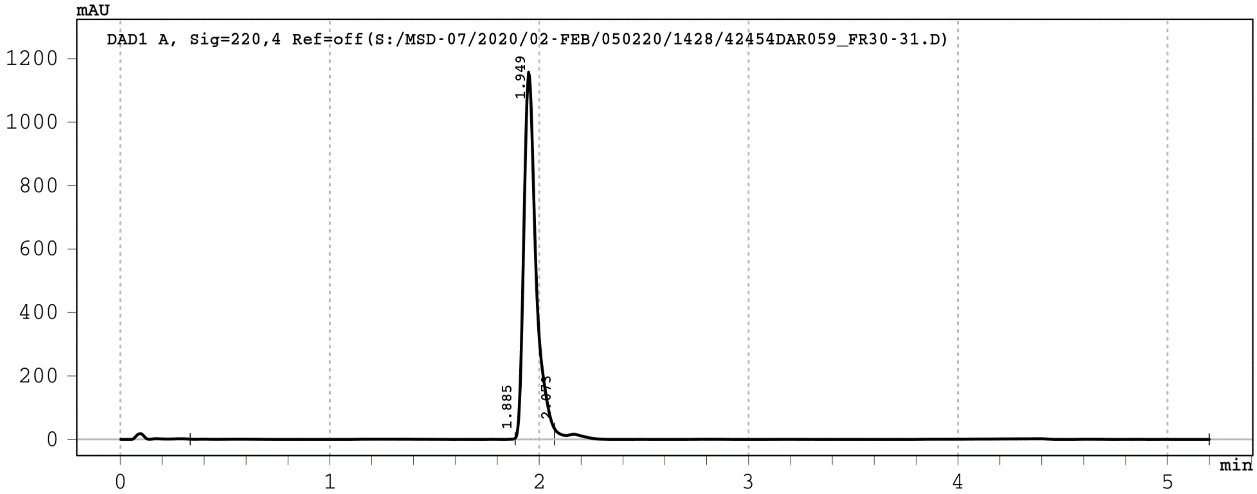


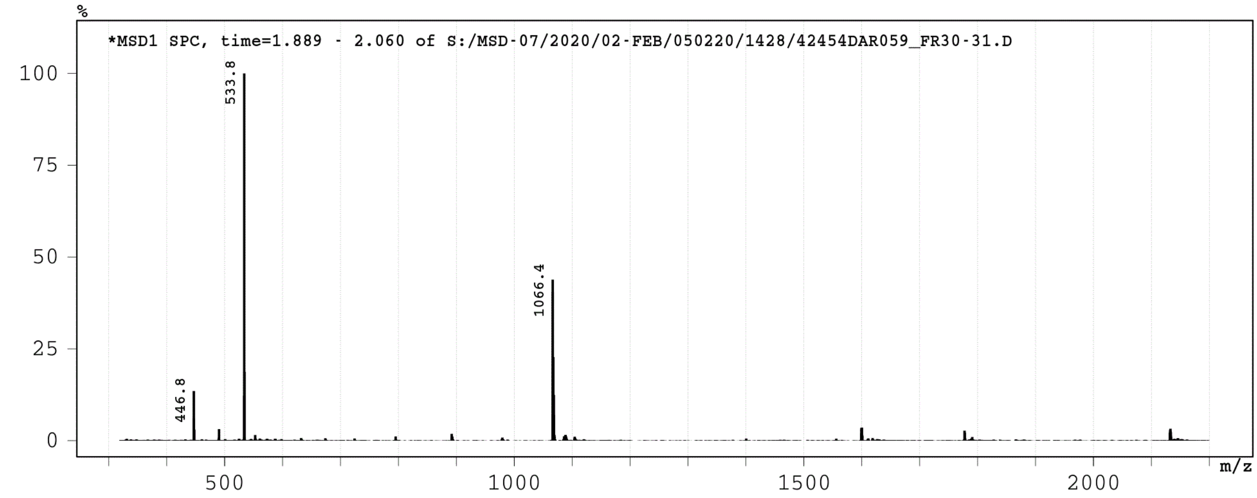


Signal 1: DAD1 A, Sig=220,4 Ref=off

| Peak # | RT [min] | Width [min] | Area | Area % |
|---|---|---|---|---|
| 1 | 1.885 | 1.55 | 27.8 | 0.6 |
| 2 | 1.949 | 0.19 | 4694.9 | 95.3 |
| 3 | 2.073 | 3.13 | 203.8 | 4.1 |

Data File : S:/MSD-07/2020/02-FEB/050220/1428/42454DAR060_FR5->

Injection Date : 05-Feb-20, 17:01:30
Sample Name : 42454DAR060_FR57
Remarks : Baseline corrected
Acq. Method : D:/METHODS/5-95-6M.M
column used : Je_559
Analysis Method : C:/CHEM32/1/METHODS/JPT-LCMS.M

Sample Info : 42454_060-Dr. Martine Darwish H-KNWINYARF-OH

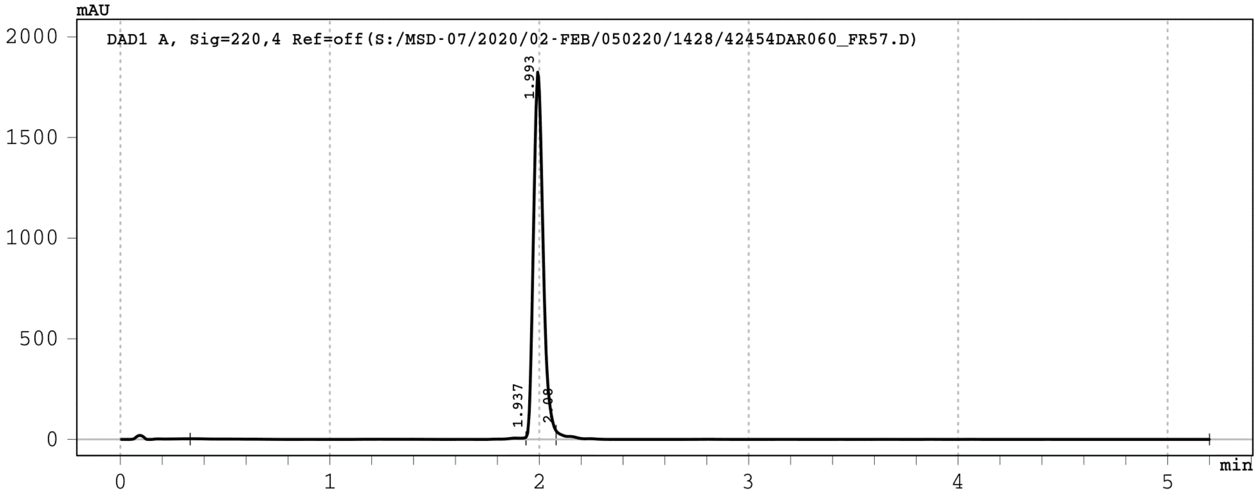


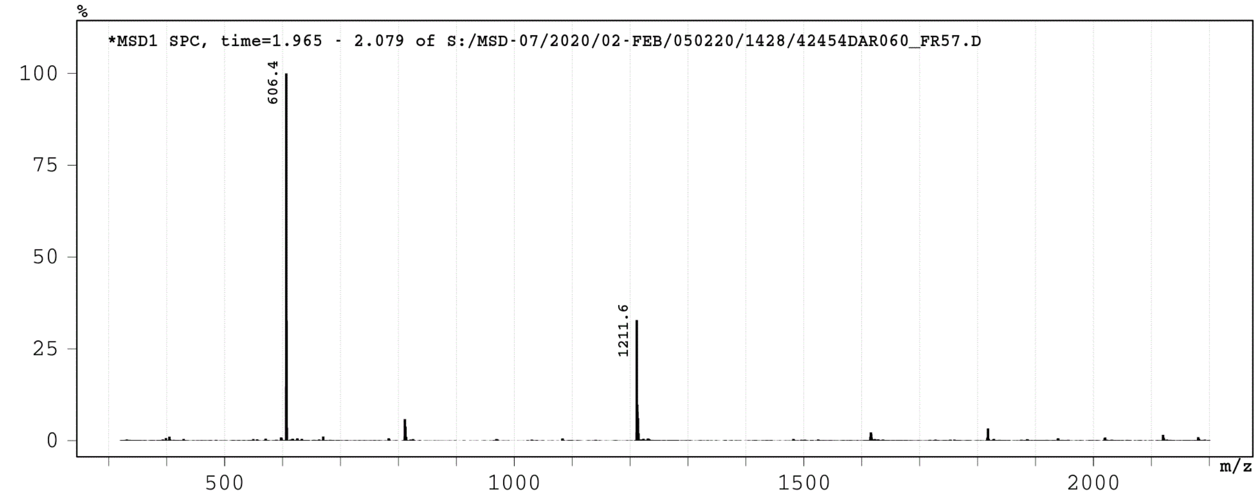


Signal 1: DAD1 A, Sig=220,4 Ref=off

| Peak # | RT [min] | Width [min] | Area | Area % |
|---|---|---|---|---|
| 1 | 1.937 | 1.60 | 94.6 | 1.5 |
| 2 | 1.993 | 0.14 | 5869.0 | 96.0 |
| 3 | 2.080 | 3.12 | 147.3 | 2.4 |

Data File : S:/MSD-07/2020/02-FEB/070220/1430/42454DAR061_FR4->

Injection Date : 07-Feb-20, 13:38:03
Sample Name : 42454DAR061_FR46-49
Remarks : Baseline corrected
Acq. Method : D:/METHODS/5-95-6M.M
column used : Je_559
Analysis Method : C:/CHEM32/1/METHODS/JPT-LCMS.M

Sample Info : 42454_061-Dr. Martine Darwish H-QFFHCYCPL-OH

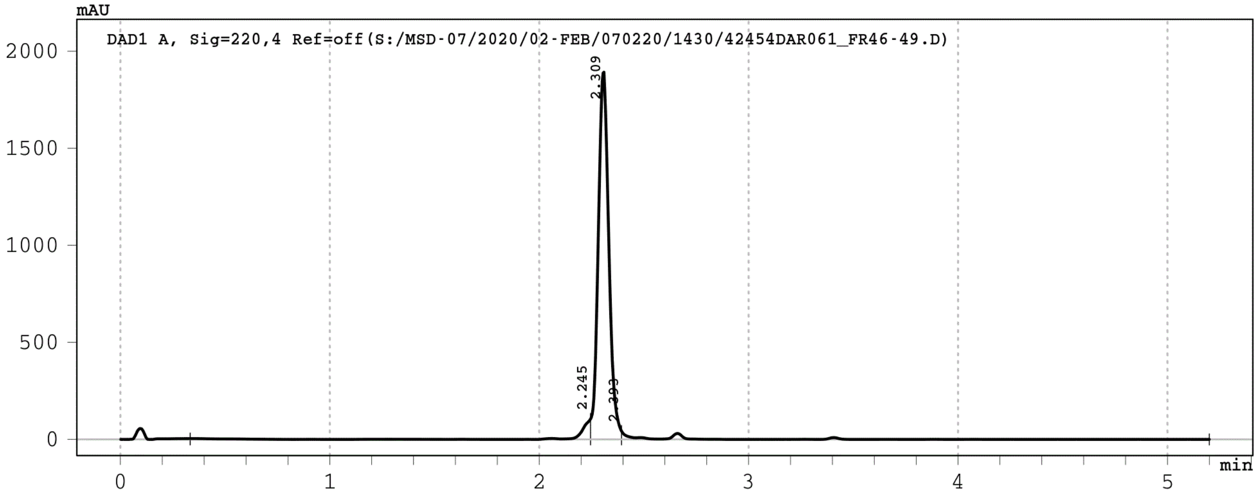


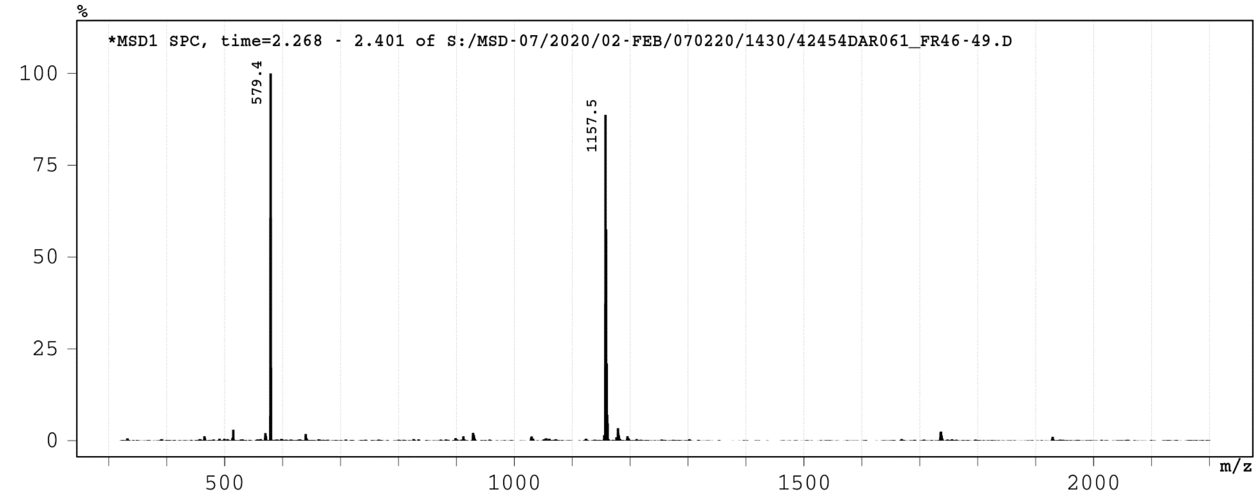


Signal 1: DAD1 A, Sig=220,4 Ref=off

| Peak # | RT [min] | Width [min] | Area | Area % |
|---|---|---|---|---|
| 1 | 2.245 | 1.91 | 344.0 | 4.7 |
| 2 | 2.309 | 0.15 | 6658.7 | 91.8 |
| 3 | 2.393 | 2.81 | 249.4 | 3.4 |

Data File : S:/MSD-07/2020/02-FEB/110220/1433/42454DAR062_FR1->

Injection Date : 11-Feb-20, 23:53:13
Sample Name : 42454DAR062_FR15-17
Remarks : Baseline corrected
Acq. Method : D:/METHODS/5-95-6M.M
column used : Je_559
Analysis Method : C:/CHEM32/1/METHODS/JPT-LCMS.M

Sample Info : 42454_062-Dr. Martine Darwish H-IVYVHRGV-OH

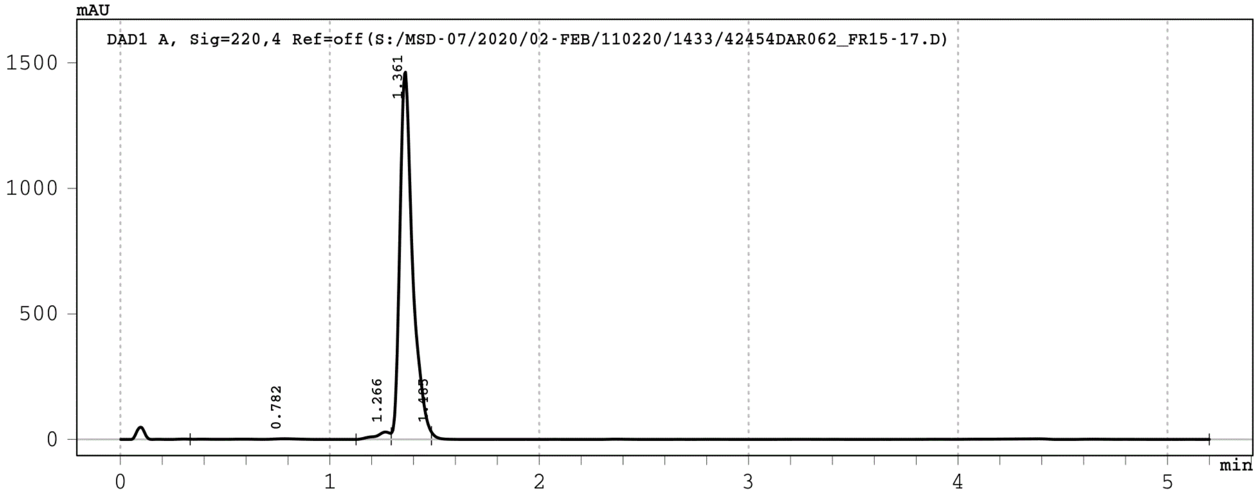


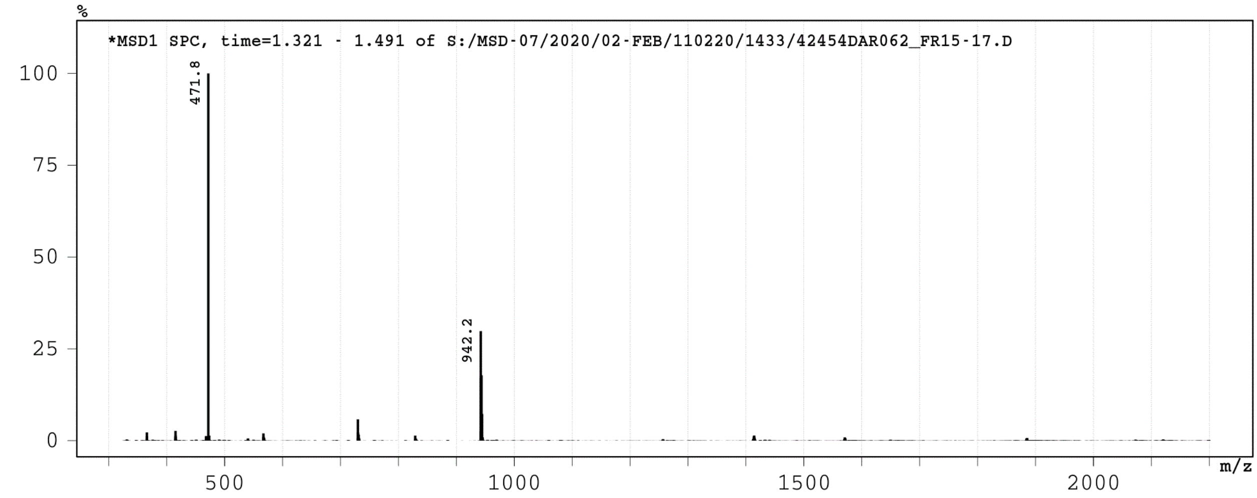


Signal 1: DAD1 A, Sig=220,4 Ref=off

| Peak # | RT [min] | Width [min] | Area | Area % |
|---|---|---|---|---|
| 1 | 0.782 | 0.79 | 32.6 | 0.5 |
| 2 | 1.266 | 0.17 | 139.3 | 2.1 |
| 3 | 1.361 | 0.19 | 6220.3 | 95.9 |
| 4 | 1.485 | 3.71 | 96.8 | 1.5 |

Data File : S:/MSD-07/2020/02-FEB/050220/1428/42454DAR063_FR7->

Injection Date : 05-Feb-20, 17:09:51
Sample Name : 42454DAR063_FR77
Remarks : Baseline corrected
Acq. Method : D:/METHODS/5-95-6M.M
column used : Je_559
Analysis Method : C:/CHEM32/1/METHODS/JPT-LCMS.M

Sample Info : 42454_063-Dr. Martine Darwish H-WMNKHLNLL-OH

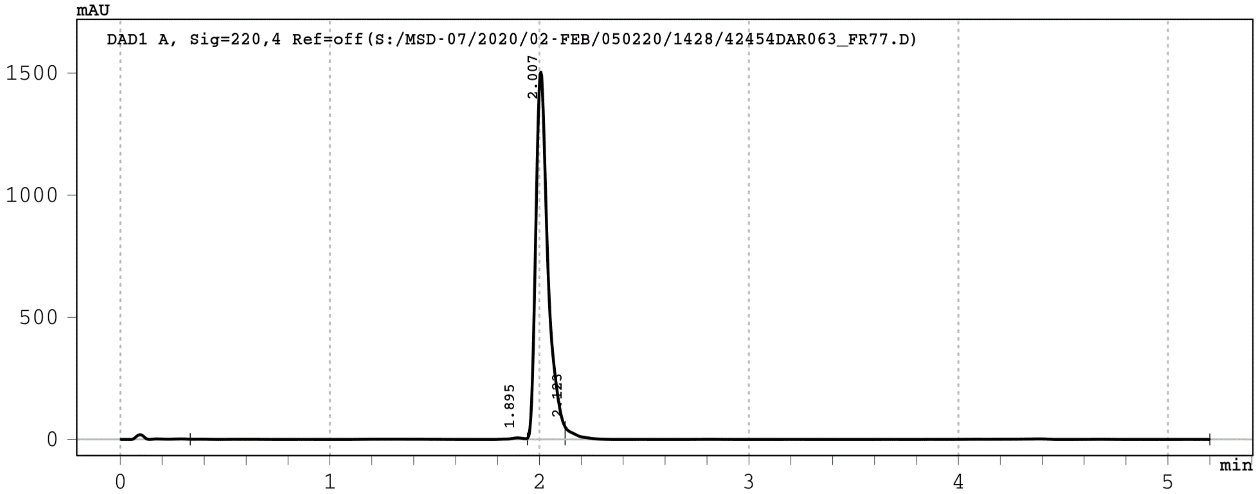


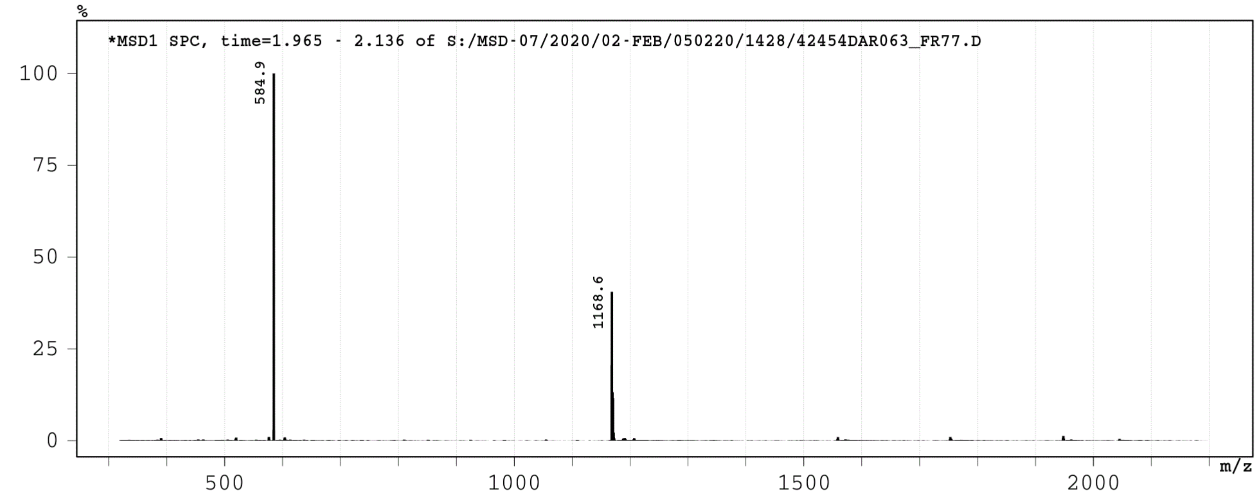


Signal 1: DAD1 A, Sig=220,4 Ref=off

| Peak # | RT [min] | Width [min] | Area | Area % |
|---|---|---|---|---|
| 1 | 1.895 | 1.61 | 57.5 | 0.9 |
| 2 | 2.007 | 0.18 | 6095.9 | 96.1 |
| 3 | 2.123 | 3.08 | 187.6 | 3.0 |

Data File : S:/MSD-07/2020/02-FEB/050220/1428/42454DAR064_FR6->

Injection Date : 05-Feb-20, 17:26:34
Sample Name : 42454DAR064_FR6-7
Remarks : Baseline corrected
Acq. Method : D:/METHODS/5-95-6M.M
column used : Je_559
Analysis Method : C:/CHEM32/1/METHODS/JPT-LCMS.M

Sample Info : 42454_064-Dr. Martine Darwish H-SAAERYKWM-OH

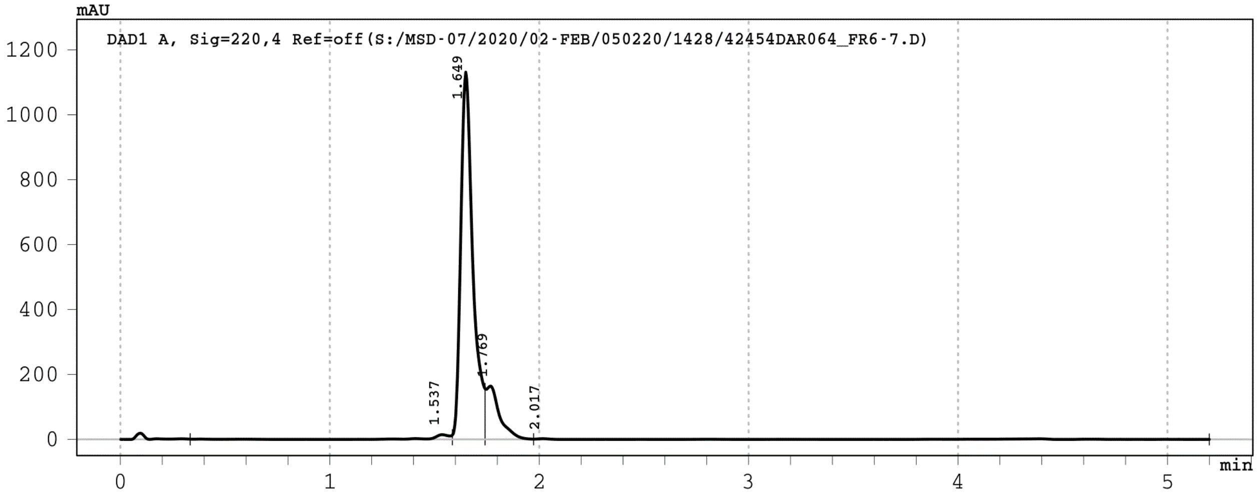


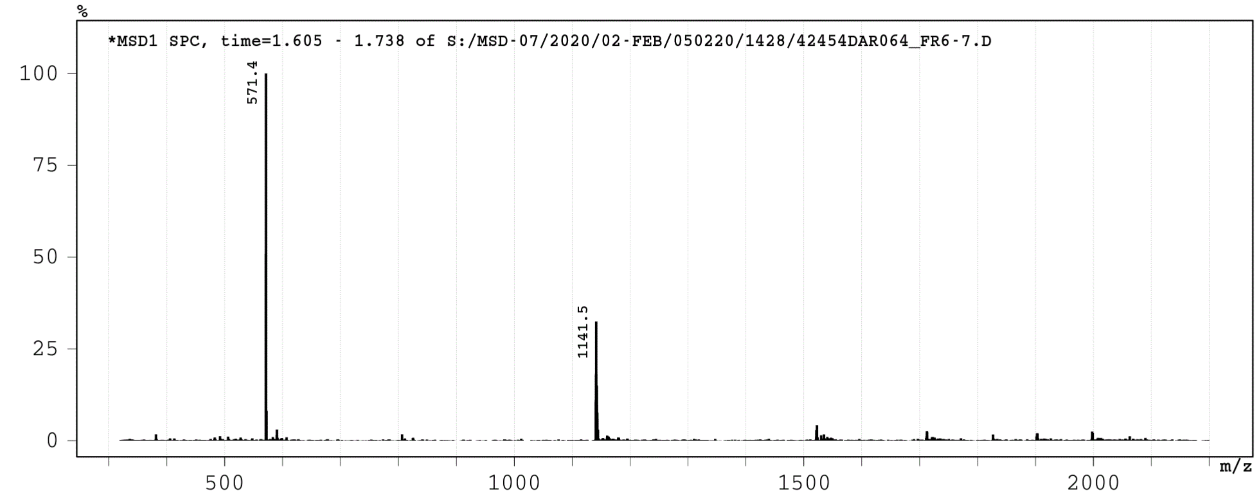


Signal 1: DAD1 A, Sig=220,4 Ref=off

| Peak # | RT [min] | Width [min] | Area | Area % |
|---|---|---|---|---|
| 1 | 1.537 | 1.25 | 97.6 | 1.8 |
| 2 | 1.649 | 0.16 | 4566.8 | 83.3 |
| 3 | 1.769 | 0.23 | 764.7 | 14.0 |
| 4 | 2.017 | 3.23 | 52.5 | 1.0 |

Data File : S:/MSD-07/2020/02-FEB/050220/1428/42454DAR065_FR3->

Injection Date : 05-Feb-20, 17:34:59
Sample Name : 42454DAR065_FR30-31
Remarks : Baseline corrected
Acq. Method : D:/METHODS/5-95-6M.M
column used : Je_559
Analysis Method : C:/CHEM32/1/METHODS/JPT-LCMS.M

Sample Info : 42454_065-Dr. Martine Darwish H-PNAVYFPLV-OH

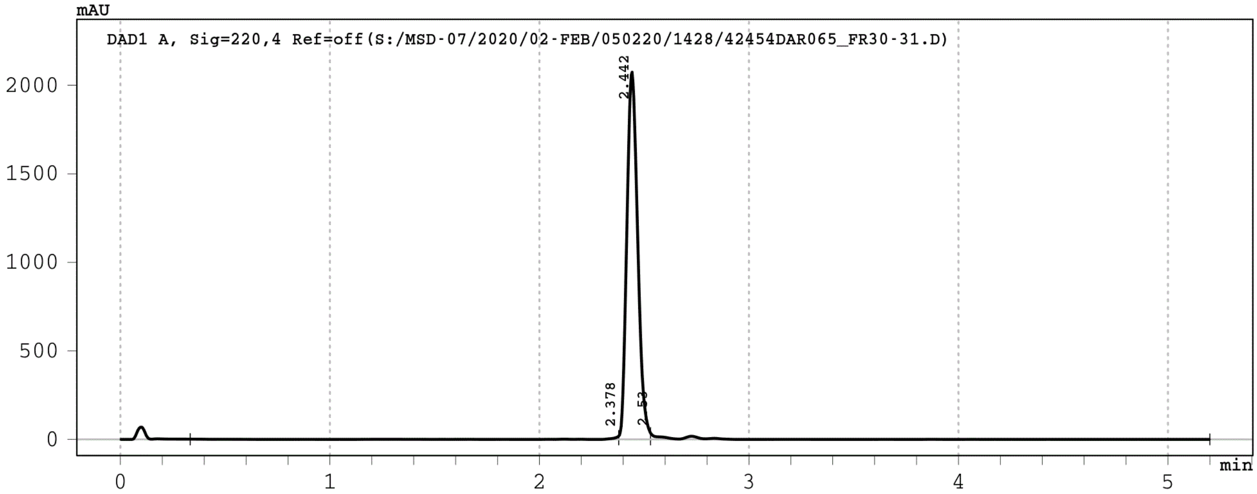


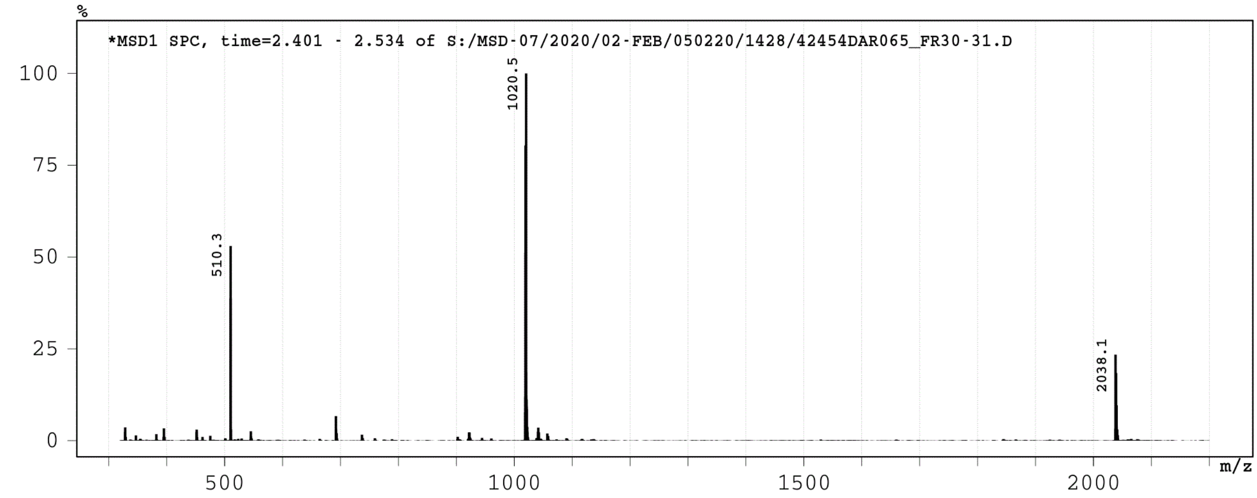


Signal 1: DAD1 A, Sig=220,4 Ref=off

| Peak # | RT [min] | Width [min] | Area | Area % |
|---|---|---|---|---|
| 1 | 2.378 | 2.05 | 60.6 | 0.8 |
| 2 | 2.442 | 0.15 | 7488.2 | 96.7 |
| 3 | 2.530 | 2.67 | 197.2 | 2.5 |

Data File : S:/MSD-07/2020/02-FEB/110220/1433/42454DAR066_FR2->

Injection Date : 12-Feb-20, 00:01:36
Sample Name : 42454DAR066_FR24-28
Remarks : Baseline corrected
Acq. Method : D:/METHODS/5-95-6M.M
column used : Je_559
Analysis Method : C:/CHEM32/1/METHODS/JPT-LCMS.M

Sample Info : 42454_066-Dr. Martine Darwish H-TKILYPRV-OH

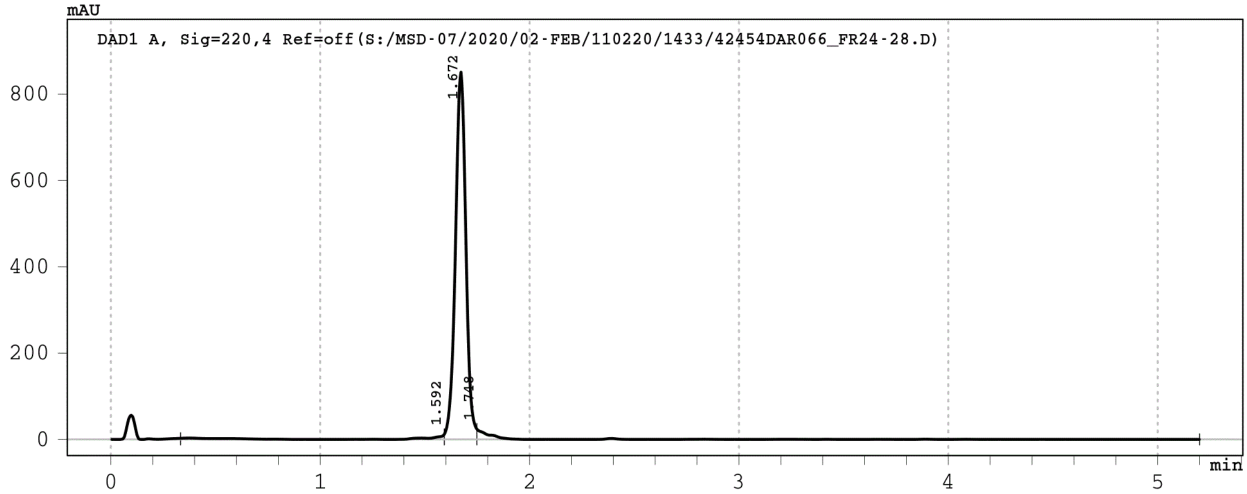


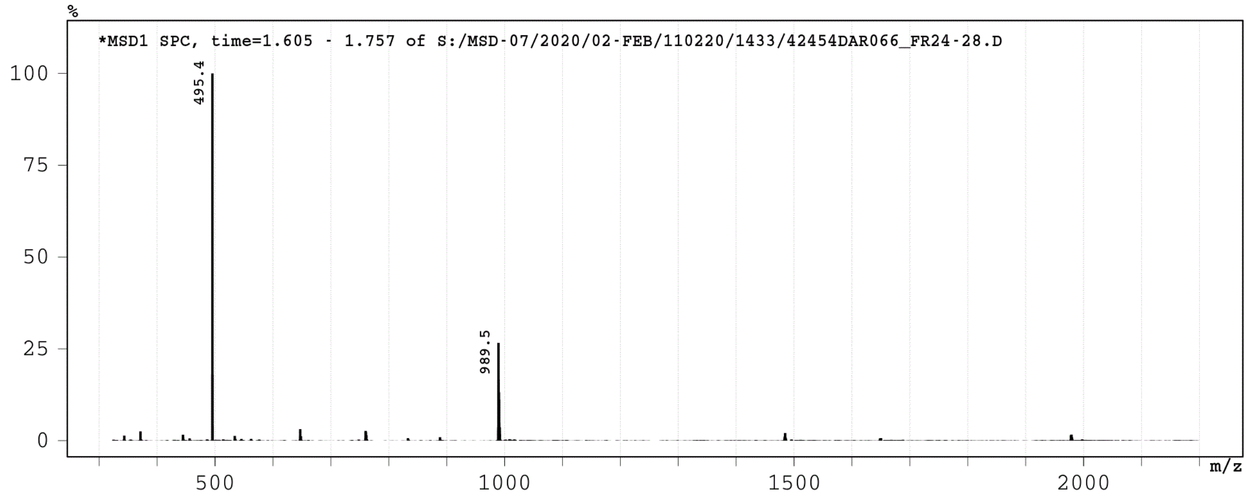


Signal 1: DAD1 A, Sig=220,4 Ref=off

| Peak # | RT [min] | Width [min] | Area | Area % |
|---|---|---|---|---|
| 1 | 1.592 | 1.26 | 91.4 | 2.9 |
| 2 | 1.672 | 0.16 | 2954.1 | 93.7 |
| 3 | 1.748 | 3.45 | 107.6 | 3.4 |

Data File : S:/MSD-07/2020/02-FEB/050220/1428/42454DAR067_FR4->

Injection Date : 05-Feb-20, 17:43:23
Sample Name : 42454DAR067_FR44-46
Remarks : Baseline corrected
Acq. Method : D:/METHODS/5-95-6M.M
column used : Je_559
Analysis Method : C:/CHEM32/1/METHODS/JPT-LCMS.M

Sample Info : 42454_067-Dr. Martine Darwish H-ALYVCGSL-OH

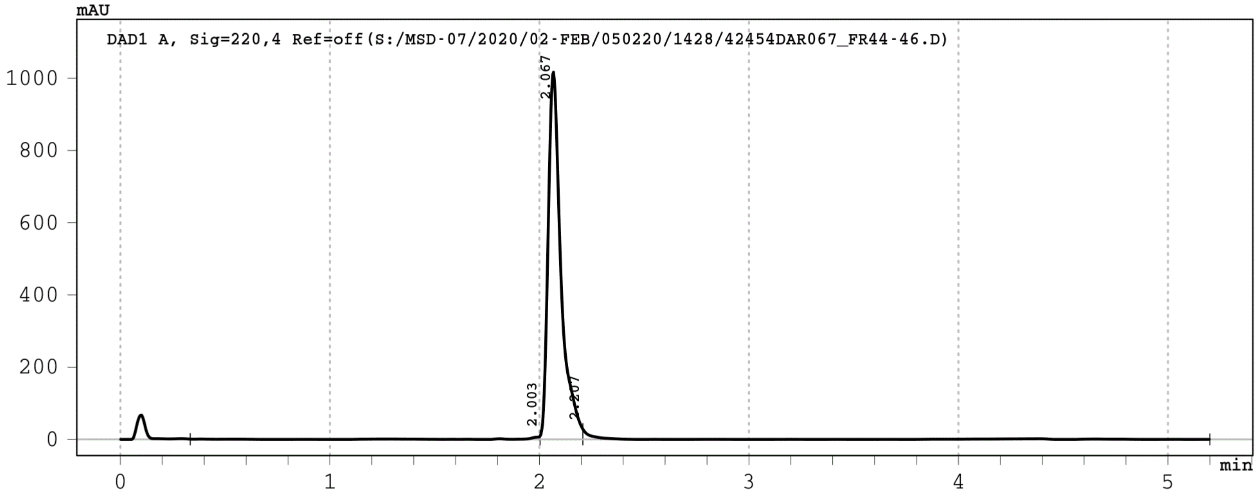


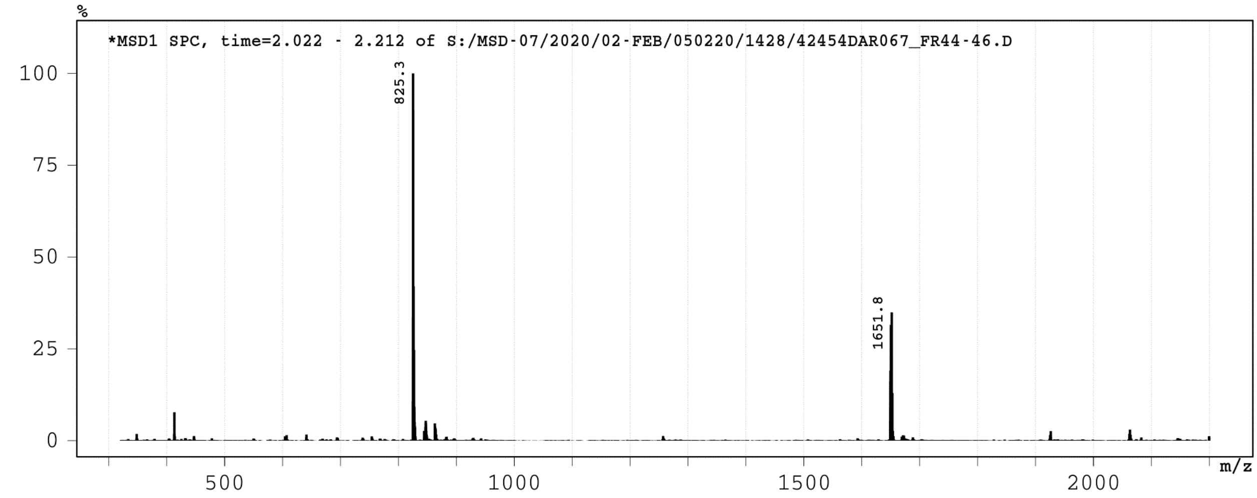


Signal 1: DAD1 A, Sig=220,4 Ref=off

| Peak # | RT [min] | Width [min] | Area | Area % |
|---|---|---|---|---|
| 1 | 2.003 | 1.67 | 57.1 | 1.3 |
| 2 | 2.067 | 0.20 | 4349.5 | 95.9 |
| 3 | 2.207 | 2.99 | 128.9 | 2.8 |

Data File : S:/MSD-07/2020/02-FEB/050220/1428/42454DAR068_FR5->

Injection Date : 05-Feb-20, 17:51:47
Sample Name : 42454DAR068_FR53-54
Remarks : Baseline corrected
Acq. Method : D:/METHODS/5-95-6M.M
column used : Je_559
Analysis Method : C:/CHEM32/1/METHODS/JPT-LCMS.M

Sample Info : 42454_068-Dr. Martine Darwish H-KALLNGDGAI-OH

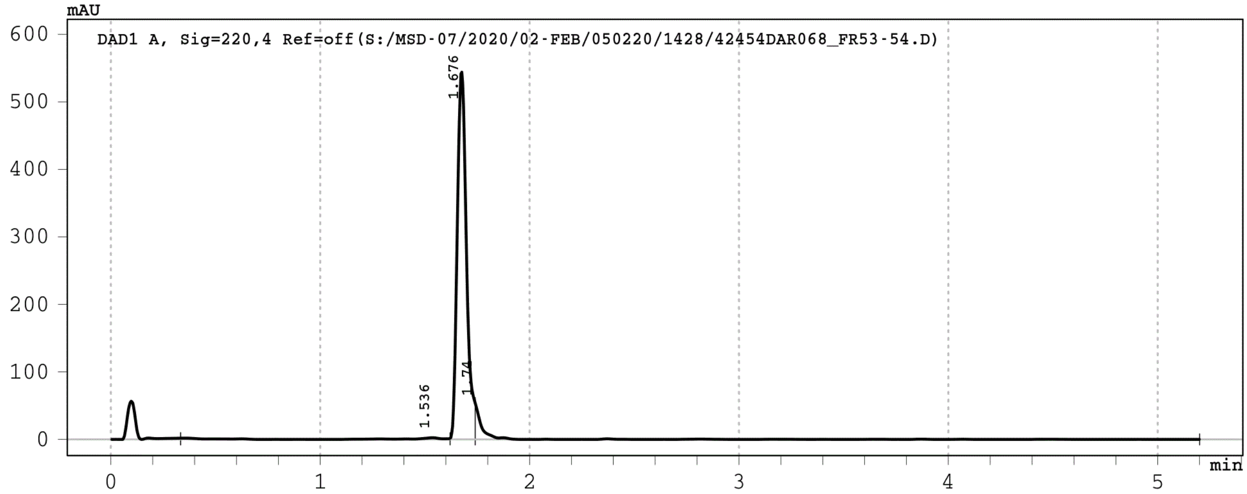


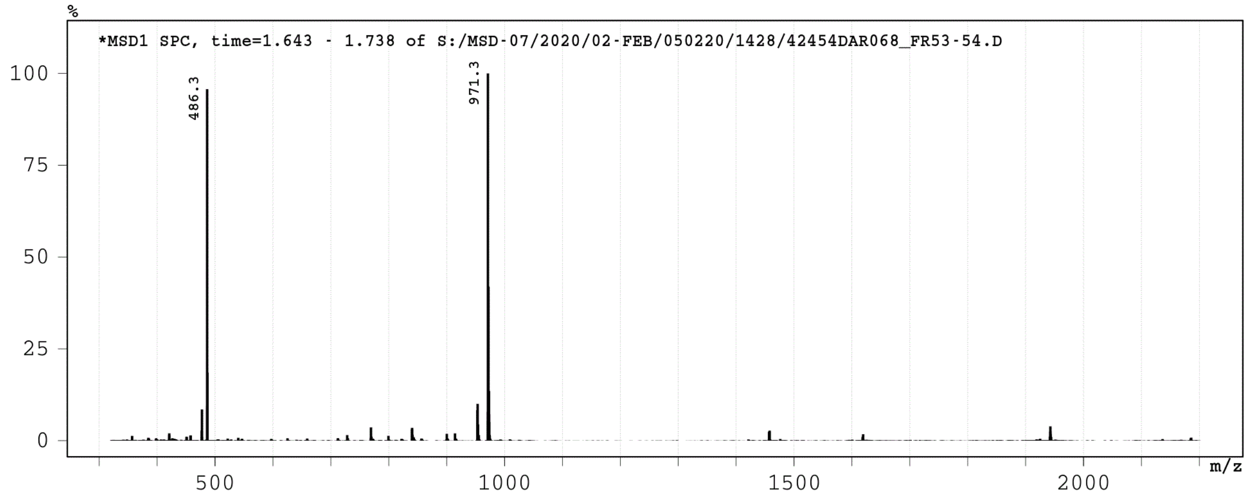


Signal 1: DAD1 A, Sig=220,4 Ref=off

| Peak # | RT [min] | Width [min] | Area | Area % |
|---|---|---|---|---|
| 1 | 1.536 | 1.29 | 43.5 | 2.4 |
| 2 | 1.676 | 0.12 | 1667.5 | 90.9 |
| 3 | 1.740 | 3.46 | 122.6 | 6.7 |

| | | |
|---|---|---|
| Data File | : | S:/MSD-07/2020/02-FEB/050220/1428/42454DAR070_FR6-> |
| Injection Date | : | 05-Feb-20, 18:00:09 |
| Sample Name | : | 42454DAR070_FR64 |
| Remarks | : | Baseline corrected |
| Acq. Method | : | D:/METHODS/5-95-6M.M |
| column used | : | Je_559 |
| Analysis Method | : | C:/CHEM32/1/METHODS/JPT-LCMS.M |
| Sample Info | : | 42454_070-Dr. Martine Darwish H-NNEDYYSLL-OH |

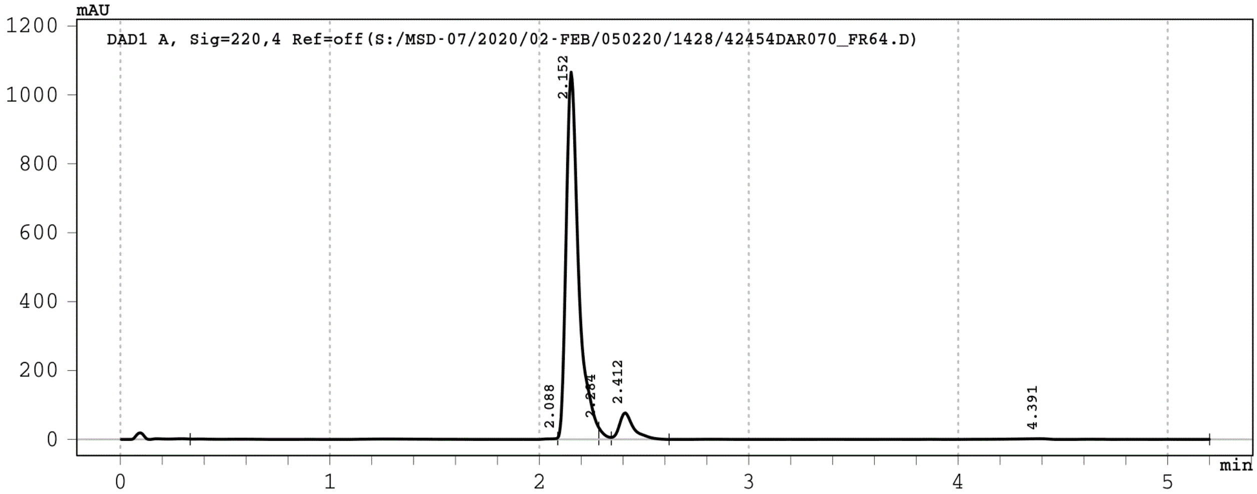


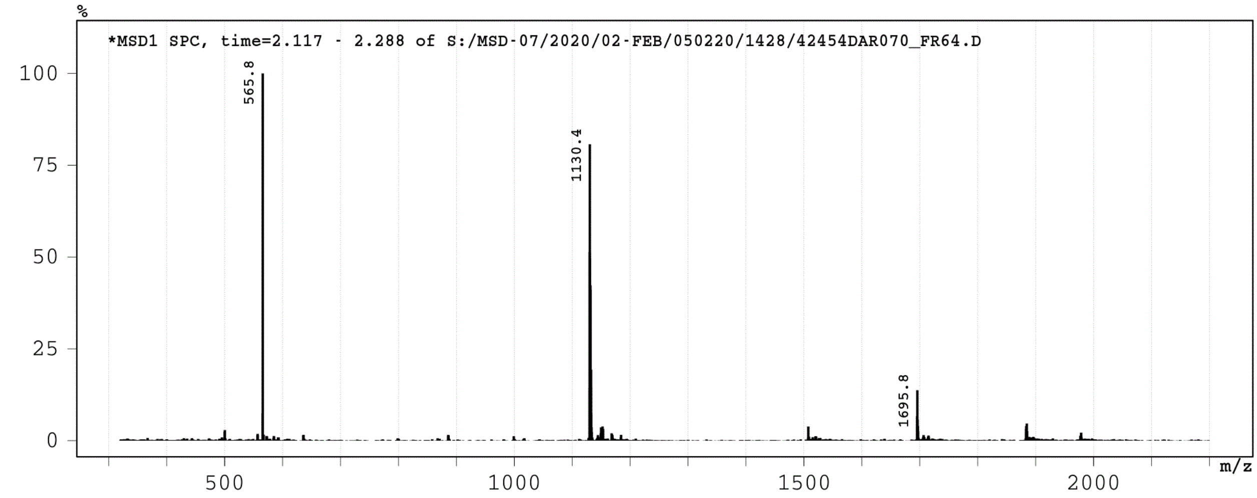


Signal 1: DAD1 A, Sig=220,4 Ref=off

| Peak # | RT [min] | Width [min] | Area | Area % |
|---|---|---|---|---|
| 1 | 2.088 | 1.75 | 44.6 | 0.9 |
| 2 | 2.152 | 0.20 | 4420.5 | 89.9 |
| 3 | 2.284 | 0.06 | 53.8 | 1.1 |
| 4 | 2.412 | 0.28 | 366.4 | 7.4 |
| 5 | 4.391 | 2.58 | 33.8 | 0.7 |

Data File : S:/MSD-07/2020/02-FEB/050220/1428/42454DAR071_FR7->

Injection Date : 05-Feb-20, 18:08:32
Sample Name : 42454DAR071_FR78
Remarks : Baseline corrected
Acq. Method : D:/METHODS/5-95-6M.M
column used : Je_559
Analysis Method : C:/CHEM32/1/METHODS/JPT-LCMS.M

Sample Info : 42454_071-Dr. Martine Darwish H-AAHRARYFL-OH

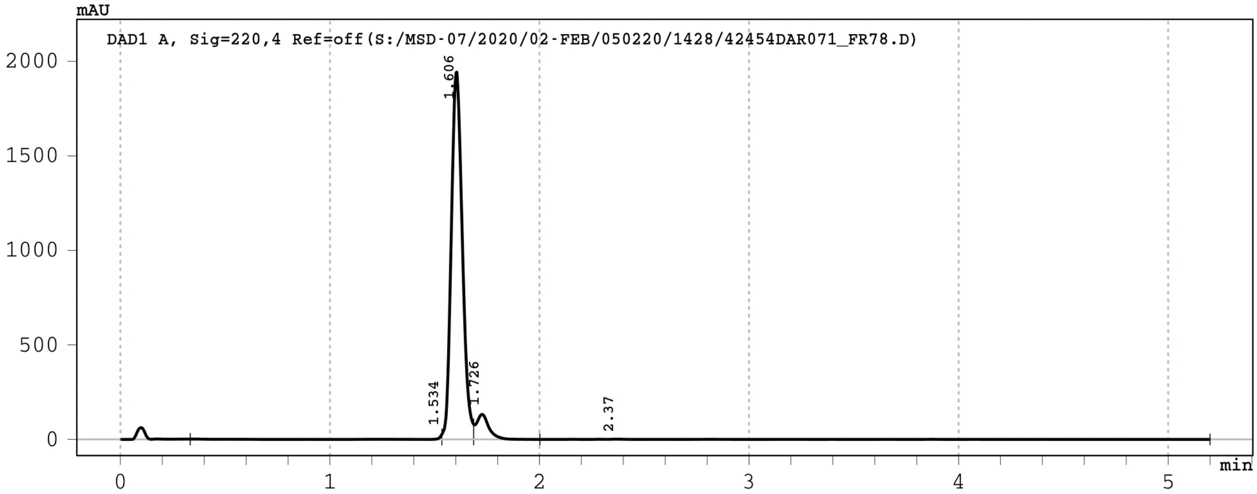


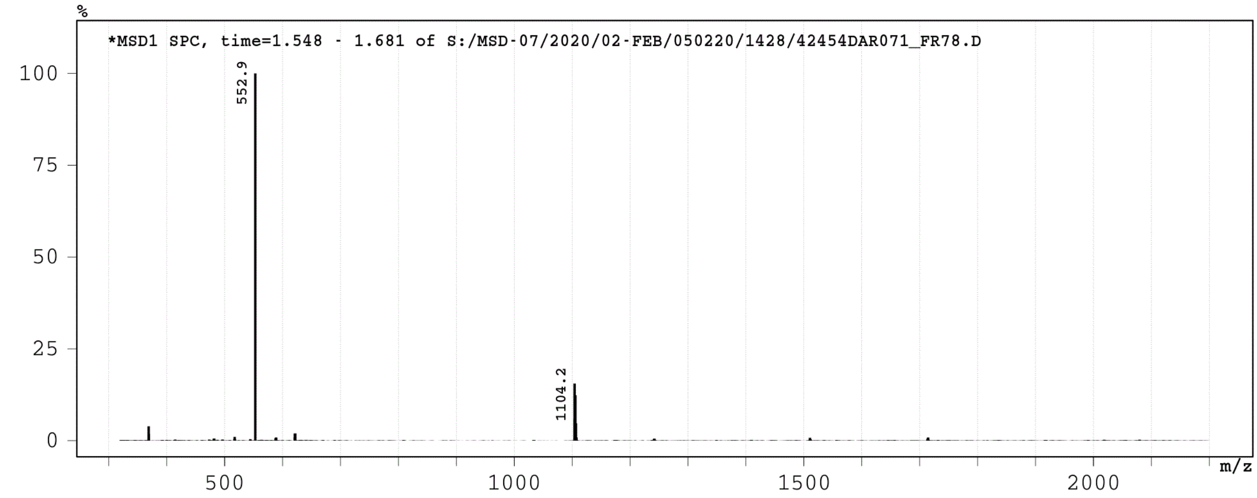


Signal 1: DAD1 A, Sig=220,4 Ref=off

| Peak # | RT [min] | Width [min] | Area | Area % |
|---|---|---|---|---|
| 1 | 1.534 | 1.20 | 39.3 | 0.5 |
| 2 | 1.606 | 0.15 | 7126.9 | 92.1 |
| 3 | 1.726 | 0.32 | 557.8 | 7.2 |
| 4 | 2.370 | 3.20 | 18.4 | 0.2 |

Data File : S:/MSD-07/2020/02-FEB/050220/1428/42454DAR072_FR4->

Injection Date : 06-Feb-20, 02:08:27
Sample Name : 42454DAR072_FR4-7
Remarks : Baseline corrected
Acq. Method : D:/METHODS/5-95-6M.M
column used : Je_559
Analysis Method : C:/CHEM32/1/METHODS/JPT-LCMS.M

Sample Info : 42454_072-Dr. Martine Darwish H-SIIVFNLL-OH

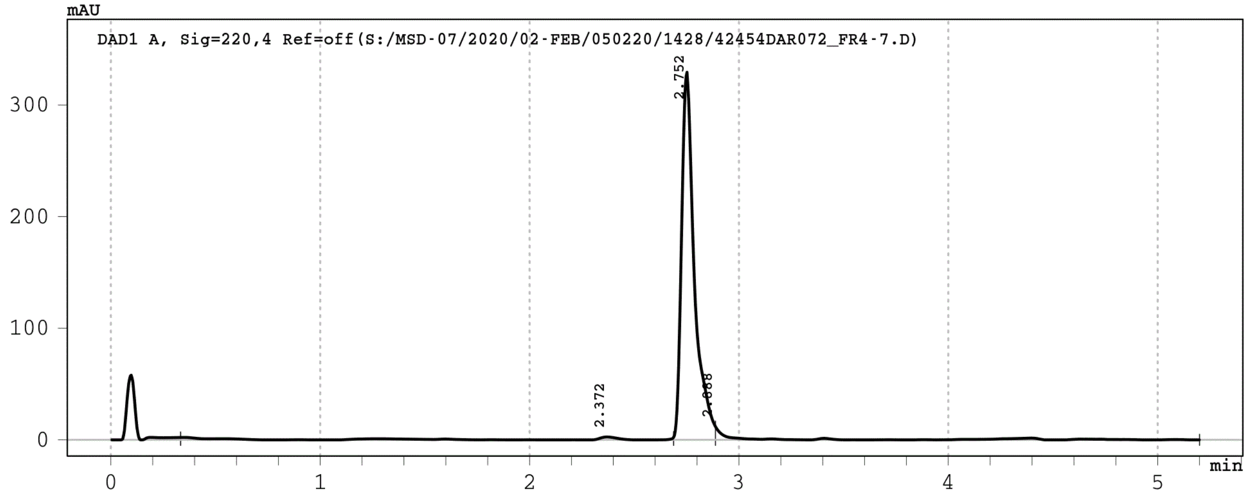


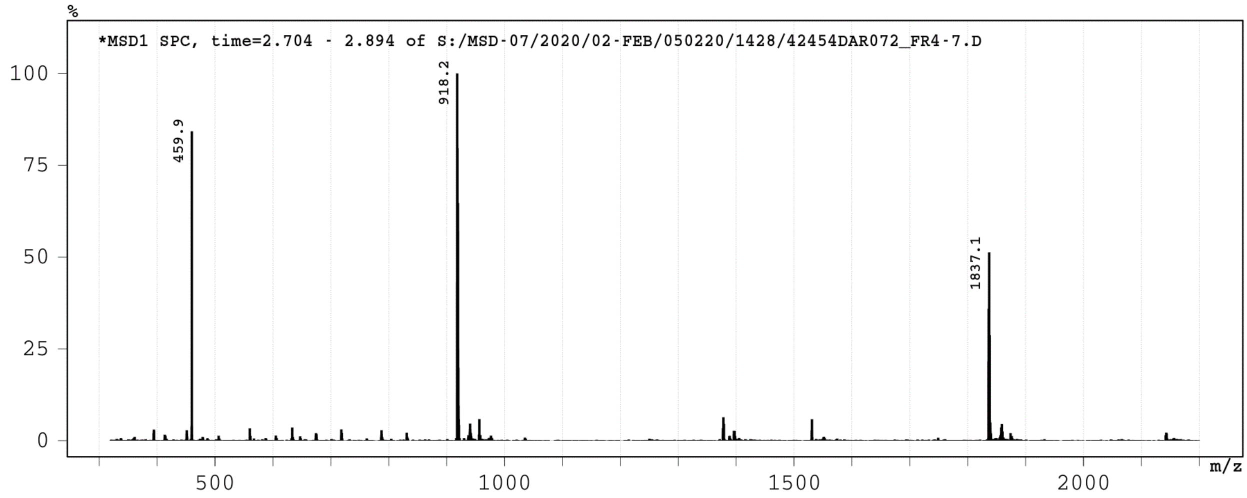


Signal 1: DAD1 A, Sig=220,4 Ref=off

| Peak # | RT [min] | Width [min] | Area | Area % |
|---|---|---|---|---|
| 1 | 2.372 | 2.35 | 62.3 | 4.0 |
| 2 | 2.752 | 0.20 | 1406.3 | 91.1 |
| 3 | 2.888 | 2.31 | 74.8 | 4.8 |

Data File : S:/MSD-07/2020/02-FEB/050220/1428/42454DAR073_FR6->

Injection Date : 06-Feb-20, 02:33:40
Sample Name : 42454DAR073_FR60-69
Remarks : Baseline corrected
Acq. Method : D:/METHODS/5-95-6M.M
column used : Je_559
Analysis Method : C:/CHEM32/1/METHODS/JPT-LCMS.M

Sample Info : 42454_073-Dr. Martine Darwish H-PVYFYIAIV-OH

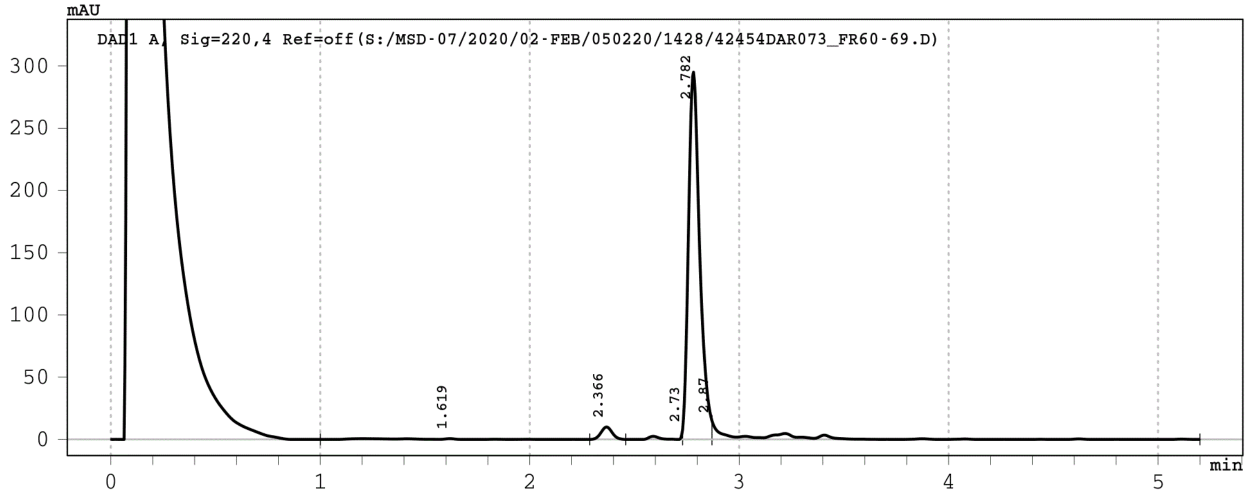


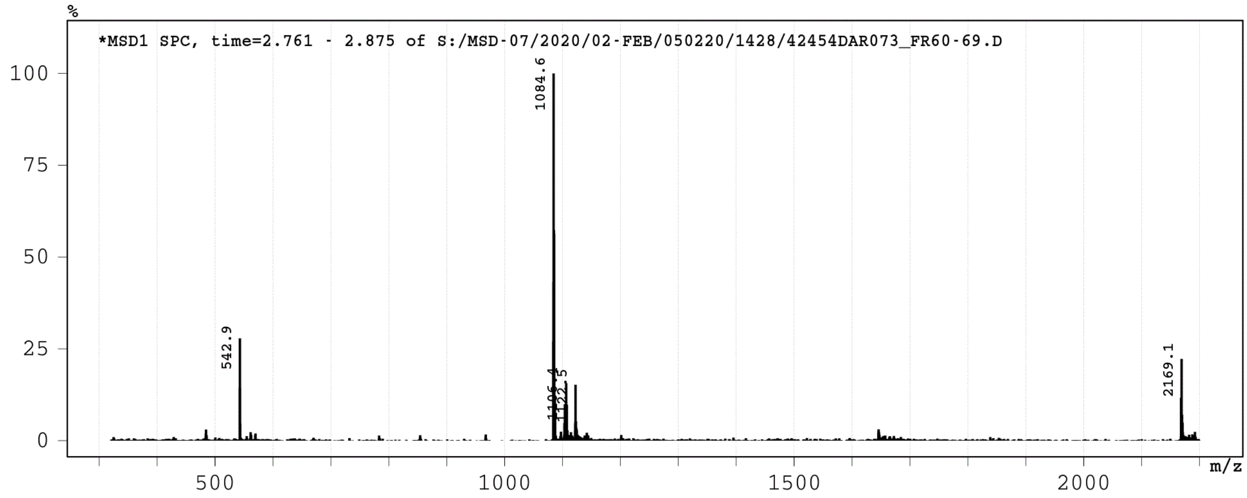


Signal 1: DAD1 A, Sig=220,4 Ref=off

| Peak # | RT [min] | Width [min] | Area | Area % |
|---|---|---|---|---|
| 1 | 1.619 | 1.29 | 12.7 | 1.0 |
| 2 | 2.366 | 0.17 | 36.9 | 2.9 |
| 3 | 2.730 | 0.27 | 10.7 | 0.8 |
| 4 | 2.782 | 0.14 | 1105.1 | 86.8 |
| 5 | 2.870 | 2.33 | 108.2 | 8.5 |

Data File : S:/MSD-07/2020/02-FEB/050220/1428/42454DAR074_FR7->

Injection Date : 06-Feb-20, 02:42:04
Sample Name : 42454DAR074_FR77-78
Remarks : Baseline corrected
Acq. Method : D:/METHODS/5-95-6M.M
column used : Je_559
Analysis Method : C:/CHEM32/1/METHODS/JPT-LCMS.M

Sample Info : 42454_074-Dr. Martine Darwish H-SSISKAML-OH

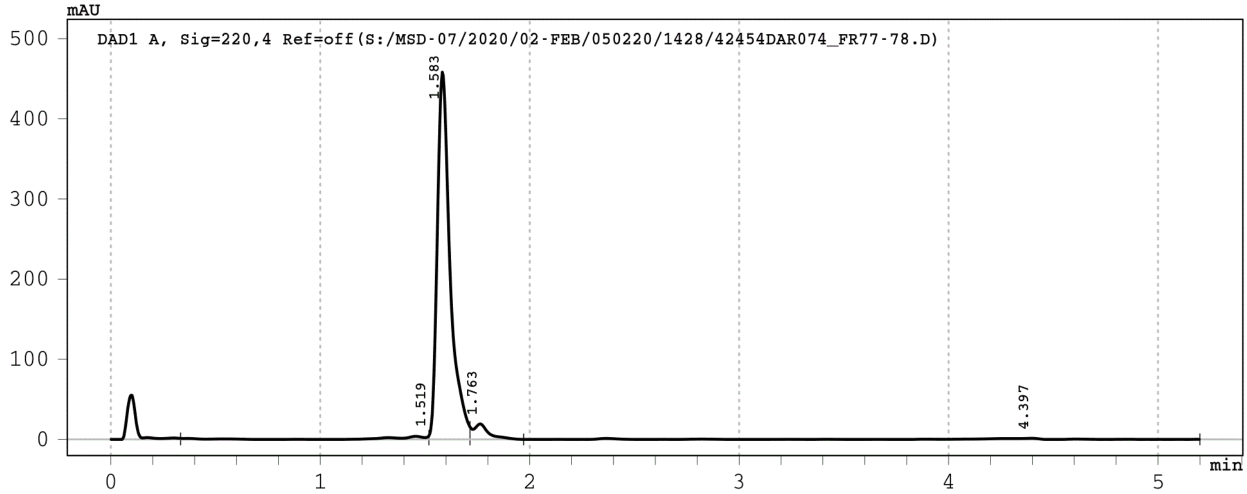


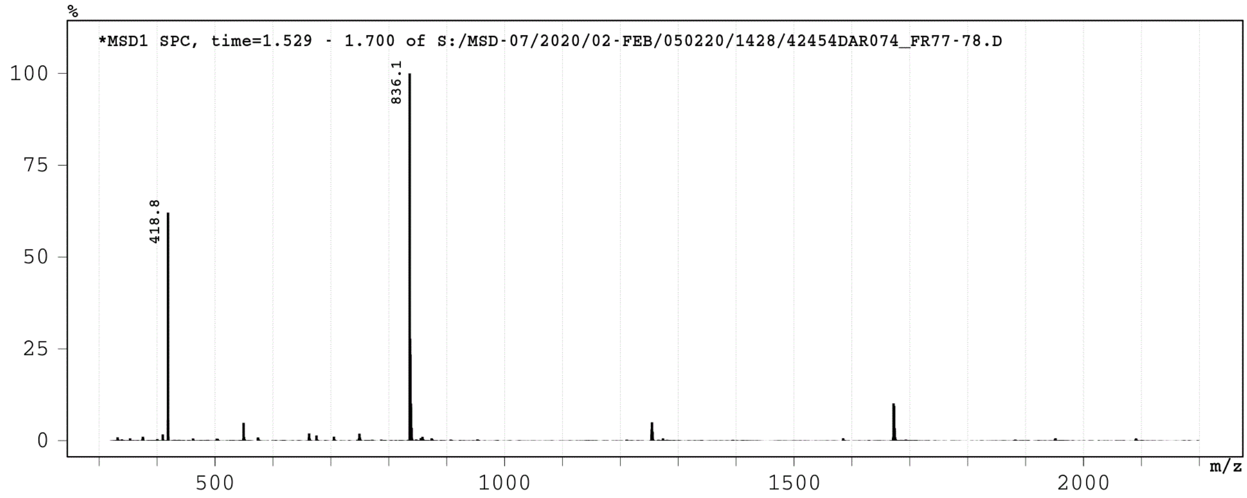


Signal 1: DAD1 A, Sig=220,4 Ref=off

| Peak # | RT [min] | Width [min] | Area | Area % |
|---|---|---|---|---|
| 1 | 1.519 | 1.19 | 50.9 | 2.4 |
| 2 | 1.583 | 0.20 | 1956.9 | 91.1 |
| 3 | 1.763 | 0.26 | 102.4 | 4.8 |
| 4 | 4.397 | 3.23 | 38.4 | 1.8 |

Data File : S:/MSD-07/2020/02-FEB/050220/1428/42454DAR075_FR8->

Injection Date : 06-Feb-20, 02:50:30
Sample Name : 42454DAR075_FR88-90
Remarks : Baseline corrected
Acq. Method : D:/METHODS/5-95-6M.M
column used : Je_559
Analysis Method : C:/CHEM32/1/METHODS/JPT-LCMS.M

Sample Info : 42454_075-Dr. Martine Darwish H-SSLTSSVPV-OH

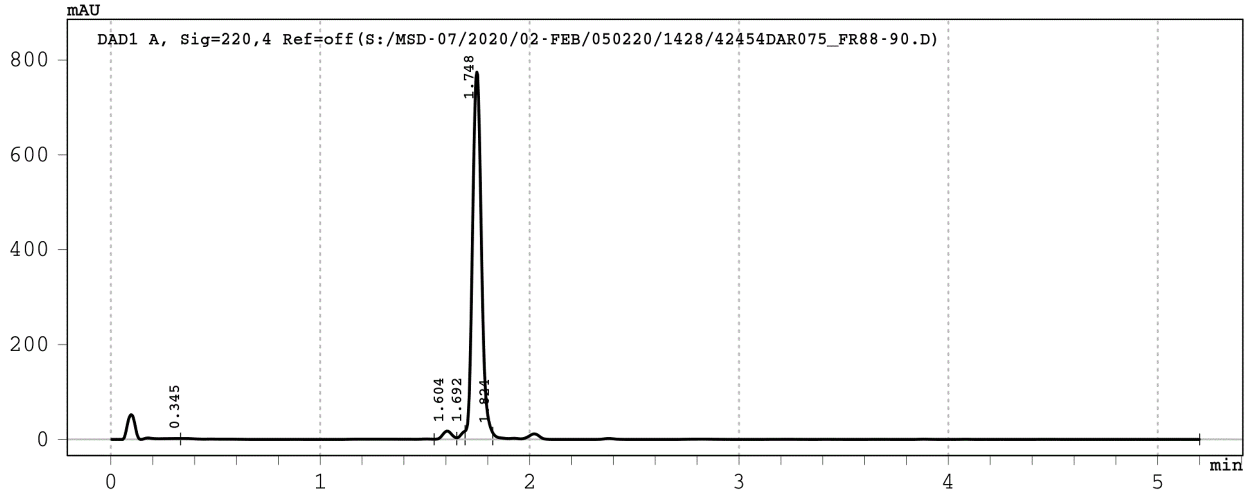


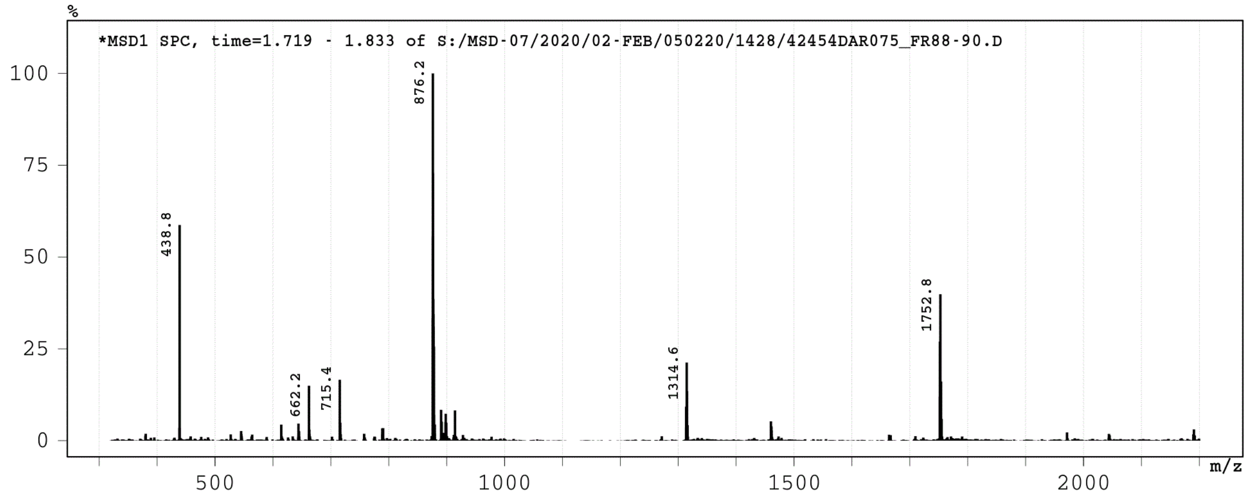


Signal 1: DAD1 A, Sig=220,4 Ref=off

| Peak # | RT [min] | Width [min] | Area | Area % |
|---|---|---|---|---|
| 1 | 0.345 | 1.21 | 26.7 | 1.1 |
| 2 | 1.604 | 0.11 | 54.1 | 2.2 |
| 3 | 1.692 | 0.04 | 24.0 | 1.0 |
| 4 | 1.748 | 0.13 | 2280.9 | 92.3 |
| 5 | 1.824 | 3.38 | 85.9 | 3.5 |

Data File : S:/MSD-07/2020/02-FEB/050220/1428/42454DAR076_FR3->

Injection Date : 06-Feb-20, 02:58:53
Sample Name : 42454DAR076_FR3-4
Remarks : Baseline corrected
Acq. Method : D:/METHODS/5-95-6M.M
column used : Je_559
Analysis Method : C:/CHEM32/1/METHODS/JPT-LCMS.M

Sample Info : 42454_076-Dr. Martine Darwish H-KAYTKPELL-OH

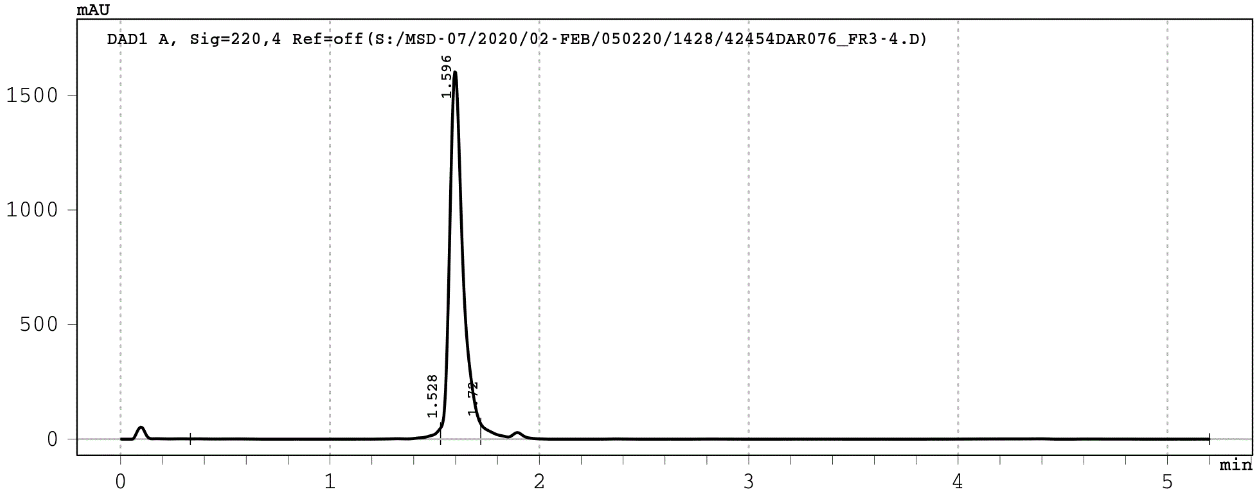


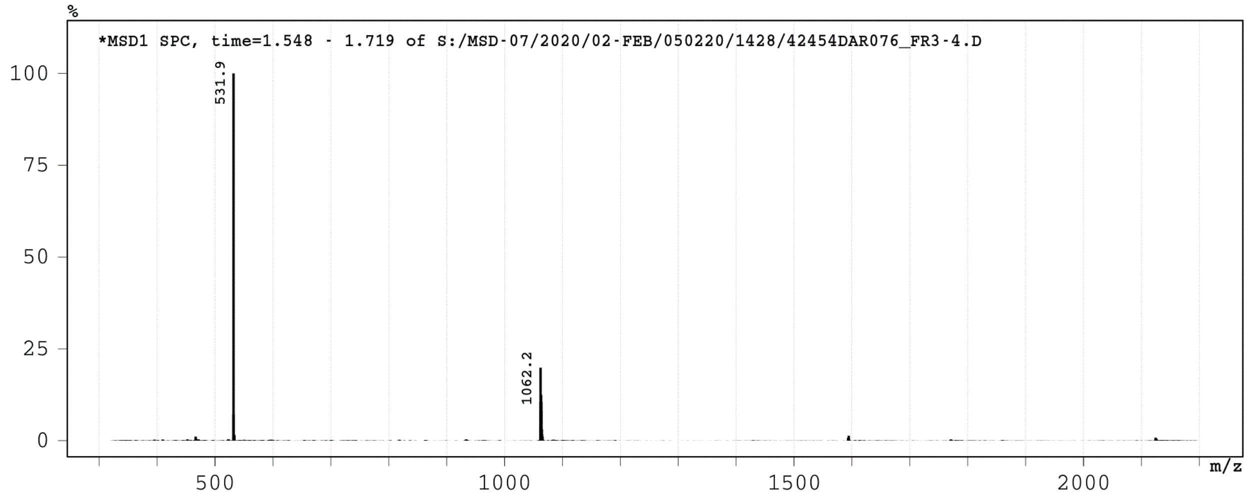


Signal 1: DAD1 A, Sig=220,4 Ref=off

| Peak # | RT [min] | Width [min] | Area | Area % |
|---|---|---|---|---|
| 1 | 1.528 | 1.19 | 145.4 | 1.9 |
| 2 | 1.596 | 0.19 | 7085.9 | 93.1 |
| 3 | 1.720 | 3.48 | 380.8 | 5.0 |

Data File : S:/MSD-07/2020/02-FEB/180220/1438/42454DAR077_FR9->

Injection Date : 18-Feb-20, 09:41:19
Sample Name : 42454DAR077_FR9
Remarks : Baseline corrected
Acq. Method : D:/METHODS/5-95-6M.M
column used : Je_559
Analysis Method : C:/CHEM32/1/METHODS/JPT-LCMS.M

Sample Info : 42454_077-Dr. Martine Darwish H-QIFDYNYNGL-OH

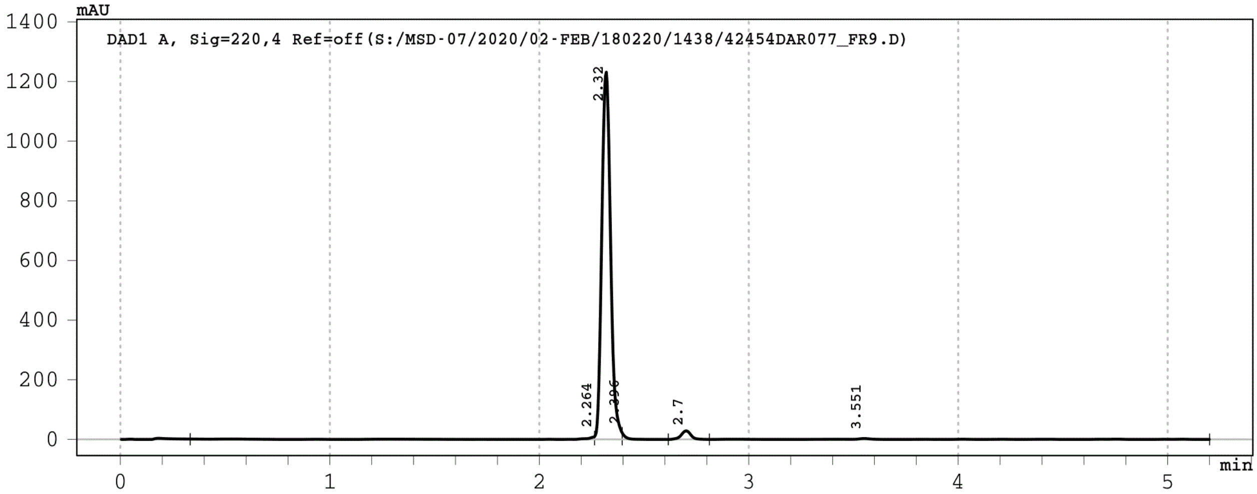


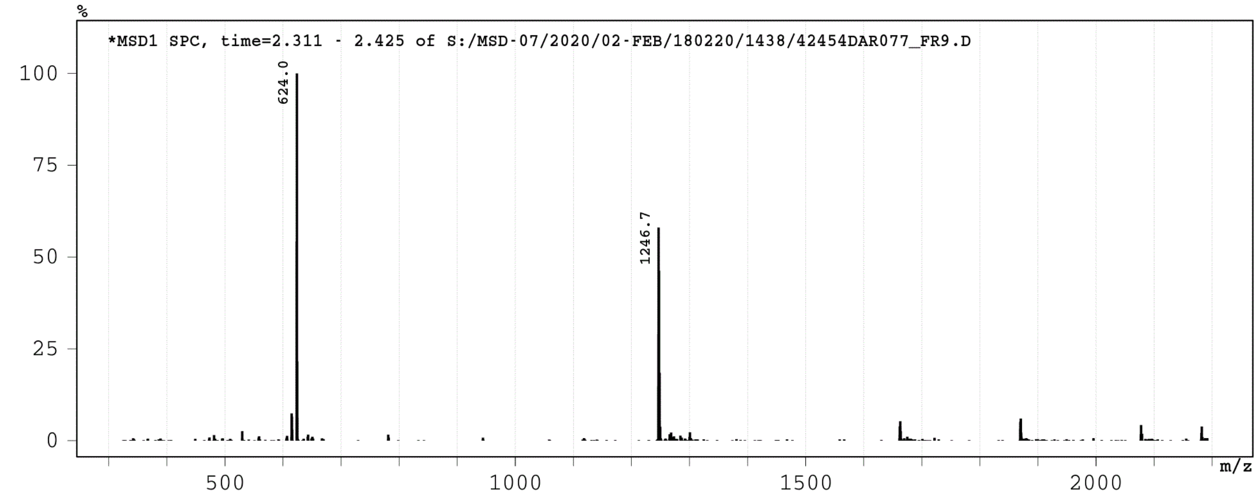


Signal 1: DAD1 A, Sig=220,4 Ref=off

| Peak # | RT [min] | Width [min] | Area | Area % |
|---|---|---|---|---|
| 1 | 2.264 | 1.93 | 58.0 | 1.5 |
| 2 | 2.320 | 0.13 | 3604.2 | 94.6 |
| 3 | 2.396 | 0.22 | 21.8 | 0.6 |
| 4 | 2.700 | 0.20 | 92.0 | 2.4 |
| 5 | 3.551 | 2.39 | 35.6 | 0.9 |

Data File : S:/MSD-07/2020/02-FEB/050220/1428/42454DAR078_FR1->

Injection Date : 06-Feb-20, 03:07:17
Sample Name : 42454DAR078_FR16
Remarks : Baseline corrected
Acq. Method : D:/METHODS/5-95-6M.M
column used : Je_559
Analysis Method : C:/CHEM32/1/METHODS/JPT-LCMS.M

Sample Info : 42454_078-Dr. Martine Darwish H-SCRTFLSPL-OH

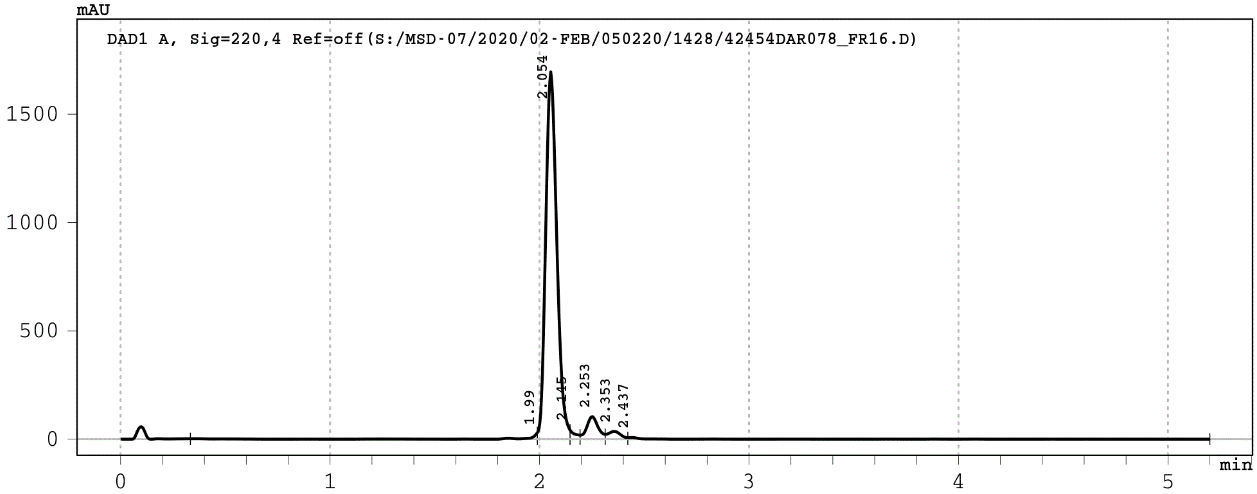


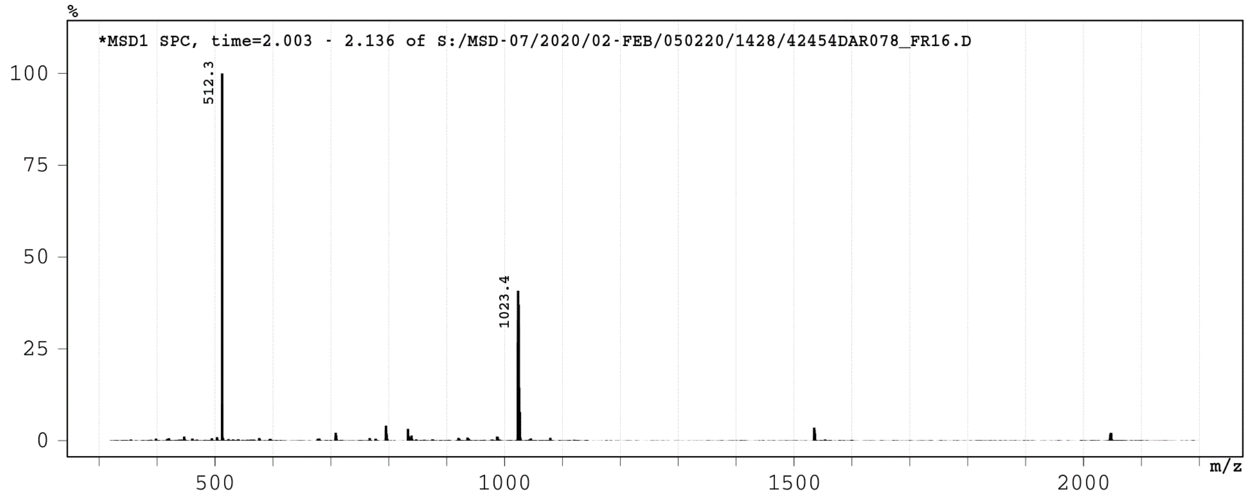


Signal 1: DAD1 A, Sig=220,4 Ref=off

| Peak # | RT [min] | Width [min] | Area | Area % |
|---|---|---|---|---|
| 1 | 1.990 | 1.66 | 88.8 | 1.3 |
| 2 | 2.054 | 0.16 | 6253.6 | 89.2 |
| 3 | 2.145 | 0.05 | 72.9 | 1.0 |
| 4 | 2.253 | 0.12 | 400.1 | 5.7 |
| 5 | 2.353 | 0.11 | 153.9 | 2.2 |
| 6 | 2.437 | 2.78 | 39.0 | 0.6 |

Data File : S:/MSD-07/2020/02-FEB/050220/1428/42454DAR079_FR4->

Injection Date : 06-Feb-20, 03:24:08
Sample Name : 42454DAR079_FR40-41
Remarks : Baseline corrected
Acq. Method : D:/METHODS/5-95-6M.M
column used : Je_559
Analysis Method : C:/CHEM32/1/METHODS/JPT-LCMS.M

Sample Info : 42454_079-Dr. Martine Darwish H-TSYCPSDL-OH

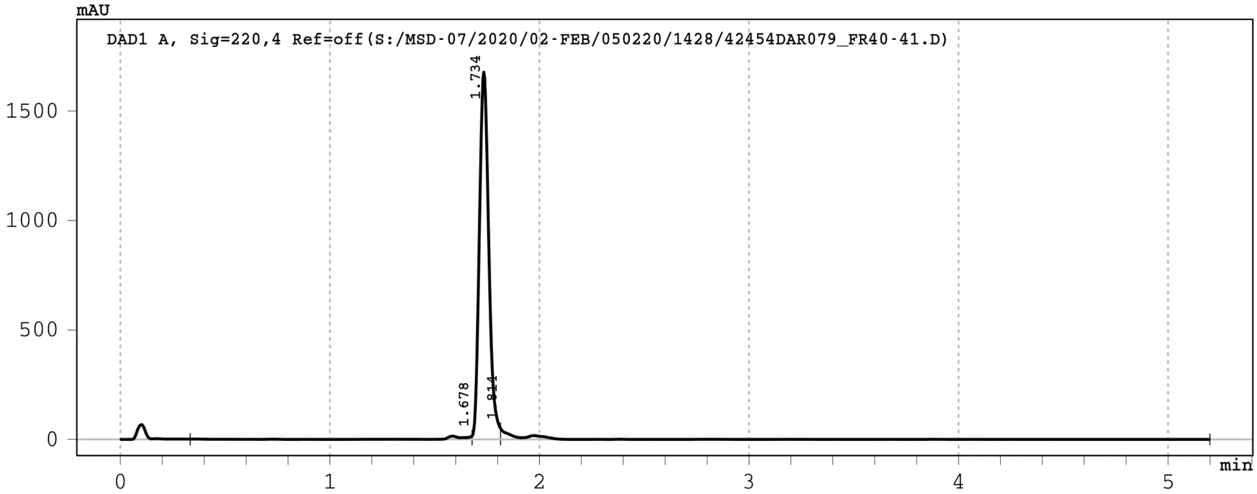


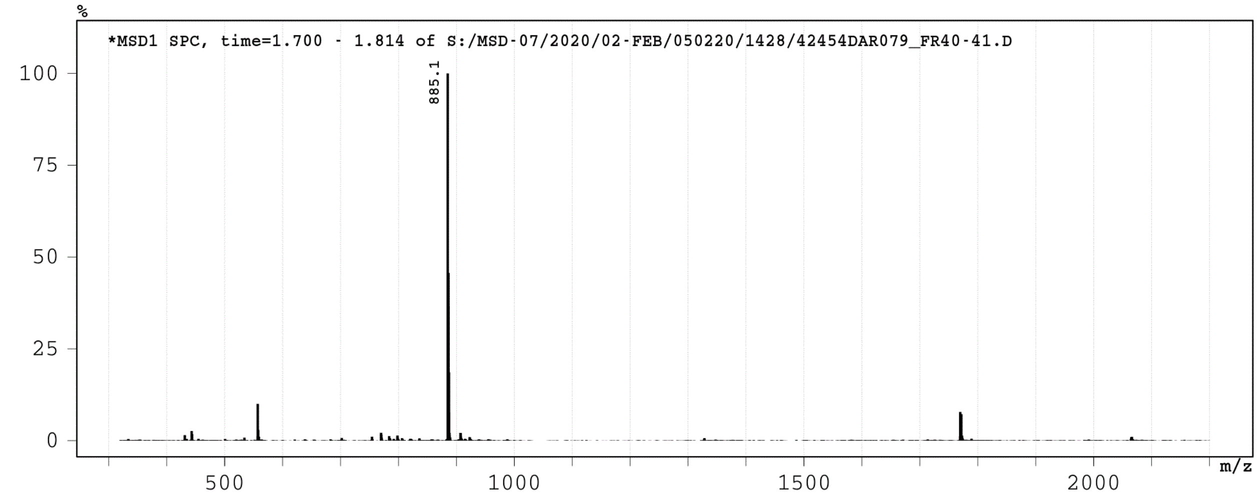


Signal 1: DAD1 A, Sig=220,4 Ref=off

| Peak # | RT [min] | Width [min] | Area | Area % |
|---|---|---|---|---|
| 1 | 1.678 | 1.35 | 108.3 | 1.9 |
| 2 | 1.734 | 0.14 | 5254.0 | 93.5 |
| 3 | 1.814 | 3.39 | 254.4 | 4.5 |

Data File : S:/MSD-07/2020/02-FEB/060220/1429/42454DAR080_FR1->

Injection Date : 06-Feb-20, 14:51:25
Sample Name : 42454DAR080_FR10-11
Remarks : Baseline corrected
Acq. Method : D:/METHODS/5-95-6M.M
column used : Je_559
Analysis Method : C:/CHEM32/1/METHODS/JPT-LCMS.M

Sample Info : 42454_080-Dr. Martine Darwish H-LQPLFYAL-OH

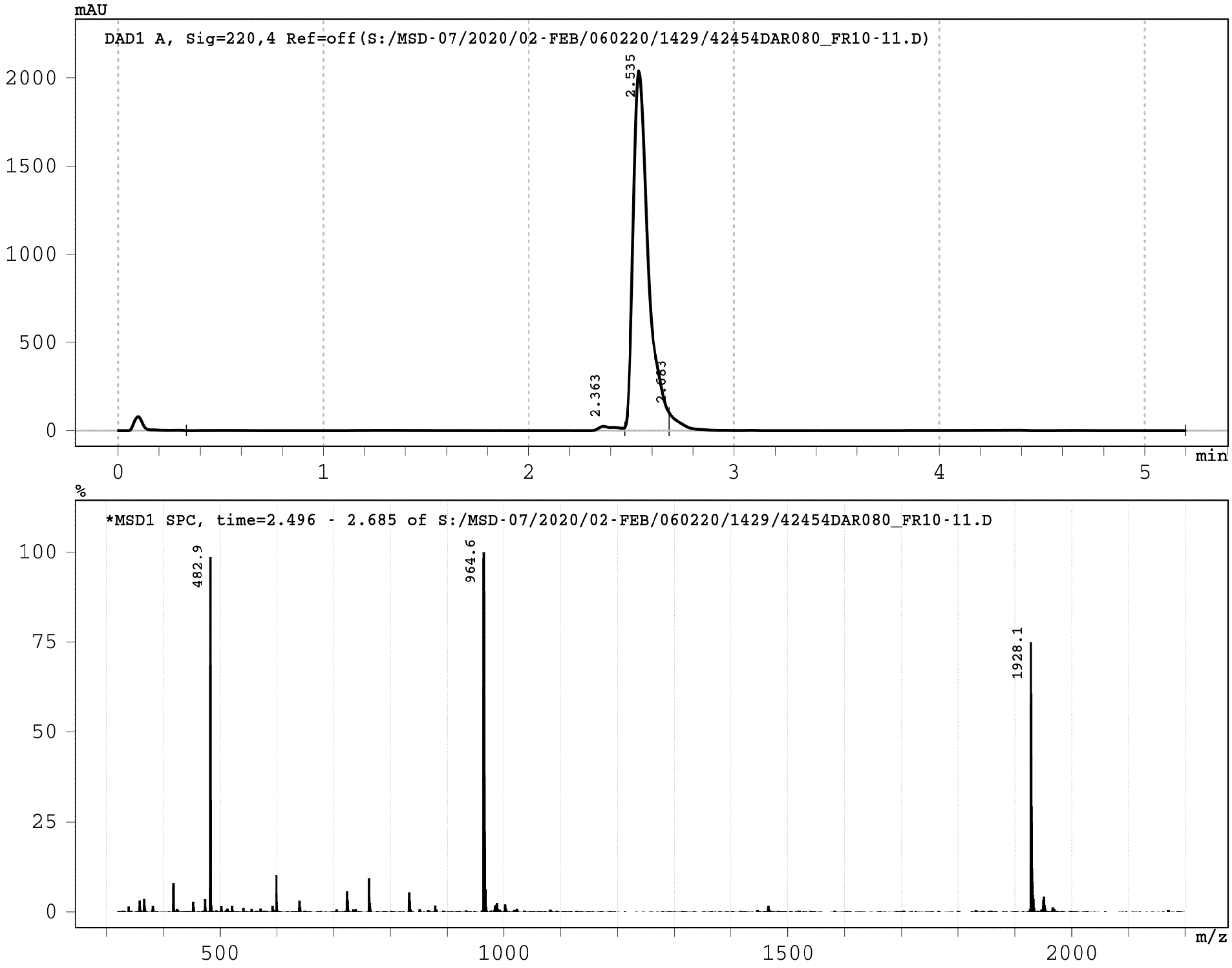


Signal 1: DAD1 A, Sig=220,4 Ref=off

| Peak # | RT [min] | Width [min] | Area | Area % |
|---|---|---|---|---|
| 1 | 2.363 | 2.13 | 184.7 | 1.7 |
| 2 | 2.535 | 0.22 | 10389.0 | 94.7 |
| 3 | 2.683 | 2.52 | 400.5 | 3.6 |

Data File: S:\MSD-07\2020\02-FEB\050220\142->

Injection Date : Thu, 6. Feb. 2020
Sample Name : 240120SG1-72|42454Dar081
Acq Operator : SYSTEM
Acq. Method : D:\Data\2020\02-FEB\050220\1428\5-95-6M.M
column used :
Analysis Method : C:\Chem32\1\METHODS\DEF_LC.M

Sample Info : 42454_081-Dr. Martine Darwish|H-LKYCHLLVL-OH

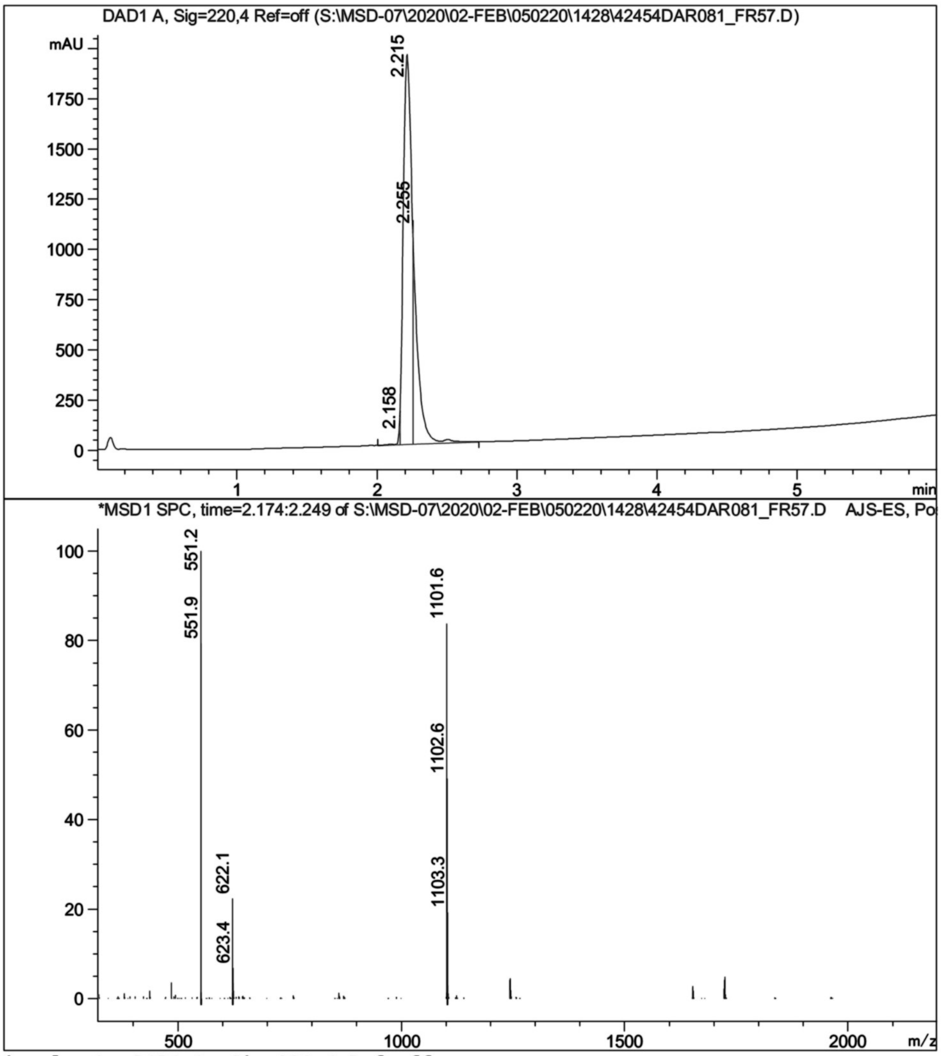


Signal 1: DAD1 A, Sig=220,4 Ref=off

| Peak # | RT [min] | Width [min] | Area | Area % |
|---|---|---|---|---|
| 1 | 2.158 | 0.0 | 105.4 | 1.0 |
| 2 | 2.215 | 0.1 | 7570.3 | 74.5 |
| 3 | 2.255 | 0.0 | 2480.5 | 24.4 |

*** End of Report ***

Data File : S:/MSD-07/2020/02-FEB/050220/1428/42454DAR082_FR8->

Injection Date : 06-Feb-20, 03:40:59
Sample Name : 42454DAR082_FR80-81
Remarks : Baseline corrected
Acq. Method : D:/METHODS/5-95-6M.M
column used : Je_559
Analysis Method : C:/CHEM32/1/METHODS/JPT-LCMS.M

Sample Info : 42454_082-Dr. Martine Darwish H-FAINMETCWI-OH

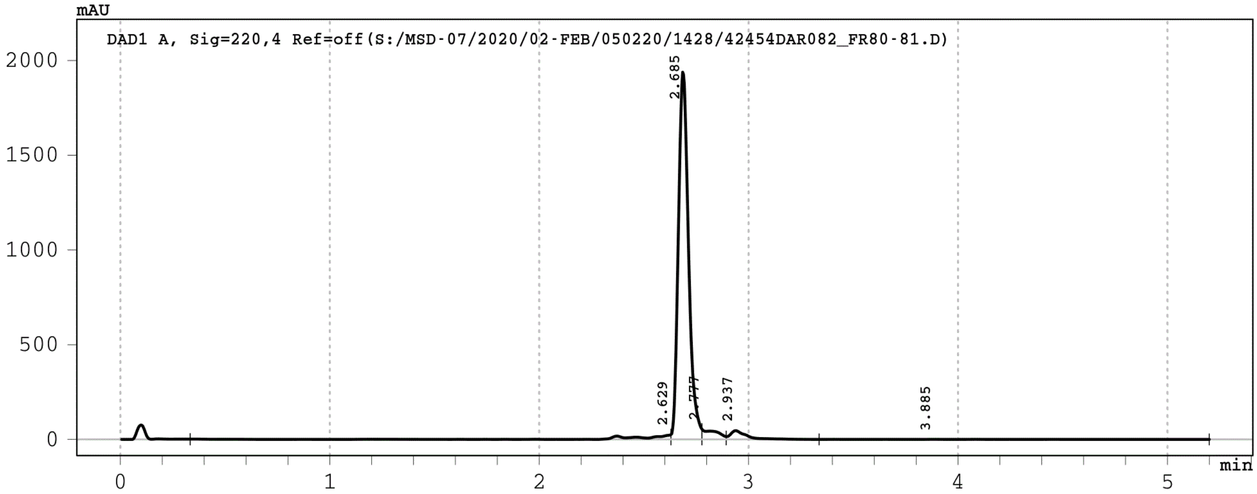


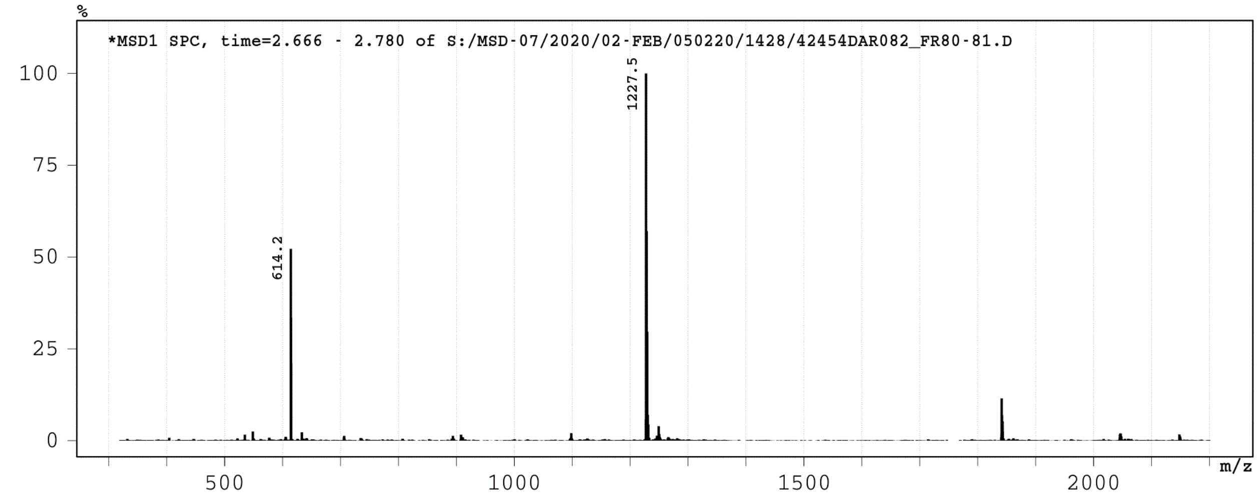


Signal 1: DAD1 A, Sig=220,4 Ref=off

| Peak # | RT [min] | Width [min] | Area | Area % |
|---|---|---|---|---|
| 1 | 2.629 | 2.30 | 254.6 | 3.5 |
| 2 | 2.685 | 0.15 | 6563.4 | 89.5 |
| 3 | 2.777 | 0.12 | 257.7 | 3.5 |
| 4 | 2.937 | 0.44 | 253.4 | 3.5 |
| 5 | 3.885 | 1.86 | 4.5 | 0.1 |

Data File : S:/MSD-07/2020/02-FEB/050220/1428/42454DAR083_FR1->

Injection Date : 06-Feb-20, 03:49:22
Sample Name : 42454DAR083_FR100+1-2
Remarks : Baseline corrected
Acq. Method : D:/METHODS/5-95-6M.M
column used : Je_559
Analysis Method : C:/CHEM32/1/METHODS/JPT-LCMS.M

Sample Info : 42454_083-Dr. Martine Darwish H-TSLECLSNL-OH

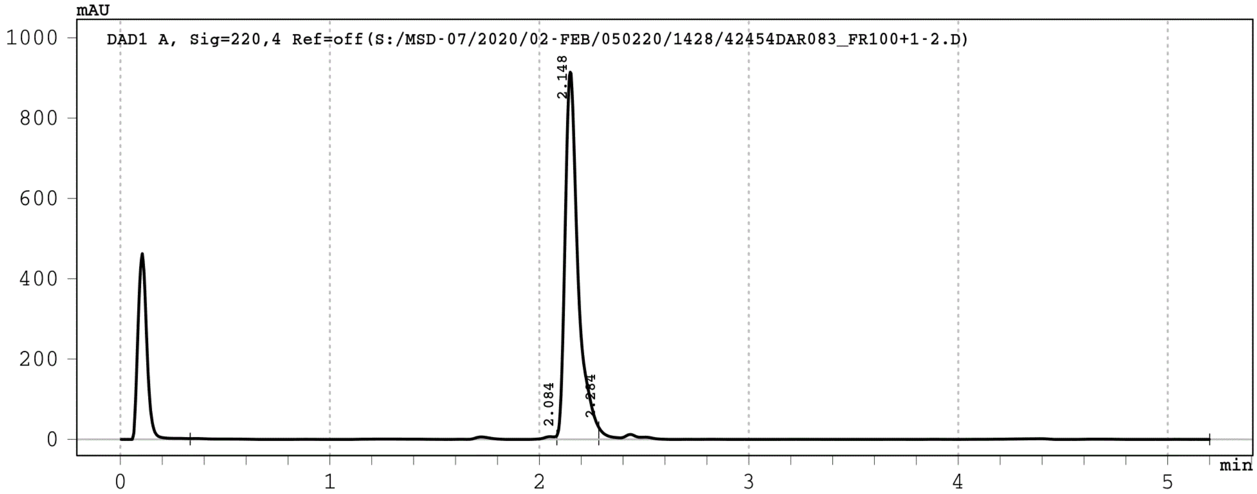


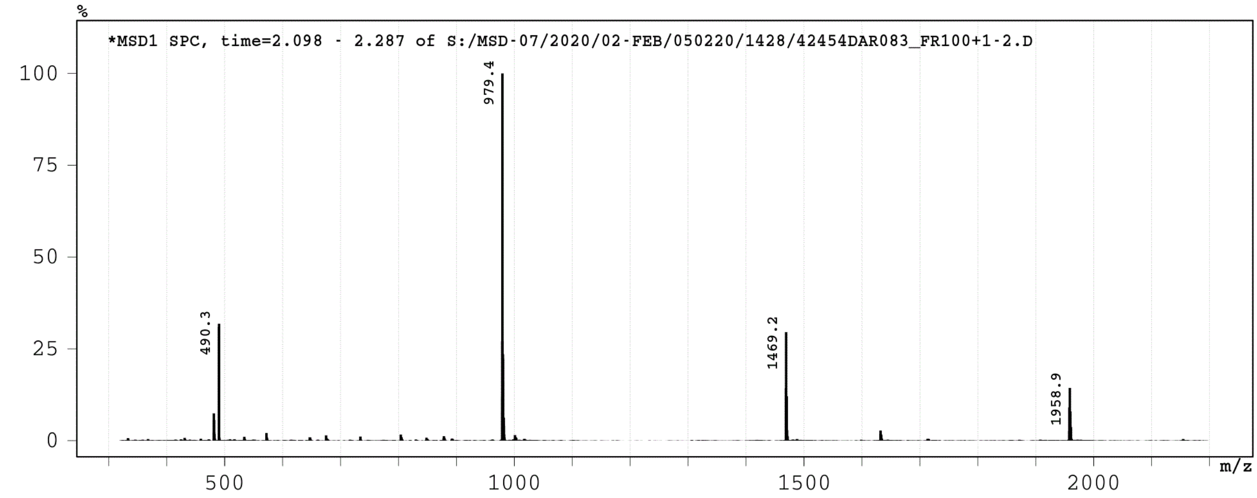


Signal 1: DAD1 A, Sig=220,4 Ref=off

| Peak # | RT [min] | Width [min] | Area | Area % |
|---|---|---|---|---|
| 1 | 2.084 | 1.75 | 92.8 | 2.2 |
| 2 | 2.148 | 0.20 | 3944.1 | 93.9 |
| 3 | 2.284 | 2.92 | 164.8 | 3.9 |

Data File: S:\MSD-07\2020\02-FEB\050220\142->

Injection Date : Thu, 6. Feb. 2020
Sample Name : 240120SG1-74|42454Dar084
Acq Operator : SYSTEM
Acq. Method : D:\Data\2020\02-FEB\050220\1428\5-95-6MM
column used :
Analysis Method : C:\Chem32\1\METHODS\DEF_LC.M

Sample Info : 42454_084-Dr. Martine Darwish|H-CTFSHLTKL-OH

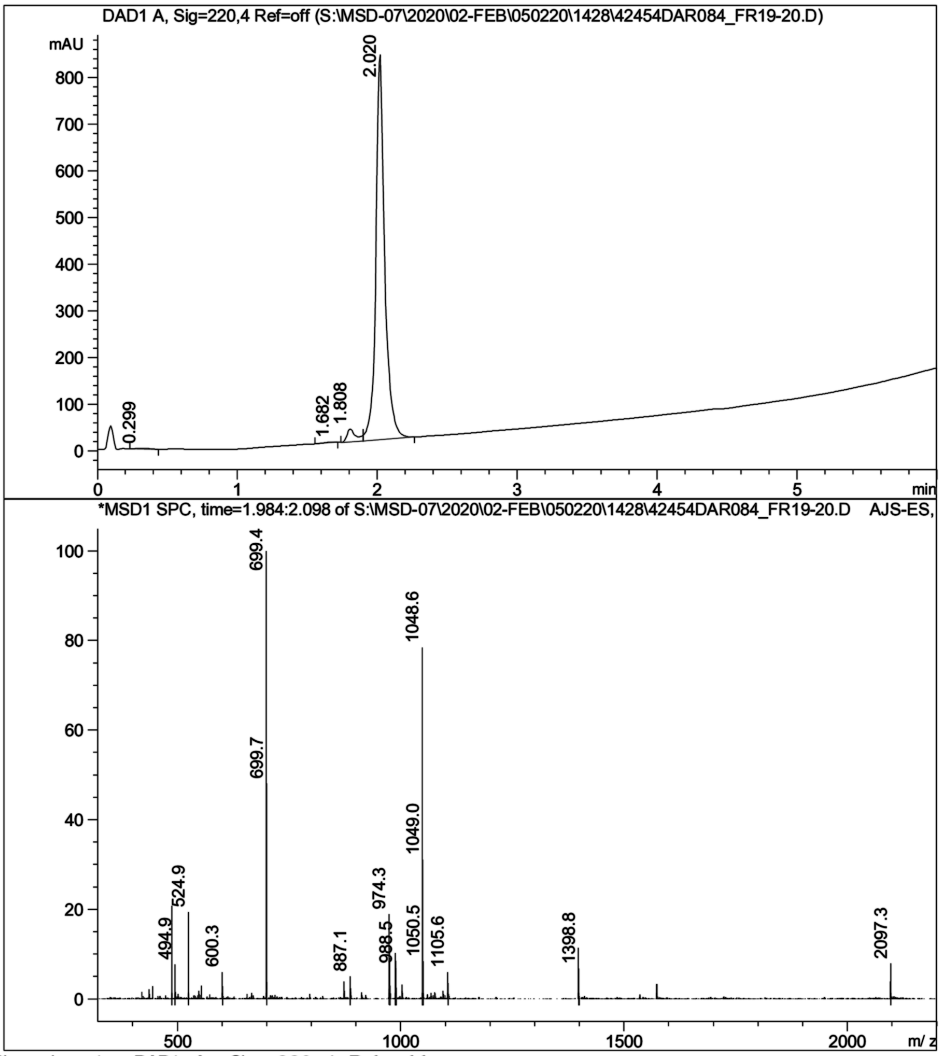


Signal 1: DAD1 A, Sig=220,4 Ref=off

| Peak # | RT [min] | Width [min] | Area | Area % |
|---|---|---|---|---|
| 1 | 0.299 | 0.1 | 8.1 | 0.2 |
| 2 | 1.682 | 0.1 | 6.5 | 0.2 |
| 3 | 1.808 | 0.1 | 125.8 | 3.4 |
| 4 | 2.020 | 0.1 | 3612.3 | 96.3 |

*** End of Report ***

Data File : S:/MSD-07/2020/02-FEB/060220/1429/42454DAR085_FR2->

Injection Date : 06-Feb-20, 14:59:48
Sample Name : 42454DAR085_FR24-25
Remarks : Baseline corrected
Acq. Method : D:/METHODS/5-95-6M.M
column used : Je_559
Analysis Method : C:/CHEM32/1/METHODS/JPT-LCMS.M

Sample Info : 42454_085-Dr. Martine Darwish H-SIYSSRRFHV-OH

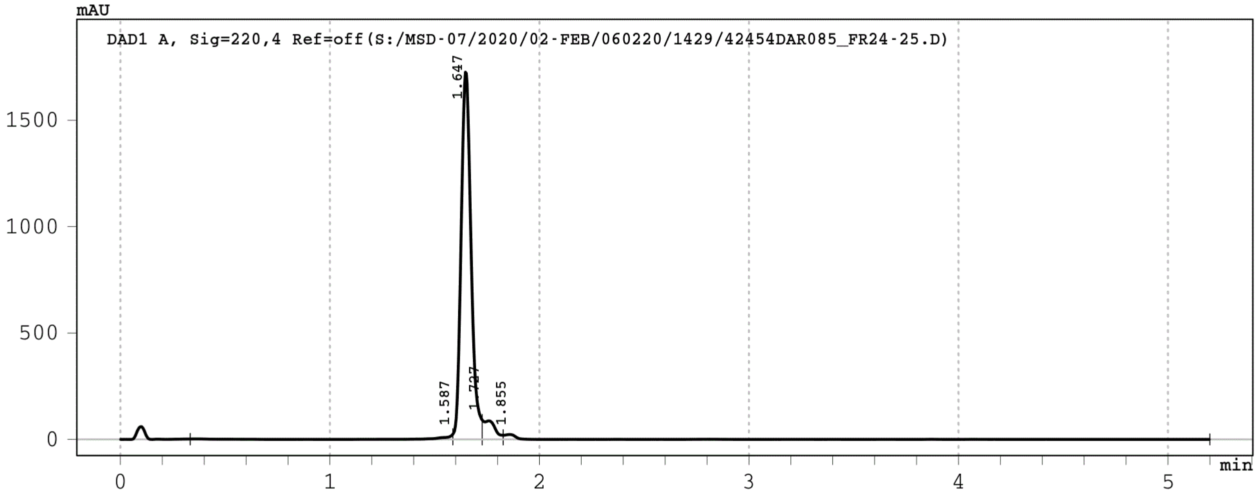


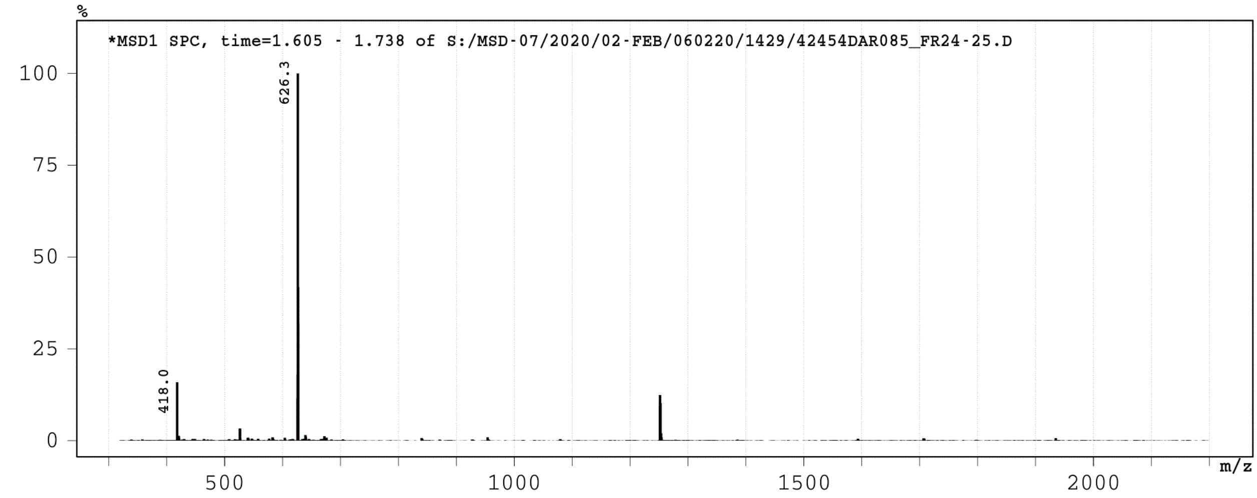


Signal 1: DAD1 A, Sig=220,4 Ref=off

| Peak # | RT [min] | Width [min] | Area | Area % |
|---|---|---|---|---|
| 1 | 1.587 | 1.25 | 93.9 | 1.5 |
| 2 | 1.647 | 0.14 | 5693.9 | 91.1 |
| 3 | 1.727 | 0.10 | 361.7 | 5.8 |
| 4 | 1.855 | 3.37 | 97.6 | 1.6 |

| | | |
|---|---|---|
| Data File | : | S:/MSD-07/2020/02-FEB/060220/1429/42454DAR086_FR4-> |
| Injection Date | : | 06-Feb-20, 15:08:11 |
| Sample Name | : | 42454DAR086_FR41-42 |
| Remarks | : | Baseline corrected |
| Acq. Method | : | D:/METHODS/5-95-6M.M |
| column used | : | Je_559 |
| Analysis Method | : | C:/CHEM32/1/METHODS/JPT-LCMS.M |
| Sample Info | : | 42454_086-Dr. Martine Darwish H-ISYLKGNATI-OH |

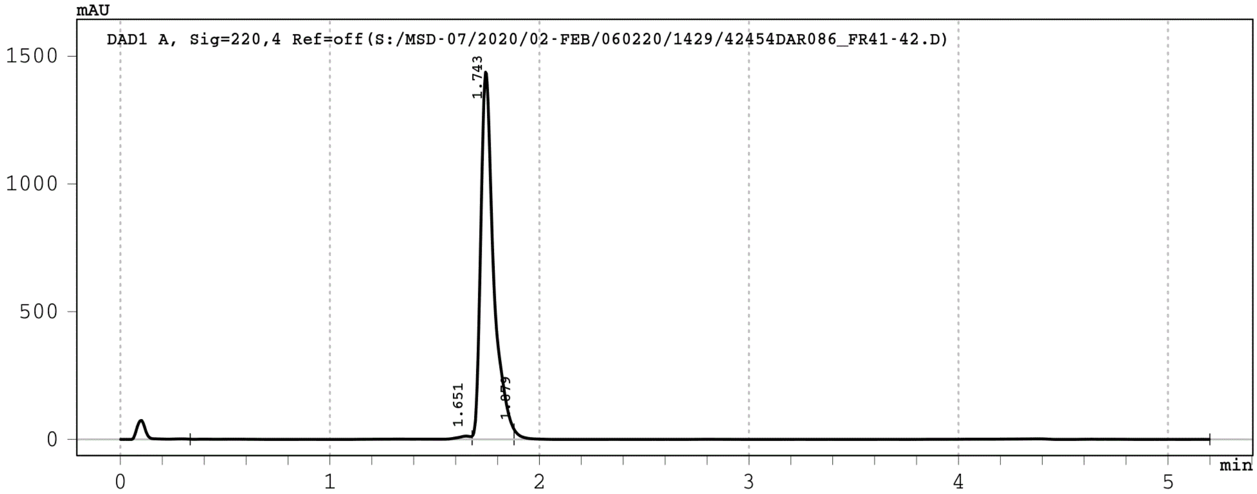


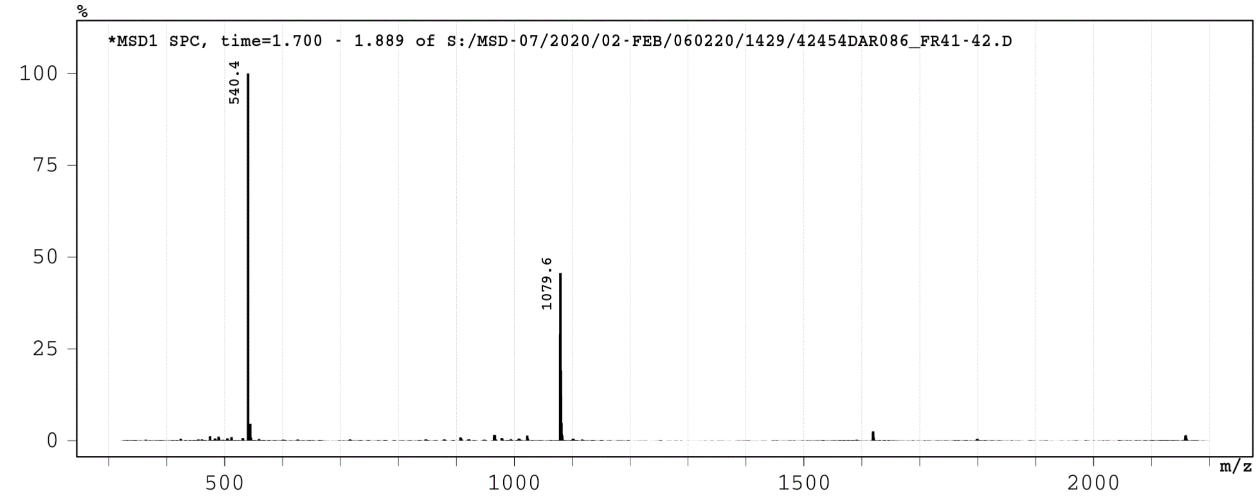


Signal 1: DAD1 A, Sig=220,4 Ref=off

| Peak # | RT [min] | Width [min] | Area | Area % |
|---|---|---|---|---|
| 1 | 1.651 | 1.35 | 94.5 | 1.5 |
| 2 | 1.743 | 0.20 | 6153.8 | 96.6 |
| 3 | 1.879 | 3.32 | 123.6 | 1.9 |

Data File : S:/MSD-07/2020/02-FEB/060220/1429/42454DAR087_FR5->

Injection Date : 06-Feb-20, 15:16:34
Sample Name : 42454DAR087_FR51-52
Remarks : Baseline corrected
Acq. Method : D:/METHODS/5-95-6M.M
column used : Je_559
Analysis Method : C:/CHEM32/1/METHODS/JPT-LCMS.M

Sample Info : 42454_087-Dr. Martine Darwish H-SHYRAYIL-OH

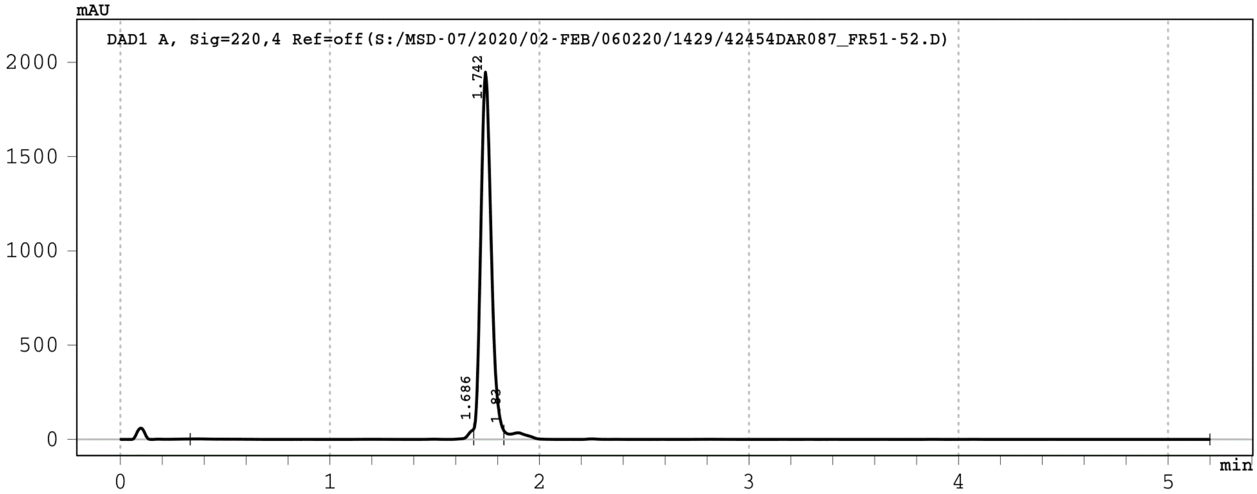


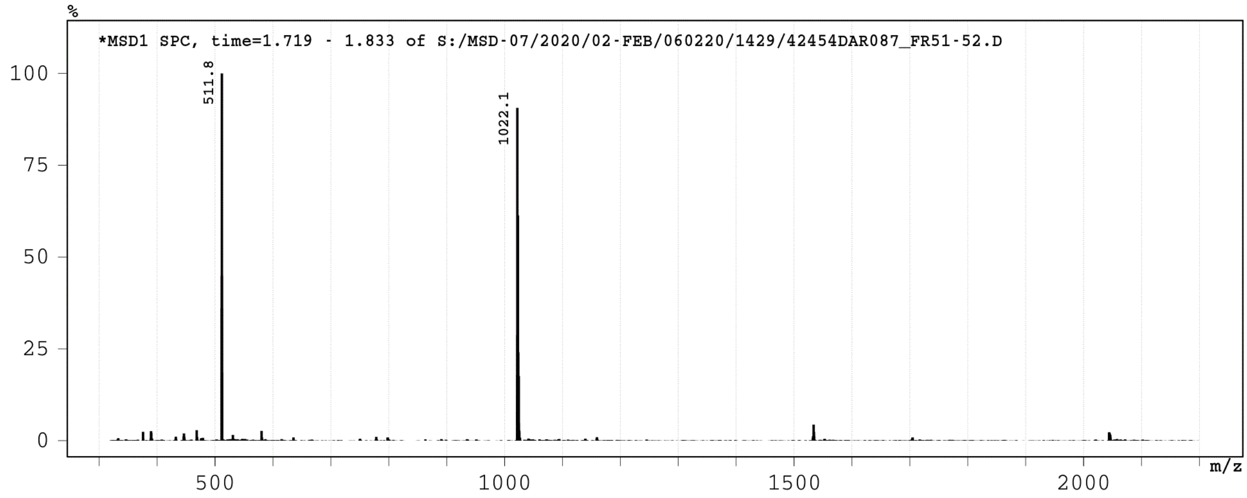


Signal 1: DAD1 A, Sig=220,4 Ref=off

| Peak # | RT [min] | Width [min] | Area | Area % |
|---|---|---|---|---|
| 1 | 1.686 | 1.35 | 126.3 | 1.8 |
| 2 | 1.742 | 0.14 | 6739.1 | 94.5 |
| 3 | 1.830 | 3.37 | 265.2 | 3.7 |

Data File : S:/MSD-07/2020/02-FEB/060220/1429/42454DAR088_FR7->

Injection Date : 06-Feb-20, 15:33:58
Sample Name : 42454DAR088_FR72-73
Remarks : Baseline corrected
Acq. Method : D:/METHODS/5-95-6M.M
column used : Je_559
Analysis Method : C:/CHEM32/1/METHODS/JPT-LCMS.M

Sample Info : 42454_088-Dr. Martine Darwish H-VFFPRHREL-OH

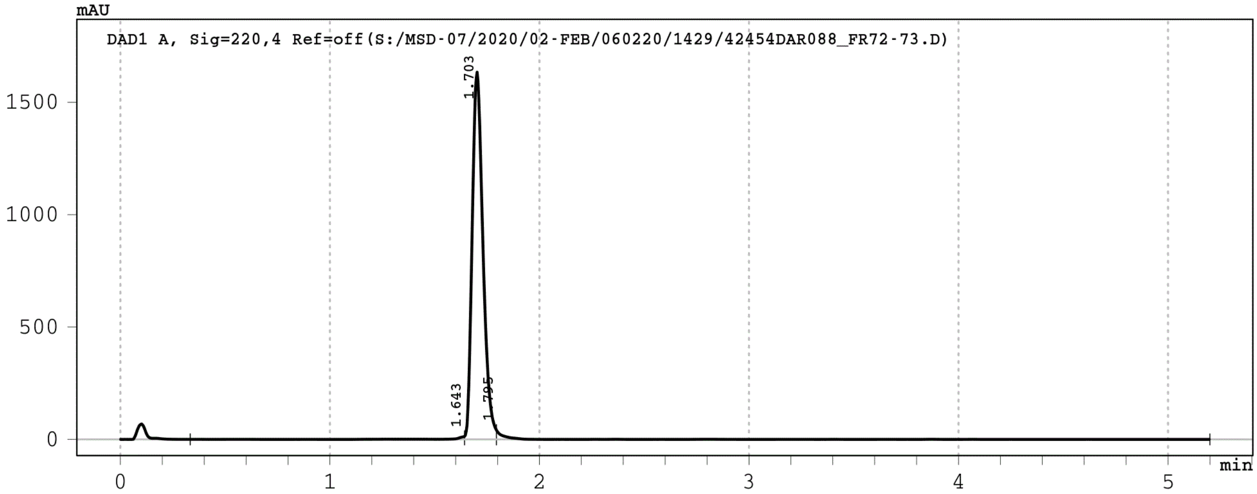


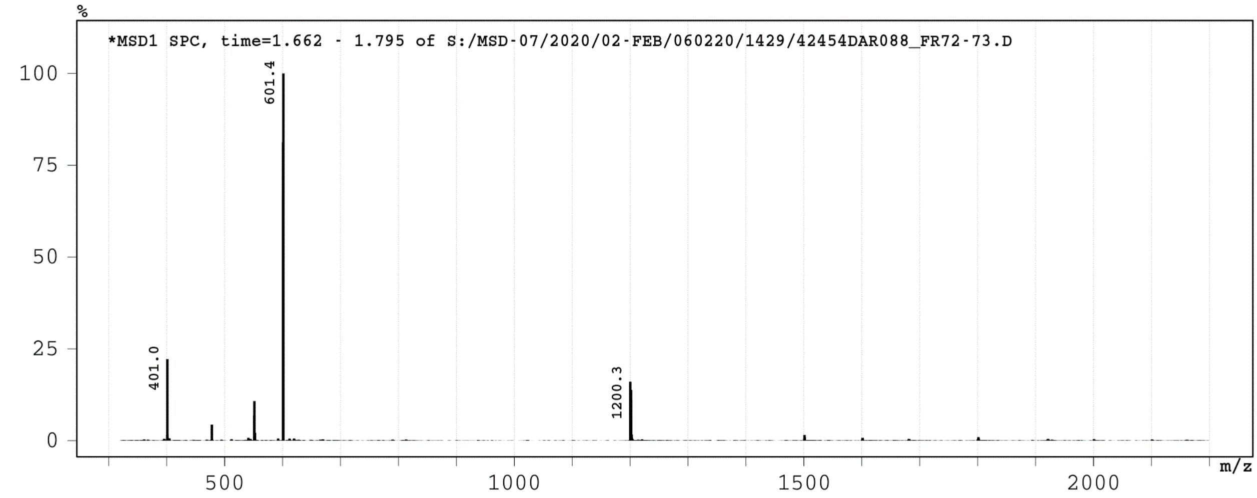


Signal 1: DAD1 A, Sig=220,4 Ref=off

| Peak # | RT [min] | Width [min] | Area | Area % |
|---|---|---|---|---|
| 1 | 1.643 | 1.31 | 47.6 | 0.8 |
| 2 | 1.703 | 0.15 | 5855.9 | 97.5 |
| 3 | 1.795 | 3.41 | 104.2 | 1.7 |

Data File : S:/MSD-07/2020/02-FEB/050220/1428/42454DAR089_FR3->

Injection Date : 06-Feb-20, 04:14:36
Sample Name : 42454DAR089_FR38-39
Remarks : Baseline corrected
Acq. Method : D:/METHODS/5-95-6M.M
column used : Je_559
Analysis Method : C:/CHEM32/1/METHODS/JPT-LCMS.M

Sample Info : 42454_089-Dr. Martine Darwish H-MTYLHNLEM-OH

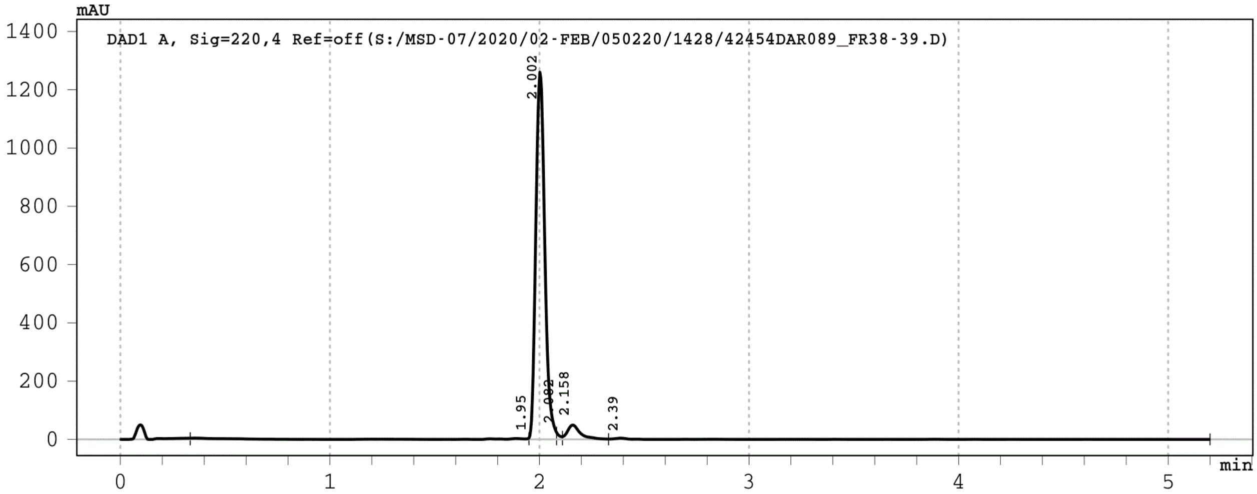


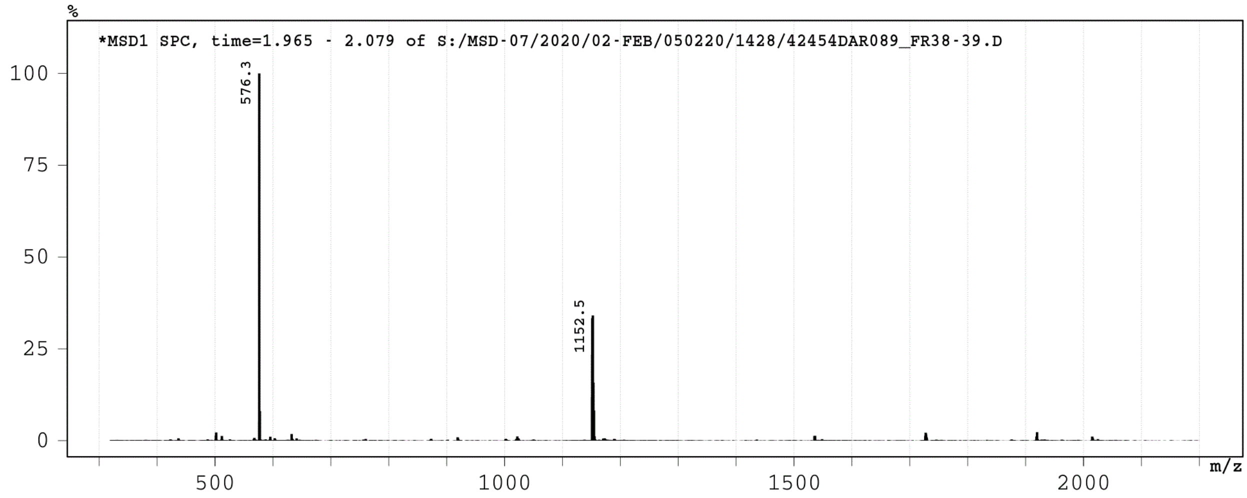


Signal 1: DAD1 A, Sig=220,4 Ref=off

| Peak # | RT [min] | Width [min] | Area | Area % |
|---|---|---|---|---|
| 1 | 1.950 | 1.62 | 93.1 | 2.3 |
| 2 | 2.002 | 0.13 | 3754.4 | 91.2 |
| 3 | 2.082 | 0.03 | 22.5 | 0.5 |
| 4 | 2.158 | 0.22 | 217.7 | 5.3 |
| 5 | 2.390 | 2.87 | 30.4 | 0.7 |

Data File : S:/MSD-07/2020/02-FEB/050220/1428/42454DAR090_FR4->

Injection Date : 06-Feb-20, 04:23:01
Sample Name : 42454DAR090_FR45-46
Remarks : Baseline corrected
Acq. Method : D:/METHODS/5-95-6M.M
column used : Je_559
Analysis Method : C:/CHEM32/1/METHODS/JPT-LCMS.M

Sample Info : 42454_090-Dr. Martine Darwish H-IVLEHMNI-OH

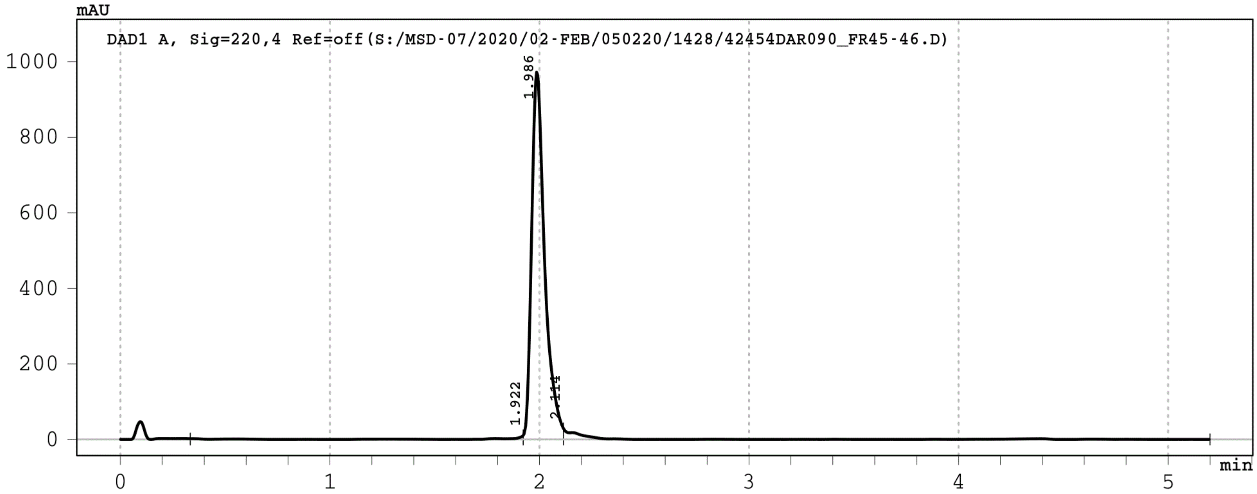


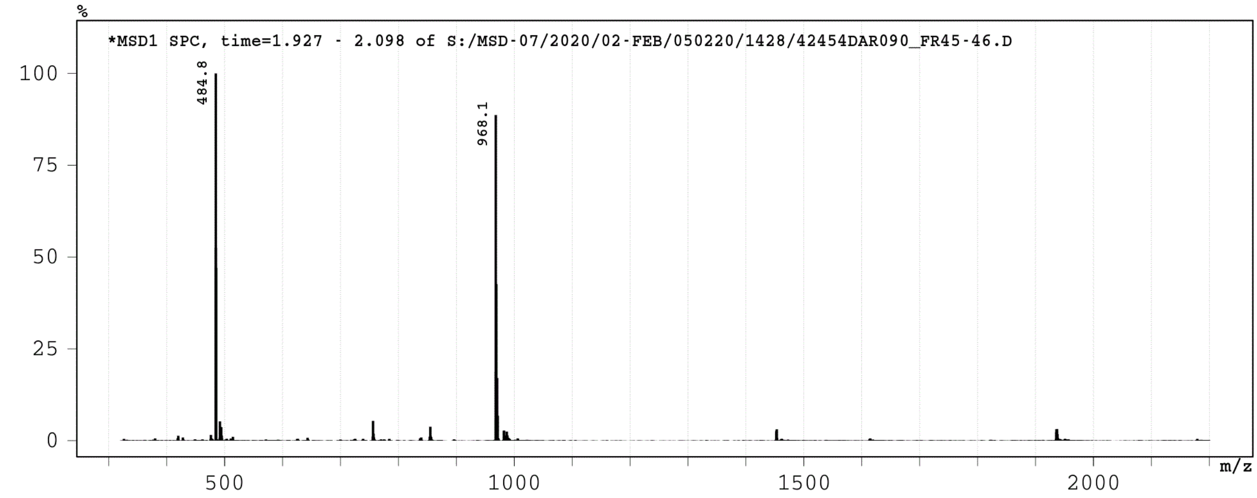


Signal 1: DAD1 A, Sig=220,4 Ref=off

| Peak # | RT [min] | Width [min] | Area | Area % |
|---|---|---|---|---|
| 1 | 1.922 | 1.59 | 60.4 | 1.4 |
| 2 | 1.986 | 0.19 | 4225.9 | 94.7 |
| 3 | 2.114 | 3.09 | 174.1 | 3.9 |

Data File : S:/MSD-07/2020/02-FEB/050220/1428/42454DAR091_FR1->

Injection Date : 05-Feb-20, 18:16:57
Sample Name : 42454DAR091_FR12
Remarks : Baseline corrected
Acq. Method : D:/METHODS/5-95-6M.M
column used : Je_559
Analysis Method : C:/CHEM32/1/METHODS/JPT-LCMS.M

Sample Info : 42454_091-Dr. Martine Darwish H-LSFAQRTLYM-OH

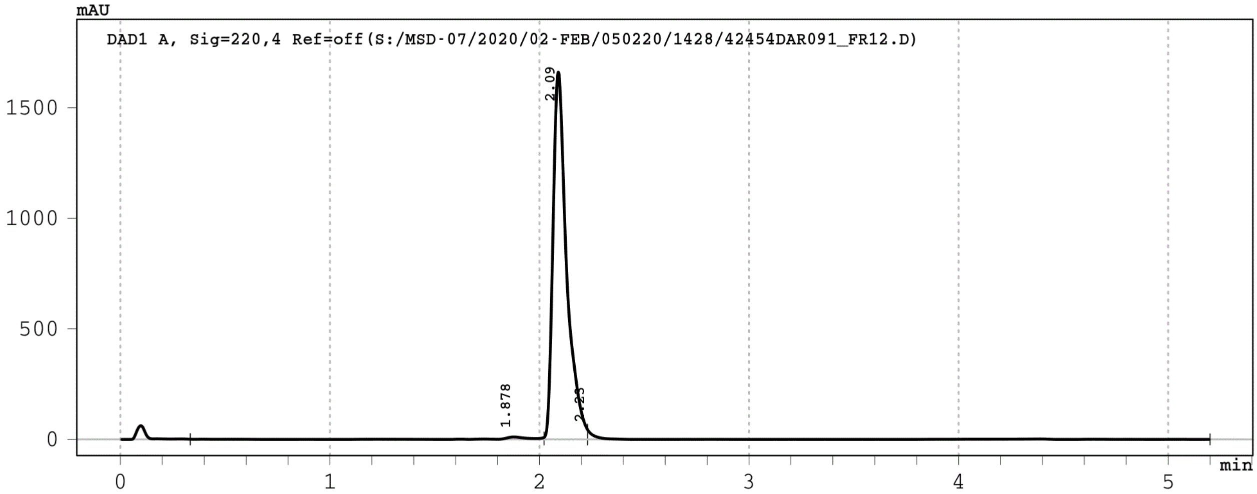


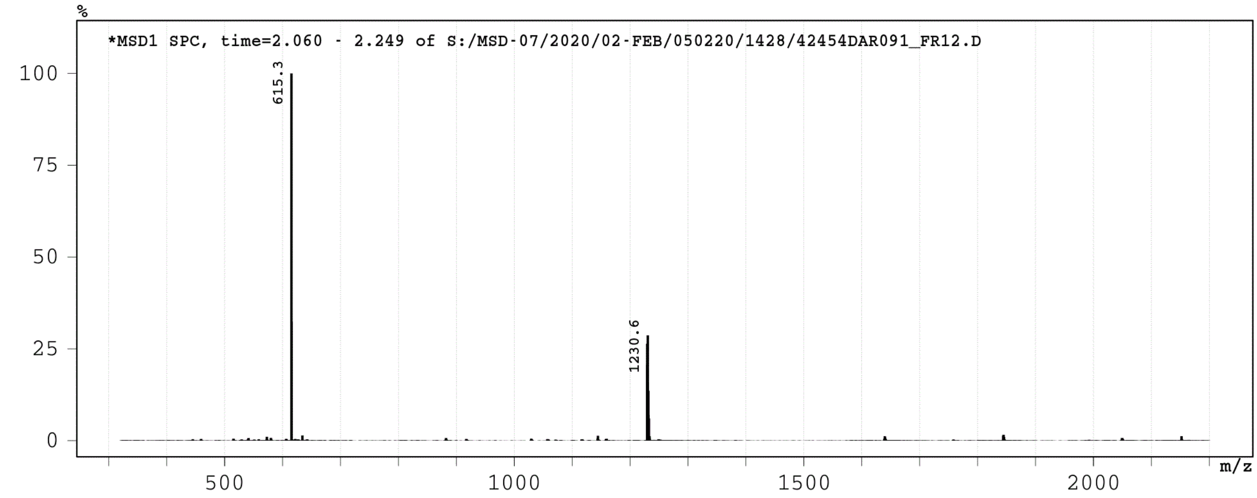


Signal 1: DAD1 A, Sig=220,4 Ref=off

| Peak # | RT [min] | Width [min] | Area | Area % |
|---|---|---|---|---|
| 1 | 1.878 | 1.69 | 114.6 | 1.5 |
| 2 | 2.090 | 0.21 | 7561.6 | 96.8 |
| 3 | 2.230 | 2.97 | 138.3 | 1.8 |

Data File : S:/MSD-07/2020/02-FEB/050220/1428/42454DAR092_FR2->

Injection Date : 05-Feb-20, 18:25:21
Sample Name : 42454DAR092_FR27-28
Remarks : Baseline corrected
Acq. Method : D:/METHODS/5-95-6M.M
column used : Je_559
Analysis Method : C:/CHEM32/1/METHODS/JPT-LCMS.M

Sample Info : 42454_092-Dr. Martine Darwish H-MNIAHLHL-OH

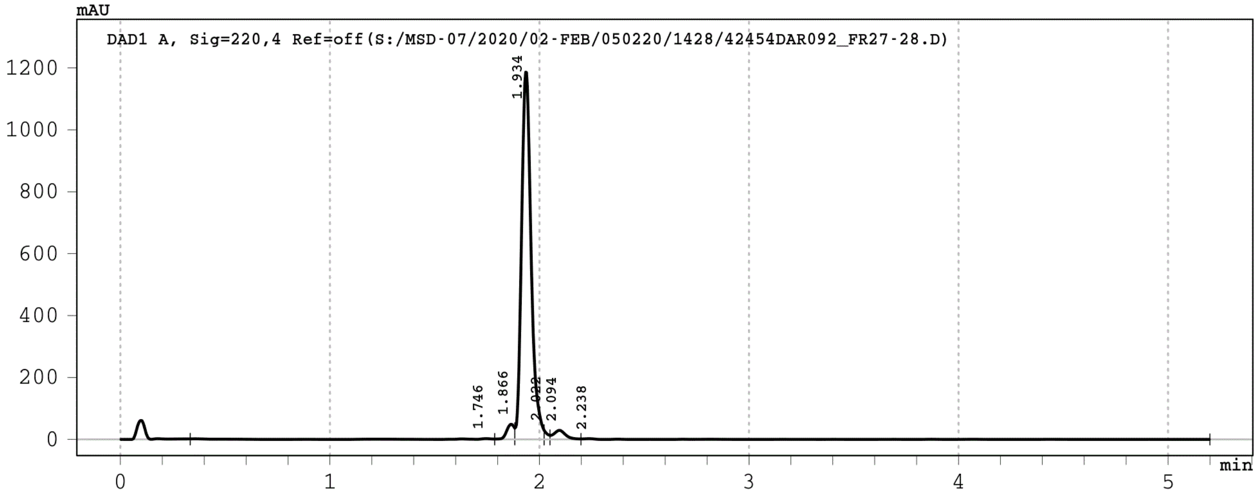


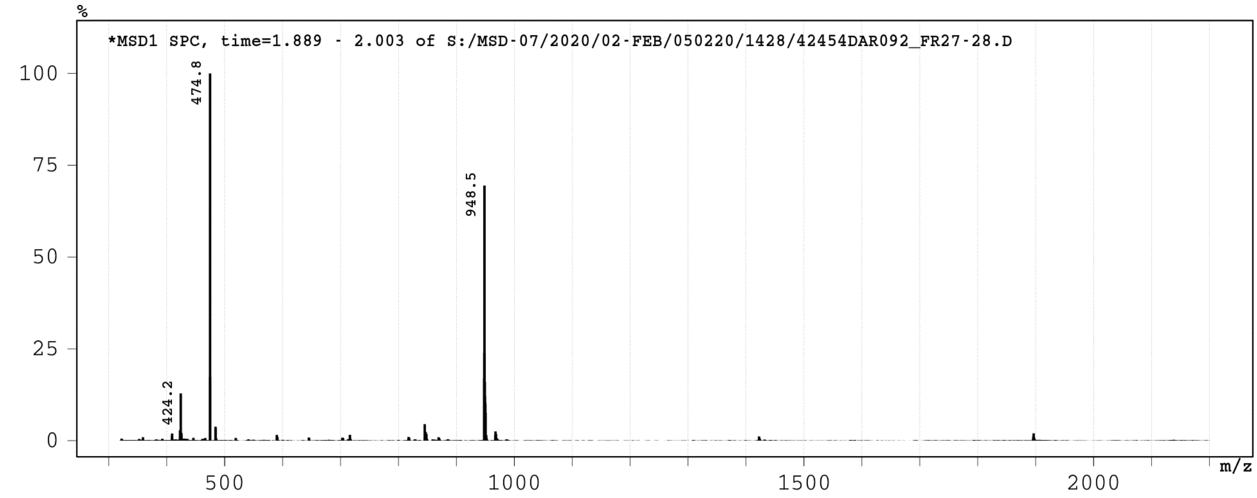


Signal 1: DAD1 A, Sig=220,4 Ref=off

| Peak # | RT [min] | Width [min] | Area | Area % |
|---|---|---|---|---|
| 1 | 1.746 | 1.45 | 44.4 | 1.1 |
| 2 | 1.866 | 0.10 | 119.8 | 2.9 |
| 3 | 1.934 | 0.14 | 3786.9 | 91.8 |
| 4 | 2.022 | 0.03 | 32.0 | 0.8 |
| 5 | 2.094 | 0.15 | 118.5 | 2.9 |
| 6 | 2.238 | 3.00 | 21.8 | 0.5 |

Data File : S:/MSD-07/2020/02-FEB/050220/1428/42454DAR093_FR3->

Injection Date : 05-Feb-20, 18:33:44
Sample Name : 42454DAR093_FR35-36
Remarks : Baseline corrected
Acq. Method : D:/METHODS/5-95-6M.M
column used : Je_559
Analysis Method : C:/CHEM32/1/METHODS/JPT-LCMS.M

Sample Info : 42454_093-Dr. Martine Darwish H-CAYSISSGI-OH

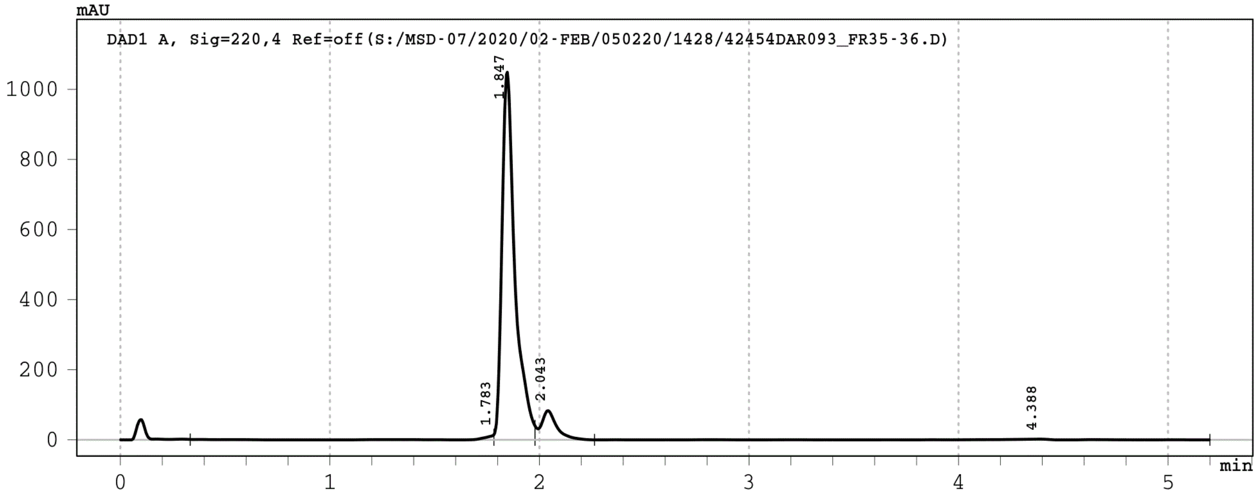


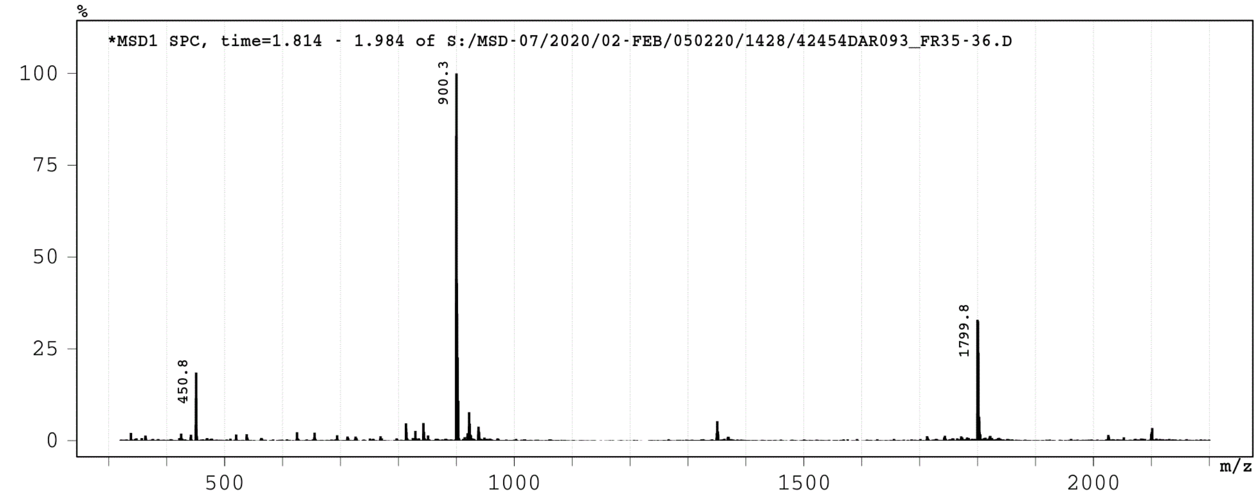


Signal 1: DAD1 A, Sig=220,4 Ref=off

| Peak # | RT [min] | Width [min] | Area | Area % |
|---|---|---|---|---|
| 1 | 1.783 | 1.45 | 62.4 | 1.2 |
| 2 | 1.847 | 0.20 | 4551.2 | 89.3 |
| 3 | 2.043 | 0.28 | 443.0 | 8.7 |
| 4 | 4.388 | 2.94 | 42.1 | 0.8 |

Data File : S:/MSD-07/2020/02-FEB/050220/1428/42454DAR094_FR5->

Injection Date : 05-Feb-20, 18:42:10
Sample Name : 42454DAR094_FR53
Remarks : Baseline corrected
Acq. Method : D:/METHODS/5-95-6M.M
column used : Je_559
Analysis Method : C:/CHEM32/1/METHODS/JPT-LCMS.M

Sample Info : 42454_094-Dr. Martine Darwish H-DVYPFHMIL-OH

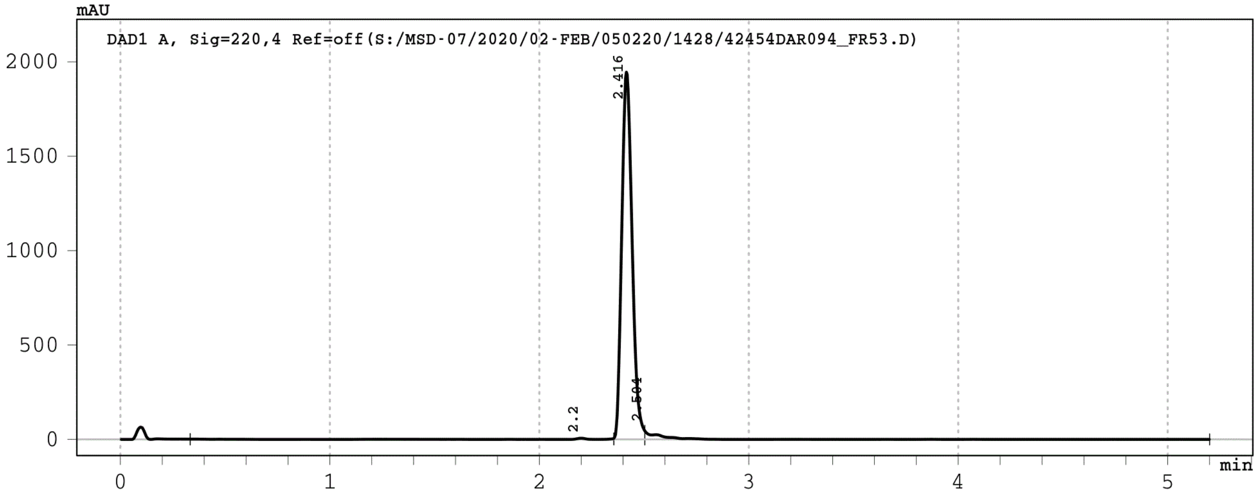


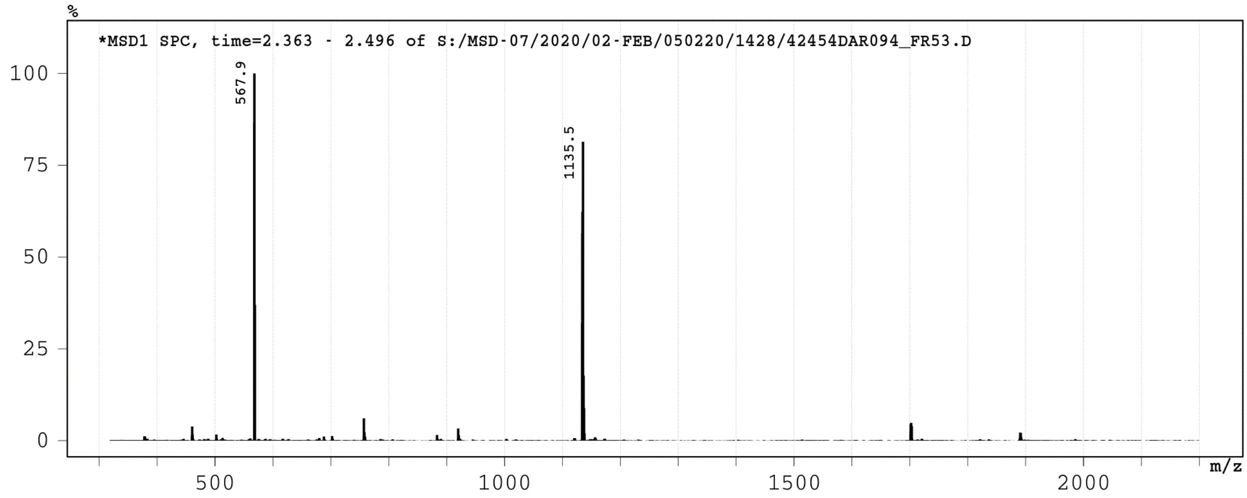


Signal 1: DAD1 A, Sig=220,4 Ref=off

| Peak # | RT [min] | Width [min] | Area | Area % |
|---|---|---|---|---|
| 1 | 2.200 | 2.02 | 54.2 | 0.8 |
| 2 | 2.416 | 0.15 | 6700.8 | 96.1 |
| 3 | 2.504 | 2.70 | 214.7 | 3.1 |

Data File : S:/MSD-07/2020/02-FEB/180220/1438/42454DAR095_FR1->

Injection Date : 18-Feb-20, 09:49:42
Sample Name : 42454DAR095_FR17
Remarks : Baseline corrected
Acq. Method : D:/METHODS/5-95-6M.M
column used : Je_559
Analysis Method : C:/CHEM32/1/METHODS/JPT-LCMS.M

Sample Info : 42454_095-Dr. Martine Darwish H-QIYAFLQGF-OH

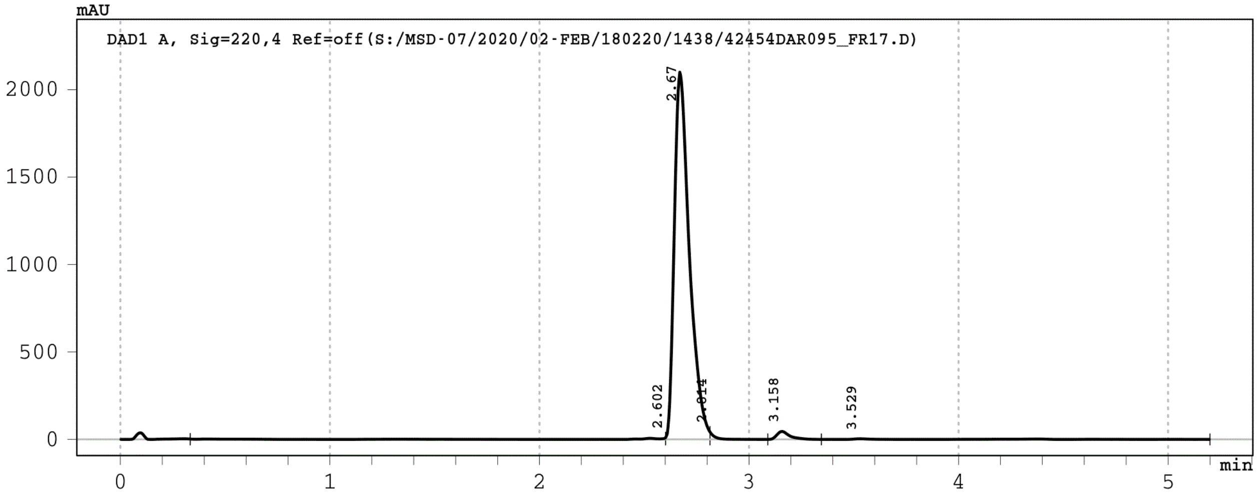


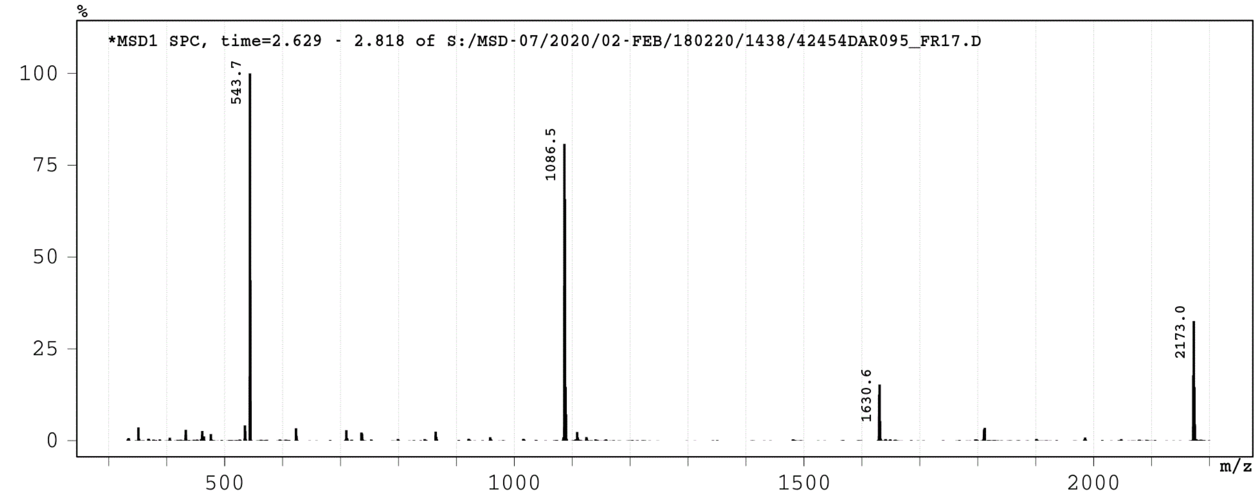


Signal 1: DAD1 A, Sig=220,4 Ref=off

| Peak # | RT [min] | Width [min] | Area | Area % |
|---|---|---|---|---|
| 1 | 2.602 | 2.27 | 101.7 | 0.9 |
| 2 | 2.670 | 0.21 | 10521.8 | 96.2 |
| 3 | 2.814 | 0.28 | 60.1 | 0.5 |
| 4 | 3.158 | 0.26 | 195.7 | 1.8 |
| 5 | 3.529 | 1.85 | 60.0 | 0.5 |

Data File : S:/MSD-07/2020/02-FEB/060220/1429/42454DAR096_FR2->

Injection Date : 06-Feb-20, 15:42:21
Sample Name : 42454DAR096_FR2-3
Remarks : Baseline corrected
Acq. Method : D:/METHODS/5-95-6M.M
column used : Je_559
Analysis Method : C:/CHEM32/1/METHODS/JPT-LCMS.M

Sample Info : 42454_096-Dr. Martine Darwish H-PSLLNWTRV-OH

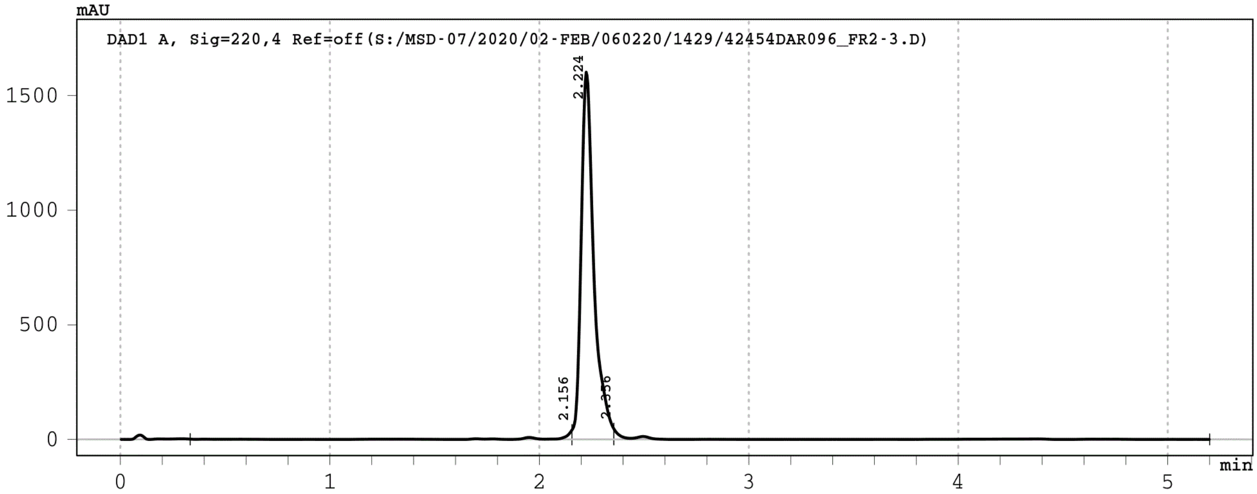


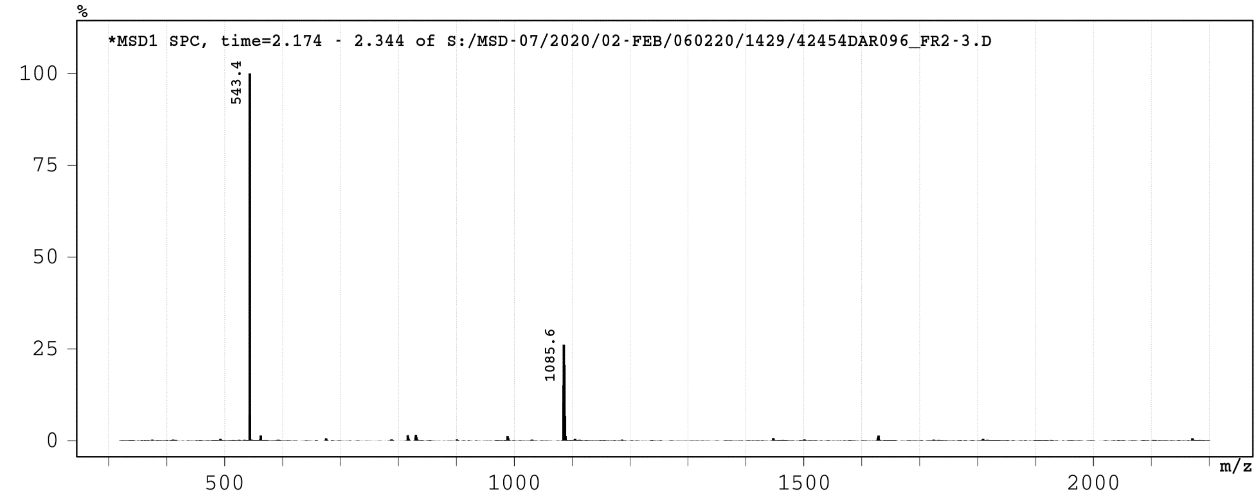


Signal 1: DAD1 A, Sig=220,4 Ref=off

| Peak # | RT [min] | Width [min] | Area | Area % |
|---|---|---|---|---|
| 1 | 2.156 | 1.82 | 142.8 | 2.0 |
| 2 | 2.224 | 0.20 | 6782.2 | 95.3 |
| 3 | 2.356 | 2.84 | 193.8 | 2.7 |

Data File : S:/MSD-07/2020/02-FEB/060220/1429/42454DAR097_FR3->

Injection Date : 06-Feb-20, 15:50:41
Sample Name : 42454DAR097_FR35-36
Remarks : Baseline corrected
Acq. Method : D:/METHODS/5-95-6M.M
column used : Je_559
Analysis Method : C:/CHEM32/1/METHODS/JPT-LCMS.M

Sample Info : 42454_097-Dr. Martine Darwish H-KGYPHWPAL-OH

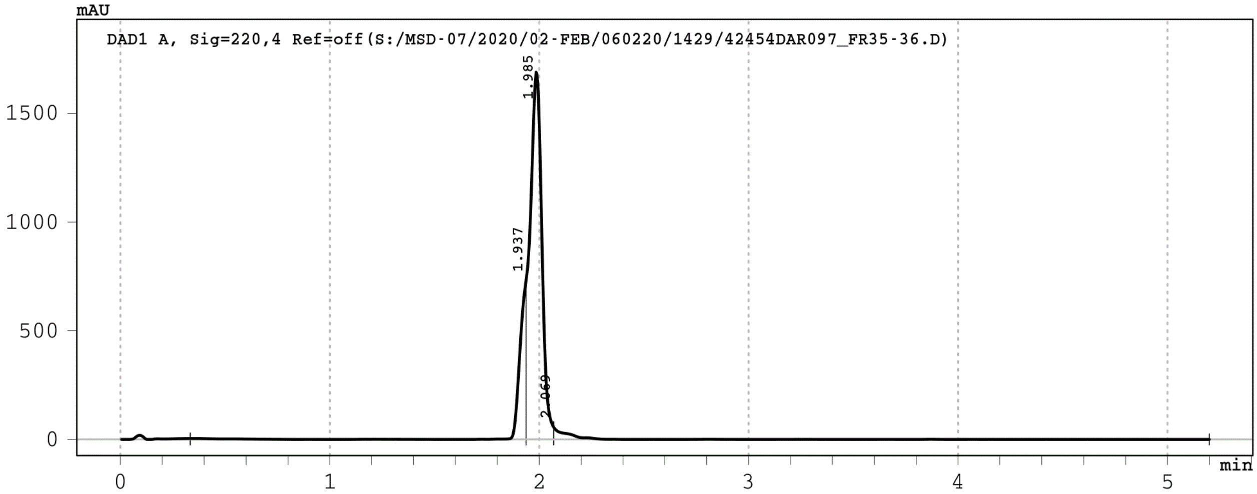


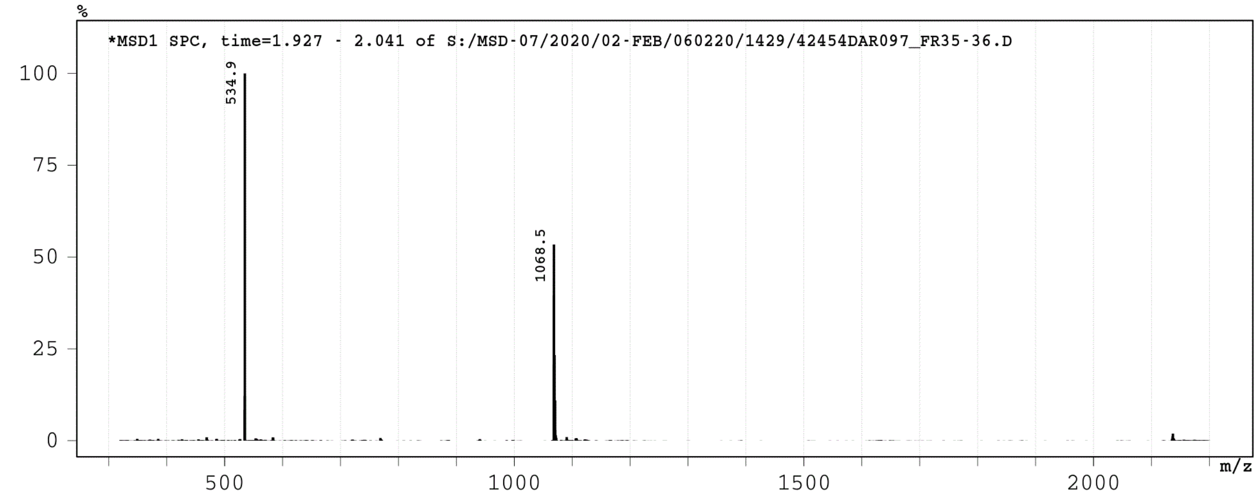


Signal 1: DAD1 A, Sig=220,4 Ref=off

| Peak # | RT [min] | Width [min] | Area | Area % |
|---|---|---|---|---|
| 1 | 1.937 | 1.60 | 1482.1 | 17.7 |
| 2 | 1.985 | 0.13 | 6633.3 | 79.3 |
| 3 | 2.069 | 3.13 | 247.9 | 3.0 |

Data File : S:/MSD-07/2020/02-FEB/050220/1428/42454DAR098_FR6->

Injection Date : 05-Feb-20, 18:50:35
Sample Name : 42454DAR098_FR63-64
Remarks : Baseline corrected
Acq. Method : D:/METHODS/5-95-6M.M
column used : Je_559
Analysis Method : C:/CHEM32/1/METHODS/JPT-LCMS.M

Sample Info : 42454_098-Dr. Martine Darwish H-CVYEHTAVL-OH

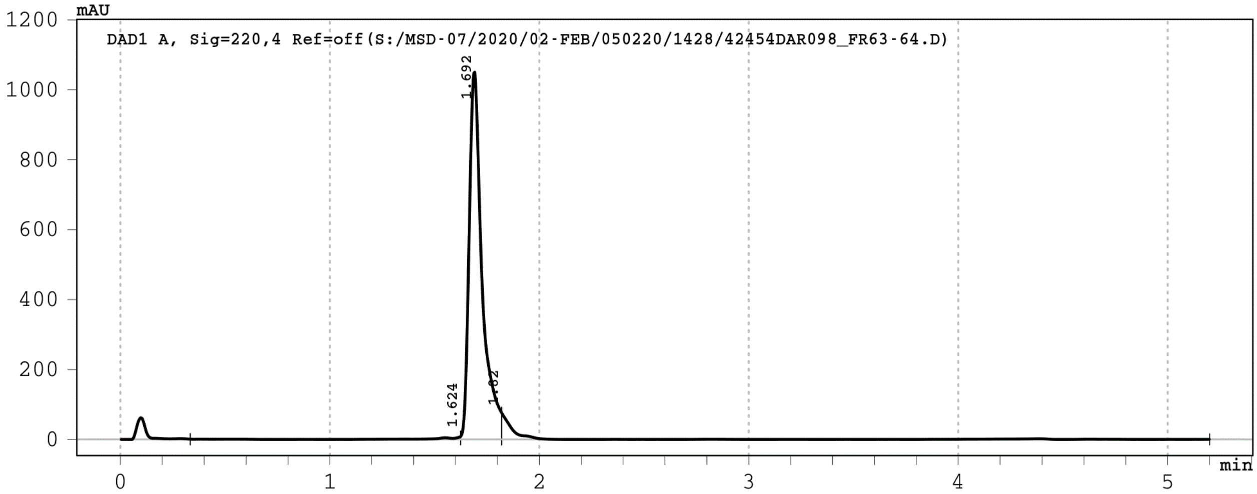


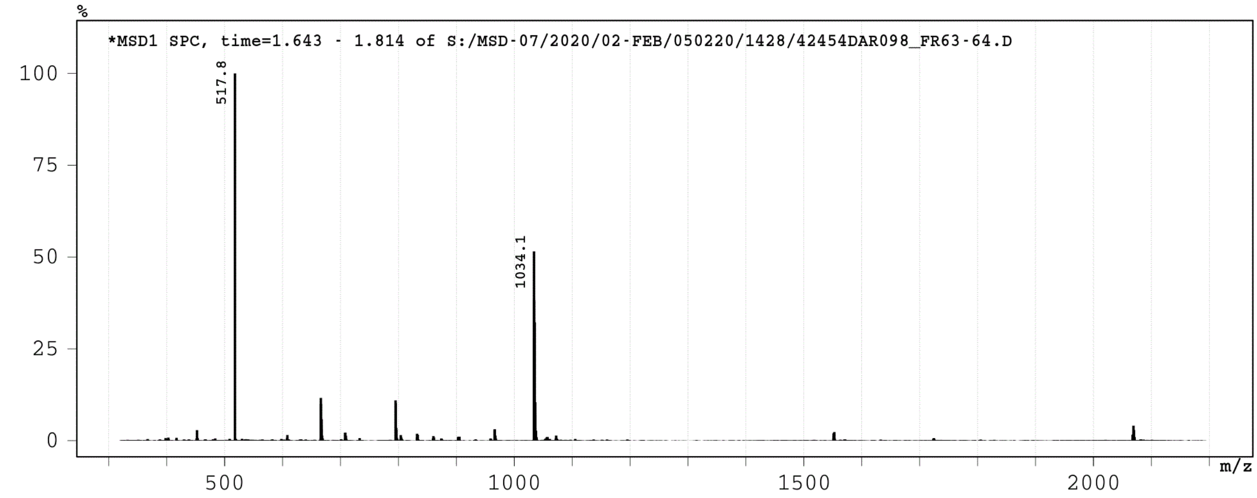


Signal 1: DAD1 A, Sig=220,4 Ref=off

| Peak # | RT [min] | Width [min] | Area | Area % |
|---|---|---|---|---|
| 1 | 1.624 | 1.29 | 57.4 | 1.2 |
| 2 | 1.692 | 0.20 | 4529.8 | 92.8 |
| 3 | 1.820 | 3.38 | 291.6 | 6.0 |

Data File : S:/MSD-07/2020/02-FEB/060220/1429/42454DAR099_FR5->

Injection Date : 06-Feb-20, 15:59:04
Sample Name : 42454DAR099_FR53
Remarks : Baseline corrected
Acq. Method : D:/METHODS/5-95-6M.M
column used : Je_559
Analysis Method : C:/CHEM32/1/METHODS/JPT-LCMS.M

Sample Info : 42454_099-Dr. Martine Darwish H-ESRSRFQTL-OH

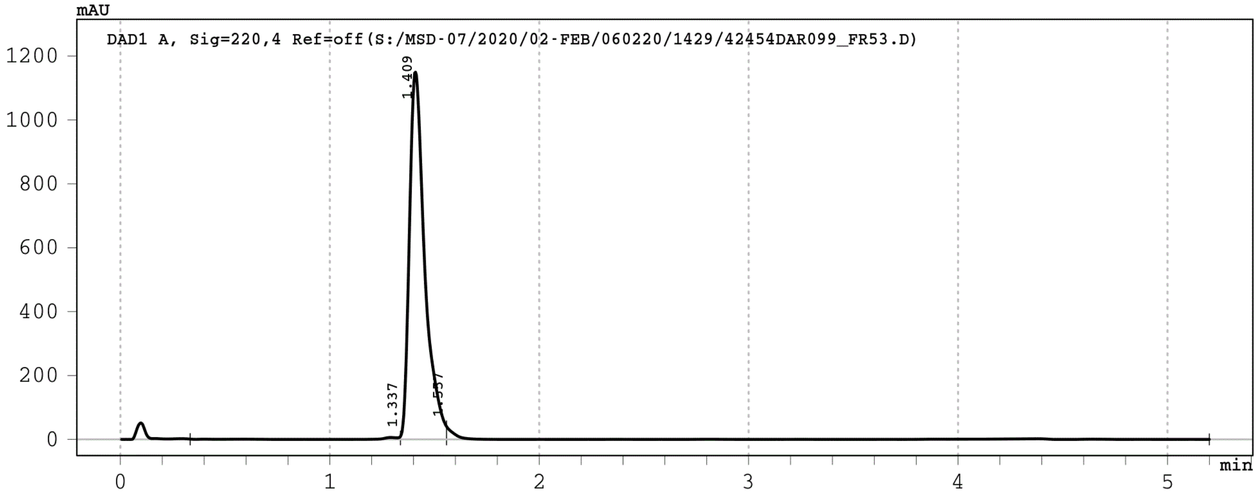


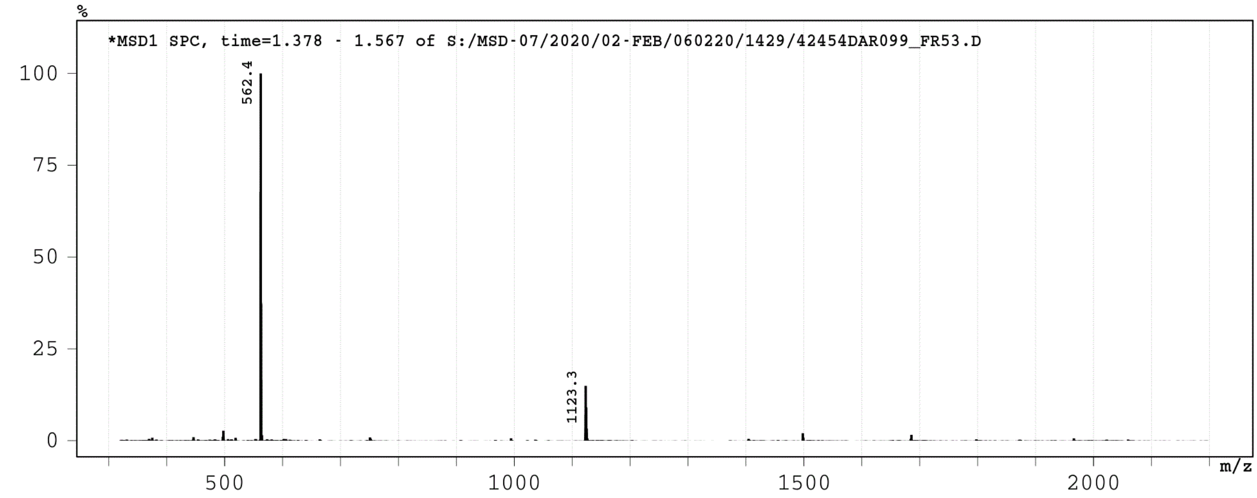


Signal 1: DAD1 A, Sig=220,4 Ref=off

| Peak # | RT [min] | Width [min] | Area | Area % |
|---|---|---|---|---|
| 1 | 1.337 | 1.00 | 41.5 | 0.7 |
| 2 | 1.409 | 0.22 | 5869.5 | 96.9 |
| 3 | 1.557 | 3.64 | 148.9 | 2.5 |

Data File : S:/MSD-07/2020/02-FEB/050220/1428/42454DAR100_FR8->

Injection Date : 05-Feb-20, 18:59:00
Sample Name : 42454DAR100_FR80-81
Remarks : Baseline corrected
Acq. Method : D:/METHODS/5-95-6M.M
column used : Je_559
Analysis Method : C:/CHEM32/1/METHODS/JPT-LCMS.M

Sample Info : 42454_100-Dr. Martine Darwish H-KMSSNAFFV-OH

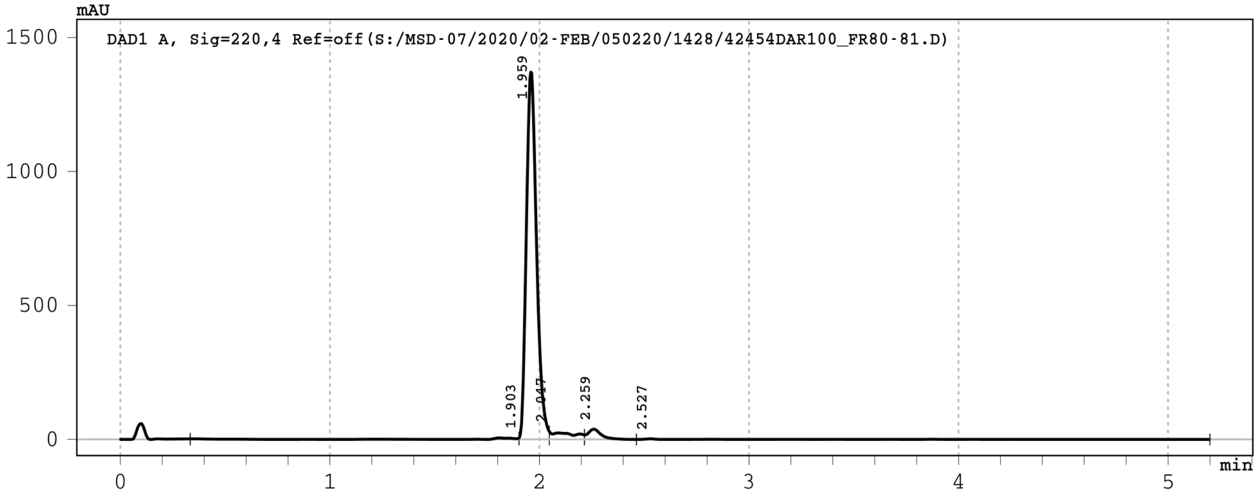


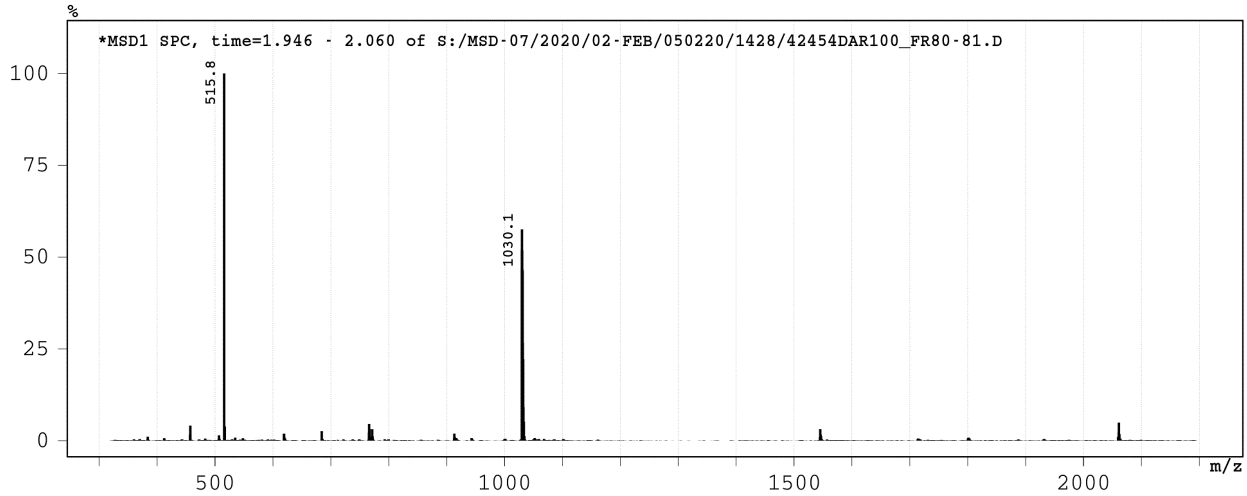


Signal 1: DAD1 A, Sig=220,4 Ref=off

| Peak # | RT [min] | Width [min] | Area | Area % |
|---|---|---|---|---|
| 1 | 1.903 | 1.57 | 65.5 | 1.3 |
| 2 | 1.959 | 0.14 | 4650.2 | 91.0 |
| 3 | 2.047 | 0.17 | 208.5 | 4.1 |
| 4 | 2.259 | 0.25 | 170.0 | 3.3 |
| 5 | 2.527 | 2.74 | 13.7 | 0.3 |

Data File : S:/MSD-07/2020/02-FEB/060220/1429/42454DAR101_FR6->

Injection Date : 06-Feb-20, 16:41:34
Sample Name : 42454DAR101_FR60
Remarks : Baseline corrected
Acq. Method : D:/METHODS/5-95-6M.M
column used : Je_559
Analysis Method : C:/CHEM32/1/METHODS/JPT-LCMS.M

Sample Info : 42454_101-Dr. Martine Darwish H-SYVVKLRGL-OH

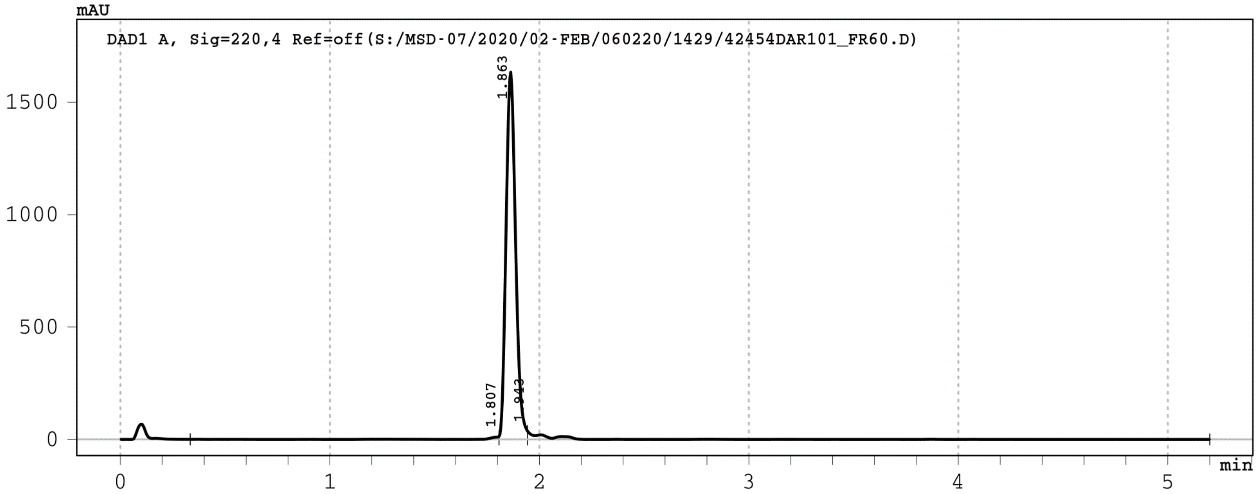


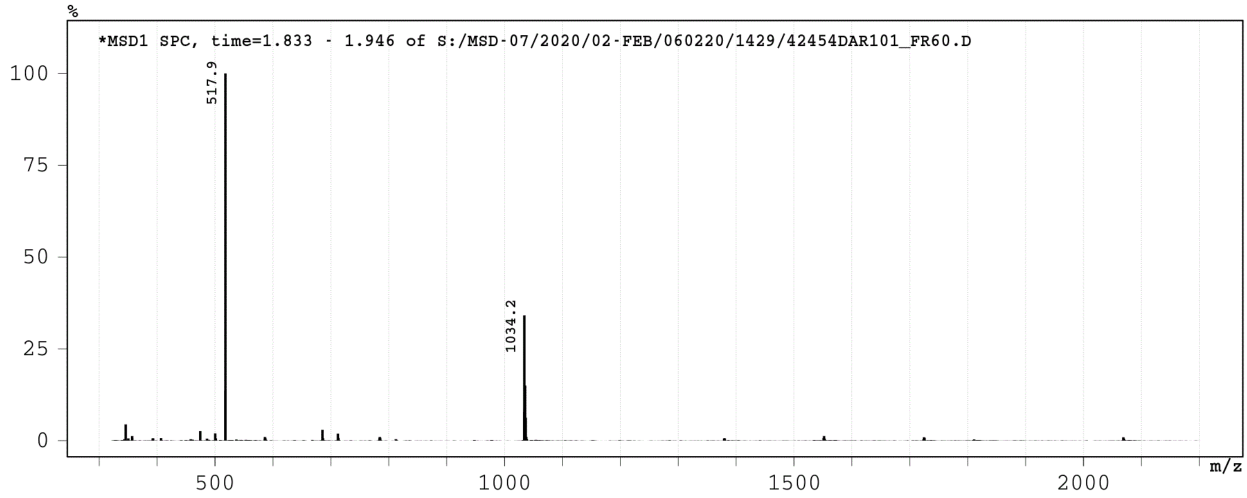


Signal 1: DAD1 A, Sig=220,4 Ref=off

| Peak # | RT [min] | Width [min] | Area | Area % |
|---|---|---|---|---|
| 1 | 1.807 | 1.47 | 50.5 | 0.9 |
| 2 | 1.863 | 0.14 | 5174.5 | 95.4 |
| 3 | 1.943 | 3.26 | 200.0 | 3.7 |

Data File : S:/MSD-07/2020/02-FEB/060220/1429/42454DAR102_FR6->

Injection Date : 06-Feb-20, 16:49:58
Sample Name : 42454DAR102_FR69-71
Remarks : Baseline corrected
Acq. Method : D:/METHODS/5-95-6M.M
column used : Je_559
Analysis Method : C:/CHEM32/1/METHODS/JPT-LCMS.M

Sample Info : 42454_102-Dr. Martine Darwish H-IHPKAFPL-OH

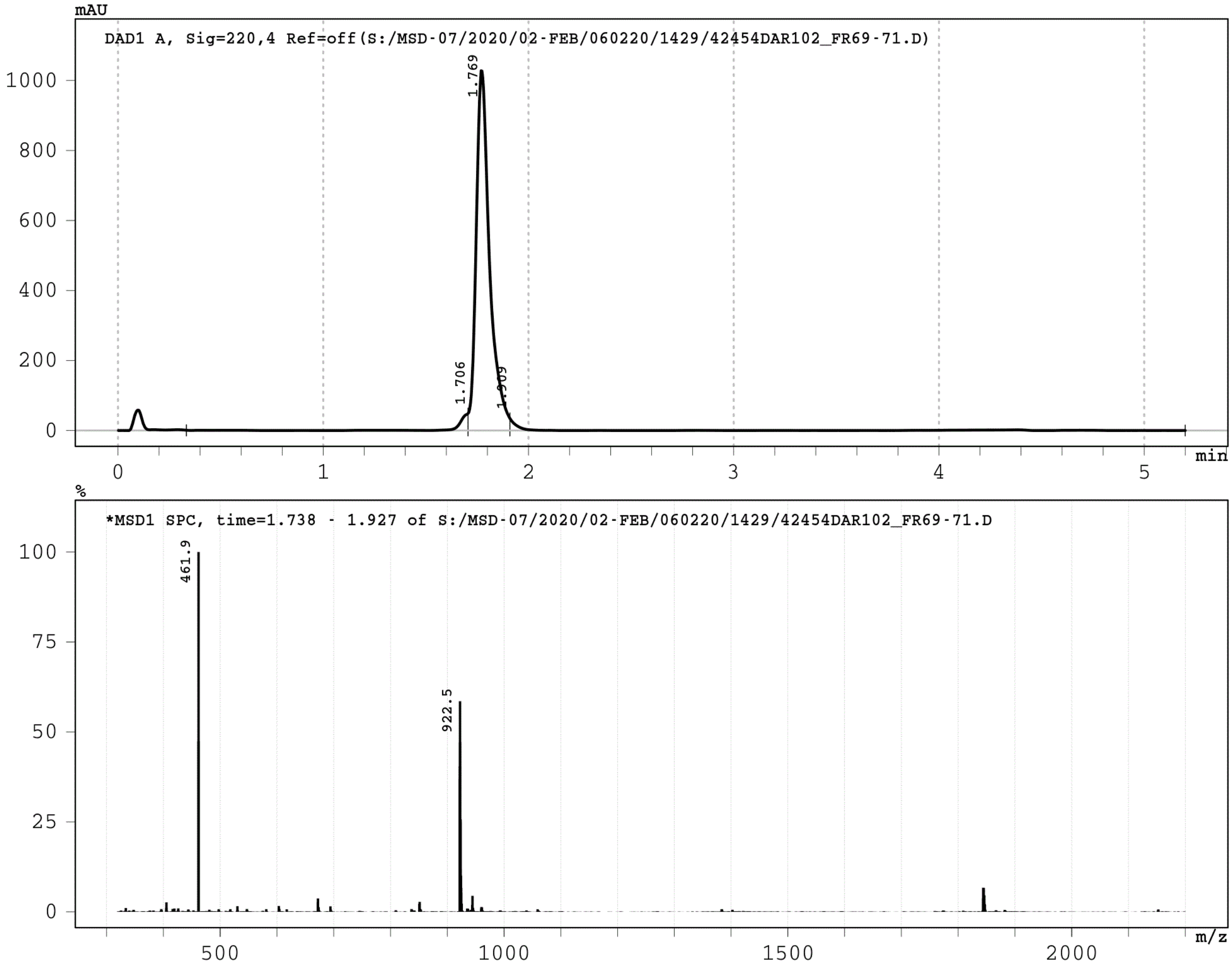


Signal 1: DAD1 A, Sig=220,4 Ref=off

| Peak # | RT [min] | Width [min] | Area | Area % |
|---|---|---|---|---|
| 1 | 1.706 | 1.37 | 144.3 | 2.9 |
| 2 | 1.769 | 0.20 | 4684.6 | 94.4 |
| 3 | 1.909 | 3.29 | 131.4 | 2.6 |

Data File : S:/MSD-07/2020/02-FEB/070220/1430/42454DAR103_FR4->

Injection Date : 07-Feb-20, 15:52:26
Sample Name : 42454DAR103_FR4-6
Remarks : Baseline corrected
Acq. Method : D:/METHODS/5-95-6M.M
column used : Je_559
Analysis Method : C:/CHEM32/1/METHODS/JPT-LCMS.M

Sample Info : 42454_103-Dr. Martine Darwish H-IHPKAFPF-OH

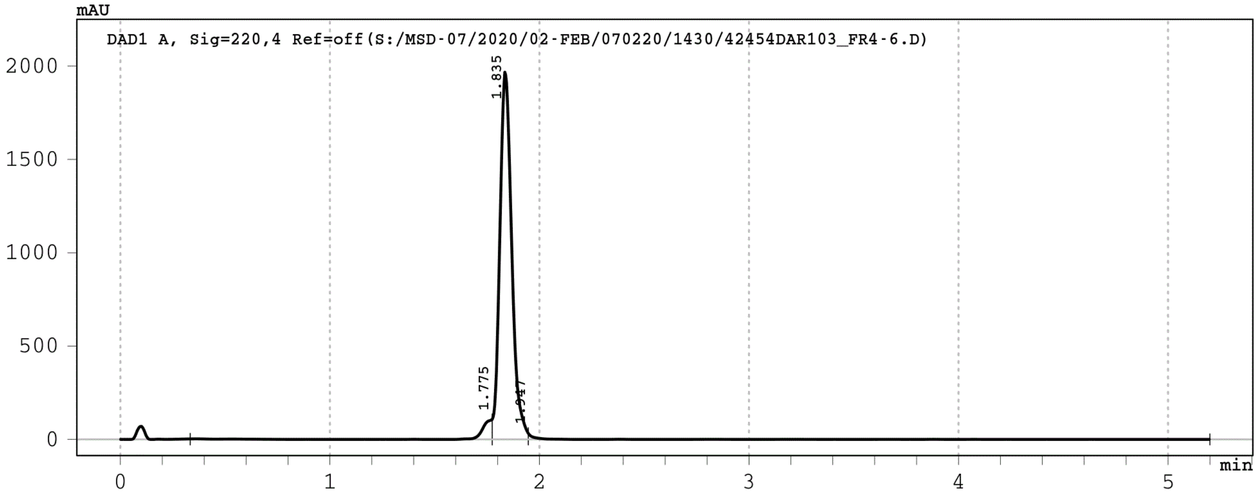


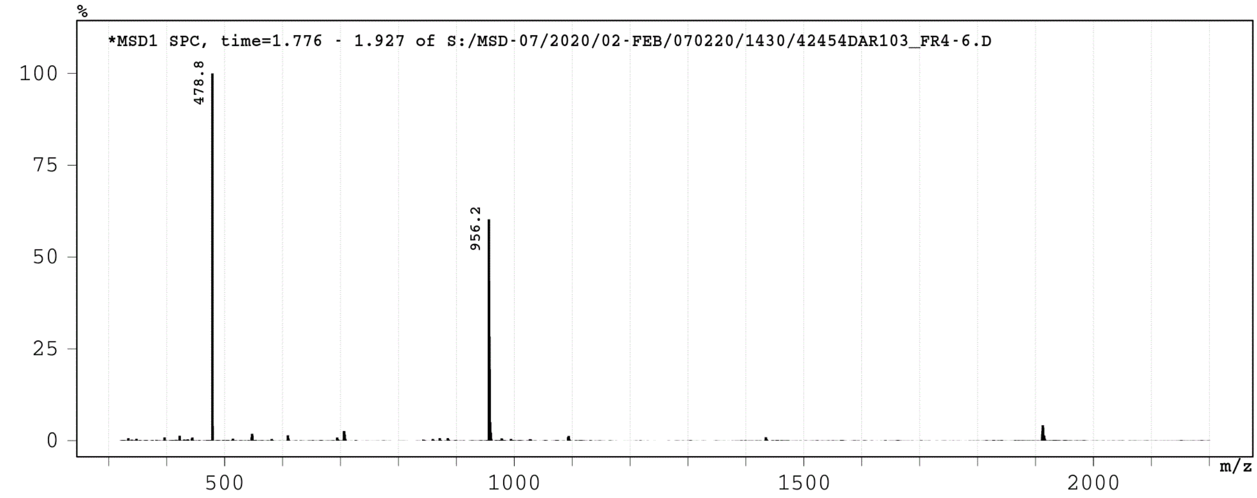


Signal 1: DAD1 A, Sig=220,4 Ref=off

| Peak # | RT [min] | Width [min] | Area | Area % |
|---|---|---|---|---|
| 1 | 1.775 | 1.44 | 364.1 | 4.4 |
| 2 | 1.835 | 0.17 | 7770.7 | 94.7 |
| 3 | 1.947 | 3.25 | 70.9 | 0.9 |

Data File : S:/MSD-07/2020/02-FEB/070220/1430/42454DAR104_FR1->

Injection Date : 07-Feb-20, 16:00:47
Sample Name : 42454DAR104_FR19-21
Remarks : Baseline corrected
Acq. Method : D:/METHODS/5-95-6M.M
column used : Je_559
Analysis Method : C:/CHEM32/1/METHODS/JPT-LCMS.M

Sample Info : 42454_104-Dr. Martine Darwish H-SLLAWRAL-OH

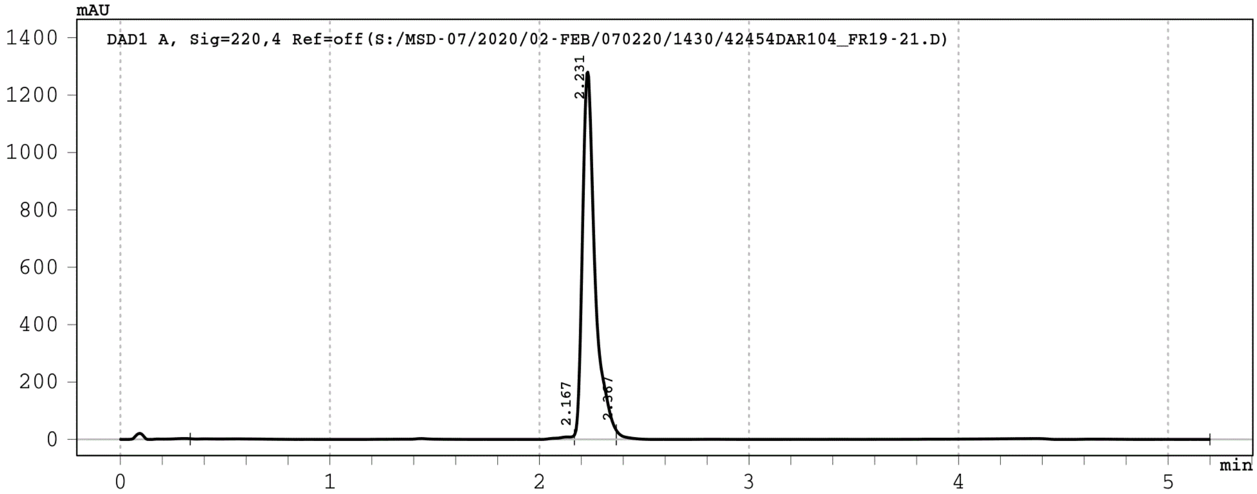


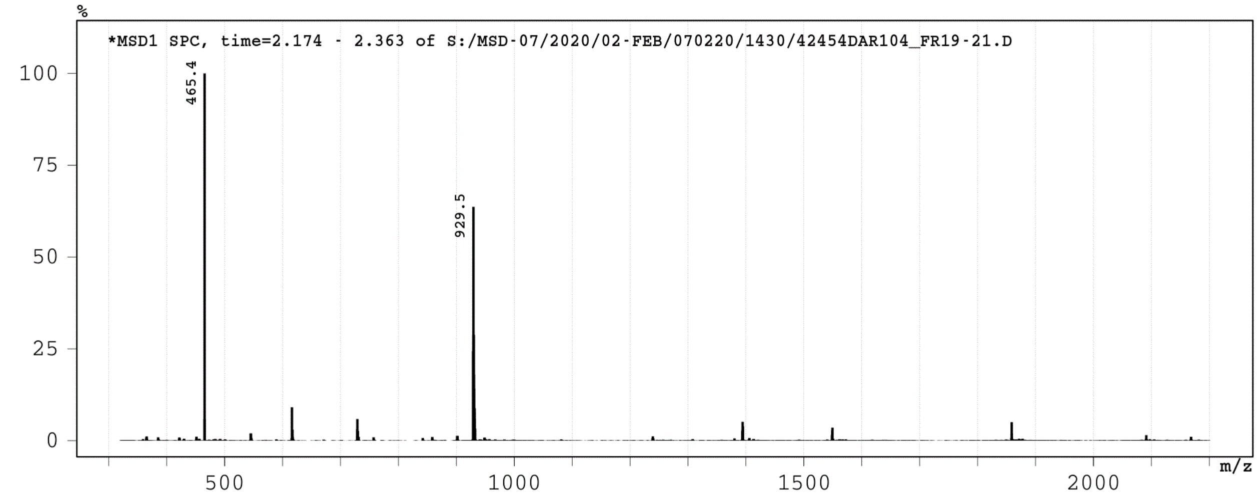


Signal 1: DAD1 A, Sig=220,4 Ref=off

| Peak # | RT [min] | Width [min] | Area | Area % |
|---|---|---|---|---|
| 1 | 2.167 | 1.83 | 108.6 | 1.9 |
| 2 | 2.231 | 0.20 | 5364.8 | 95.5 |
| 3 | 2.367 | 2.83 | 143.4 | 2.6 |

Data File : S:/MSD-07/2020/02-FEB/070220/1430/42454DAR105_FR4->

Injection Date : 07-Feb-20, 16:09:09
Sample Name : 42454DAR105_FR43
Remarks : Baseline corrected
Acq. Method : D:/METHODS/5-95-6M.M
column used : Je_559
Analysis Method : C:/CHEM32/1/METHODS/JPT-LCMS.M

Sample Info : 42454_105-Dr. Martine Darwish H-AWLSKVSRL-OH

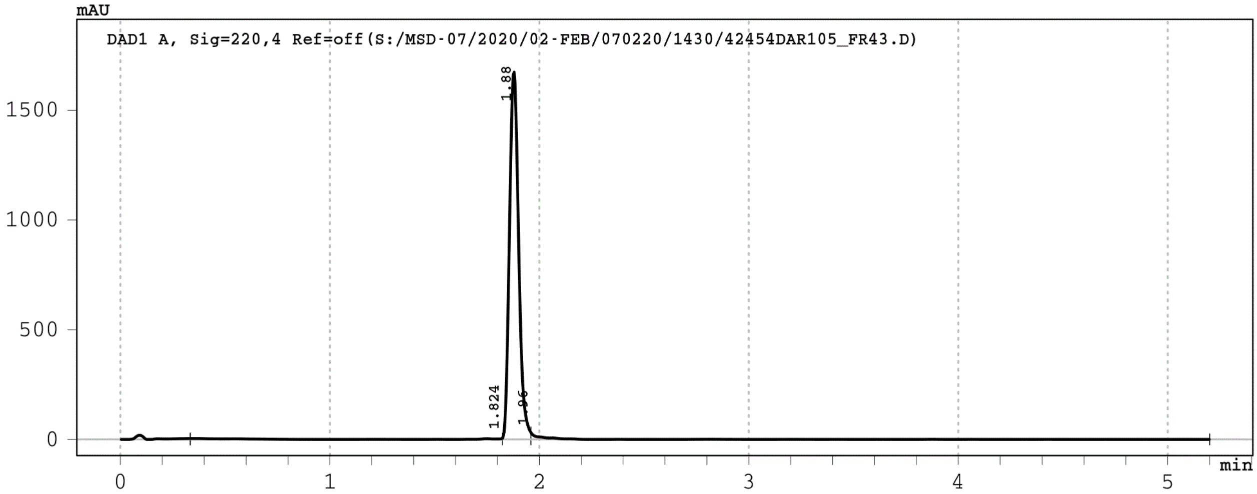


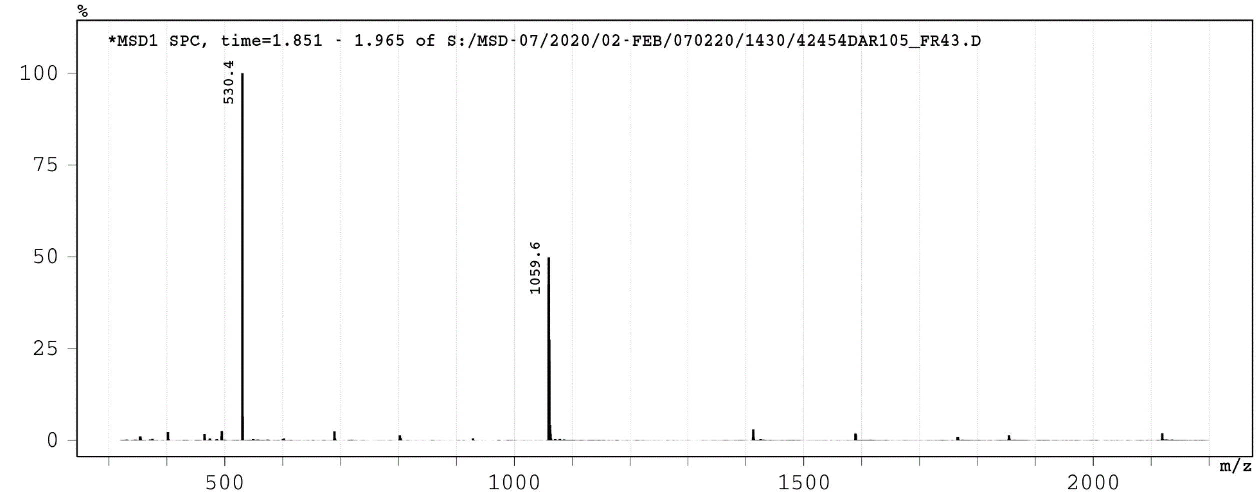


Signal 1: DAD1 A, Sig=220,4 Ref=off

| Peak # | RT [min] | Width [min] | Area | Area % |
|---|---|---|---|---|
| 1 | 1.824 | 1.49 | 84.1 | 1.6 |
| 2 | 1.880 | 0.14 | 5111.6 | 96.4 |
| 3 | 1.960 | 3.24 | 107.7 | 2.0 |

Data File : S:/MSD-07/2020/02-FEB/070220/1430/42454DAR106_FR6->

Injection Date : 07-Feb-20, 16:17:31
Sample Name : 42454DAR106_FR60
Remarks : Baseline corrected
Acq. Method : D:/METHODS/5-95-6M.M
column used : Je_559
Analysis Method : C:/CHEM32/1/METHODS/JPT-LCMS.M

Sample Info : 42454_106-Dr. Martine Darwish H-SSLKSYVQL-OH

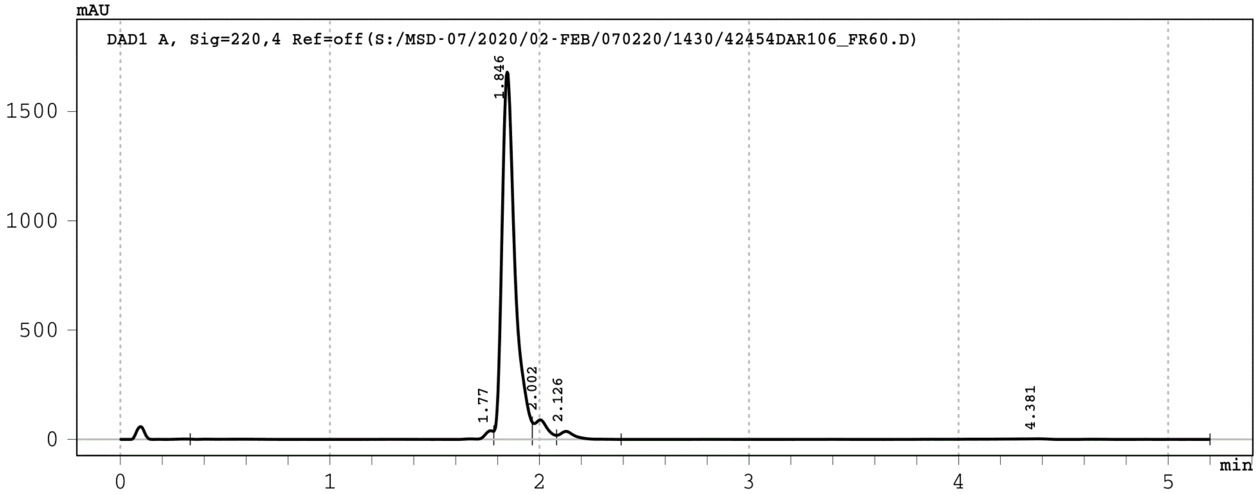


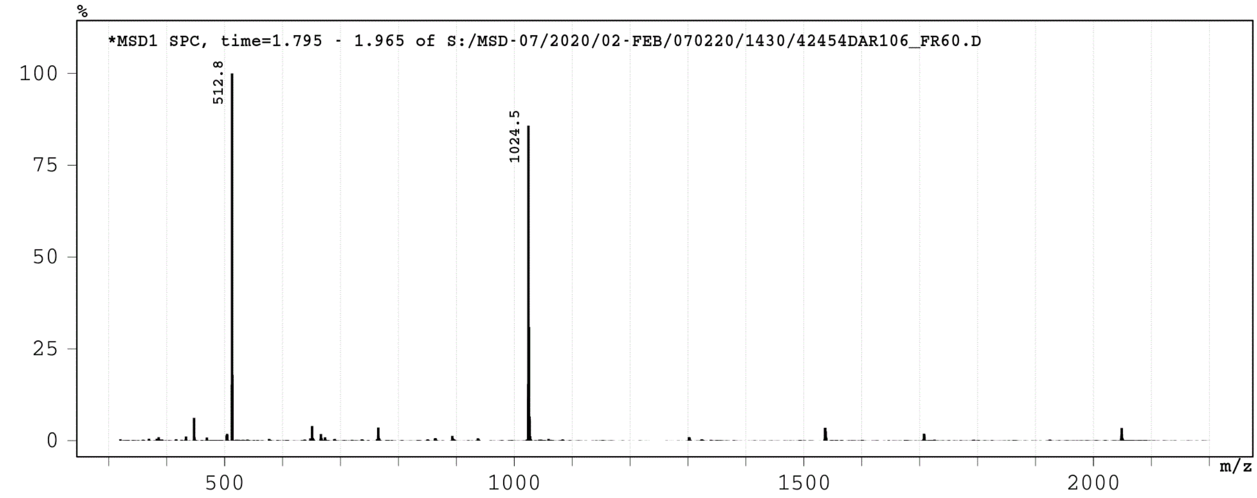


Signal 1: DAD1 A, Sig=220,4 Ref=off

| Peak # | RT [min] | Width [min] | Area | Area % |
|---|---|---|---|---|
| 1 | 1.770 | 1.45 | 132.4 | 1.6 |
| 2 | 1.846 | 0.18 | 7261.2 | 90.3 |
| 3 | 2.002 | 0.12 | 407.1 | 5.1 |
| 4 | 2.126 | 0.31 | 182.7 | 2.3 |
| 5 | 4.381 | 2.81 | 61.9 | 0.8 |

Data File : S:/MSD-07/2020/02-FEB/070220/1430/42454DAR107_FR7->

Injection Date : 07-Feb-20, 16:25:56
Sample Name : 42454DAR107_FR72-73
Remarks : Baseline corrected
Acq. Method : D:/METHODS/5-95-6M.M
column used : Je_559
Analysis Method : C:/CHEM32/1/METHODS/JPT-LCMS.M

Sample Info : 42454_107-Dr. Martine Darwish H-VNGLLYVSV-OH

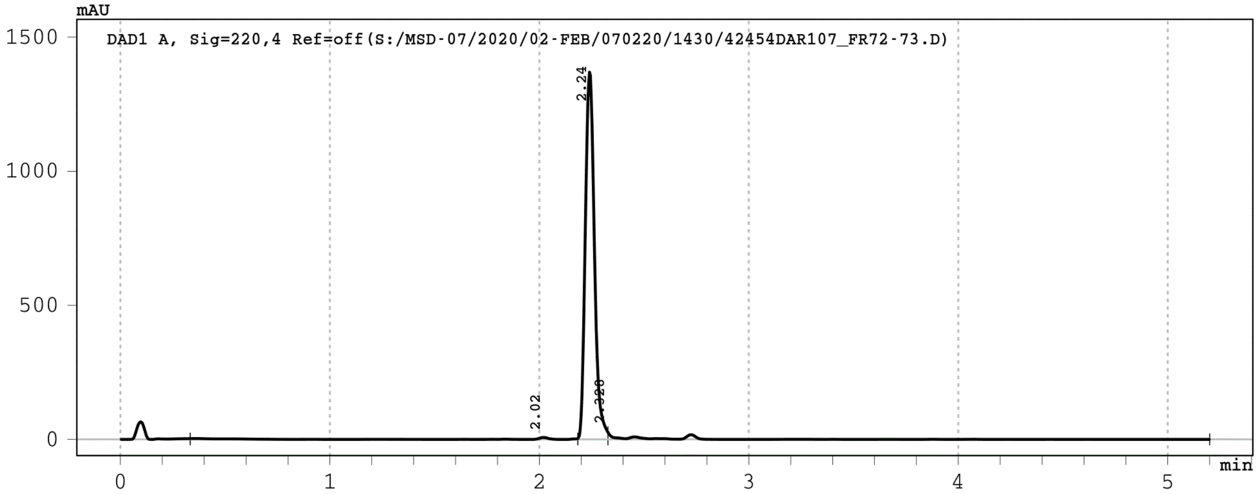


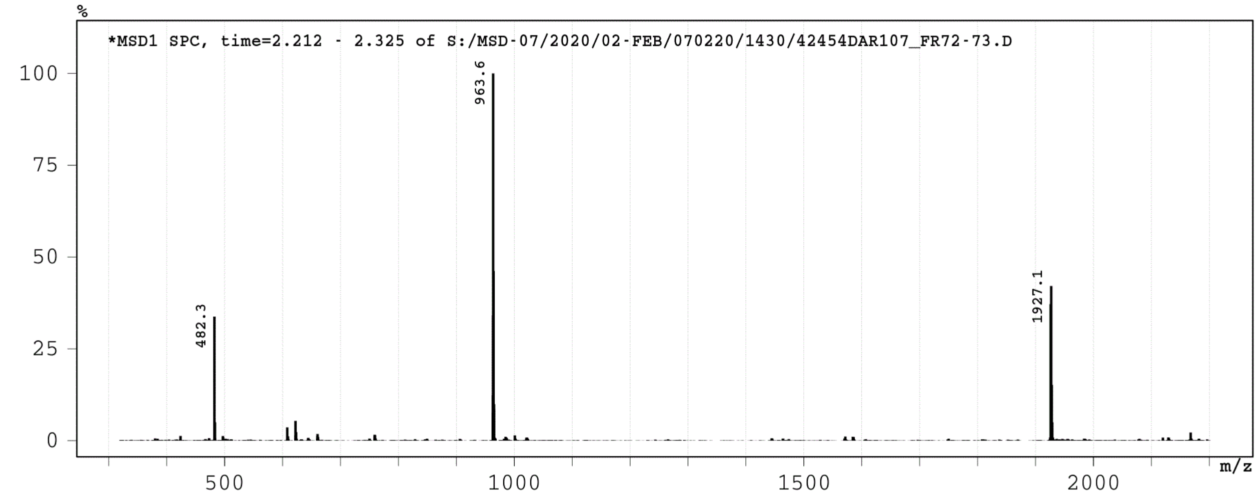


Signal 1: DAD1 A, Sig=220,4 Ref=off

| Peak # | RT [min] | Width [min] | Area | Area % |
|---|---|---|---|---|
| 1 | 2.020 | 1.85 | 78.8 | 1.8 |
| 2 | 2.240 | 0.14 | 4157.4 | 94.6 |
| 3 | 2.328 | 2.87 | 160.0 | 3.6 |

Innovative Peptide Solutions

Data File : S:/MSD-07/2020/02-FEB/050220/1428/42454DAR108_FR1->

Injection Date : 05-Feb-20, 19:07:23
Sample Name : 42454DAR108_FR17-18
Remarks : Baseline corrected
Acq. Method : D:/METHODS/5-95-6M.M
column used : Je_559
Analysis Method : C:/CHEM32/1/METHODS/JPT-LCMS.M

Sample Info : 42454_108-Dr. Martine Darwish H-RVYLMLPL-OH

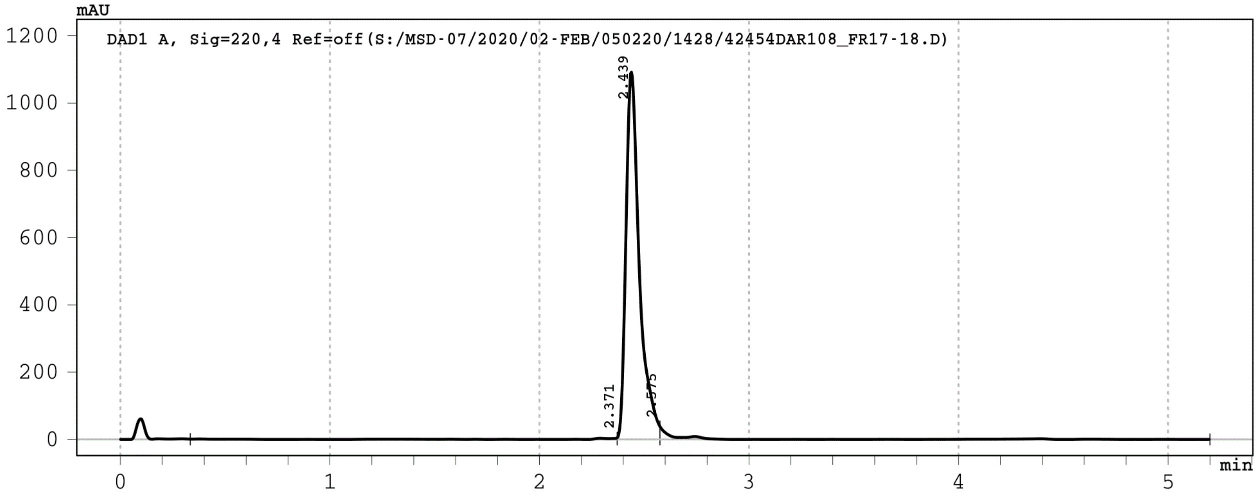


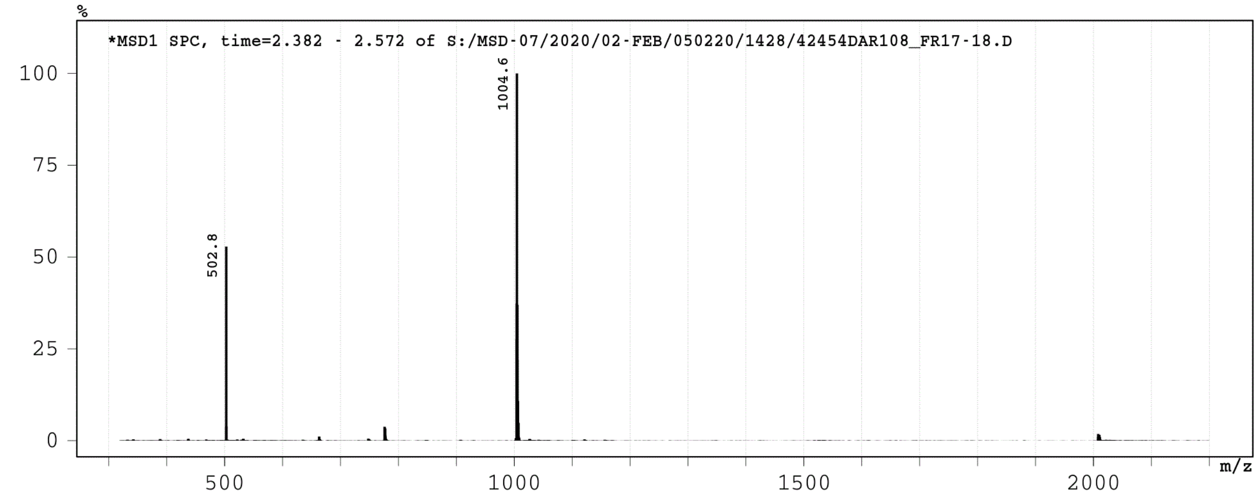


Signal 1: DAD1 A, Sig=220,4 Ref=off

| Peak # | RT [min] | Width [min] | Area | Area % |
|---|---|---|---|---|
| 1 | 2.371 | 2.04 | 54.0 | 1.1 |
| 2 | 2.439 | 0.20 | 4875.2 | 95.5 |
| 3 | 2.575 | 2.63 | 176.4 | 3.5 |

Data File : S:/MSD-07/2020/02-FEB/070220/1430/42454DAR109_FR8->

Injection Date : 07-Feb-20, 16:34:18
Sample Name : 42454DAR109_FR82
Remarks : Baseline corrected
Acq. Method : D:/METHODS/5-95-6M.M
column used : Je_559
Analysis Method : C:/CHEM32/1/METHODS/JPT-LCMS.M

Sample Info : 42454_109-Dr. Martine Darwish H-AALLNSAVL-OH

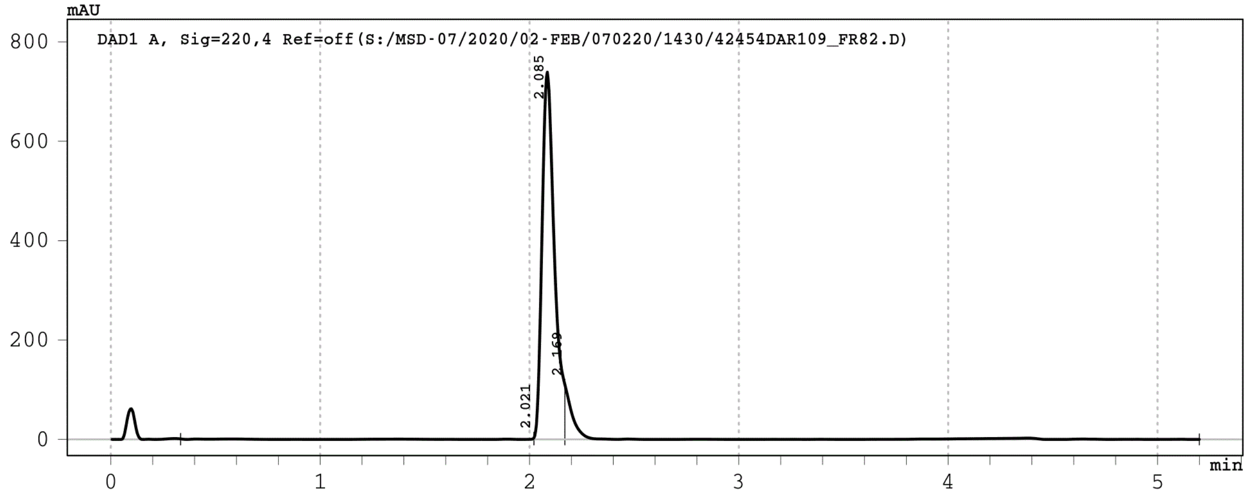


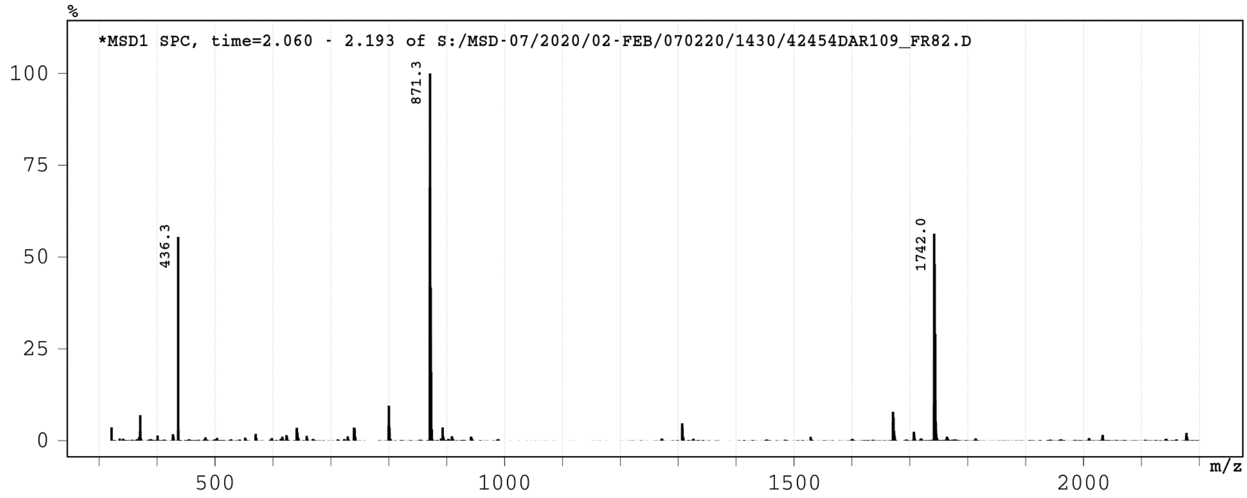


Signal 1: DAD1 A, Sig=220,4 Ref=off

| Peak # | RT [min] | Width [min] | Area | Area % |
|---|---|---|---|---|
| 1 | 2.021 | 1.69 | 25.5 | 0.8 |
| 2 | 2.085 | 0.15 | 3029.1 | 89.5 |
| 3 | 2.169 | 3.03 | 331.0 | 9.8 |

Data File : S:/MSD-07/2020/02-FEB/070220/1430/42454DAR110_FR2->

Injection Date : 07-Feb-20, 16:42:43
Sample Name : 42454DAR110_FR2
Remarks : Baseline corrected
Acq. Method : D:/METHODS/5-95-6M.M
column used : Je_559
Analysis Method : C:/CHEM32/1/METHODS/JPT-LCMS.M

Sample Info : 42454_110-Dr. Martine Darwish H-VQINNPVTFL-OH

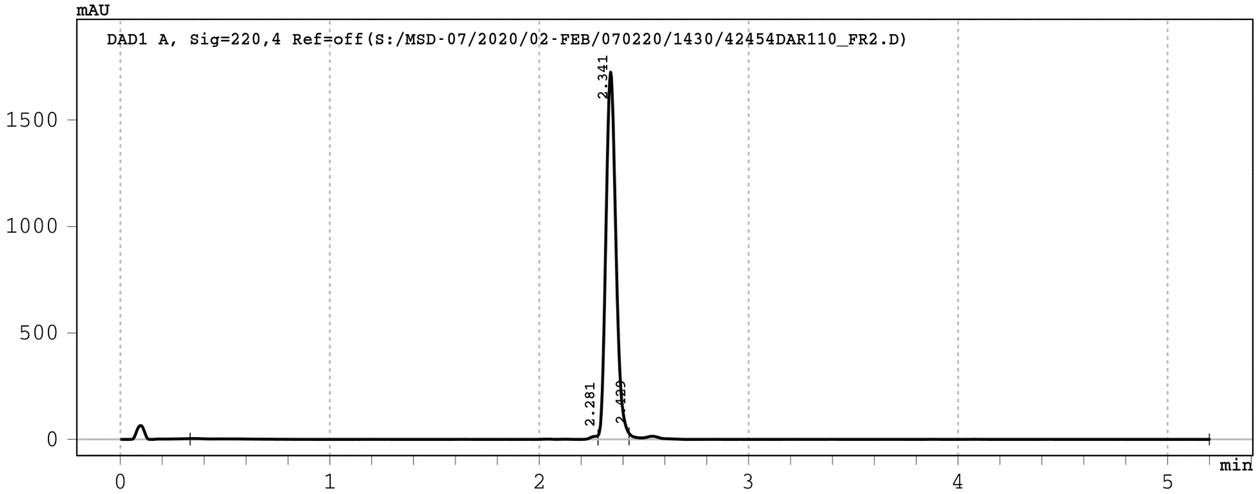


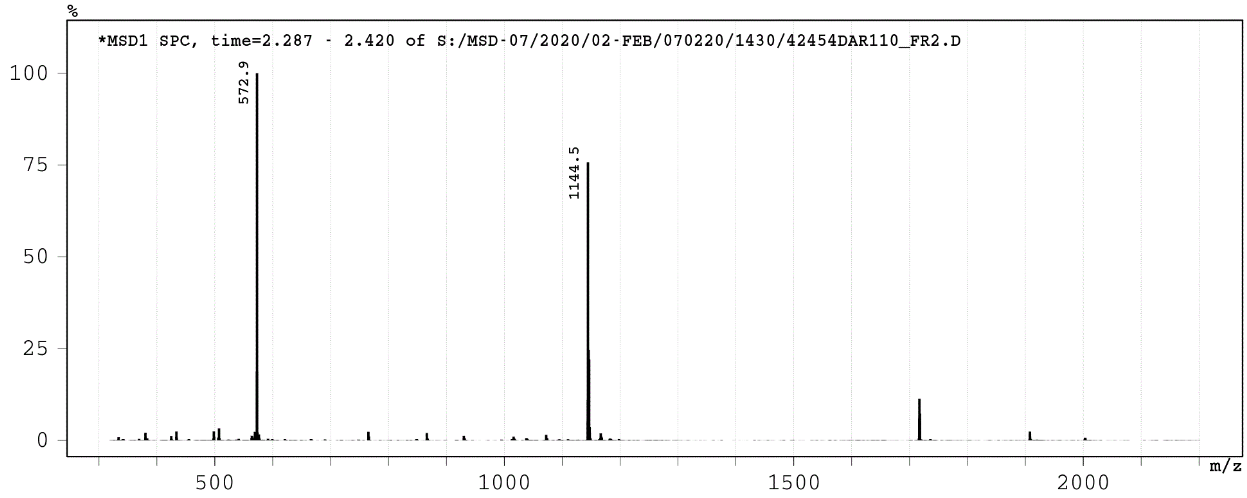


Signal 1: DAD1 A, Sig=220,4 Ref=off

| Peak # | RT [min] | Width [min] | Area | Area % |
|---|---|---|---|---|
| 1 | 2.281 | 1.95 | 100.8 | 1.7 |
| 2 | 2.341 | 0.15 | 5749.2 | 96.2 |
| 3 | 2.429 | 2.77 | 128.3 | 2.1 |

Data File : S:/MSD-07/2020/02-FEB/070220/1430/42454DAR112_FR7->

Injection Date : 07-Feb-20, 16:51:05
Sample Name : 42454DAR112_FR7-9
Remarks : Baseline corrected
Acq. Method : D:/METHODS/5-95-6M.M
column used : Je_559
Analysis Method : C:/CHEM32/1/METHODS/JPT-LCMS.M

Sample Info : 42454_112-Dr. Martine Darwish H-SQESNGVYV-OH

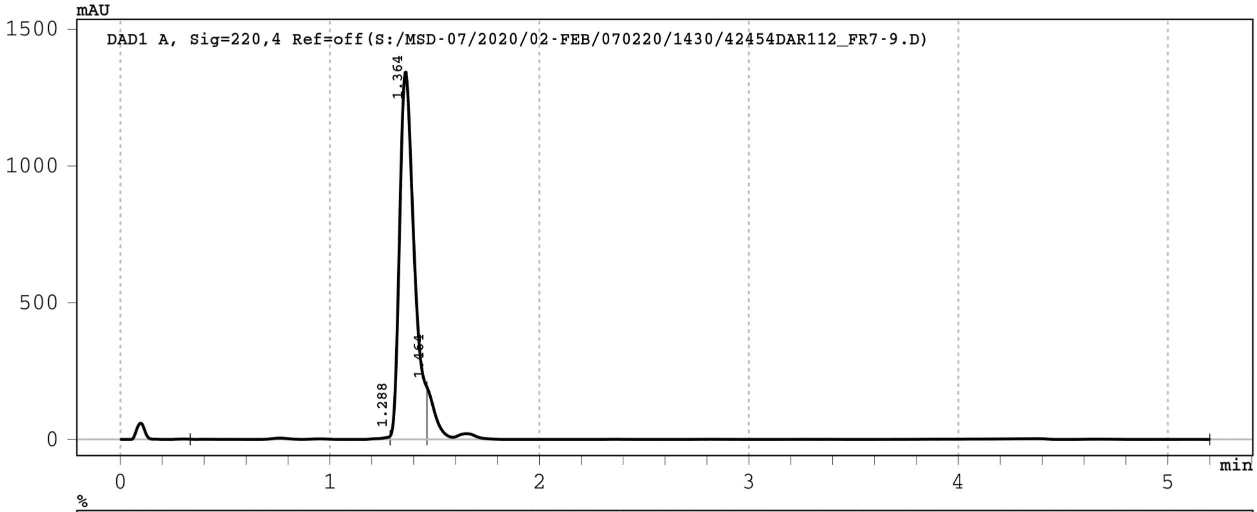


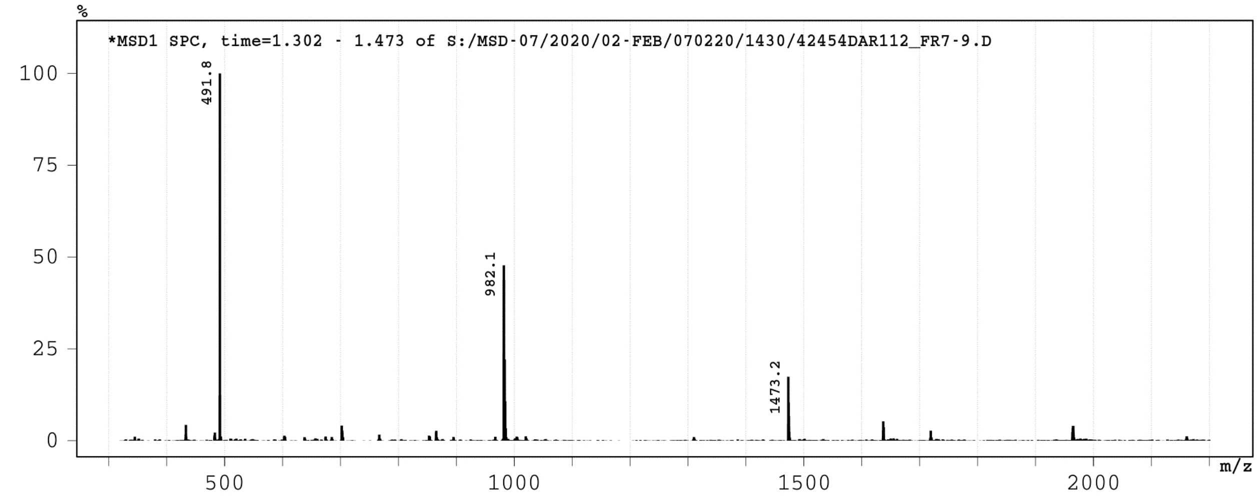


Signal 1: DAD1 A, Sig=220,4 Ref=off

| Peak # | RT [min] | Width [min] | Area | Area % |
|---|---|---|---|---|
| 1 | 1.288 | 0.95 | 64.8 | 0.9 |
| 2 | 1.364 | 0.18 | 6496.3 | 89.4 |
| 3 | 1.464 | 3.74 | 709.3 | 9.8 |

Data File : S:/MSD-07/2020/02-FEB/050220/1428/42454DAR113_FR3->

Injection Date : 05-Feb-20, 19:15:47
Sample Name : 42454DAR113_FR36-37
Remarks : Baseline corrected
Acq. Method : D:/METHODS/5-95-6M.M
column used : Je_559
Analysis Method : C:/CHEM32/1/METHODS/JPT-LCMS.M

Sample Info : 42454_113-Dr. Martine Darwish H-IVYLYVVC-OH

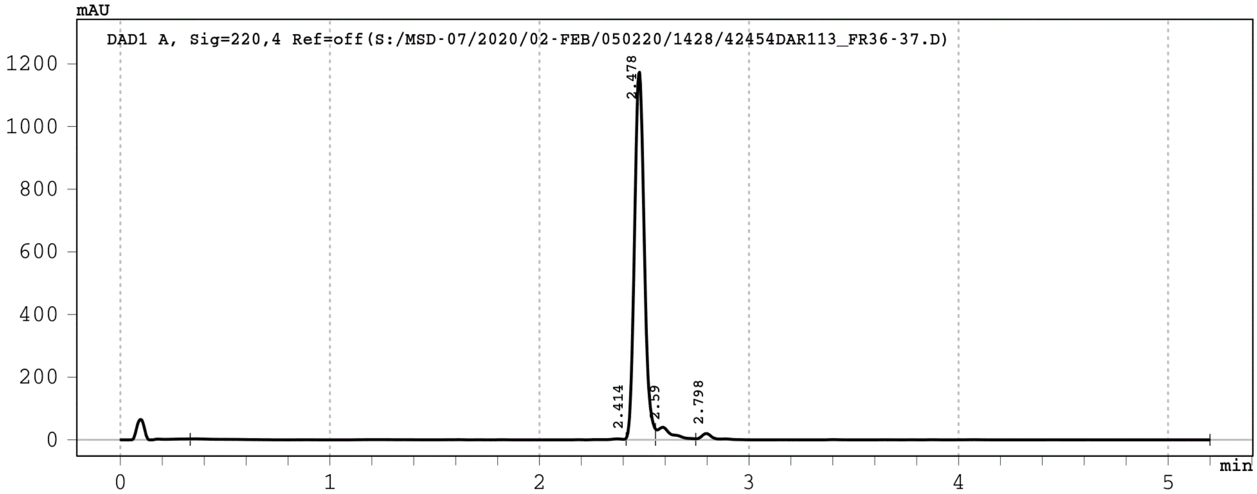


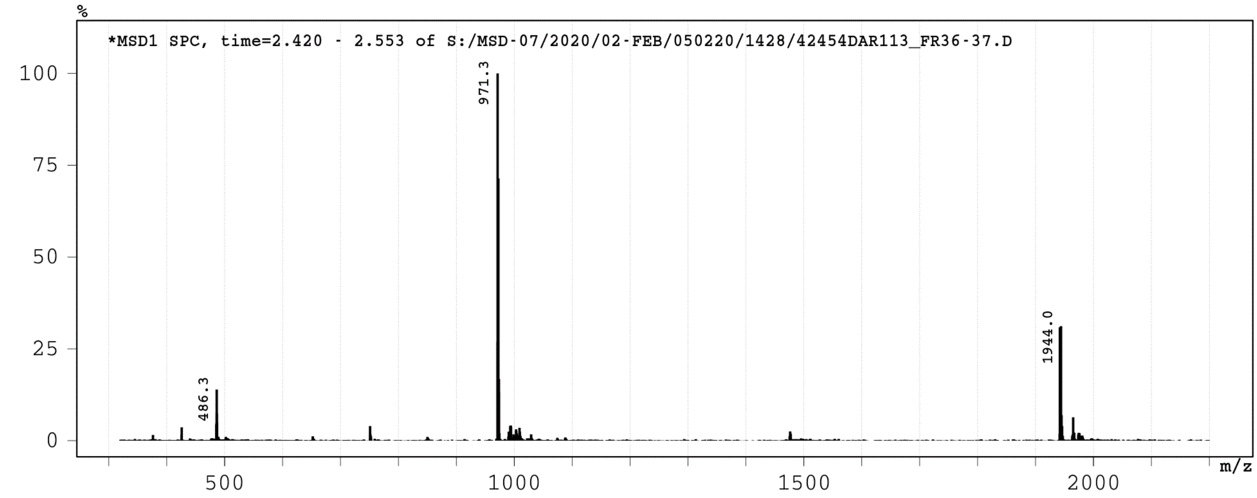


Signal 1: DAD1 A, Sig=220,4 Ref=off

| Peak # | RT [min] | Width [min] | Area | Area % |
|---|---|---|---|---|
| 1 | 2.414 | 2.08 | 69.8 | 1.7 |
| 2 | 2.478 | 0.14 | 3806.3 | 91.2 |
| 3 | 2.590 | 0.19 | 207.4 | 5.0 |
| 4 | 2.798 | 2.45 | 89.5 | 2.1 |

Data File : S:/MSD-07/2020/02-FEB/070220/1430/42454DAR114_FR2->

Injection Date : 07-Feb-20, 16:59:28
Sample Name : 42454DAR114_FR20-22
Remarks : Baseline corrected
Acq. Method : D:/METHODS/5-95-6M.M
column used : Je_559
Analysis Method : C:/CHEM32/1/METHODS/JPT-LCMS.M

Sample Info : 42454_114-Dr. Martine Darwish H-AFFEAASI-OH

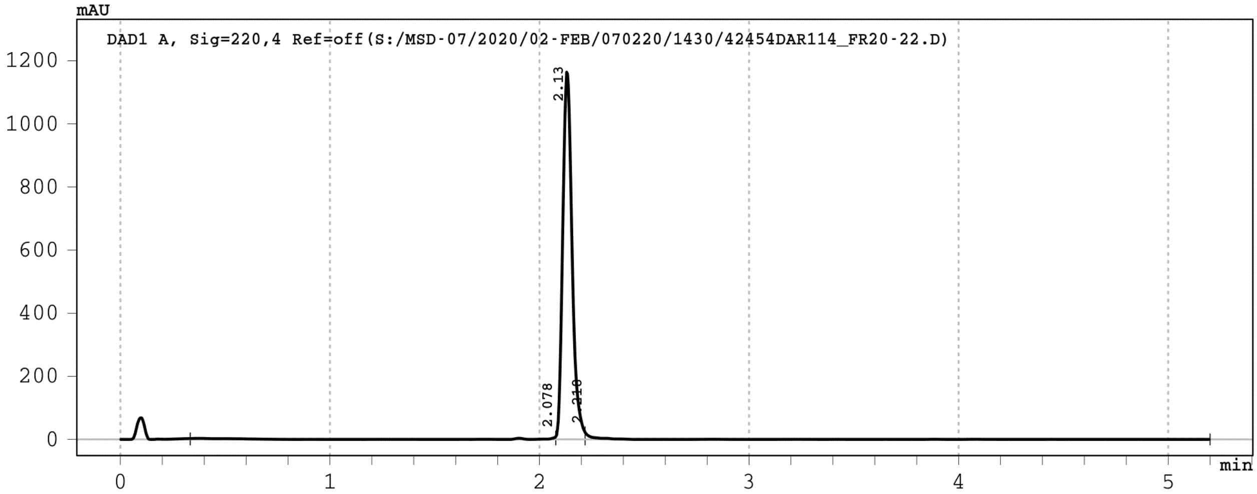


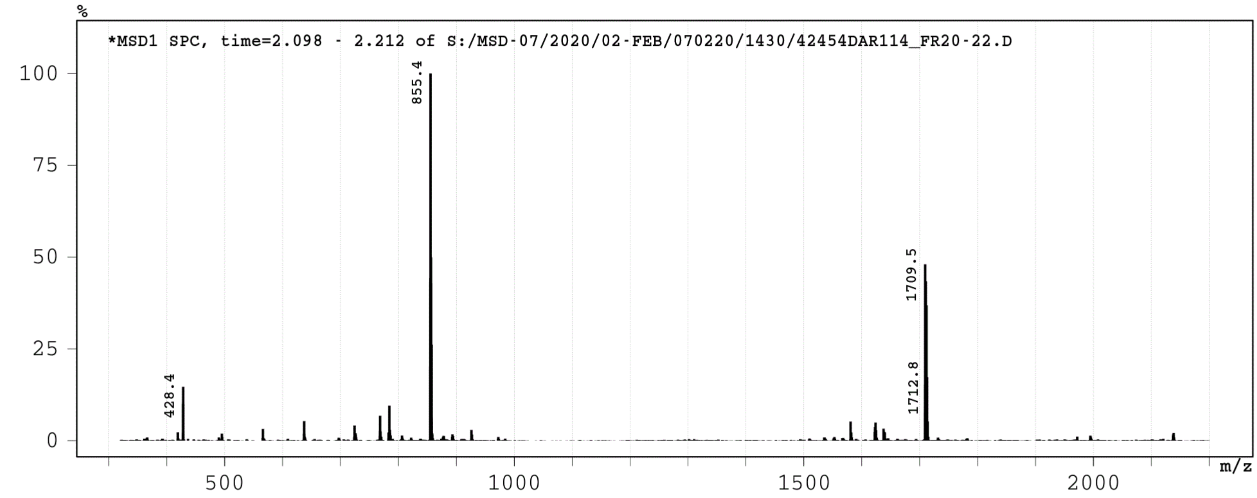


Signal 1: DAD1 A, Sig=220,4 Ref=off

| Peak # | RT [min] | Width [min] | Area | Area % |
|---|---|---|---|---|
| 1 | 2.078 | 1.74 | 73.9 | 2.0 |
| 2 | 2.130 | 0.14 | 3617.5 | 96.4 |
| 3 | 2.218 | 2.98 | 63.1 | 1.7 |

Data File : S:/MSD-07/2020/02-FEB/140220/1435/42454DAR115_FR1->

Injection Date : 14-Feb-20, 23:53:14
Sample Name : 42454DAR115_FR18-19
Remarks : Baseline corrected
Acq. Method : D:/METHODS/5-95-6M.M
column used : Je_559
Analysis Method : C:/CHEM32/1/METHODS/JPT-LCMS.M

Sample Info : 42454_115-Dr. Martine Darwish H-IQIAKEMAL-OH

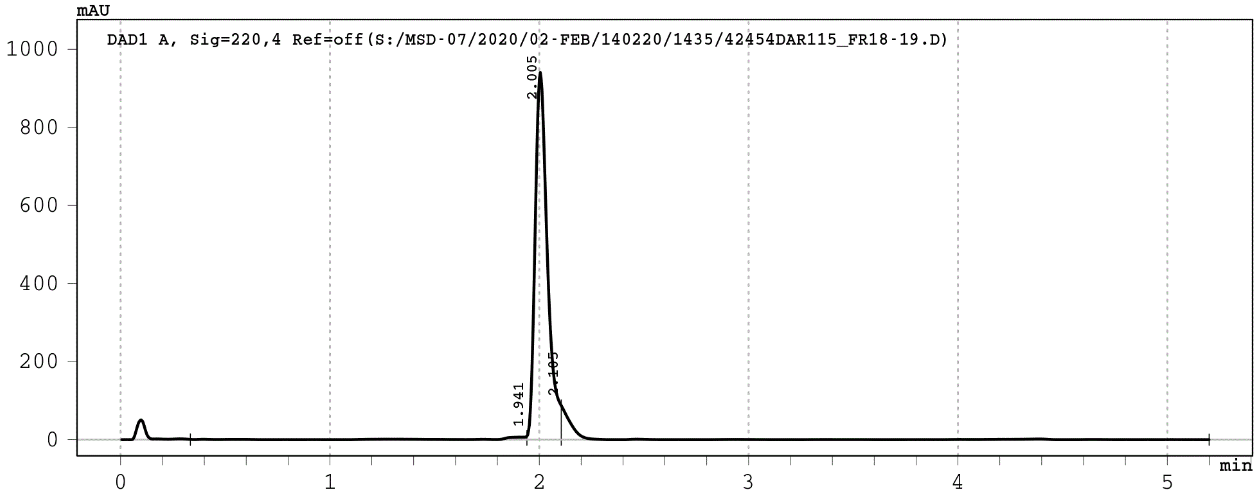


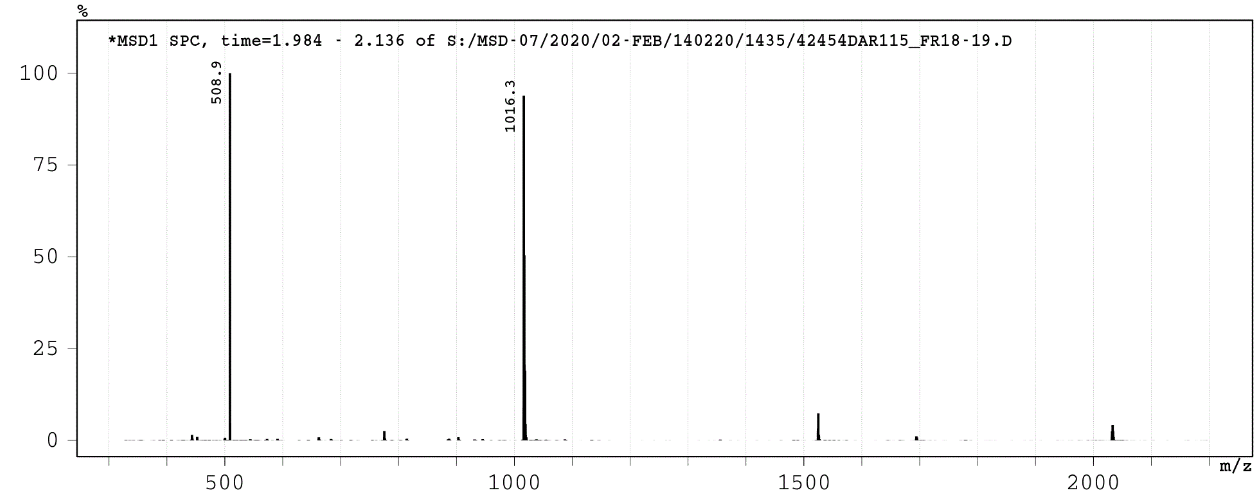


Signal 1: DAD1 A, Sig=220,4 Ref=off

| Peak # | RT [min] | Width [min] | Area | Area % |
|---|---|---|---|---|
| 1 | 1.941 | 1.61 | 79.3 | 1.9 |
| 2 | 2.005 | 0.16 | 3857.2 | 91.1 |
| 3 | 2.105 | 3.10 | 298.9 | 7.1 |

Data File : S:/MSD-07/2020/02-FEB/070220/1430/42454DAR116_FR2->

Injection Date : 07-Feb-20, 17:07:50
Sample Name : 42454DAR116_FR25-26
Remarks : Baseline corrected
Acq. Method : D:/METHODS/5-95-6M.M
column used : Je_559
Analysis Method : C:/CHEM32/1/METHODS/JPT-LCMS.M

Sample Info : 42454_116-Dr. Martine Darwish H-ASISTATPV-OH

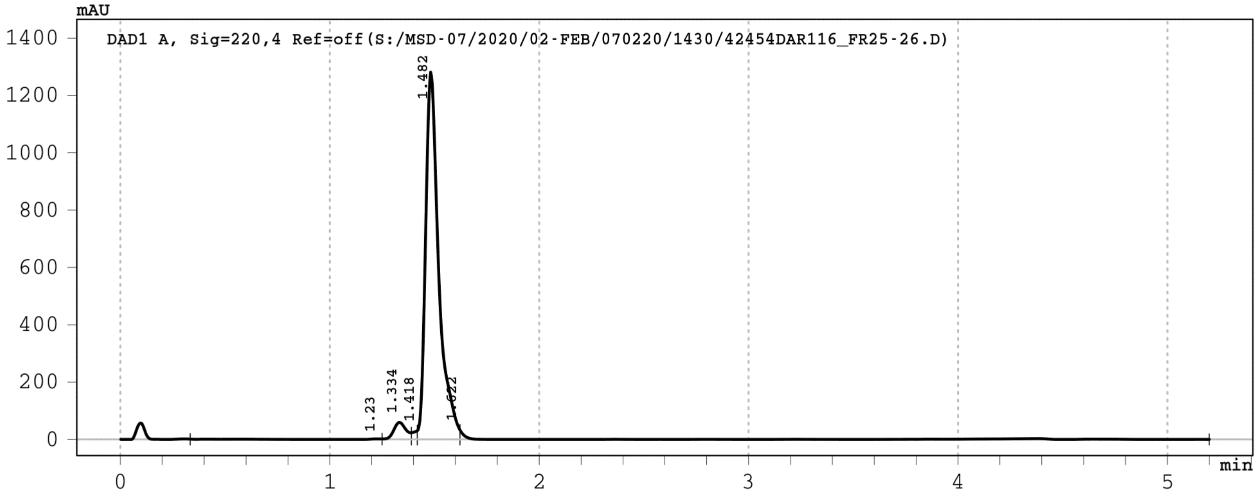


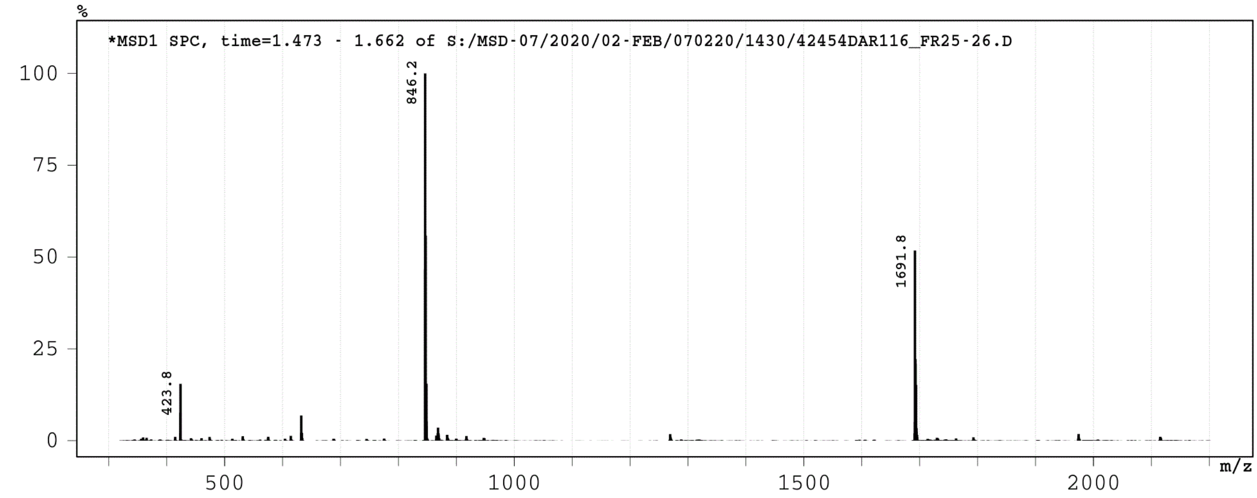


Signal 1: DAD1 A, Sig=220,4 Ref=off

| Peak # | RT [min] | Width [min] | Area | Area % |
|---|---|---|---|---|
| 1 | 1.230 | 0.92 | 21.4 | 0.4 |
| 2 | 1.334 | 0.14 | 244.1 | 4.1 |
| 3 | 1.418 | 0.03 | 43.8 | 0.7 |
| 4 | 1.482 | 0.20 | 5592.2 | 92.9 |
| 5 | 1.622 | 3.58 | 120.9 | 2.0 |

Data File : S:/MSD-07/2020/02-FEB/050220/1428/42454DAR117_FR5->

Injection Date : 05-Feb-20, 19:24:10
Sample Name : 42454DAR117_FR52
Remarks : Baseline corrected
Acq. Method : D:/METHODS/5-95-6M.M
column used : Je_559
Analysis Method : C:/CHEM32/1/METHODS/JPT-LCMS.M

Sample Info : 42454_117-Dr. Martine Darwish H-WARCNFNLL-OH

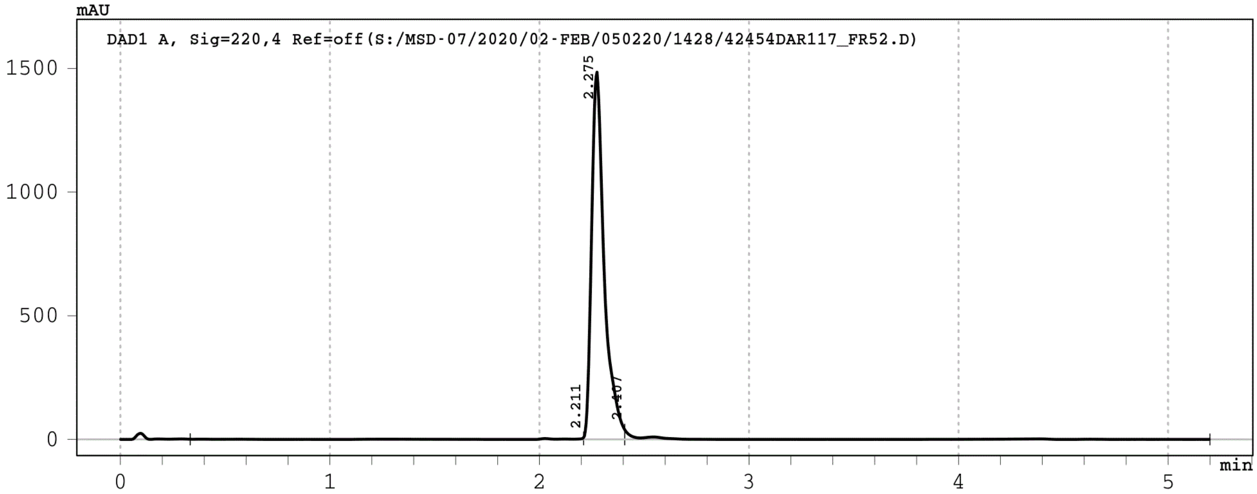


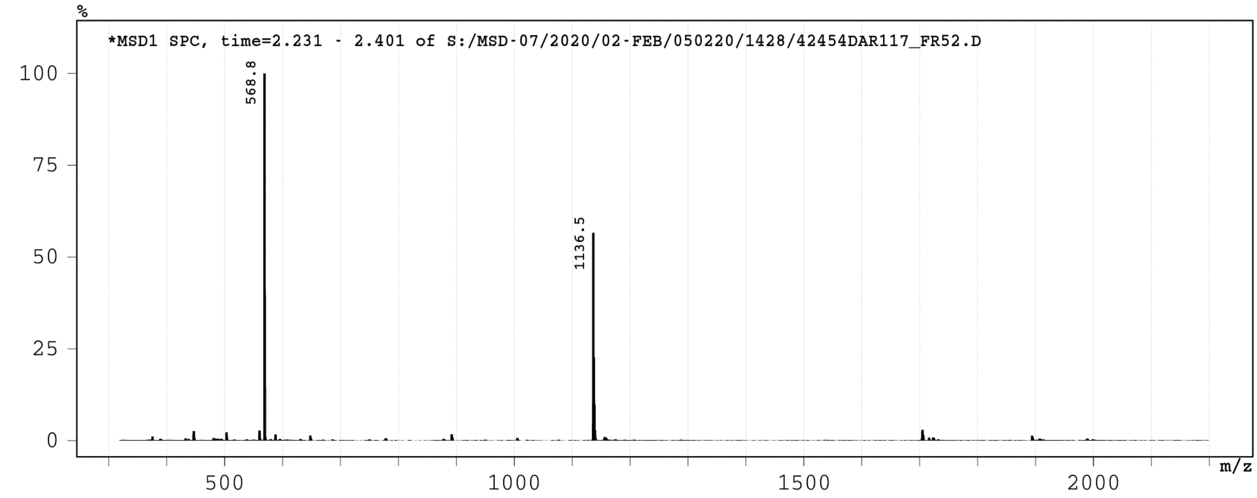


Signal 1: DAD1 A, Sig=220,4 Ref=off

| Peak # | RT [min] | Width [min] | Area | Area % |
|---|---|---|---|---|
| 1 | 2.211 | 1.88 | 56.2 | 0.9 |
| 2 | 2.275 | 0.20 | 6240.0 | 96.4 |
| 3 | 2.407 | 2.79 | 178.7 | 2.8 |

Data File : S:/MSD-07/2020/02-FEB/140220/1435/42454DAR118_FR2->

Injection Date : 15-Feb-20, 00:01:36
Sample Name : 42454DAR118_FR23-24
Remarks : Baseline corrected
Acq. Method : D:/METHODS/5-95-6M.M
column used : Je_559
Analysis Method : C:/CHEM32/1/METHODS/JPT-LCMS.M

Sample Info : 42454_118-Dr. Martine Darwish H-CTLQFEAAL-OH

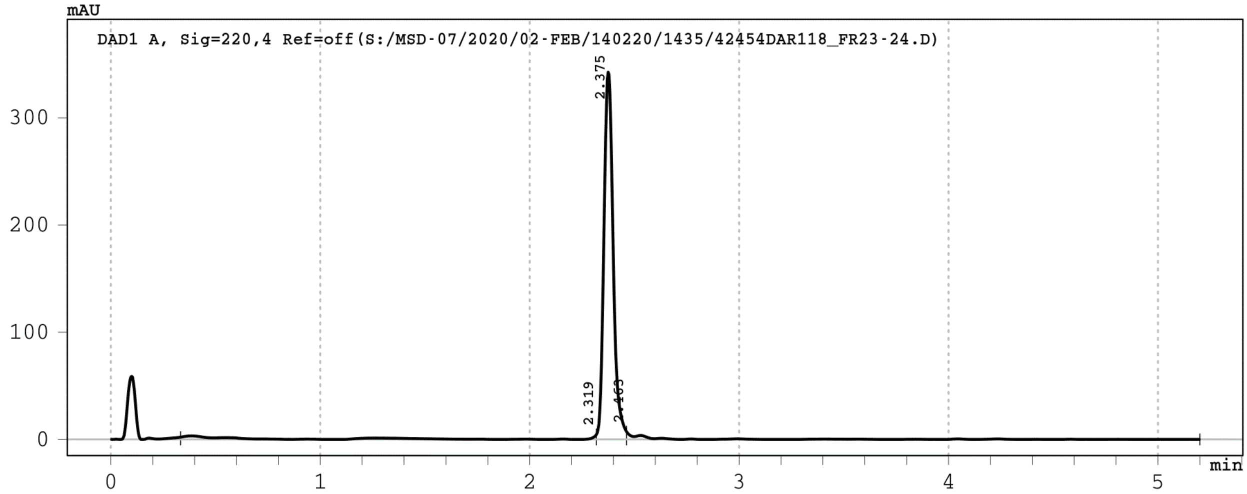


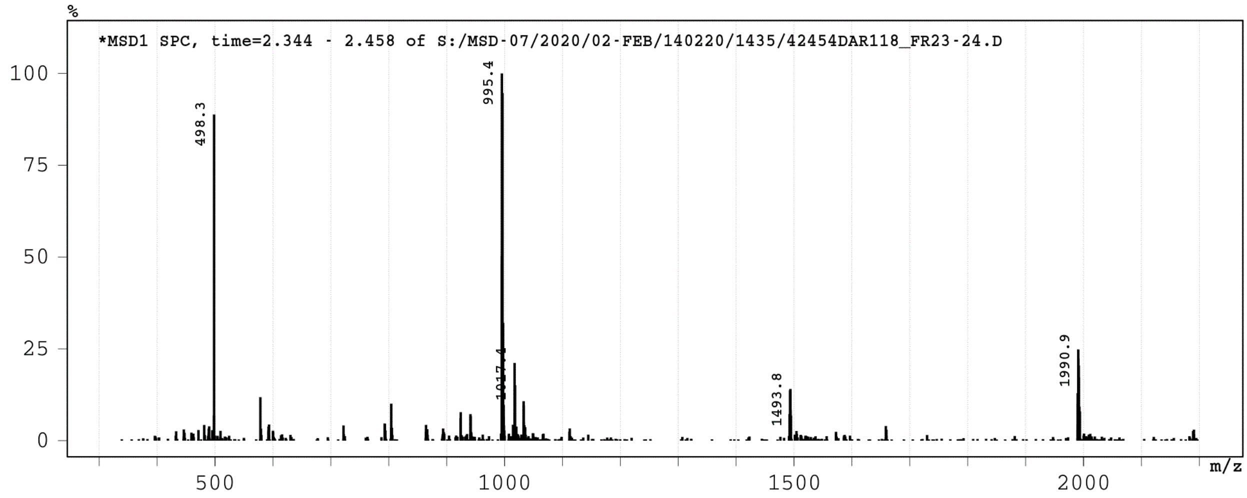


Signal 1: DAD1 A, Sig=220,4 Ref=off

| Peak # | RT [min] | Width [min] | Area | Area % |
|---|---|---|---|---|
| 1 | 2.319 | 1.99 | 74.9 | 6.4 |
| 2 | 2.375 | 0.14 | 1057.0 | 90.1 |
| 3 | 2.463 | 2.74 | 40.7 | 3.5 |

Data File : S:/MSD-07/2020/02-FEB/140220/1435/42454DAR119_FR2->

Injection Date : 15-Feb-20, 00:09:59
Sample Name : 42454DAR119_FR27-28
Remarks : Baseline corrected
Acq. Method : D:/METHODS/5-95-6M.M
column used : Je_559
Analysis Method : C:/CHEM32/1/METHODS/JPT-LCMS.M

Sample Info : 42454_119-Dr. Martine Darwish H-AALTFRRL-OH

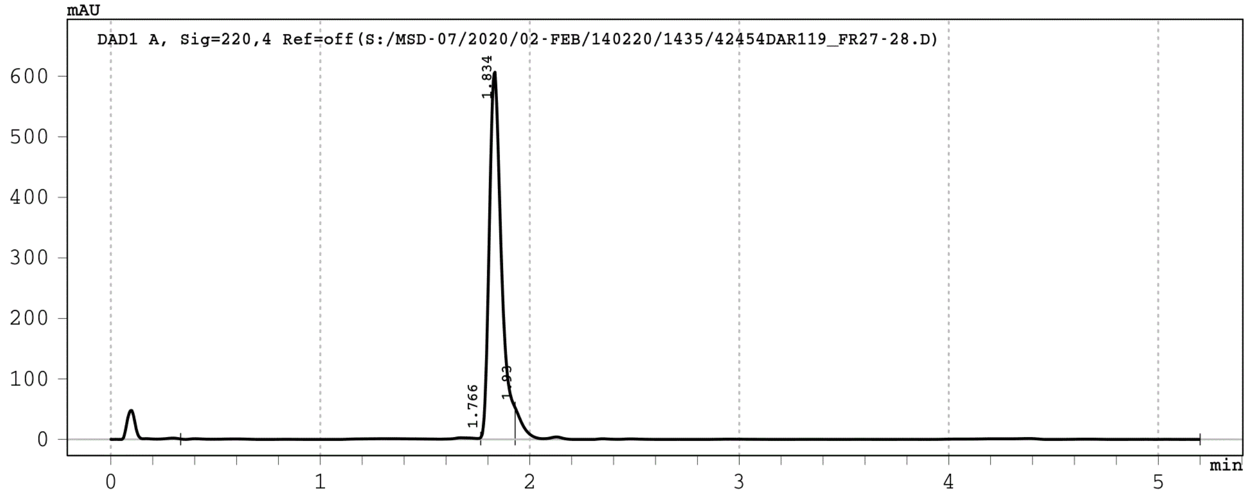


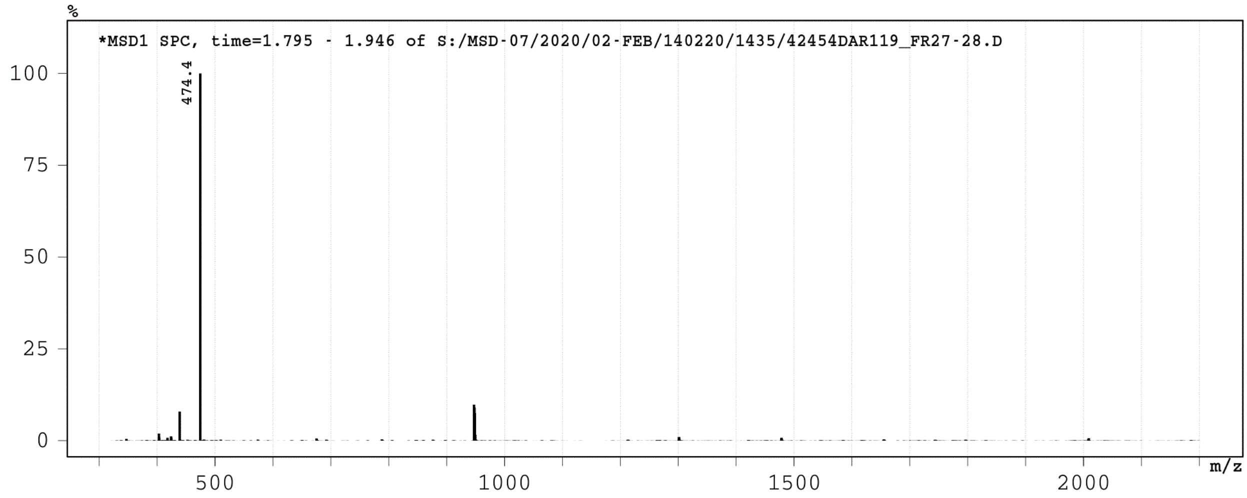


Signal 1: DAD1 A, Sig=220,4 Ref=off

| Peak # | RT [min] | Width [min] | Area | Area % |
|---|---|---|---|---|
| 1 | 1.766 | 1.43 | 55.5 | 2.1 |
| 2 | 1.834 | 0.16 | 2440.1 | 91.0 |
| 3 | 1.930 | 3.27 | 185.5 | 6.9 |

Data File: S:\MSD-07\2020\02-FEB\060220\142->

Injection Date : Thu, 6. Feb. 2020
Sample Name : 240120SG1-87|42454Dar120
Acq Operator : SYSTEM
Acq. Method : D:\Data\2020\02-FEB\060220\1429\5-95-6M.M
column used :
Analysis Method : C:\Chem32\1\METHODS\DEF_LC.M

Sample Info : 42454_120-Dr. Martine Darwish|H-CGLVPLARL-OH

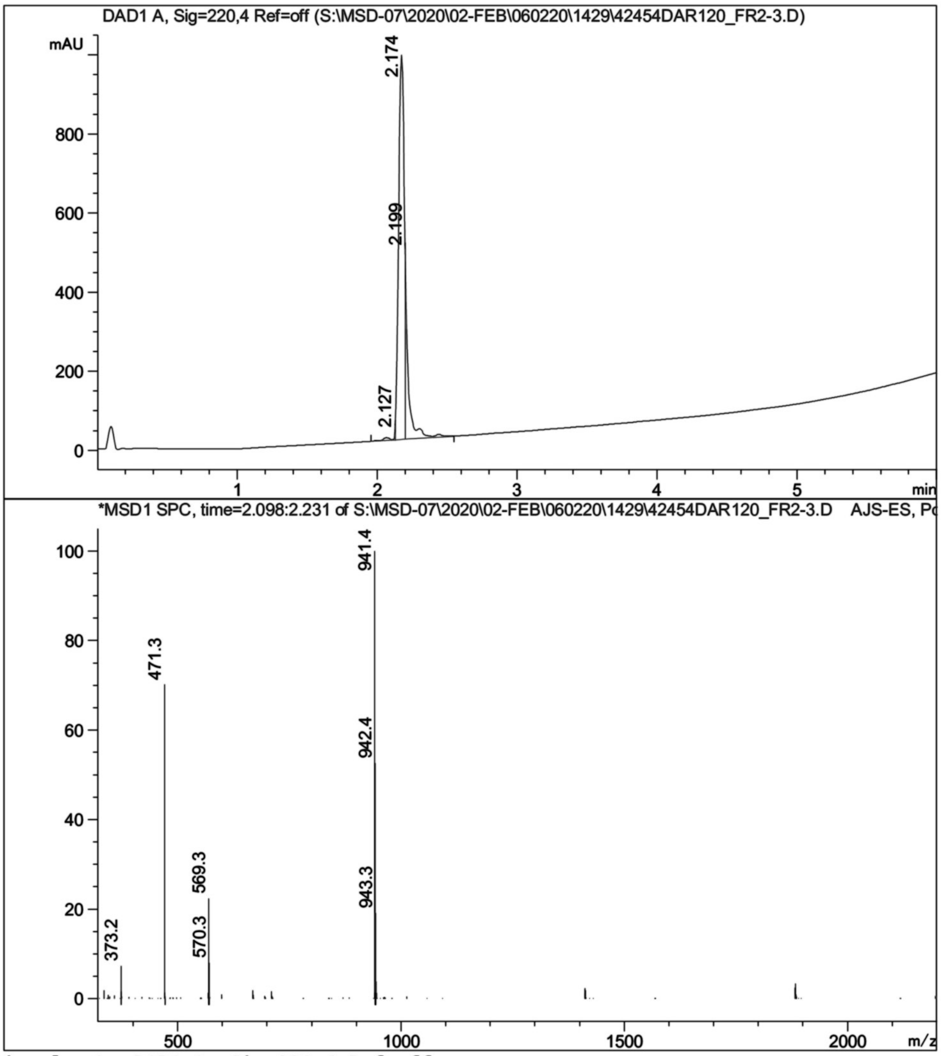


Signal 1: DAD1 A, Sig=220,4 Ref=off

| Peak # | RT [min] | Width [min] | Area | Area % |
|---|---|---|---|---|
| 1 | 2.127 | 0.0 | 47.8 | 1.4 |
| 2 | 2.174 | 0.0 | 2664.3 | 80.6 |
| 3 | 2.199 | 0.0 | 595.3 | 18.0 |

*** End of Report ***

| | | |
|---|---|---|
| Data File | : | S:/MSD-07/2020/02-FEB/070220/1430/42454DAR121_FR3-> |
| Injection Date | : | 07-Feb-20, 17:16:14 |
| Sample Name | : | 42454DAR121_FR35-36 |
| Remarks | : | Baseline corrected |
| Acq. Method | : | D:/METHODS/5-95-6M.M |
| column used | : | Je_559 |
| Analysis Method | : | C:/CHEM32/1/METHODS/JPT-LCMS.M |
| Sample Info | : | 42454_121-Dr. Martine Darwish H-LNFLHNFITRI-OH |

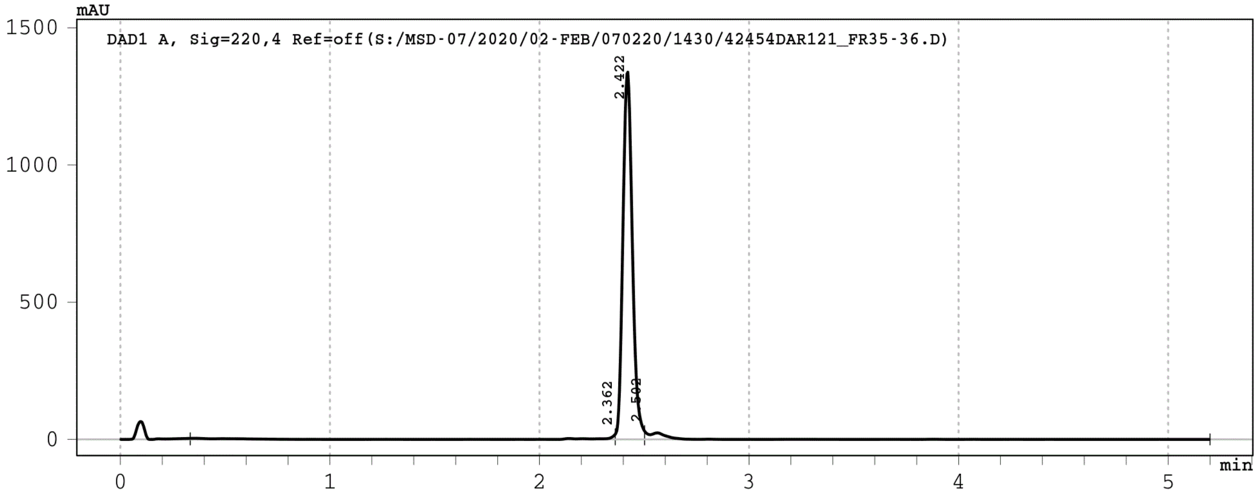


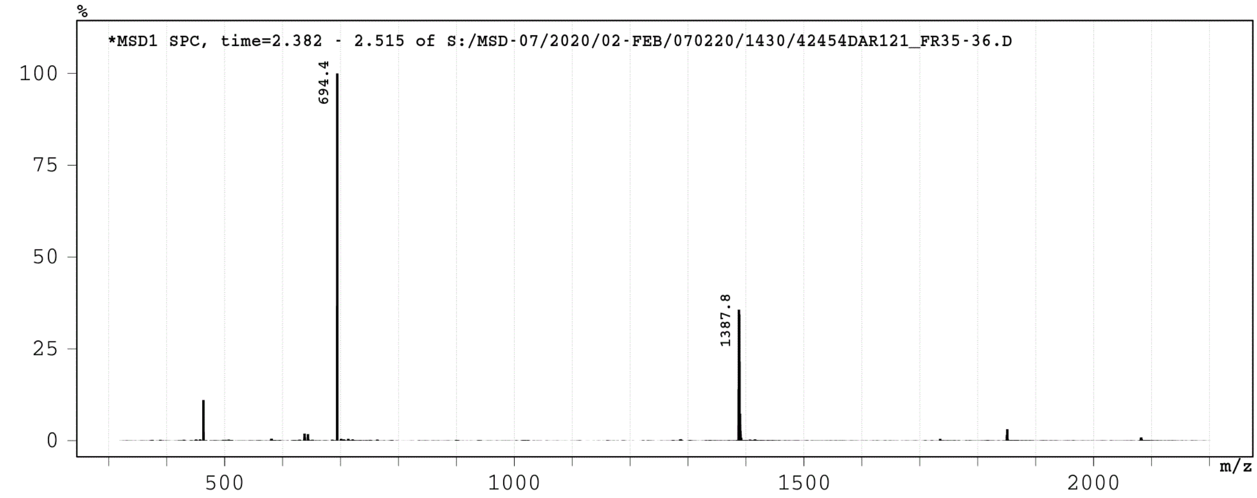


Signal 1: DAD1 A, Sig=220,4 Ref=off

| Peak # | RT [min] | Width [min] | Area | Area % |
|---|---|---|---|---|
| 1 | 2.362 | 2.03 | 109.8 | 2.4 |
| 2 | 2.422 | 0.14 | 4220.9 | 93.9 |
| 3 | 2.502 | 2.70 | 163.2 | 3.6 |

Data File : S:/MSD-07/2020/02-FEB/070220/1430/42454DAR122_FR4->

Injection Date : 07-Feb-20, 17:24:38
Sample Name : 42454DAR122_FR41
Remarks : Baseline corrected
Acq. Method : D:/METHODS/5-95-6M.M
column used : Je_559
Analysis Method : C:/CHEM32/1/METHODS/JPT-LCMS.M

Sample Info : 42454_122-Dr. Martine Darwish H-EALVNLNKL-OH

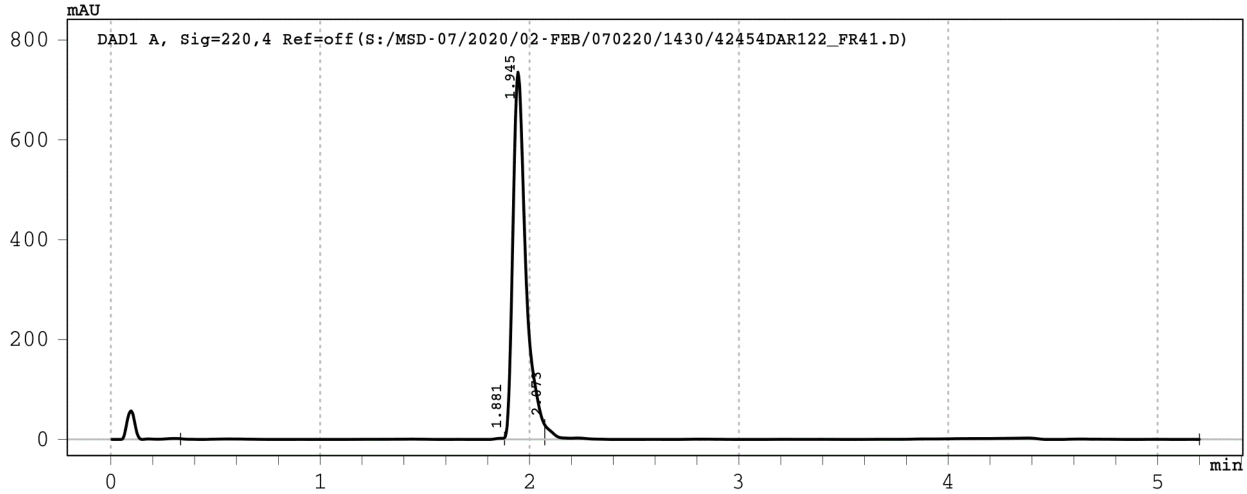


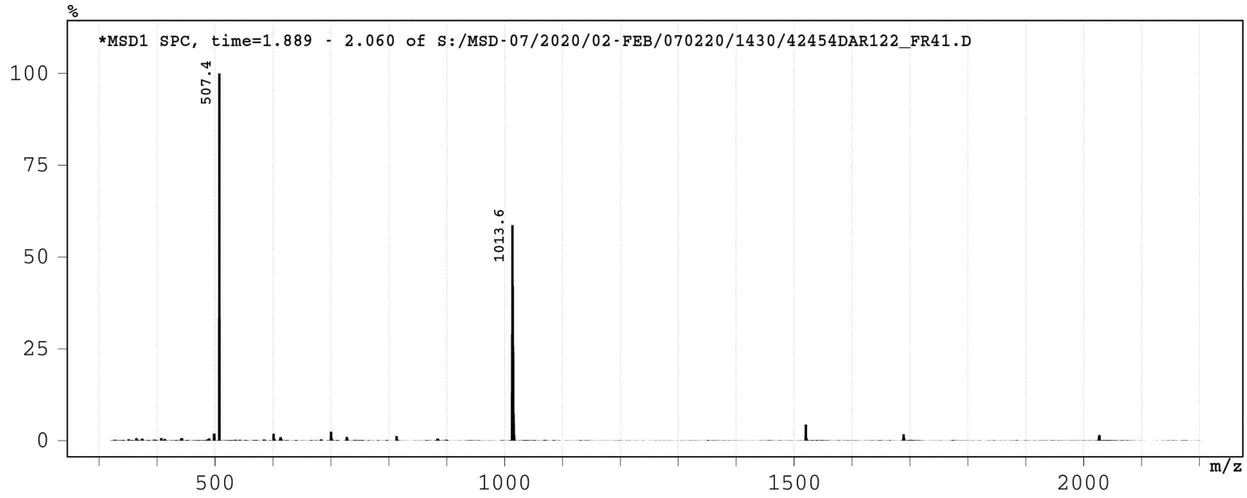


Signal 1: DAD1 A, Sig=220,4 Ref=off

| Peak # | RT [min] | Width [min] | Area | Area % |
|---|---|---|---|---|
| 1 | 1.881 | 1.55 | 24.8 | 0.7 |
| 2 | 1.945 | 0.19 | 3171.3 | 94.4 |
| 3 | 2.073 | 3.13 | 163.3 | 4.9 |

Data File : S:/MSD-07/2020/02-FEB/060220/1429/42454DAR123_FR7->

Injection Date : 06-Feb-20, 18:30:33
Sample Name : 42454DAR123_FR7-8
Remarks : Baseline corrected
Acq. Method : D:/METHODS/5-95-6M.M
column used : Je_559
Analysis Method : C:/CHEM32/1/METHODS/JPT-LCMS.M

Sample Info : 42454_123-Dr. Martine Darwish H-VMKAHRTSL-OH

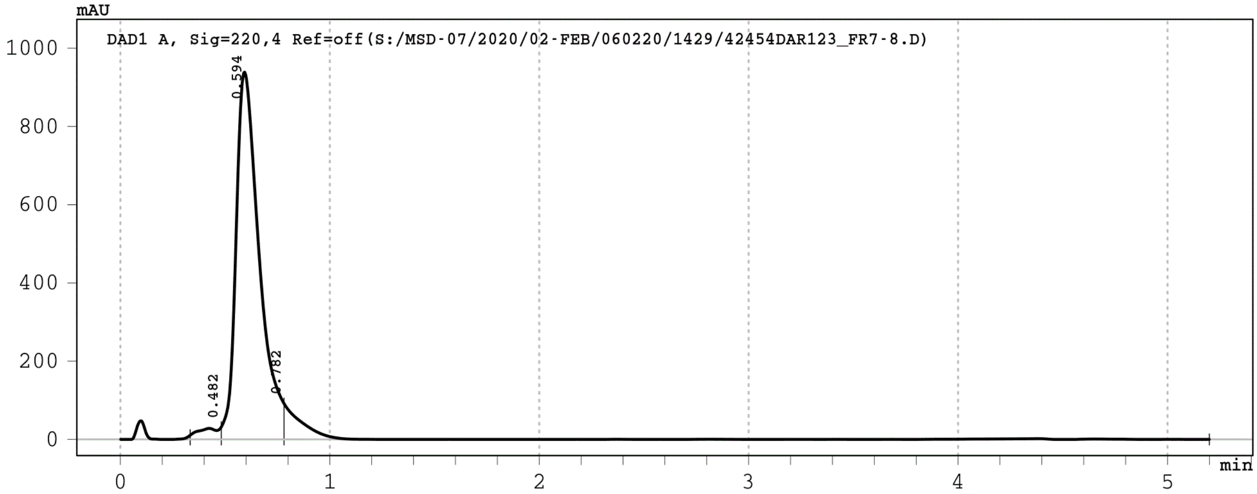


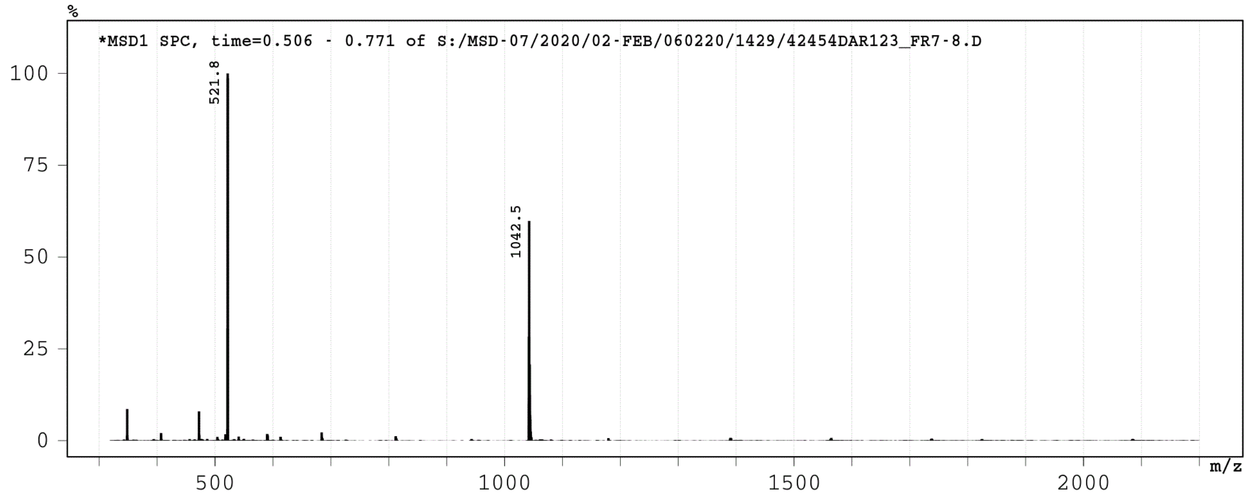


Signal 1: DAD1 A, Sig=220,4 Ref=off

| Peak # | RT [min] | Width [min] | Area | Area % |
|---|---|---|---|---|
| 1 | 0.482 | 0.15 | 204.9 | 2.6 |
| 2 | 0.594 | 0.30 | 7228.1 | 90.2 |
| 3 | 0.782 | 4.42 | 583.7 | 7.3 |

Data File : S:/MSD-07/2020/02-FEB/070220/1430/42454DAR124_FR4->

Injection Date : 07-Feb-20, 17:33:02
Sample Name : 42454DAR124_FR48-50
Remarks : Baseline corrected
Acq. Method : D:/METHODS/5-95-6M.M
column used : Je_559
Analysis Method : C:/CHEM32/1/METHODS/JPT-LCMS.M

Sample Info : 42454_124-Dr. Martine Darwish H-SVFGQSNDL-OH

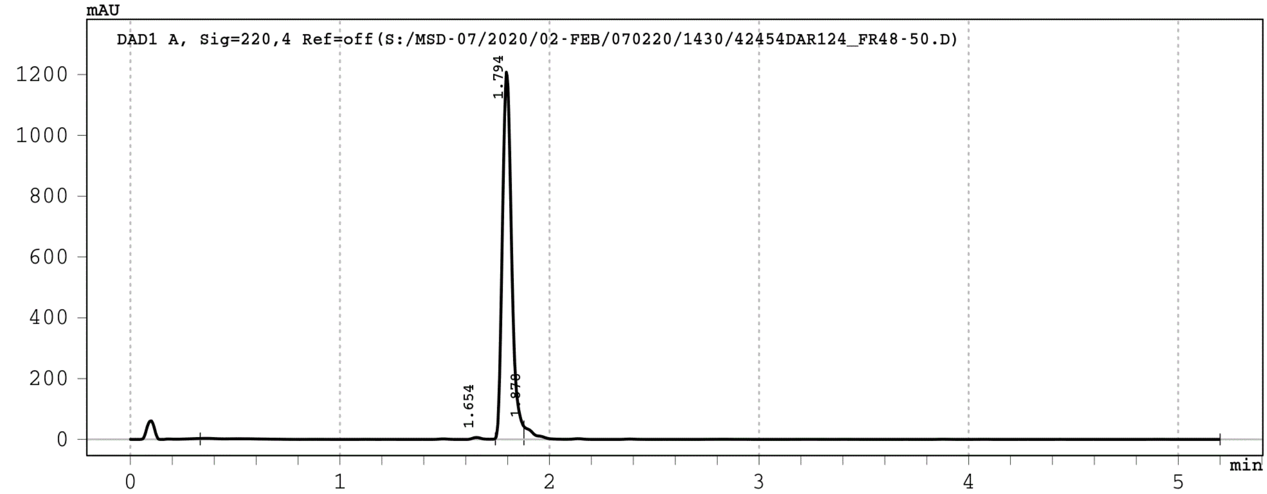


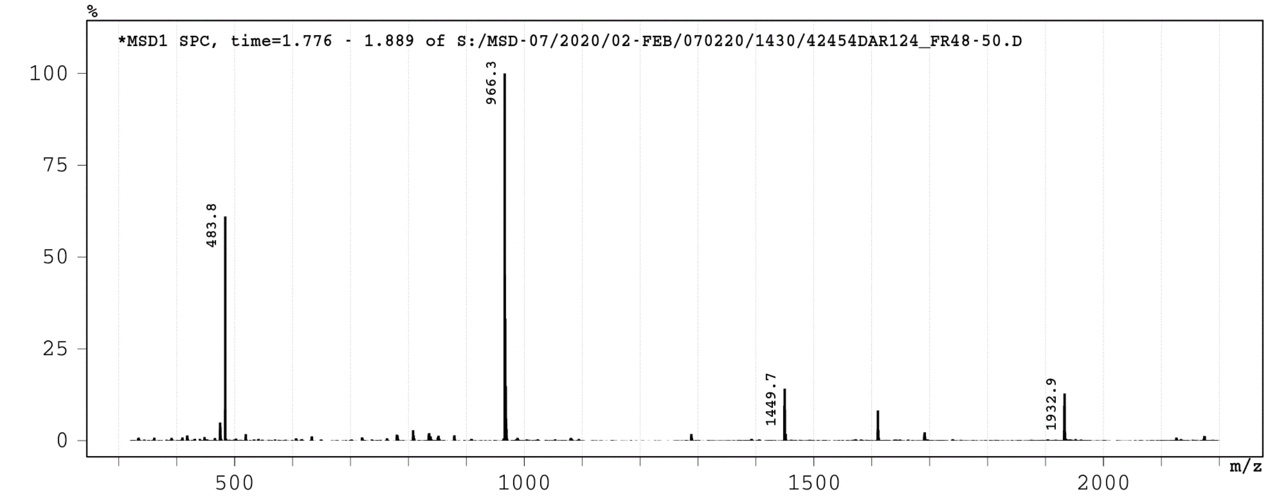


Signal 1: DAD1 A, Sig=220,4 Ref=off

| Peak # | RT [min] | Width [min] | Area | Area % |
|---|---|---|---|---|
| 1 | 1.654 | 1.41 | 71.7 | 1.8 |
| 2 | 1.794 | 0.14 | 3706.5 | 94.2 |
| 3 | 1.878 | 3.32 | 156.3 | 4.0 |

Data File : S:/MSD-07/2020/02-FEB/110220/1433/42454DAR125_FR3->

Injection Date : 12-Feb-20, 00:09:59
Sample Name : 42454DAR125_FR39-41
Remarks : Baseline corrected
Acq. Method : D:/METHODS/5-95-6M.M
column used : Je_559
Analysis Method : C:/CHEM32/1/METHODS/JPT-LCMS.M

Sample Info : 42454_125-Dr. Martine Darwish H-SLRSFQAL-OH

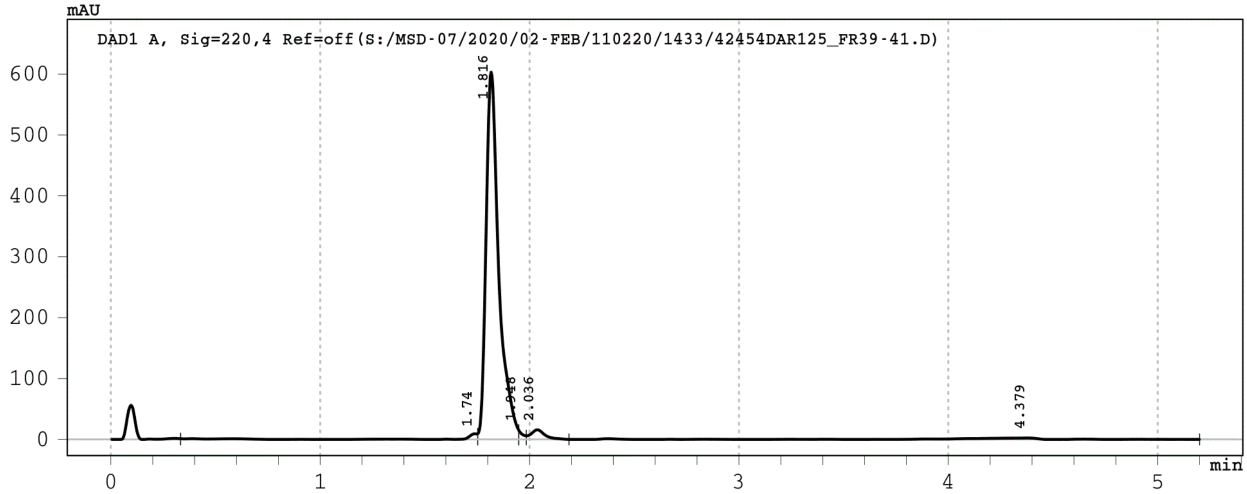


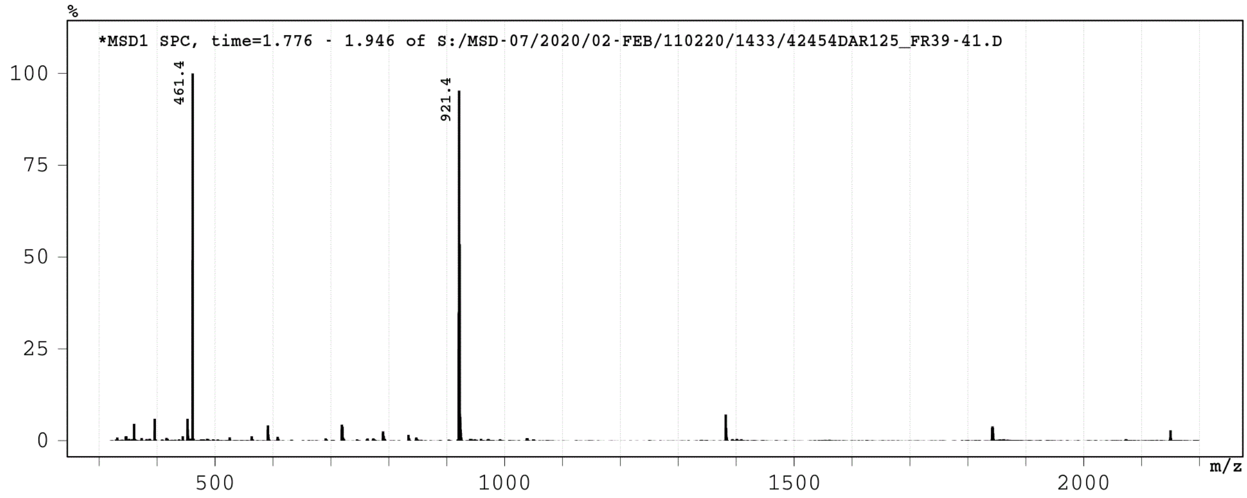


Signal 1: DAD1 A, Sig=220,4 Ref=off

| Peak # | RT [min] | Width [min] | Area | Area % |
|---|---|---|---|---|
| 1 | 1.740 | 1.42 | 51.4 | 1.9 |
| 2 | 1.816 | 0.20 | 2561.4 | 92.4 |
| 3 | 1.948 | 0.04 | 19.1 | 0.7 |
| 4 | 2.036 | 0.20 | 73.9 | 2.7 |
| 5 | 4.379 | 3.01 | 67.6 | 2.4 |

Data File : S:/MSD-07/2020/02-FEB/070220/1430/42454DAR126_FR6->

Injection Date : 07-Feb-20, 17:41:24
Sample Name : 42454DAR126_FR62
Remarks : Baseline corrected
Acq. Method : D:/METHODS/5-95-6M.M
column used : Je_559
Analysis Method : C:/CHEM32/1/METHODS/JPT-LCMS.M

Sample Info : 42454_126-Dr. Martine Darwish H-FSLSFQHPV-OH

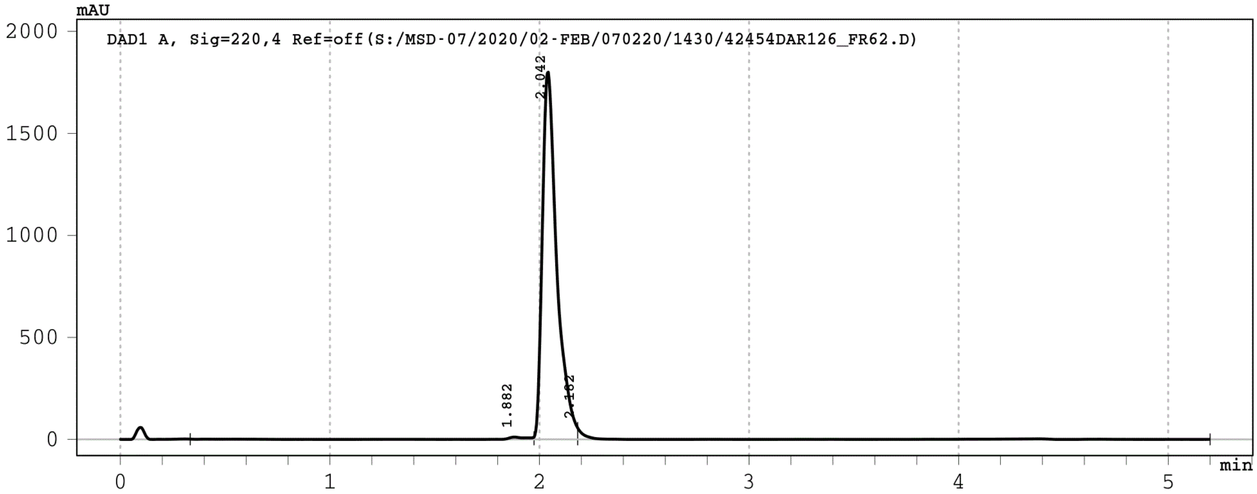


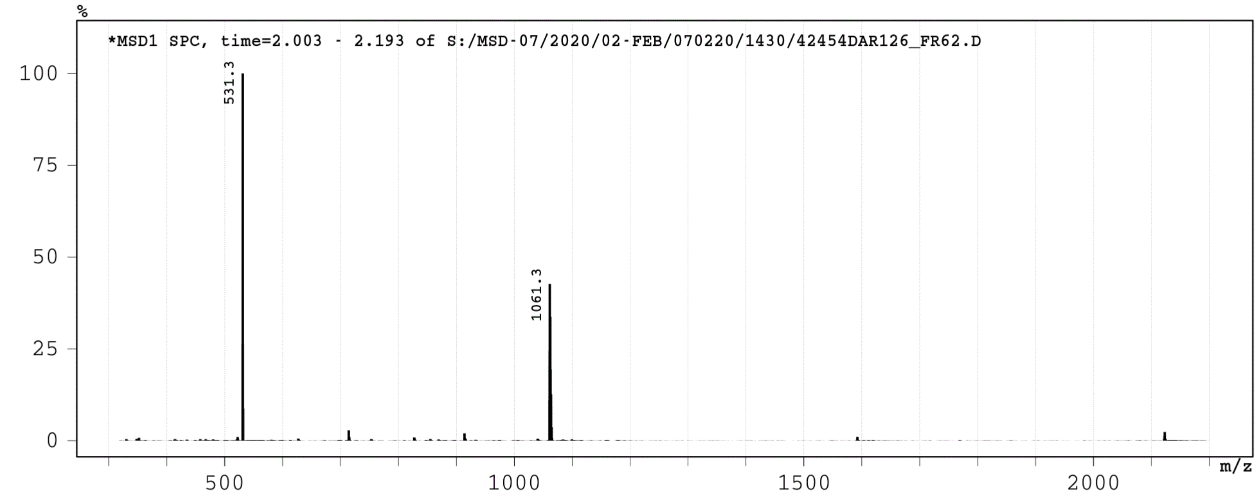


Signal 1: DAD1 A, Sig=220,4 Ref=off

| Peak # | RT [min] | Width [min] | Area | Area % |
|---|---|---|---|---|
| 1 | 1.882 | 1.64 | 95.8 | 1.1 |
| 2 | 2.042 | 0.21 | 8656.1 | 96.9 |
| 3 | 2.182 | 3.02 | 182.3 | 2.0 |

Data File : S:/MSD-07/2020/02-FEB/110220/1433/42454DAR127_FR5->

Injection Date : 12-Feb-20, 00:18:23
Sample Name : 42454DAR127_FR53-55
Remarks : Baseline corrected
Acq. Method : D:/METHODS/5-95-6M.M
column used : Je_559
Analysis Method : C:/CHEM32/1/METHODS/JPT-LCMS.M

Sample Info : 42454_127-Dr. Martine Darwish H-SSVLFEYM-OH

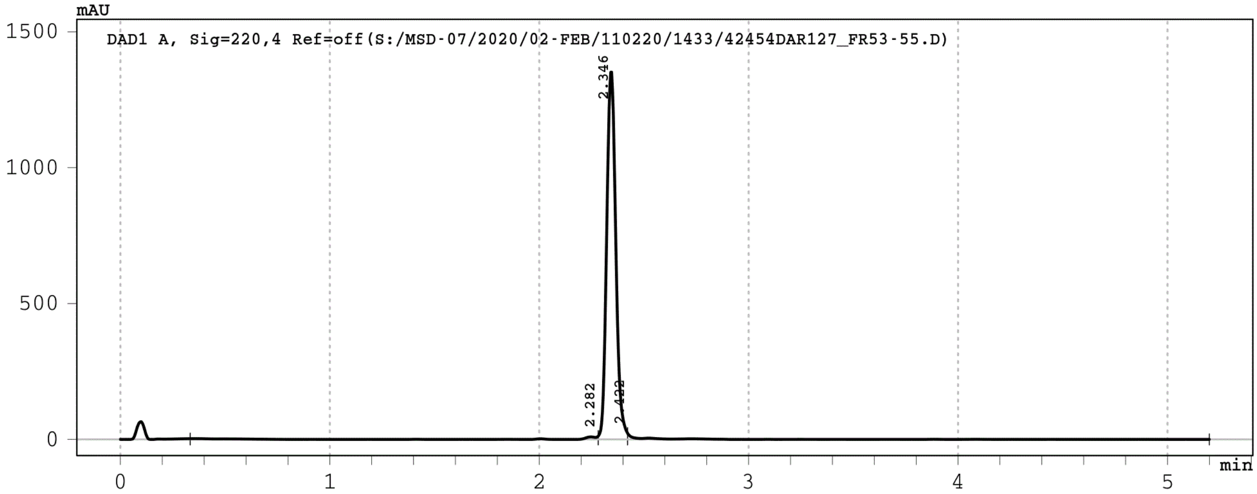


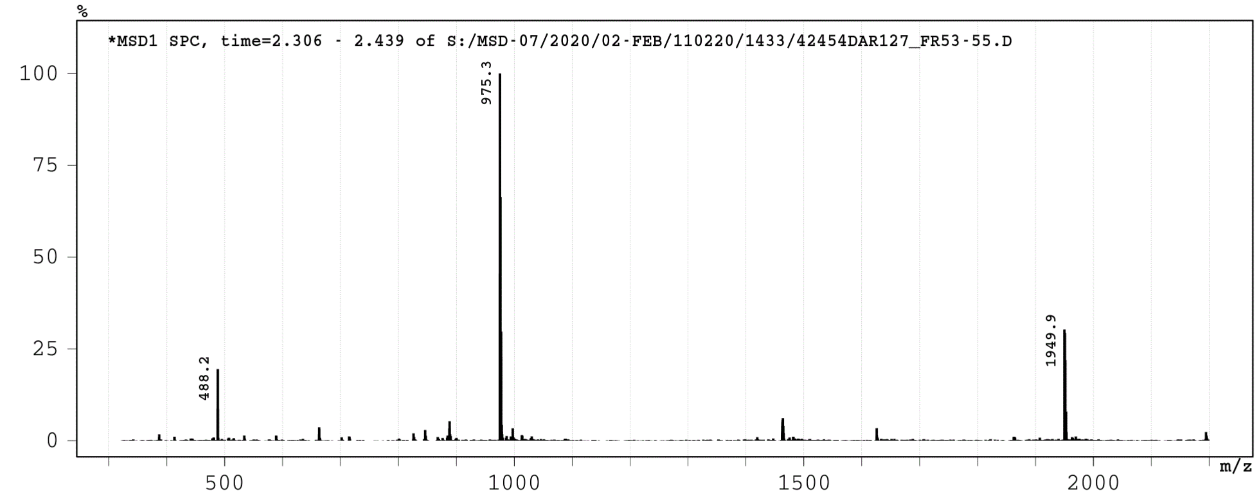


Signal 1: DAD1 A, Sig=220,4 Ref=off

| Peak # | RT [min] | Width [min] | Area | Area % |
|---|---|---|---|---|
| 1 | 2.282 | 1.95 | 83.8 | 2.0 |
| 2 | 2.346 | 0.14 | 4103.7 | 96.0 |
| 3 | 2.422 | 2.78 | 87.3 | 2.0 |

Data File : S:/MSD-07/2020/02-FEB/070220/1430/42454DAR128_FR7->

Injection Date : 07-Feb-20, 17:49:46
Sample Name : 42454DAR128_FR73-74
Remarks : Baseline corrected
Acq. Method : D:/METHODS/5-95-6M.M
column used : Je_559
Analysis Method : C:/CHEM32/1/METHODS/JPT-LCMS.M

Sample Info : 42454_128-Dr. Martine Darwish H-SVYRHQQPT-OH

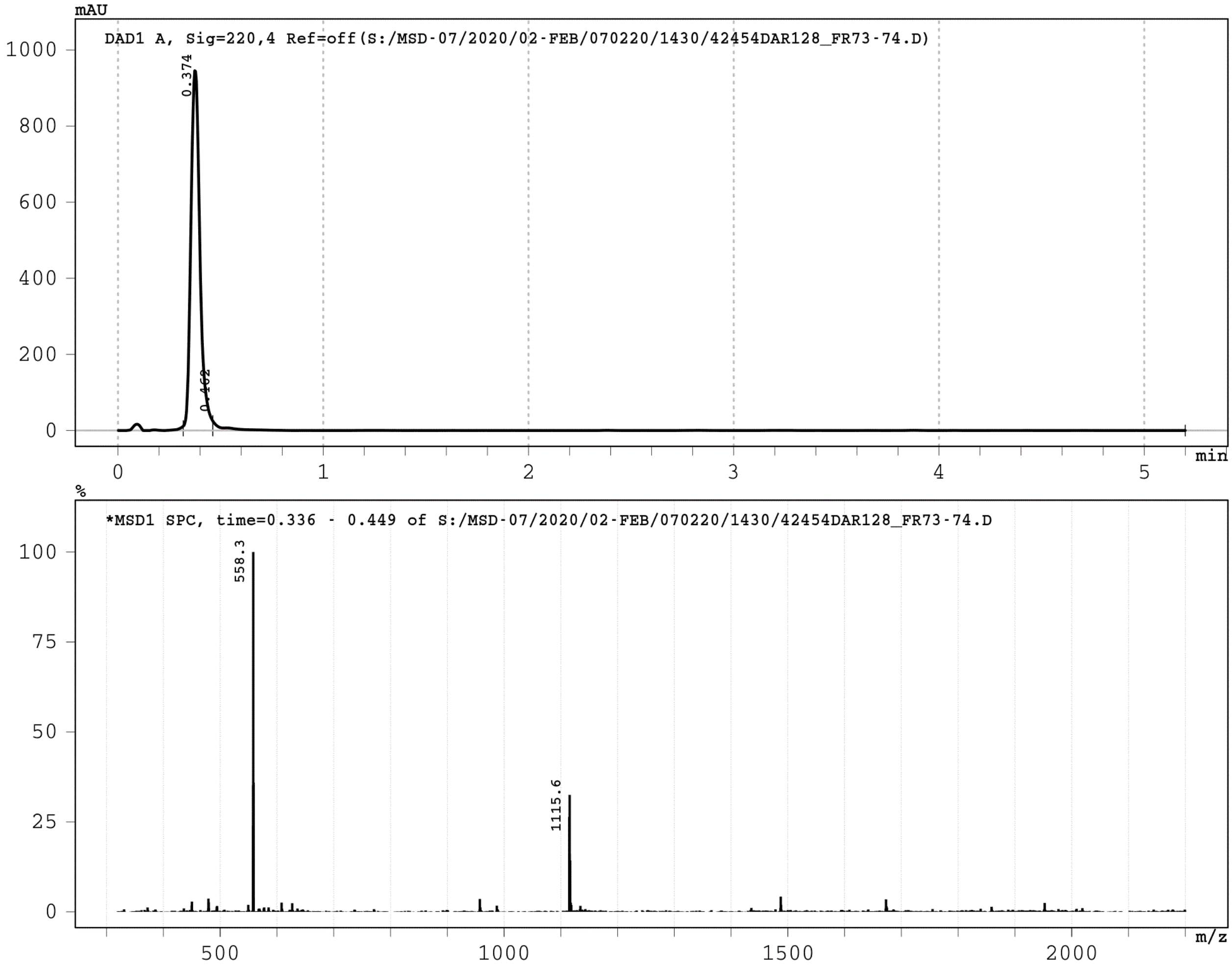


Signal 1: DAD1 A, Sig=220,4 Ref=off

| Peak # | RT [min] | Width [min] | Area | Area % |
|---|---|---|---|---|
| 1 | 0.374 | 0.14 | 2910.7 | 96.7 |
| 2 | 0.462 | 4.74 | 99.1 | 3.3 |

Data File : S:/MSD-07/2020/02-FEB/060220/1429/42454DAR129_FR1->

Injection Date : 06-Feb-20, 18:38:56
Sample Name : 42454DAR129_FR16-18
Remarks : Baseline corrected
Acq. Method : D:/METHODS/5-95-6M.M
column used : Je_559
Analysis Method : C:/CHEM32/1/METHODS/JPT-LCMS.M

Sample Info : 42454_129-Dr. Martine Darwish H-CAMGFFLQI-OH

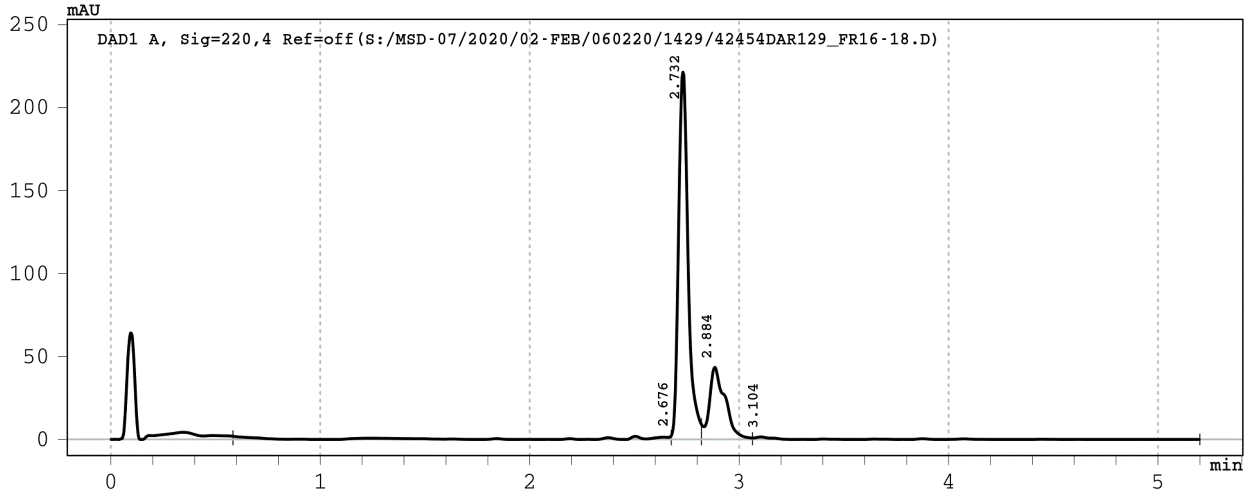


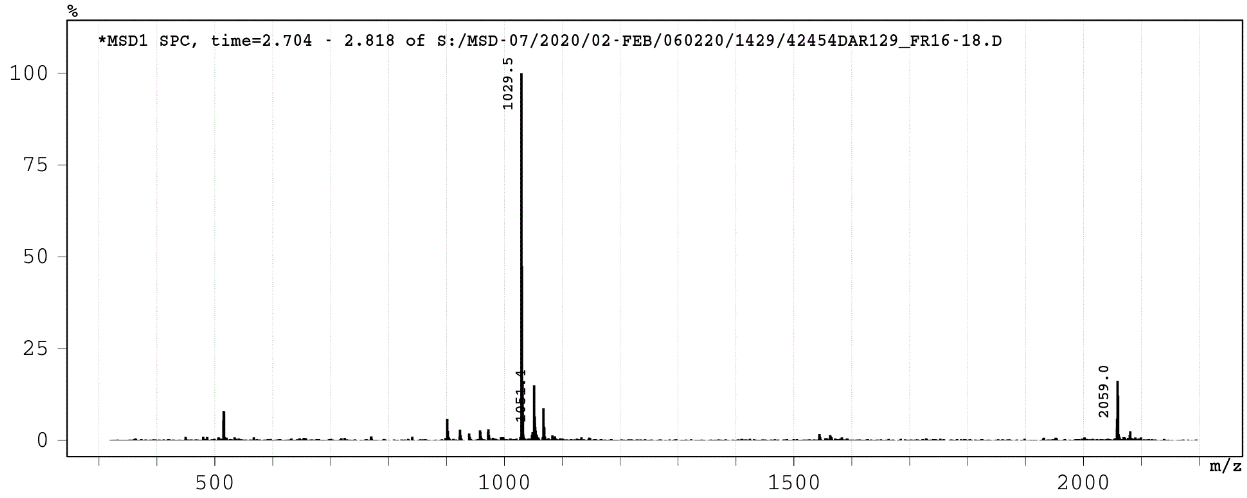


Signal 1: DAD1 A, Sig=220,4 Ref=off

| Peak # | RT [min] | Width [min] | Area | Area % |
|---|---|---|---|---|
| 1 | 2.676 | 2.09 | 45.9 | 4.6 |
| 2 | 2.732 | 0.14 | 704.9 | 71.0 |
| 3 | 2.884 | 0.24 | 229.4 | 23.1 |
| 4 | 3.104 | 2.14 | 13.0 | 1.3 |

Data File : S:/MSD-07/2020/02-FEB/180220/1438/42454DAR130_FR3->

Injection Date : 18-Feb-20, 09:58:03
Sample Name : 42454DAR130_FR30
Remarks : Baseline corrected
Acq. Method : D:/METHODS/5-95-6M.M
column used : Je_559
Analysis Method : C:/CHEM32/1/METHODS/JPT-LCMS.M

Sample Info : 42454_130-Dr. Martine Darwish H-QQHTFAQV-OH

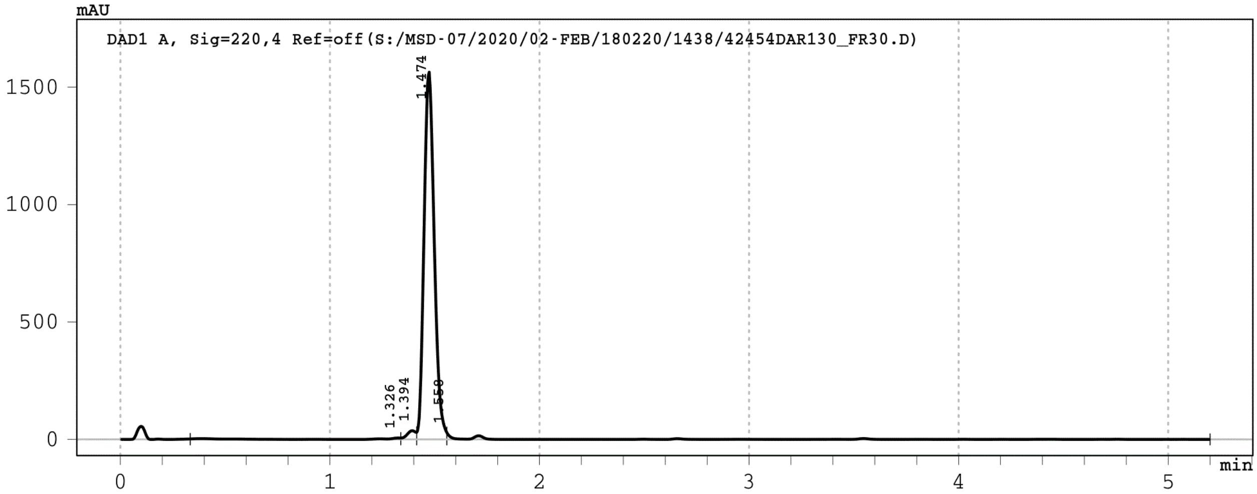


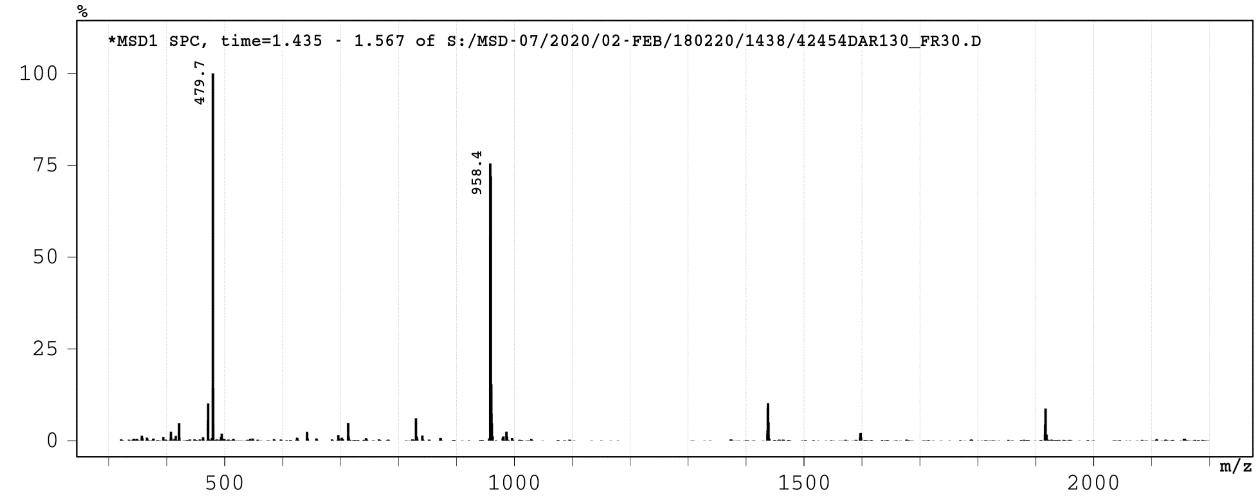


Signal 1: DAD1 A, Sig=220,4 Ref=off

| Peak # | RT [min] | Width [min] | Area | Area % |
|---|---|---|---|---|
| 1 | 1.326 | 1.00 | 68.7 | 1.2 |
| 2 | 1.394 | 0.08 | 107.7 | 1.9 |
| 3 | 1.474 | 0.14 | 5317.9 | 94.7 |
| 4 | 1.558 | 3.64 | 123.6 | 2.2 |

Data File : S:/MSD-07/2020/02-FEB/110220/1433/42454DAR131_FR6->

Injection Date : 12-Feb-20, 00:26:47
Sample Name : 42454DAR131_FR64-65
Remarks : Baseline corrected
Acq. Method : D:/METHODS/5-95-6M.M
column used : Je_559
Analysis Method : C:/CHEM32/1/METHODS/JPT-LCMS.M

Sample Info : 42454_131-Dr. Martine Darwish H-VGYCETLL-OH

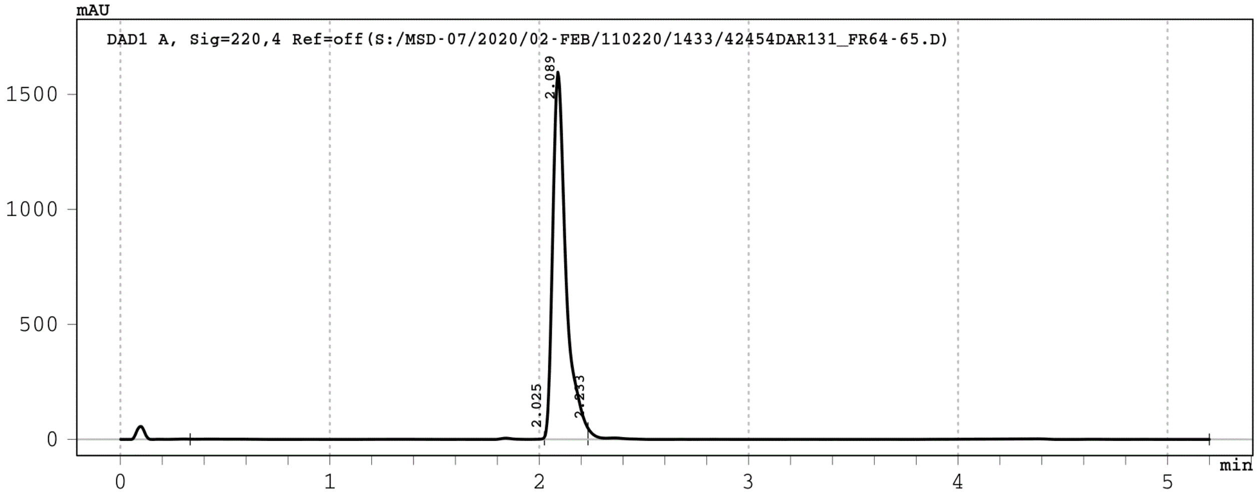


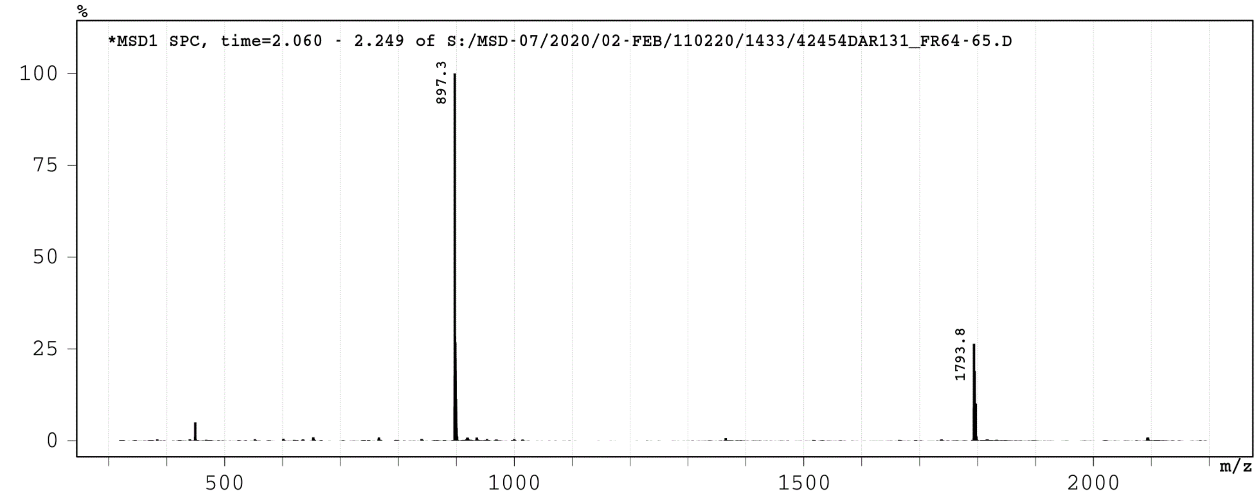


Signal 1: DAD1 A, Sig=220,4 Ref=off

| Peak # | RT [min] | Width [min] | Area | Area % |
|---|---|---|---|---|
| 1 | 2.025 | 1.69 | 52.9 | 0.7 |
| 2 | 2.089 | 0.21 | 7053.4 | 96.8 |
| 3 | 2.233 | 2.97 | 183.4 | 2.5 |

Data File : S:/MSD-07/2020/02-FEB/070220/1430/42454DAR132_FR3->

Injection Date : 07-Feb-20, 17:58:10
Sample Name : 42454DAR132_FR3-4
Remarks : Baseline corrected
Acq. Method : D:/METHODS/5-95-6M.M
column used : Je_559
Analysis Method : C:/CHEM32/1/METHODS/JPT-LCMS.M

Sample Info : 42454_132-Dr. Martine Darwish H-TGYVERSPL-OH

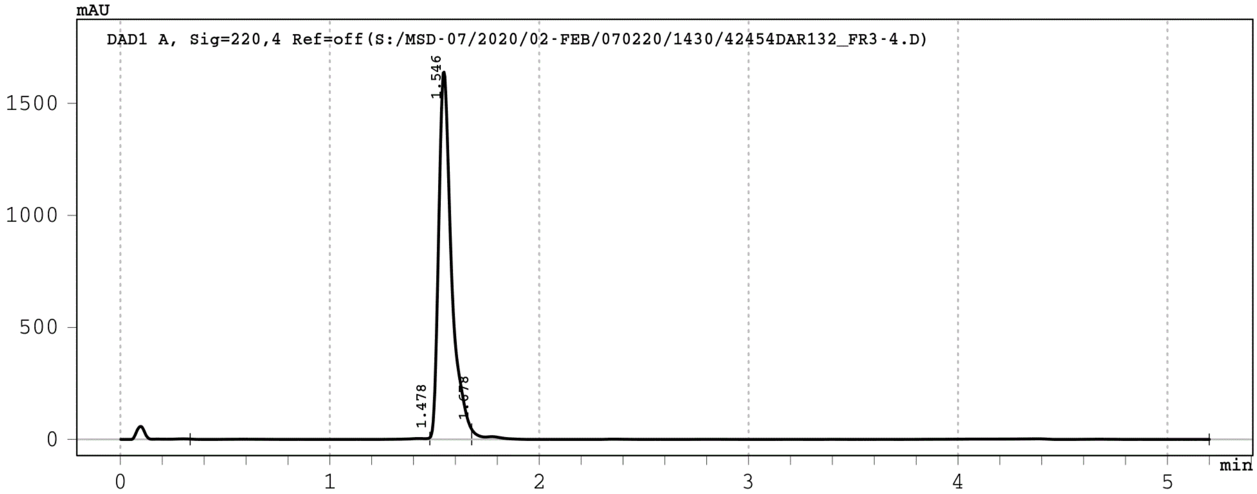


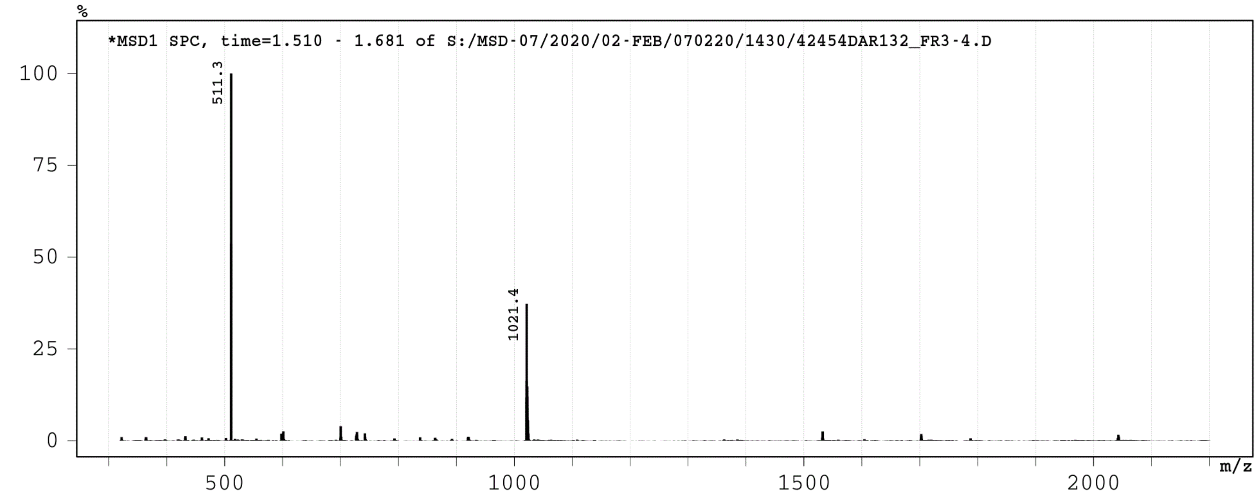


Signal 1: DAD1 A, Sig=220,4 Ref=off

| Peak # | RT [min] | Width [min] | Area | Area % |
|---|---|---|---|---|
| 1 | 1.478 | 1.14 | 39.1 | 0.5 |
| 2 | 1.546 | 0.20 | 7033.6 | 96.5 |
| 3 | 1.678 | 3.52 | 219.3 | 3.0 |

Data File : S:/MSD-07/2020/02-FEB/110220/1433/42454DAR133_FR8->

Injection Date : 12-Feb-20, 00:35:14
Sample Name : 42454DAR133_FR81-83
Remarks : Baseline corrected
Acq. Method : D:/METHODS/5-95-6M.M
column used : Je_559
Analysis Method : C:/CHEM32/1/METHODS/JPT-LCMS.M

Sample Info : 42454_133-Dr. Martine Darwish H-ATINFRRL-OH

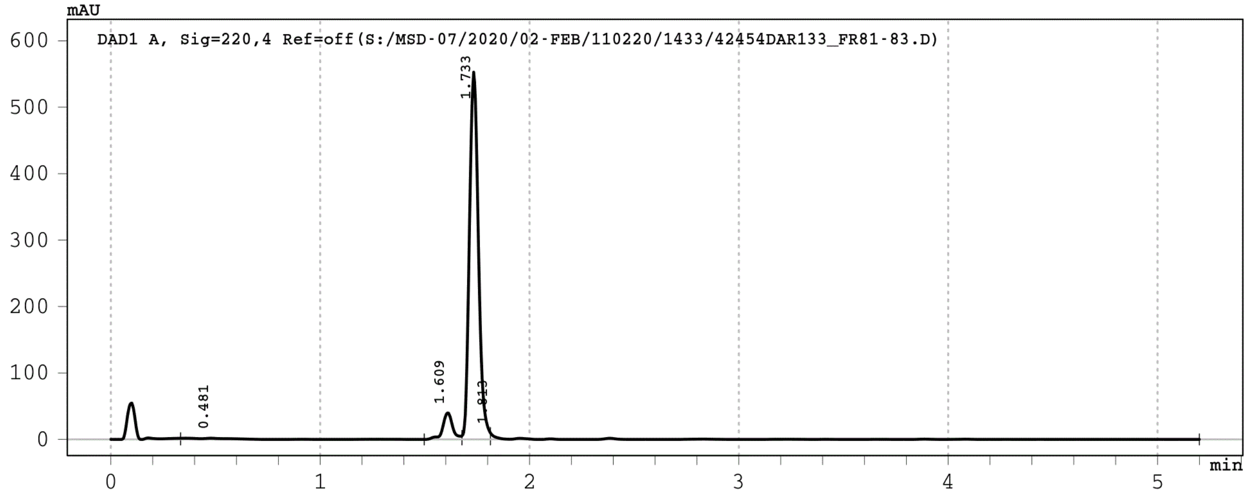


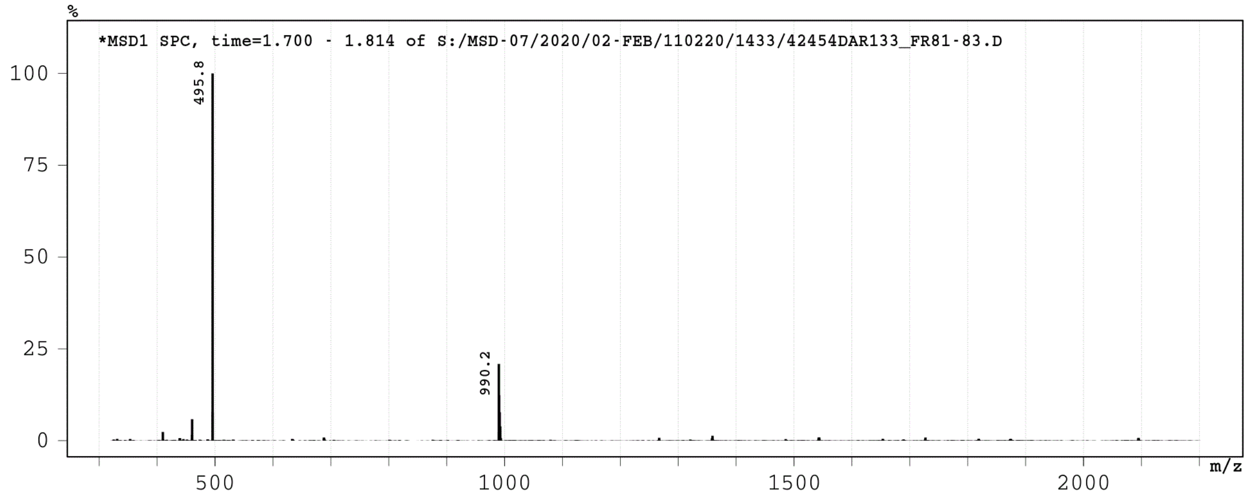


Signal 1: DAD1 A, Sig=220,4 Ref=off

| Peak # | RT [min] | Width [min] | Area | Area % |
|---|---|---|---|---|
| 1 | 0.481 | 1.16 | 28.3 | 1.5 |
| 2 | 1.609 | 0.18 | 130.8 | 7.1 |
| 3 | 1.733 | 0.14 | 1653.6 | 89.4 |
| 4 | 1.813 | 3.39 | 37.9 | 2.0 |

Data File : S:/MSD-07/2020/02-FEB/070220/1430/42454DAR134_FR1->

Injection Date : 07-Feb-20, 18:06:33
Sample Name : 42454DAR134_FR18-19
Remarks : Baseline corrected
Acq. Method : D:/METHODS/5-95-6M.M
column used : Je_559
Analysis Method : C:/CHEM32/1/METHODS/JPT-LCMS.M

Sample Info : 42454_134-Dr. Martine Darwish H-YSLSHLKRV-OH

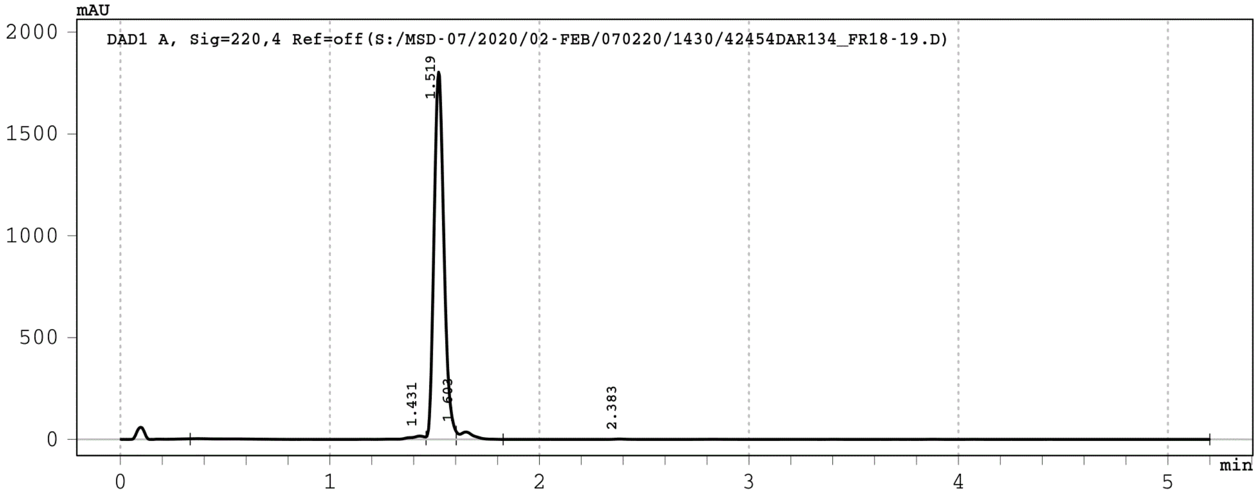


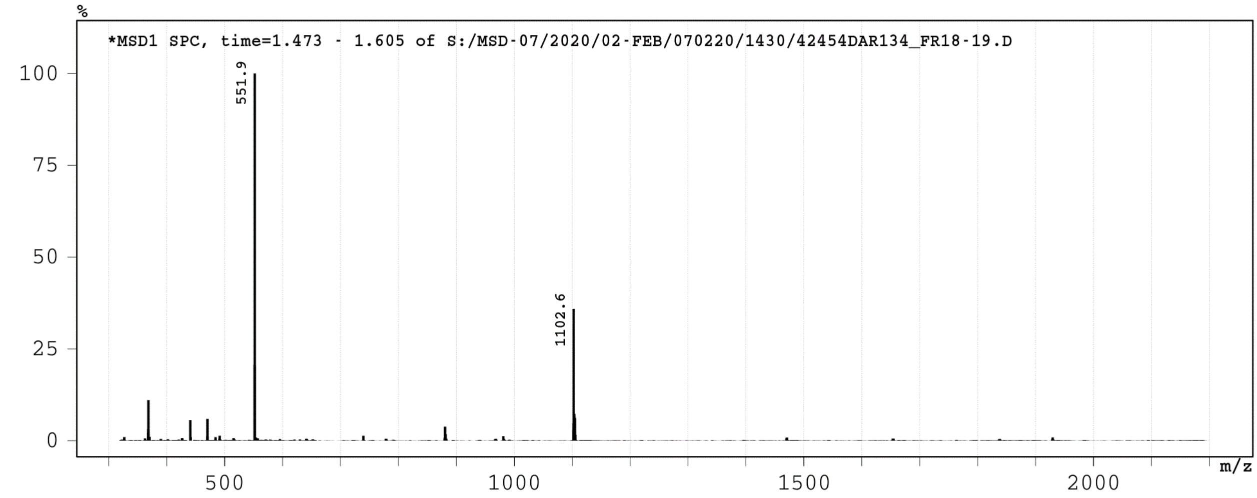


Signal 1: DAD1 A, Sig=220,4 Ref=off

| Peak # | RT [min] | Width [min] | Area | Area % |
|---|---|---|---|---|
| 1 | 1.431 | 1.13 | 134.5 | 2.1 |
| 2 | 1.519 | 0.14 | 5934.0 | 94.7 |
| 3 | 1.603 | 0.22 | 181.5 | 2.9 |
| 4 | 2.383 | 3.37 | 16.8 | 0.3 |

Data File : S:/MSD-07/2020/02-FEB/140220/1435/42454DAR135_FR4->

Injection Date : 15-Feb-20, 00:18:27
Sample Name : 42454DAR135_FR40-41
Remarks : Baseline corrected
Acq. Method : D:/METHODS/5-95-6M.M
column used : Je_559
Analysis Method : C:/CHEM32/1/METHODS/JPT-LCMS.M

Sample Info : 42454_135-Dr. Martine Darwish H-VLIQWLPAL-OH

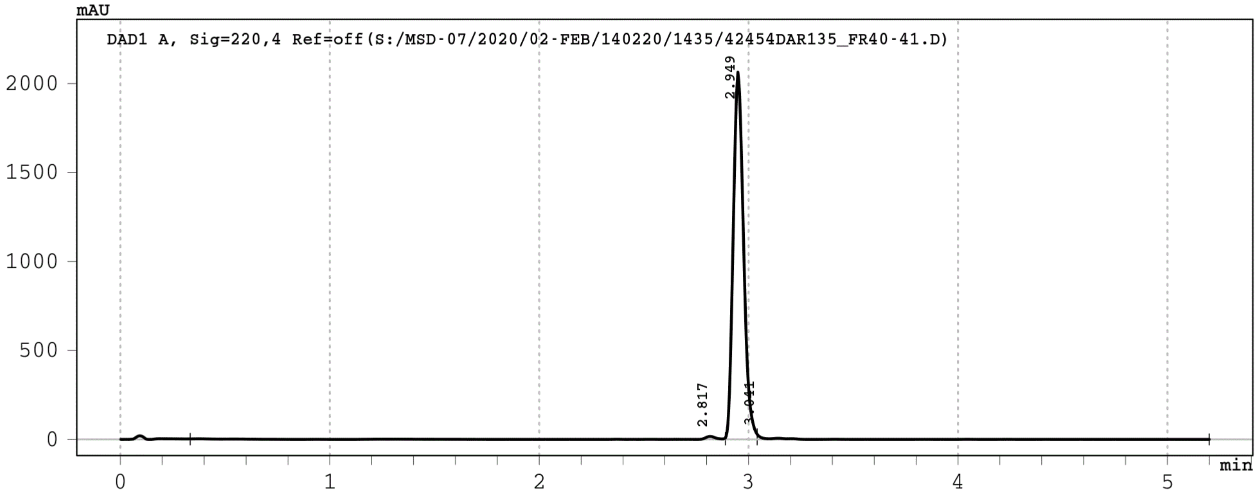


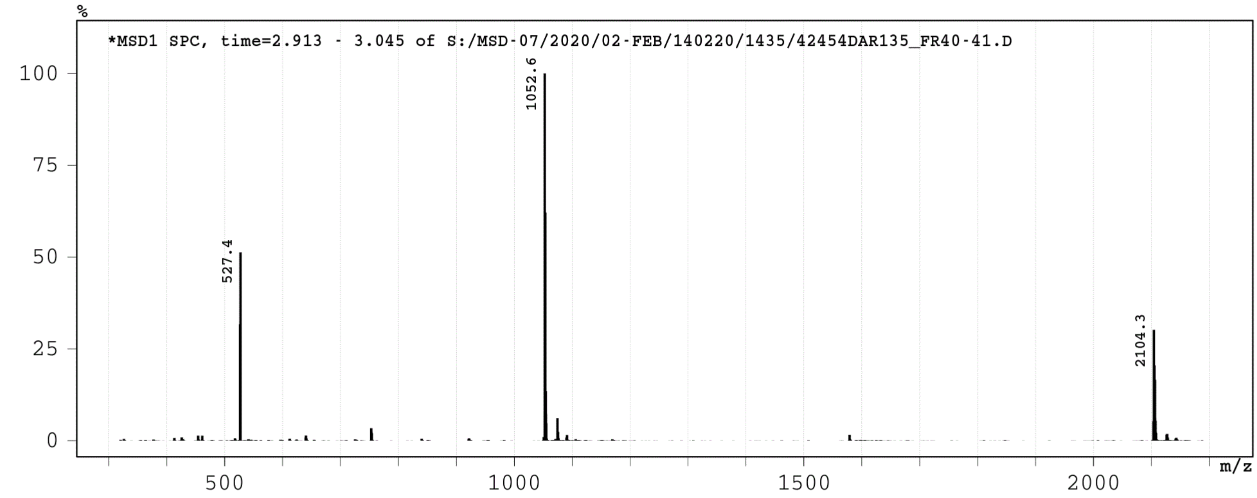


Signal 1: DAD1 A, Sig=220,4 Ref=off

| Peak # | RT [min] | Width [min] | Area | Area % |
|---|---|---|---|---|
| 1 | 2.817 | 2.56 | 136.8 | 1.9 |
| 2 | 2.949 | 0.15 | 7060.5 | 97.0 |
| 3 | 3.041 | 2.16 | 79.0 | 1.1 |

Data File : S:/MSD-07/2020/02-FEB/070220/1430/42454DAR136_FR3->

Injection Date : 07-Feb-20, 18:14:56
Sample Name : 42454DAR136_FR35-37
Remarks : Baseline corrected
Acq. Method : D:/METHODS/5-95-6M.M
column used : Je_559
Analysis Method : C:/CHEM32/1/METHODS/JPT-LCMS.M

Sample Info : 42454_136-Dr. Martine Darwish H-FNFWLHKLI-OH

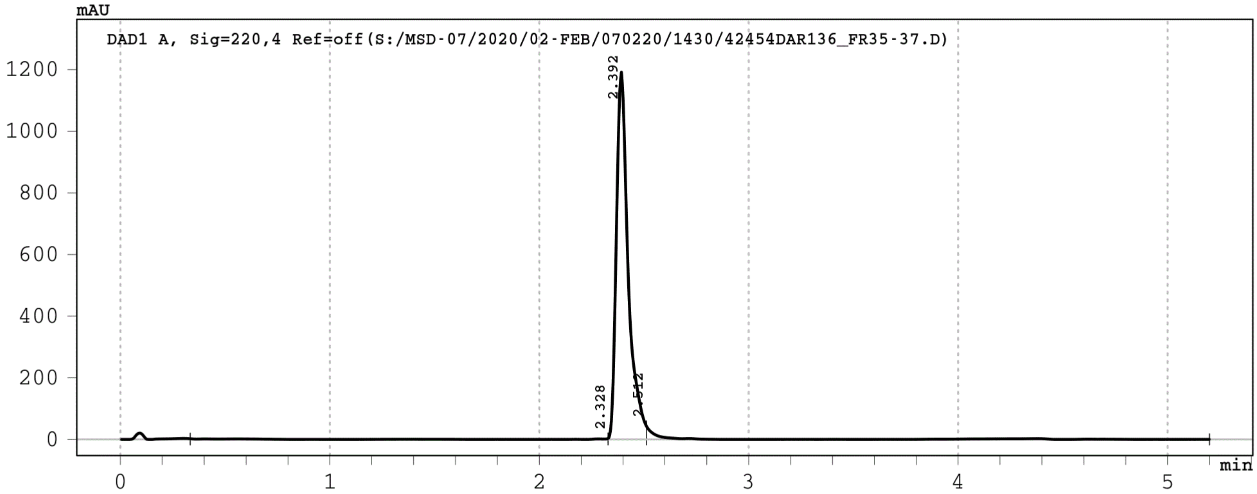


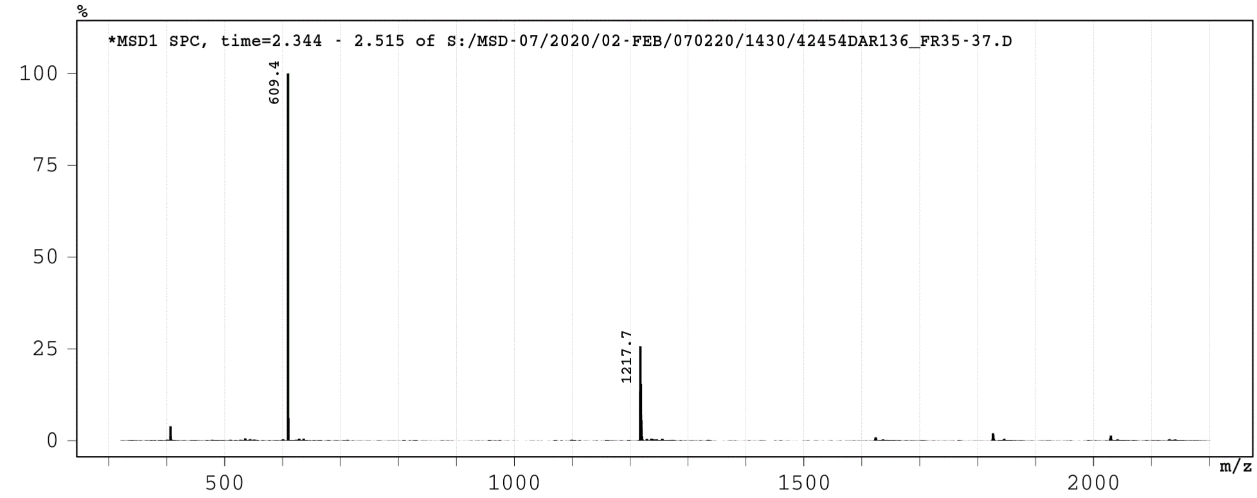


Signal 1: DAD1 A, Sig=220,4 Ref=off

| Peak # | RT [min] | Width [min] | Area | Area % |
|---|---|---|---|---|
| 1 | 2.328 | 2.00 | 48.8 | 1.0 |
| 2 | 2.392 | 0.18 | 4676.0 | 95.1 |
| 3 | 2.512 | 2.69 | 191.8 | 3.9 |

Data File : S:/MSD-07/2020/02-FEB/070220/1430/42454DAR137_FR5->

Injection Date : 07-Feb-20, 18:23:17
Sample Name : 42454DAR137_FR51-52
Remarks : Baseline corrected
Acq. Method : D:/METHODS/5-95-6M.M
column used : Je_559
Analysis Method : C:/CHEM32/1/METHODS/JPT-LCMS.M

Sample Info : 42454_137-Dr. Martine Darwish H-ASYSLVAHI-OH

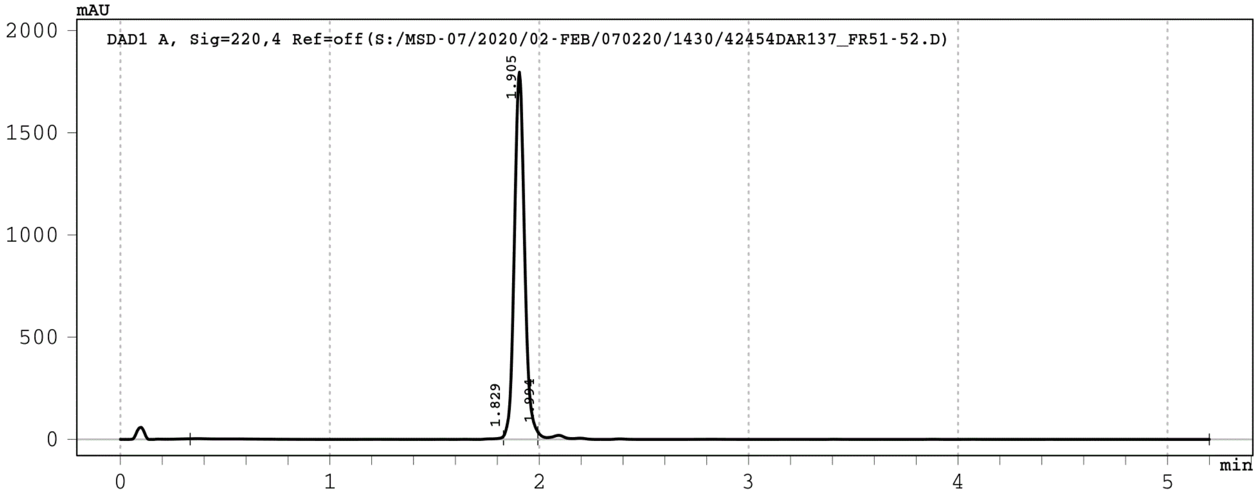


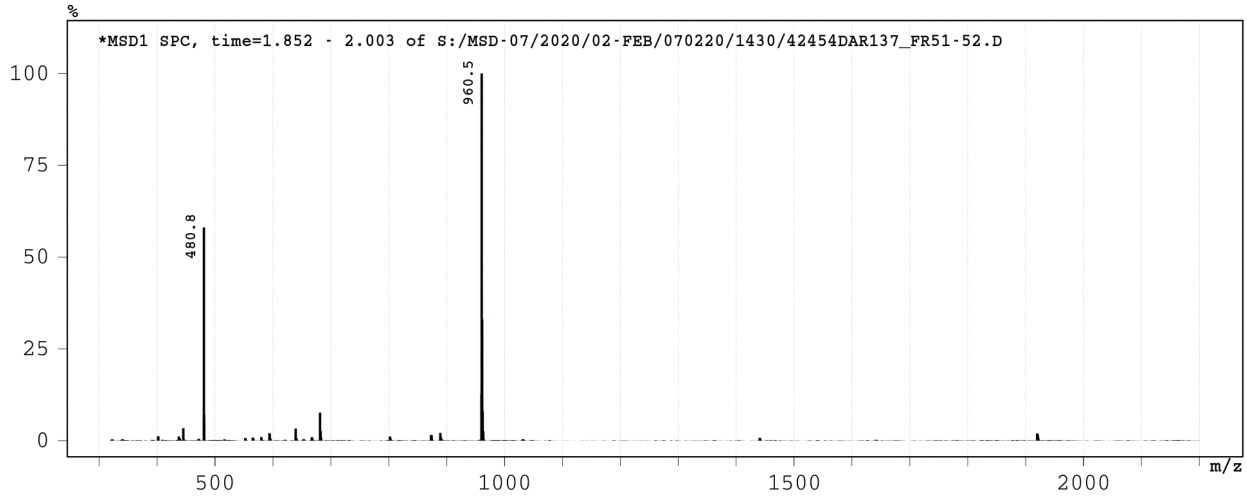


Signal 1: DAD1 A, Sig=220,4 Ref=off

| Peak # | RT [min] | Width [min] | Area | Area % |
|---|---|---|---|---|
| 1 | 1.829 | 1.50 | 86.2 | 1.4 |
| 2 | 1.905 | 0.16 | 6074.7 | 95.9 |
| 3 | 1.994 | 3.21 | 172.5 | 2.7 |

Data File : S:/MSD-07/2020/02-FEB/110220/1433/42454DAR138_FR1->

Injection Date : 12-Feb-20, 00:43:40
Sample Name : 42454DAR138_FR10-11
Remarks : Baseline corrected
Acq. Method : D:/METHODS/5-95-6M.M
column used : Je_559
Analysis Method : C:/CHEM32/1/METHODS/JPT-LCMS.M

Sample Info : 42454_138-Dr. Martine Darwish H-TSIMQLYL-OH

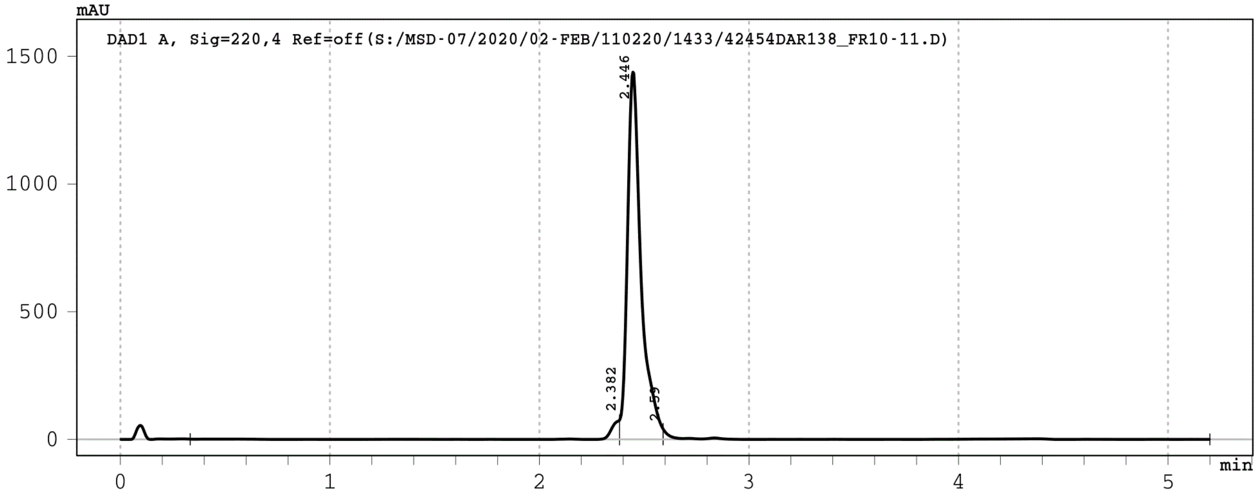


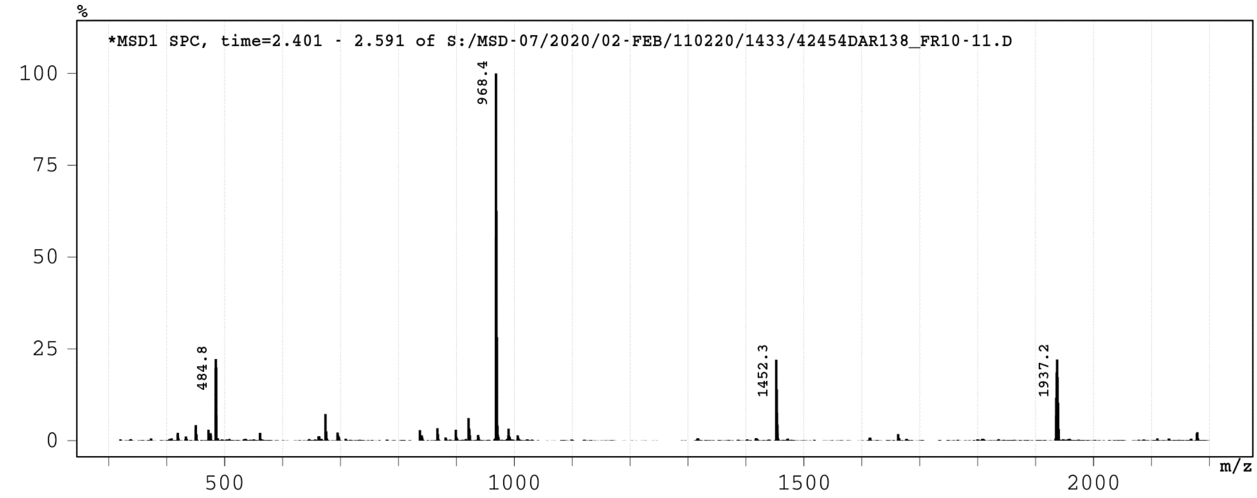


Signal 1: DAD1 A, Sig=220,4 Ref=off

| Peak # | RT [min] | Width [min] | Area | Area % |
|---|---|---|---|---|
| 1 | 2.382 | 2.05 | 221.2 | 3.2 |
| 2 | 2.446 | 0.21 | 6522.4 | 94.3 |
| 3 | 2.590 | 2.61 | 172.3 | 2.5 |

Data File : S:/MSD-07/2020/02-FEB/060220/1429/42454DAR139_FR2->

Injection Date : 06-Feb-20, 18:47:18
Sample Name : 42454DAR139_FR25-26
Remarks : Baseline corrected
Acq. Method : D:/METHODS/5-95-6M.M
column used : Je_559
Analysis Method : C:/CHEM32/1/METHODS/JPT-LCMS.M

Sample Info : 42454_139-Dr. Martine Darwish H-MALSTYYAL-OH

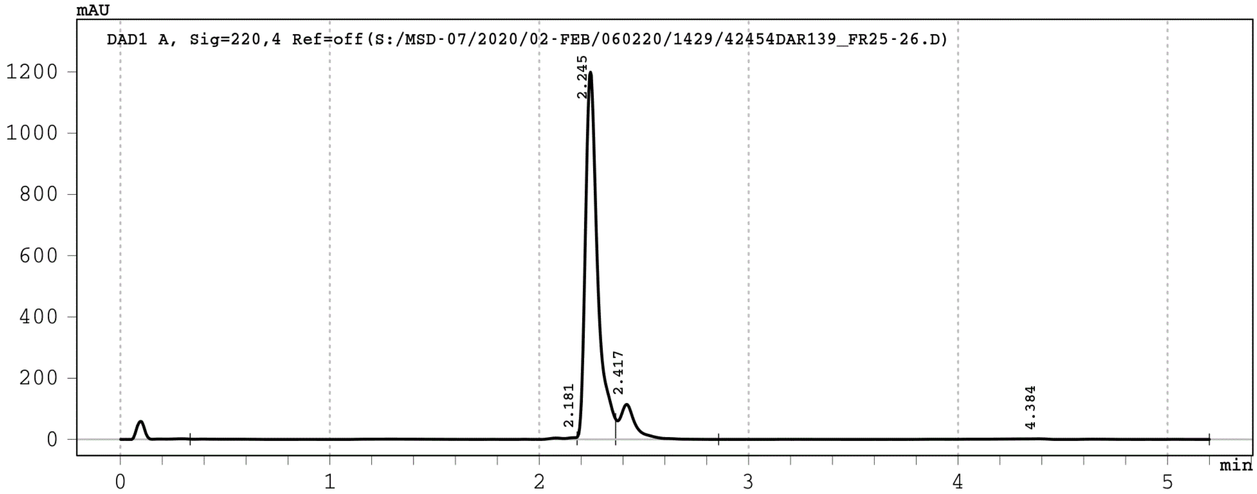


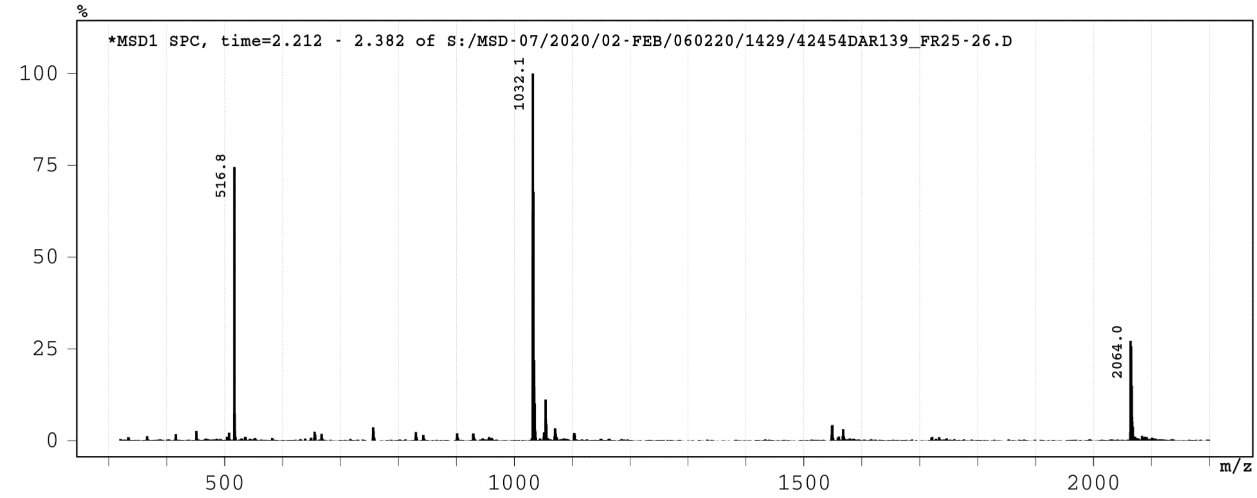


Signal 1: DAD1 A, Sig=220,4 Ref=off

| Peak # | RT [min] | Width [min] | Area | Area % |
|---|---|---|---|---|
| 1 | 2.181 | 1.85 | 69.4 | 1.2 |
| 2 | 2.245 | 0.18 | 4875.6 | 86.9 |
| 3 | 2.417 | 0.49 | 616.1 | 11.0 |
| 4 | 4.384 | 2.34 | 46.8 | 0.8 |

Inj. Date : 14-Feb-20, 18:09:02
Sample Name : 42454DAR140_FR60-61_A
Column used : Je-2502
Remarks : Baseline corrected
Acq. Method : D:/METHODS/5-95-6M.M

Data File : S:/MSD-07/2020/02-FEB/140220/1435/42454DAR140_FR6->

Sample Info : 42454_140-Dr. Martine Darwish
H-KGYRHKVPL-OH

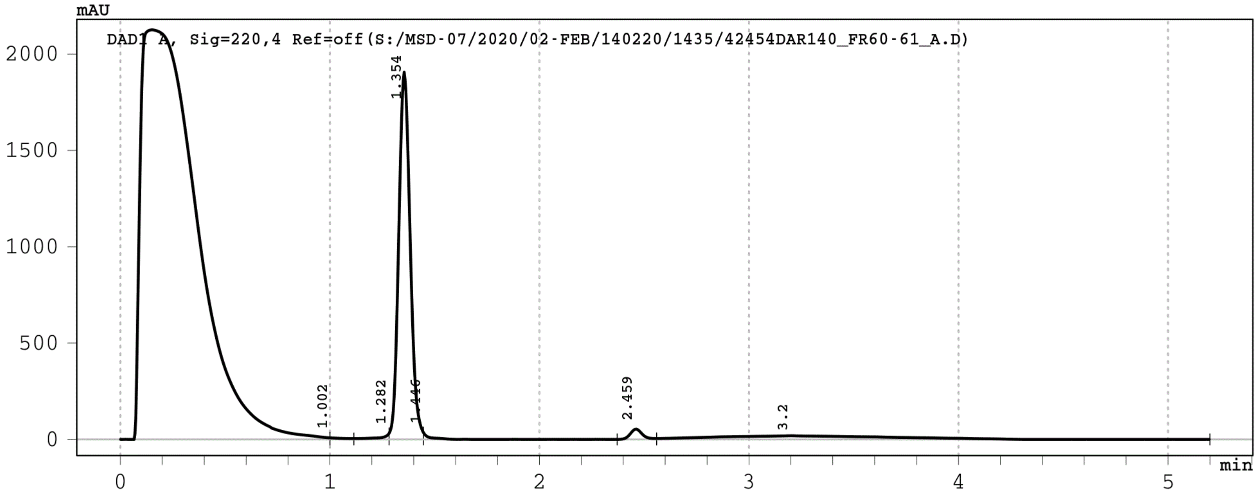


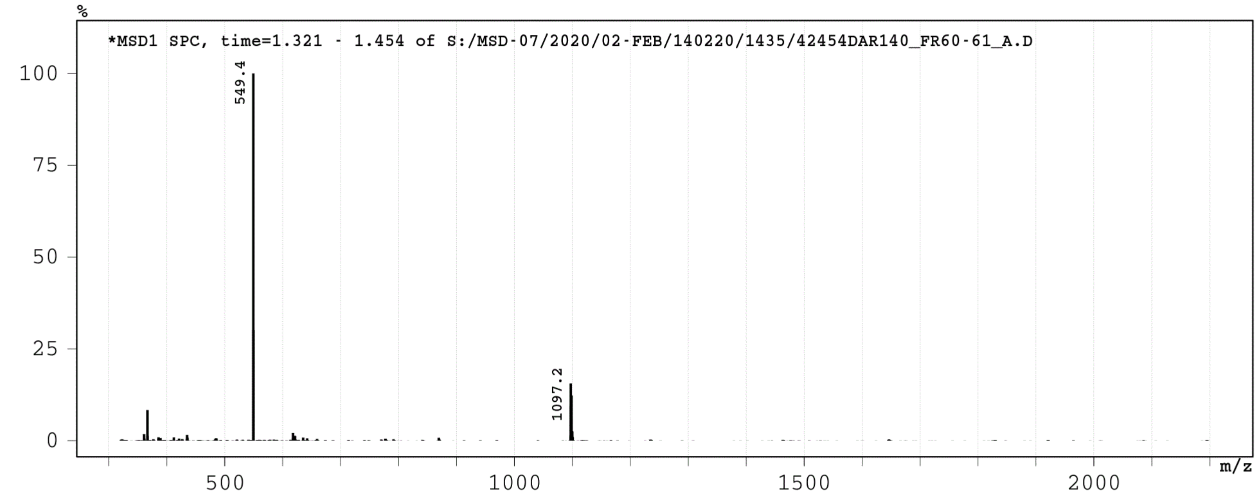


Signal 1: DAD1 A, Sig=220,4 Ref=off

| Peak # | RT [min] | Width [min] | Area | Area % |
|---|---|---|---|---|
| 1 | 1.002 | 0.11 | 38.7 | 0.4 |
| 2 | 1.282 | 0.17 | 84.5 | 1.0 |
| 3 | 1.354 | 0.16 | 7231.5 | 82.4 |
| 4 | 1.446 | 0.92 | 68.9 | 0.8 |
| 5 | 2.459 | 0.19 | 217.7 | 2.5 |
| 6 | 3.200 | 2.64 | 1129.8 | 12.9 |

Data File : S:/MSD-07/2020/02-FEB/060220/1429/42454DAR141_FR3->

Injection Date : 06-Feb-20, 18:55:42
Sample Name : 42454DAR141_FR36-37
Remarks : Baseline corrected
Acq. Method : D:/METHODS/5-95-6M.M
column used : Je_559
Analysis Method : C:/CHEM32/1/METHODS/JPT-LCMS.M

Sample Info : 42454_141-Dr. Martine Darwish H-INPHWNFEKM-OH

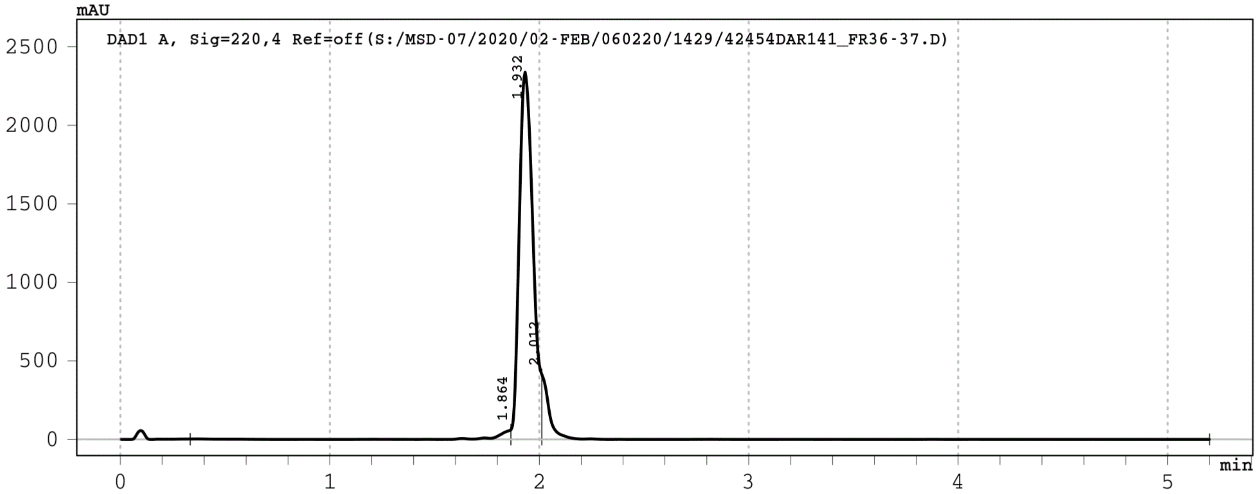


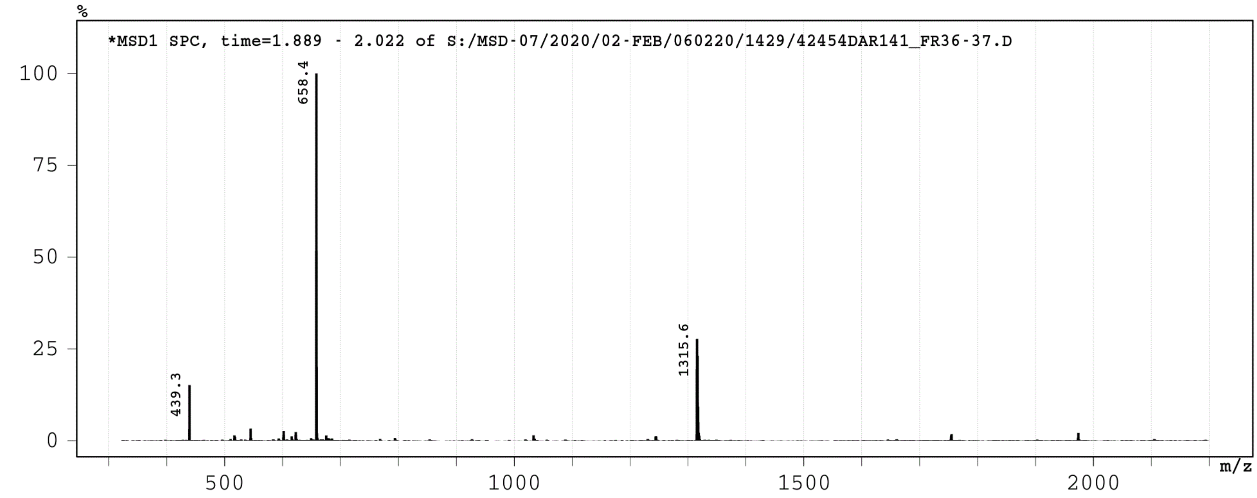


Signal 1: DAD1 A, Sig=220,4 Ref=off

| Peak # | RT [min] | Width [min] | Area | Area % |
|---|---|---|---|---|
| 1 | 1.864 | 1.53 | 277.2 | 2.3 |
| 2 | 1.932 | 0.15 | 10657.2 | 89.3 |
| 3 | 2.012 | 3.19 | 1006.2 | 8.4 |

Data File : S:/MSD-07/2020/02-FEB/060220/1429/42454DAR142_FR4->

Injection Date : 06-Feb-20, 19:04:07
Sample Name : 42454DAR142_FR47-48
Remarks : Baseline corrected
Acq. Method : D:/METHODS/5-95-6M.M
column used : Je_559
Analysis Method : C:/CHEM32/1/METHODS/JPT-LCMS.M

Sample Info : 42454_142-Dr. Martine Darwish H-VIYQLCAF-OH

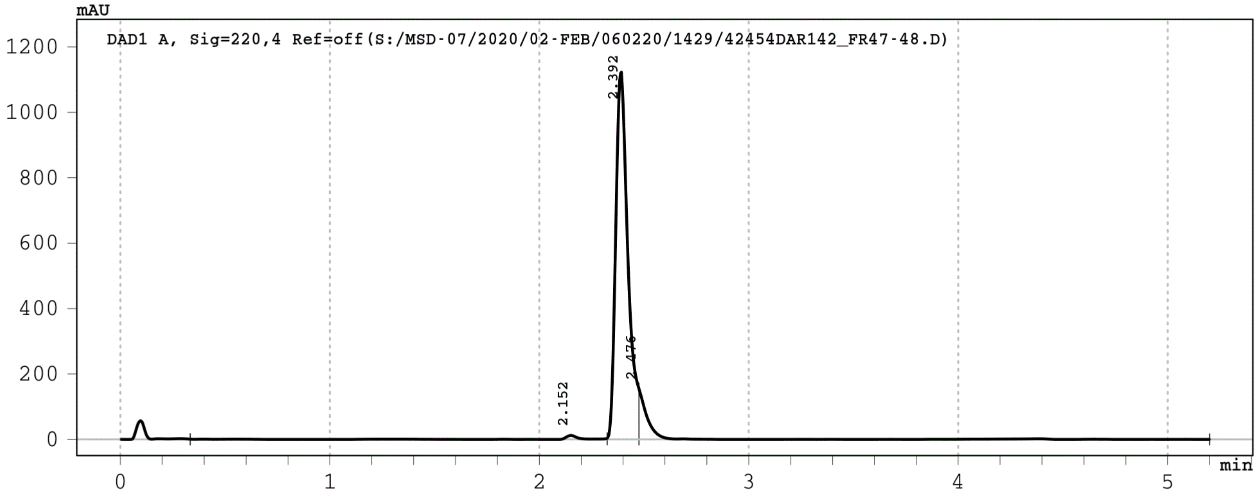


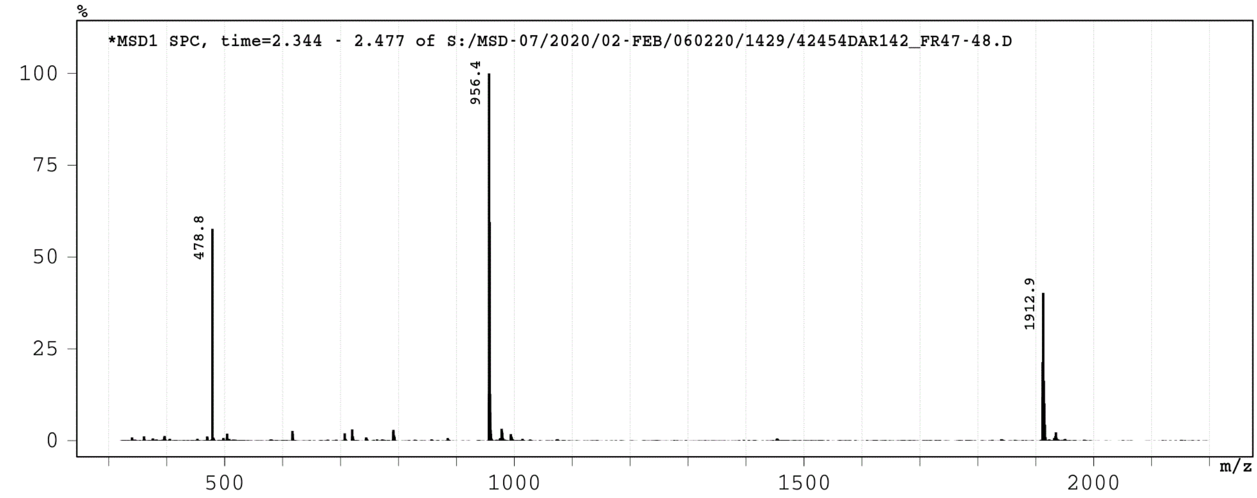


Signal 1: DAD1 A, Sig=220,4 Ref=off

| Peak # | RT [min] | Width [min] | Area | Area % |
|---|---|---|---|---|
| 1 | 2.152 | 1.99 | 77.8 | 1.5 |
| 2 | 2.392 | 0.15 | 4535.7 | 89.6 |
| 3 | 2.476 | 2.72 | 446.9 | 8.8 |

Data File : S:/MSD-07/2020/02-FEB/060220/1429/42454DAR143_FR5->

Injection Date : 06-Feb-20, 19:12:32
Sample Name : 42454DAR143_FR51-52
Remarks : Baseline corrected
Acq. Method : D:/METHODS/5-95-6M.M
column used : Je_559
Analysis Method : C:/CHEM32/1/METHODS/JPT-LCMS.M

Sample Info : 42454_143-Dr. Martine Darwish H-ICAVKFLKL-OH

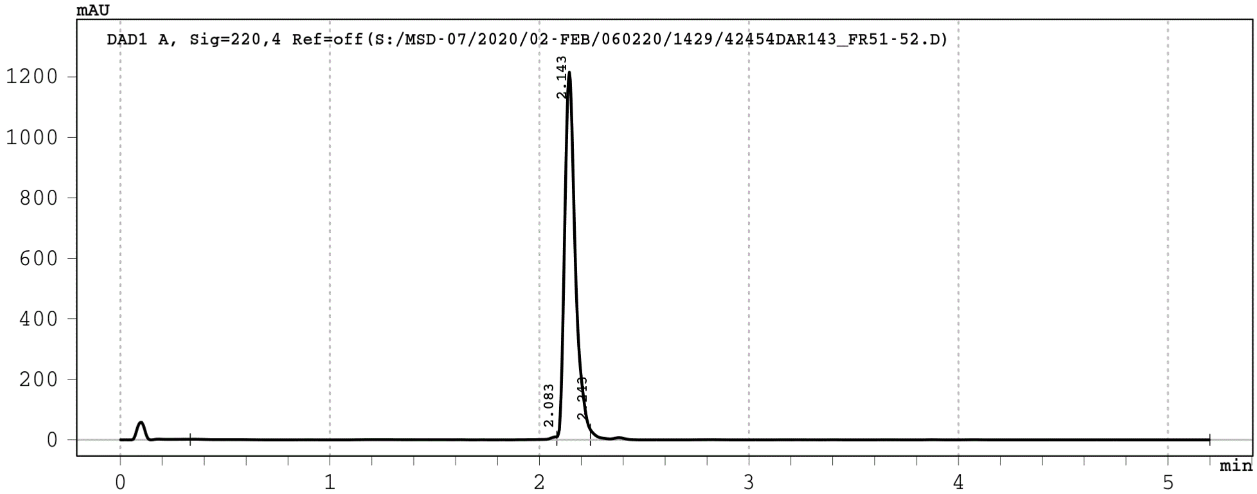


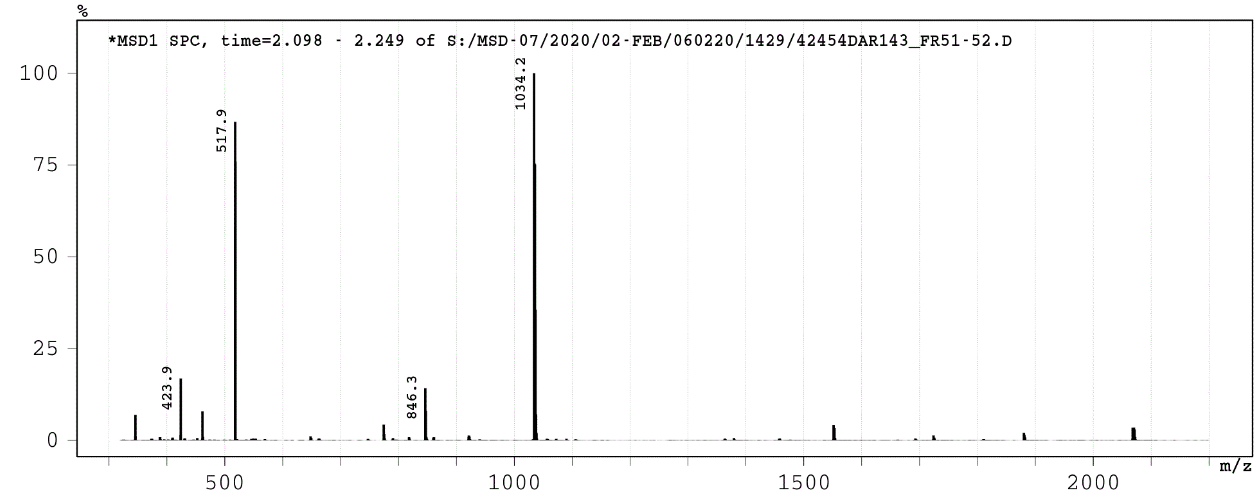


Signal 1: DAD1 A, Sig=220,4 Ref=off

| Peak # | RT [min] | Width [min] | Area | Area % |
|---|---|---|---|---|
| 1 | 2.083 | 1.75 | 61.0 | 1.4 |
| 2 | 2.143 | 0.16 | 4335.9 | 96.4 |
| 3 | 2.243 | 2.96 | 99.2 | 2.2 |

Data File : S:/MSD-07/2020/02-FEB/140220/1435/42454DAR144_FR5->

Injection Date : 15-Feb-20, 00:26:49
Sample Name : 42454DAR144_FR53
Remarks : Baseline corrected
Acq. Method : D:/METHODS/5-95-6M.M
column used : Je_559
Analysis Method : C:/CHEM32/1/METHODS/JPT-LCMS.M

Sample Info : 42454_144-Dr. Martine Darwish H-TWHEWRYYFF-OH

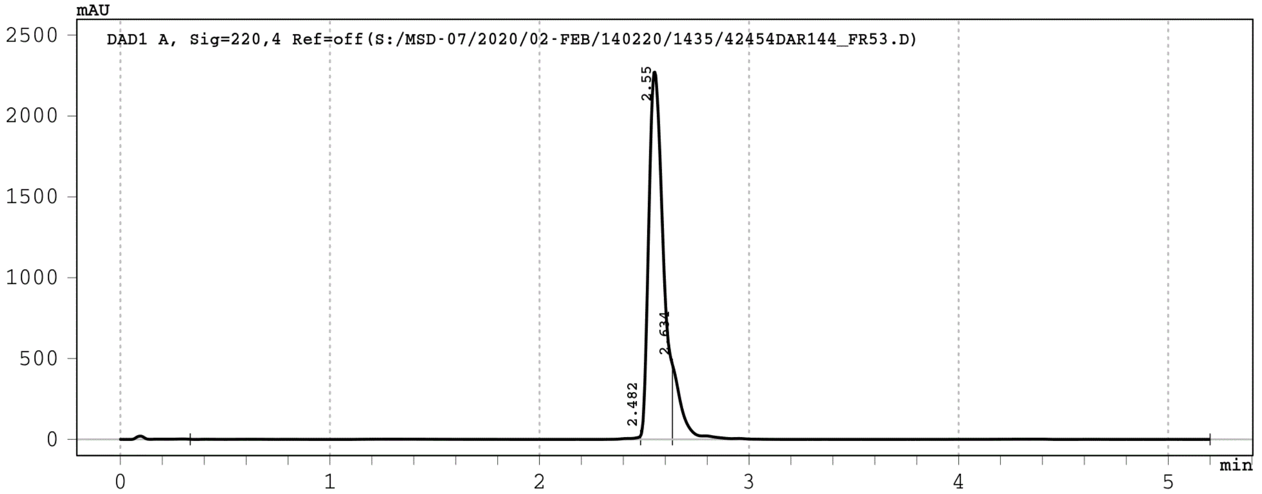


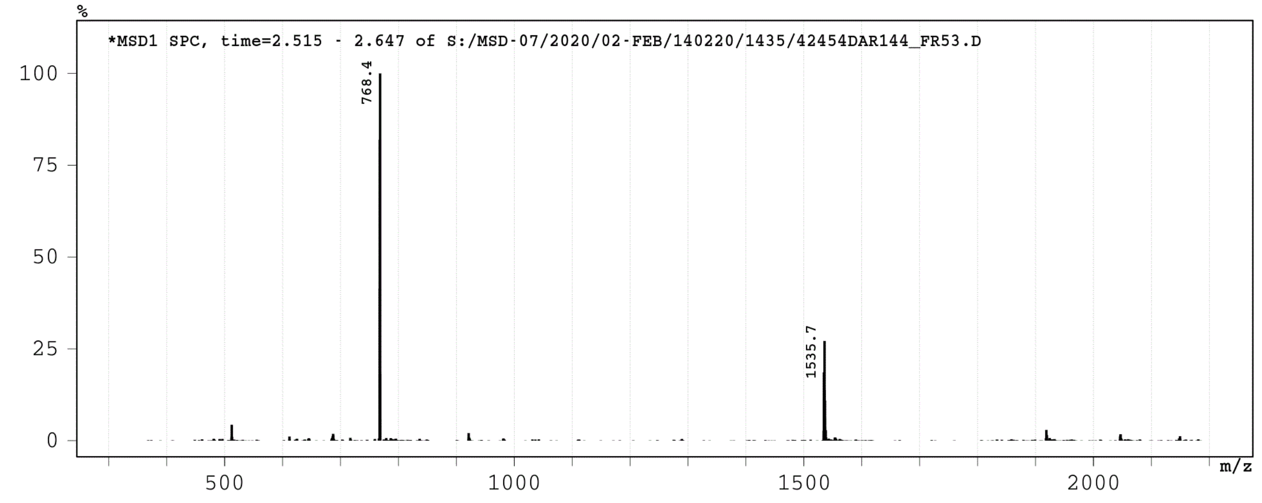


Signal 1: DAD1 A, Sig=220,4 Ref=off

| Peak # | RT [min] | Width [min] | Area | Area % |
|---|---|---|---|---|
| 1 | 2.482 | 2.15 | 95.8 | 0.8 |
| 2 | 2.550 | 0.15 | 10709.4 | 87.3 |
| 3 | 2.634 | 2.57 | 1463.0 | 11.9 |

Data File : S:/MSD-07/2020/02-FEB/140220/1435/42454DAR145_FR8->

Injection Date : 15-Feb-20, 00:35:11
Sample Name : 42454DAR145_FR81-83
Remarks : Baseline corrected
Acq. Method : D:/METHODS/5-95-6M.M
column used : Je_559
Analysis Method : C:/CHEM32/1/METHODS/JPT-LCMS.M

Sample Info : 42454_145-Dr. Martine Darwish H-IILETIWL-OH

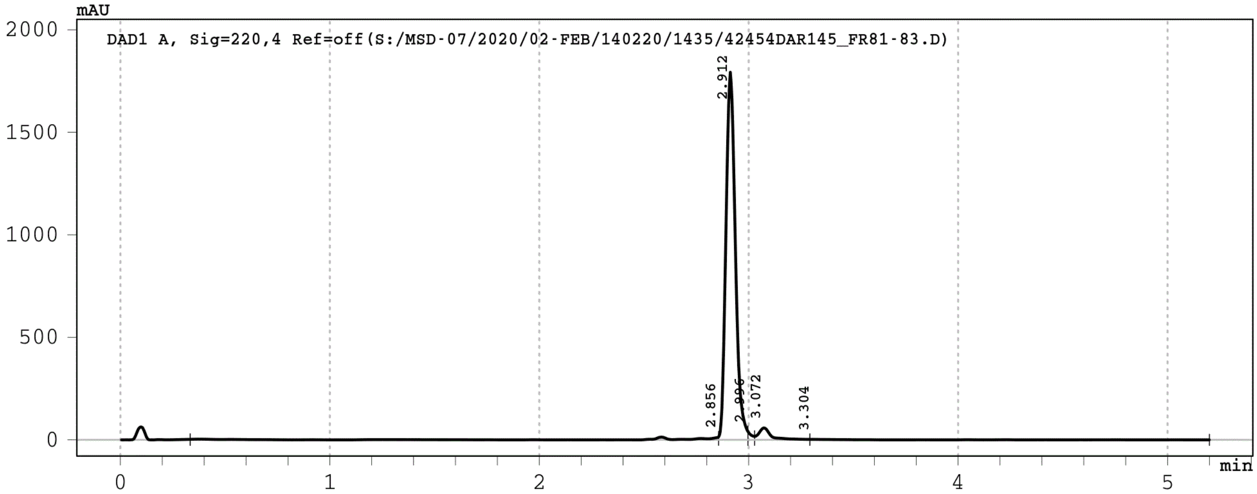


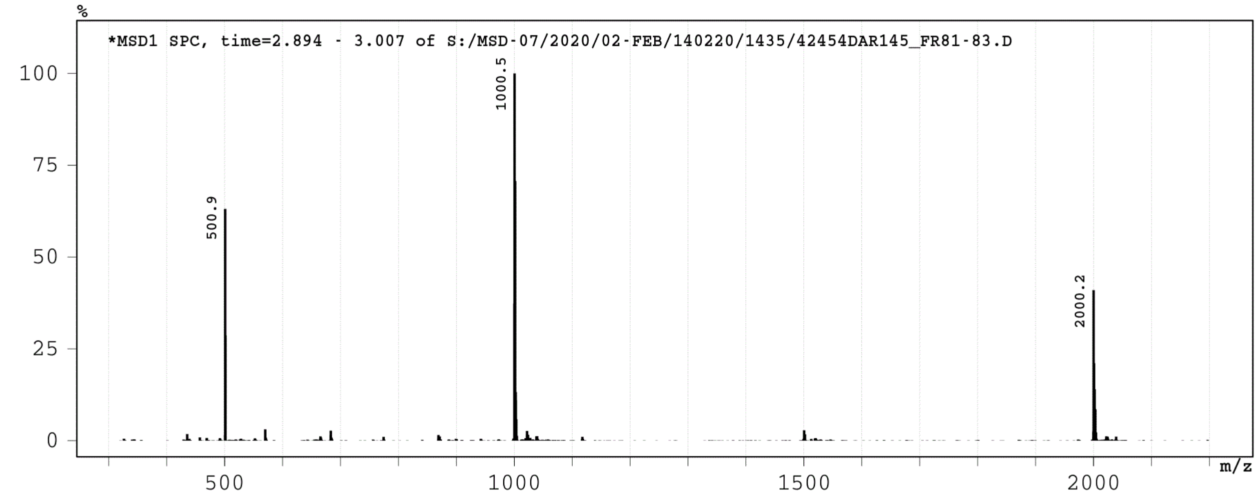


Signal 1: DAD1 A, Sig=220,4 Ref=off

| Peak # | RT [min] | Width [min] | Area | Area % |
|---|---|---|---|---|
| 1 | 2.856 | 2.52 | 191.3 | 3.1 |
| 2 | 2.912 | 0.14 | 5619.2 | 91.6 |
| 3 | 2.996 | 0.03 | 47.8 | 0.8 |
| 4 | 3.072 | 0.26 | 246.1 | 4.0 |
| 5 | 3.304 | 1.91 | 31.4 | 0.5 |

Data File : S:/MSD-07/2020/02-FEB/140220/1435/42454DAR146_FR2->

Injection Date : 15-Feb-20, 00:43:33
Sample Name : 42454DAR146_FR2-4
Remarks : Baseline corrected
Acq. Method : D:/METHODS/5-95-6M.M
column used : Je_559
Analysis Method : C:/CHEM32/1/METHODS/JPT-LCMS.M

Sample Info : 42454_146-Dr. Martine Darwish H-SASTNPNAM-OH

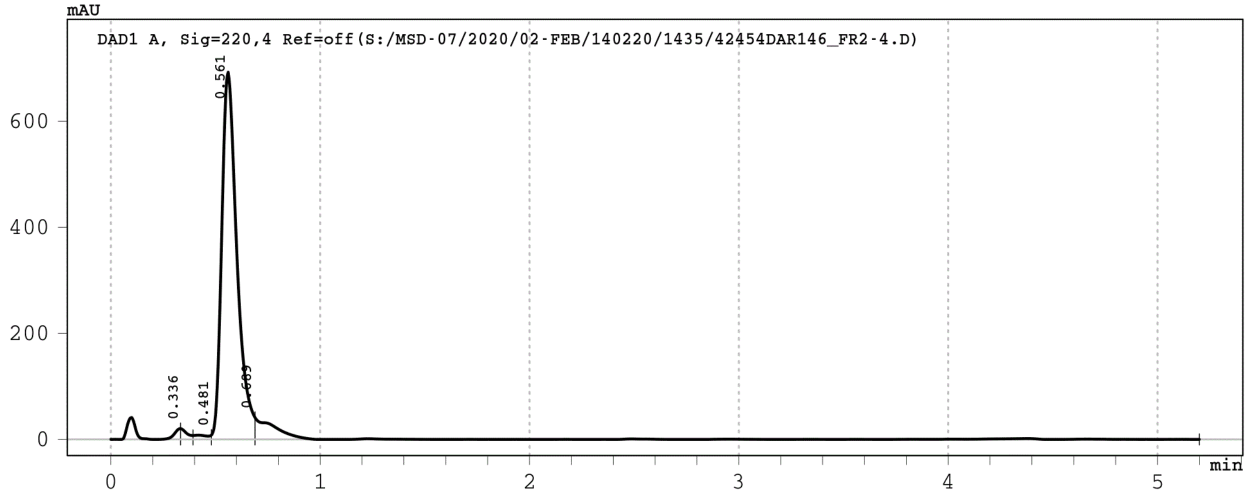


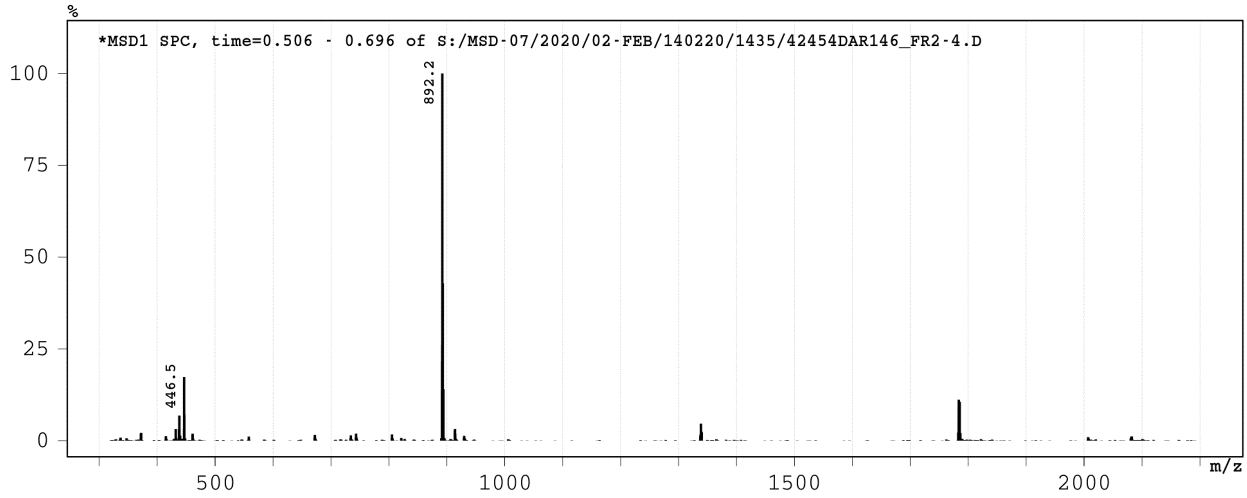


Signal 1: DAD1 A, Sig=220,4 Ref=off

| Peak # | RT [min] | Width [min] | Area | Area % |
|---|---|---|---|---|
| 1 | 0.336 | 0.06 | 40.5 | 1.0 |
| 2 | 0.481 | 0.09 | 38.2 | 1.0 |
| 3 | 0.561 | 0.21 | 3497.7 | 89.6 |
| 4 | 0.689 | 4.51 | 327.5 | 8.4 |

Data File : S:/MSD-07/2020/02-FEB/070220/1430/42454DAR147_FR5->

Injection Date : 07-Feb-20, 18:48:25
Sample Name : 42454DAR147_FR5-6
Remarks : Baseline corrected
Acq. Method : D:/METHODS/5-95-6M.M
column used : Je_559
Analysis Method : C:/CHEM32/1/METHODS/JPT-LCMS.M

Sample Info : 42454_147-Dr. Martine Darwish H-KGYYFNKGTL-OH

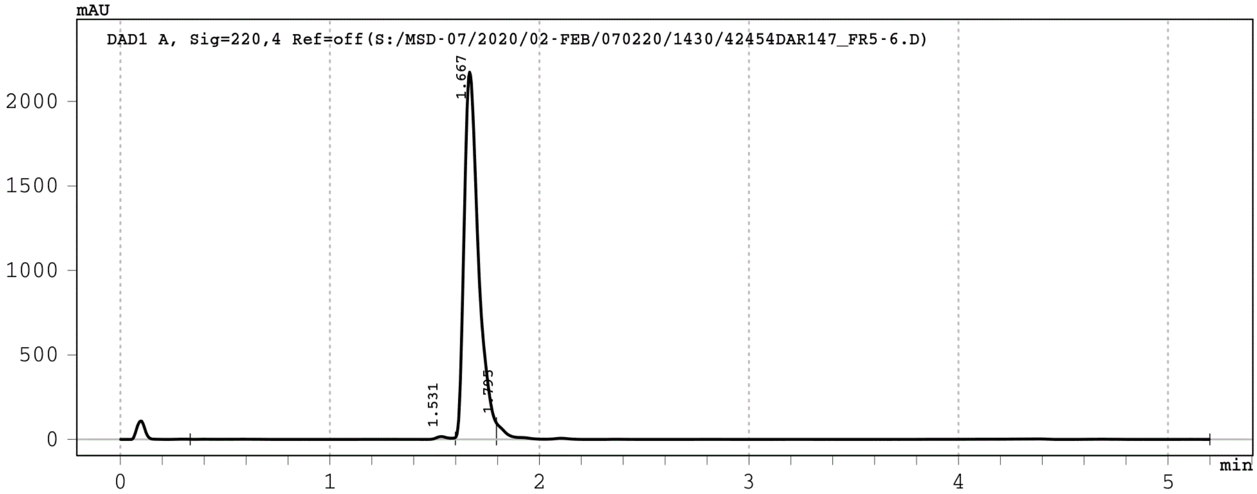


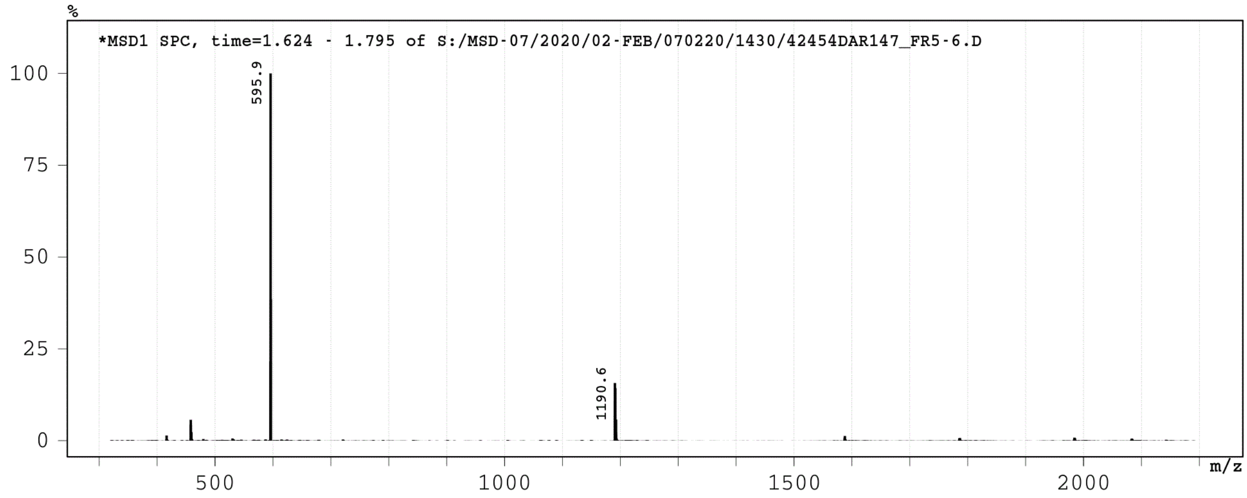


Signal 1: DAD1 A, Sig=220,4 Ref=off

| Peak # | RT [min] | Width [min] | Area | Area % |
|---|---|---|---|---|
| 1 | 1.531 | 1.27 | 89.1 | 0.8 |
| 2 | 1.667 | 0.20 | 10377.3 | 95.5 |
| 3 | 1.795 | 3.41 | 397.0 | 3.7 |

Data File : S:/MSD-07/2020/02-FEB/070220/1430/42454DAR148_FR1->

Injection Date : 07-Feb-20, 18:56:49
Sample Name : 42454DAR148_FR17
Remarks : Baseline corrected
Acq. Method : D:/METHODS/5-95-6M.M
column used : Je_559
Analysis Method : C:/CHEM32/1/METHODS/JPT-LCMS.M

Sample Info : 42454_148-Dr. Martine Darwish H-VRFSLPSHL-OH

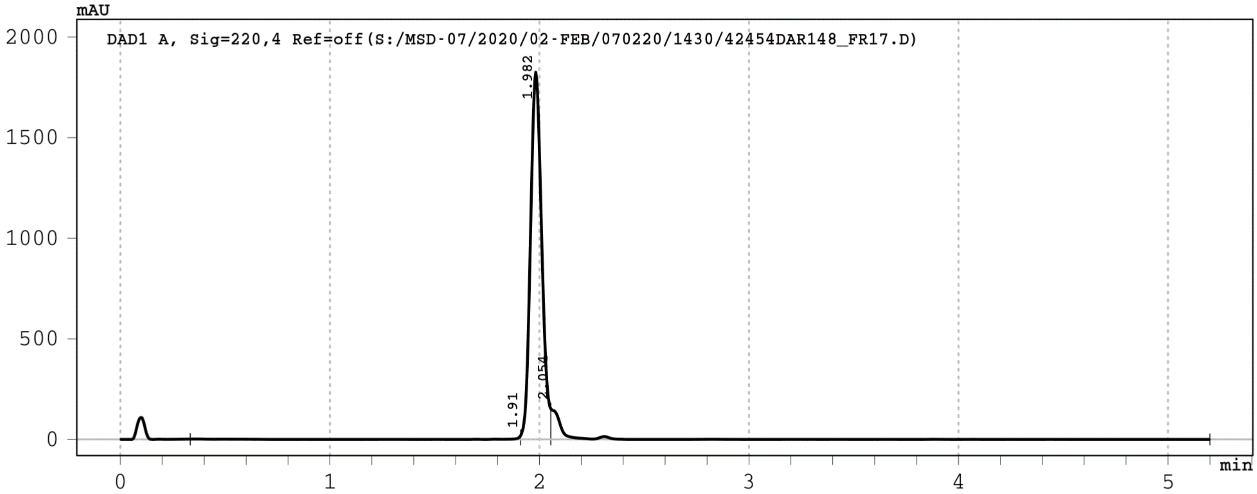


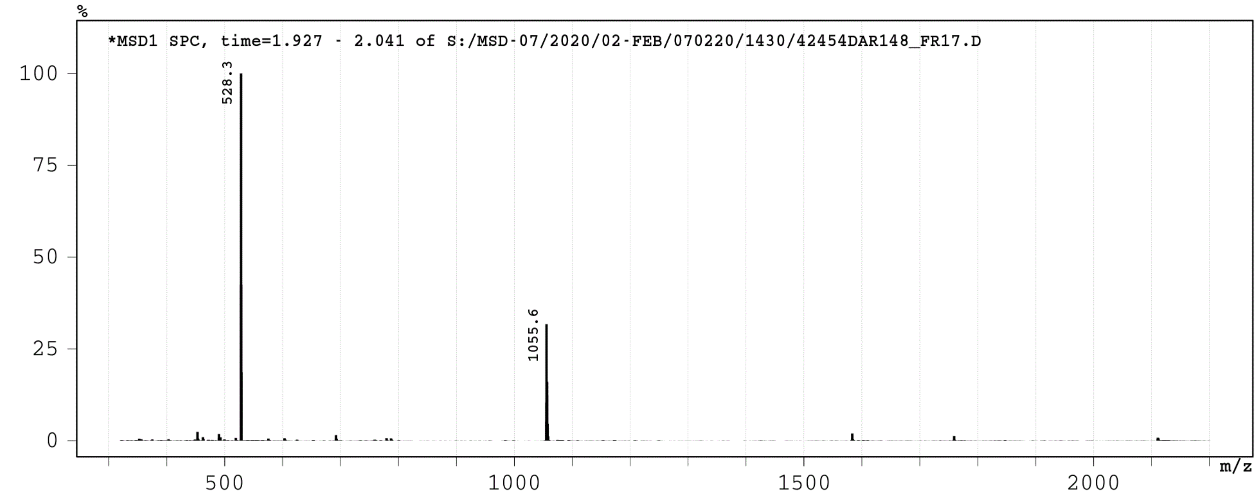


Signal 1: DAD1 A, Sig=220,4 Ref=off

| Peak # | RT [min] | Width [min] | Area | Area % |
|---|---|---|---|---|
| 1 | 1.910 | 1.58 | 47.1 | 0.6 |
| 2 | 1.982 | 0.14 | 6783.1 | 92.2 |
| 3 | 2.054 | 3.15 | 524.0 | 7.1 |

Data File : S:/MSD-07/2020/02-FEB/070220/1430/42454DAR149_FR3->

Injection Date : 07-Feb-20, 19:05:14
Sample Name : 42454DAR149_FR31-32
Remarks : Baseline corrected
Acq. Method : D:/METHODS/5-95-6M.M
column used : Je_559
Analysis Method : C:/CHEM32/1/METHODS/JPT-LCMS.M

Sample Info : 42454_149-Dr. Martine Darwish H-CWYLALVSAL-OH

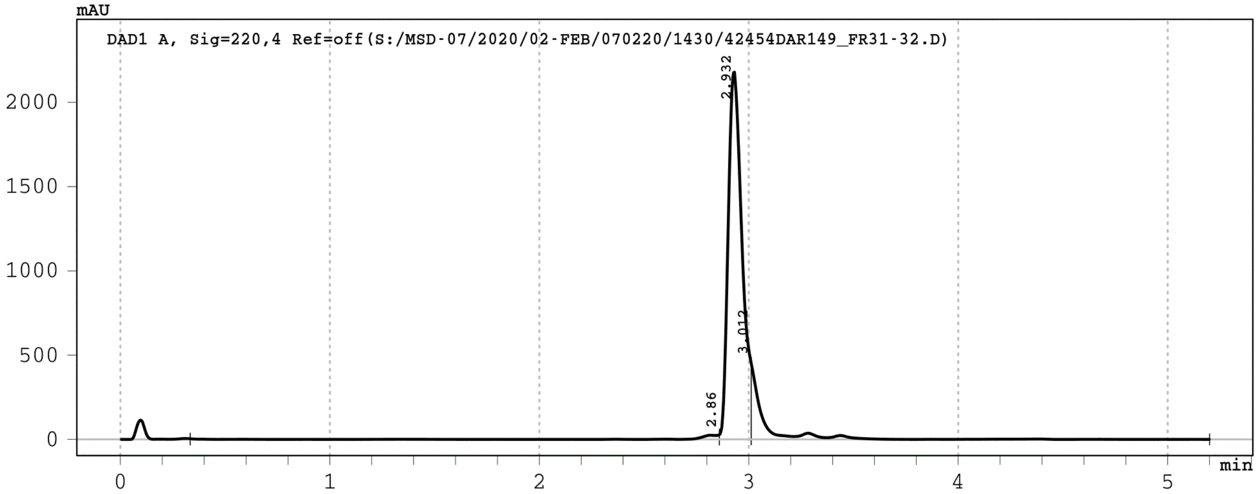


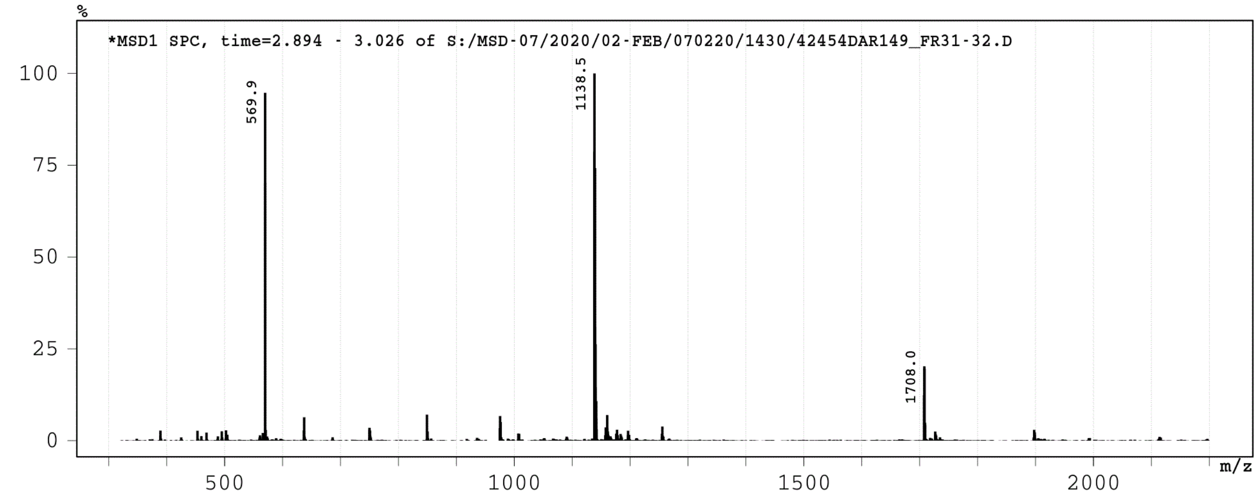


Signal 1: DAD1 A, Sig=220,4 Ref=off

| Peak # | RT [min] | Width [min] | Area | Area % |
|---|---|---|---|---|
| 1 | 2.860 | 2.53 | 172.2 | 1.5 |
| 2 | 2.932 | 0.15 | 10018.0 | 84.6 |
| 3 | 3.012 | 2.19 | 1653.6 | 14.0 |

Data File: S:\MSD-07\2020\02-FEB\070220\143->

Injection Date : Fri, 7. Feb. 2020
Sample Name : 240120SG2-72|42454Dar150
Acq Operator : SYSTEM
Acq. Method : D:\Data\2020\02-FEB\070220\1430\5-95-6M.M
column used :
Analysis Method : C:\Chem32\1\METHODS\DEF_LC.M

Sample Info : 42454_150-Dr. Martine Darwish|H-SSPSFAKI-OH

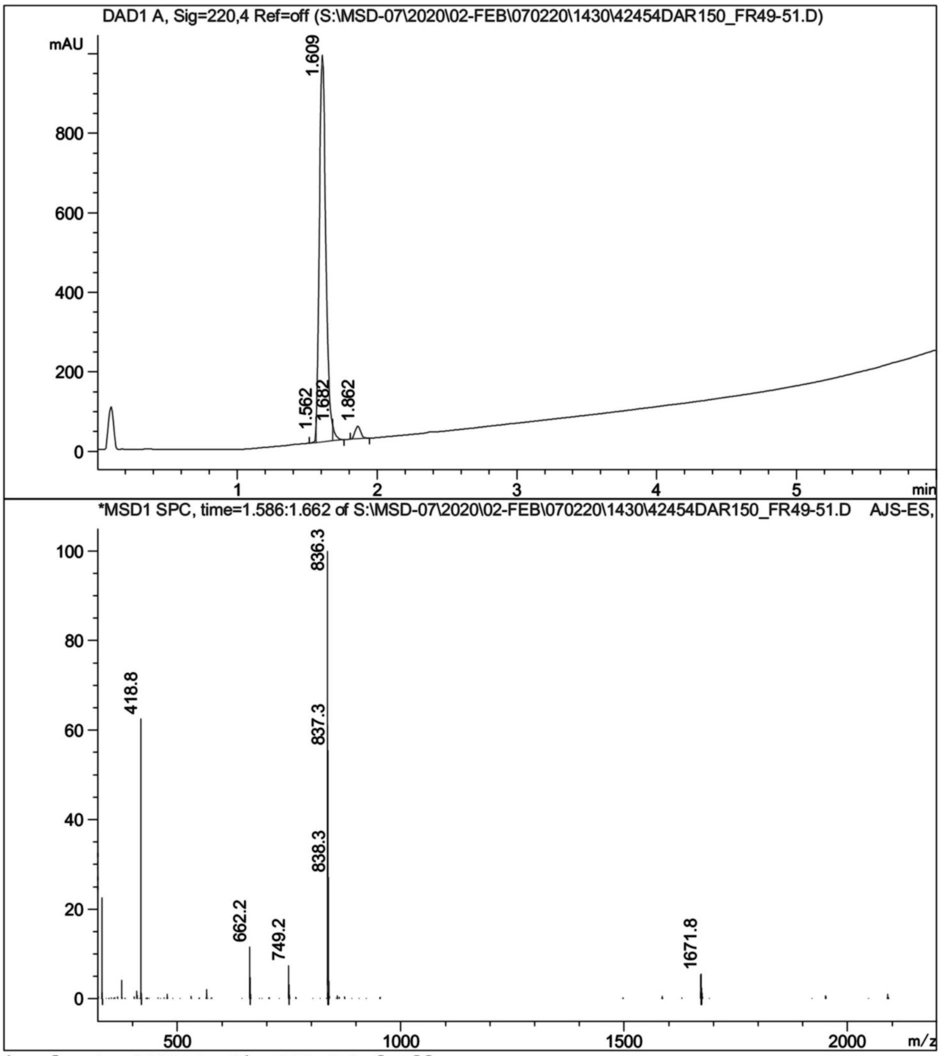


Signal 1 : DAD1 A, Sig=220,4 Ref=off

| Peak # | RT [min] | Width [min] | Area | Area % |
|---|---|---|---|---|
| 1 | 1.562 | 0.0 | 30.0 | 0.9 |
| 2 | 1.609 | 0.1 | 3102.6 | 95.1 |
| 3 | 1.682 | 0.0 | 41.3 | 1.3 |
| 4 | 1.862 | 0.0 | 89.4 | 2.7 |

*** End of Report ***

Data File : S:/MSD-07/2020/02-FEB/060220/1429/42454DAR151_FR5->

Injection Date : 06-Feb-20, 19:20:54
Sample Name : 42454DAR151_FR58-59
Remarks : Baseline corrected
Acq. Method : D:/METHODS/5-95-6M.M
column used : Je_559
Analysis Method : C:/CHEM32/1/METHODS/JPT-LCMS.M

Sample Info : 42454_151-Dr. Martine Darwish H-SSPGFCNTFL-OH

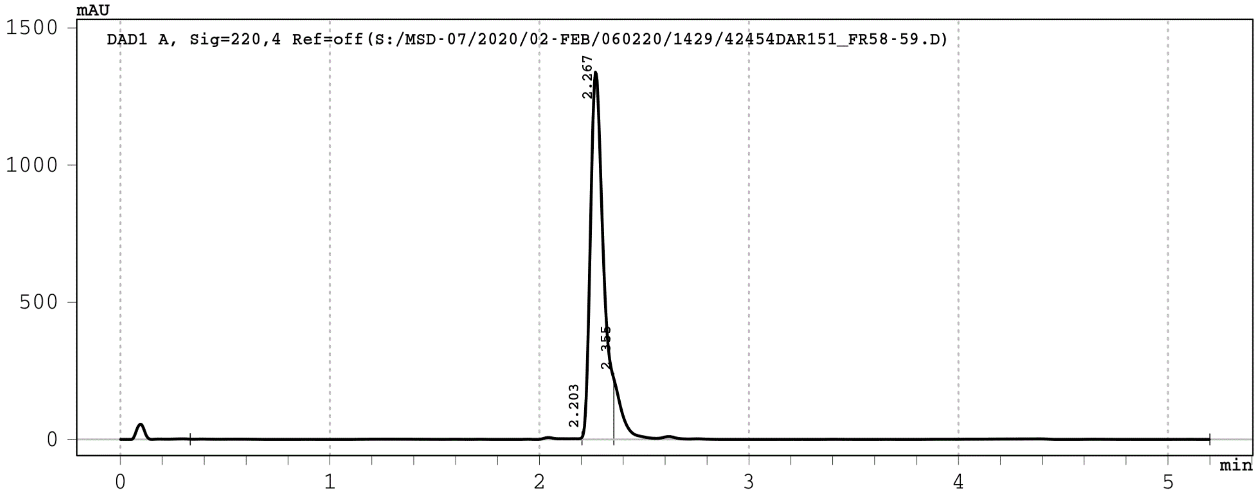


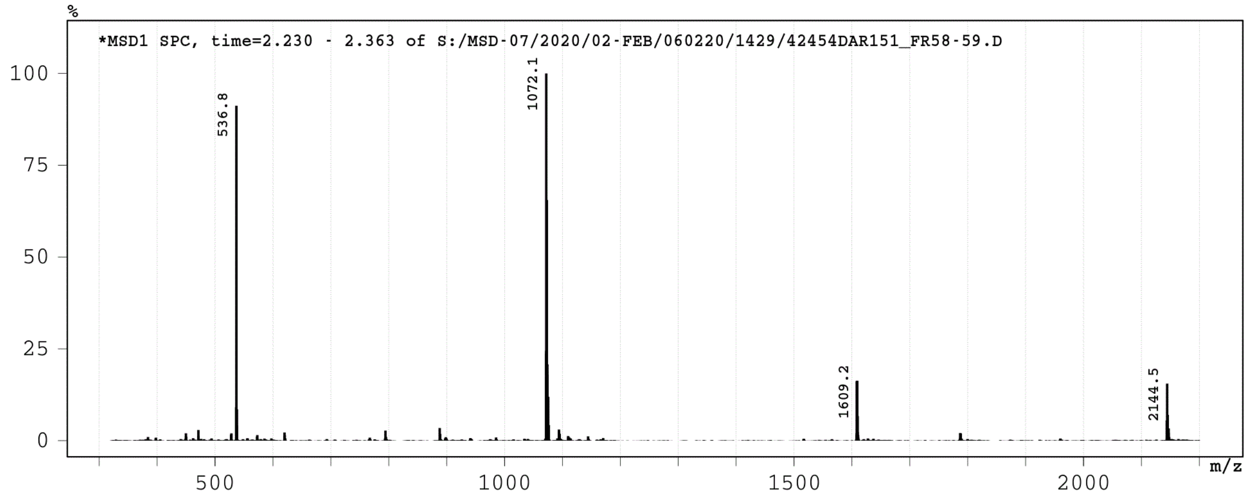


Signal 1: DAD1 A, Sig=220,4 Ref=off

| Peak # | RT [min] | Width [min] | Area | Area % |
|---|---|---|---|---|
| 1 | 2.203 | 1.87 | 73.4 | 1.1 |
| 2 | 2.267 | 0.15 | 5713.6 | 88.1 |
| 3 | 2.355 | 2.84 | 701.2 | 10.8 |

Data File : S:/MSD-07/2020/02-FEB/110220/1433/42454DAR152_FR1->

Injection Date : 12-Feb-20, 00:52:04
Sample Name : 42454DAR152_FR19
Remarks : Baseline corrected
Acq. Method : D:/METHODS/5-95-6M.M
column used : Je_559
Analysis Method : C:/CHEM32/1/METHODS/JPT-LCMS.M

Sample Info : 42454_152-Dr. Martine Darwish H-FSLQFALL-OH

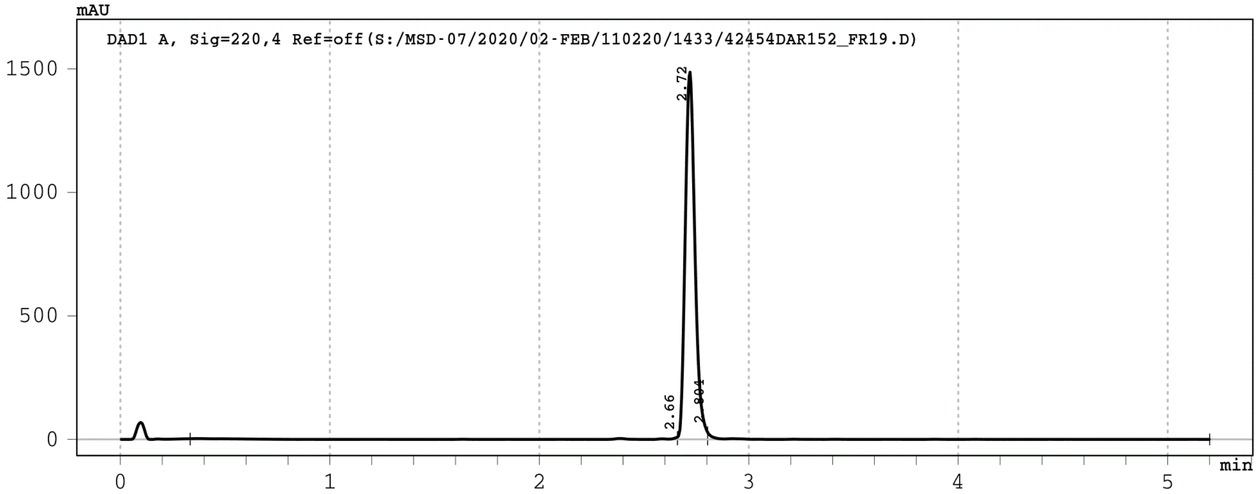


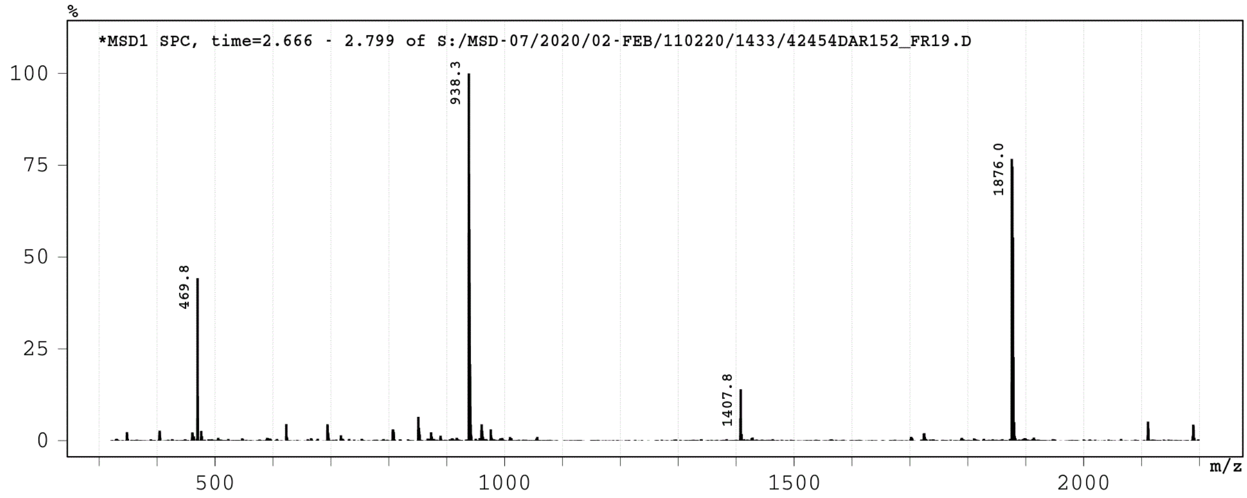


Signal 1: DAD1 A, Sig=220,4 Ref=off

| Peak # | RT [min] | Width [min] | Area | Area % |
|---|---|---|---|---|
| 1 | 2.660 | 2.33 | 85.2 | 1.7 |
| 2 | 2.720 | 0.14 | 4775.8 | 96.9 |
| 3 | 2.804 | 2.40 | 66.2 | 1.3 |

Data File : S:/MSD-07/2020/02-FEB/110220/1433/42454DAR153_FR2->

Injection Date : 12-Feb-20, 01:00:30
Sample Name : 42454DAR153_FR26-27
Remarks : Baseline corrected
Acq. Method : D:/METHODS/5-95-6M.M
column used : Je_559
Analysis Method : C:/CHEM32/1/METHODS/JPT-LCMS.M

Sample Info : 42454_153-Dr. Martine Darwish H-SSMVPSAL-OH

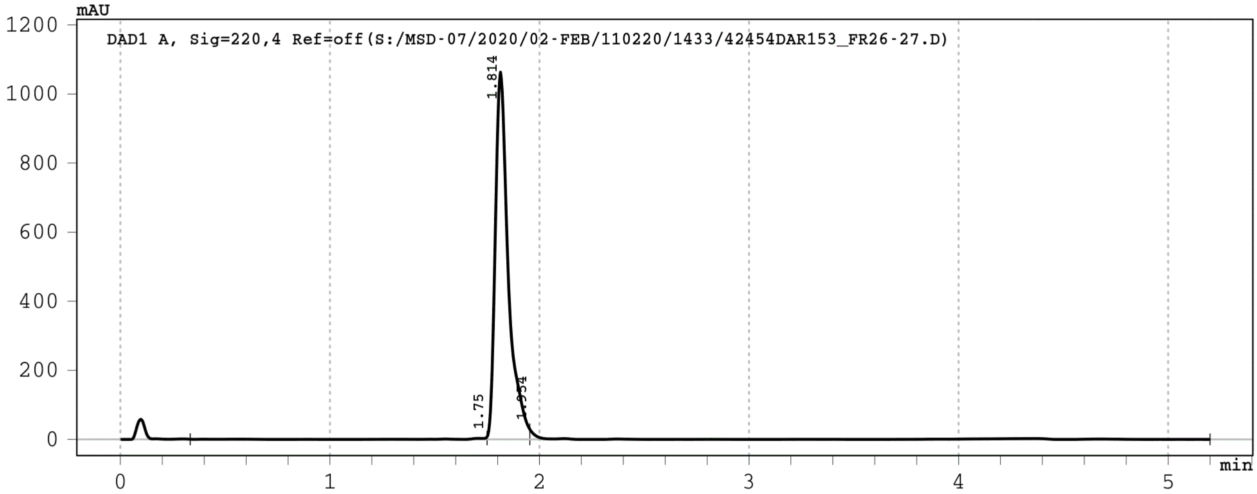


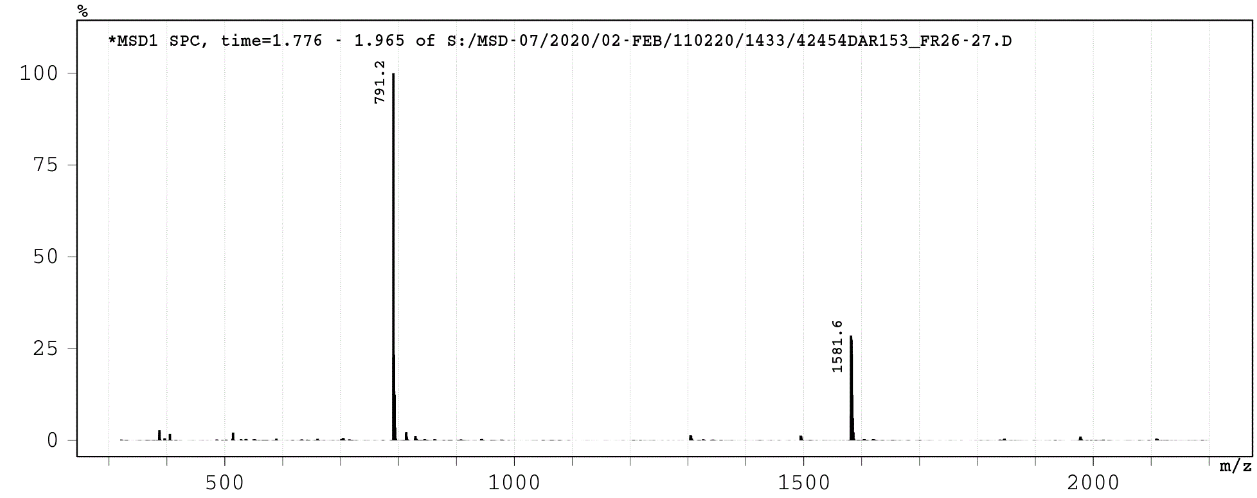


Signal 1: DAD1 A, Sig=220,4 Ref=off

| Peak # | RT [min] | Width [min] | Area | Area % |
|---|---|---|---|---|
| 1 | 1.750 | 1.42 | 34.4 | 0.7 |
| 2 | 1.814 | 0.20 | 4597.2 | 96.3 |
| 3 | 1.954 | 3.25 | 140.1 | 2.9 |

Data File : S:/MSD-07/2020/02-FEB/070220/1430/42454DAR154_FR5->

Injection Date : 07-Feb-20, 19:22:01
Sample Name : 42454DAR154_FR54
Remarks : Baseline corrected
Acq. Method : D:/METHODS/5-95-6M.M
column used : Je_559
Analysis Method : C:/CHEM32/1/METHODS/JPT-LCMS.M

Sample Info : 42454_154-Dr. Martine Darwish H-YSGANATAT-OH

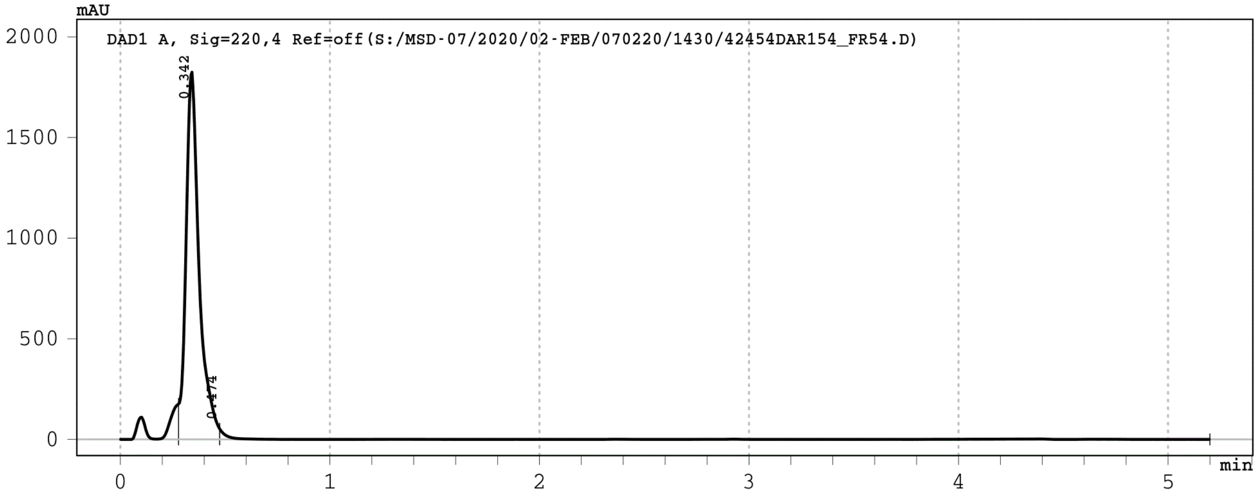


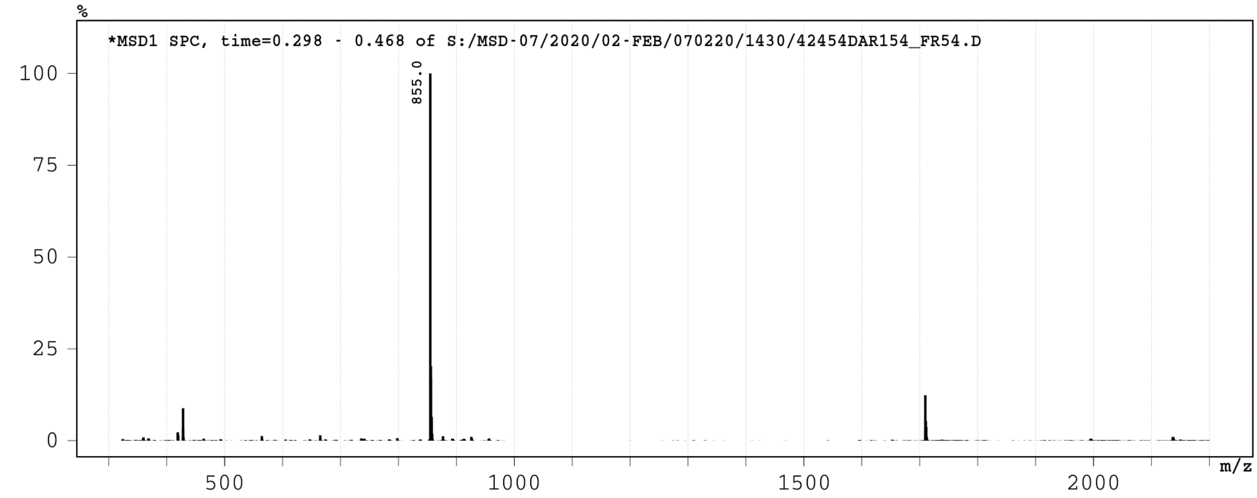


Signal 1: DAD1 A, Sig=220,4 Ref=off

| Peak # | RT [min] | Width [min] | Area | Area % |
|---|---|---|---|---|
| 1 | 0.342 | 0.20 | 7968.3 | 97.7 |
| 2 | 0.474 | 4.73 | 184.4 | 2.3 |

Data File : S:/MSD-07/2020/02-FEB/060220/1429/42454DAR155_FR6->

Injection Date : 06-Feb-20, 19:29:17
Sample Name : 42454DAR155_FR69-70
Remarks : Baseline corrected
Acq. Method : D:/METHODS/5-95-6M.M
column used : Je_559
Analysis Method : C:/CHEM32/1/METHODS/JPT-LCMS.M

Sample Info : 42454_155-Dr. Martine Darwish H-ASLPMGYYI-OH

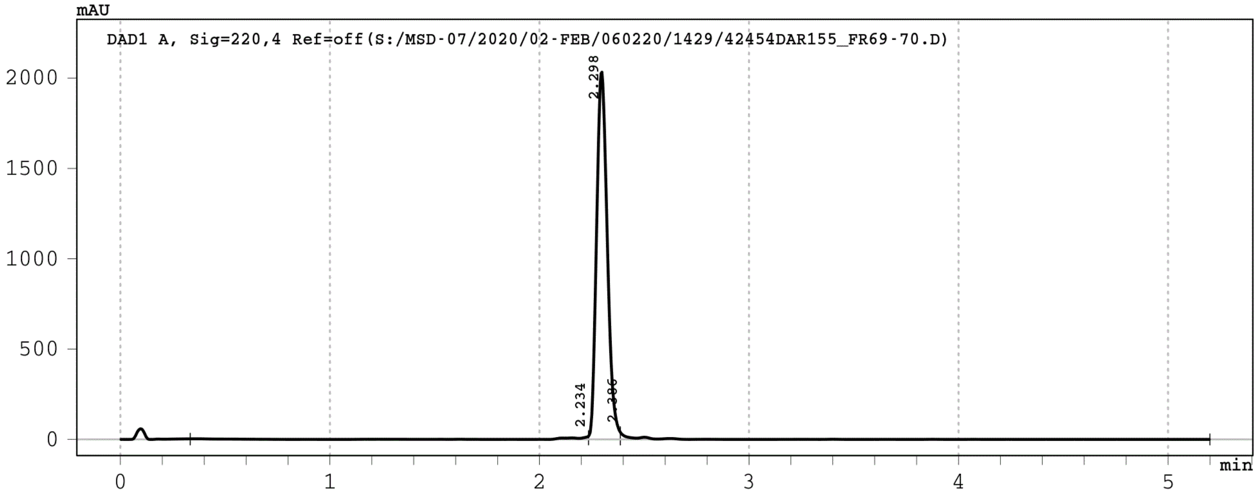


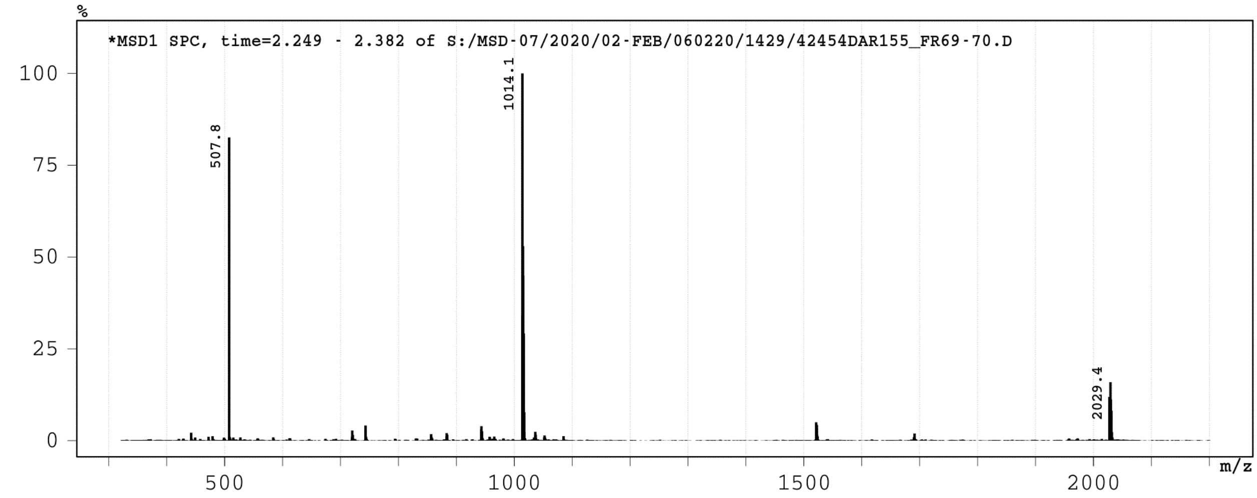


Signal 1: DAD1 A, Sig=220,4 Ref=off

| Peak # | RT [min] | Width [min] | Area | Area % |
|---|---|---|---|---|
| 1 | 2.234 | 1.90 | 137.7 | 1.8 |
| 2 | 2.298 | 0.15 | 7243.8 | 96.2 |
| 3 | 2.386 | 2.81 | 148.4 | 2.0 |

Data File : S:/MSD-07/2020/02-FEB/070220/1430/42454DAR156_FR5->

Injection Date : 07-Feb-20, 19:30:24
Sample Name : 42454DAR156_FR59
Remarks : Baseline corrected
Acq. Method : D:/METHODS/5-95-6M.M
column used : Je_559
Analysis Method : C:/CHEM32/1/METHODS/JPT-LCMS.M

Sample Info : 42454_156-Dr. Martine Darwish H-RNRRIFALL-OH

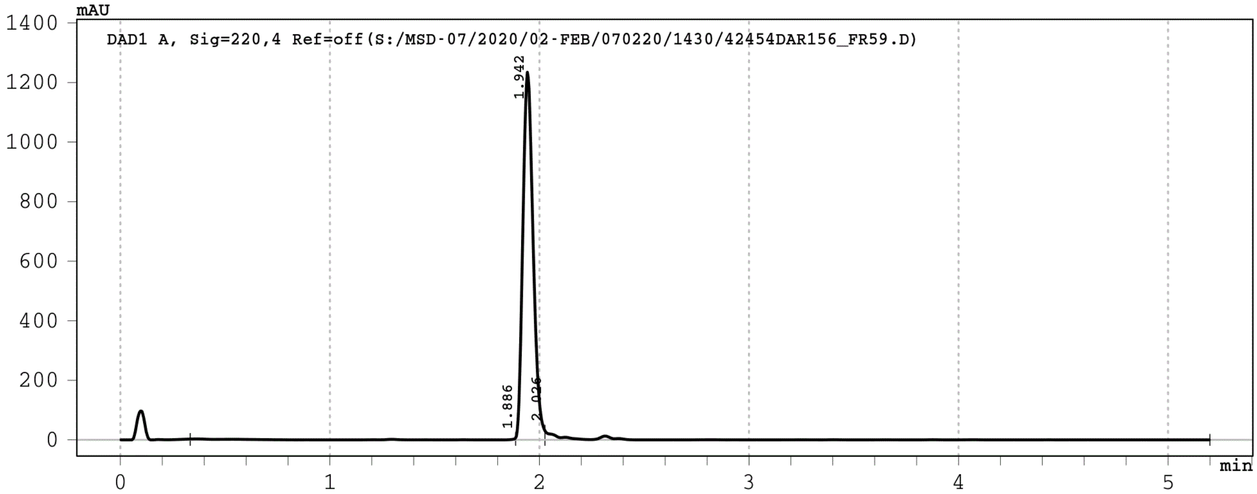


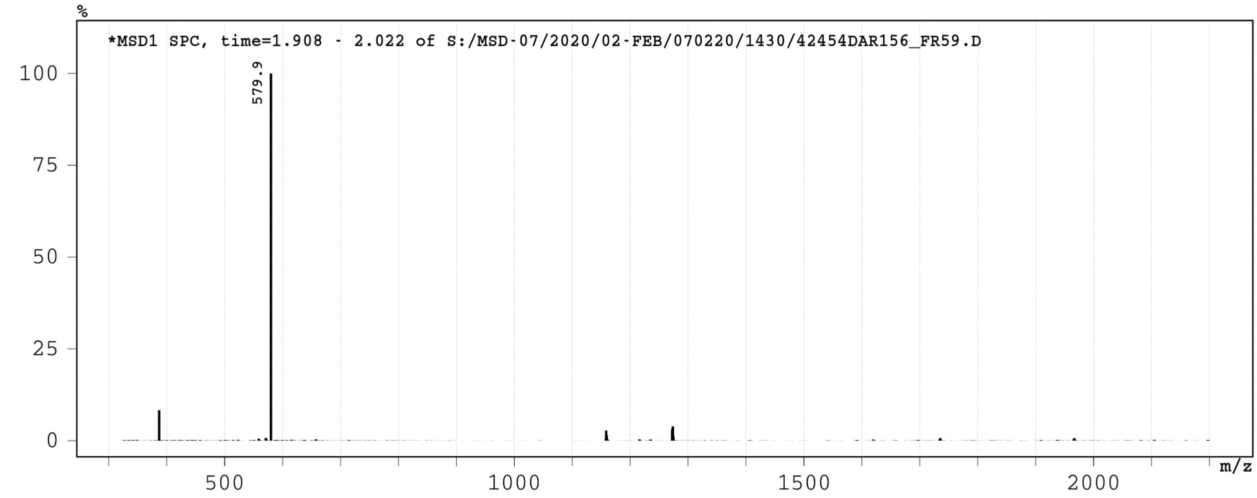


Signal 1: DAD1 A, Sig=220,4 Ref=off

| Peak # | RT [min] | Width [min] | Area | Area % |
|---|---|---|---|---|
| 1 | 1.886 | 1.55 | 57.3 | 1.3 |
| 2 | 1.942 | 0.14 | 4210.2 | 94.7 |
| 3 | 2.026 | 3.17 | 178.4 | 4.0 |

Data File : S:/MSD-07/2020/02-FEB/070220/1430/42454DAR157_FR6->

Injection Date : 07-Feb-20, 19:38:48
Sample Name : 42454DAR157_FR69-70
Remarks : Baseline corrected
Acq. Method : D:/METHODS/5-95-6M.M
column used : Je_559
Analysis Method : C:/CHEM32/1/METHODS/JPT-LCMS.M

Sample Info : 42454_157-Dr. Martine Darwish H-VRYVYGGI-OH

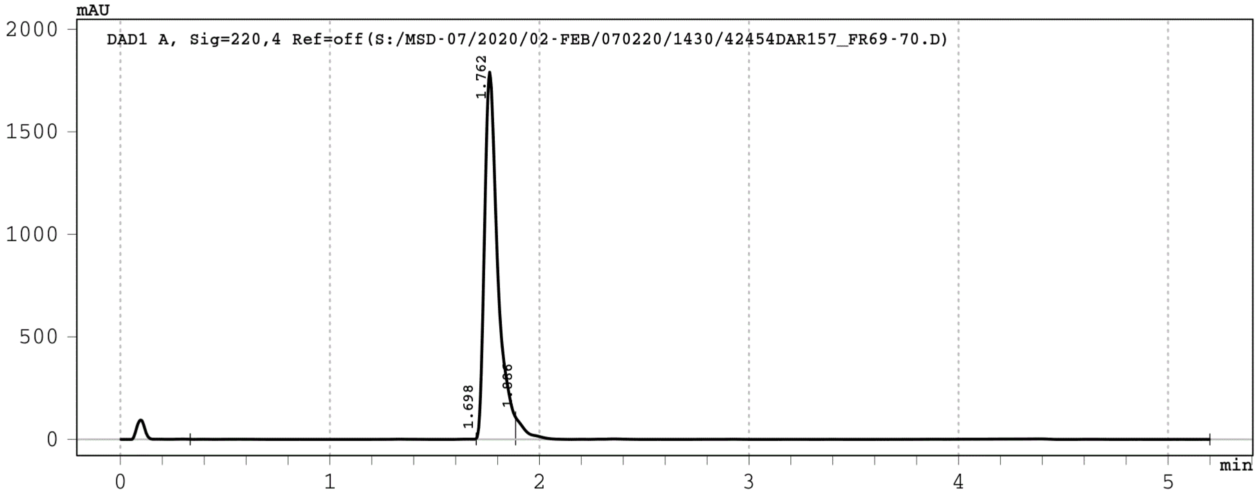


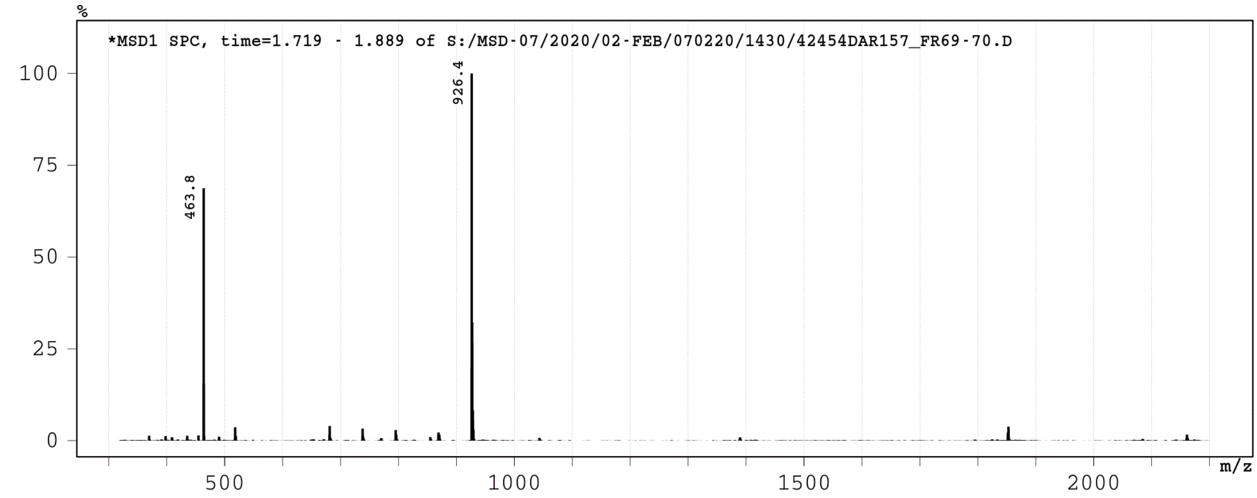


Signal 1: DAD1 A, Sig=220,4 Ref=off

| Peak # | RT [min] | Width [min] | Area | Area % |
|---|---|---|---|---|
| 1 | 1.698 | 1.37 | 23.4 | 0.3 |
| 2 | 1.762 | 0.19 | 7855.0 | 94.7 |
| 3 | 1.886 | 3.31 | 420.0 | 5.1 |

Data File : S:/MSD-07/2020/02-FEB/110220/1433/42454DAR158_FR3->

Injection Date : 12-Feb-20, 01:08:57
Sample Name : 42454DAR158_FR31
Remarks : Baseline corrected
Acq. Method : D:/METHODS/5-95-6M.M
column used : Je_559
Analysis Method : C:/CHEM32/1/METHODS/JPT-LCMS.M

Sample Info : 42454_158-Dr. Martine Darwish H-SPFLLITL-OH

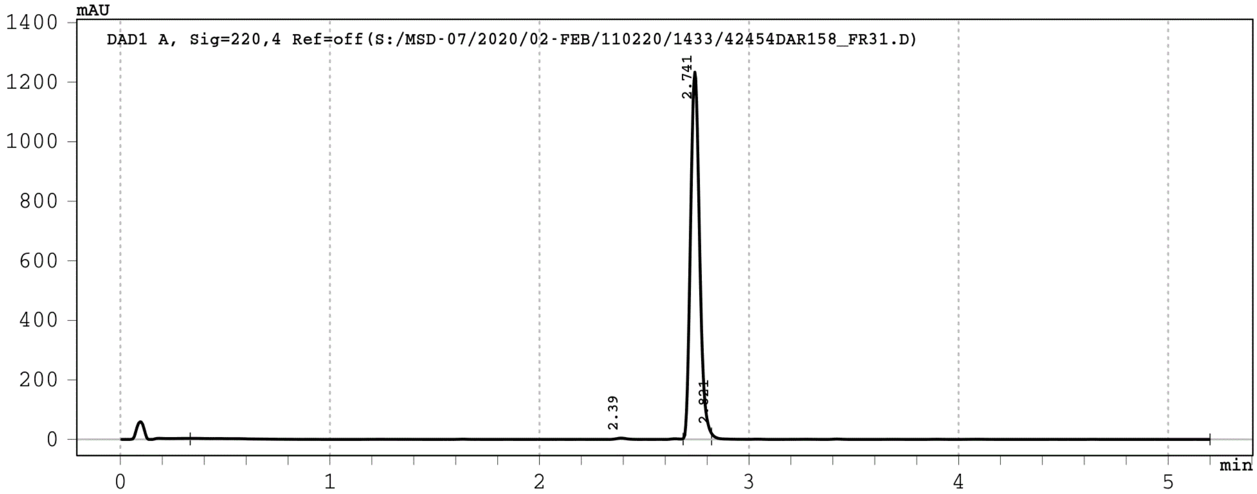


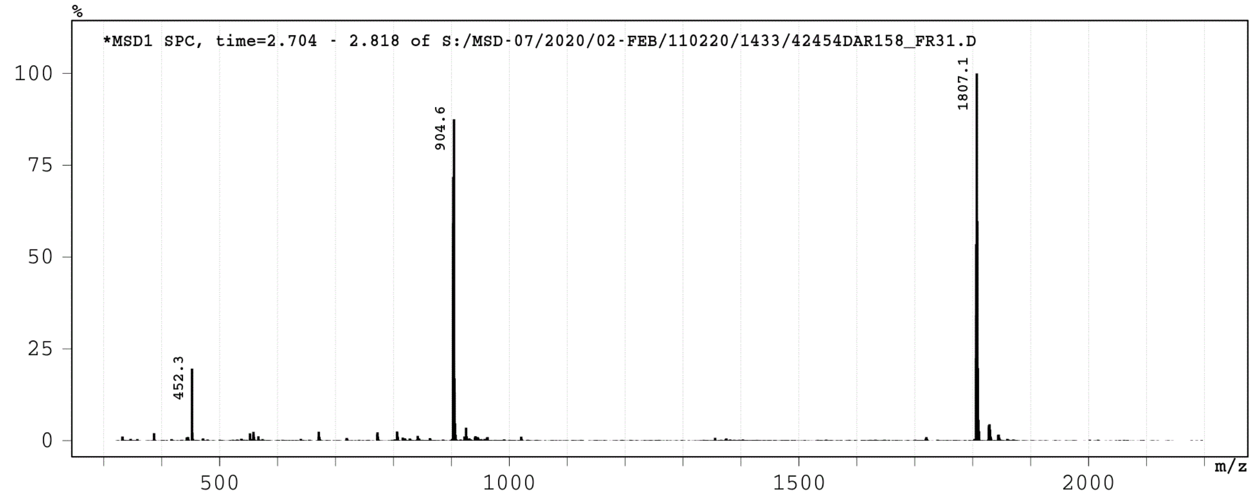


Signal 1: DAD1 A, Sig=220,4 Ref=off

| Peak # | RT [min] | Width [min] | Area | Area % |
|---|---|---|---|---|
| 1 | 2.390 | 2.35 | 86.3 | 2.2 |
| 2 | 2.741 | 0.14 | 3722.1 | 96.7 |
| 3 | 2.821 | 2.38 | 40.7 | 1.1 |

Data File : S:/MSD-07/2020/02-FEB/070220/1430/42454DAR159_FR7->

Injection Date : 07-Feb-20, 19:47:12
Sample Name : 42454DAR159_FR74
Remarks : Baseline corrected
Acq. Method : D:/METHODS/5-95-6M.M
column used : Je_559
Analysis Method : C:/CHEM32/1/METHODS/JPT-LCMS.M

Sample Info : 42454_159-Dr. Martine Darwish H-SAPPLLEGPL-OH

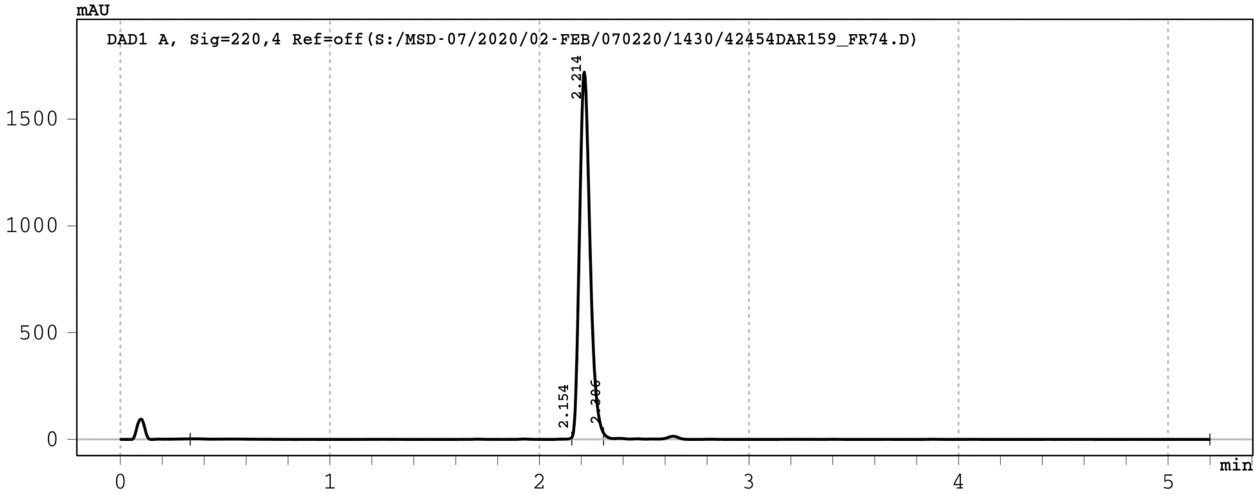


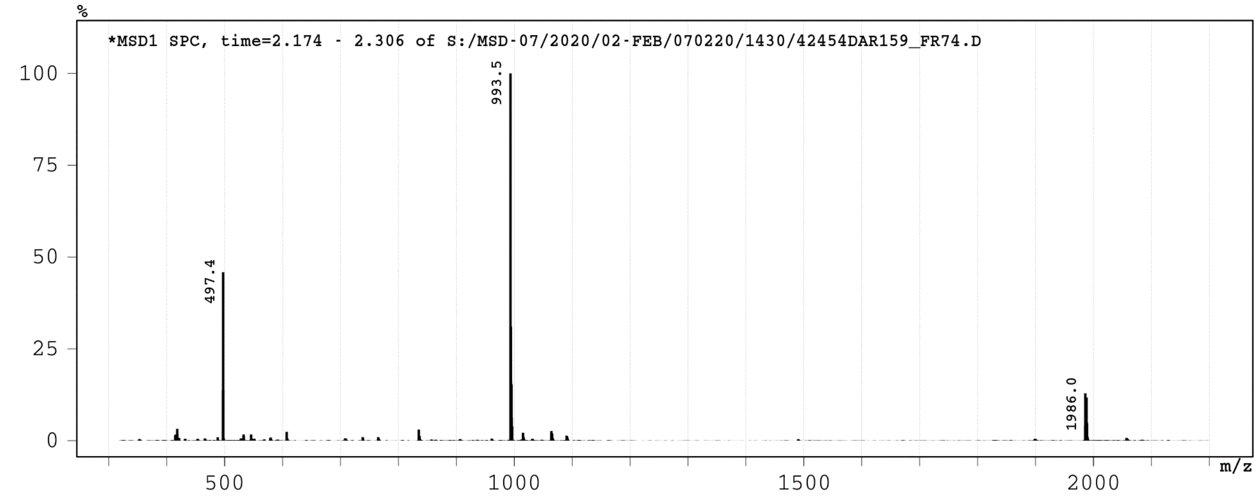


Signal 1: DAD1 A, Sig=220,4 Ref=off

| Peak # | RT [min] | Width [min] | Area | Area % |
|---|---|---|---|---|
| 1 | 2.154 | 1.82 | 55.7 | 0.9 |
| 2 | 2.214 | 0.15 | 6019.0 | 97.0 |
| 3 | 2.306 | 2.89 | 130.6 | 2.1 |

Data File : S:/MSD-07/2020/02-FEB/070220/1430/42454DAR160_FR7->

Injection Date : 07-Feb-20, 19:55:37
Sample Name : 42454DAR160_FR78-79
Remarks : Baseline corrected
Acq. Method : D:/METHODS/5-95-6M.M
column used : Je_559
Analysis Method : C:/CHEM32/1/METHODS/JPT-LCMS.M

Sample Info : 42454_160-Dr. Martine Darwish H-IQLVNGSLA-OH

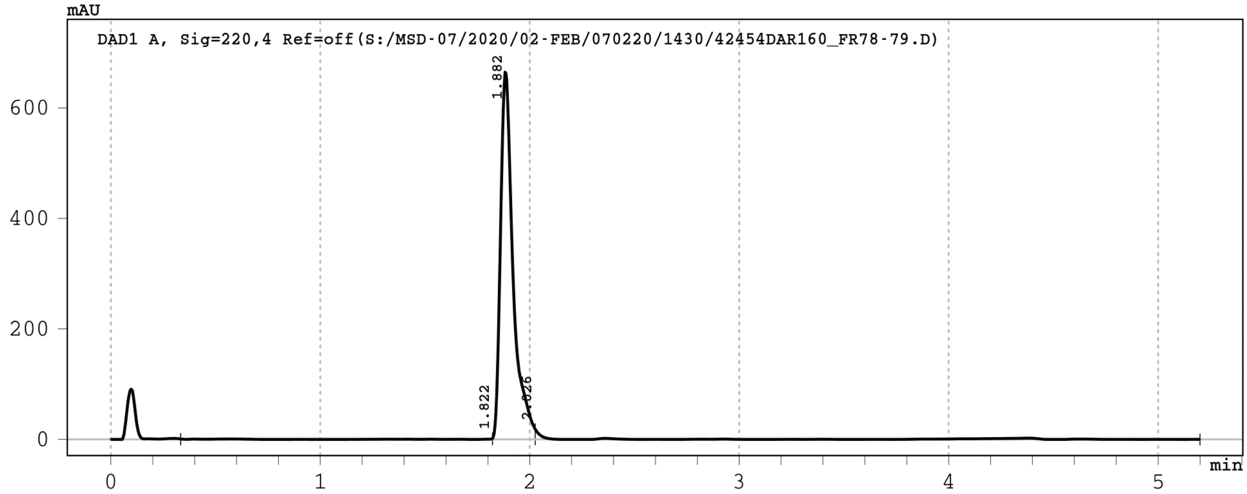


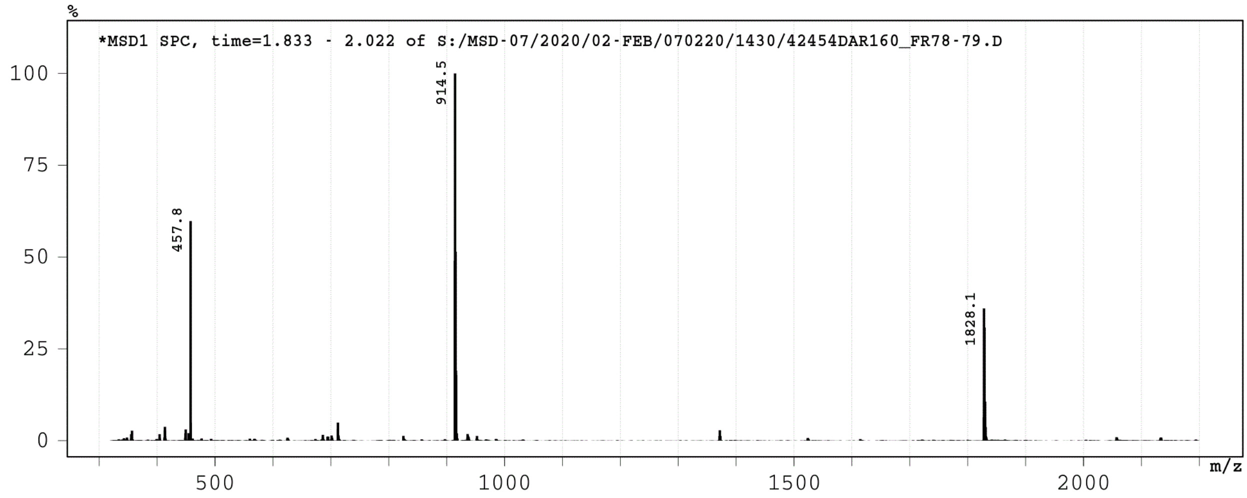


Signal 1: DAD1 A, Sig=220,4 Ref=off

| Peak # | RT [min] | Width [min] | Area | Area % |
|---|---|---|---|---|
| 1 | 1.822 | 1.49 | 20.4 | 0.7 |
| 2 | 1.882 | 0.20 | 2786.4 | 95.8 |
| 3 | 2.026 | 3.17 | 100.6 | 3.5 |

Data File : S:/MSD-07/2020/02-FEB/060220/1429/42454DAR161_FR7->

Injection Date : 06-Feb-20, 19:37:40
Sample Name : 42454DAR161_FR78-80
Remarks : Baseline corrected
Acq. Method : D:/METHODS/5-95-6M.M
column used : Je_559
Analysis Method : C:/CHEM32/1/METHODS/JPT-LCMS.M

Sample Info : 42454_161-Dr. Martine Darwish H-IFVRHMVGV-OH

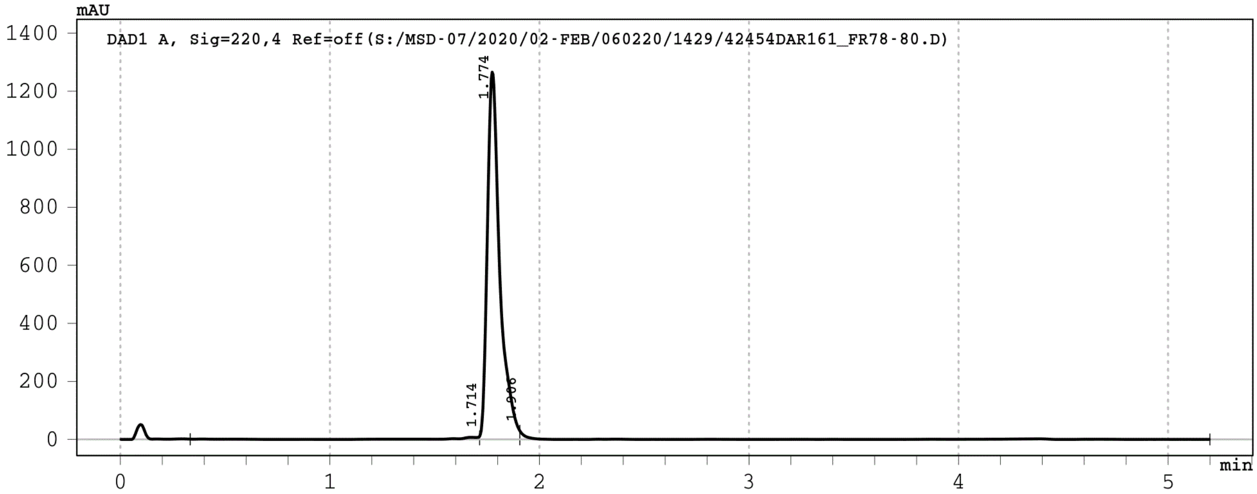


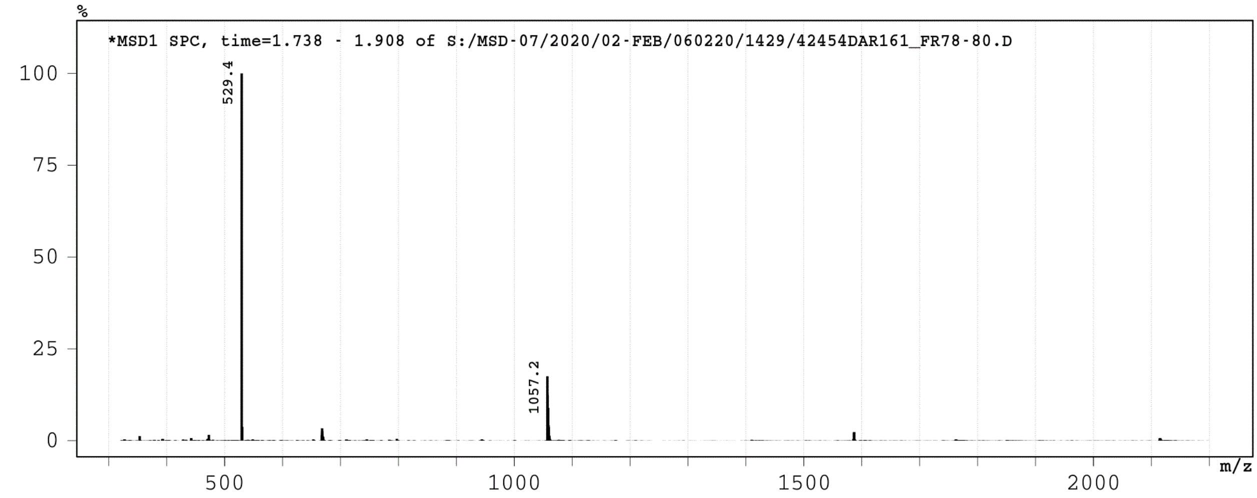


Signal 1: DAD1 A, Sig=220,4 Ref=off

| Peak # | RT [min] | Width [min] | Area | Area % |
|---|---|---|---|---|
| 1 | 1.714 | 1.38 | 76.3 | 1.4 |
| 2 | 1.774 | 0.19 | 5245.0 | 96.7 |
| 3 | 1.906 | 3.29 | 100.9 | 1.9 |

Data File : S:/MSD-07/2020/02-FEB/110220/1433/42454DAR162_FR4->

Injection Date : 12-Feb-20, 01:17:38
Sample Name : 42454DAR162_FR41-42
Remarks : Baseline corrected
Acq. Method : D:/METHODS/5-95-6M.M
column used : Je_559
Analysis Method : C:/CHEM32/1/METHODS/JPT-LCMS.M

Sample Info : 42454_162-Dr. Martine Darwish H-ICYGSAPL-OH

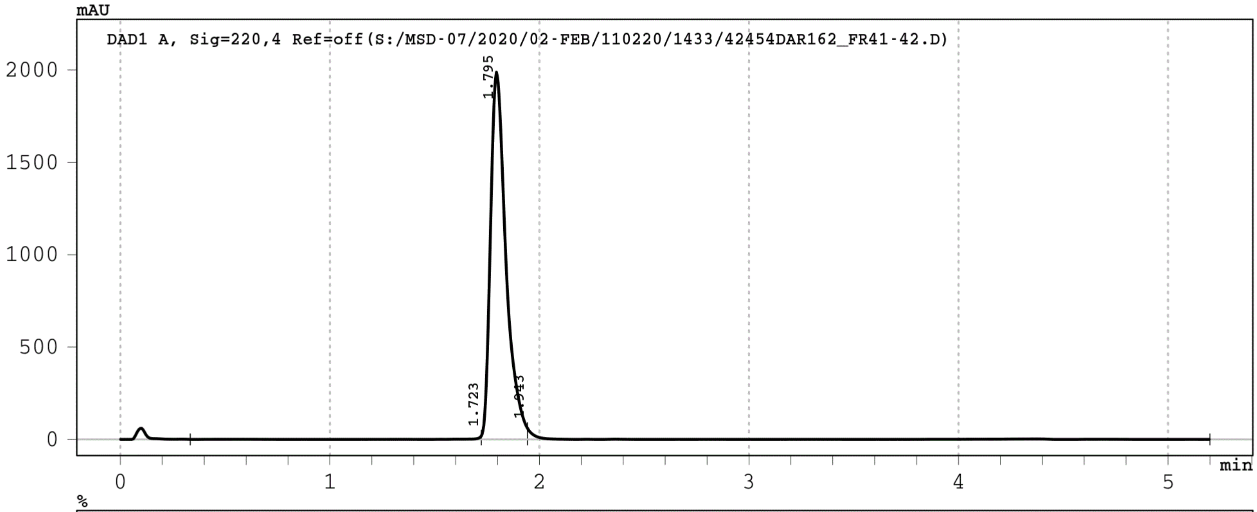


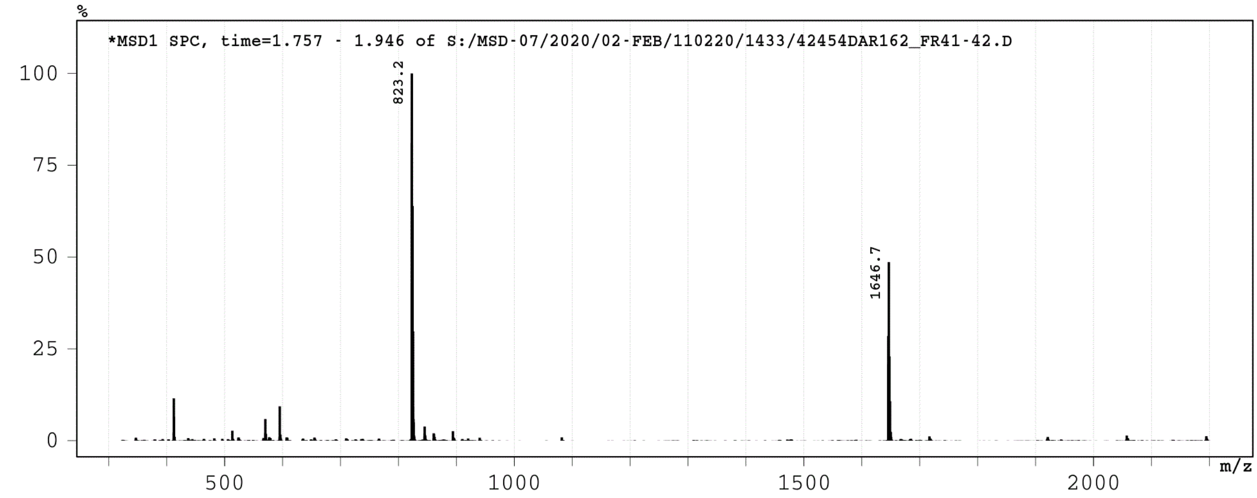


Signal 1: DAD1 A, Sig=220,4 Ref=off

| Peak # | RT [min] | Width [min] | Area | Area % |
|---|---|---|---|---|
| 1 | 1.723 | 1.39 | 37.5 | 0.4 |
| 2 | 1.795 | 0.22 | 10228.6 | 97.8 |
| 3 | 1.943 | 3.26 | 187.6 | 1.8 |

Data File : S:/MSD-07/2020/02-FEB/110220/1433/42454DAR163_FR5->

Injection Date : 12-Feb-20, 01:26:47
Sample Name : 42454DAR163_FR59-61
Remarks : Baseline corrected
Acq. Method : D:/METHODS/5-95-6M.M
column used : Je_559
Analysis Method : C:/CHEM32/1/METHODS/JPT-LCMS.M

Sample Info : 42454_163-Dr. Martine Darwish H-YIPLFPHL-OH

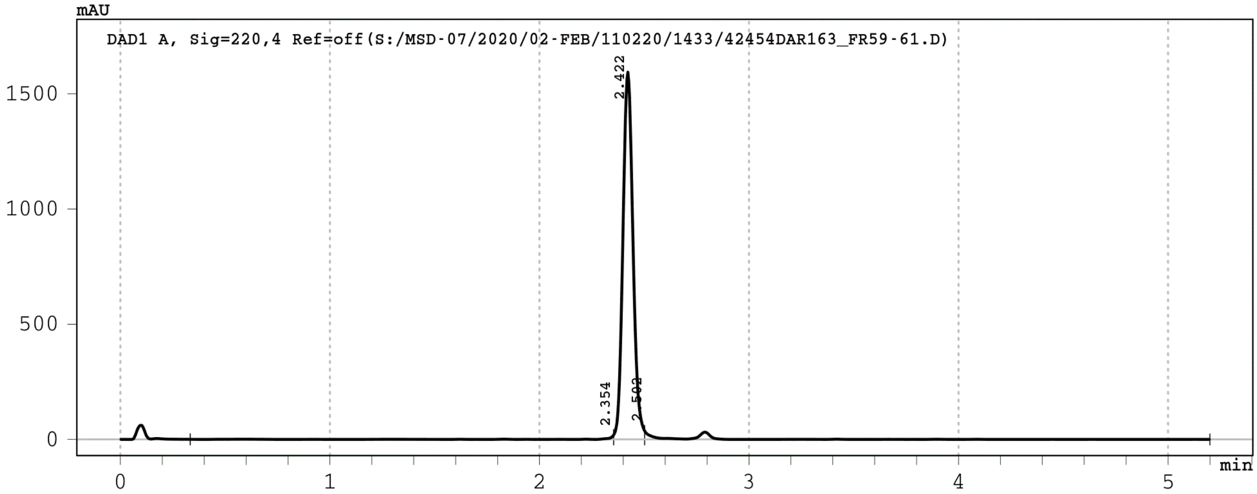


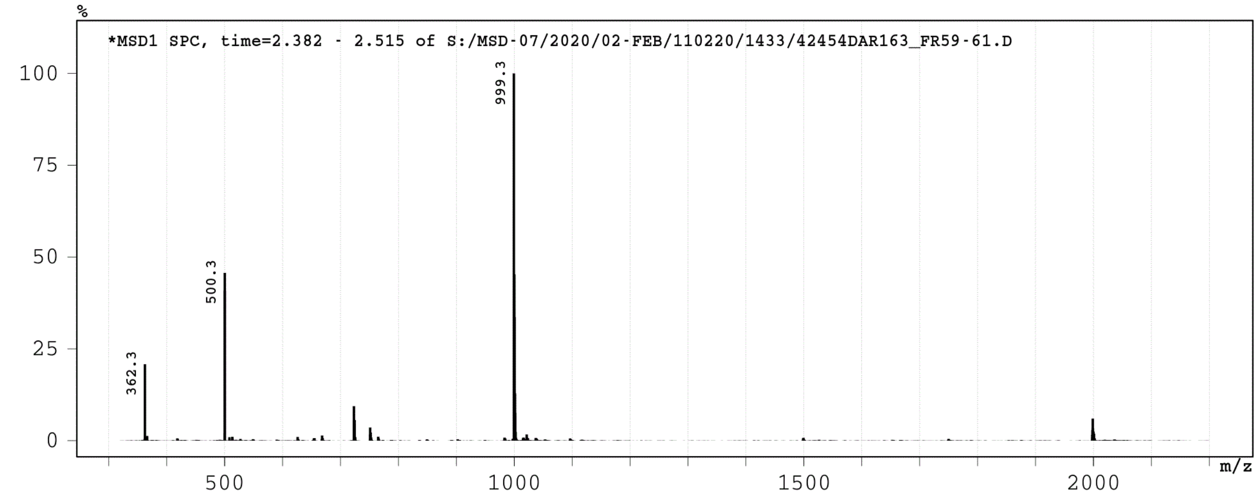


Signal 1: DAD1 A, Sig=220,4 Ref=off

| Peak # | RT [min] | Width [min] | Area | Area % |
|---|---|---|---|---|
| 1 | 2.354 | 2.02 | 46.2 | 0.8 |
| 2 | 2.422 | 0.15 | 5337.2 | 95.3 |
| 3 | 2.502 | 2.70 | 214.9 | 3.8 |

Data File : S:/MSD-07/2020/02-FEB/070220/1430/42454DAR164_FR8->

Injection Date : 07-Feb-20, 20:03:59
Sample Name : 42454DAR164_FR82
Remarks : Baseline corrected
Acq. Method : D:/METHODS/5-95-6M.M
column used : Je_559
Analysis Method : C:/CHEM32/1/METHODS/JPT-LCMS.M

Sample Info : 42454_164-Dr. Martine Darwish H-TAVVRQSTL-OH

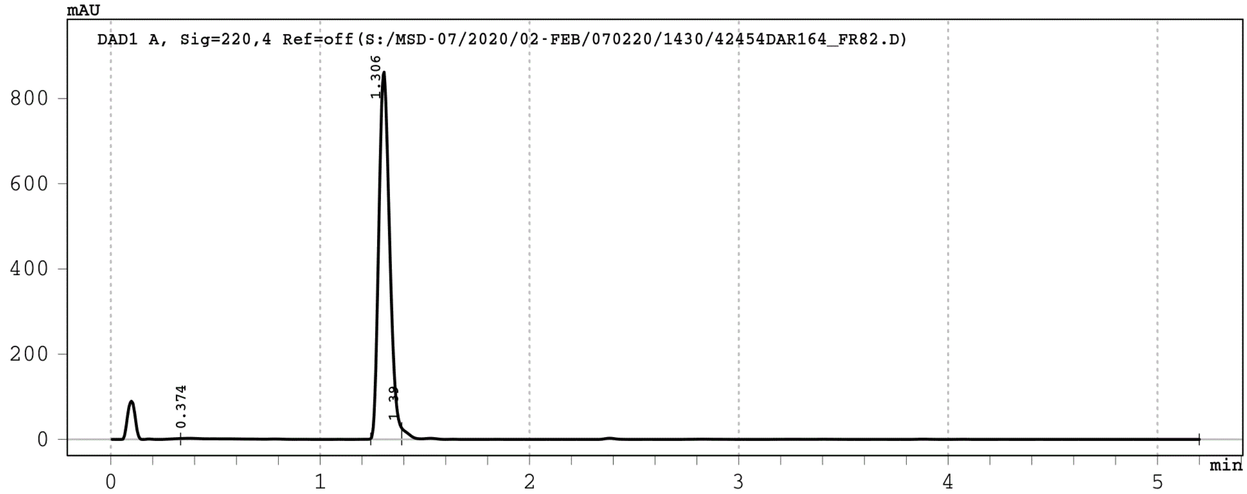


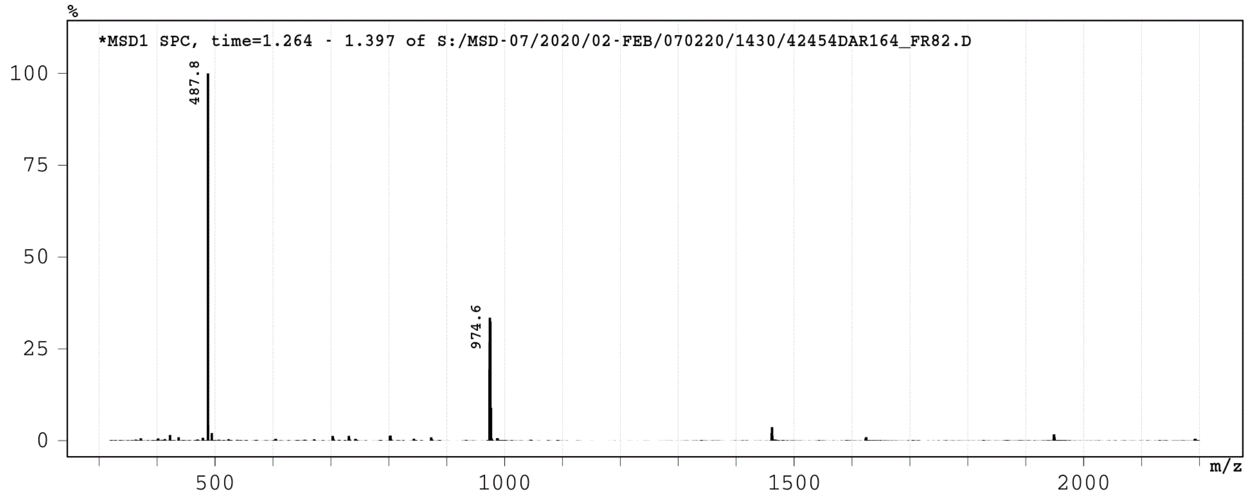


Signal 1: DAD1 A, Sig=220,4 Ref=off

| Peak # | RT [min] | Width [min] | Area | Area % |
|---|---|---|---|---|
| 1 | 0.374 | 0.91 | 41.1 | 1.3 |
| 2 | 1.306 | 0.15 | 3103.1 | 96.0 |
| 3 | 1.390 | 3.81 | 87.3 | 2.7 |

Data File : S:/MSD-07/2020/02-FEB/070220/1430/42454DAR165_FR9->

Injection Date : 07-Feb-20, 20:12:21
Sample Name : 42454DAR165_FR9-10
Remarks : Baseline corrected
Acq. Method : D:/METHODS/5-95-6M.M
column used : Je_559
Analysis Method : C:/CHEM32/1/METHODS/JPT-LCMS.M

Sample Info : 42454_165-Dr. Martine Darwish H-SYVSLWPDL-OH

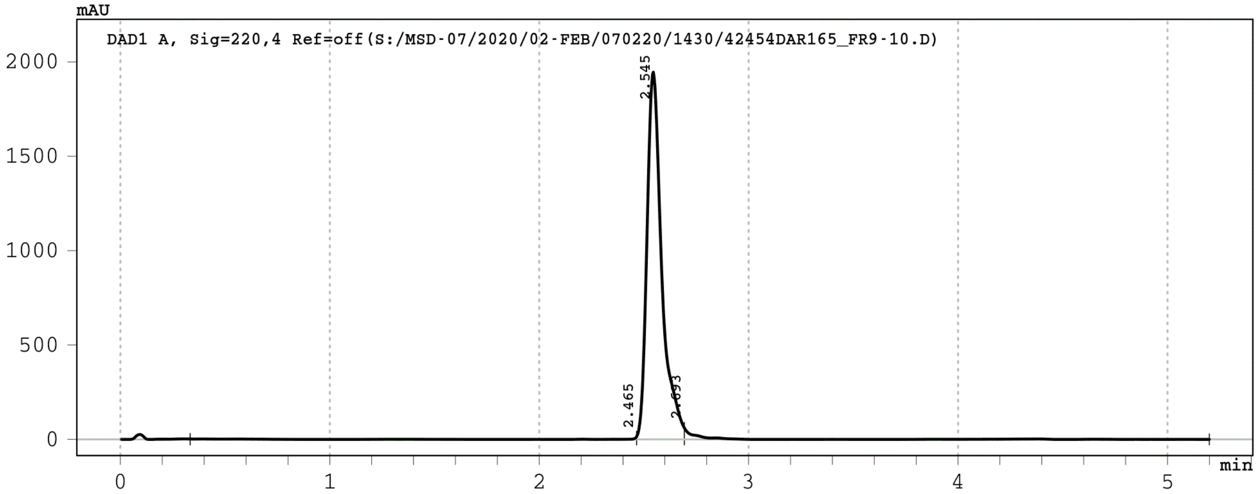


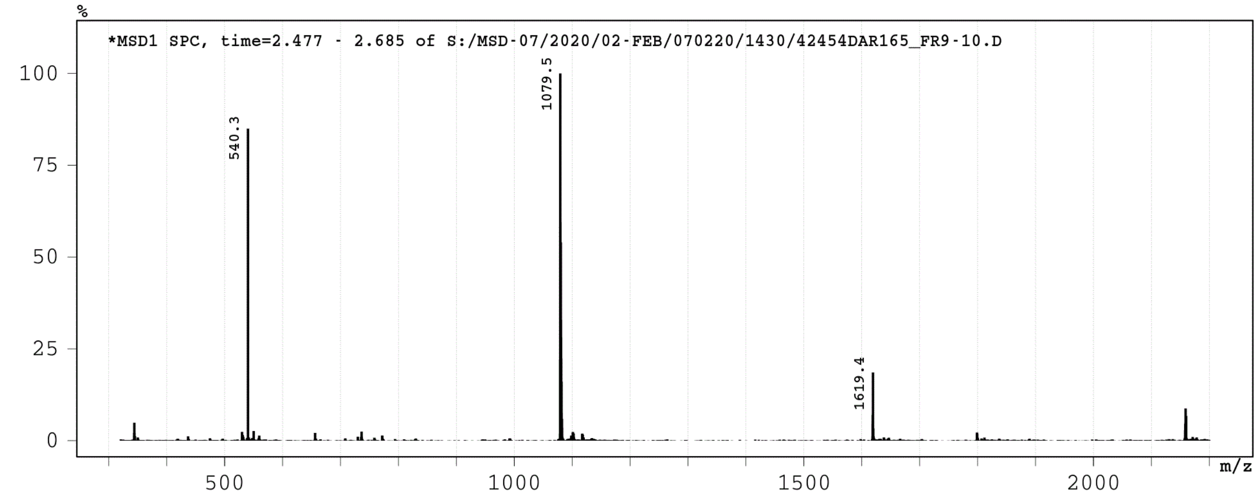


Signal 1: DAD1 A, Sig=220,4 Ref=off

| Peak # | RT [min] | Width [min] | Area | Area % |
|---|---|---|---|---|
| 1 | 2.465 | 2.13 | 72.3 | 0.7 |
| 2 | 2.545 | 0.23 | 9574.7 | 96.7 |
| 3 | 2.693 | 2.51 | 258.8 | 2.6 |

Data File : S:/MSD-07/2020/02-FEB/110220/1433/42454DAR166_FR7->

Injection Date : 12-Feb-20, 01:35:20
Sample Name : 42454DAR166_FR72-74
Remarks : Baseline corrected
Acq. Method : D:/METHODS/5-95-6M.M
column used : Je_559
Analysis Method : C:/CHEM32/1/METHODS/JPT-LCMS.M

Sample Info : 42454_166-Dr. Martine Darwish H-STPQLLPL-OH

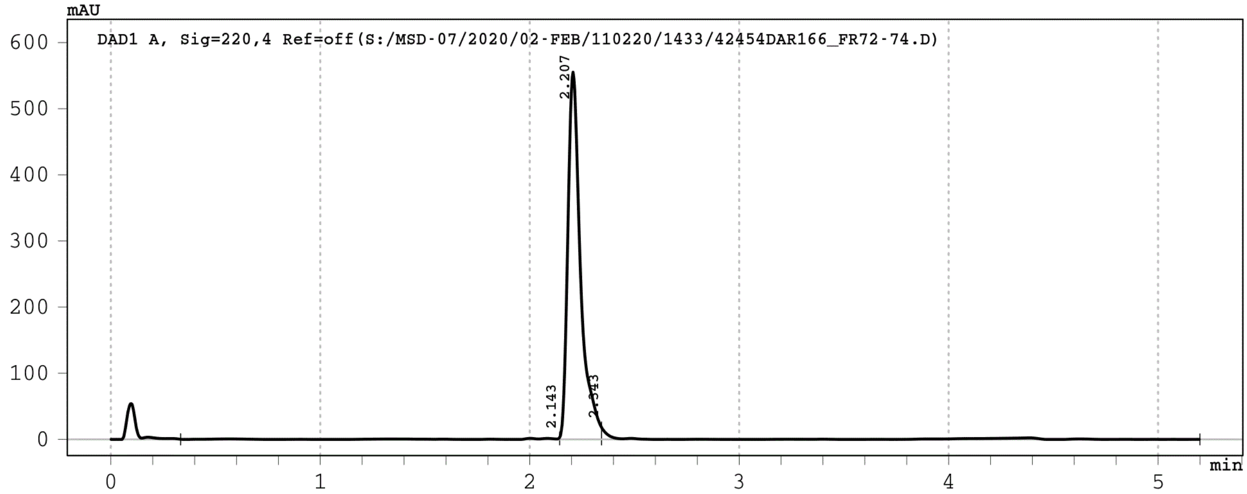


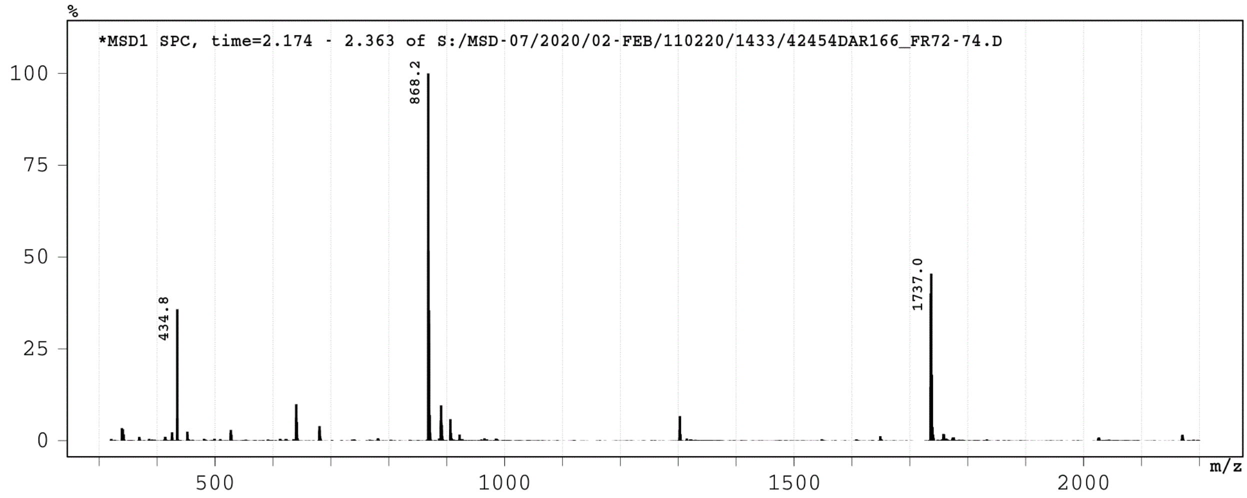


Signal 1: DAD1 A, Sig=220,4 Ref=off

| Peak # | RT [min] | Width [min] | Area | Area % |
|---|---|---|---|---|
| 1 | 2.143 | 1.81 | 33.7 | 1.4 |
| 2 | 2.207 | 0.20 | 2352.2 | 94.7 |
| 3 | 2.343 | 2.86 | 98.0 | 3.9 |

Data File : S:/MSD-07/2020/02-FEB/070220/1430/42454DAR167_FR2->

Injection Date : 07-Feb-20, 20:20:44
Sample Name : 42454DAR167_FR22-23
Remarks : Baseline corrected
Acq. Method : D:/METHODS/5-95-6M.M
column used : Je_559
Analysis Method : C:/CHEM32/1/METHODS/JPT-LCMS.M

Sample Info : 42454_167-Dr. Martine Darwish H-FVLESYLNL-OH

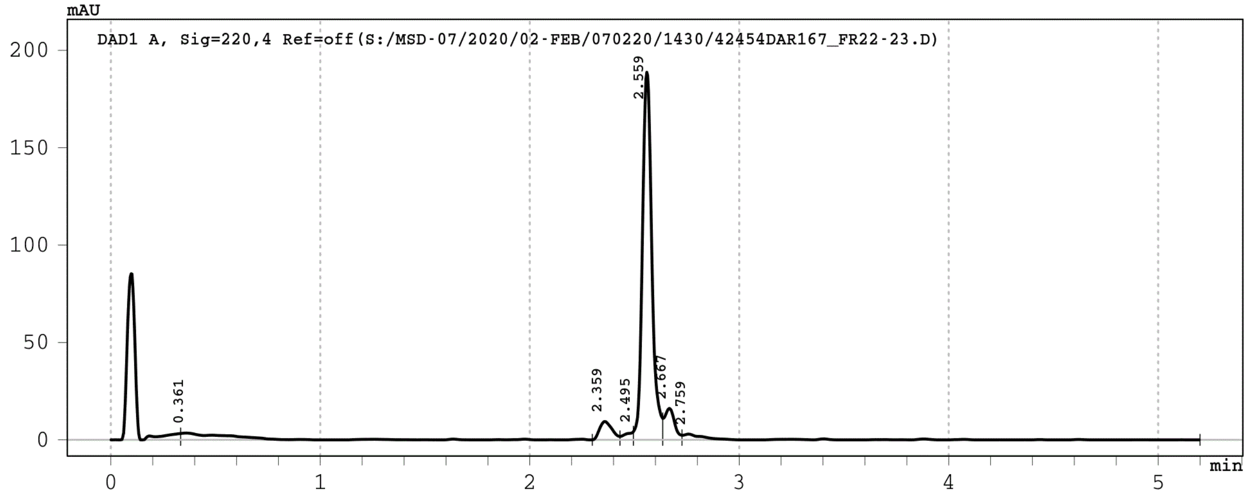


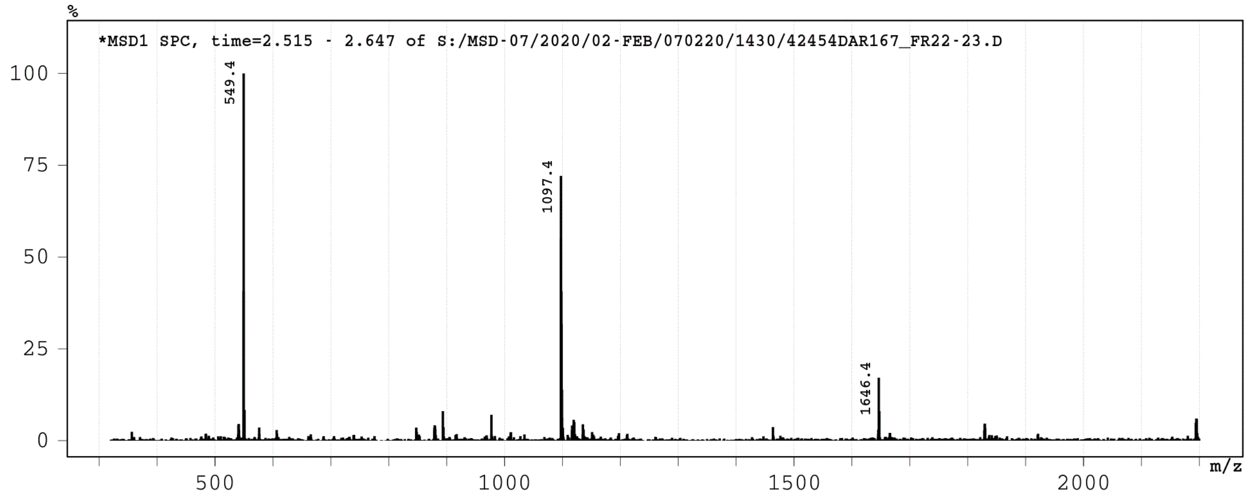


Signal 1: DAD1 A, Sig=220,4 Ref=off

| Peak # | RT [min] | Width [min] | Area | Area % |
|---|---|---|---|---|
| 1 | 0.361 | 1.97 | 59.6 | 7.6 |
| 2 | 2.359 | 0.13 | 39.4 | 5.0 |
| 3 | 2.495 | 0.06 | 11.2 | 1.4 |
| 4 | 2.559 | 0.14 | 593.8 | 75.5 |
| 5 | 2.667 | 0.09 | 54.8 | 7.0 |
| 6 | 2.759 | 2.47 | 27.5 | 3.5 |

Data File : S:/MSD-07/2020/02-FEB/060220/1429/42454DAR168_FR8->

Injection Date : 06-Feb-20, 19:46:02
Sample Name : 42454DAR168_FR84+1
Remarks : Baseline corrected
Acq. Method : D:/METHODS/5-95-6M.M
column used : Je_559
Analysis Method : C:/CHEM32/1/METHODS/JPT-LCMS.M

Sample Info : 42454_168-Dr. Martine Darwish H-MSRCNHNRL-OH

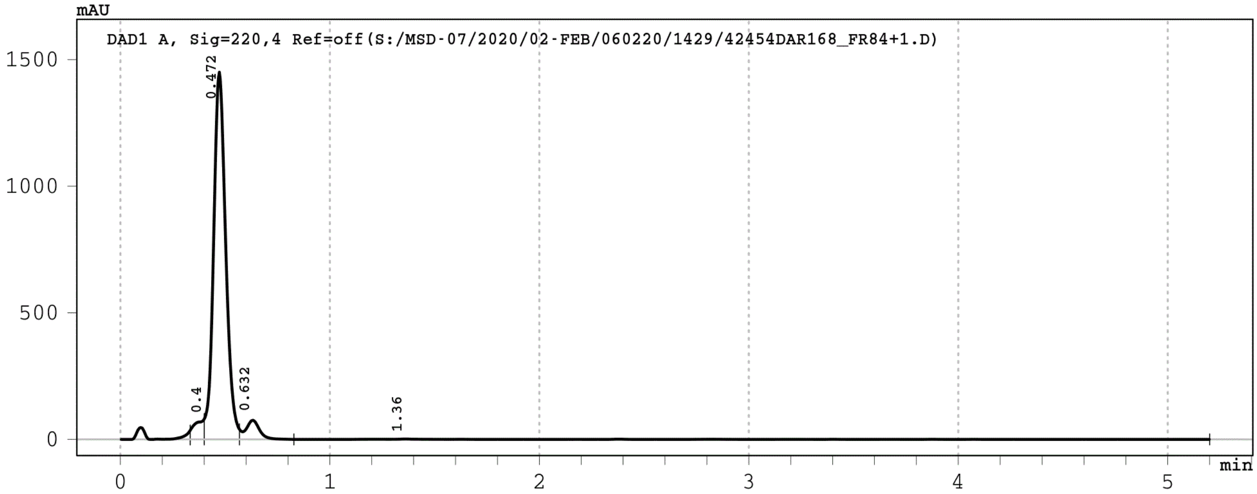


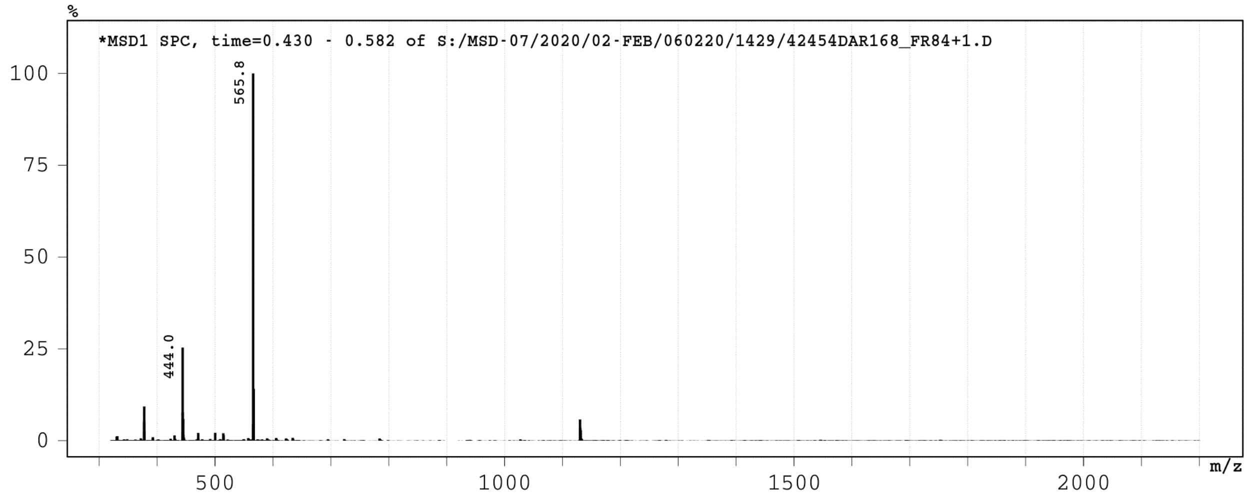


Signal 1: DAD1 A, Sig=220,4 Ref=off

| Peak # | RT [min] | Width [min] | Area | Area % |
|---|---|---|---|---|
| 1 | 0.400 | 0.07 | 244.5 | 3.8 |
| 2 | 0.472 | 0.17 | 5847.2 | 90.2 |
| 3 | 0.632 | 0.26 | 358.2 | 5.5 |
| 4 | 1.360 | 4.37 | 32.9 | 0.5 |

Data File : S:/MSD-07/2020/02-FEB/070220/1430/42454DAR169_FR3->

Injection Date : 07-Feb-20, 20:29:05
Sample Name : 42454DAR169_FR31
Remarks : Baseline corrected
Acq. Method : D:/METHODS/5-95-6M.M
column used : Je_559
Analysis Method : C:/CHEM32/1/METHODS/JPT-LCMS.M

Sample Info : 42454_169-Dr. Martine Darwish H-SAWEVLIPL-OH

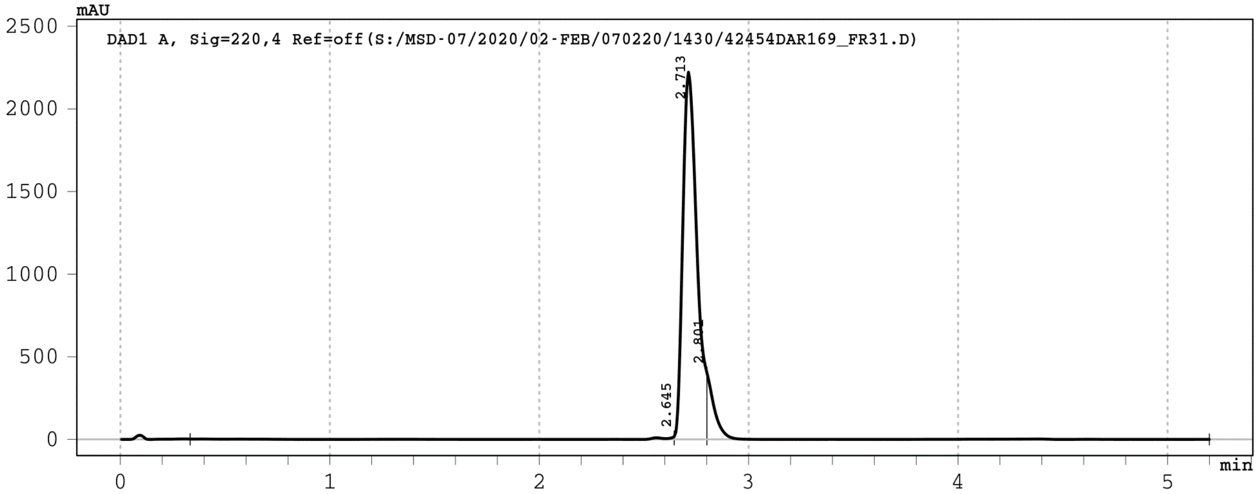


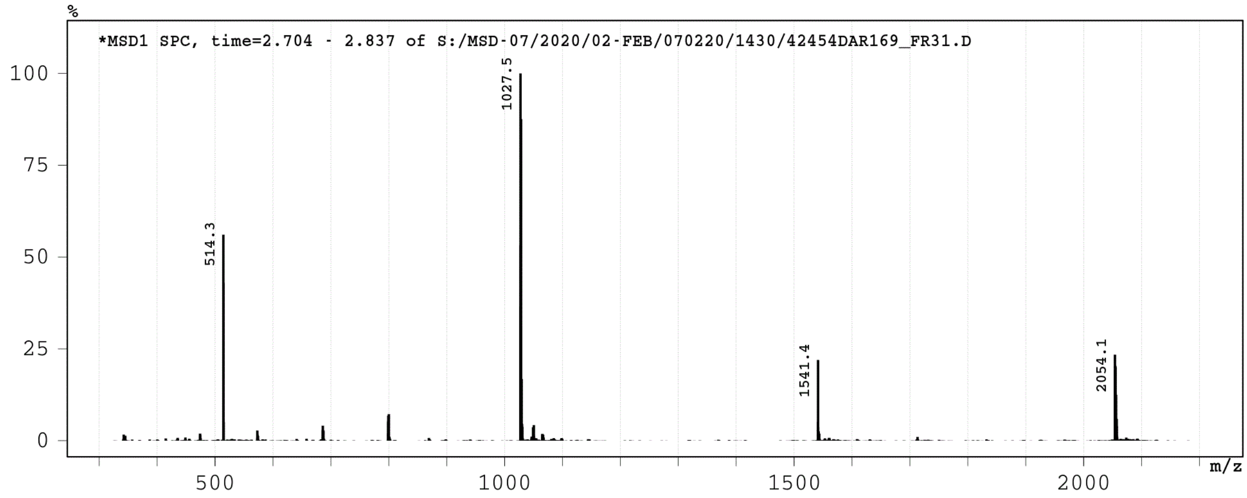


Signal 1: DAD1 A, Sig=220,4 Ref=off

| Peak # | RT [min] | Width [min] | Area | Area % |
|---|---|---|---|---|
| 1 | 2.645 | 2.31 | 112.2 | 1.0 |
| 2 | 2.713 | 0.16 | 10343.1 | 90.0 |
| 3 | 2.801 | 2.40 | 1034.6 | 9.0 |

Data File : S:/MSD-07/2020/02-FEB/070220/1430/42454DAR170_FR4->

Injection Date : 07-Feb-20, 20:37:27
Sample Name : 42454DAR170_FR41-42
Remarks : Baseline corrected
Acq. Method : D:/METHODS/5-95-6M.M
column used : Je_559
Analysis Method : C:/CHEM32/1/METHODS/JPT-LCMS.M

Sample Info : 42454_170-Dr. Martine Darwish H-ATLAQFNTV-OH

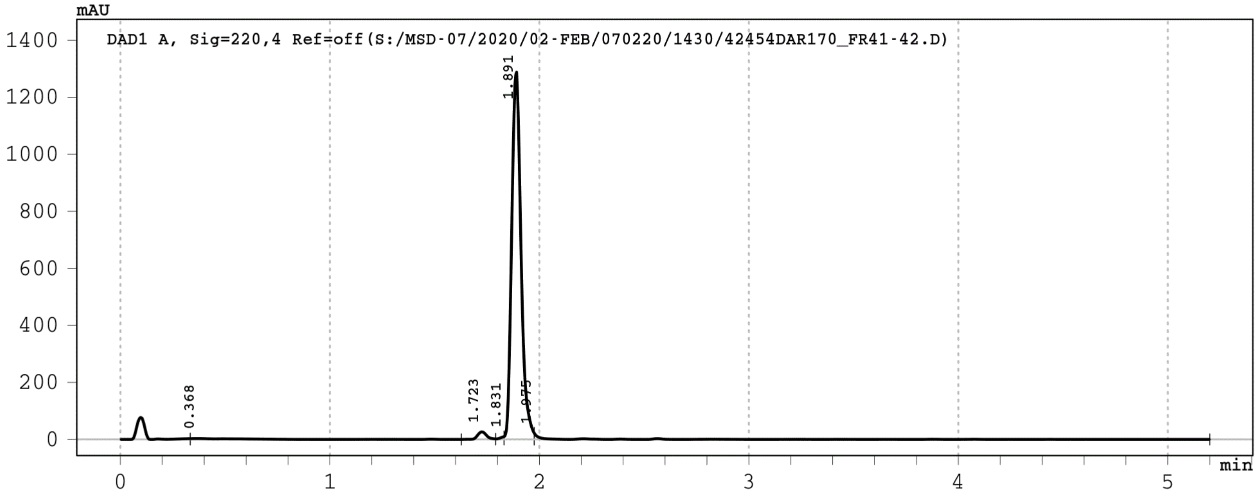


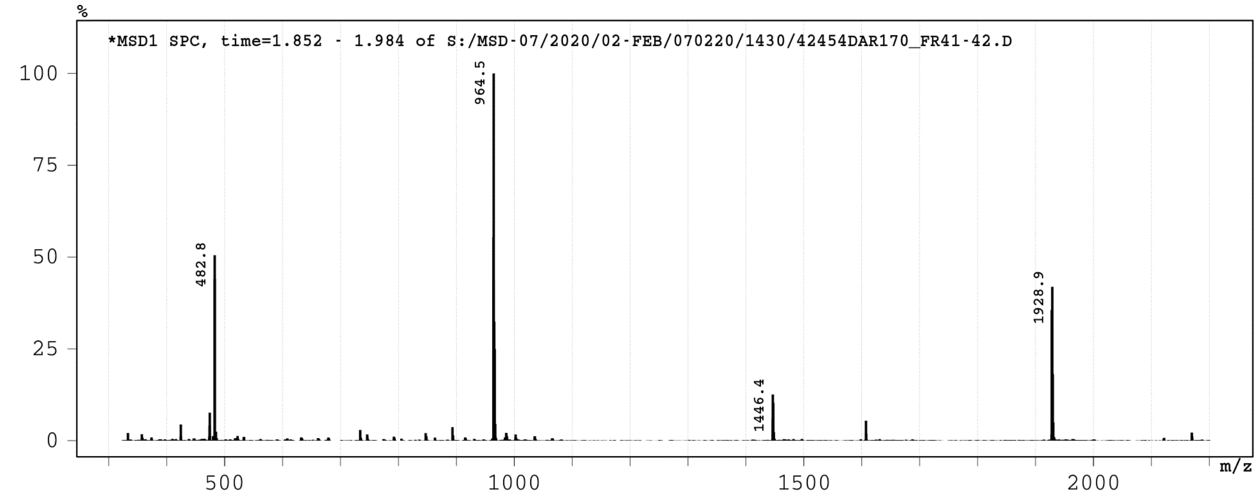


Signal 1: DAD1 A, Sig=220,4 Ref=off

| Peak # | RT [min] | Width [min] | Area | Area % |
|---|---|---|---|---|
| 1 | 0.368 | 1.29 | 51.7 | 1.2 |
| 2 | 1.723 | 0.16 | 79.4 | 1.9 |
| 3 | 1.831 | 0.04 | 12.1 | 0.3 |
| 4 | 1.891 | 0.14 | 3979.0 | 95.0 |
| 5 | 1.975 | 3.22 | 65.4 | 1.6 |

Data File : S:/MSD-07/2020/02-FEB/060220/1429/42454DAR171_FR8->

Injection Date : 06-Feb-20, 19:54:27
Sample Name : 42454DAR171_FR8-9
Remarks : Baseline corrected
Acq. Method : D:/METHODS/5-95-6M.M
column used : Je_559
Analysis Method : C:/CHEM32/1/METHODS/JPT-LCMS.M

Sample Info : 42454_171-Dr. Martine Darwish H-FAMLGTIFL-OH

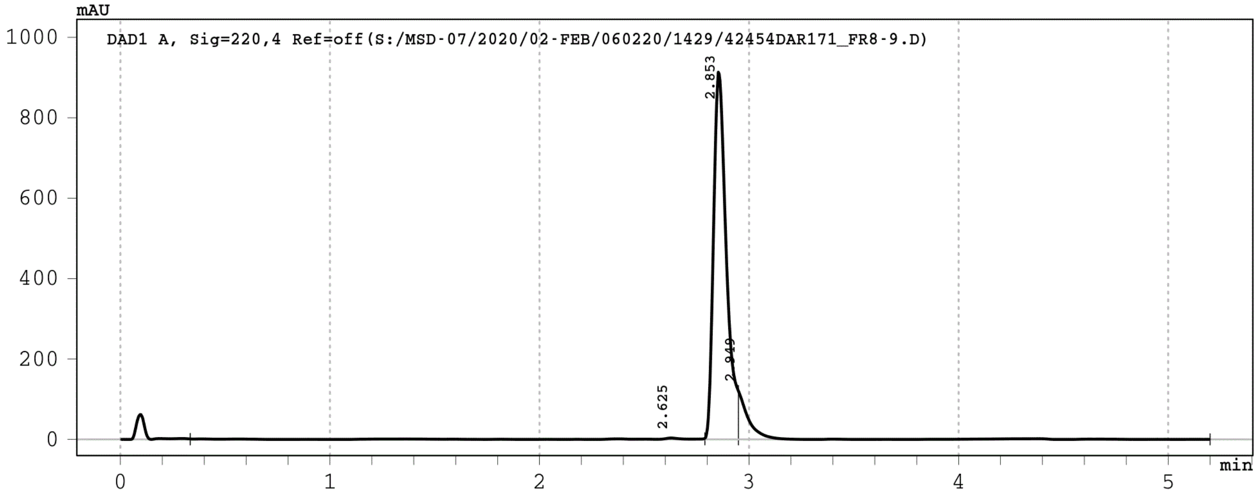


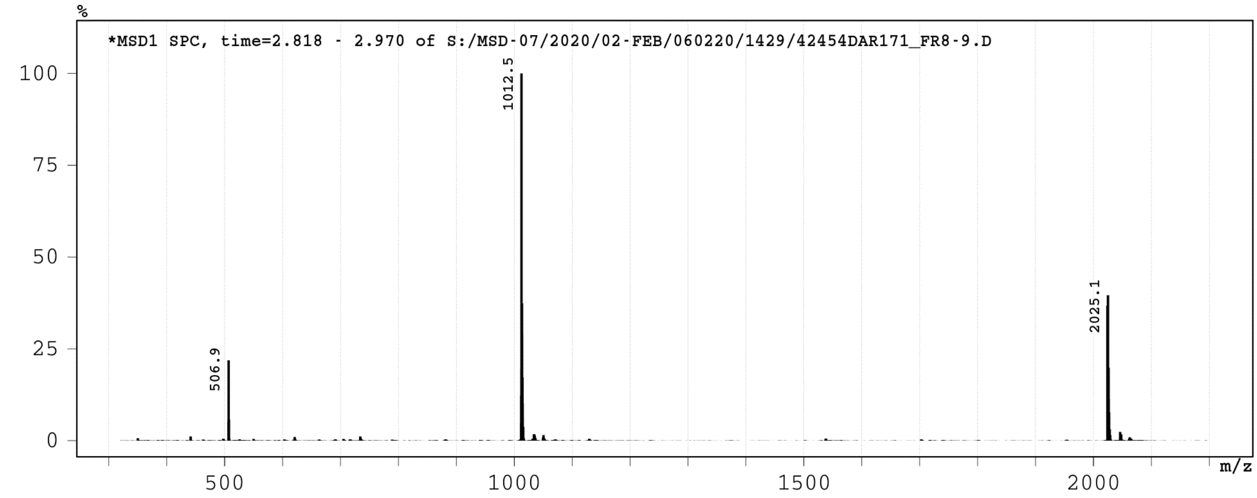


Signal 1: DAD1 A, Sig=220,4 Ref=off

| Peak # | RT [min] | Width [min] | Area | Area % |
|---|---|---|---|---|
| 1 | 2.625 | 2.46 | 74.0 | 1.7 |
| 2 | 2.853 | 0.16 | 3886.1 | 88.4 |
| 3 | 2.949 | 2.25 | 433.5 | 9.9 |

Data File : S:/MSD-07/2020/02-FEB/060220/1429/42454DAR172_FR1->

Injection Date : 06-Feb-20, 20:02:50
Sample Name : 42454DAR172_FR16-17
Remarks : Baseline corrected
Acq. Method : D:/METHODS/5-95-6M.M
column used : Je_559
Analysis Method : C:/CHEM32/1/METHODS/JPT-LCMS.M

Sample Info : 42454_172-Dr. Martine Darwish H-VAQNRHCEL-OH

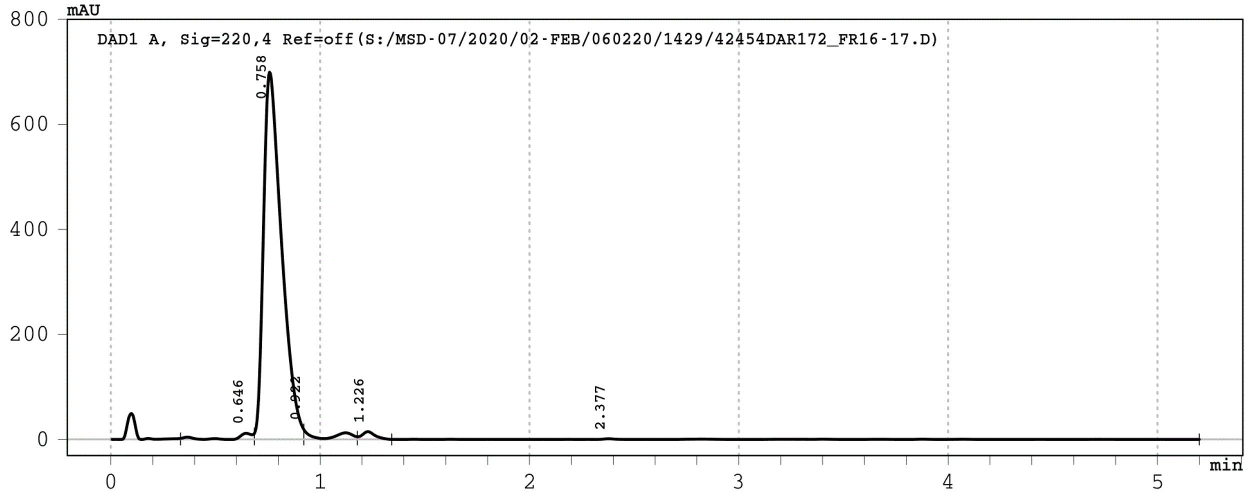


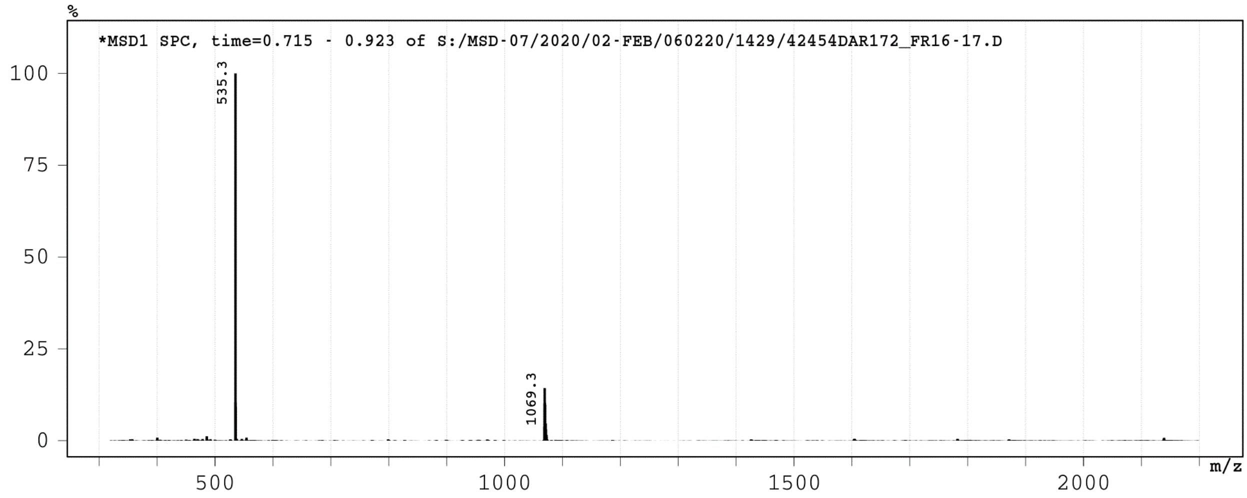


Signal 1: DAD1 A, Sig=220,4 Ref=off

| Peak # | RT [min] | Width [min] | Area | Area % |
|---|---|---|---|---|
| 1 | 0.646 | 0.35 | 63.2 | 1.4 |
| 2 | 0.758 | 0.24 | 4215.8 | 94.4 |
| 3 | 0.922 | 0.26 | 105.1 | 2.4 |
| 4 | 1.226 | 0.16 | 64.6 | 1.4 |
| 5 | 2.377 | 3.86 | 19.1 | 0.4 |

Data File : S:/MSD-07/2020/02-FEB/070220/1430/42454DAR173_FR5->

Injection Date : 07-Feb-20, 20:45:50
Sample Name : 42454DAR173_FR50
Remarks : Baseline corrected
Acq. Method : D:/METHODS/5-95-6M.M
column used : Je_559
Analysis Method : C:/CHEM32/1/METHODS/JPT-LCMS.M

Sample Info : 42454_173-Dr. Martine Darwish H-YVWGRYDFL-OH

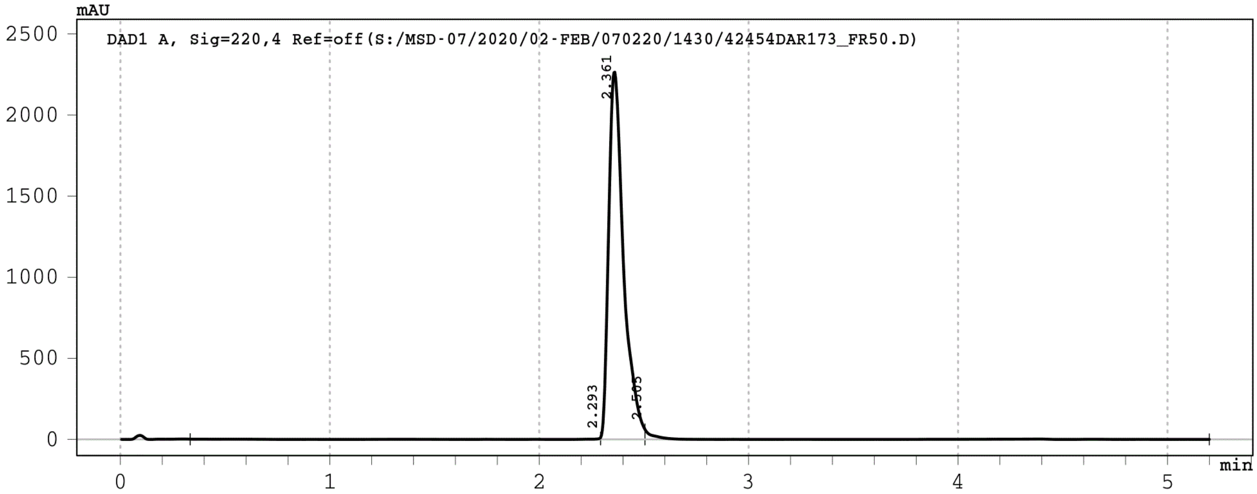


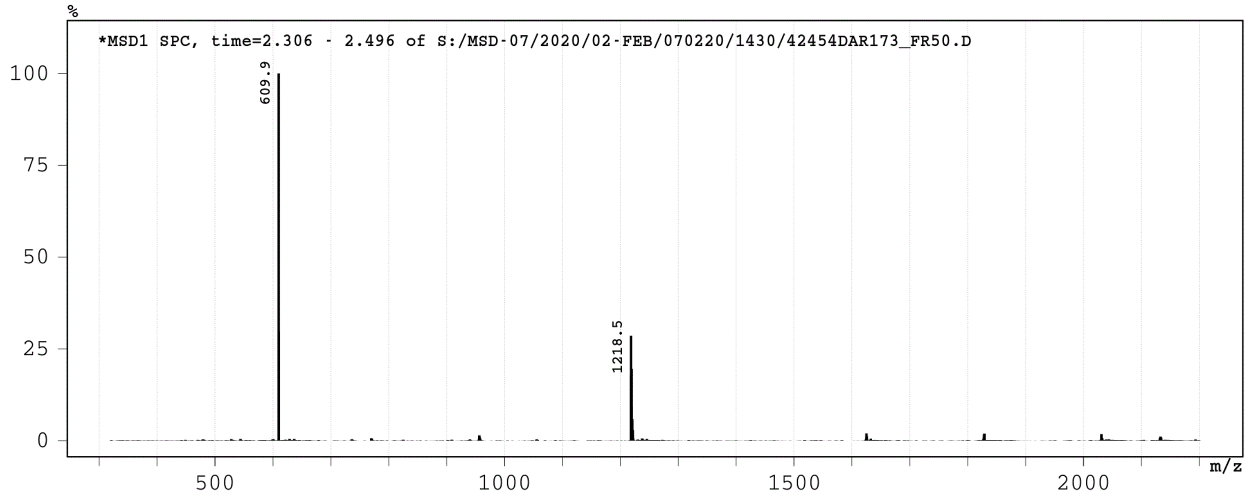


Signal 1: DAD1 A, Sig=220,4 Ref=off

| Peak # | RT [min] | Width [min] | Area | Area % |
|---|---|---|---|---|
| 1 | 2.293 | 1.96 | 52.7 | 0.5 |
| 2 | 2.361 | 0.21 | 11232.2 | 97.7 |
| 3 | 2.505 | 2.70 | 211.2 | 1.8 |

Data File : S:/MSD-07/2020/02-FEB/060220/1429/42454DAR174_FR2->

Injection Date : 06-Feb-20, 20:19:38
Sample Name : 42454DAR174_FR22-23
Remarks : Baseline corrected
Acq. Method : D:/METHODS/5-95-6M.M
column used : Je_559
Analysis Method : C:/CHEM32/1/METHODS/JPT-LCMS.M

Sample Info : 42454_174-Dr. Martine Darwish H-MNMGQFLEF-OH

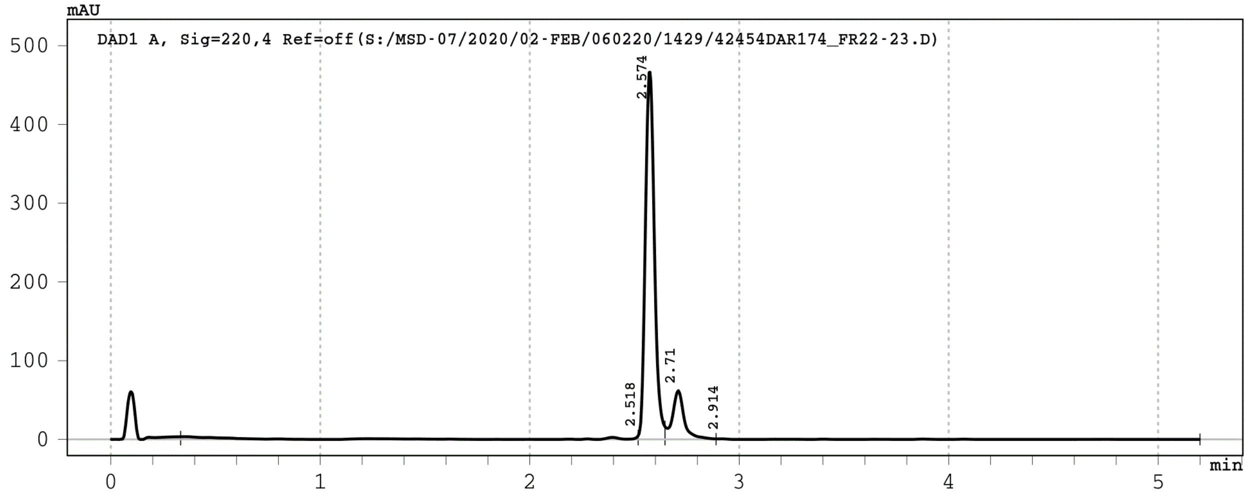


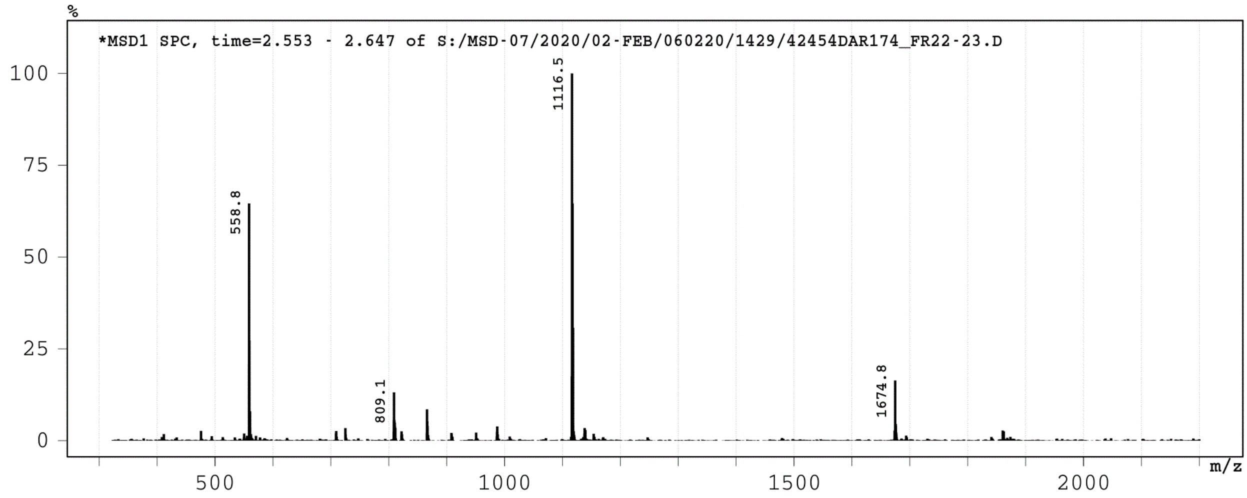


Signal 1: DAD1 A, Sig=220,4 Ref=off

| Peak # | RT [min] | Width [min] | Area | Area % |
|---|---|---|---|---|
| 1 | 2.518 | 2.18 | 83.7 | 4.8 |
| 2 | 2.574 | 0.13 | 1408.3 | 80.7 |
| 3 | 2.710 | 0.24 | 244.8 | 14.0 |
| 4 | 2.914 | 2.31 | 8.6 | 0.5 |

Data File : S:/MSD-07/2020/02-FEB/070220/1430/42454DAR175_FR5->

Injection Date : 07-Feb-20, 20:54:13
Sample Name : 42454DAR175_FR58-59
Remarks : Baseline corrected
Acq. Method : D:/METHODS/5-95-6M.M
column used : Je_559
Analysis Method : C:/CHEM32/1/METHODS/JPT-LCMS.M

Sample Info : 42454_175-Dr. Martine Darwish H-SVTVFVNNL-OH

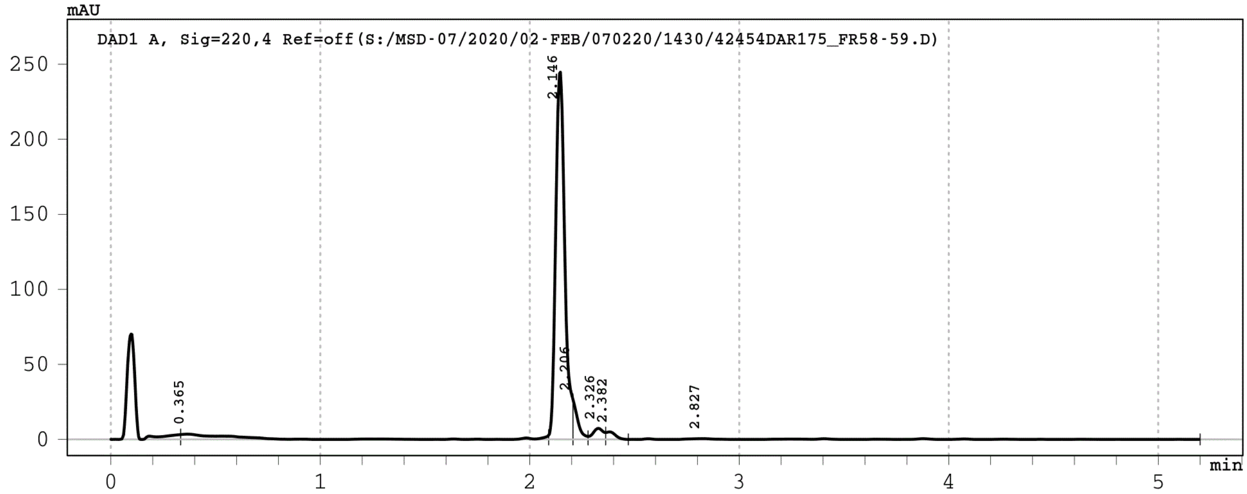


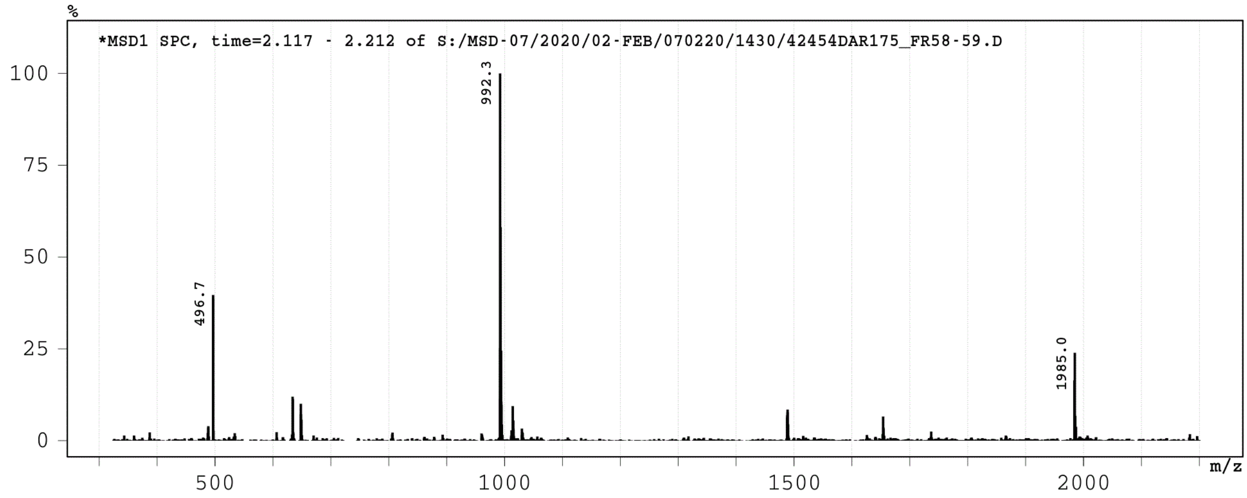


Signal 1: DAD1 A, Sig=220,4 Ref=off

| Peak # | RT [min] | Width [min] | Area | Area % |
|---|---|---|---|---|
| 1 | 0.365 | 1.76 | 64.5 | 7.2 |
| 2 | 2.146 | 0.12 | 739.8 | 82.1 |
| 3 | 2.206 | 0.07 | 42.0 | 4.7 |
| 4 | 2.326 | 0.08 | 25.7 | 2.9 |
| 5 | 2.382 | 0.11 | 15.3 | 1.7 |
| 6 | 2.827 | 2.73 | 13.7 | 1.5 |

Data File : S:/MSD-07/2020/02-FEB/060220/1429/42454DAR176_FR2->

Injection Date : 06-Feb-20, 20:28:02
Sample Name : 42454DAR176_FR26-27
Remarks : Baseline corrected
Acq. Method : D:/METHODS/5-95-6M.M
column used : Je_559
Analysis Method : C:/CHEM32/1/METHODS/JPT-LCMS.M

Sample Info : 42454_176-Dr. Martine Darwish H-KVLQKLCQL-OH

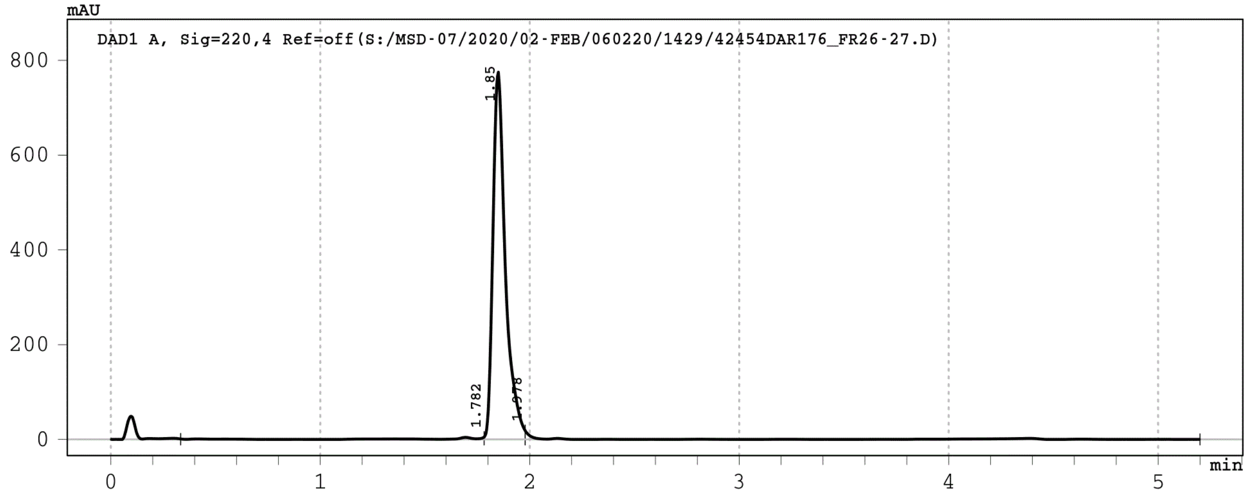


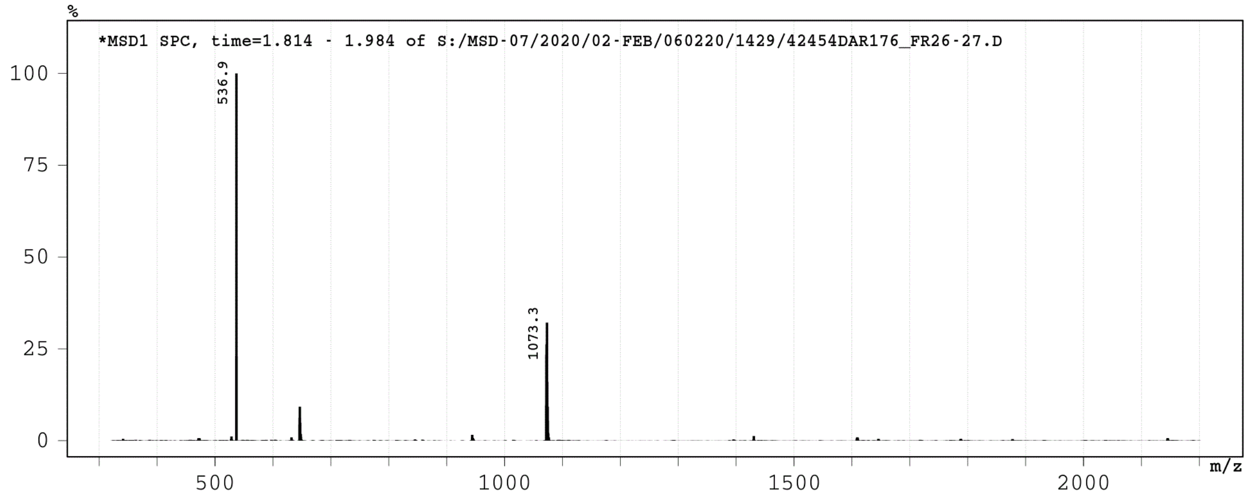


Signal 1: DAD1 A, Sig=220,4 Ref=off

| Peak # | RT [min] | Width [min] | Area | Area % |
|---|---|---|---|---|
| 1 | 1.782 | 1.45 | 55.5 | 1.7 |
| 2 | 1.850 | 0.20 | 3210.6 | 95.7 |
| 3 | 1.978 | 3.22 | 88.3 | 2.6 |

Data File : S:/MSD-07/2020/02-FEB/060220/1429/42454DAR177_FR3->

Injection Date : 06-Feb-20, 20:36:26
Sample Name : 42454DAR177_FR30-31
Remarks : Baseline corrected
Acq. Method : D:/METHODS/5-95-6M.M
column used : Je_559
Analysis Method : C:/CHEM32/1/METHODS/JPT-LCMS.M

Sample Info : 42454_177-Dr. Martine Darwish H-HSLTNSSSDFM-OH

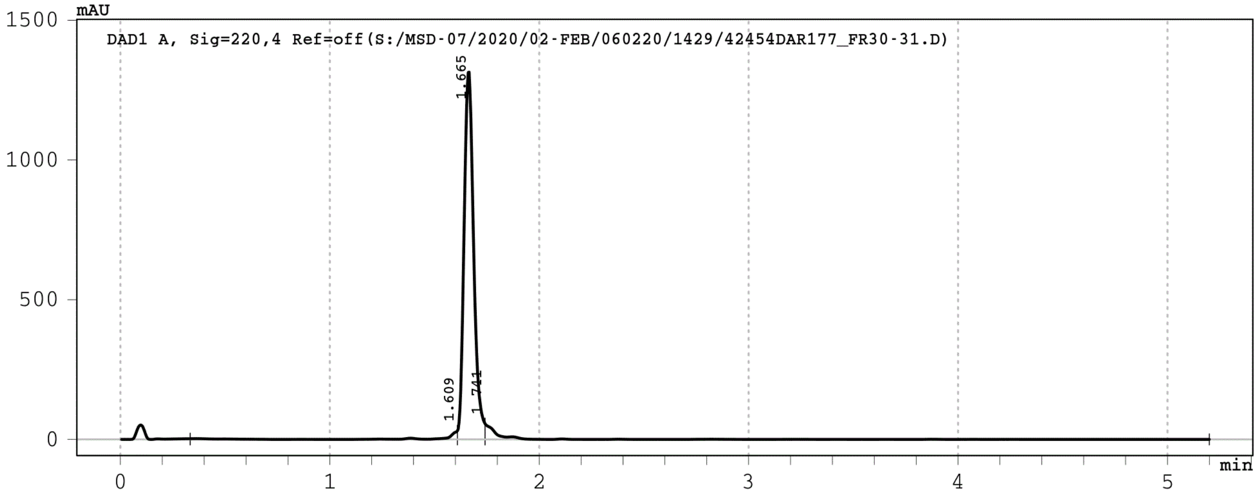


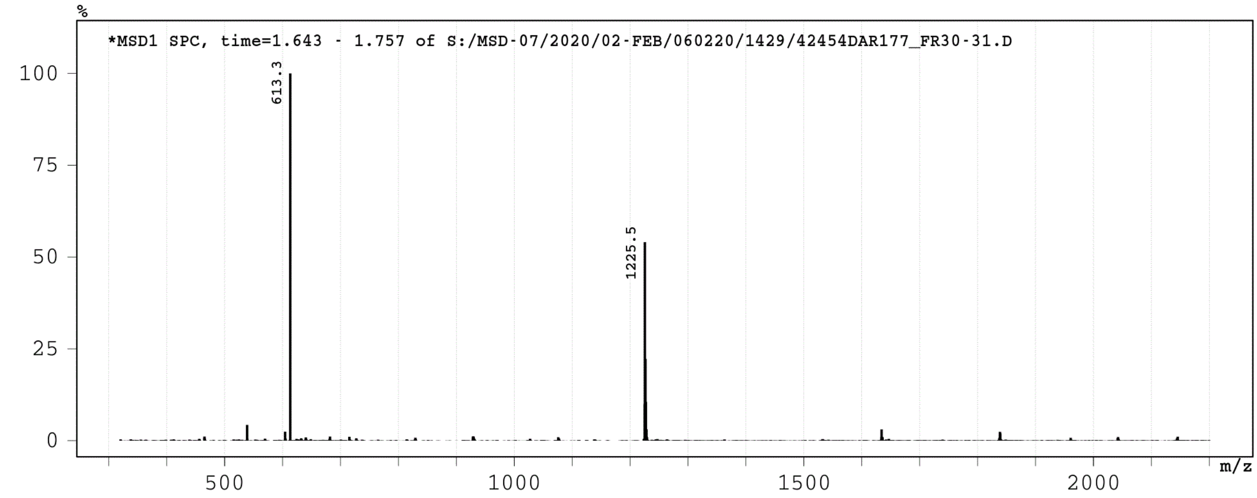


Signal 1: DAD1 A, Sig=220,4 Ref=off

| Peak # | RT [min] | Width [min] | Area | Area % |
|---|---|---|---|---|
| 1 | 1.609 | 1.28 | 120.5 | 2.6 |
| 2 | 1.665 | 0.13 | 4212.4 | 92.6 |
| 3 | 1.741 | 3.46 | 217.0 | 4.8 |

Data File : S:/MSD-07/2020/02-FEB/070220/1430/42454DAR178_FR6->

Injection Date : 07-Feb-20, 21:02:36
Sample Name : 42454DAR178_FR65-67
Remarks : Baseline corrected
Acq. Method : D:/METHODS/5-95-6M.M
column used : Je_559
Analysis Method : C:/CHEM32/1/METHODS/JPT-LCMS.M

Sample Info : 42454_178-Dr. Martine Darwish H-AQHHRWQLL-OH

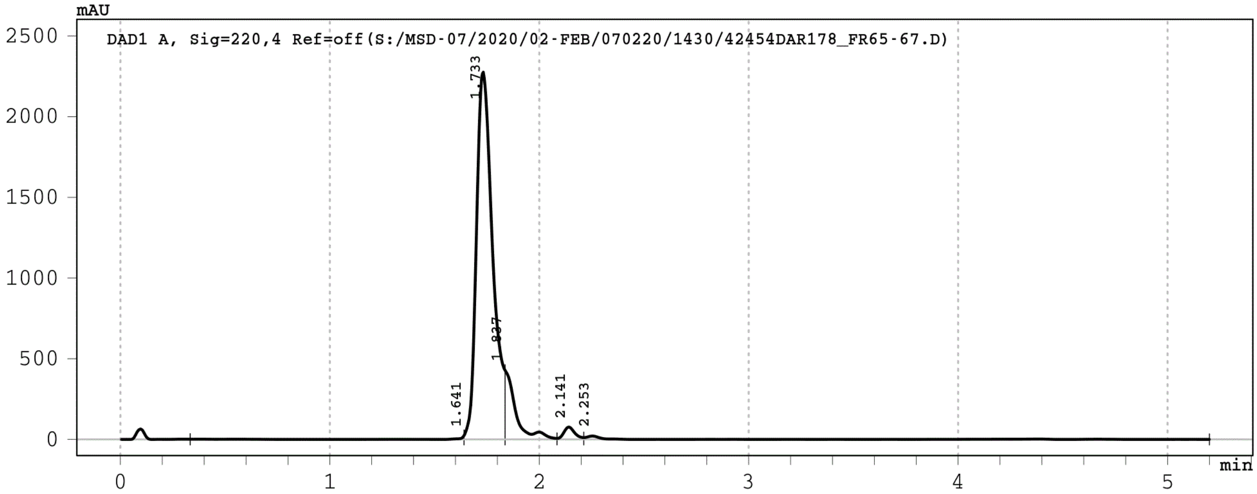


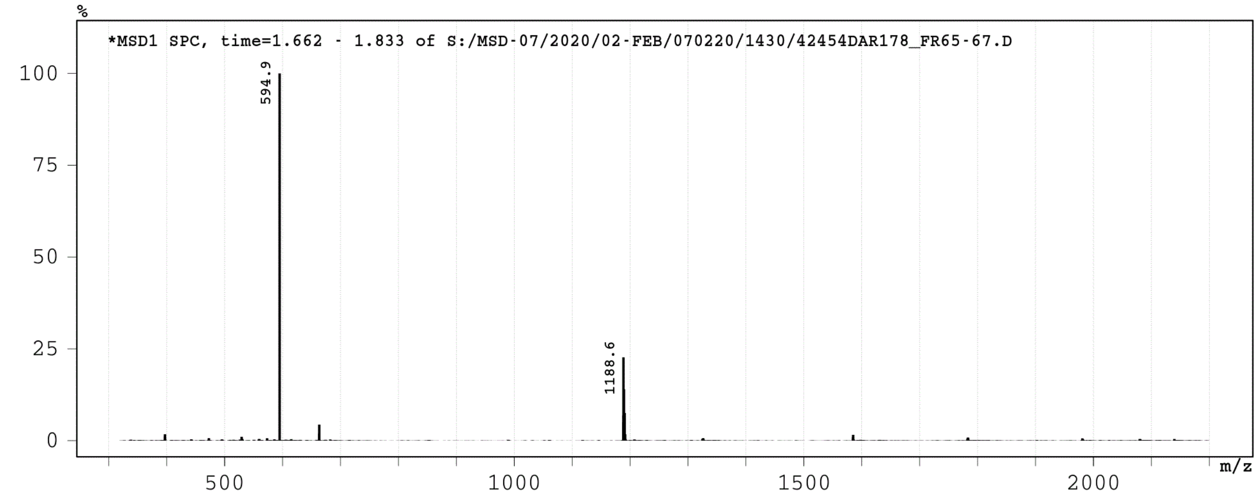


Signal 1: DAD1 A, Sig=220,4 Ref=off

| Peak # | RT [min] | Width [min] | Area | Area % |
|---|---|---|---|---|
| 1 | 1.641 | 1.31 | 50.7 | 0.4 |
| 2 | 1.733 | 0.20 | 12301.3 | 86.0 |
| 3 | 1.837 | 0.25 | 1506.2 | 10.5 |
| 4 | 2.141 | 0.13 | 284.7 | 2.0 |
| 5 | 2.253 | 2.99 | 152.8 | 1.1 |

Data File : S:/MSD-07/2020/02-FEB/060220/1429/42454DAR179_FR3->

Injection Date : 06-Feb-20, 20:44:48
Sample Name : 42454DAR179_FR37-39
Remarks : Baseline corrected
Acq. Method : D:/METHODS/5-95-6M.M
column used : Je_559
Analysis Method : C:/CHEM32/1/METHODS/JPT-LCMS.M

Sample Info : 42454_179-Dr. Martine Darwish H-MQQLNPEFV-OH

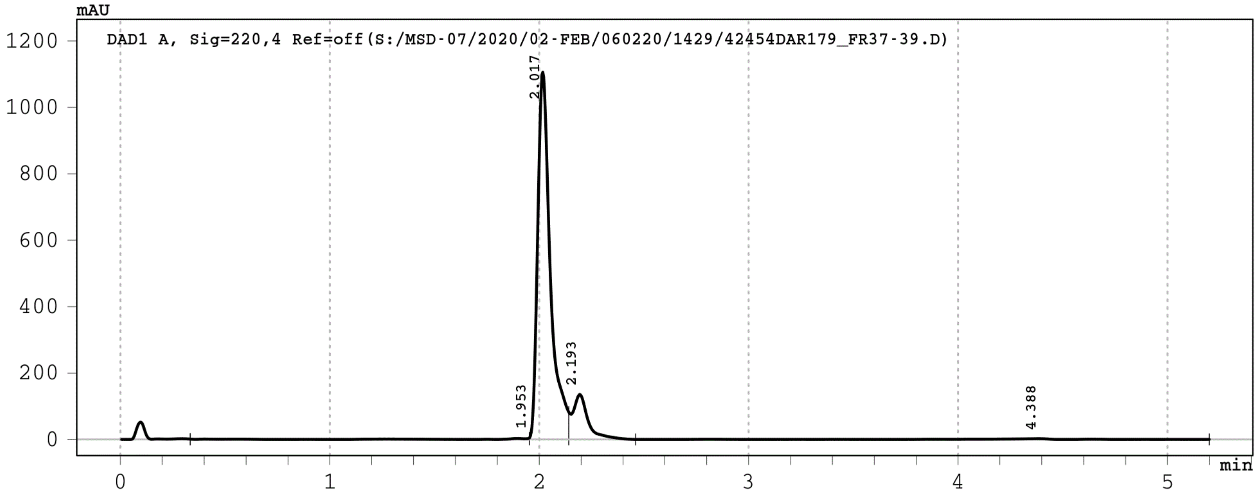


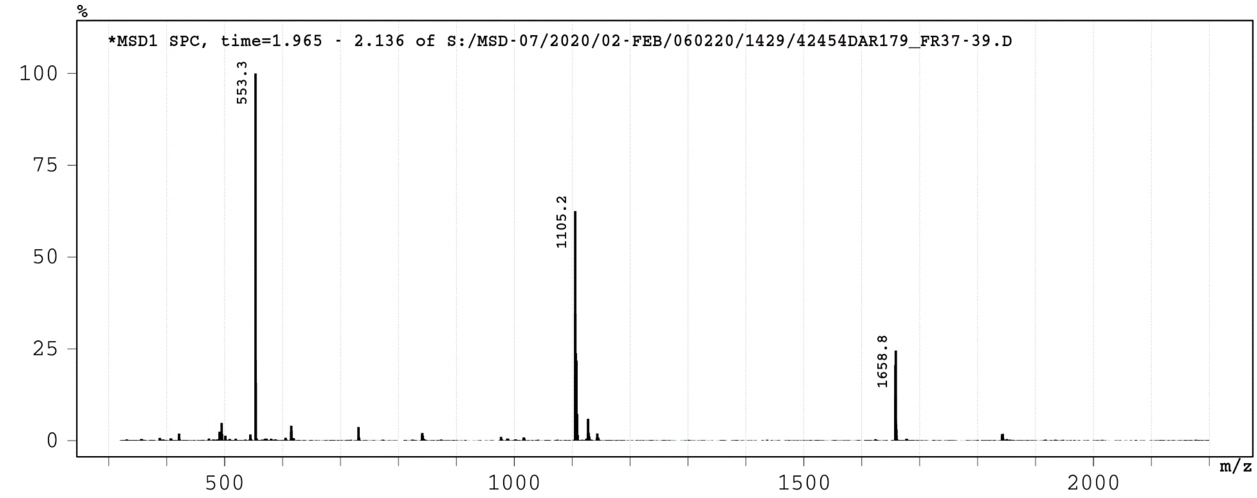


Signal 1: DAD1 A, Sig=220,4 Ref=off

| Peak # | RT [min] | Width [min] | Area | Area % |
|---|---|---|---|---|
| 1 | 1.953 | 1.62 | 45.7 | 0.8 |
| 2 | 2.017 | 0.19 | 4683.9 | 85.6 |
| 3 | 2.193 | 0.32 | 700.8 | 12.8 |
| 4 | 4.388 | 2.74 | 44.1 | 0.8 |

Data File : S:/MSD-07/2020/02-FEB/070220/1430/42454DAR180_FR8->

Injection Date : 07-Feb-20, 21:10:59
Sample Name : 42454DAR180_FR83-84
Remarks : Baseline corrected
Acq. Method : D:/METHODS/5-95-6M.M
column used : Je_559
Analysis Method : C:/CHEM32/1/METHODS/JPT-LCMS.M

Sample Info : 42454_180-Dr. Martine Darwish H-LSASRYALL-OH

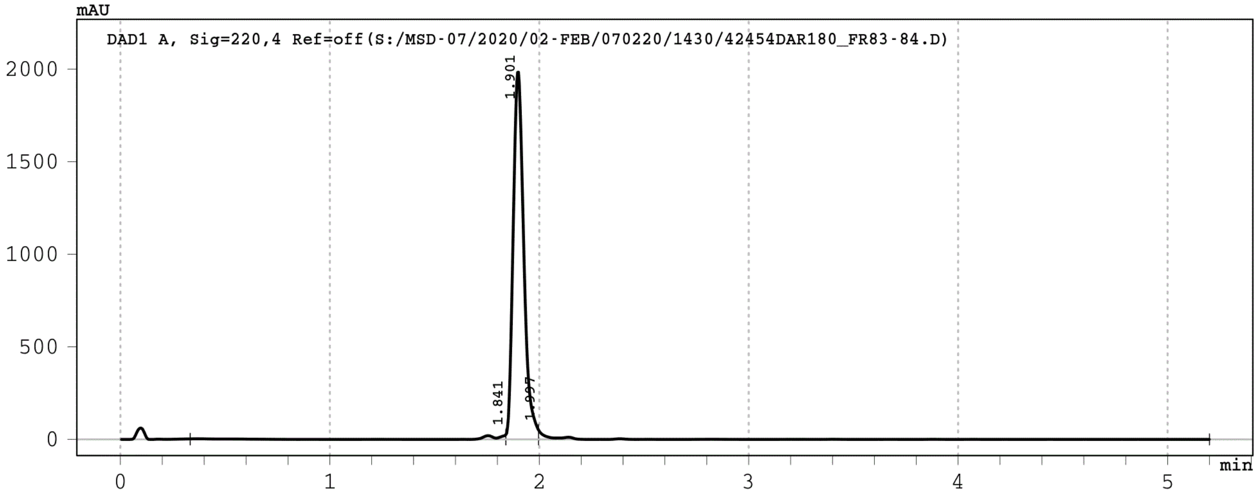


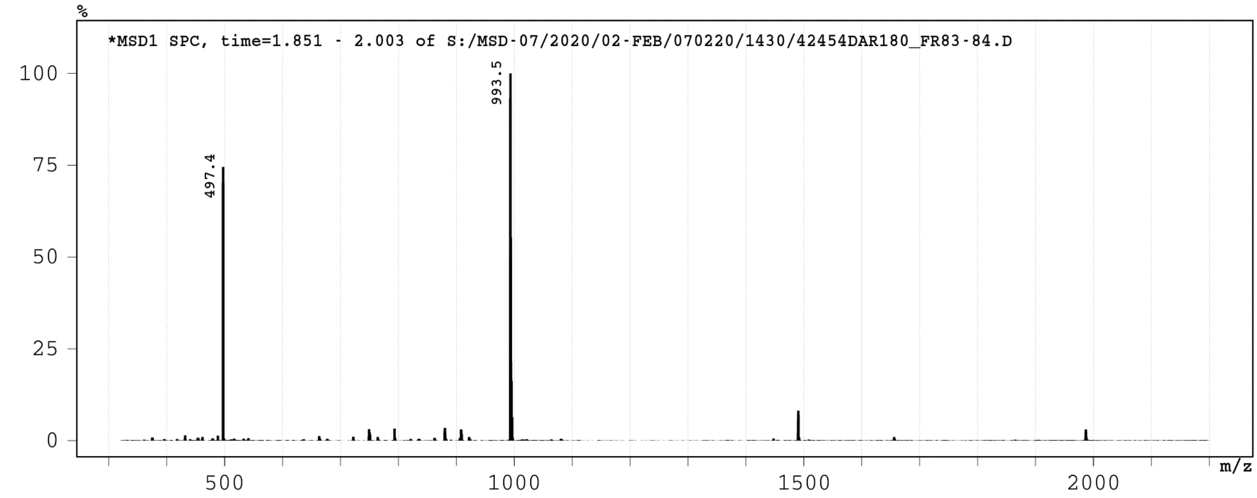


Signal 1: DAD1 A, Sig=220,4 Ref=off

| Peak # | RT [min] | Width [min] | Area | Area % |
|---|---|---|---|---|
| 1 | 1.841 | 1.51 | 161.2 | 2.2 |
| 2 | 1.901 | 0.16 | 7014.3 | 95.5 |
| 3 | 1.997 | 3.20 | 168.2 | 2.3 |

Data File : S:/MSD-07/2020/02-FEB/110220/1433/42454DAR181_FR7->

Injection Date : 12-Feb-20, 01:43:48
Sample Name : 42454DAR181_FR79-81
Remarks : Baseline corrected
Acq. Method : D:/METHODS/5-95-6M.M
column used : Je_559
Analysis Method : C:/CHEM32/1/METHODS/JPT-LCMS.M

Sample Info : 42454_181-Dr. Martine Darwish H-SSIKVVGL-OH

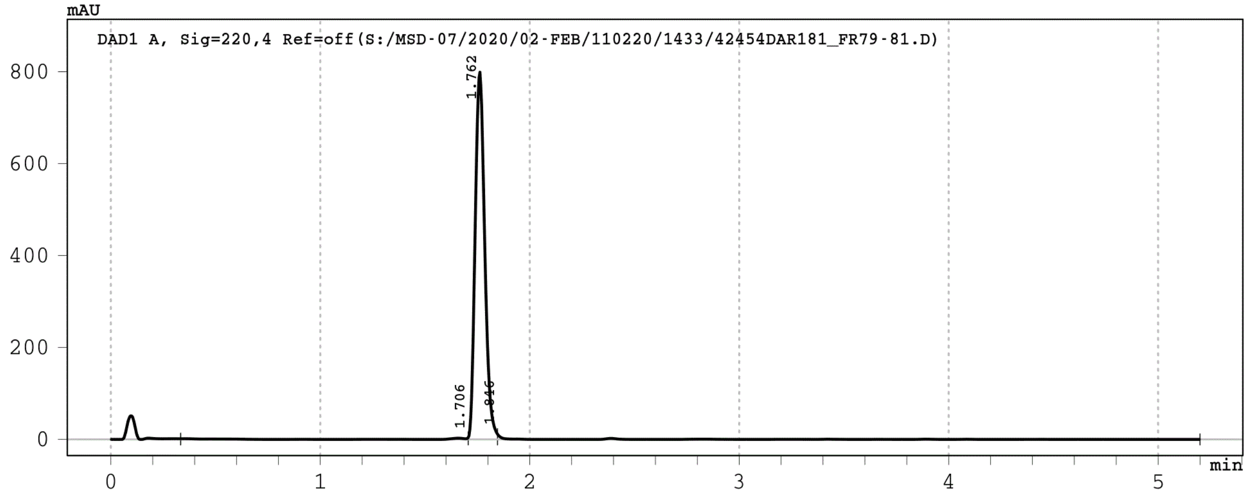


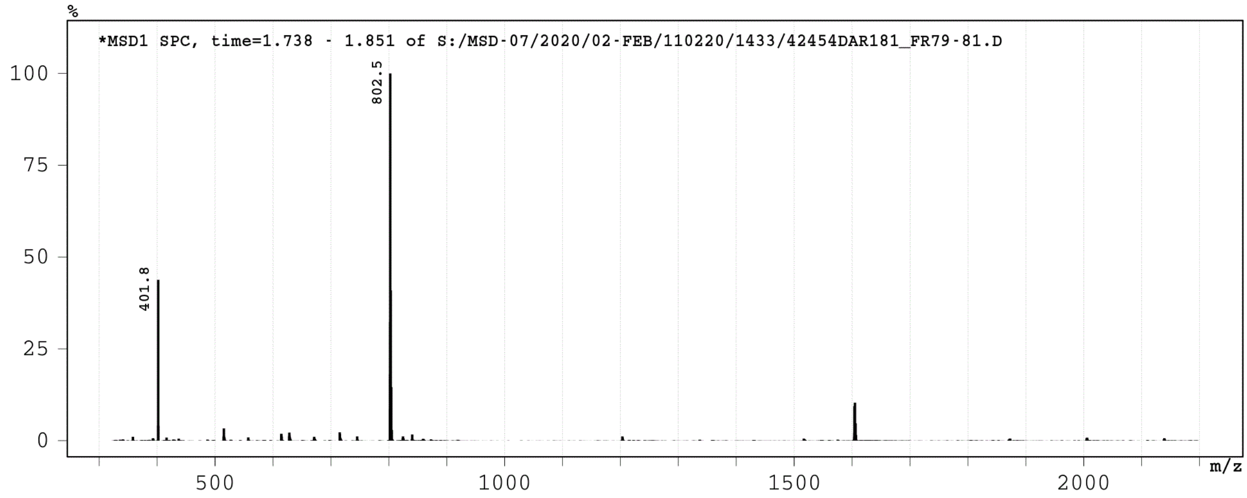


Signal 1: DAD1 A, Sig=220,4 Ref=off

| Peak # | RT [min] | Width [min] | Area | Area % |
|---|---|---|---|---|
| 1 | 1.706 | 1.37 | 32.6 | 1.3 |
| 2 | 1.762 | 0.14 | 2538.2 | 97.4 |
| 3 | 1.846 | 3.35 | 34.5 | 1.3 |

Data File : S:/MSD-07/2020/02-FEB/110220/1433/42454DAR182_FR6->

Injection Date : 12-Feb-20, 01:52:15
Sample Name : 42454DAR182_FR6-7
Remarks : Baseline corrected
Acq. Method : D:/METHODS/5-95-6M.M
column used : Je_559
Analysis Method : C:/CHEM32/1/METHODS/JPT-LCMS.M

Sample Info : 42454_182-Dr. Martine Darwish H-TGSVFGEL-OH

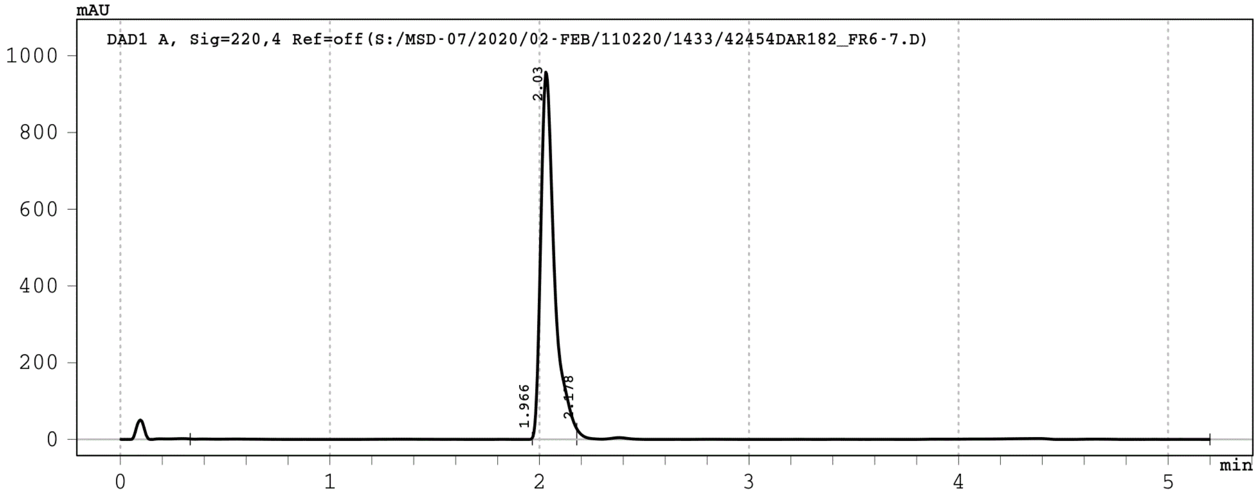


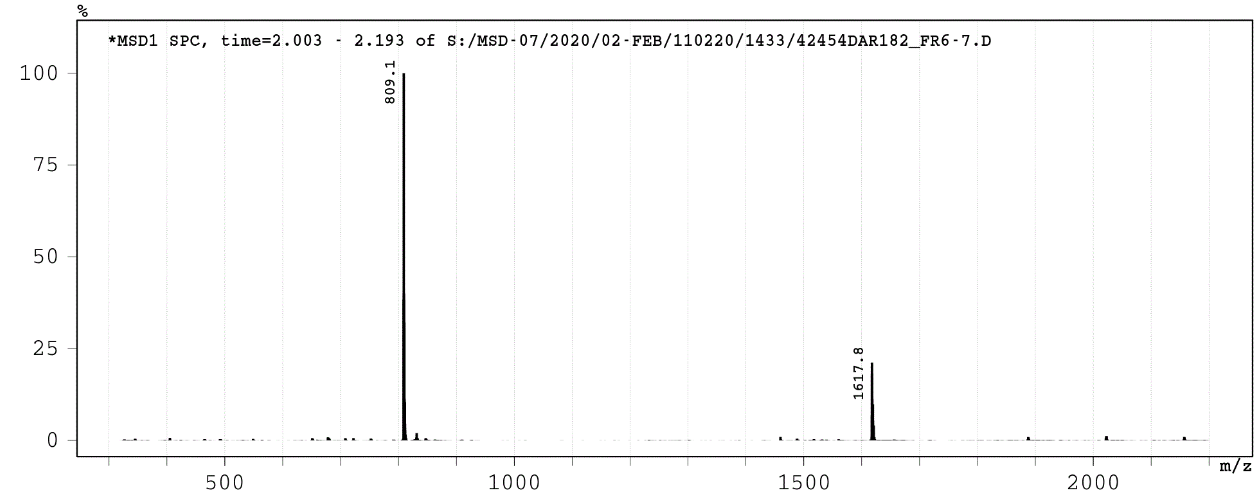


Signal 1: DAD1 A, Sig=220,4 Ref=off

| Peak # | RT [min] | Width [min] | Area | Area % |
|---|---|---|---|---|
| 1 | 1.966 | 1.63 | 26.5 | 0.6 |
| 2 | 2.030 | 0.21 | 4355.1 | 96.6 |
| 3 | 2.178 | 3.02 | 127.1 | 2.8 |

Data File : S:/MSD-07/2020/02-FEB/110220/1433/42454DAR183_FR1->

Injection Date : 12-Feb-20, 02:00:39
Sample Name : 42454DAR183_FR12
Remarks : Baseline corrected
Acq. Method : D:/METHODS/5-95-6M.M
column used : Je_559
Analysis Method : C:/CHEM32/1/METHODS/JPT-LCMS.M

Sample Info : 42454_183-Dr. Martine Darwish H-LGVLFSQL-OH

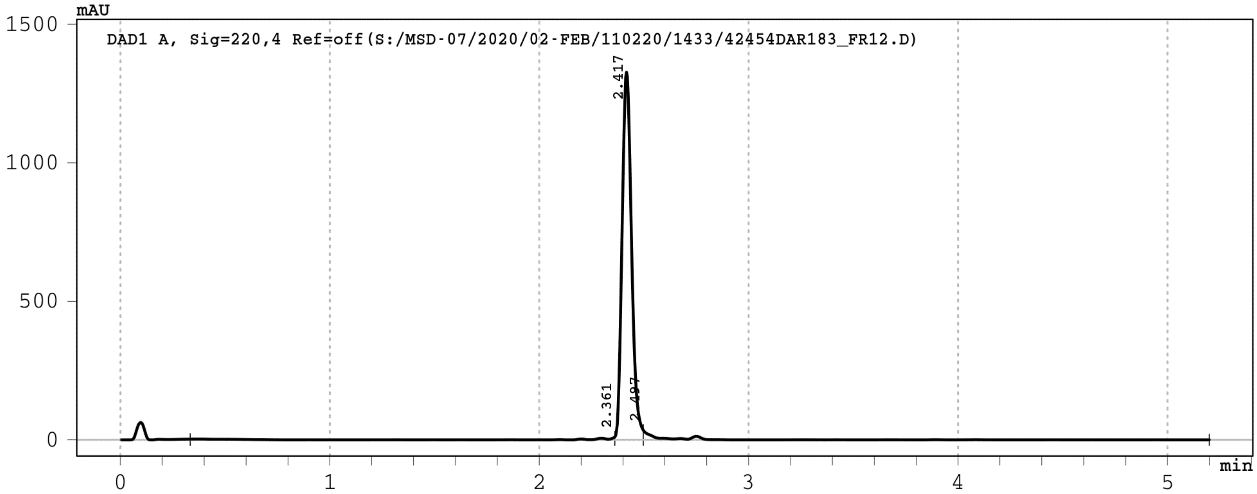


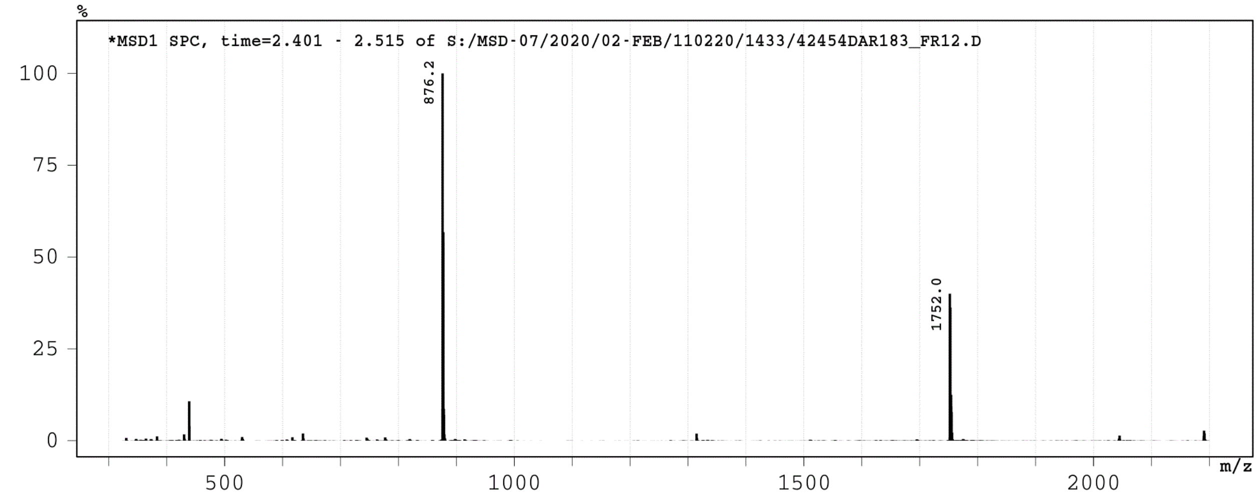


Signal 1: DAD1 A, Sig=220,4 Ref=off

| Peak # | RT [min] | Width [min] | Area | Area % |
|---|---|---|---|---|
| 1 | 2.361 | 2.03 | 81.7 | 1.8 |
| 2 | 2.417 | 0.14 | 4179.3 | 94.6 |
| 3 | 2.497 | 2.70 | 158.9 | 3.6 |

Data File : S:/MSD-07/2020/02-FEB/060220/1429/42454DAR184_FR4->

Injection Date : 06-Feb-20, 20:53:12
Sample Name : 42454DAR184_FR47-48
Remarks : Baseline corrected
Acq. Method : D:/METHODS/5-95-6M.M
column used : Je_559
Analysis Method : C:/CHEM32/1/METHODS/JPT-LCMS.M

Sample Info : 42454_184-Dr. Martine Darwish H-LFFPCSNPL-OH

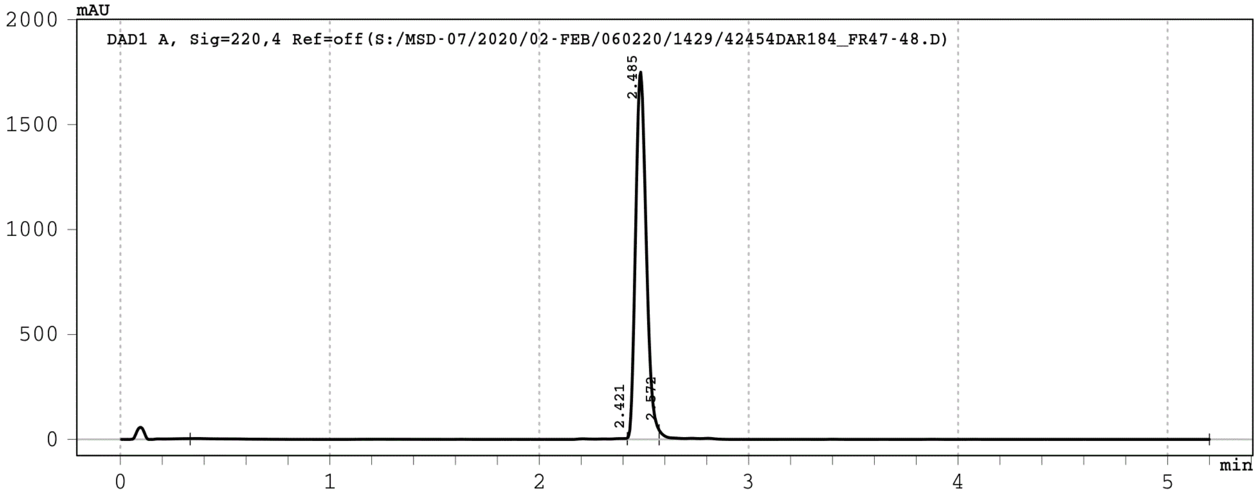


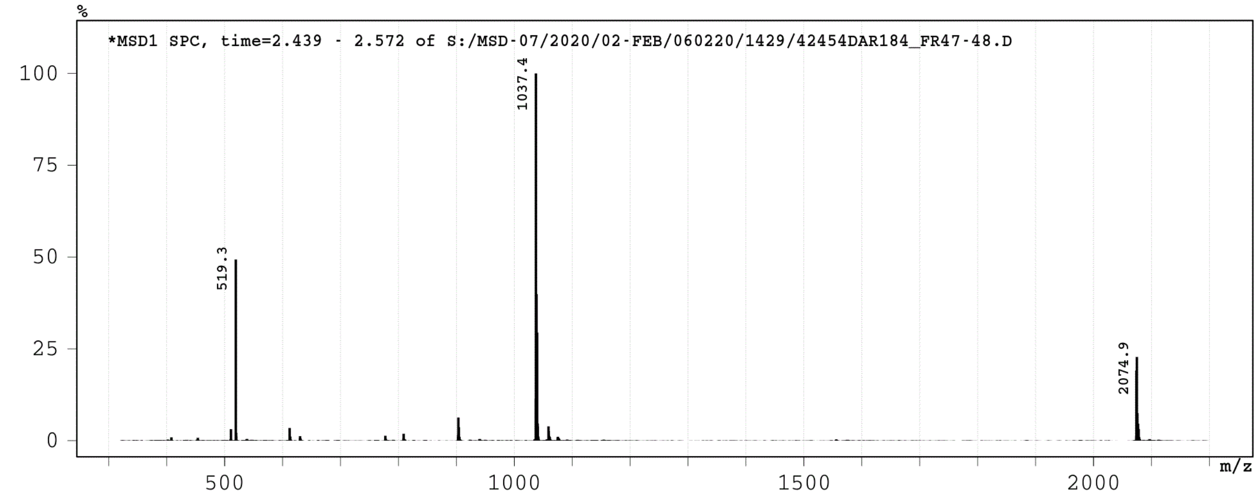


Signal 1: DAD1 A, Sig=220,4 Ref=off

| Peak # | RT [min] | Width [min] | Area | Area % |
|---|---|---|---|---|
| 1 | 2.421 | 2.09 | 107.4 | 1.7 |
| 2 | 2.485 | 0.15 | 6197.6 | 96.2 |
| 3 | 2.572 | 2.63 | 135.1 | 2.1 |

Data File : S:/MSD-07/2020/02-FEB/070220/1430/42454DAR185_FR7->

Injection Date : 07-Feb-20, 21:19:24
Sample Name : 42454DAR185_FR7-9
Remarks : Baseline corrected
Acq. Method : D:/METHODS/5-95-6M.M
column used : Je_559
Analysis Method : C:/CHEM32/1/METHODS/JPT-LCMS.M

Sample Info : 42454_185-Dr. Martine Darwish H-SNLNRNATV-OH

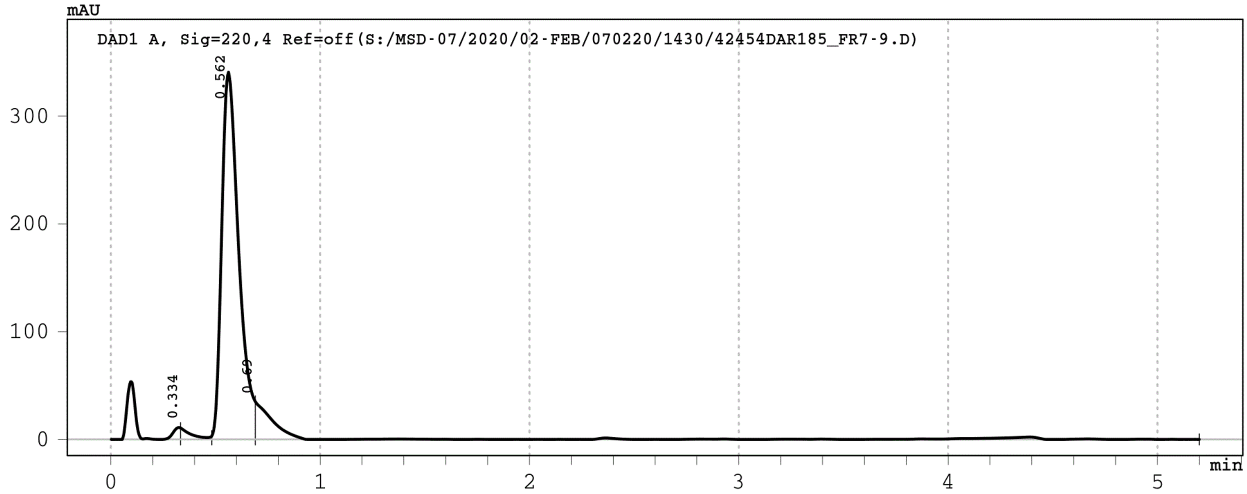


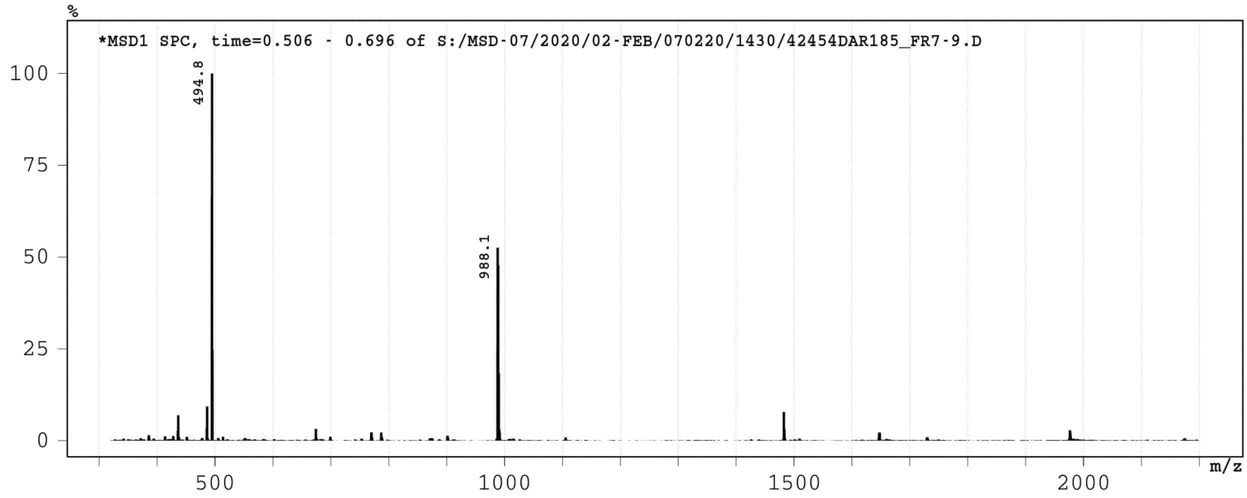


Signal 1: DAD1 A, Sig=220,4 Ref=off

| Peak # | RT [min] | Width [min] | Area | Area % |
|---|---|---|---|---|
| 1 | 0.334 | 0.15 | 37.0 | 1.7 |
| 2 | 0.562 | 0.21 | 1922.6 | 86.7 |
| 3 | 0.690 | 4.51 | 257.0 | 11.6 |

Data File : S:/MSD-07/2020/02-FEB/140220/1435/42454DAR186_FR7->

Injection Date : 15-Feb-20, 00:51:56
Sample Name : 42454DAR186_FR7-8
Remarks : Baseline corrected
Acq. Method : D:/METHODS/5-95-6M.M
column used : Je_559
Analysis Method : C:/CHEM32/1/METHODS/JPT-LCMS.M

Sample Info : 42454_186-Dr. Martine Darwish H-LGSIFSTL-OH

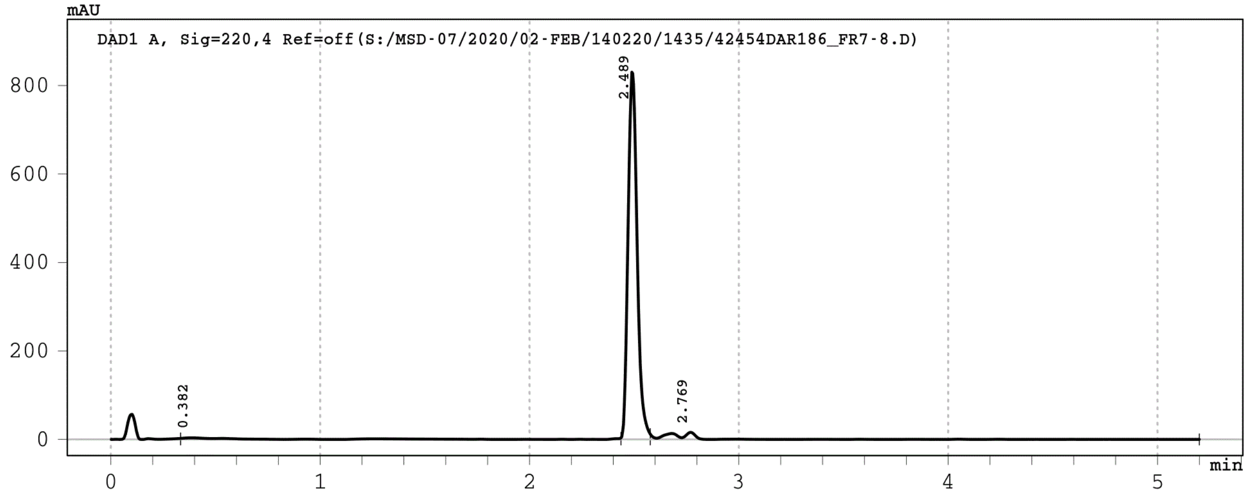


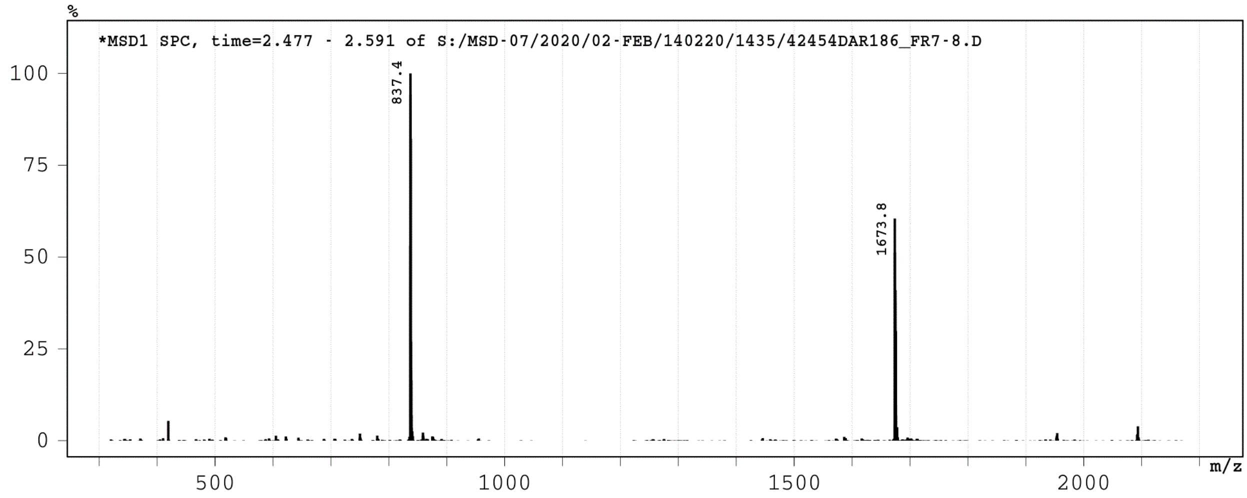


Signal 1: DAD1 A, Sig=220,4 Ref=off

| Peak # | RT [min] | Width [min] | Area | Area % |
|---|---|---|---|---|
| 1 | 0.382 | 2.10 | 88.3 | 3.2 |
| 2 | 2.489 | 0.14 | 2521.6 | 91.9 |
| 3 | 2.769 | 2.62 | 132.9 | 4.8 |

Data File : S:/MSD-07/2020/02-FEB/070220/1430/42454DAR187_FR1->

Injection Date : 07-Feb-20, 21:36:11
Sample Name : 42454DAR187_FR13-15
Remarks : Baseline corrected
Acq. Method : D:/METHODS/5-95-6M.M
column used : Je_559
Analysis Method : C:/CHEM32/1/METHODS/JPT-LCMS.M

Sample Info : 42454_187-Dr. Martine Darwish H-HSFVYSVGF-OH

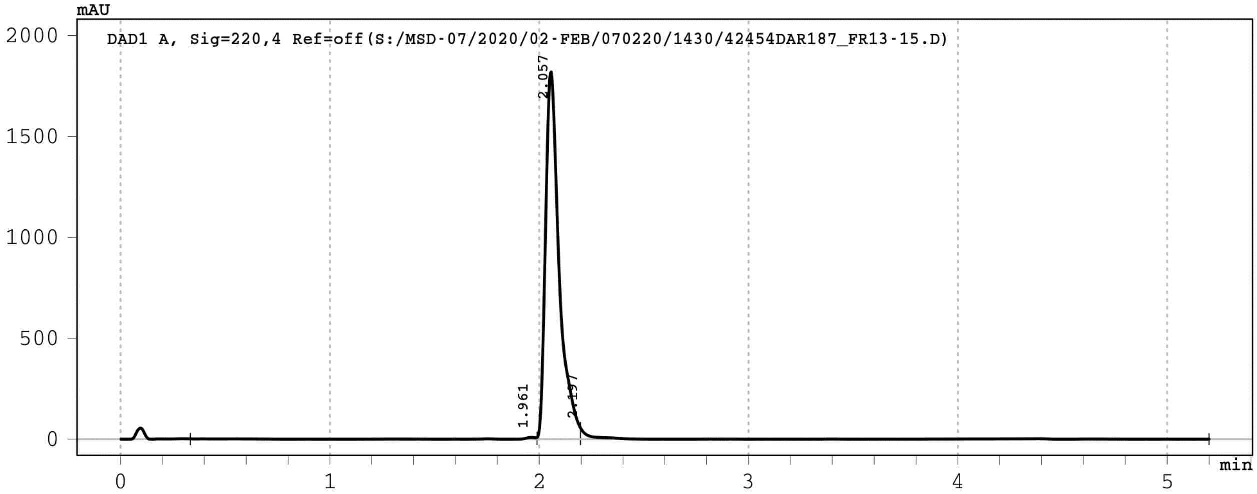


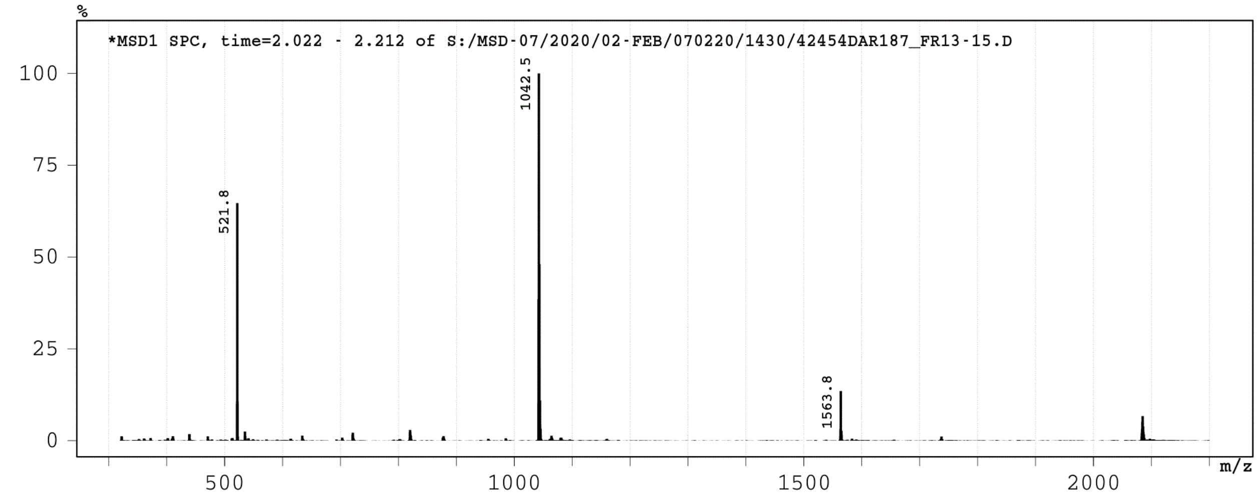


Signal 1: DAD1 A, Sig=220,4 Ref=off

| Peak # | RT [min] | Width [min] | Area | Area % |
|---|---|---|---|---|
| 1 | 1.961 | 1.66 | 68.9 | 0.8 |
| 2 | 2.057 | 0.21 | 8269.3 | 96.7 |
| 3 | 2.197 | 3.00 | 216.5 | 2.5 |

Data File : S:/MSD-07/2020/02-FEB/060220/1429/42454DAR188_FR5->

Injection Date : 06-Feb-20, 21:01:34
Sample Name : 42454DAR188_FR52-53
Remarks : Baseline corrected
Acq. Method : D:/METHODS/5-95-6M.M
column used : Je_559
Analysis Method : C:/CHEM32/1/METHODS/JPT-LCMS.M

Sample Info : 42454_188-Dr. Martine Darwish H-PNAGFMSQL-OH

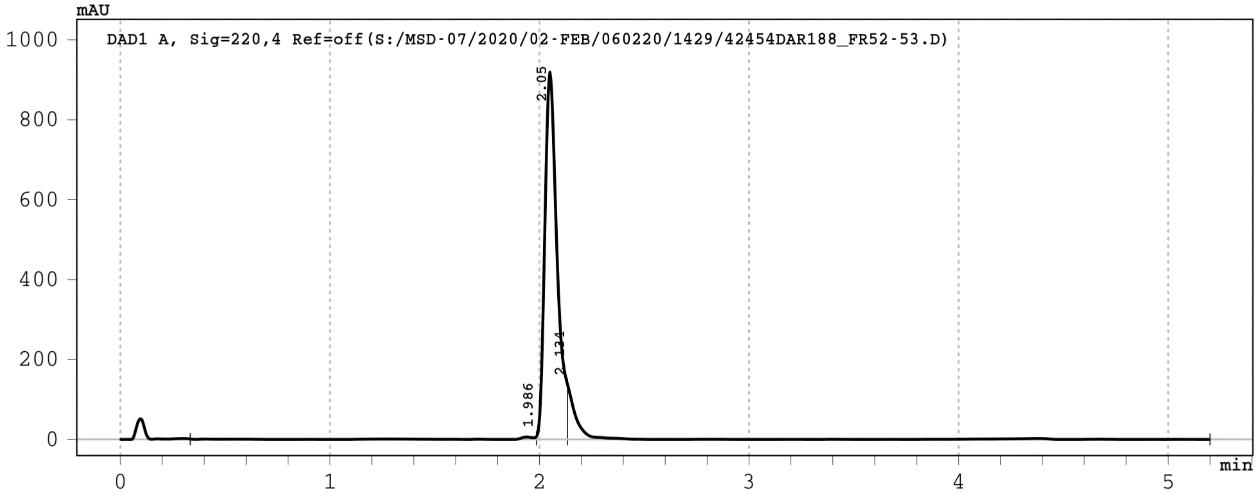


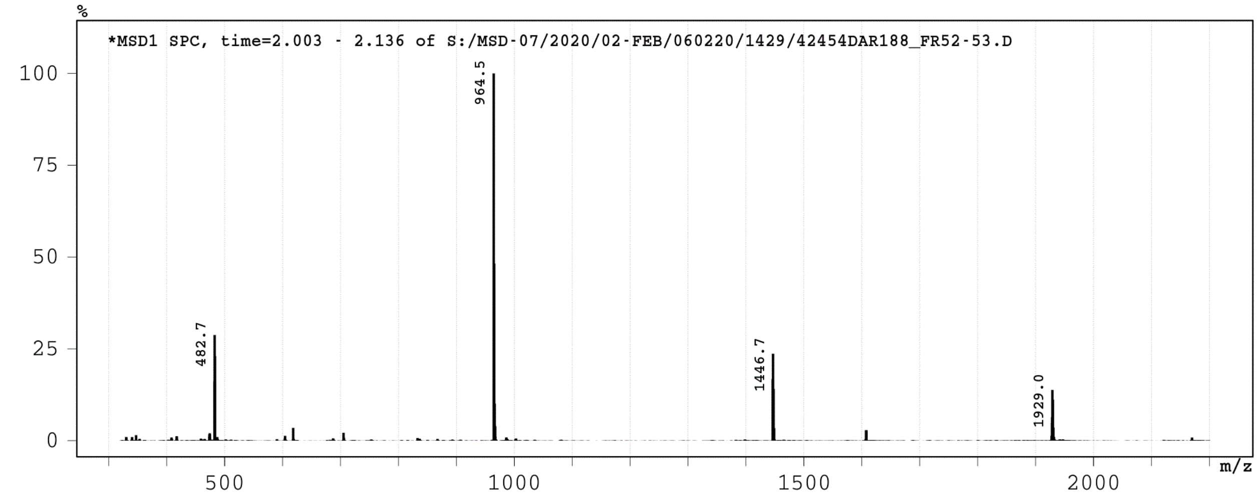


Signal 1: DAD1 A, Sig=220,4 Ref=off

| Peak # | RT [min] | Width [min] | Area | Area % |
|---|---|---|---|---|
| 1 | 1.986 | 1.65 | 56.2 | 1.3 |
| 2 | 2.050 | 0.15 | 3734.8 | 88.6 |
| 3 | 2.134 | 3.07 | 423.1 | 10.0 |

Data File : S:/MSD-07/2020/02-FEB/060220/1429/42454DAR189_FR6->

Injection Date : 06-Feb-20, 21:09:56
Sample Name : 42454DAR189_FR60-61
Remarks : Baseline corrected
Acq. Method : D:/METHODS/5-95-6M.M
column used : Je_559
Analysis Method : C:/CHEM32/1/METHODS/JPT-LCMS.M

Sample Info : 42454_189-Dr. Martine Darwish H-SWGGLCSNL-OH

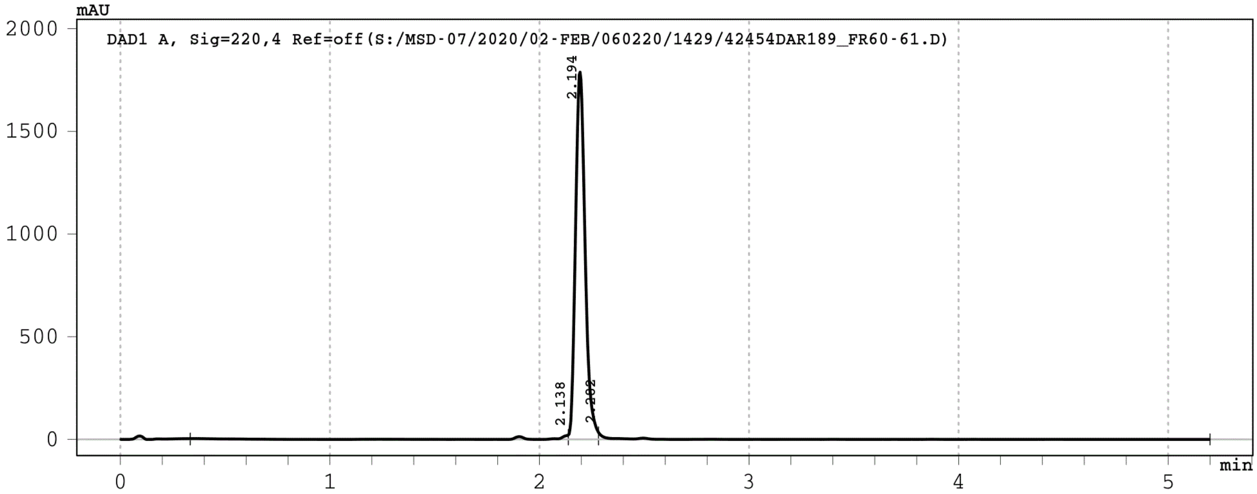


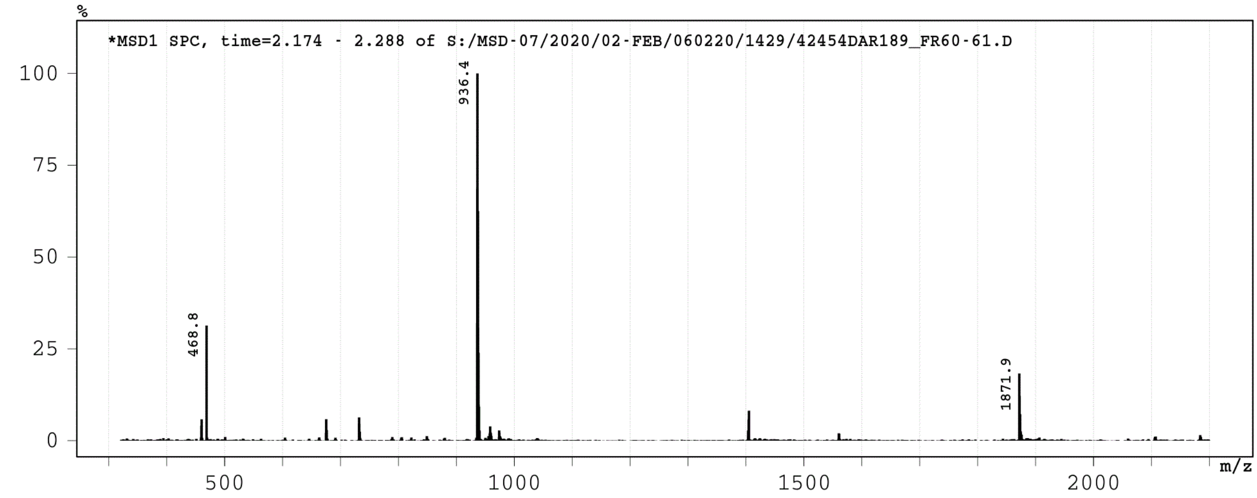


Signal 1: DAD1 A, Sig=220,4 Ref=off

| Peak # | RT [min] | Width [min] | Area | Area % |
|---|---|---|---|---|
| 1 | 2.138 | 1.80 | 146.8 | 2.5 |
| 2 | 2.194 | 0.14 | 5719.3 | 95.8 |
| 3 | 2.282 | 2.92 | 101.6 | 1.7 |

Data File : S:/MSD-07/2020/02-FEB/110220/1433/42454DAR190_FR2->

Injection Date : 12-Feb-20, 02:09:04
Sample Name : 42454DAR190_FR22-24
Remarks : Baseline corrected
Acq. Method : D:/METHODS/5-95-6M.M
column used : Je_559
Analysis Method : C:/CHEM32/1/METHODS/JPT-LCMS.M

Sample Info : 42454_190-Dr. Martine Darwish H-IHPVMSTL-OH

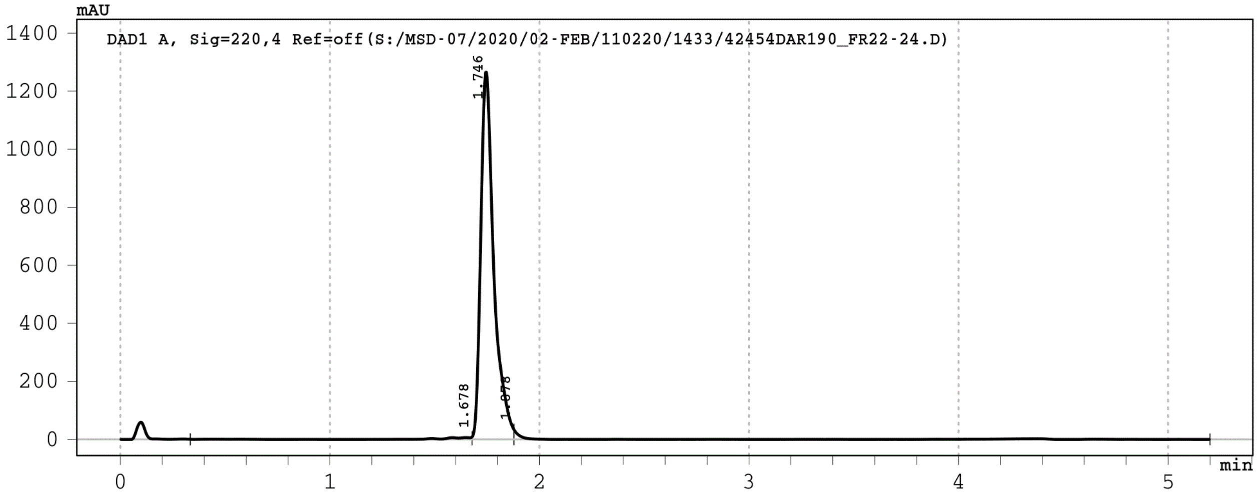


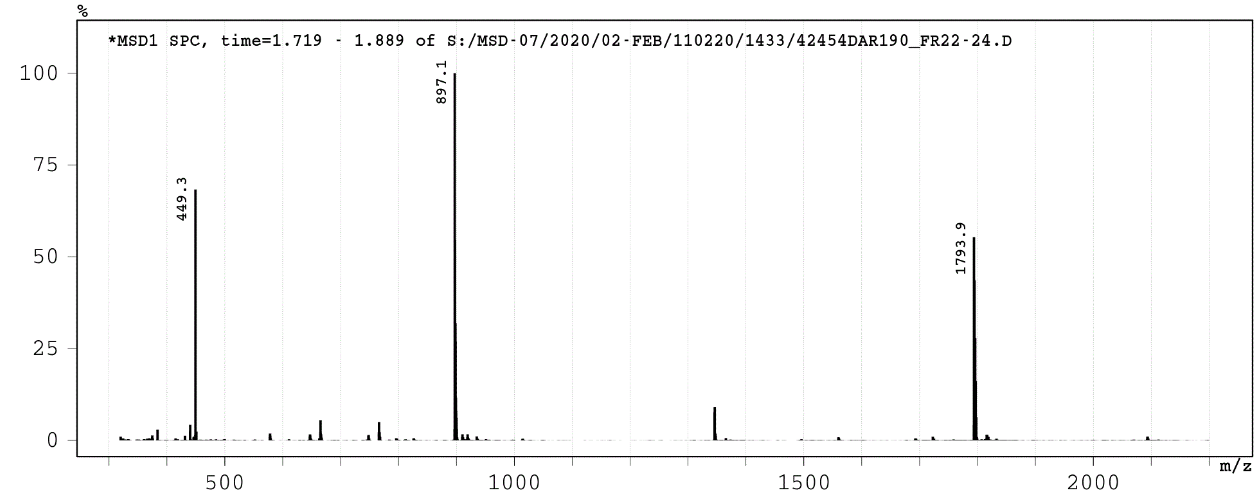


Signal 1: DAD1 A, Sig=220,4 Ref=off

| Peak # | RT [min] | Width [min] | Area | Area % |
|---|---|---|---|---|
| 1 | 1.678 | 1.34 | 69.5 | 1.2 |
| 2 | 1.746 | 0.20 | 5490.0 | 96.7 |
| 3 | 1.878 | 3.32 | 115.1 | 2.0 |

Data File : S:/MSD-07/2020/02-FEB/100220/1432/42454DAR191_FR6->

Injection Date : 10-Feb-20, 18:17:13
Sample Name : 42454DAR191_FR6
Remarks : Baseline corrected
Acq. Method : D:/METHODS/5-95-6M.M
column used : Je_559
Analysis Method : C:/CHEM32/1/METHODS/JPT-LCMS.M

Sample Info : 42454_191-Dr. Martine Darwish H-RAWKNHKAYI-OH

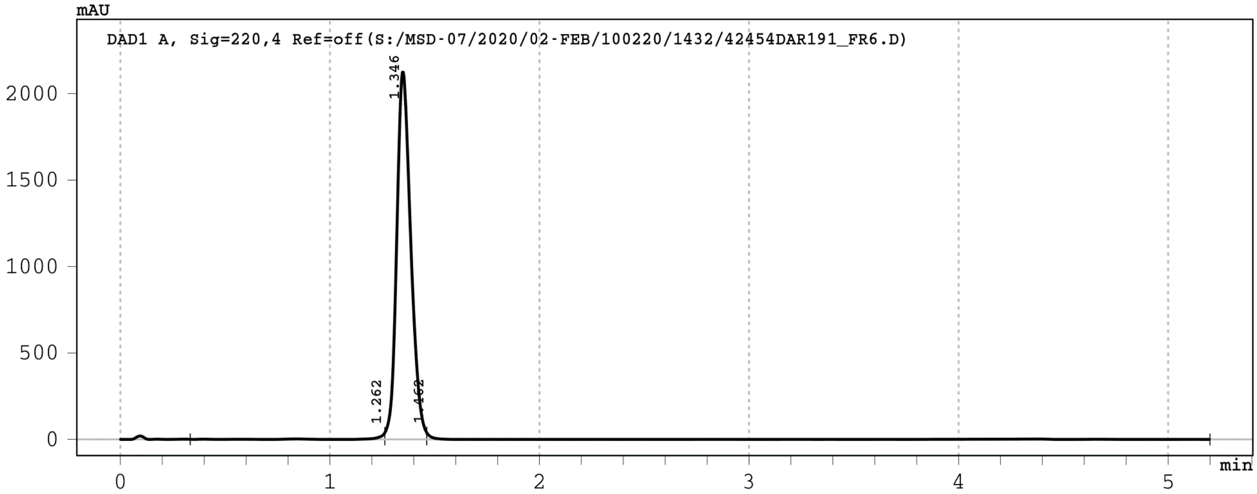


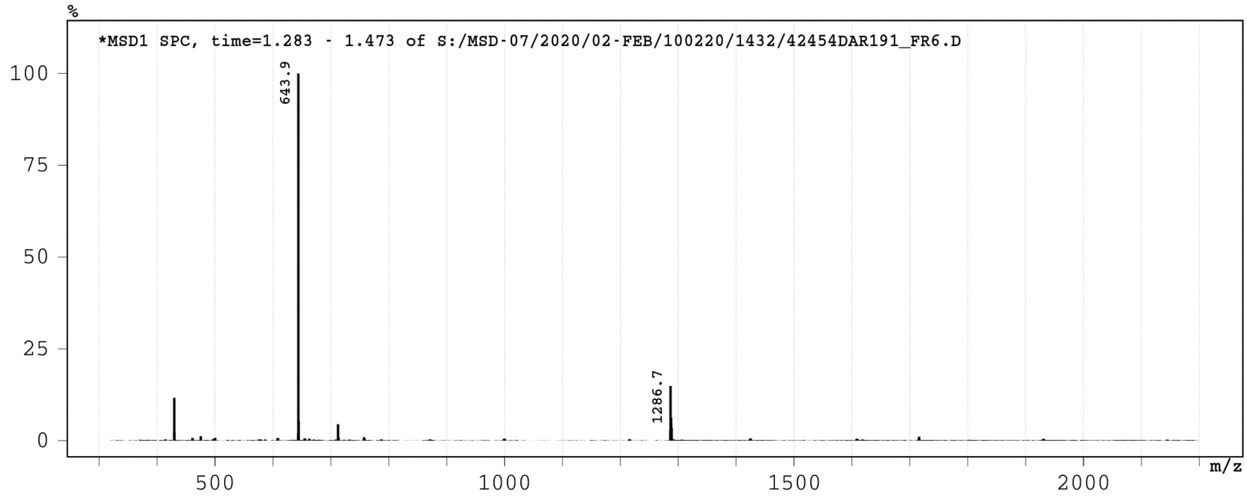


Signal 1: DAD1 A, Sig=220,4 Ref=off

| Peak # | RT [min] | Width [min] | Area | Area % |
|---|---|---|---|---|
| 1 | 1.262 | 0.93 | 92.2 | 0.9 |
| 2 | 1.346 | 0.20 | 9939.1 | 98.0 |
| 3 | 1.462 | 3.74 | 109.8 | 1.1 |

Data File : S:/MSD-07/2020/02-FEB/100220/1432/42454DAR192_FR2->

Injection Date : 10-Feb-20, 18:25:34
Sample Name : 42454DAR192_FR27-28
Remarks : Baseline corrected
Acq. Method : D:/METHODS/5-95-6M.M
column used : Je_559
Analysis Method : C:/CHEM32/1/METHODS/JPT-LCMS.M

Sample Info : 42454_192-Dr. Martine Darwish H-SAVDFKLHI-OH

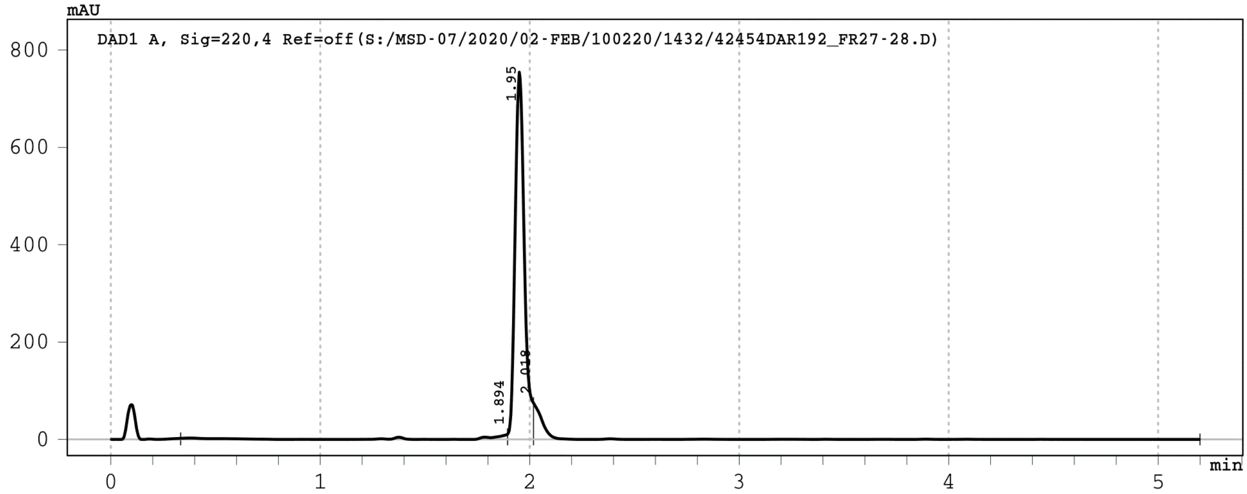


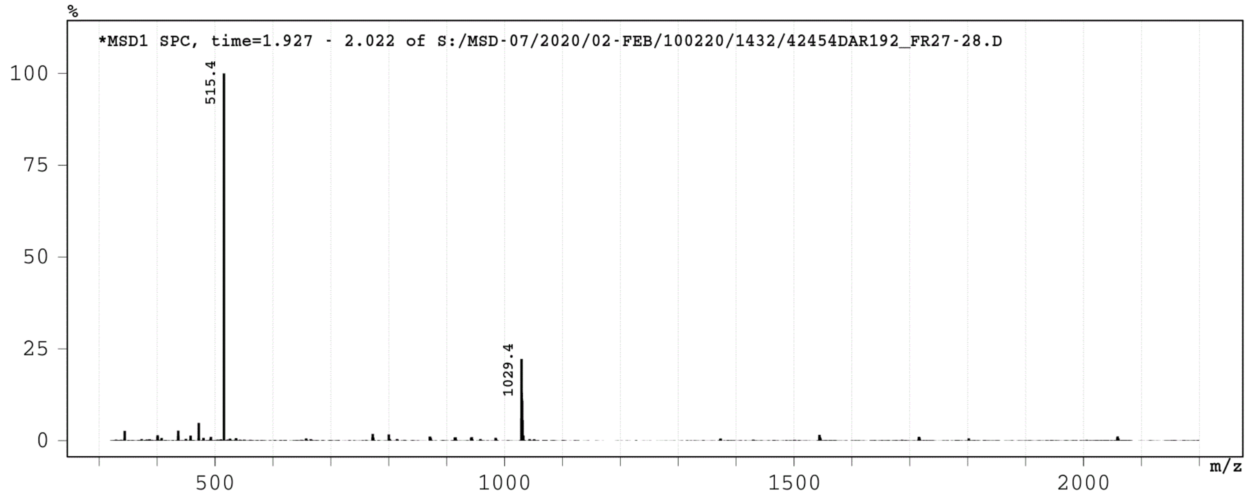


Signal 1: DAD1 A, Sig=220,4 Ref=off

| Peak # | RT [min] | Width [min] | Area | Area % |
|---|---|---|---|---|
| 1 | 1.894 | 1.56 | 99.9 | 3.8 |
| 2 | 1.950 | 0.12 | 2326.1 | 87.9 |
| 3 | 2.018 | 3.18 | 221.0 | 8.3 |

Data File : S:/MSD-07/2020/02-FEB/100220/1432/42454DAR193_FR4->

Injection Date : 10-Feb-20, 18:42:21
Sample Name : 42454DAR193_FR46
Remarks : Baseline corrected
Acq. Method : D:/METHODS/5-95-6M.M
column used : Je_559
Analysis Method : C:/CHEM32/1/METHODS/JPT-LCMS.M

Sample Info : 42454_193-Dr. Martine Darwish H-HGYERYPAL-OH

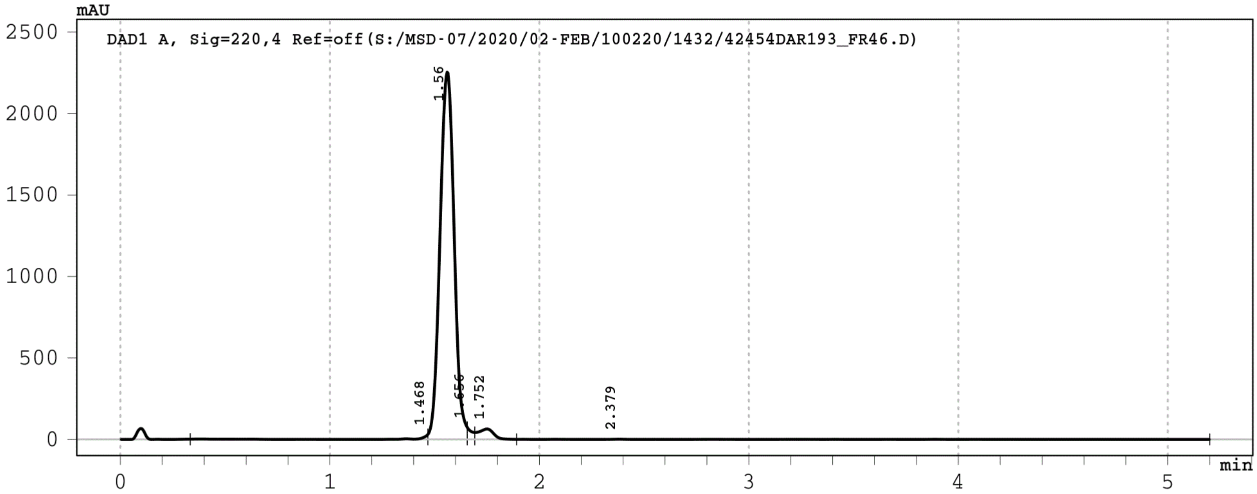


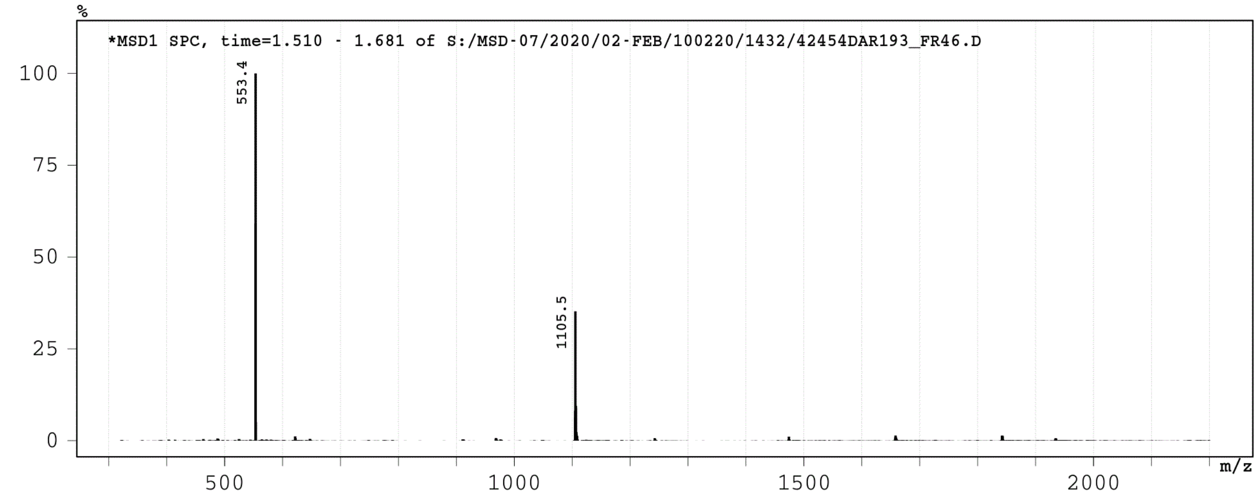


Signal 1: DAD1 A, Sig=220,4 Ref=off

| Peak # | RT [min] | Width [min] | Area | Area % |
|---|---|---|---|---|
| 1 | 1.468 | 1.13 | 82.8 | 0.7 |
| 2 | 1.560 | 0.19 | 10545.9 | 95.1 |
| 3 | 1.656 | 0.04 | 112.9 | 1.0 |
| 4 | 1.752 | 0.20 | 328.9 | 3.0 |
| 5 | 2.379 | 3.31 | 19.5 | 0.2 |

Data File : S:/MSD-07/2020/02-FEB/100220/1432/42454DAR194_FR5->

Injection Date : 10-Feb-20, 18:50:44
Sample Name : 42454DAR194_FR57-58
Remarks : Baseline corrected
Acq. Method : D:/METHODS/5-95-6M.M
column used : Je_559
Analysis Method : C:/CHEM32/1/METHODS/JPT-LCMS.M

Sample Info : 42454_194-Dr. Martine Darwish H-SSASRRLLV-OH

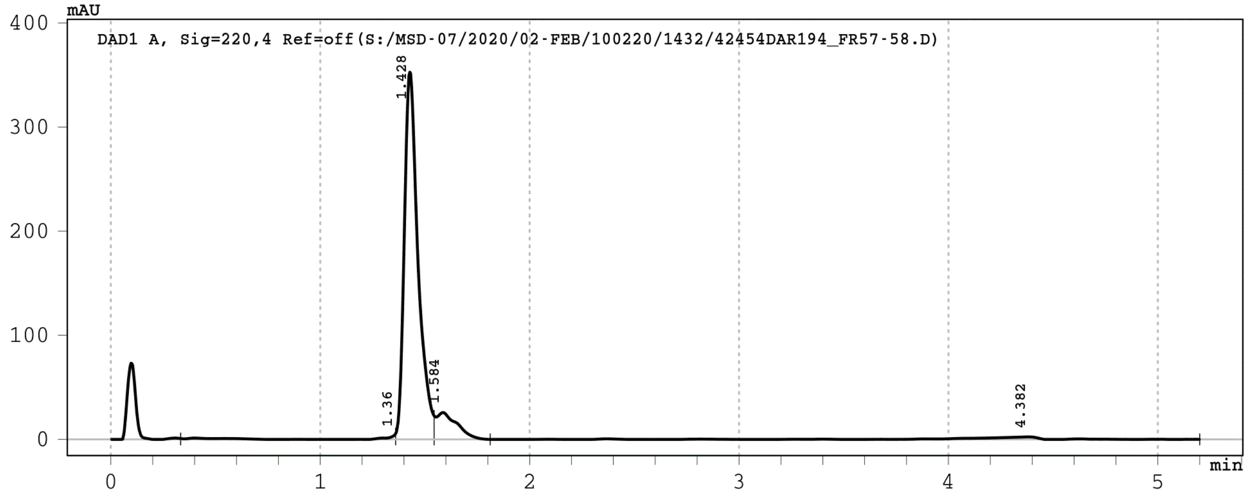


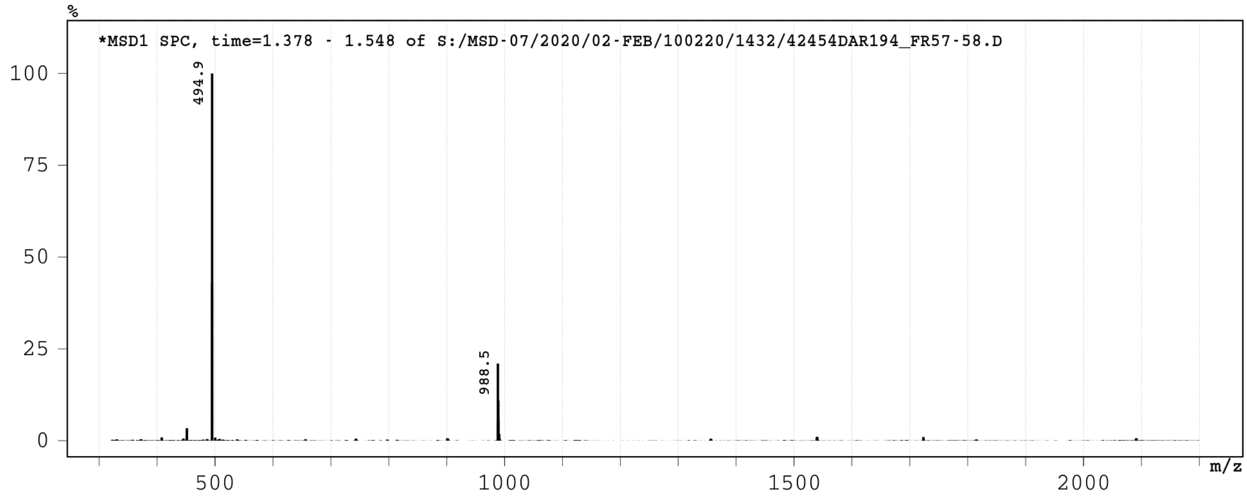


Signal 1: DAD1 A, Sig=220,4 Ref=off

| Peak # | RT [min] | Width [min] | Area | Area % |
|---|---|---|---|---|
| 1 | 1.360 | 1.03 | 28.6 | 1.5 |
| 2 | 1.428 | 0.18 | 1614.3 | 85.6 |
| 3 | 1.584 | 0.27 | 183.1 | 9.7 |
| 4 | 4.382 | 3.39 | 60.1 | 3.2 |

Data File : S:/MSD-07/2020/02-FEB/110220/1433/42454DAR195_FR3->

Injection Date : 12-Feb-20, 02:17:27
Sample Name : 42454DAR195_FR34-36
Remarks : Baseline corrected
Acq. Method : D:/METHODS/5-95-6M.M
column used : Je_559
Analysis Method : C:/CHEM32/1/METHODS/JPT-LCMS.M

Sample Info : 42454_195-Dr. Martine Darwish H-VQFMSCNL-OH

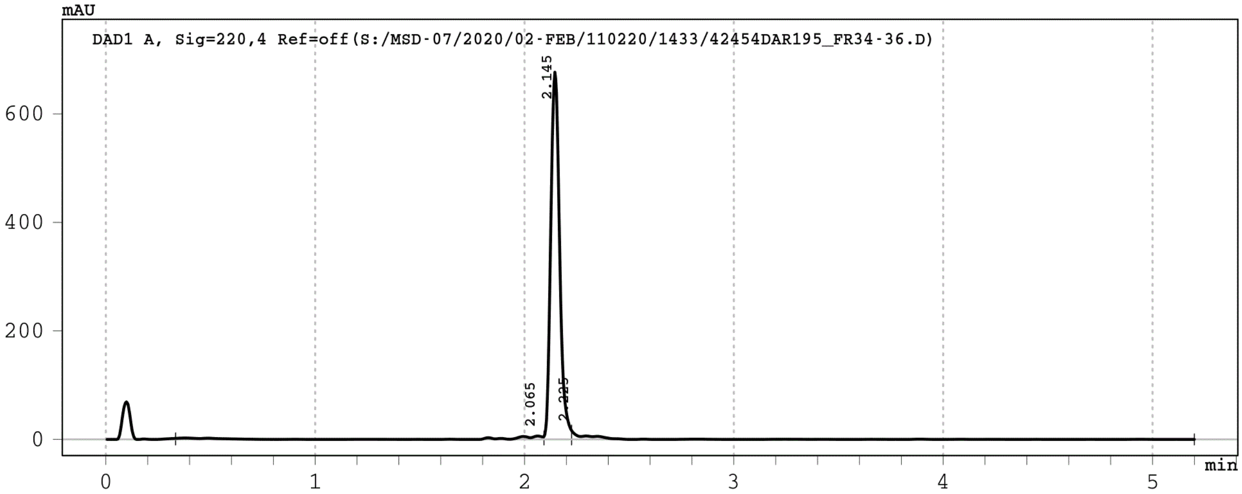


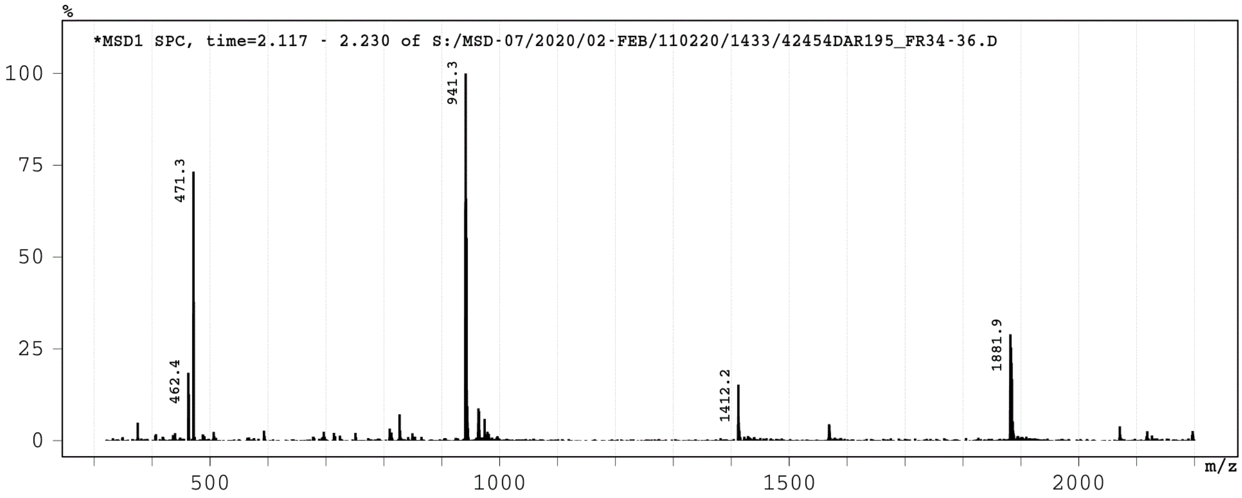


Signal 1: DAD1 A, Sig=220,4 Ref=off

| Peak # | RT [min] | Width [min] | Area | Area % |
|---|---|---|---|---|
| 1 | 2.065 | 1.76 | 89.4 | 4.1 |
| 2 | 2.145 | 0.13 | 1998.9 | 92.1 |
| 3 | 2.225 | 2.97 | 81.6 | 3.8 |

Data File : S:/MSD-07/2020/02-FEB/060220/1429/42454DAR196_FR4->

Injection Date : 06-Feb-20, 16:58:21
Sample Name : 42454DAR196_FR4-5
Remarks : Baseline corrected
Acq. Method : D:/METHODS/5-95-6M.M
column used : Je_559
Analysis Method : C:/CHEM32/1/METHODS/JPT-LCMS.M

Sample Info : 42454_196-Dr. Martine Darwish H-ASYHNMGDLF-OH

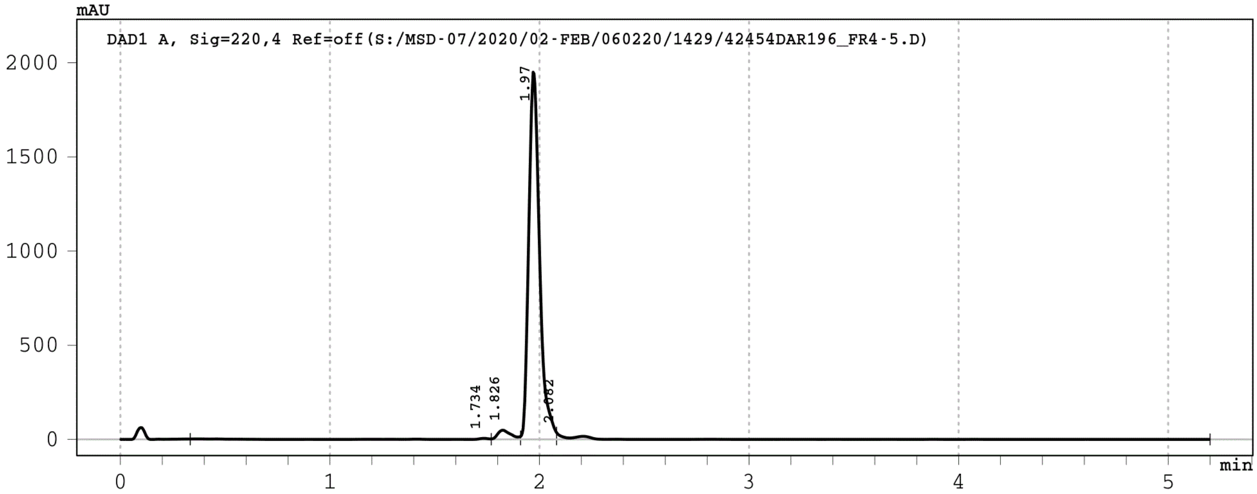


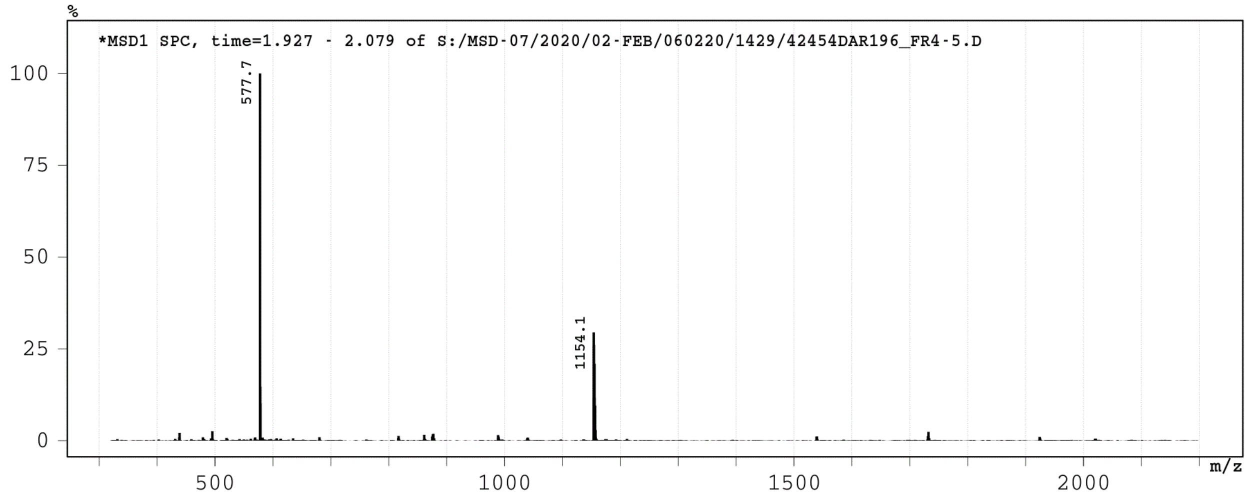


Signal 1: DAD1 A, Sig=220,4 Ref=off

| Peak # | RT [min] | Width [min] | Area | Area % |
|---|---|---|---|---|
| 1 | 1.734 | 1.44 | 59.1 | 0.8 |
| 2 | 1.826 | 0.14 | 218.7 | 2.9 |
| 3 | 1.970 | 0.17 | 7004.7 | 94.2 |
| 4 | 2.082 | 3.12 | 156.6 | 2.1 |

Data File : S:/MSD-07/2020/02-FEB/110220/1433/42454DAR197_FR4->

Injection Date : 12-Feb-20, 02:25:51
Sample Name : 42454DAR197_FR42-44
Remarks : Baseline corrected
Acq. Method : D:/METHODS/5-95-6M.M
column used : Je_559
Analysis Method : C:/CHEM32/1/METHODS/JPT-LCMS.M

Sample Info : 42454_197-Dr. Martine Darwish H-SSFVPVGL-OH

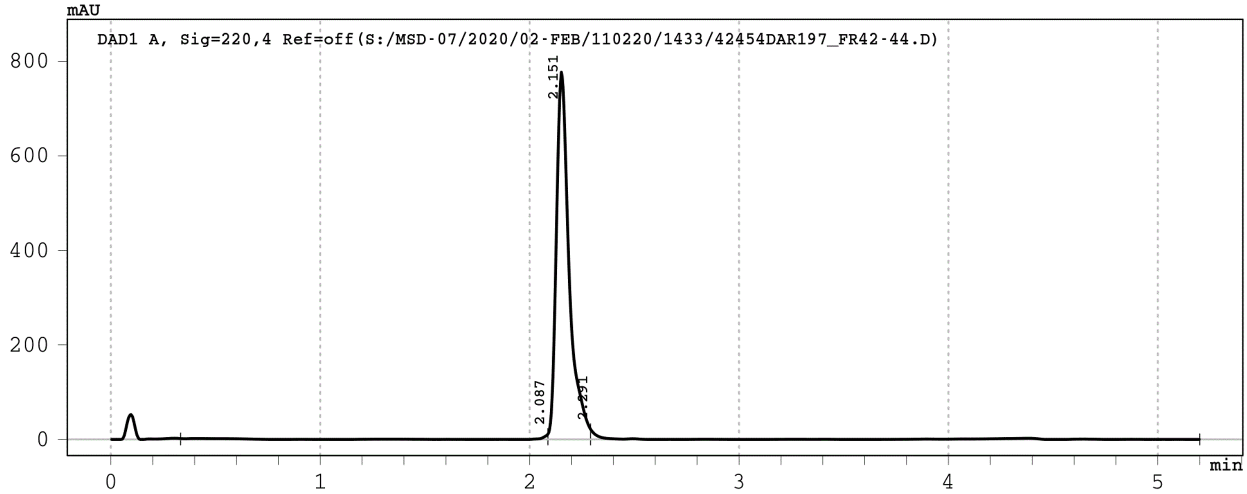


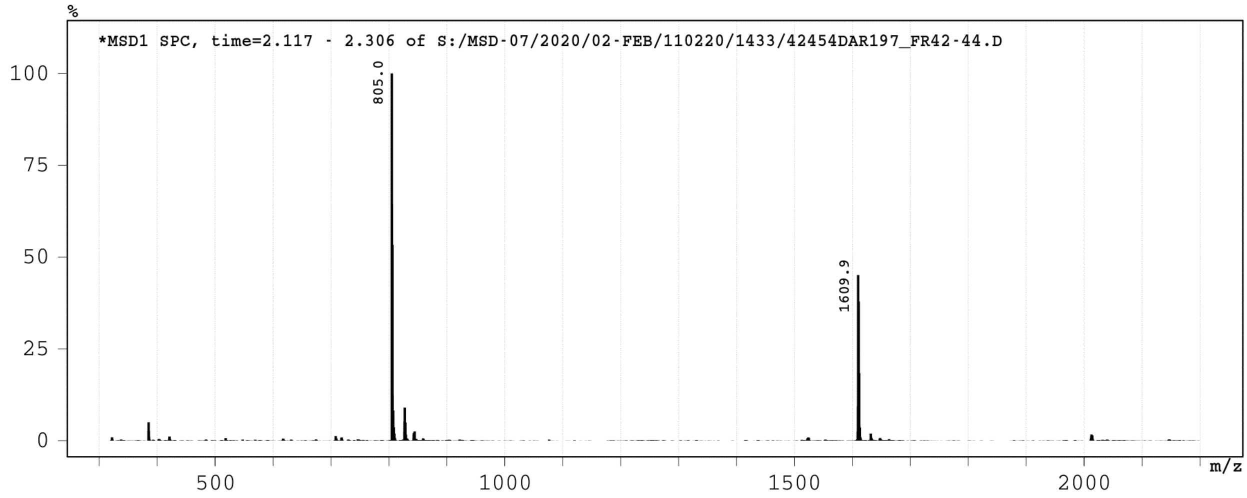


Signal 1: DAD1 A, Sig=220,4 Ref=off

| Peak # | RT [min] | Width [min] | Area | Area % |
|---|---|---|---|---|
| 1 | 2.087 | 1.75 | 55.9 | 1.6 |
| 2 | 2.151 | 0.20 | 3317.8 | 95.4 |
| 3 | 2.291 | 2.91 | 104.2 | 3.0 |

Data File : S:/MSD-07/2020/02-FEB/100220/1432/42454DAR198_FR6->

Injection Date : 10-Feb-20, 18:59:06
Sample Name : 42454DAR198_FR67
Remarks : Baseline corrected
Acq. Method : D:/METHODS/5-95-6M.M
column used : Je_559
Analysis Method : C:/CHEM32/1/METHODS/JPT-LCMS.M

Sample Info : 42454_198-Dr. Martine Darwish H-VLYLLDTSL-OH

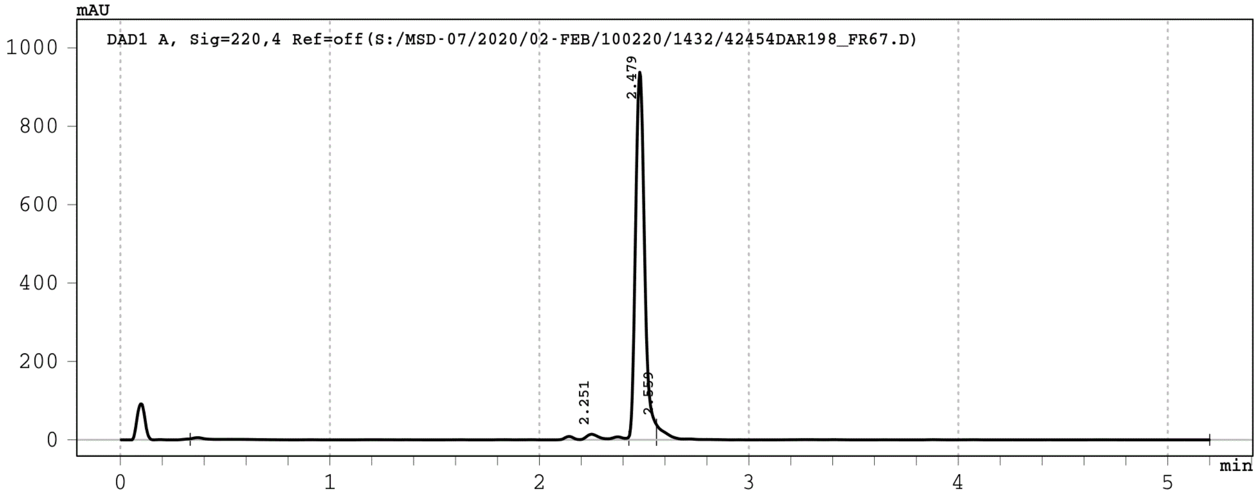


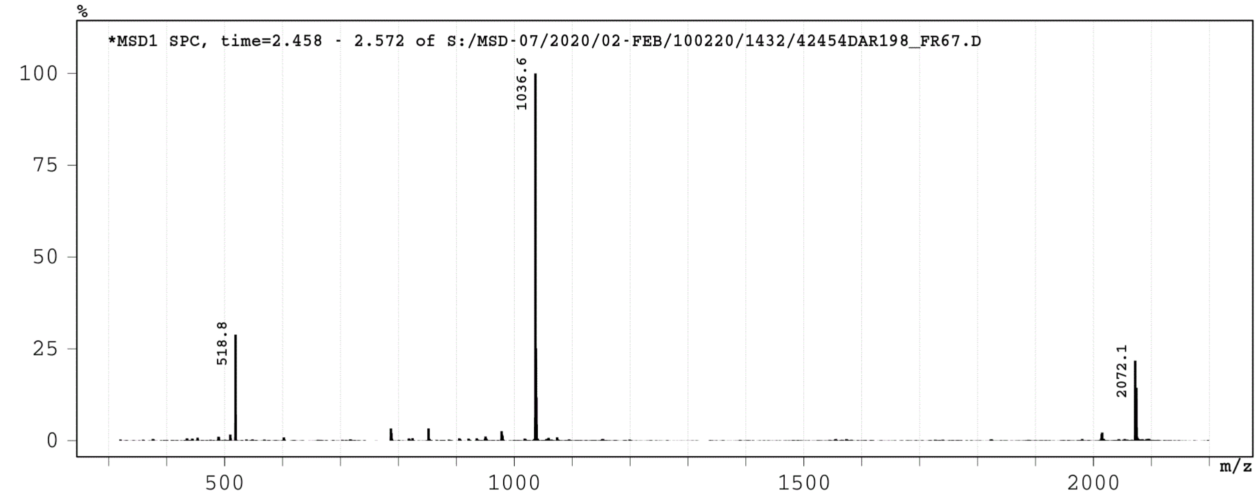


Signal 1: DAD1 A, Sig=220,4 Ref=off

| Peak # | RT [min] | Width [min] | Area | Area % |
|---|---|---|---|---|
| 1 | 2.251 | 2.09 | 161.5 | 5.2 |
| 2 | 2.479 | 0.13 | 2821.4 | 90.7 |
| 3 | 2.559 | 2.64 | 129.4 | 4.2 |

Data File : S:/MSD-07/2020/02-FEB/100220/1432/42454DAR199_FR7->

Injection Date : 10-Feb-20, 19:07:30
Sample Name : 42454DAR199_FR79-80
Remarks : Baseline corrected
Acq. Method : D:/METHODS/5-95-6M.M
column used : Je_559
Analysis Method : C:/CHEM32/1/METHODS/JPT-LCMS.M

Sample Info : 42454_199-Dr. Martine Darwish H-IGAGFFSDV-OH

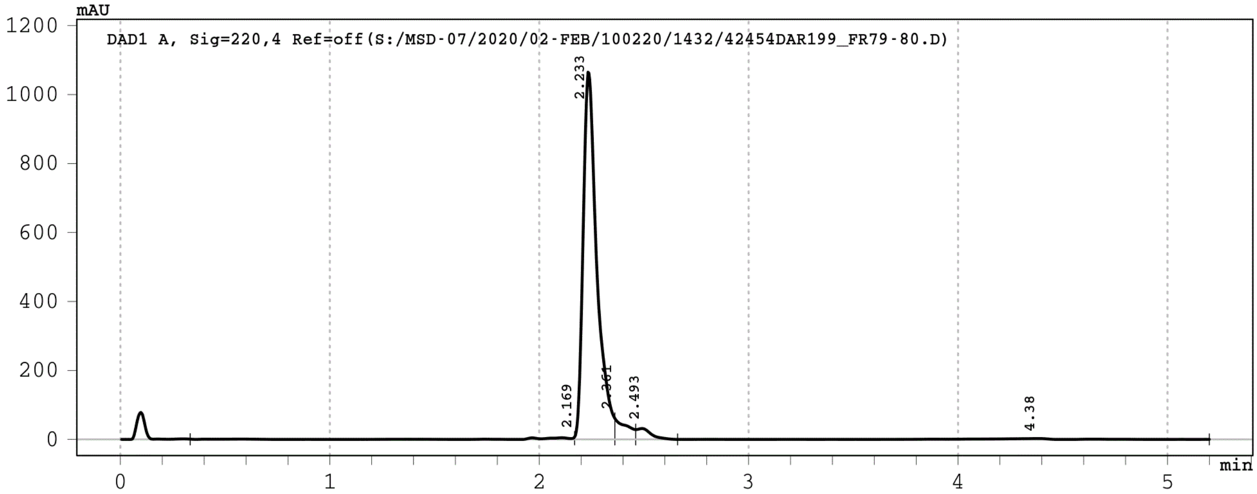


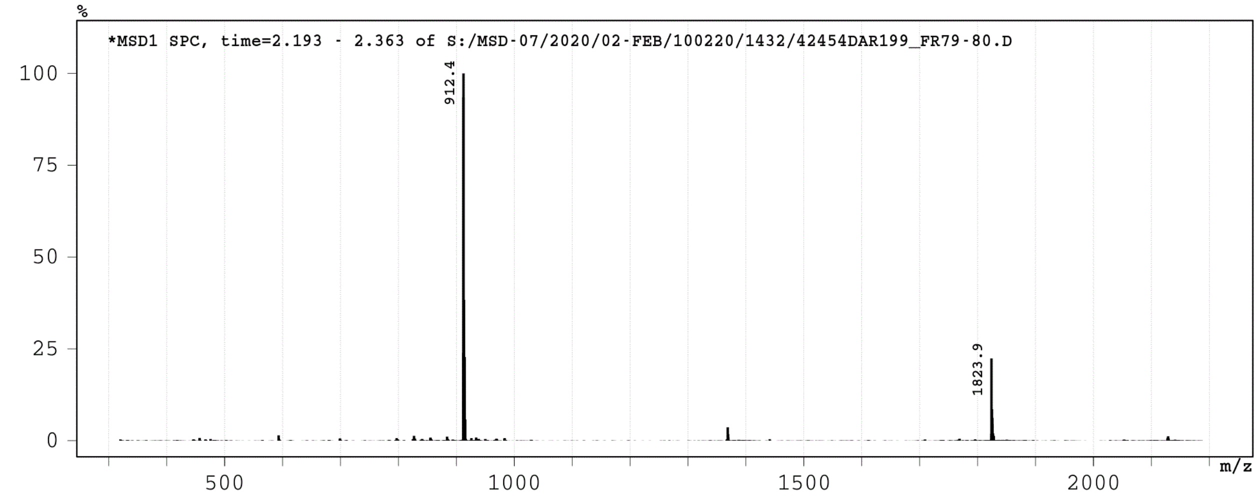


Signal 1: DAD1 A, Sig=220,4 Ref=off

| Peak # | RT [min] | Width [min] | Area | Area % |
|---|---|---|---|---|
| 1 | 2.169 | 1.84 | 70.2 | 1.3 |
| 2 | 2.233 | 0.19 | 4849.7 | 90.2 |
| 3 | 2.361 | 0.10 | 244.9 | 4.6 |
| 4 | 2.493 | 0.20 | 152.5 | 2.8 |
| 5 | 4.380 | 2.54 | 60.2 | 1.1 |

Data File : S:/MSD-07/2020/02-FEB/140220/1435/42454DAR200_FR1->

Injection Date : 15-Feb-20, 01:01:27
Sample Name : 42454DAR200_FR19-21
Remarks : Baseline corrected
Acq. Method : D:/METHODS/5-95-6M.M
column used : Je_559
Analysis Method : C:/CHEM32/1/METHODS/JPT-LCMS.M

Sample Info : 42454_200-Dr. Martine Darwish H-FTPSHPPL-OH

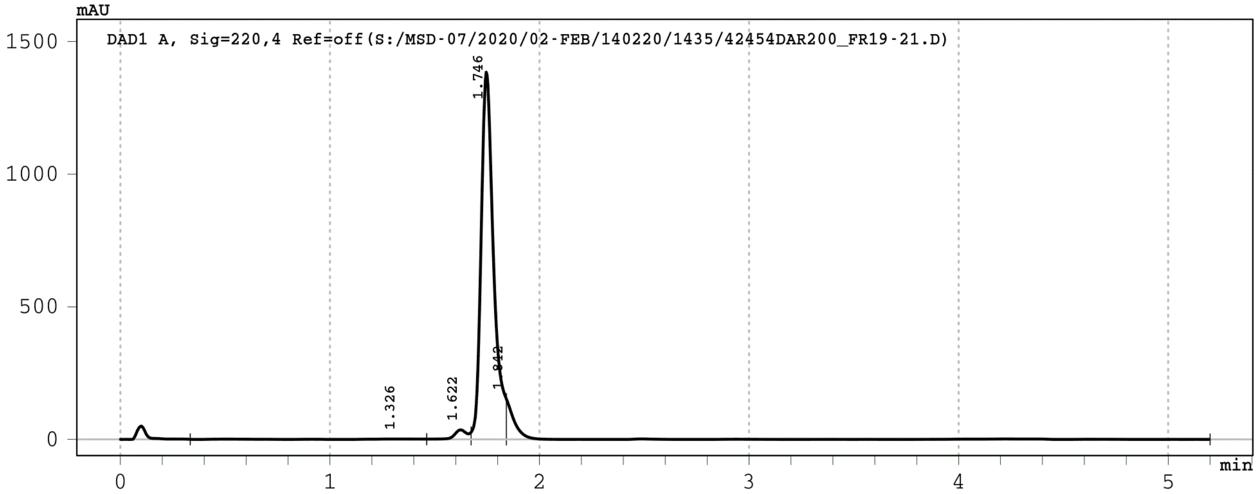


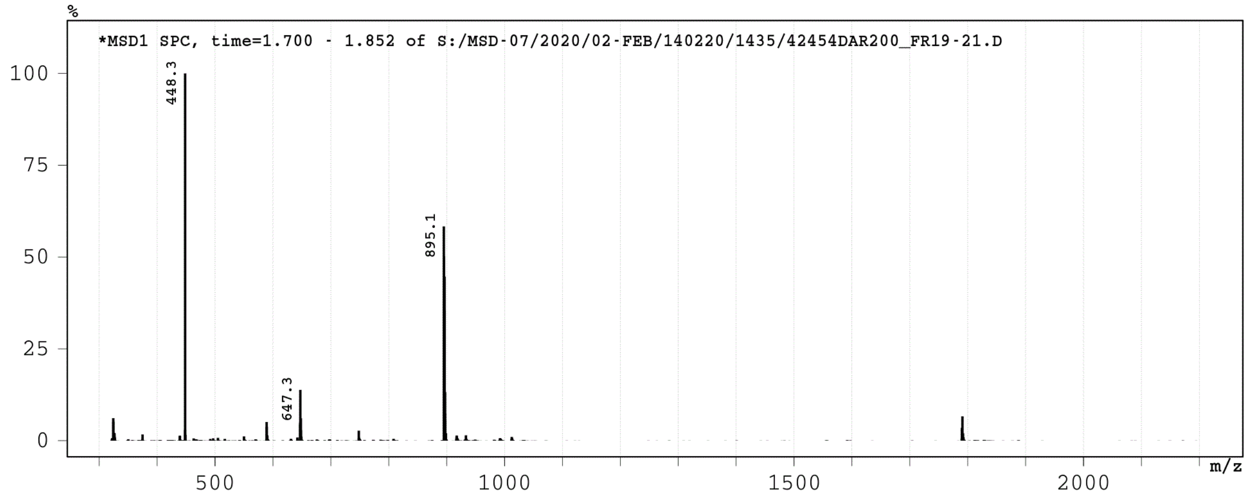


Signal 1: DAD1 A, Sig=220,4 Ref=off

| Peak # | RT [min] | Width [min] | Area | Area % |
|---|---|---|---|---|
| 1 | 1.326 | 1.13 | 42.9 | 0.7 |
| 2 | 1.622 | 0.21 | 159.8 | 2.5 |
| 3 | 1.746 | 0.17 | 5733.2 | 89.7 |
| 4 | 1.842 | 3.36 | 455.8 | 7.1 |

Data File : S:/MSD-07/2020/02-FEB/060220/1429/42454DAR201_FR1->

Injection Date : 06-Feb-20, 17:06:43
Sample Name : 42454DAR201_FR12-13
Remarks : Baseline corrected
Acq. Method : D:/METHODS/5-95-6M.M
column used : Je_559
Analysis Method : C:/CHEM32/1/METHODS/JPT-LCMS.M

Sample Info : 42454_201-Dr. Martine Darwish H-VAMCRLGIF-OH

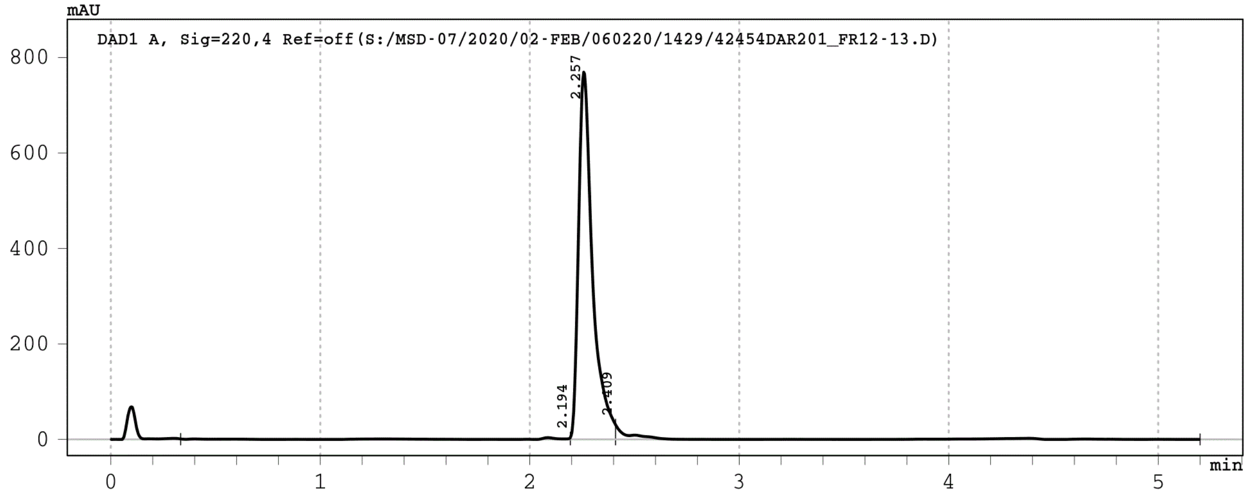


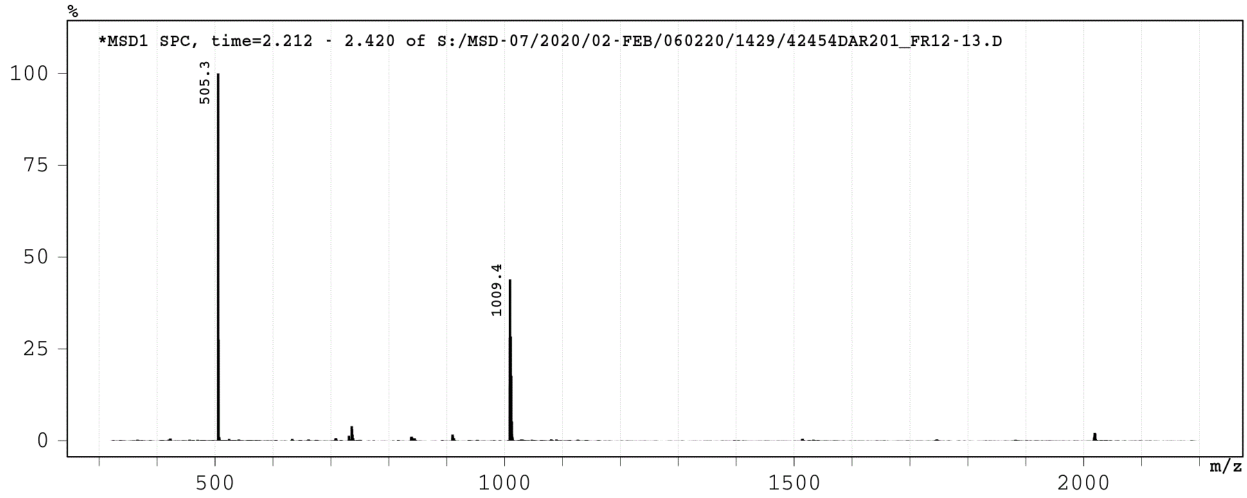


Signal 1: DAD1 A, Sig=220,4 Ref=off

| Peak # | RT [min] | Width [min] | Area | Area % |
|---|---|---|---|---|
| 1 | 2.194 | 1.86 | 48.3 | 1.3 |
| 2 | 2.257 | 0.22 | 3582.2 | 94.2 |
| 3 | 2.409 | 2.79 | 171.7 | 4.5 |

Data File : S:/MSD-07/2020/02-FEB/060220/1429/42454DAR202_FR2->

Injection Date : 06-Feb-20, 17:15:06
Sample Name : 42454DAR202_FR23-24
Remarks : Baseline corrected
Acq. Method : D:/METHODS/5-95-6M.M
column used : Je_559
Analysis Method : C:/CHEM32/1/METHODS/JPT-LCMS.M

Sample Info : 42454_202-Dr. Martine Darwish H-IMTQHLEPI-OH

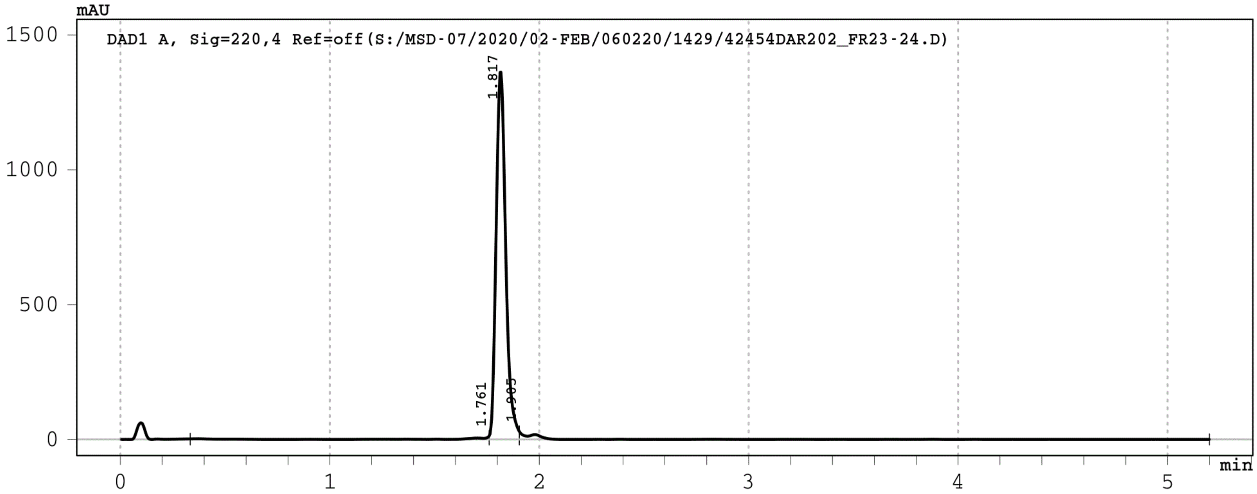


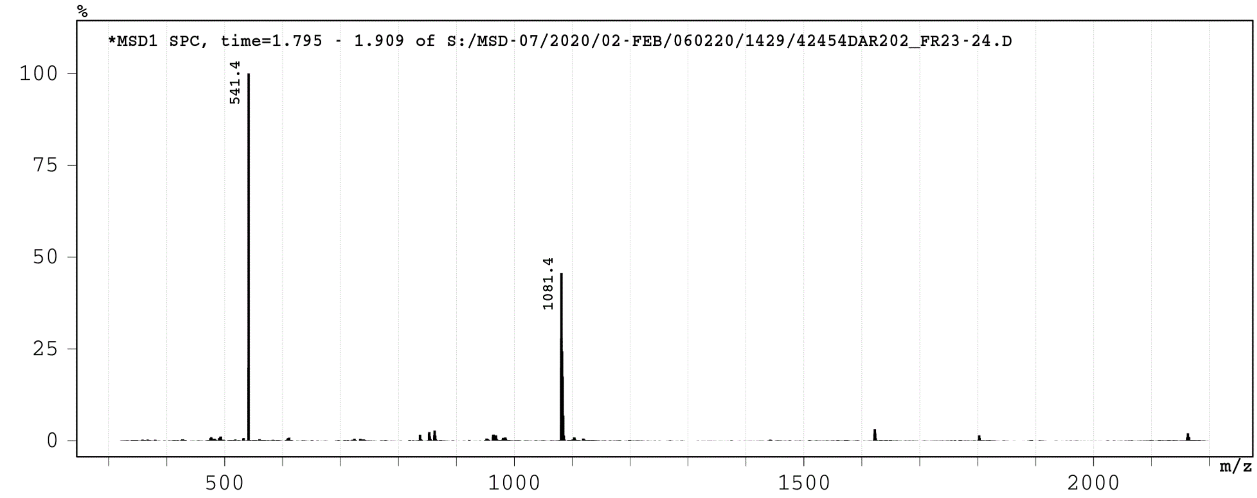


Signal 1: DAD1 A, Sig=220,4 Ref=off

| Peak # | RT [min] | Width [min] | Area | Area % |
|---|---|---|---|---|
| 1 | 1.761 | 1.43 | 70.4 | 1.5 |
| 2 | 1.817 | 0.14 | 4351.4 | 95.7 |
| 3 | 1.905 | 3.30 | 125.7 | 2.8 |

Data File : S:/MSD-07/2020/02-FEB/100220/1432/42454DAR203_FR7->

Injection Date : 10-Feb-20, 19:15:53
Sample Name : 42454DAR203_FR7-8
Remarks : Baseline corrected
Acq. Method : D:/METHODS/5-95-6M.M
column used : Je_559
Analysis Method : C:/CHEM32/1/METHODS/JPT-LCMS.M

Sample Info : 42454_203-Dr. Martine Darwish H-SAWVPFGGL-OH

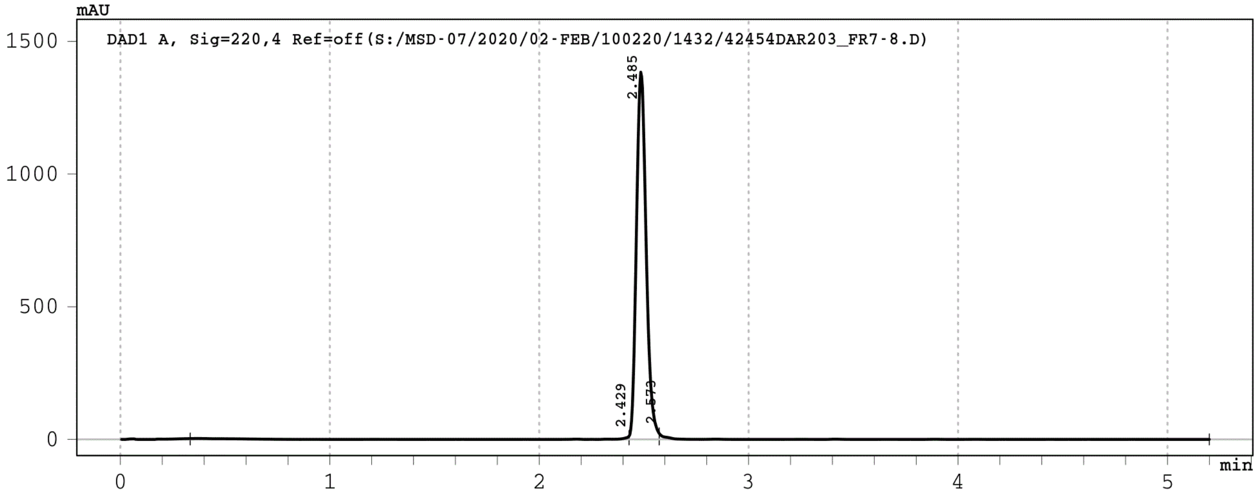


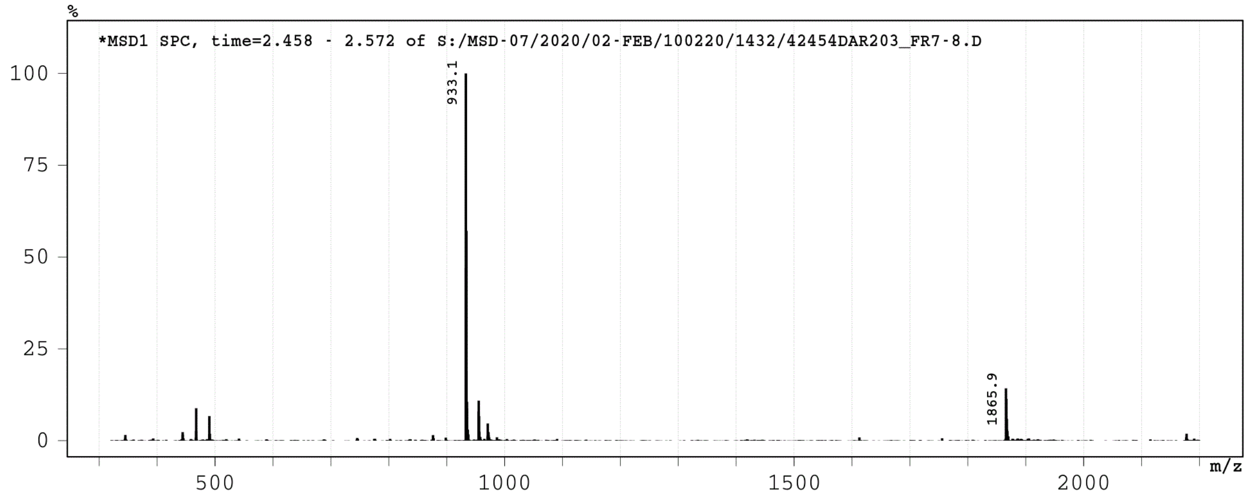


Signal 1: DAD1 A, Sig=220,4 Ref=off

| Peak # | RT [min] | Width [min] | Area | Area % |
|---|---|---|---|---|
| 1 | 2.429 | 2.10 | 77.5 | 1.7 |
| 2 | 2.485 | 0.14 | 4484.8 | 97.1 |
| 3 | 2.573 | 2.63 | 55.8 | 1.2 |

Data File : S:/MSD-07/2020/02-FEB/140220/1435/42454DAR204_FR2->

Injection Date : 15-Feb-20, 01:10:01
Sample Name : 42454DAR204_FR25-28
Remarks : Baseline corrected
Acq. Method : D:/METHODS/5-95-6M.M
column used : Je_559
Analysis Method : C:/CHEM32/1/METHODS/JPT-LCMS.M

Sample Info : 42454_204-Dr. Martine Darwish H-ILLRLKFL-OH

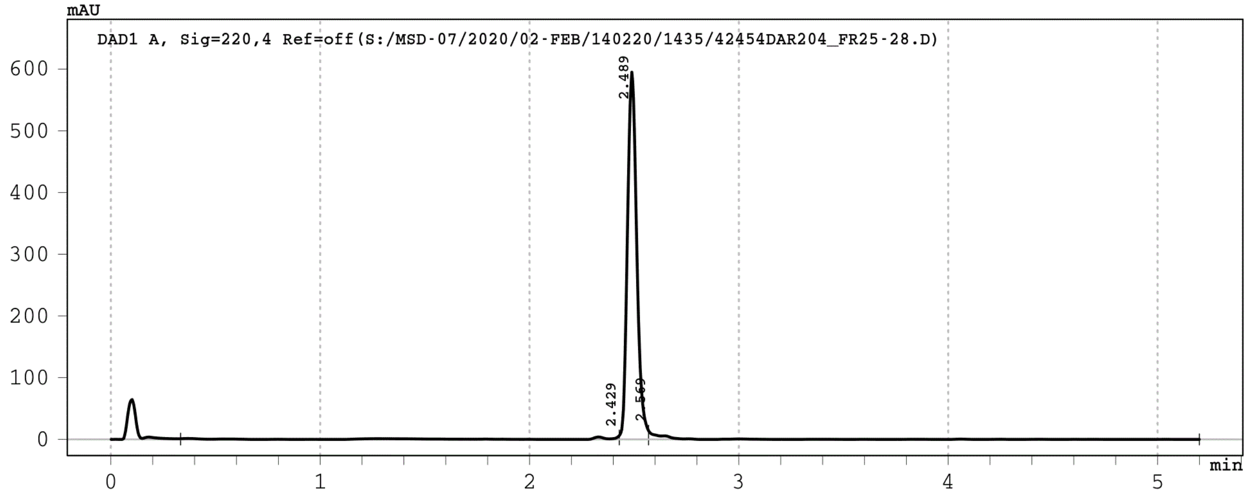


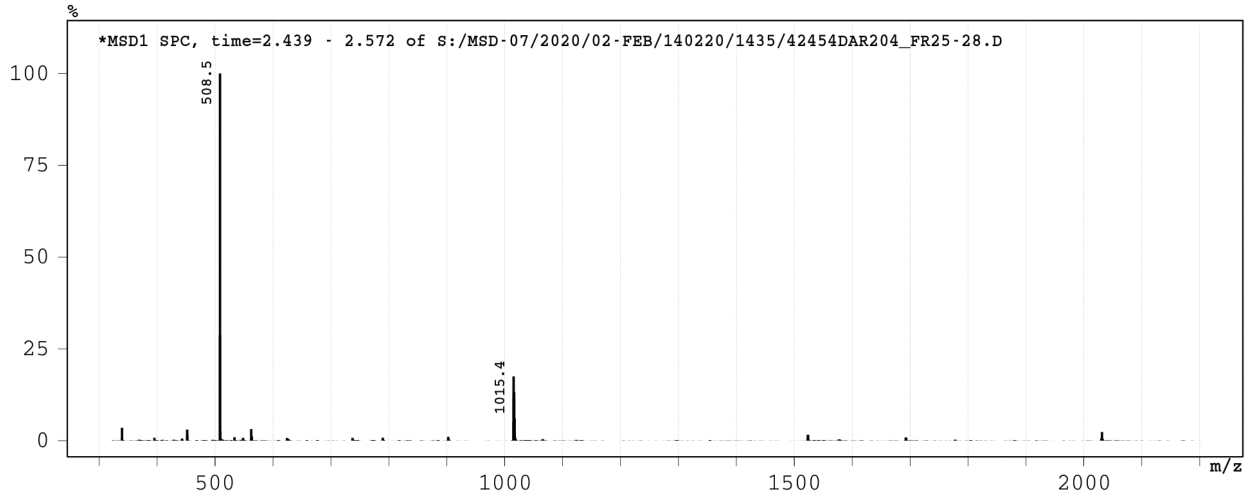


Signal 1: DAD1 A, Sig=220,4 Ref=off

| Peak # | RT [min] | Width [min] | Area | Area % |
|---|---|---|---|---|
| 1 | 2.429 | 2.10 | 63.9 | 3.2 |
| 2 | 2.489 | 0.14 | 1844.3 | 93.5 |
| 3 | 2.569 | 2.63 | 63.6 | 3.2 |

Data File : S:/MSD-07/2020/02-FEB/100220/1432/42454DAR205_FR2->

Injection Date : 10-Feb-20, 19:24:17
Sample Name : 42454DAR205_FR22
Remarks : Baseline corrected
Acq. Method : D:/METHODS/5-95-6M.M
column used : Je_559
Analysis Method : C:/CHEM32/1/METHODS/JPT-LCMS.M

Sample Info : 42454_205-Dr. Martine Darwish H-VVFNHFYNI-OH

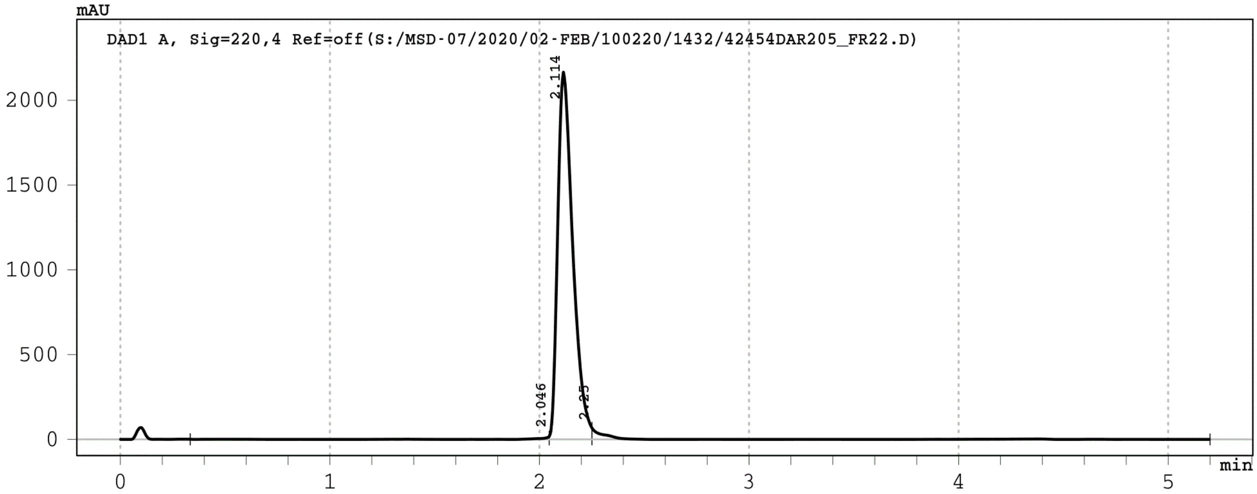


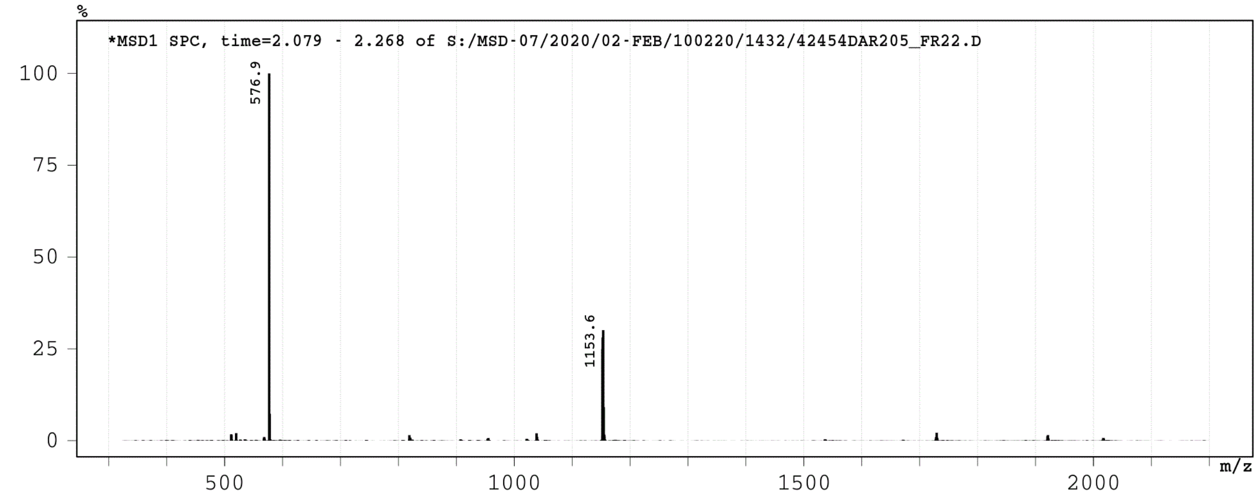


Signal 1: DAD1 A, Sig=220,4 Ref=off

| Peak # | RT [min] | Width [min] | Area | Area % |
|---|---|---|---|---|
| 1 | 2.046 | 1.71 | 64.2 | 0.6 |
| 2 | 2.114 | 0.20 | 10876.3 | 96.8 |
| 3 | 2.250 | 2.95 | 291.9 | 2.6 |

Data File : S:/MSD-07/2020/02-FEB/100220/1432/42454DAR206_FR3->

Injection Date : 10-Feb-20, 19:32:40
Sample Name : 42454DAR206_FR34-35
Remarks : Baseline corrected
Acq. Method : D:/METHODS/5-95-6M.M
column used : Je_559
Analysis Method : C:/CHEM32/1/METHODS/JPT-LCMS.M

Sample Info : 42454_206-Dr. Martine Darwish H-SVAGFNPAL-OH

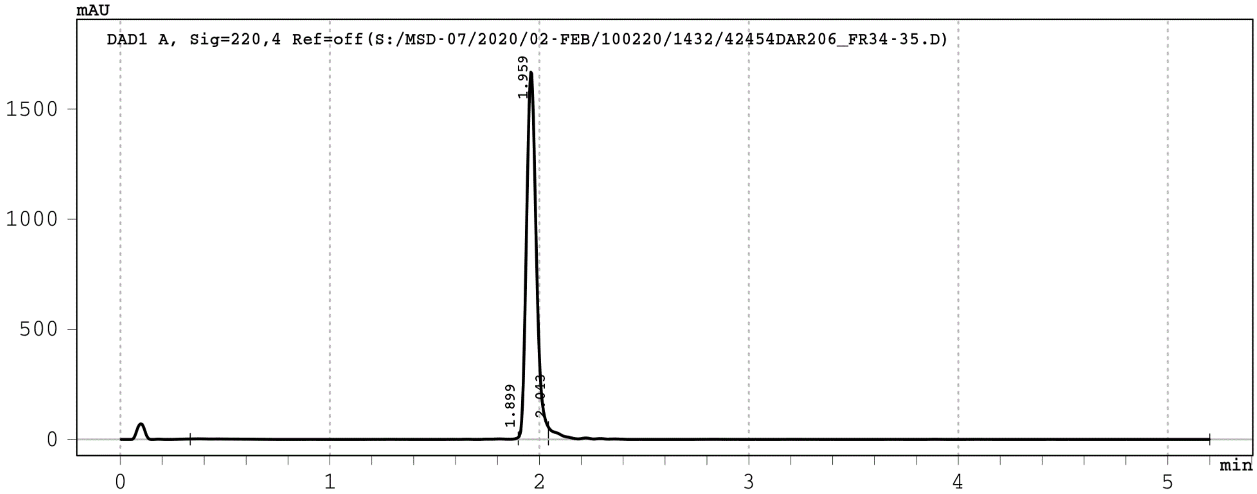


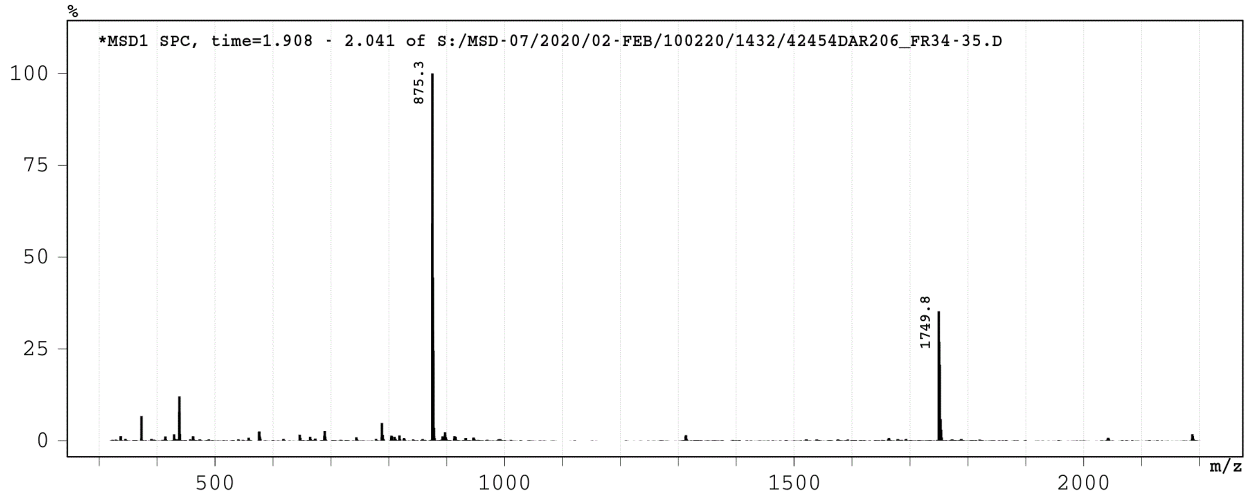


Signal 1: DAD1 A, Sig=220,4 Ref=off

| Peak # | RT [min] | Width [min] | Area | Area % |
|---|---|---|---|---|
| 1 | 1.899 | 1.57 | 55.7 | 1.0 |
| 2 | 1.959 | 0.14 | 5421.6 | 95.3 |
| 3 | 2.043 | 3.16 | 212.9 | 3.7 |

Data File : S:/MSD-07/2020/02-FEB/060220/1429/42454DAR207_FR3->

Injection Date : 06-Feb-20, 17:23:29
Sample Name : 42454DAR207_FR34-36
Remarks : Baseline corrected
Acq. Method : D:/METHODS/5-95-6M.M
column used : Je_559
Analysis Method : C:/CHEM32/1/METHODS/JPT-LCMS.M

Sample Info : 42454_207-Dr. Martine Darwish H-MGVMNRRPI-OH

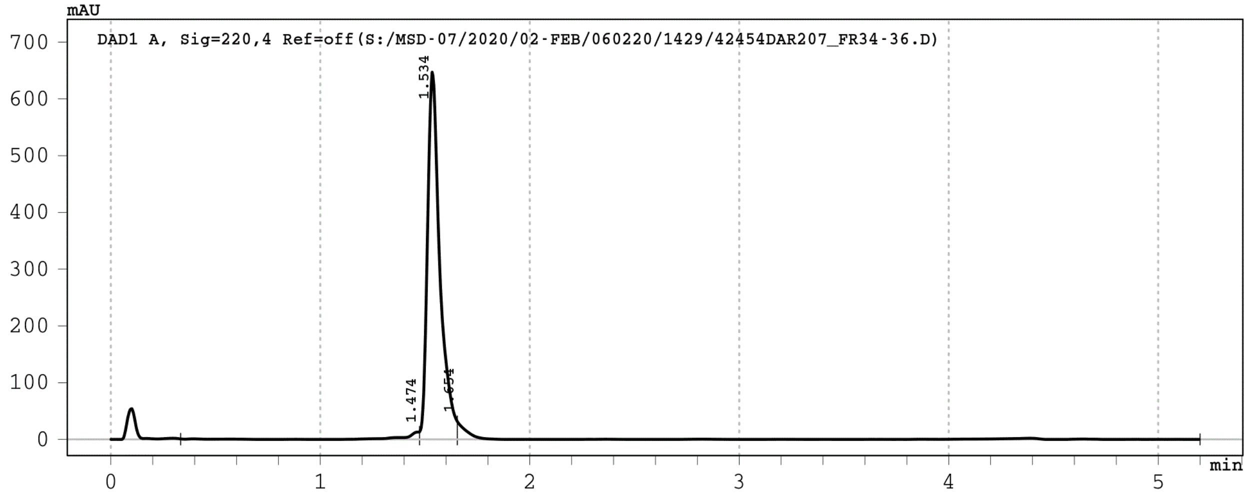


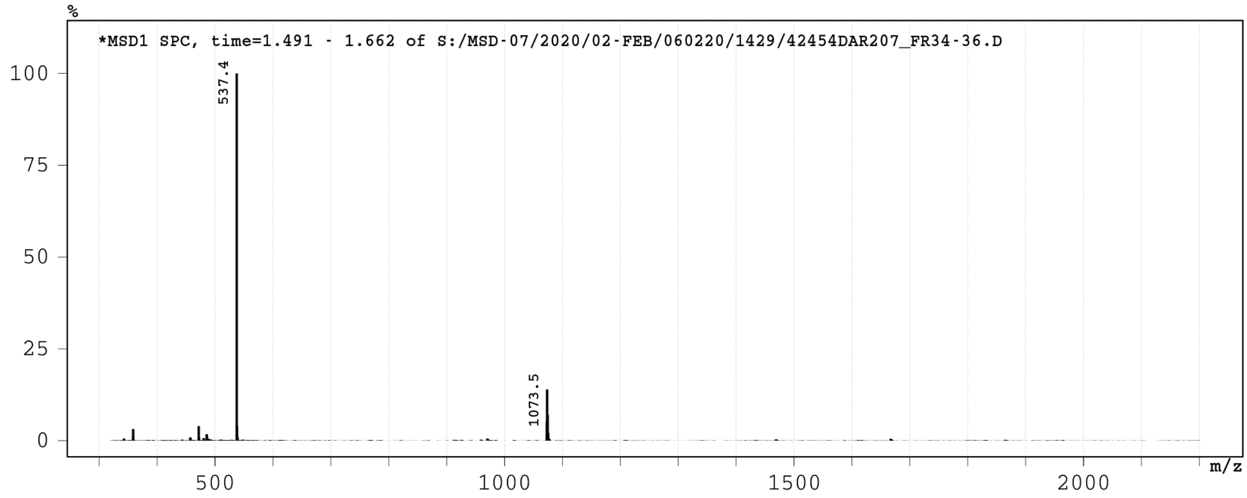


Signal 1: DAD1 A, Sig=220,4 Ref=off

| Peak # | RT [min] | Width [min] | Area | Area % |
|---|---|---|---|---|
| 1 | 1.474 | 1.14 | 72.4 | 2.5 |
| 2 | 1.534 | 0.18 | 2670.6 | 92.8 |
| 3 | 1.654 | 3.55 | 135.4 | 4.7 |

Data File : S:/MSD-07/2020/02-FEB/060220/1429/42454DAR208_FR3->

Injection Date : 06-Feb-20, 17:31:51
Sample Name : 42454DAR208_FR38-39
Remarks : Baseline corrected
Acq. Method : D:/METHODS/5-95-6M.M
column used : Je_559
Analysis Method : C:/CHEM32/1/METHODS/JPT-LCMS.M

Sample Info : 42454_208-Dr. Martine Darwish H-MSLGTLLLL-OH

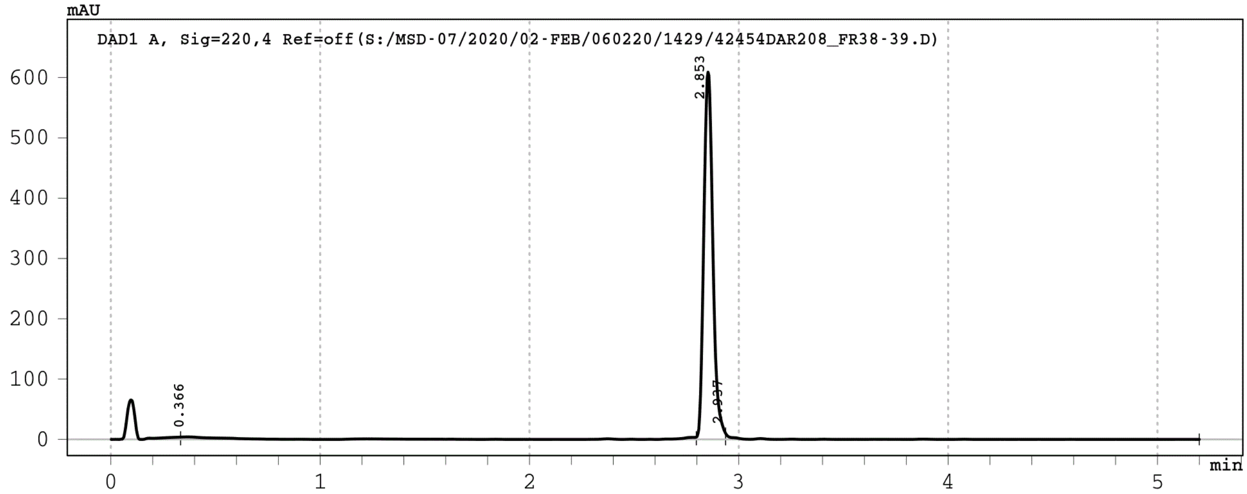


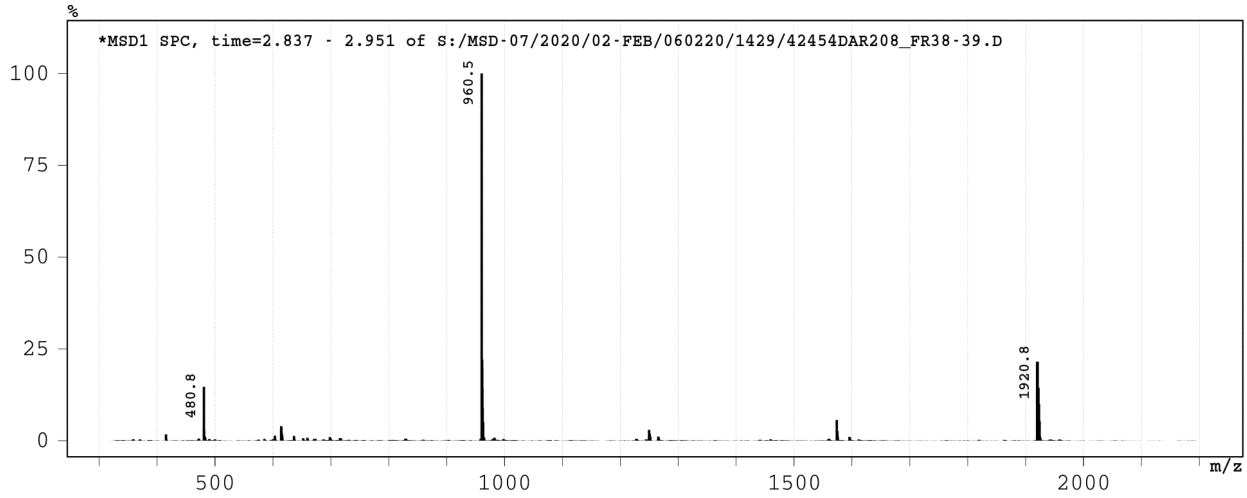


Signal 1: DAD1 A, Sig=220,4 Ref=off

| Peak # | RT [min] | Width [min] | Area | Area % |
|---|---|---|---|---|
| 1 | 0.366 | 2.46 | 95.6 | 4.9 |
| 2 | 2.853 | 0.14 | 1836.9 | 93.7 |
| 3 | 2.937 | 2.26 | 26.8 | 1.4 |

Data File : S:/MSD-07/2020/02-FEB/100220/1432/42454DAR209_FR4->

Injection Date : 10-Feb-20, 19:41:05
Sample Name : 42454DAR209_FR43-44
Remarks : Baseline corrected
Acq. Method : D:/METHODS/5-95-6M.M
column used : Je_559
Analysis Method : C:/CHEM32/1/METHODS/JPT-LCMS.M

Sample Info : 42454_209-Dr. Martine Darwish H-AQARNSLEL-OH

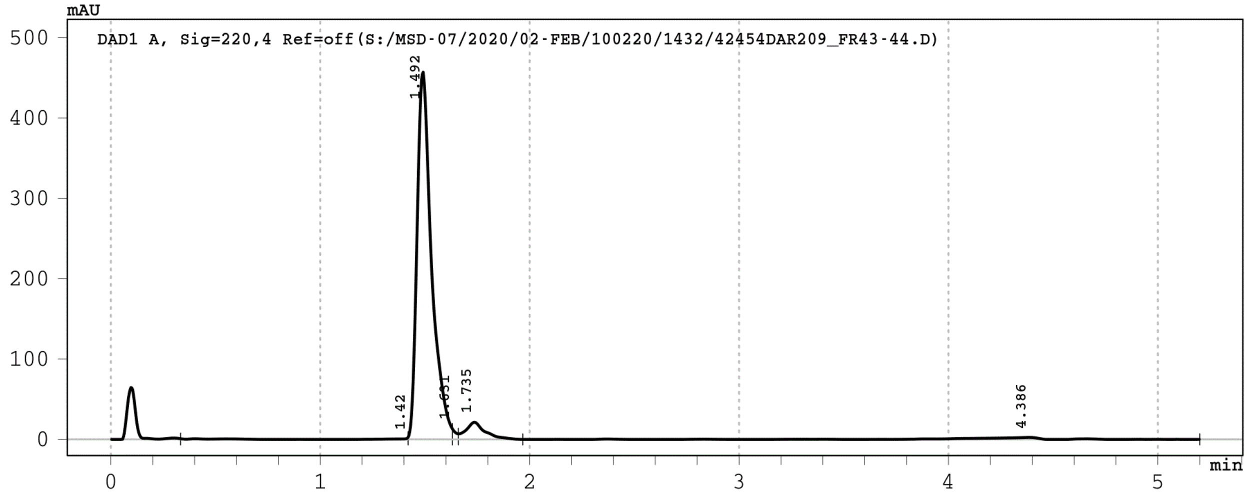


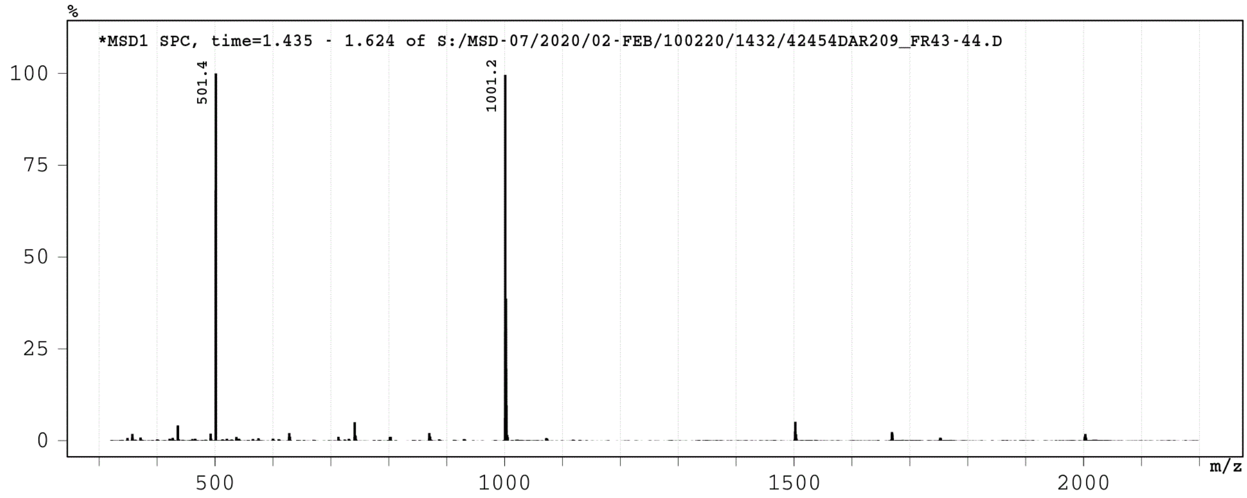


Signal 1: DAD1 A, Sig=220,4 Ref=off

| Peak # | RT [min] | Width [min] | Area | Area % |
|---|---|---|---|---|
| 1 | 1.420 | 1.09 | 16.2 | 0.7 |
| 2 | 1.492 | 0.21 | 2210.7 | 90.5 |
| 3 | 1.631 | 0.03 | 15.2 | 0.6 |
| 4 | 1.735 | 0.31 | 138.9 | 5.7 |
| 5 | 4.386 | 3.23 | 62.9 | 2.6 |

Data File : S:/MSD-07/2020/02-FEB/140220/1435/42454DAR210_FR3->

Injection Date : 15-Feb-20, 01:18:29
Sample Name : 42454DAR210_FR33
Remarks : Baseline corrected
Acq. Method : D:/METHODS/5-95-6M.M
column used : Je_559
Analysis Method : C:/CHEM32/1/METHODS/JPT-LCMS.M

Sample Info : 42454_210-Dr. Martine Darwish H-FSYIVELL-OH

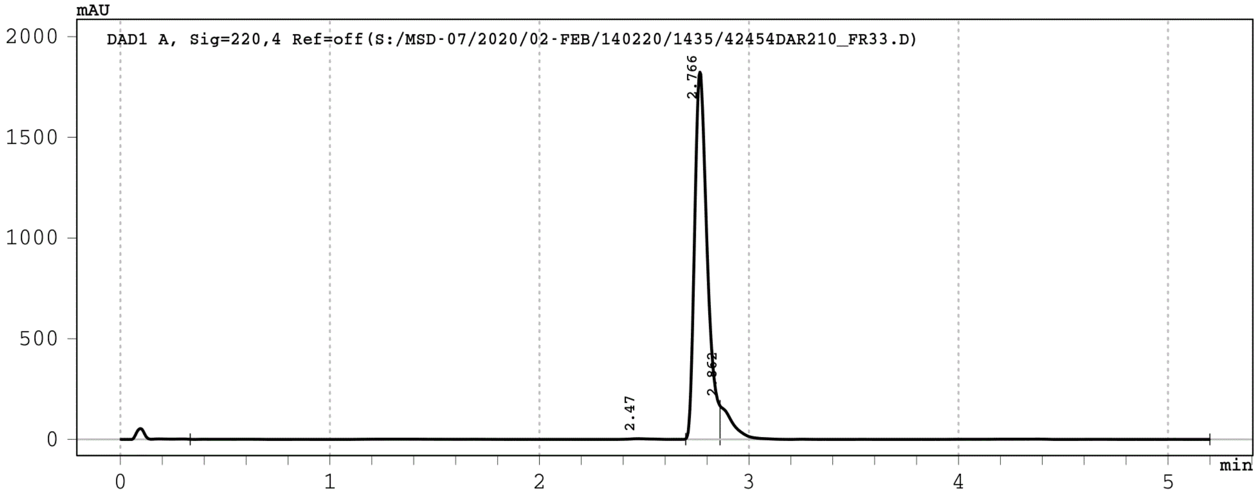


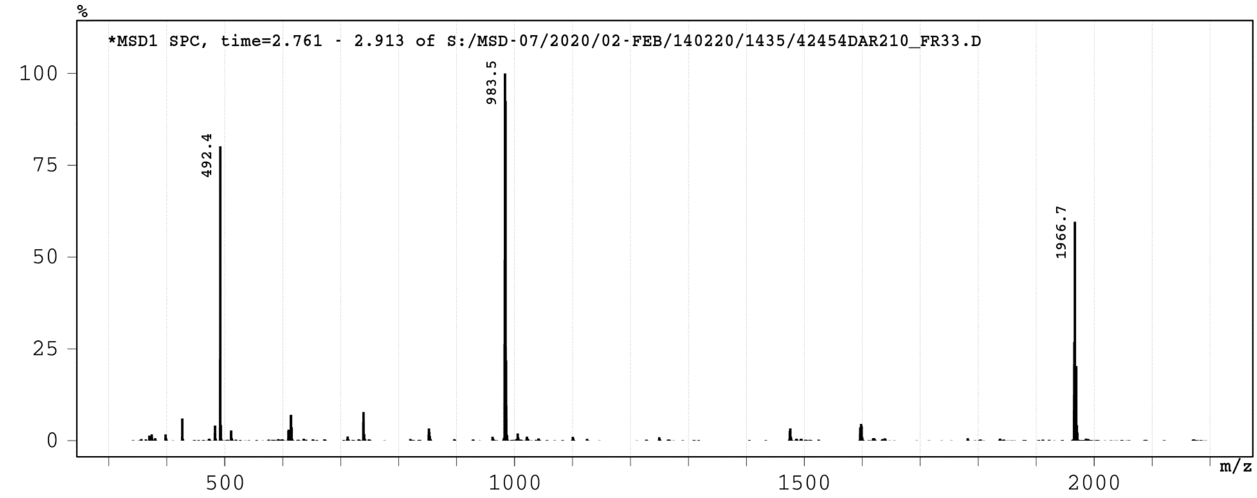


Signal 1: DAD1 A, Sig=220,4 Ref=off

| Peak # | RT [min] | Width [min] | Area | Area % |
|---|---|---|---|---|
| 1 | 2.470 | 2.36 | 71.0 | 0.8 |
| 2 | 2.766 | 0.16 | 7638.0 | 90.5 |
| 3 | 2.862 | 2.34 | 734.0 | 8.7 |

Data File : S:/MSD-07/2020/02-FEB/100220/1432/42454DAR211_FR5->

Injection Date : 10-Feb-20, 19:49:28
Sample Name : 42454DAR211_FR52-53
Remarks : Baseline corrected
Acq. Method : D:/METHODS/5-95-6M.M
column used : Je_559
Analysis Method : C:/CHEM32/1/METHODS/JPT-LCMS.M

Sample Info : 42454_211-Dr. Martine Darwish H-VAALREFLV-OH

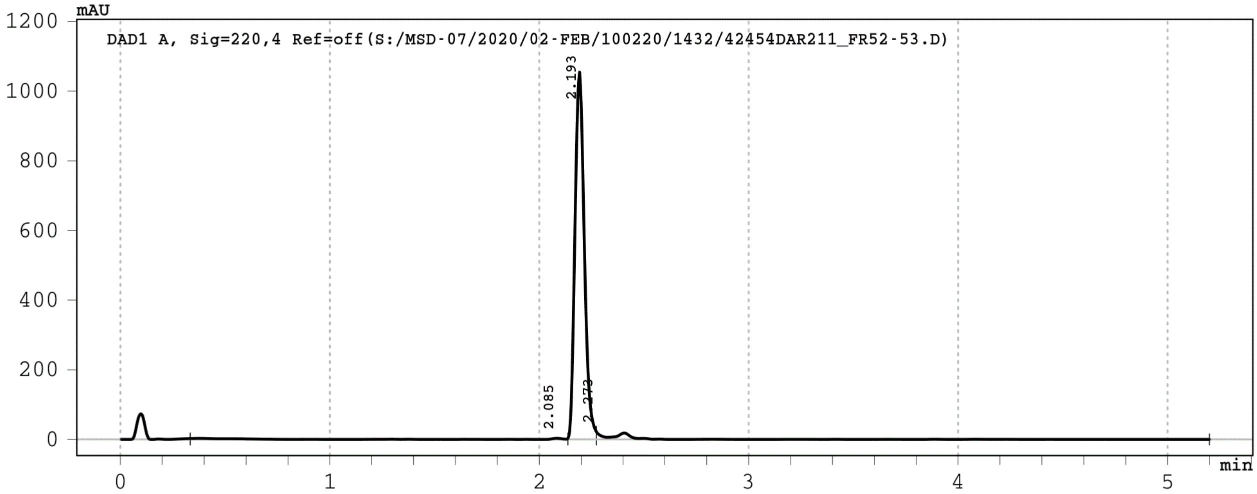


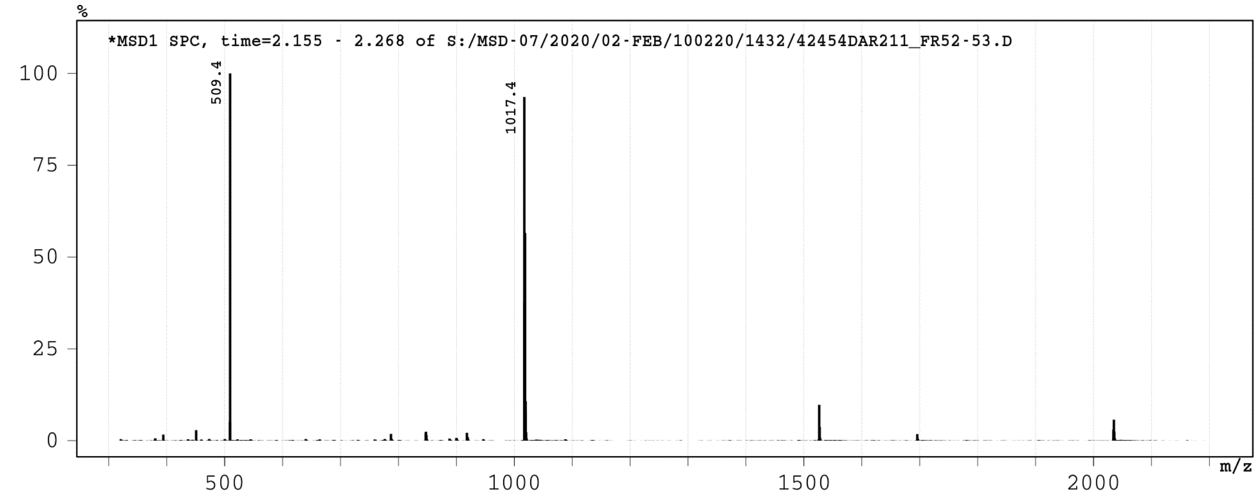


Signal 1: DAD1 A, Sig=220,4 Ref=off

| Peak # | RT [min] | Width [min] | Area | Area % |
|---|---|---|---|---|
| 1 | 2.085 | 1.80 | 62.3 | 1.8 |
| 2 | 2.193 | 0.14 | 3278.7 | 94.1 |
| 3 | 2.273 | 2.93 | 141.5 | 4.1 |

Data File : S:/MSD-07/2020/02-FEB/180220/1438/42454DAR212_FR4->

Injection Date : 18-Feb-20, 10:23:16
Sample Name : 42454DAR212_FR48
Remarks : Baseline corrected
Acq. Method : D:/METHODS/5-95-6M.M
column used : Je_559
Analysis Method : C:/CHEM32/1/METHODS/JPT-LCMS.M

Sample Info : 42454_212-Dr. Martine Darwish H-QVHWFTEL-OH

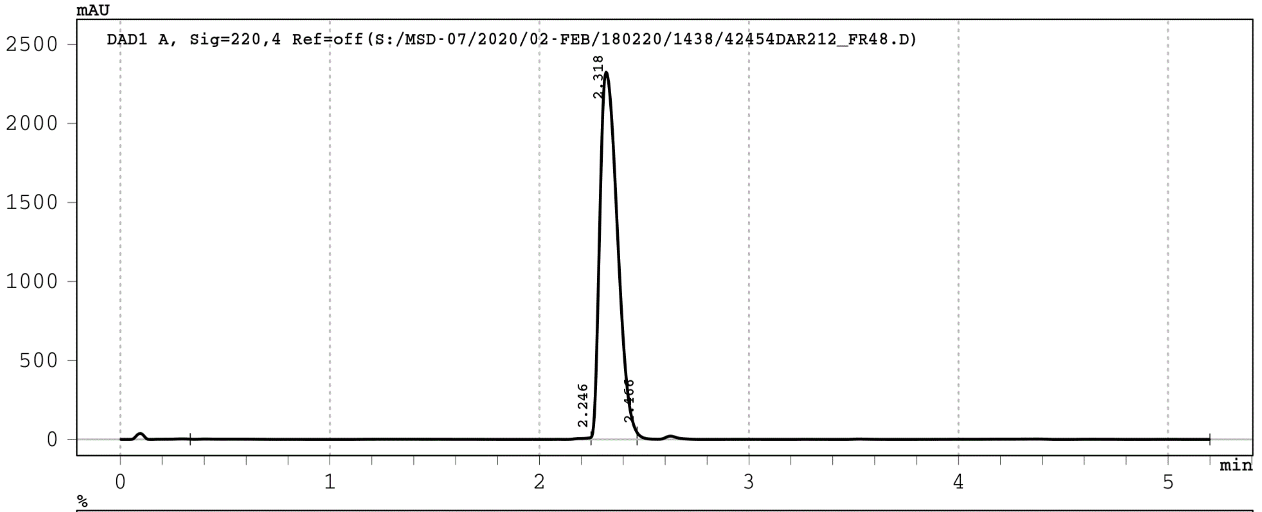


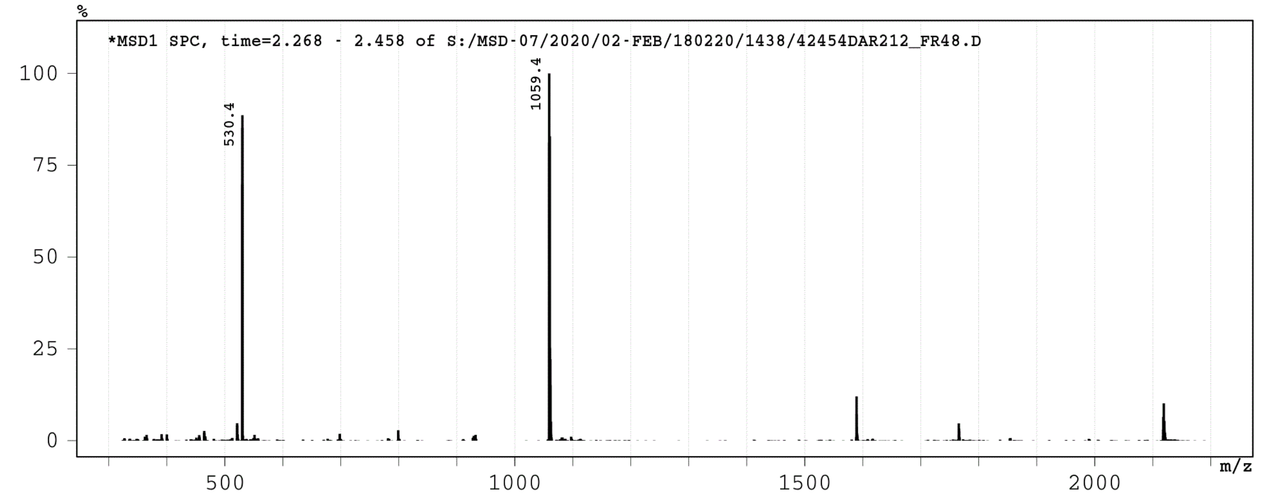


Signal 1: DAD1 A, Sig=220,4 Ref=off

| Peak # | RT [min] | Width [min] | Area | Area % |
|---|---|---|---|---|
| 1 | 2.246 | 1.91 | 95.5 | 0.7 |
| 2 | 2.318 | 0.22 | 13619.7 | 97.8 |
| 3 | 2.466 | 2.73 | 206.8 | 1.5 |

Data File : S:/MSD-07/2020/02-FEB/100220/1432/42454DAR213_FR6->

Injection Date : 10-Feb-20, 19:57:50
Sample Name : 42454DAR213_FR61-62
Remarks : Baseline corrected
Acq. Method : D:/METHODS/5-95-6M.M
column used : Je_559
Analysis Method : C:/CHEM32/1/METHODS/JPT-LCMS.M

Sample Info : 42454_213-Dr. Martine Darwish H-RALSGLEPF-OH

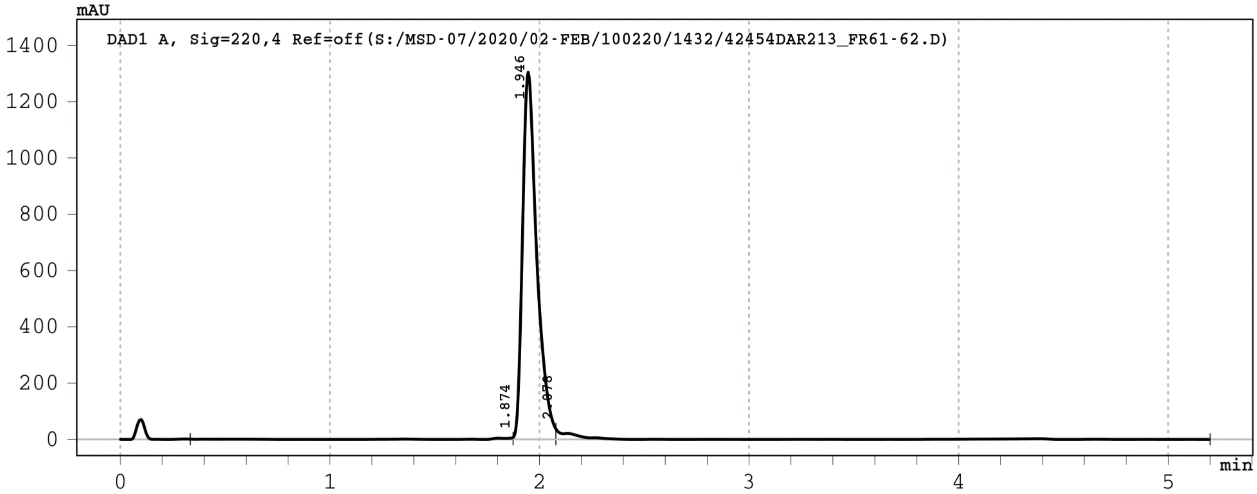


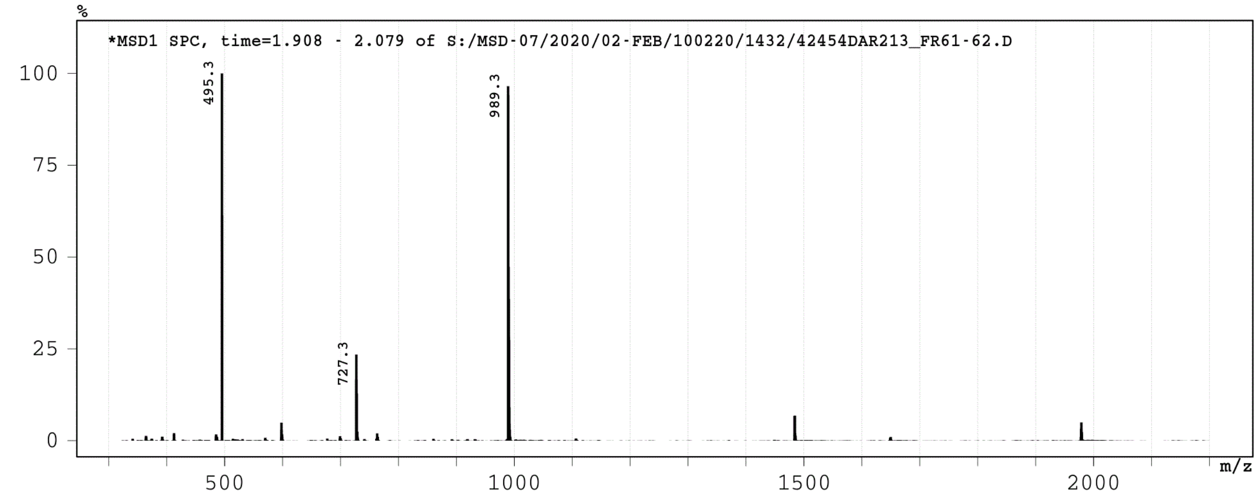


Signal 1: DAD1 A, Sig=220,4 Ref=off

| Peak # | RT [min] | Width [min] | Area | Area % |
|---|---|---|---|---|
| 1 | 1.874 | 1.54 | 53.6 | 0.8 |
| 2 | 1.946 | 0.20 | 6182.7 | 95.3 |
| 3 | 2.078 | 3.12 | 252.4 | 3.9 |

Data File : S:/MSD-07/2020/02-FEB/140220/1435/42454DAR215_FR3->

Injection Date : 15-Feb-20, 01:26:57
Sample Name : 42454DAR215_FR39-41
Remarks : Baseline corrected
Acq. Method : D:/METHODS/5-95-6M.M
column used : Je_559
Analysis Method : C:/CHEM32/1/METHODS/JPT-LCMS.M

Sample Info : 42454_215-Dr. Martine Darwish H-VSPQHRPV-OH

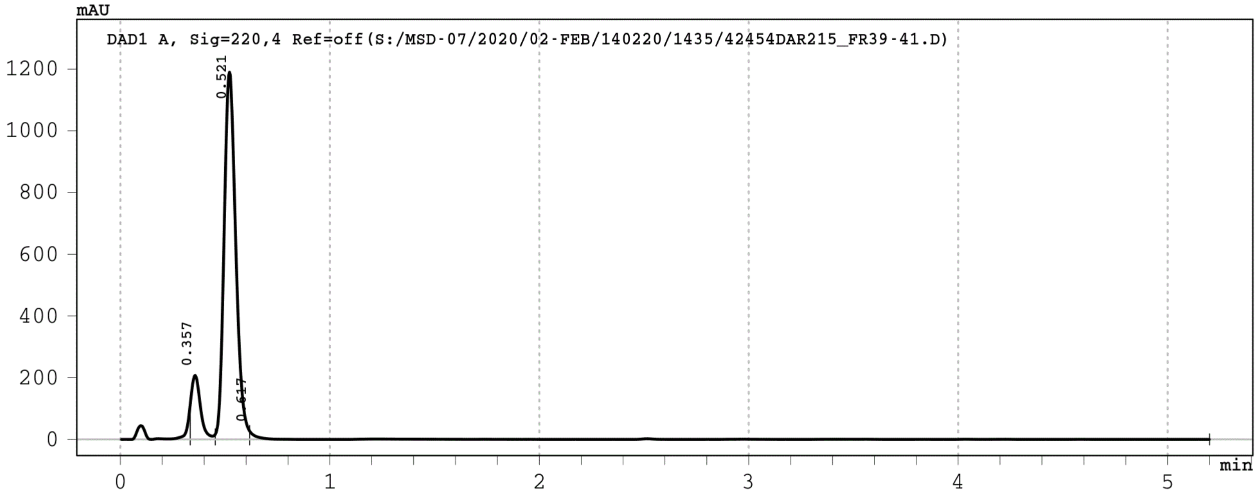


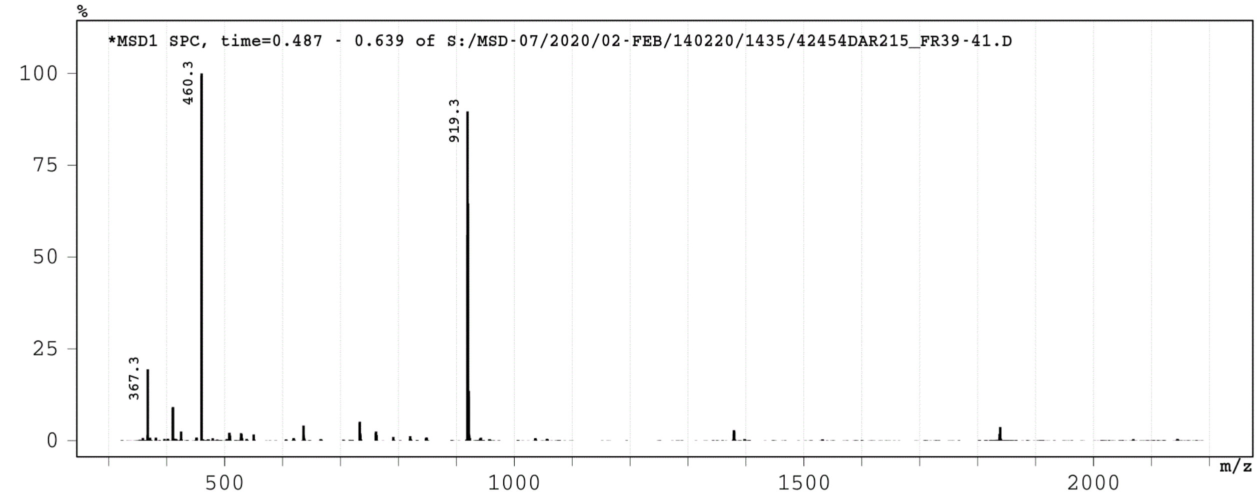


Signal 1: DAD1 A, Sig=220,4 Ref=off

| Peak # | RT [min] | Width [min] | Area | Area % |
|---|---|---|---|---|
| 1 | 0.357 | 0.12 | 599.0 | 11.3 |
| 2 | 0.521 | 0.16 | 4601.4 | 86.7 |
| 3 | 0.617 | 4.58 | 106.2 | 2.0 |

Data File : S:/MSD-07/2020/02-FEB/100220/1432/42454DAR216_FR8->

Injection Date : 10-Feb-20, 20:14:38
Sample Name : 42454DAR216_FR80-81
Remarks : Baseline corrected
Acq. Method : D:/METHODS/5-95-6M.M
column used : Je_559
Analysis Method : C:/CHEM32/1/METHODS/JPT-LCMS.M

Sample Info : 42454_216-Dr. Martine Darwish H-IGDLRLATL-OH

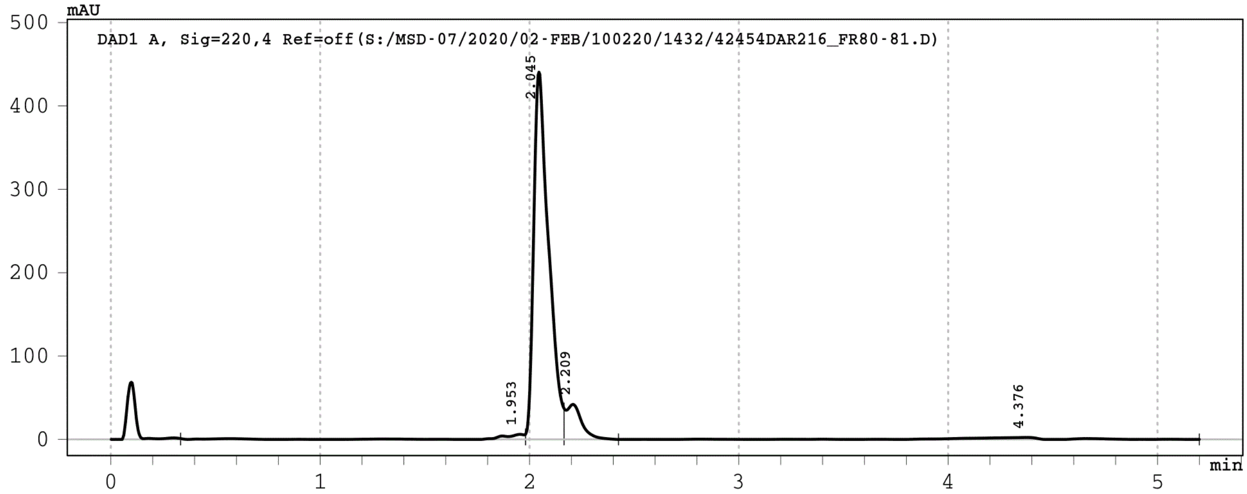


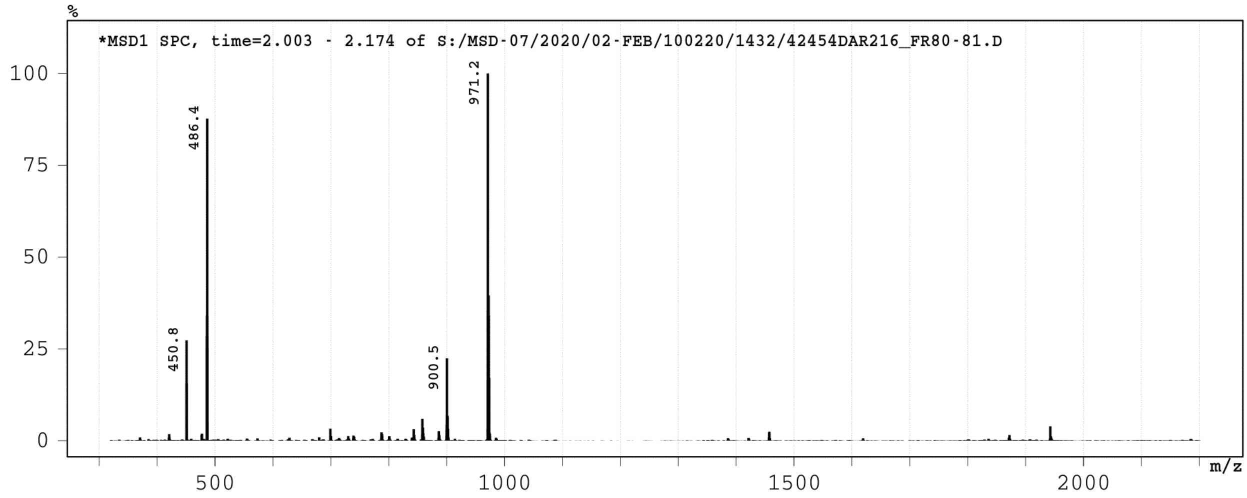


Signal 1: DAD1 A, Sig=220,4 Ref=off

| Peak # | RT [min] | Width [min] | Area | Area % |
|---|---|---|---|---|
| 1 | 1.953 | 1.65 | 60.3 | 2.4 |
| 2 | 2.045 | 0.18 | 2192.8 | 86.1 |
| 3 | 2.209 | 0.26 | 225.2 | 8.8 |
| 4 | 4.376 | 2.77 | 67.1 | 2.6 |

Data File : S:/MSD-07/2020/02-FEB/100220/1432/42454DAR217_FR1->

Injection Date : 10-Feb-20, 20:23:00
Sample Name : 42454DAR217_FR10-11
Remarks : Baseline corrected
Acq. Method : D:/METHODS/5-95-6M.M
column used : Je_559
Analysis Method : C:/CHEM32/1/METHODS/JPT-LCMS.M

Sample Info : 42454_217-Dr. Martine Darwish H-LNQLYELAL-OH

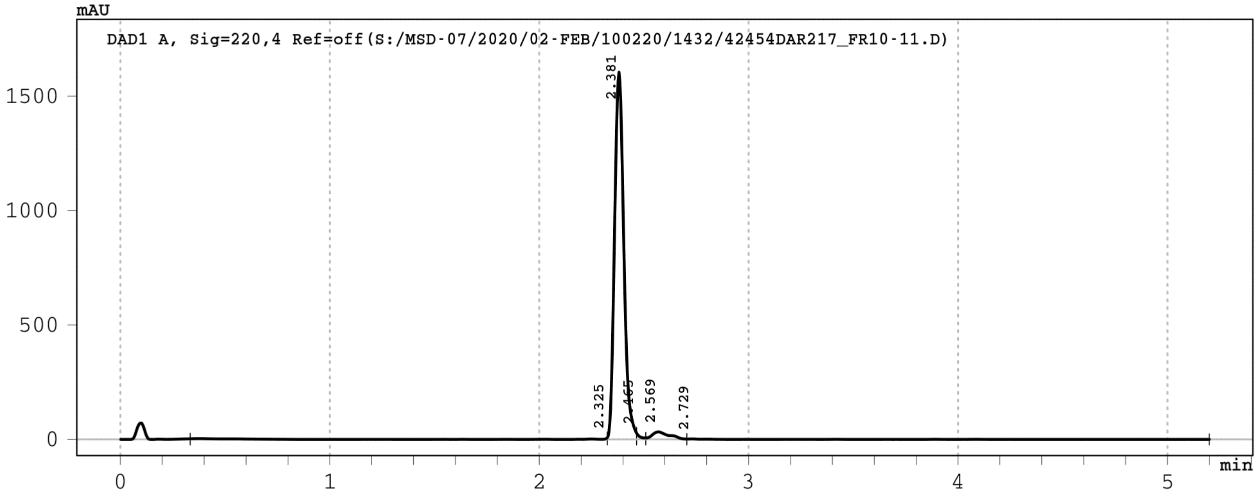


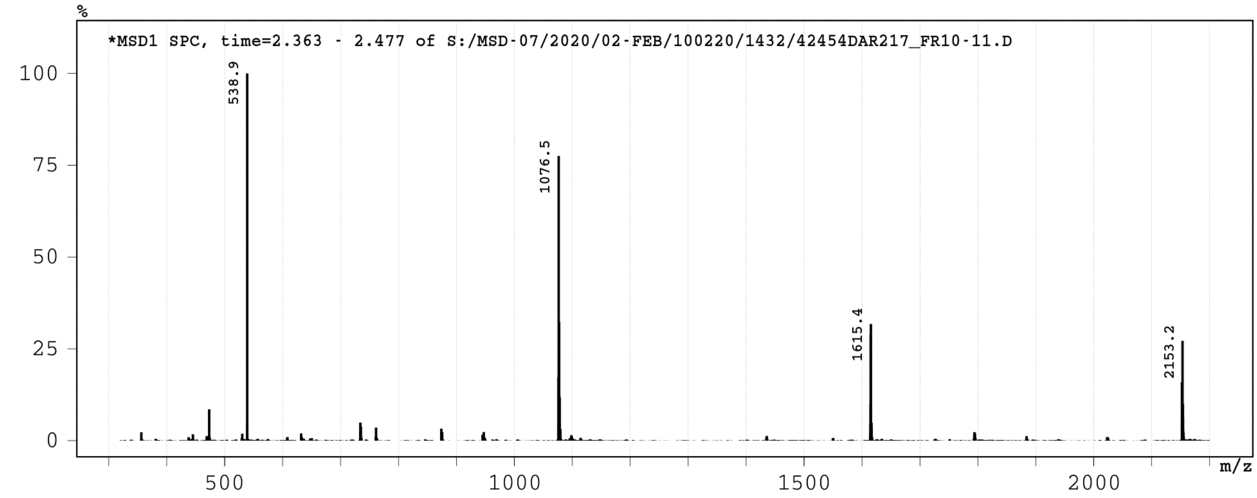


Signal 1: DAD1 A, Sig=220,4 Ref=off

| Peak # | RT [min] | Width [min] | Area | Area % |
|---|---|---|---|---|
| 1 | 2.325 | 1.99 | 69.1 | 1.3 |
| 2 | 2.381 | 0.14 | 4930.9 | 94.1 |
| 3 | 2.465 | 0.04 | 30.5 | 0.6 |
| 4 | 2.569 | 0.20 | 192.4 | 3.7 |
| 5 | 2.729 | 2.50 | 18.5 | 0.4 |

Data File : S:/MSD-07/2020/02-FEB/140220/1435/42454DAR218_FR4->

Injection Date : 15-Feb-20, 01:35:22
Sample Name : 42454DAR218_FR44-46
Remarks : Baseline corrected
Acq. Method : D:/METHODS/5-95-6M.M
column used : Je_559
Analysis Method : C:/CHEM32/1/METHODS/JPT-LCMS.M

Sample Info : 42454_218-Dr. Martine Darwish H-AVIGYSLL-OH

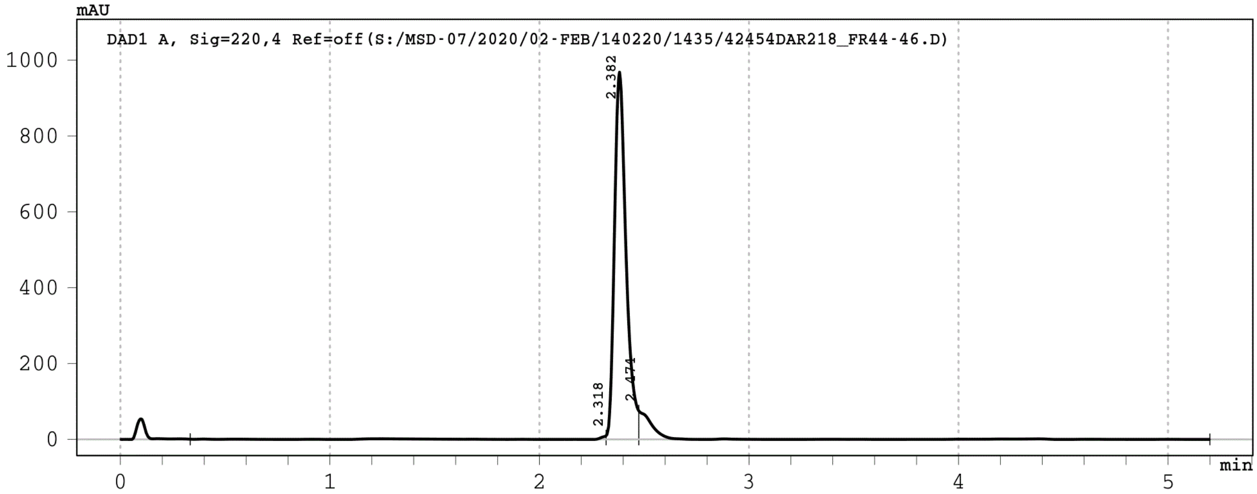


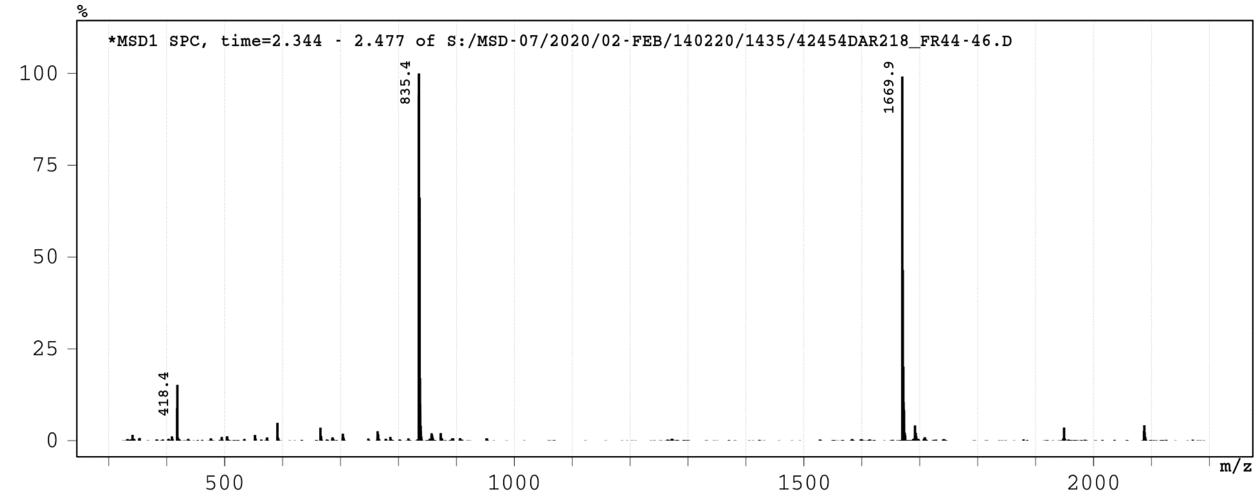


Signal 1: DAD1 A, Sig=220,4 Ref=off

| Peak # | RT [min] | Width [min] | Area | Area % |
|---|---|---|---|---|
| 1 | 2.318 | 1.98 | 63.3 | 1.5 |
| 2 | 2.382 | 0.16 | 3754.1 | 89.9 |
| 3 | 2.474 | 2.73 | 357.5 | 8.6 |

Data File : S:/MSD-07/2020/02-FEB/060220/1429/42454DAR219_FR4->

Injection Date : 06-Feb-20, 17:40:14
Sample Name : 42454DAR219_FR45-46
Remarks : Baseline corrected
Acq. Method : D:/METHODS/5-95-6M.M
column used : Je_559
Analysis Method : C:/CHEM32/1/METHODS/JPT-LCMS.M

Sample Info : 42454_219-Dr. Martine Darwish H-SLLEHMSLL-OH

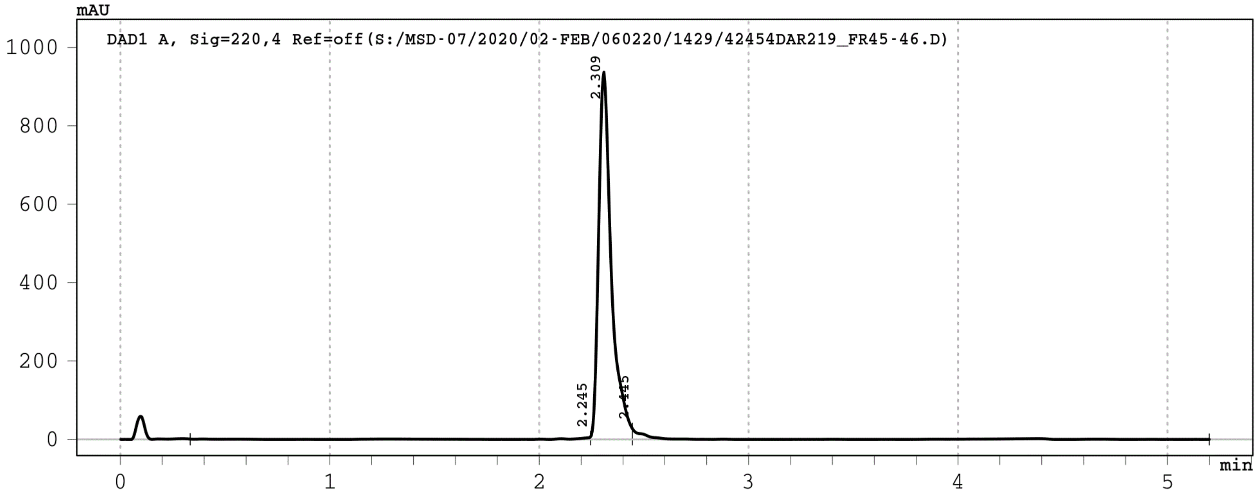


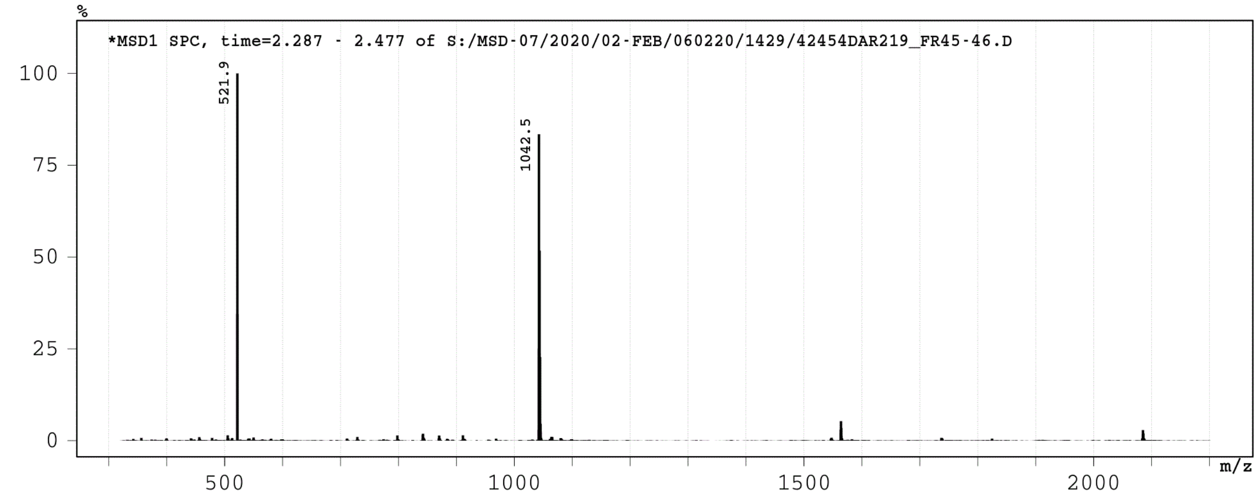


Signal 1: DAD1 A, Sig=220,4 Ref=off

| Peak # | RT [min] | Width [min] | Area | Area % |
|---|---|---|---|---|
| 1 | 2.245 | 1.91 | 54.7 | 1.3 |
| 2 | 2.309 | 0.20 | 3968.3 | 95.2 |
| 3 | 2.445 | 2.76 | 146.9 | 3.5 |

Data File: S:\MSD-07\2020\02-FEB\060220\142->

Injection Date : Thu, 6. Feb. 2020
Sample Name : 240120SG2-38|42454Dar220
Acq Operator : SYSTEM
Acq. Method : D:\Data\2020\02-FEB\060220\1429\5-95-6MM
column used :
Analysis Method : C:\Chem32\1\METHODS\DEF_LC.M

Sample Info : 42454_220-Dr. Martine Darwish|H-SAMAMFGYM-OH

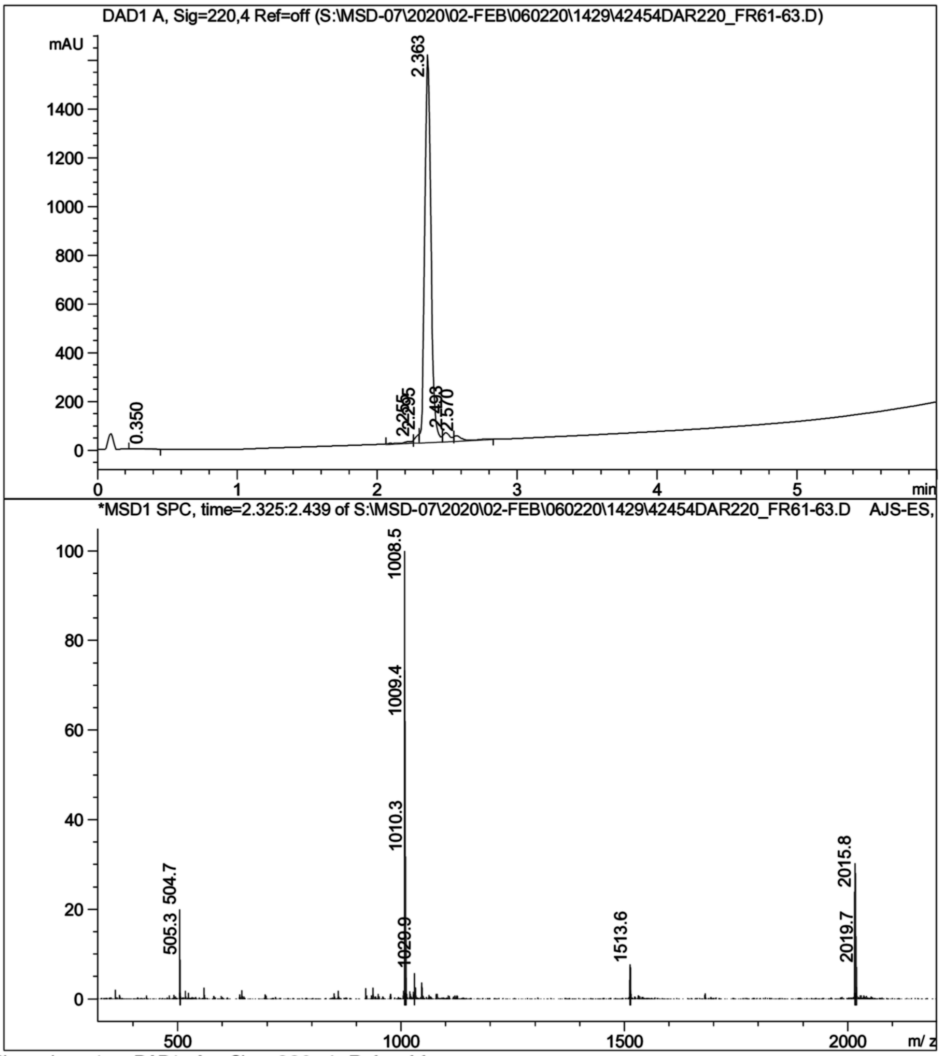


Signal 1: DAD1 A, Sig=220,4 Ref=off

| Peak # | RT [min] | Width [min] | Area | Area % |
|---|---|---|---|---|
| 1 | 0.350 | 0.1 | 7.6 | 0.1 |
| 2 | 2.255 | 0.1 | 56.7 | 1.1 |
| 3 | 2.295 | 0.0 | 65.3 | 1.2 |
| 4 | 2.363 | 0.1 | 4937.2 | 93.0 |
| 5 | 2.493 | 0.1 | 130.4 | 2.5 |
| 6 | 2.570 | 0.1 | 109.6 | 2.1 |

Data File : S:/MSD-07/2020/02-FEB/140220/1435/42454DAR221_FR5->

Injection Date : 15-Feb-20, 01:43:45
Sample Name : 42454DAR221_FR50-51
Remarks : Baseline corrected
Acq. Method : D:/METHODS/5-95-6M.M
column used : Je_559
Analysis Method : C:/CHEM32/1/METHODS/JPT-LCMS.M

Sample Info : 42454_221-Dr. Martine Darwish H-LQFIQSPL-OH

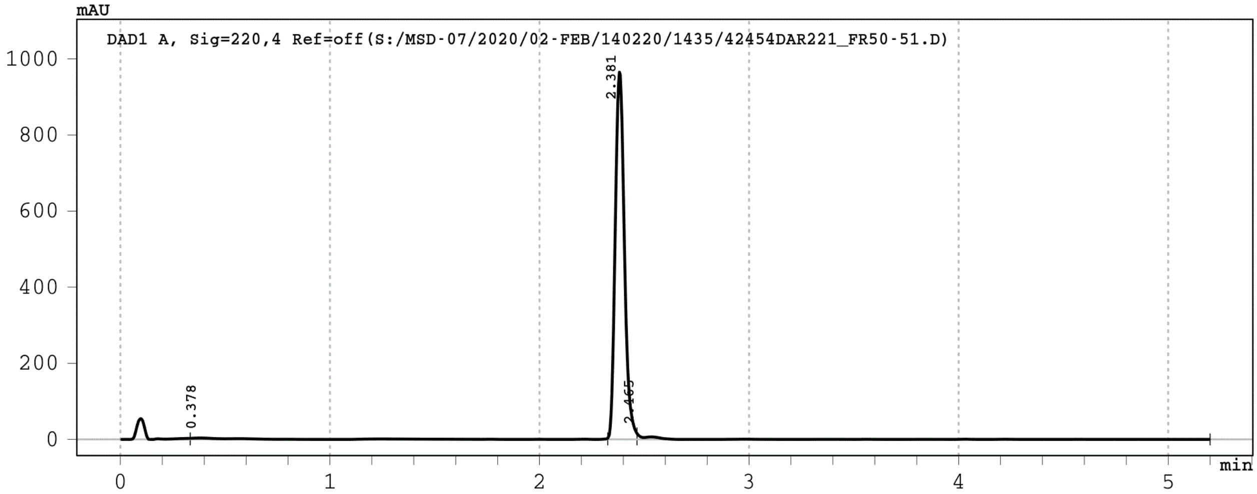


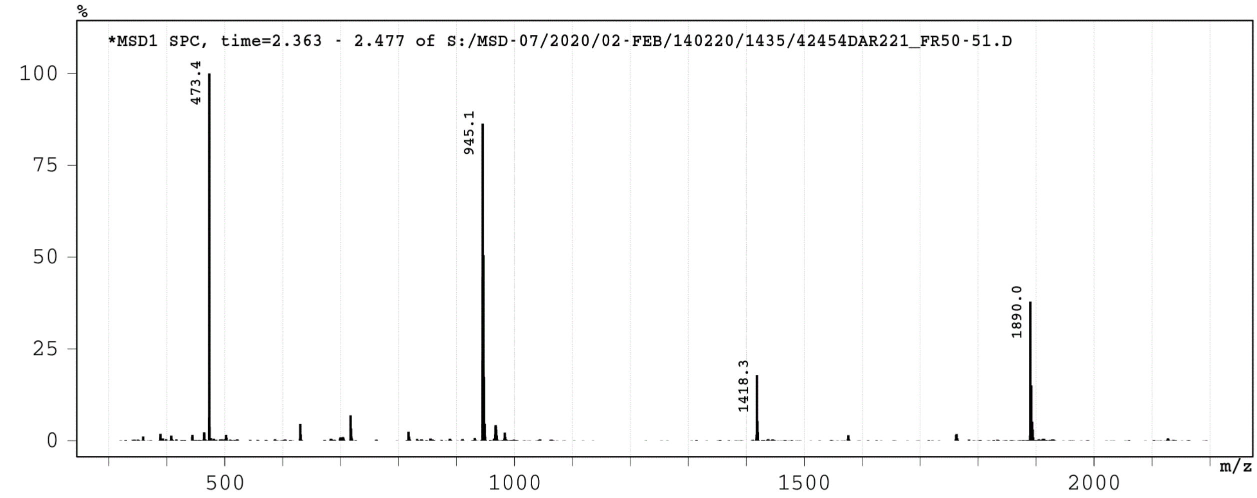


Signal 1: DAD1 A, Sig=220,4 Ref=off

| Peak # | RT [min] | Width [min] | Area | Area % |
|---|---|---|---|---|
| 1 | 0.378 | 1.99 | 82.0 | 2.7 |
| 2 | 2.381 | 0.14 | 2946.0 | 95.4 |
| 3 | 2.465 | 2.73 | 61.2 | 2.0 |

Data File : S:/MSD-07/2020/02-FEB/100220/1432/42454DAR222_FR2->

Injection Date : 10-Feb-20, 20:31:23
Sample Name : 42454DAR222_FR20-21
Remarks : Baseline corrected
Acq. Method : D:/METHODS/5-95-6M.M
column used : Je_559
Analysis Method : C:/CHEM32/1/METHODS/JPT-LCMS.M

Sample Info : 42454_222-Dr. Martine Darwish H-IIAVLKQRL-OH

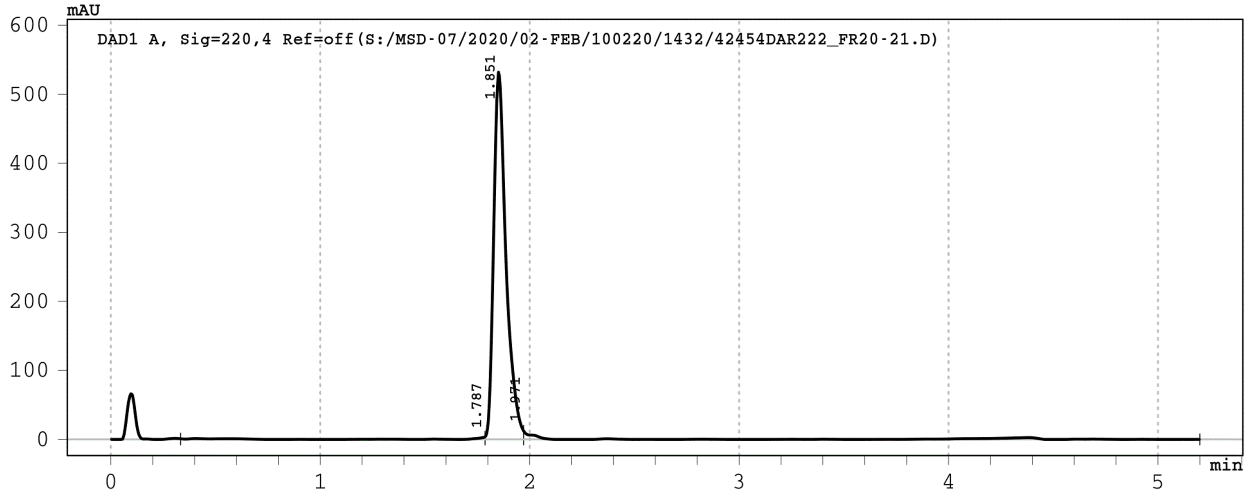


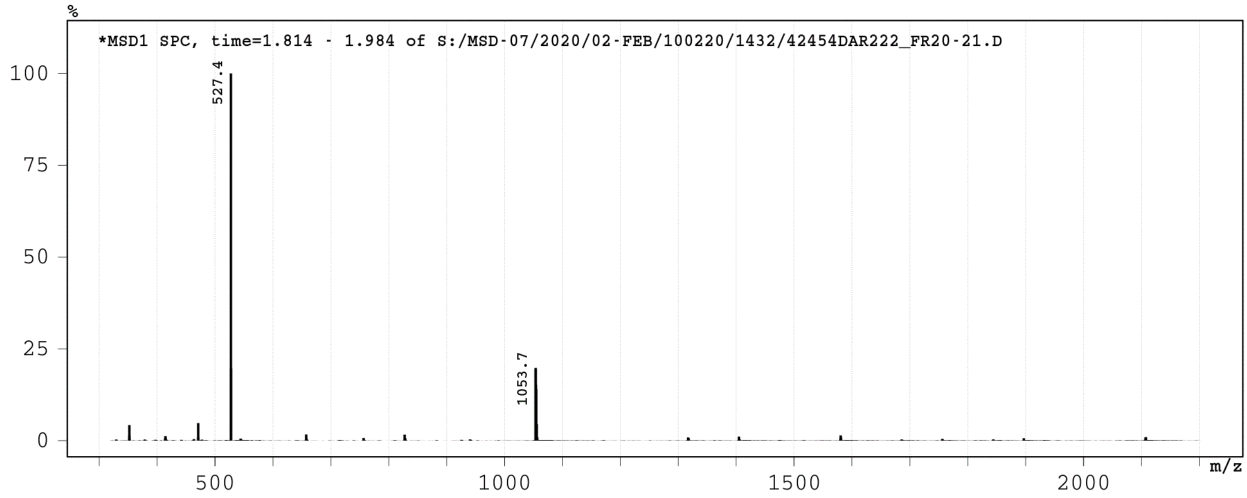


Signal 1: DAD1 A, Sig=220,4 Ref=off

| Peak # | RT [min] | Width [min] | Area | Area % |
|---|---|---|---|---|
| 1 | 1.787 | 1.45 | 32.1 | 1.4 |
| 2 | 1.851 | 0.18 | 2187.3 | 94.5 |
| 3 | 1.971 | 3.23 | 96.3 | 4.2 |

Data File : S:/MSD-07/2020/02-FEB/060220/1429/42454DAR223_FR7->

Injection Date : 06-Feb-20, 17:57:01
Sample Name : 42454DAR223_FR77-78
Remarks : Baseline corrected
Acq. Method : D:/METHODS/5-95-6M.M
column used : Je_559
Analysis Method : C:/CHEM32/1/METHODS/JPT-LCMS.M

Sample Info : 42454_223-Dr. Martine Darwish H-RGFMWLPGF-OH

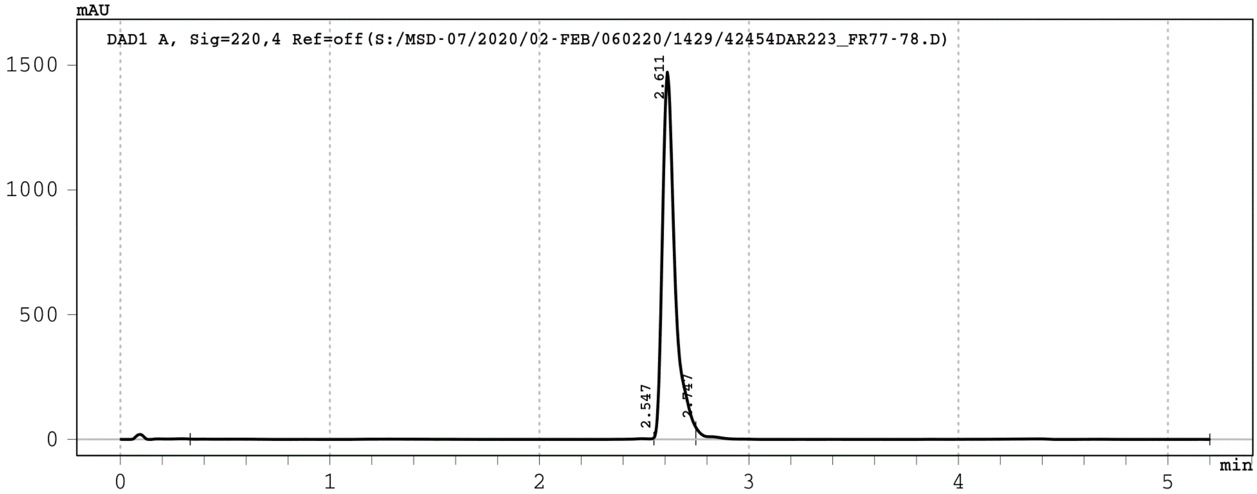


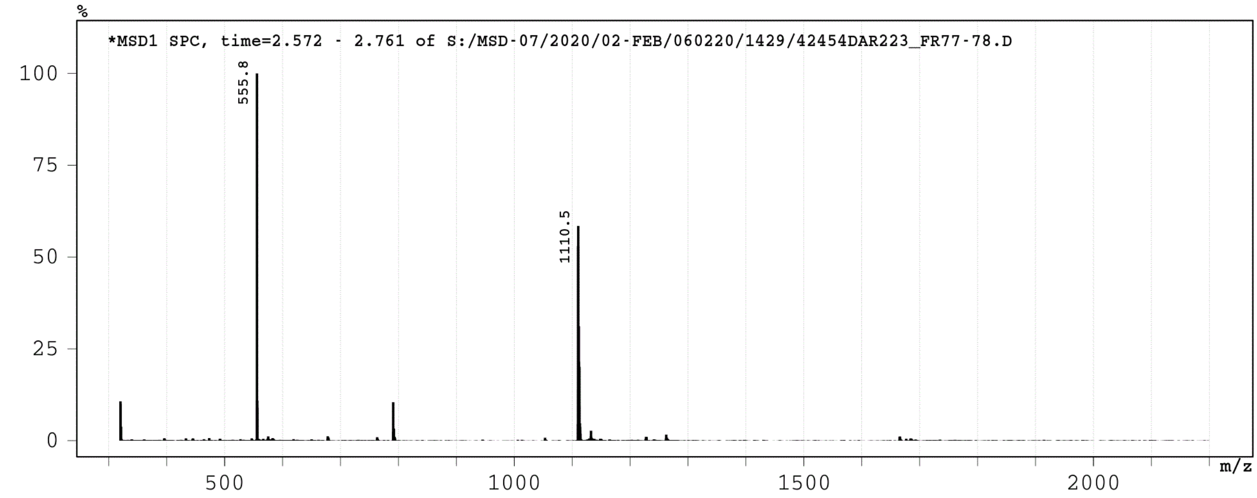


Signal 1: DAD1 A, Sig=220,4 Ref=off

| Peak # | RT [min] | Width [min] | Area | Area % |
|---|---|---|---|---|
| 1 | 2.547 | 2.21 | 63.5 | 1.0 |
| 2 | 2.611 | 0.20 | 6164.8 | 96.3 |
| 3 | 2.747 | 2.45 | 172.2 | 2.7 |

Data File : S:/MSD-07/2020/02-FEB/140220/1435/42454DAR224_FR5->

Injection Date : 15-Feb-20, 01:52:08
Sample Name : 42454DAR224_FR54-56
Remarks : Baseline corrected
Acq. Method : D:/METHODS/5-95-6M.M
column used : Je_559
Analysis Method : C:/CHEM32/1/METHODS/JPT-LCMS.M

Sample Info : 42454_224-Dr. Martine Darwish H-SNHVLGHL-OH

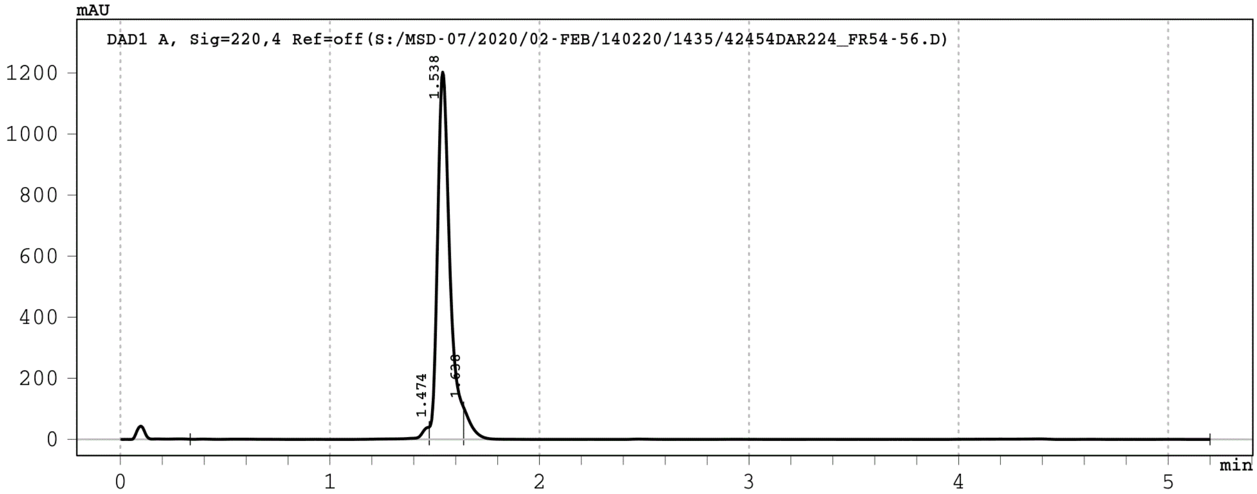


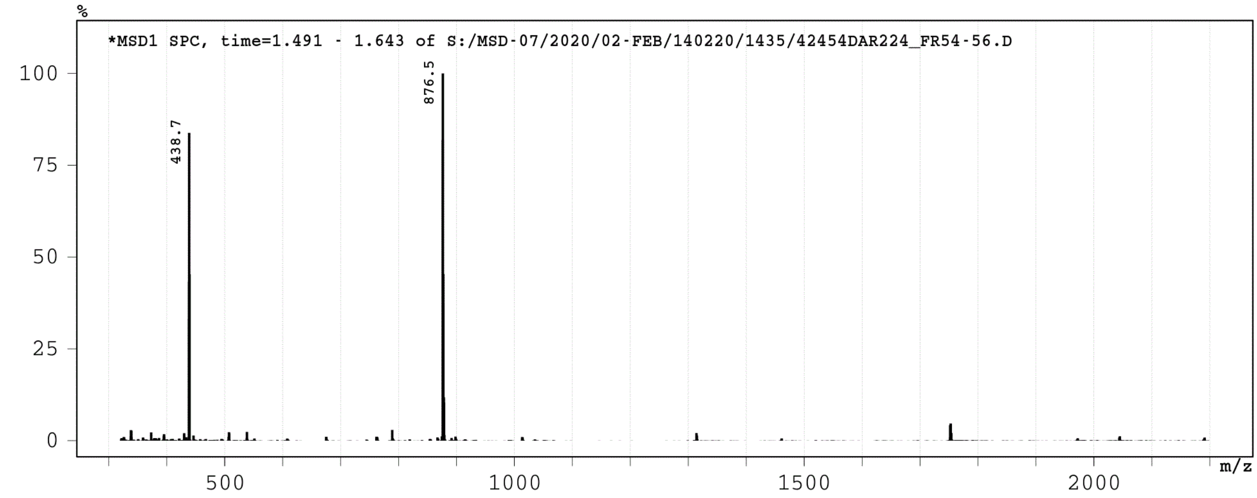


Signal 1: DAD1 A, Sig=220,4 Ref=off

| Peak # | RT [min] | Width [min] | Area | Area % |
|---|---|---|---|---|
| 1 | 1.474 | 1.14 | 127.1 | 2.4 |
| 2 | 1.538 | 0.16 | 4874.3 | 92.1 |
| 3 | 1.638 | 3.56 | 293.4 | 5.5 |

Data File : S:/MSD-07/2020/02-FEB/060220/1429/42454DAR225_FR9->

Injection Date : 06-Feb-20, 18:05:25
Sample Name : 42454DAR225_FR9-10
Remarks : Baseline corrected
Acq. Method : D:/METHODS/5-95-6M.M
column used : Je_559
Analysis Method : C:/CHEM32/1/METHODS/JPT-LCMS.M

Sample Info : 42454_225-Dr. Martine Darwish H-LMLENYNNL-OH

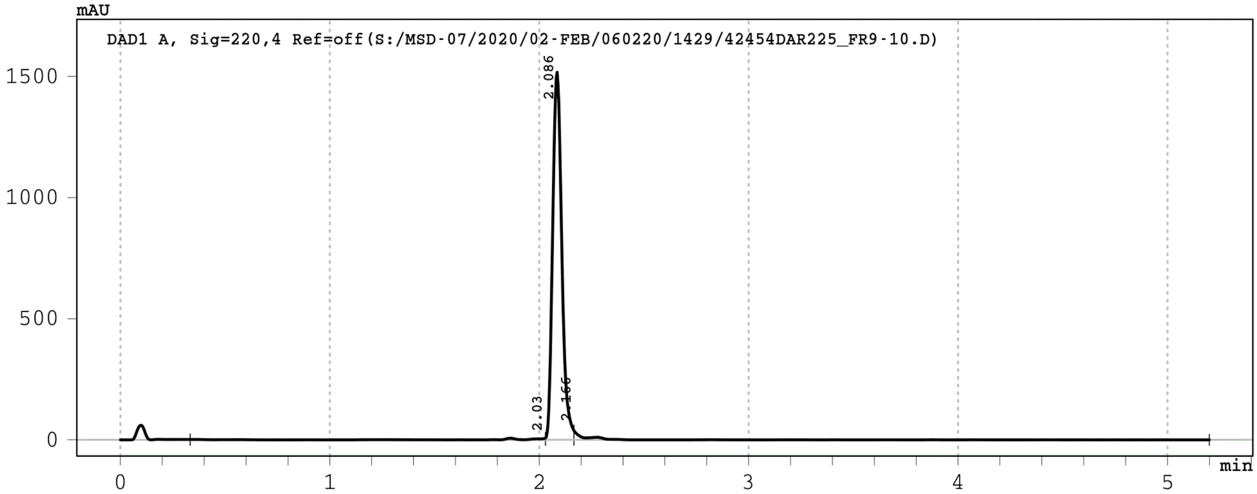


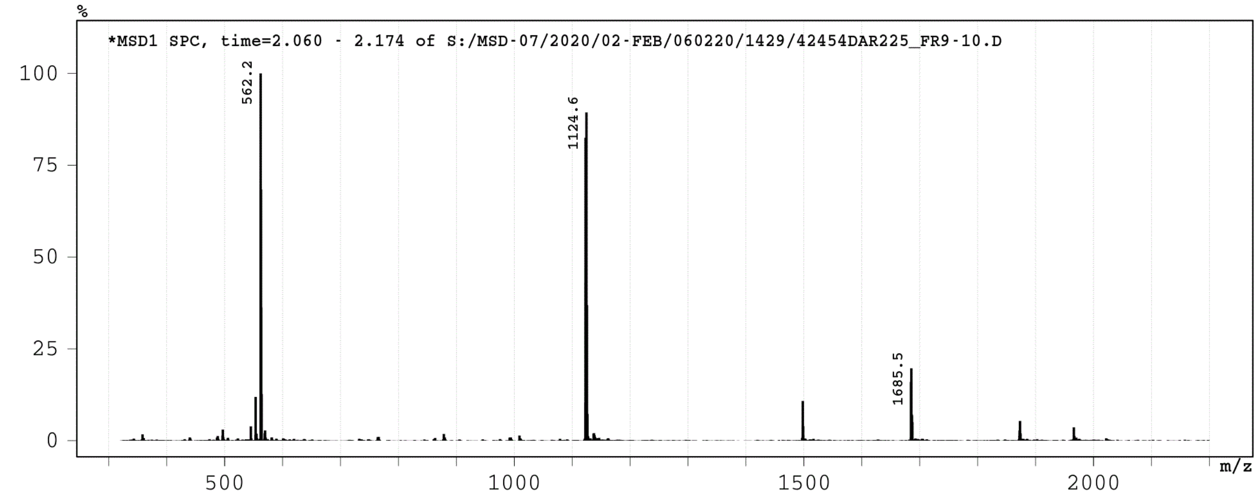


Signal 1: DAD1 A, Sig=220,4 Ref=off

| Peak # | RT [min] | Width [min] | Area | Area % |
|---|---|---|---|---|
| 1 | 2.030 | 1.70 | 70.1 | 1.5 |
| 2 | 2.086 | 0.14 | 4612.0 | 95.8 |
| 3 | 2.166 | 3.03 | 130.6 | 2.7 |

Data File : S:/MSD-07/2020/02-FEB/100220/1432/42454DAR226_FR2->

Injection Date : 10-Feb-20, 20:39:45
Sample Name : 42454DAR226_FR26-27
Remarks : Baseline corrected
Acq. Method : D:/METHODS/5-95-6M.M
column used : Je_559
Analysis Method : C:/CHEM32/1/METHODS/JPT-LCMS.M

Sample Info : 42454_226-Dr. Martine Darwish H-KSFRQKPNL-OH

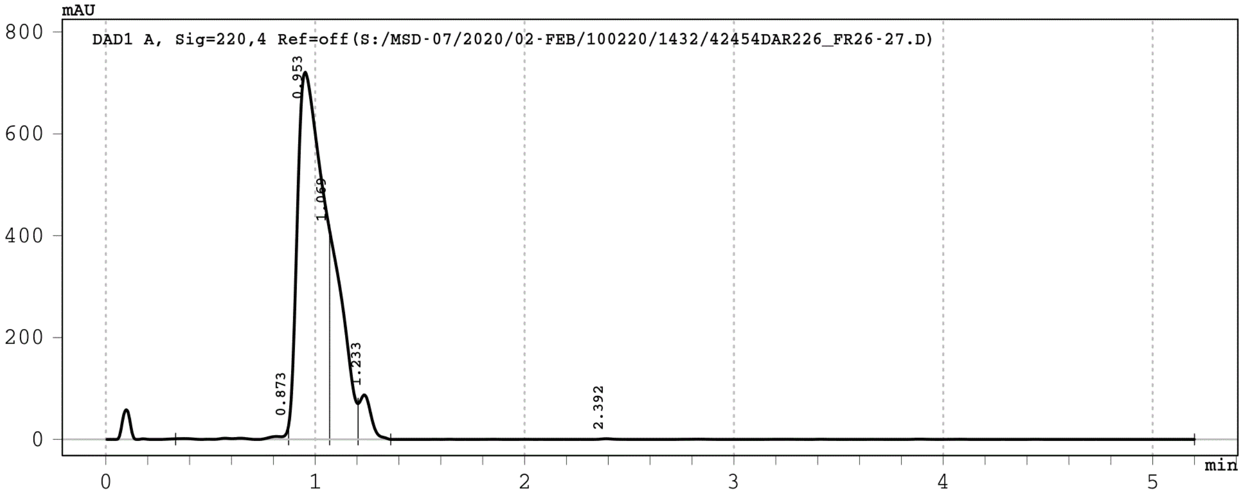


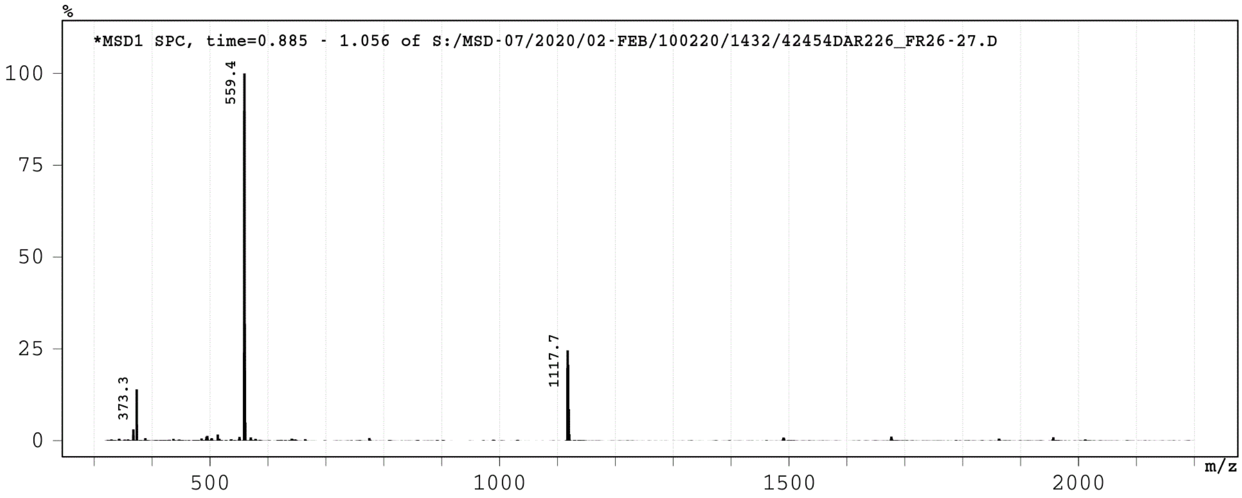


Signal 1: DAD1 A, Sig=220,4 Ref=off

| Peak # | RT [min] | Width [min] | Area | Area % |
|---|---|---|---|---|
| 1 | 0.873 | 0.54 | 68.9 | 0.8 |
| 2 | 0.953 | 0.20 | 5864.3 | 71.4 |
| 3 | 1.069 | 0.14 | 1921.7 | 23.4 |
| 4 | 1.233 | 0.16 | 343.8 | 4.2 |
| 5 | 2.392 | 3.84 | 16.1 | 0.2 |

Data File : S:/MSD-07/2020/02-FEB/060220/1429/42454DAR227_FR1->

Injection Date : 06-Feb-20, 18:13:48
Sample Name : 42454DAR227_FR17
Remarks : Baseline corrected
Acq. Method : D:/METHODS/5-95-6M.M
column used : Je_559
Analysis Method : C:/CHEM32/1/METHODS/JPT-LCMS.M

Sample Info : 42454_227-Dr. Martine Darwish H-ASACNVQHL-OH

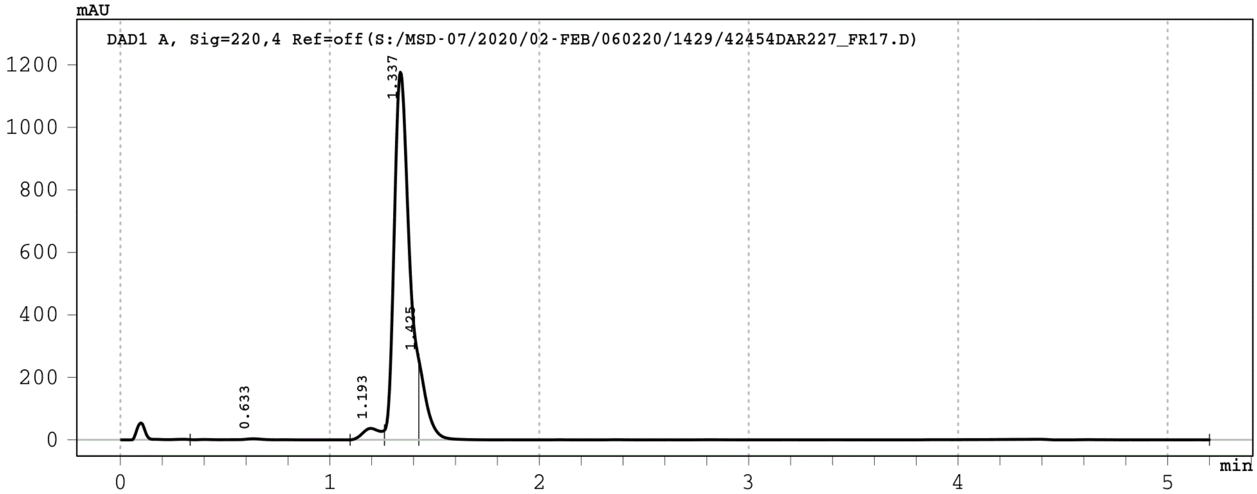


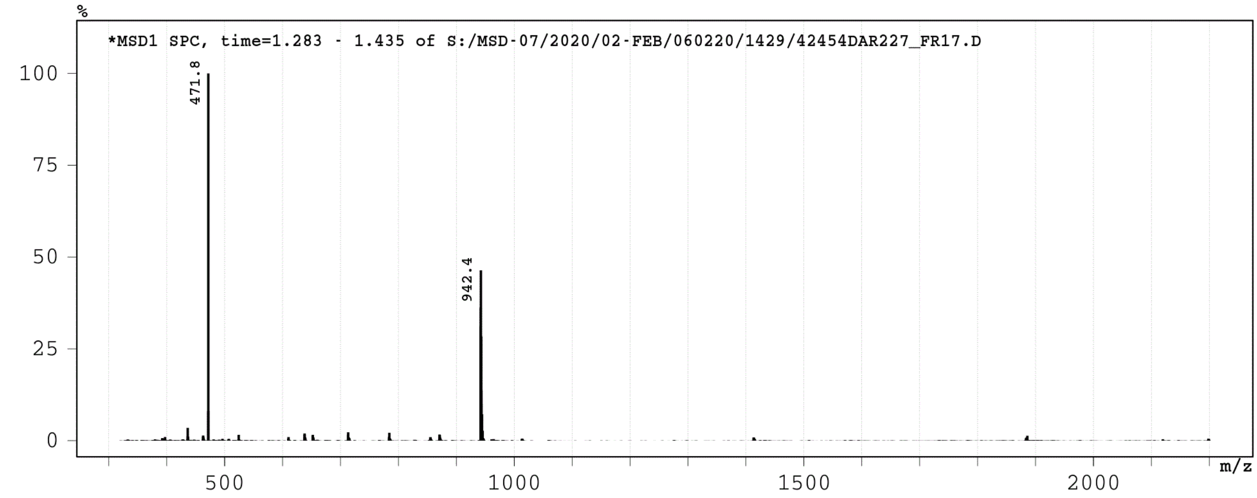


Signal 1: DAD1 A, Sig=220,4 Ref=off

| Peak # | RT [min] | Width [min] | Area | Area % |
|---|---|---|---|---|
| 1 | 0.633 | 0.76 | 29.0 | 0.4 |
| 2 | 1.193 | 0.16 | 220.6 | 3.2 |
| 3 | 1.337 | 0.16 | 5840.4 | 85.9 |
| 4 | 1.425 | 3.78 | 710.8 | 10.5 |
